# Microbial, dietary insect, and pathogen communities in fresh and decomposing guano of anthropic little brown bat (Myotis lucifugus) maternity colonies

**DOI:** 10.64898/2026.07.02.734860

**Authors:** Jagger Defenza, Laurel Eddins, Ashton Gauvin, Donal J. Heaney, Dantony Lewis, Ruiwen Lin, Ruth Martinez, Kevin Schilace, Kristiana A. Stover, Jillian Taormina, Sofia Carolina Vargas, Kyle Paist, Kendra Maas, David J. Askew, Kate Castellano, Mary Rutter, Adam Rork, Nicole Pauloski, Rachel J. O’Neill, Teisha King, Elizabeth L. Jockusch, Jill L. Wegrzyn, James Fischer, Airianna McGuire, Devaughn Fraser, Hannah Reynolds

## Abstract

The guano of insectivorous bats holds ecological information that can be assessed non-invasively to characterize the gut microbiome and diet, alongside environmental microbes and pathogens of conservation concern. Despite this potential, how guano communities change during decomposition remains understudied, particularly inside anthropic roosts rather than caves. In this study, guano from little brown bat (*Myotis lucifugus*) colonies was sampled monthly across the summer maternity season from three sites across two locations in Connecticut, USA, at fresh deposition and at 4, 8, and 12 weeks following deposition. Resolving these cross-kingdom signals required five workflows: short-read 16S and ITS2 amplicon sequencing (Illumina) for bacterial and fungal profiling, long-read CO1 metabarcoding (Oxford Nanopore) for arthropod diet in fresh samples, long-read shotgun metagenomics for viral identification in aged samples, and targeted qPCR for organisms of bat, human, and forest-health concern. Fresh guano generated a consistent bacterial signal across sites, whereas fresh fungal communities differed by site. Responses to decomposition depended on roost setting: exterior sites lost fungal diversity and shifted toward environmental aerobes over time, while the interior roost retained the fresh sample profile. Dietary composition varied temporally, was dominated by Diptera, and included the invasive emerald ash borer (*Agrilus planipennis*). *Pseudogymnoascus destructans*, the causal agent of white-nose syndrome, occurred in fresh and aged samples at all three sites but persisted for 12 weeks only at the interior roost, where antifungal bacterial taxa were depleted. Long-read shotgun metagenomics of aged guano recovered roughly 100 viral species, predominantly bacteriophages, alongside non-bacteriophage mastadenoviruses associated with humans, bats, and other mammals. These results show that anthropic structures influence the trajectory of guano microbiome succession, and that maternity colony guano enables non-invasive assessment of environmental pathogens, bat diet, and bacterial and fungal communities.

## INTRODUCTION

Bats (Chiroptera) are a highly diverse group of mammals with a wide range of life history and feeding strategies (Gerbáčová et al. 2020) that provide ecosystem services including insect pest suppression, pollination, and seed dispersal (Williams-Guillén et al. 2015). As earth’s only flying mammals, bats sample vast areas of the local landscape each night while foraging, providing ecological data across space and time, particularly when analyzed at the colony level. Because guano is deposited continuously beneath roosts, it offers a non-invasive sample type from which dietary prey, host gut microbiota, environmental microbes, and pathogens can be characterized longitudinally.

Despite the information present in bat guano, characterization of the microbiome remains understudied. Community composition varies between bat species, geographical location, and guano age, yet previous studies of guano decomposition are limited in scope. Most effort has focused on microbial communities in natural long-term aged guano, particularly in cave environments where deposits can accumulate for hundreds of years (McFarlane & Lundberg 2024). Bacterial diversity is known to shift with the age of the guano: anaerobic and aerobic taxa differ between freshly collected guano and pile surface guano, and shift even further in decaying guano, where diversity is often lowest (Newman et al. 2018). While these studies provide context for the decomposition dynamics of the guano microbiome, they infer age from pile depth and carbon aging rather than directly aging samples (McFarlane & Lundberg 2024). While Fofanov et al. (2018) used directly aged guano to assess short-term shifts in guano communities, this was conducted at the scale of hours rather than weeks or months. These studies primarily investigate the bacterial component of the gut microbiome, leaving age-related change in the fungal microbiome understudied. The few studies that have assessed fungal content of guano focused on fresh guano of a frugivorous bat species (Chaverri & Chaverri 2022) or samples from cave populations with unknown aging times (Cunha et al. 2020; Ogorek et al. 2016). The guano virome is similarly understudied; bats harbor the greatest viral diversity of any mammal (Li Y. et al. 2021), yet few studies have characterized viruses from *M. lucifugus* guano, and fewer still from aged guano in anthropic roosts (Donaldson et al. 2010).

Equally understudied is how bat guano ages inside anthropic roosts. Bats commonly roost in structures not actively inhabited by humans (Voigt et al. 2015; Lučan et al. 2024). Similar to caves, these buildings offer protection from UV radiation and predation (Voigt et al. 2015), which may be particularly important in maternity roosts, yet it is not well understood how conditions inside anthropic roosts may affect community succession of bat guano or diet. Webster and Whitaker (2005) surveyed arthropod diversity in guano from synanthropic maternity roosts but did so in samples of unknown age. Bat guano is also a potential source of pathogenic bacteria, fungi, and viruses of human health concern; while transmission from bats to humans is rare, it is important to understand how the presence and abundance of these vectors may change in human structures over time (Voigt et al. 2015).

This study used fresh and *in situ* aged guano samples in anthropic roosts (internal and external bat boxes) to better understand how the bat guano microbiome and bat diet are affected over the maternity season. Capturing host, dietary, and environmental signals from a single guano source, while also screening for pests and pathogens, depends on methods that span diverse taxa. Short-read metabarcoding has been widely applied to characterize bat diet diversity (Galan et al. 2018; Bourlat et al. 2023), but long-read approaches are preferred for more confident taxonomic assignments (Portik et al. 2022). Here, short- and long-read approaches were combined with targeted assays to profile multiple taxa and support comparisons across time and space.

## METHODS AND MATERIALS

### Experimental design overview

The temporal dynamics of microbial, fungal, and dietary insect communities in *Myotis lucifugus* guano were characterized through monthly sampling from May to September 2025 across three maternity colony bat houses at two locations in Connecticut, USA. An *in situ* aging series was established at all three roosts and sampled at 4, 8, and 12 weeks following June and August deposition. DNA from each sample was processed through five molecular workflows, characterizing the bacterial, fungal, arthropod, and viral communities and testing for five pathogens and invasive insects of bat, human, and forest health concern. Downstream analyses integrated alpha and beta diversity comparisons (ANOVA, perMANOVA), multivariate modeling, microbial community origin analysis, and indicator taxa analysis to resolve taxa specific to site, age, and month.

### Fresh and aged guano sample collection

Guano samples were collected monthly (May to September 2025) from three sites across two locations: a private residence in Ashford, CT, and White Memorial Conservation Center in Litchfield, CT (Fig. 1A-C). The two locations house maternal colonies, restricting samples predominantly to reproductive females and pups. Sampling included bat houses mounted on the interior and exterior of a barn at the Ashford residence (ASHFInt and ASHFExt; Fig. 1D) and a pole-mounted, east-facing bat house at WMCC (Fig. 1E). Fresh guano was not collected in September at the ASHFInt site due to a decrease in or absence of bats. To collect guano, aluminum foil sheets were autoclaved using the “wrapped with dry” setting (sterilization temperature 274 °F) and used to line aluminum pans under the bat houses at each site. Fresh pellets were collected from the foil within 1 minute of falling using sterilized forceps, immediately placed on dry ice, and stored at −80 °C until processing. Each sample consisted of six pellets; three biological replicates were collected during each monthly sampling and stored at −80 °C until processing.

**Figure 1:**
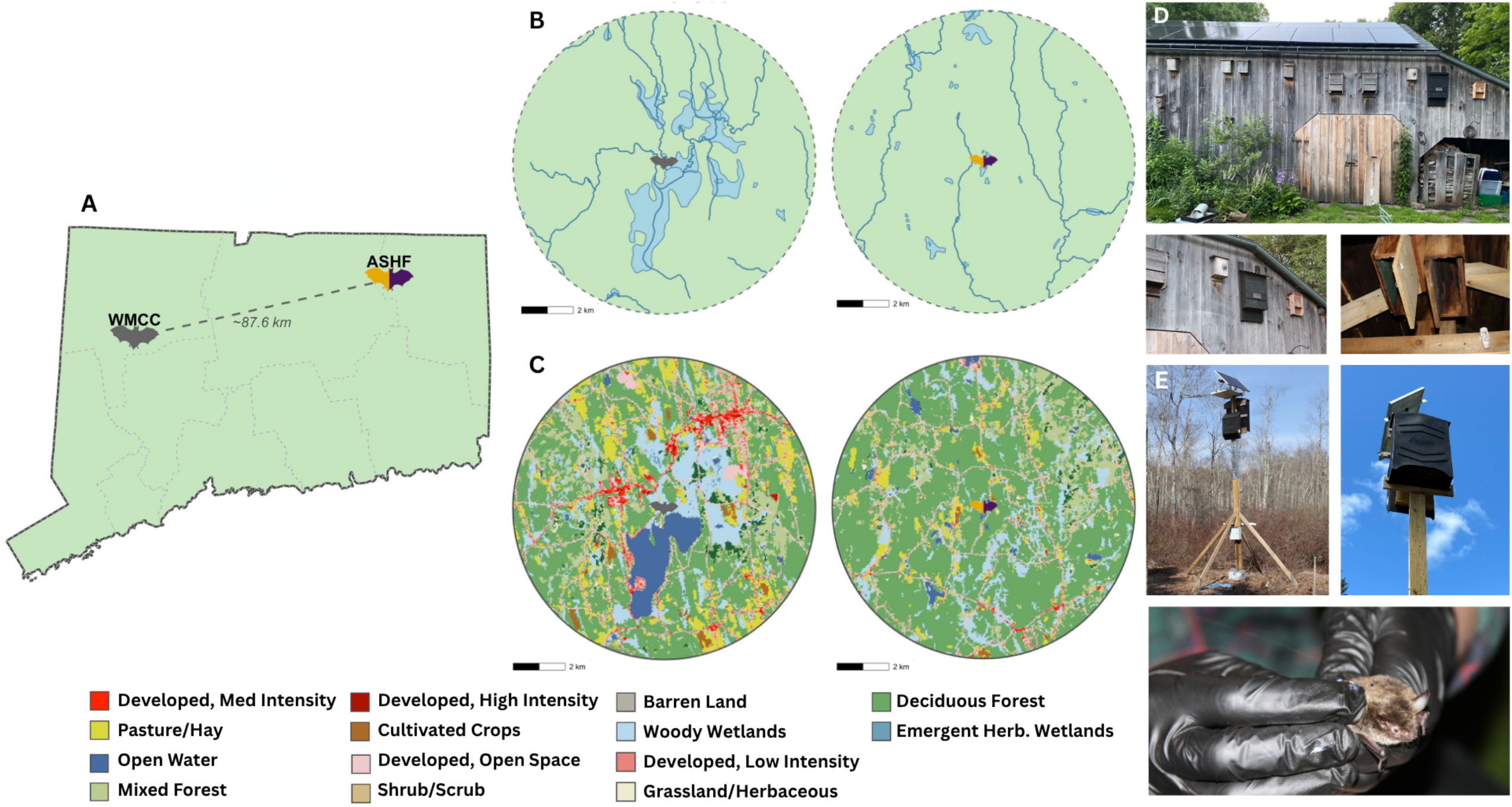
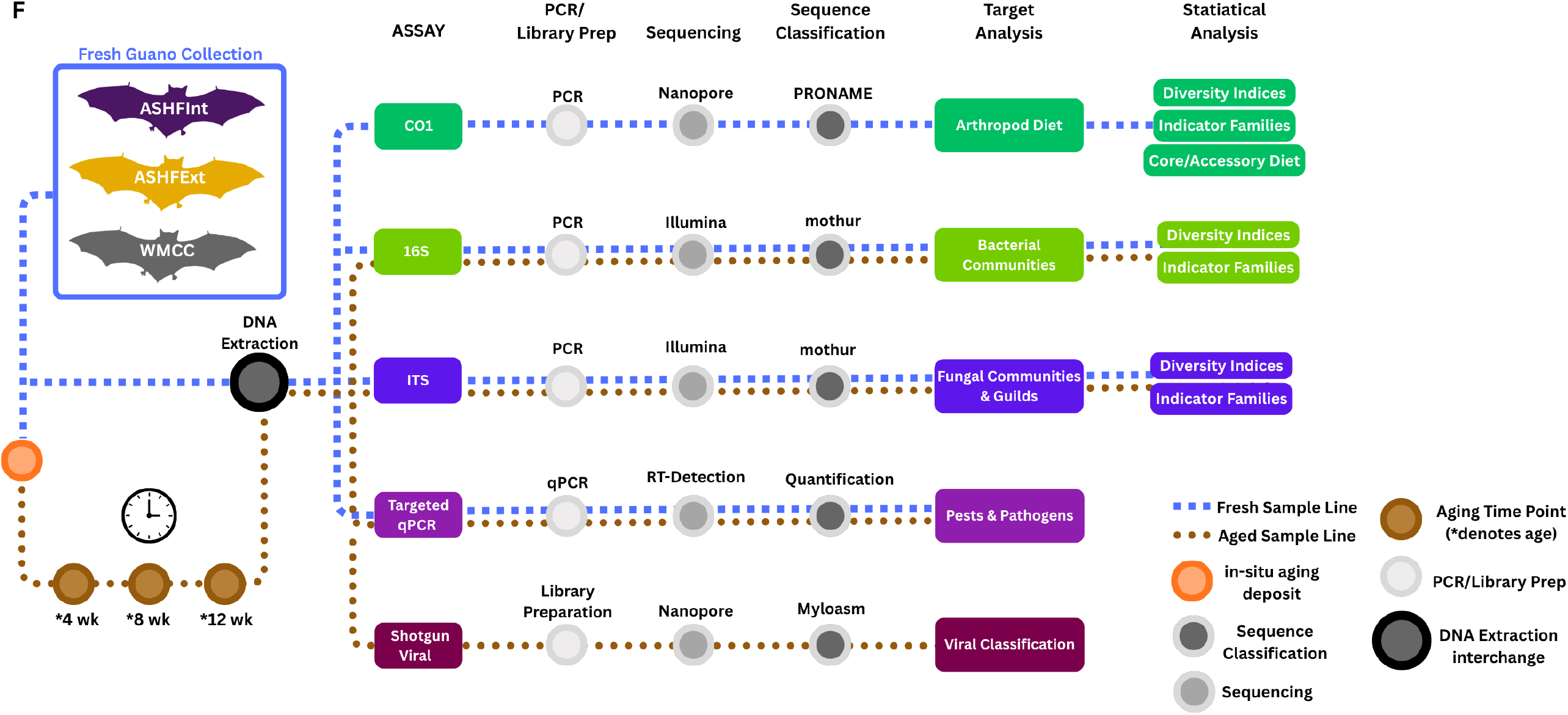
Study site locations and characteristics. **(A)** Map of Connecticut, USA, showing the two collection locations separated by ∼87.6 km. **(B)** Hydrographic features within a 5 km radius of WMCC (left) and ASHF (right), sourced from the National Hydrography Dataset. **(C)** Land cover composition within a 5 km radius of WMCC (left) and ASHF (right), derived from the 2021 National Land Cover Database. **(D)** Photographs of bat roost structures: ASHF barn (top), ASHFExt bat house (left), ASHFInt bat roost (right), **(E)** WMCC dual-roost bat house (left), WMCC close-up (right), and a little brown bat (*Myotis lucifugus*) (bottom). **(F)** Experimental Design Overview: Analytical pipeline for bat guano processing. Samples underwent five assays: CO1 metabarcoding (arthropod diet), 16S (bacterial communities), ITS (fungal communities and guilds), qPCR (pathogen detection), and shotgun viral metagenomics (viral classification). Each assay progressed through PCR/library preparation, sequencing, and sequence classification. Beta diversity and indicator taxa analyses were performed for CO1, 16S, and ITS; CO1 also included core and accessory diet classification.

*In situ* aging series were established in both June and August. After initial fresh collection, guano remaining in the foil trays was transferred to open sterile mesh boxes and left undisturbed at the collection sites to age naturally. Aged guano samples were collected at Weeks 4, 8, and 12 of aging (Fig. 1F). Because of low bat population, the August aged ASHFInt samples were collected only in Week 4; aged samples were analyzed by duration rather than by June or August deposition, so ASHFInt Week 8 and Week 12 samples remain represented. Like fresh samples, each aged sample included six pellets.

### 16S/ITS/CO1: DNA extraction, library preparation, and high throughput sequencing

DNA from approximately 20 mg of collected guano (approximately 3 pellets) was extracted using the MagAttract PowerSoil DNA kit (Qiagen) on the Kingfisher Apex processing robot. DNA was quantified with the Quant-iT PicoGreen kit (Invitrogen, ThermoFisher Scientific). The V4 region of the bacterial 16S rRNA was amplified using the 515F and 806R primers with Illumina adapters and dual indices (8 base pairs) (Kozich et al. 2013), using 30 ng of extracted DNA as a template. Fungal ITS2 regions were amplified utilizing the fITS7 (Ihrmark et al. 2012) and ITS4 (White et al. 1990) primers with the same dual indexing design. Both the 16S and ITS samples were multiplexed and sequenced on the Illumina MiSeq (V2 2X250). Amplification and regents are described in detail in File S1.

The CO1 region was amplified using long-read compatible primers (Table S4). The amplification reaction master-mix composition and thermocycler conditions implemented in this study are detailed in File S1. The PCR products were bead-cleaned twice prior to barcoding and library preparation. For barcoding, the Oxford Nanopore Technologies (ONT) Native Barcoding & Ligation kit (SQK-NBD114-96) was used following the manufacturer’s protocol. The manufacturer’s Ligation Sequencing Amplicons protocol included with the kit was also followed. CO1 PCR-ligation products were multiplexed and sequenced on a single GridION Flow Cell (R10.4).

### Shotgun metagenomics: DNA extraction, library preparation, and nanopore sequencing

DNA was quantified prior to library preparation for shotgun metagenomic sequencing. Only 48/86 samples had enough DNA to proceed. Using the same DNA extracts as were used for amplicon sequencing, sample libraries were prepared with the ONT Native Barcoding and Ligation kit (SQK-NBD114-96) for R10.4 flow-cells. Native barcodes 1 through 24 were used, resulting in two sample sets. Modifications were made to the manufacturer’s protocol to optimize barcode ligation and final library yield (File S1). Two pools (one for each set of barcodes 1–24) were made using equal volumes of each library and used for “screen” sequencing runs on the GridION. Read depth in these runs was used to calculate sample volumes that would yield pools with equal coverage. Table S1 provides details on sample input balancing during pooling. These pools were then sequenced on PromethION flow cells.

### 16S/ITS: OTU classification

The 16S and ITS Illumina short reads were independently clustered into operational taxonomic units (OTUs) with mothur (v1.48.3; Schloss et al. 2009) using the RDP taxonomy database as a reference (v19; Wang & Cole 2024). Before clustering, sequences were screened to remove ambiguous bases and were constrained to a maximum sequence length of 275 bp for 16S and 400 bp for ITS. Unique sequences were aligned to the SILVA database (SSU 138 v4) for 16S (Chuvochina et al. 2026) and the UNITE dynamic ITS reference database (v6) for ITS. Sequences that did not align or that had a homopolymer length above 8 were removed. Chimeric sequences were identified using vsearch on sequences sorted by abundance; sequences flagged as potential chimeras were removed from all samples, and the remaining sequences were classified using the respective 16S and ITS databases for reference and taxonomy with a cutoff bootstrap value of 80 for 16S and 60 for ITS. Final filters included plastids and other non-target alignments. Reads were clustered into OTUs based on uncorrected pairwise distances. The cutoff distance was 0.03 for 16S and 0.05 for ITS and ends were excluded using OptiClust (Westcott & Schloss 2017). See Tables S2 and S3 for mothur pipeline statistics for 16S and ITS respectively. Both alpha and beta diversity indices were calculated with mothur using a subsample of 10,000 reads for 16S. ITS was subsampled to 2,000 reads to retain the maximum number of samples for rarefaction given wider sample-to-sample variation in ITS read depth, while still capturing the dominant fungal taxa. Metrics included observed richness (OTUs), Shannon index, inverse Simpson, Bray-Curtis dissimilarity, Yue & Clayton dissimilarity (theta-YC), and Jaccard index. RStudio (v4.5.1) was employed for data visualization.

To estimate the proportion of aged bacterial OTUs that originated in the fresh samples, SourceTracker2 (Knights et al. 2011) was run on a subsample of 10,000 reads to allow for rarefaction. Fresh replicates were combined into a single source for each aging series (June and August), and analyses were performed separately for each site and aging series for a total of six analyses.

To assign ecological guild and trophic modes to fungal OTUs, FUNGuild (v1.1; Nguyen et al. 2016) was applied to the taxonomic assignments from mothur. To analyze the assigned guilds, relative abundances were calculated and visualized in R for guilds across sites, age status, and months.

### 16S/ITS: Diversity, community composition and indicator taxa analysis

To test for differences in bacterial alpha diversity between samples, ANOVAs were run using both alpha diversity indices (Shannon and Inverse Simpson). For fresh samples, a two-factor ANOVA with site and month as factors (including their interaction) was used to evaluate differences across these dimensions. Subsequently, the full dataset was used to test for effects of age, using a site:age model (collapsing months, which did not affect diversity of fresh samples). Site-specific decomposition effects were then assessed via per-site ANOVAs comparing fresh and aged timepoints. For significant ANOVA results, Tukey’s Honestly Significant Difference test (TukeyHSD) was used to identify significant pairwise differences in R. This workflow was repeated for the fungal (ITS) data.

Nonmetric multidimensional scaling (NMDS) based on the beta diversity distance matrices was used to visualize compositional similarity of communities across samples. To test for differences associated with sample age, month, and the interaction of site with each other variable, Permutational Multivariate Analysis of Variance (perMANOVA; Anderson 2001) models were run using the three beta diversity indices (Bray-Curtis dissimilarity, theta-YC, and Jaccard index). To test for interactions of month and age with site, we also reported results from the site:age model, while results for month are reported from the site:month model run on only fresh samples. perMANOVAs were run using the vegan package in R (v2.7-2).

To test for associations between specific OTUs in the bacterial and fungal communities and other sample features, we used MaAsLin3 (v1.1.2; Nickols et al. 2026) run with default parameters. Covariates of interest included site, age, month, the interaction of site and age, and the interaction of site and month. Age and month, as ordinal variables, were run with level contrasts so that comparisons were made to the previous timepoint rather than to a single baseline. Site, which has no inherent ordering, was run with reference-level contrasts using ASHFInt as the reference. Instead of subsampling a fixed number of reads, number of reads was included as a covariate; although MaAsLin3 normalizes read depth, samples with fewer than 1,000 reads were excluded prior to analysis to avoid spurious associations arising from extremely low sequencing depth. The three biological replicates collected per site each month were treated as a random effect to account for within-group variation.

To determine the indicator taxa from 16S and ITS community matrices, Indicspecies (v1.8.0; De Cáceres et al. 2010) was run on each site, and combinations of two of the three sites. This analysis was run separately on fresh and aged samples, including samples with a minimum of 10,000 reads. The multipatt command returns the square root of the indicator value with 999 permutations.

### CO1 Metabarcoding: OTU classification, diversity indices and indicator taxa analysis

Nanopore long reads from the fresh samples only were imported, and primer and adapter trimming were enabled via Dorado (v0.9.0) within PRONAME (v2.1.4; Dubois et al. 2024). The CO1 PRONAME run included 43 fresh samples collected in May through September 2025 (ASHFInt n = 12, ASHFExt n = 15, WMCC n = 15). The sequencing kit ID was set to SQK-NBD114-96, and the forward and reverse primer sequences were provided (Table S4). Simplex reads were filtered to remove reads with quality scores below 12 and reads outside of 350-950 bp range to ensure the full length 650 bp CO1 region was captured. Reads were clustered with a similarity threshold of 0.95 and read level error correction was performed with medaka model (v2.1.1) r1041_e82_400bps_sup_v5.0.0. Chimeras were detected using the de novo (reference-free) method and removed.

To perform the taxonomic classification, a custom Barcode of Life (BOLD) (Ratnasingham et al. 2024) database was generated for NCBI BLAST. The database consisted of arthropods collected from selected eastern states (ME, NH, VT, MA, RI, CT, NY, NJ, PA, DE, MD, WV, DC, VA, NC, SC, GA, and FL) and provinces (NL, QB, NB, and NS) of the U.S. and Canada. The database included 85,630 CO1-5P nucleotide sequences from 17,104 arthropod taxa. Sequences that contained gaps or had >1% of Ns were removed. In addition, CD-HIT (v4.8.1; Fu et al. 2012) was used to deduplicate the remaining sequences. NCBI BLAST (v2.15.0) performed sequence similarity search against the BOLD database (max target sequences 10, query coverage percentage 80, percent identity 70, e-value 1e-6).

Genus-level abundances were summed within family and order for higher rank analyses. The filtered CO1 dataset included 31 fresh samples (ASHFInt n = 12, ASHFExt n = 11, WMCC n = 8). Filters included a minimum read depth (100) as well as replicates per month/site. Reads classified only to the order or family rank but not further (coded as “Other”) were removed, and per-sample relative abundances were calculated across resolved taxa. Of the 90 detected families, 39 (43%) were retained for indicator analysis after applying a ≥10% prevalence filter (≥4 of 31 samples). Family-level NMDS analysis of Bray-Curtis, Jaccard, and theta-YC dissimilarities were computed in SciPy (v1.17.1; Virtanen et al. 2020) and ordinated by NMDS in scikit-learn (v1.8.0; Pedregosa et al. 2011) using 50 random initializations and a maximum of 2000 SMACOF iterations compared between the full 90 families and prevalence-filtered 39 family set. perMANOVA tested site and month effects under a single factor design for within-site and within-month designs, with 9,999 permutations. PERMDISP (Anderson et al. 2006) was run with every perMANOVA to distinguish centroid from dispersion effects. Indicator taxa analysis used the IndVal.g statistic (Dufrêne & Legendre 1997; De Cáceres & Legendre 2009) on the 39-family prevalence filtered subset under two contrasts (ASHFInt versus ASHFExt; ASHF versus WMCC) and per-month versus all months contrasts. P-values were corrected within each test using Benjamini-Hochberg FDR (Benjamini & Hochberg 1995) at FDR-corrected significance threshold p_critical = 0.10 (Storey 2003; Verhoeven et al. 2005; Pike 2011), with p_critical = 0.05 highlighted.

### Shotgun Metagenomics: viral identifications in aged samples

The high-throughput shotgun run was rebasecalled on Dorado (v1.4.0) using the model dna_r10.4.1_e8.2_400bps_sup@v5.2.0, then combined with the previous lower throughput run (rebasecalled on super high accuracy) to form the final read set for downstream analyses. Read quality, including number of reads, N50, and mean read length, was assessed with NanoComp (v1.25.6; De Coster & Rademakers 2023), and barcodes and primers were removed with Dorado “trim” using the kit ID SQK-NBD114-96. To identify and remove host-associated reads, trimmed reads were aligned with minimap2 (v2.28; default ONT parameters; Li 2021) to the *Myotis lucifugus carissima* reference genome (Vasquez et al. 2026), a high-quality reference for a subspecies of *M. lucifugus*. Reads without host alignment were retained using samtools (v1.21; Danecek et al. 2021) “-f 4,” and host alignment rate and read loss to host contamination were quantified with samtools “stats” and NanoComp, respectively.

Prior to assembly, reads from replicate samples were combined for co-assembly to maximize coverage and assembled with myloasm (v0.5.0; Shaw et al. 2026) using “--nano-r10” to optimize ONT R10 read assembly, with “--min-quality value-cutoff” restricting input to reads ≥97% accurate and “--min-reads-contig” set to 2 so that contigs were formed only when supported by at least two reads. For viral identification, read-level taxonomic classification was first performed with Centrifuge (v1.0.4.1; Kim et al. 2016) against the custom Centrifuge database inclusive of bacteria, fungi, archaea, and viruses, with reports visualized in Pavian (v1.0; Breitwieser & Salzberg 2020); assembled contigs from each aged sample were then analyzed with geNomad (v1.9.0; Camargo et al. 2024) under default parameters.

#### Targeted qPCR assays for human, bat, and invasive forest threats and spore count investigation

Targeted qPCR was used to identify and quantify the presence of *Histoplasma* spp *hp100, Blastomyces dermatitidis BAD1*, Pd 18S, spotted lanternfly (*Lycorma delicatula*) ITS, and emerald ash borer (*Agrilus planipennis*) CO1. Primers and fluorescent probes were designed as outlined in Table S4 and synthesized by Integrated DNA Technologies (IDT). Positive control gene fragments were synthesized by IDT to target desired amplicons. See File S1 for further details.

## RESULTS

### 16S: OTU Classification, diversity indices and indicator taxa analysis

On average, 35,102 16S reads were generated per sample. All samples were included in OTU clustering, but six samples were removed during rarefaction as they fell below the threshold of 10,000 reads. These samples were: ASHFExt n = 4 (one from July, two from August, and one Week 8 aged sample deposited in June) and WMCC n = 2 (one from May and one from September). Therefore, the final analytical dataset included 78 samples. Of these, 36 samples were fresh: ASHFInt n = 12 (three in May, June, July, and August), ASHFExt n = 11 (two in May, three in June, two in July, one in August, three in September), WMCC n = 13 (two in May, three in June, July, and August, and two in September). A total of 42 final samples were aged: ASHFInt n = 11 (five in Week 4, and three in Week 8 and 12), ASHFExt n = 15 (six in Week 4, six in Week 8, and three in Week 12), WMCC n = 16 (six in Week 4, seven in Week 8, and three in Week 12) (Table S5). After clustering, 8,381 OTUs were retained for 16S.

ANOVA tests indicated significant effects of sample age (Shannon: *F* = 63.65 *df* = 3, *p* ≤ 2e-16; Inverse Simpson: *F* = 34.227, *df* = 3, *p* ≤ 1.34e-13), site (Shannon: *F* = 26.65, *df* = 2, *p* ≤ 2.84e-9; Inverse Simpson: *F* = 12.675, *df* = 2, *p* ≤ 2.10e-5), and their interaction (Shannon: *F* = 7.22 *df* = 6, *p* ≤ 5.5e-6; Inverse Simpson: *F* = 5.141, *df* = 6, *p* ≤ 2.09e-4) on Shannon and Inverse Simpson diversity (Table S6). Tukey tests revealed a significant difference in diversity between aged and fresh samples at WMCC and ASHFExt, but neither differed significantly from ASHFInt (Fig. 2A, B). Within aged samples, further decomposition caused no significant change in diversity at any site. There was no significant difference in diversity between aged samples from exterior sites; however, there was a significant difference between aged samples from each exterior site and ASHFInt (Table S6). Within fresh samples, there was no significant effect of month (Shannon: *F* = 0.158, *df* = 3, *p* = 0.923; Inverse Simpson: *F* = 0.518 *df* = 3, *p* = 0.675), site (Shannon: *F* = 0.650, *df* = 2, *p* = 0.533; Inverse Simpson: *F* = 0.195, *df* = 2, *p* = 0.824), or their interaction (Shannon: *F* = 0.576, *df* = 6, *p* = 0.745; Inverse Simpson: *F* = 0.735, *df* = 6, *p* = 0.628) on diversity.

**Figure 2:**
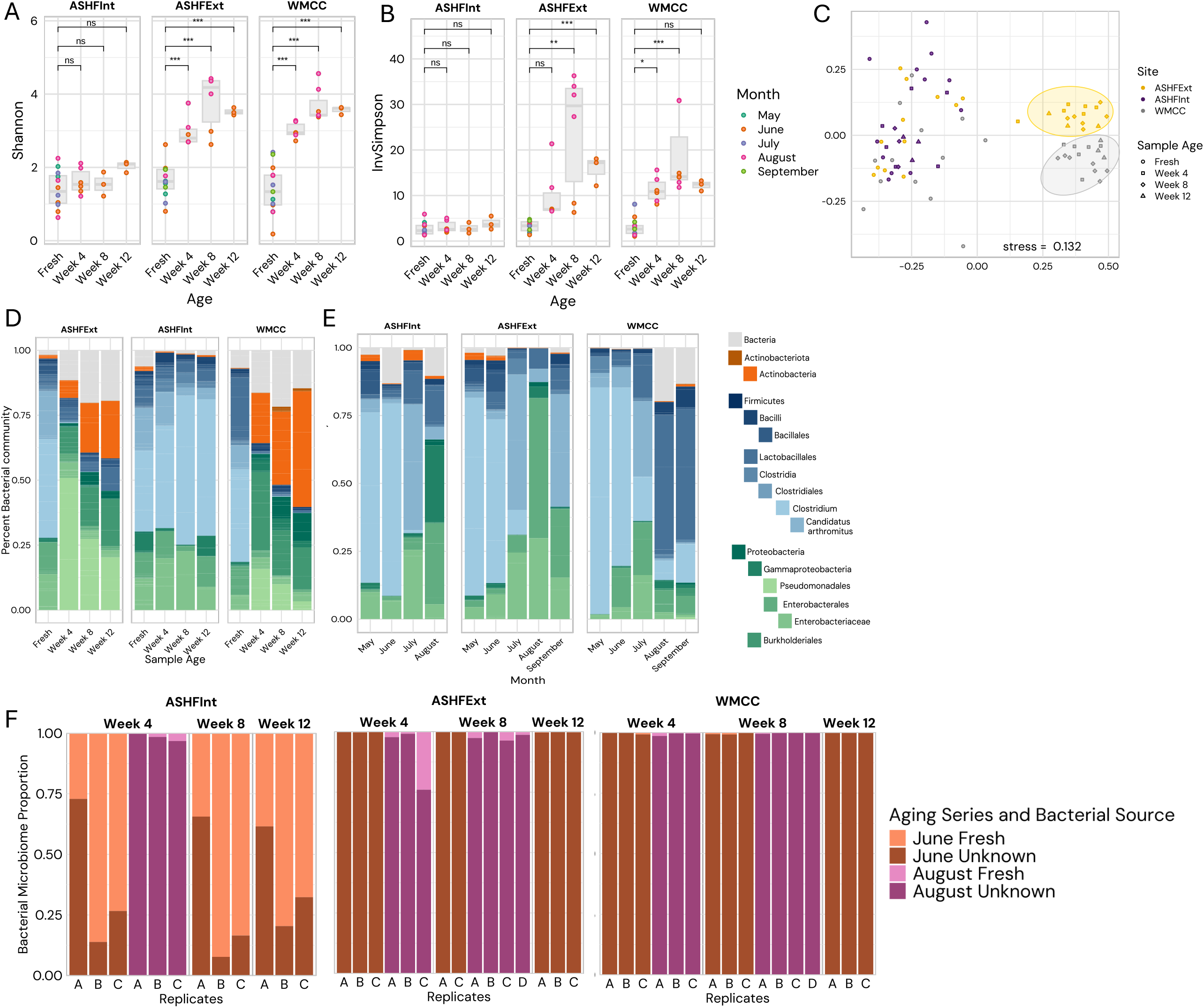
16S alpha and beta diversity, relative abundance by site and month, and bacterial source proportion in aged samples. **(A)** Shannon diversity across the aging series (Fresh, Week 4, Week 8, and Week 12) for each of the three sites. Colored dots represent the month in which the guano was deposited. Significance indicated as: * p < 0.05, ** p < 0.01, and *** p < 0.001, ns (not significant). **(B)** Changes in 16S inverse Simpson diversity are shown by site and age (Fresh, Week 4, Week 8, and Week 12). Colored dots represent the month in which the guano was deposited. **(C)** NMDS analysis of the bacterial Bray-Curtis index is shown to illustrate the similarity of the bacterial compositions for both site and age. Colored points represent the sites, while shapes indicate the age of the sample. **(D)** Bacterial (16s) taxonomic distribution and relative abundances across site and age. Represented taxa are organized by their taxonomic rank. The class, order, family, genus, and species (if applicable) are represented as indented under their respective phyla. Gray bars indicate all taxa not represented on the plot. **(E)** A fresh only bacterial (16S) taxonomic distribution is shown by site and month. **(F)** For each aging series (June and August), and site (ASHFInt, ASHFExt, and WMCC) fresh sources represent the bacteria that were present in the fresh samples from that month while unknown sources make up all other bacteria (i.e., bacteria present in an aged sample that was not present when it was collected). Fresh samples are grouped by replicates into a single source per aging series, while aged replicates are analyzed individually.

NMDS using Bray-Curtis beta diversity distances separated samples into three clusters, indicating substantial variation in community composition across groups of samples. Aged samples from WMCC and ASHFExt formed separate clusters, while a more widespread third cluster included all fresh samples along with ASHFInt aged samples (Fig. 2C). NMDS with Jaccard distances displayed a slight separation between WMCC fresh and ASHF fresh samples that was not present in when using Bray-Curtis or theta-YC; this separation likely indicates a difference in rare species between WMCC and ASHF, as the Jaccard index only considers species richness, which can provide a heavier weight to rare species compared to the other two indices given that they also account for abundance (Fig. S1C).

perMANOVA confirmed significant effects of site (Jaccard: *F* = 5.36, *df* = 2, *R*^2^ = 0.106, *p* = 0.001; Bray-Curtis: *F* = 8.76, *df* = 2, *R*^2^ = 0.141, *p* = 0.001), age (Jaccard: *F* = 3.42, *df* = 3, *R*^2^ = 0.102, *p* = 0.001; Bray-Curtis: *F* = 6.21, *df* = 3, *R*^2^ = 0.150, *p* = 0.001), and their interaction (Jaccard site:age: *F* = 2.00, *df* = 6, *R*^2^ = 0.119, *p* = 0.001; Bray-Curtis: *F* = 3.35, *df* = 7) (Fig. S2A; Fig. S2B) on bacterial community composition. Among fresh samples (the month and site:month tests), bacterial community composition also varied significantly by month under Bray-Curtis (month: *F* = 5.56, *df* = 3, *R*^2^ = 0.379, *p* = 0.001) and Jaccard (*F* = 1.66, *df* = 3, *R*^2^ = 0.136, *p* = 0.001) distances, with a significant site:month interaction under Jaccard (*F* = 1.22, *df* = 6, *R*^2^ = 0.202, *p* = 0.003) but not Bray-Curtis (*F* = 0.86, *df* = 6, *p* = 0.765; Table S7) distances (Table S7; Fig. S2A; Fig. S2B).

Fresh samples shared similar bacterial compositions when comparing the relative abundance of the most abundant taxa across all sites and comparing fresh to aged samples (Fig. 2D; Table S8). *Clostridiaceae, Enterobacteriaceae*, and *Enteroccaceae* were the most abundant families overall. ASHFExt and WMCC shared a similar pattern of diversity change between aged and fresh samples, with both seeing a loss of *Clostridiaceae* and an increase in *Oxalobacteraceae*. ASHFInt aged samples showed no change over time, maintaining the same abundant families in fresh and aged samples. The perMANOVA terms (site, age, and their interaction) together accounted for approximately 45% (Bray-Curtis) and 33% (Jaccard) of bacterial community variation (Table S7); fresh samples clustered tightly across all three sites, while aged samples diverged by site, with WMCC and ASHFExt forming distinct clusters separated from the fresh-sample cluster and from each other (Fig. 2C; Fig. S1C; Fig. S1D). Across all three sites, *Clostridiaceae* was the only family with high relative abundance in fresh guano (44.75-47.83%). *Enterobacteriaceae* abundance was high at both ASHF sites (17.66-17.75%), while *Enterococcaceae* remained relatively high only at WMCC (15.10%) (Fig. 2E; Table S8). Aged samples diverged sharply between interior and exterior sites: aged ASHFInt retained *Clostridiaceae* (55.83%) and *Enterobacteriaceae* (19.50%) as dominant, whereas aged ASHFExt was dominated by *Moraxellaceae* (30.31%) and *Oxalobacteraceae* (11.49%), and aged WMCC by *Nocardiaceae* (19.95%) and *Oxalobacteraceae* (17.85%). Full family-level relative abundances are tabulated in Table S8 (Fig. 2C).

In both exterior sites, fresh sample bacteria accounted for less than 5% of aged bacterial OTUs in all replicates except one, often accounting for less than 1% of aged OTUs. In ASHFInt samples from June, fresh sample bacteria accounted for between 27% and 92% of aged OTUs (Fig. 2F). ASHFInt August samples showed less than 5% input from fresh guano, similar to externally aged samples; however, these samples only had one fresh replicate for comparison due to rarefaction (Fig. 2F).

There were bacterial indicators for each site in the aged samples, along with several indicators of exterior sites and some combined ASHF indicators. Fresh samples had only one indicator unique to ASHFInt, while other indicators represented WMCC, combined ASHF, or exterior sites. *Corynebacteriaceae, Corynebacteriales* unclassified, and *Bogoriellaceae* were indicators of the combined ASHF sites in both fresh and aged samples (*Corynebacteriaceae* fresh: IndVal stat = 0.882, *p* = 0.029; aged: stat = 0.793, *p* = 0.002; *Bogoriellaceae* fresh: stat = 0.791, *p* = 0.004; aged: stat = 0.816, *p* = 0.001), while *Burkholderiaceae* was an indicator of WMCC in fresh and aged samples (fresh: stat = 0.562, *p* = 0.040; aged: stat = 0.962, *p* = 0.001). Five bacterial families, including *Lachnospiraceae* (aged ASHFInt: stat = 0.941, *p* = 0.001; fresh combined exterior: stat = 0.885, *p* = 0.011), serve as indicators for different sites in fresh versus aged samples (Fig. 3A; Table S9).

**Figure 3:**
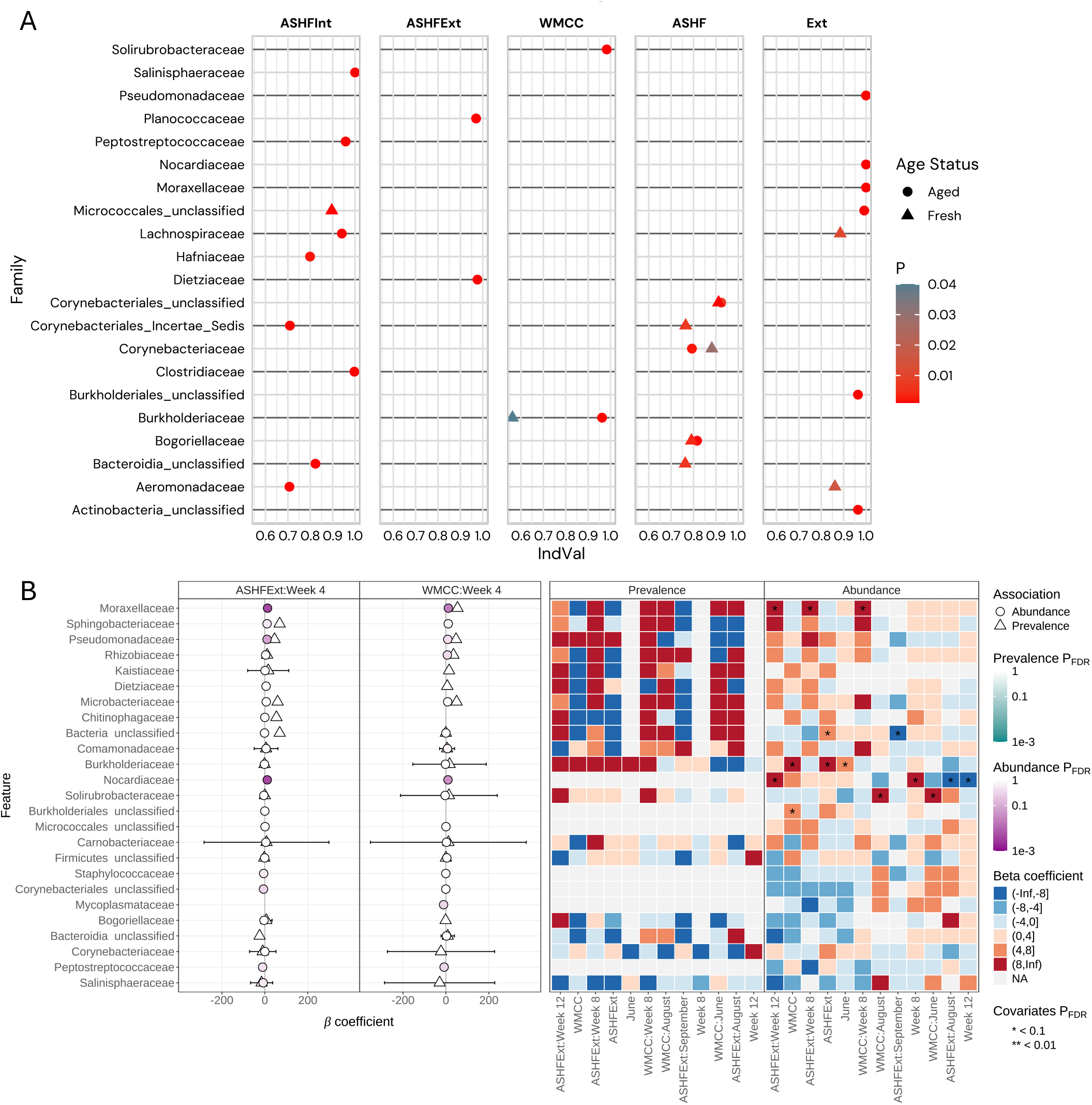
16S Indicator taxon and site associations. **(A)** Subset of significant bacterial family site indicators, separated by aged and fresh samples. ASHF represents indicators of ASHFInt+ASHFExt and Ext represents indicators of ASHFExt+WMCC. **(B)** Statistically significant relationships between taxonomic families and site metadata conducted with MaAsLin3. The two most significant interactions are shown in the left panel, ASHFExt vs. Week 4 and WMCC vs. Week 4. A single asterisk in the heat beta-coefficient panel indicates p < 0.1 and two asterisks indicate p < 0.01.

The abundances of five bacterial families (*Moraxellaceae, Nocardiaceae, Burkholderiaceae, Pseudomonadaceae*, and *Solirubrobacteraceae*) and two unclassified groups (*Burkholderiales* unclassified and *Bacteria* unclassified) were significantly associated with sample sites and characteristics. Of these families with significant associations, all but *Pseudomonadaceae* were significantly associated with more than one sample characteristic, often across multiple sites. There were no significant associations between bacterial family prevalence and sample characteristics; the lack of significance in prevalence statistics may be due to sample size (Fig. 3B).

### ITS: OTU classification, diversity indices and indicator taxa analysis

On average, 29,548 ITS reads were generated. All samples were included in OTU clustering, but four samples were removed during rarefaction as they fell below the threshold of 2,000 reads. The final analytical dataset included 82 samples. Of these, 38 samples were fresh: ASHFInt n = 12 (three in May, June, July, and August, and 0 in September), ASHFExt n = 11 (three in May and June, one in July, one in August, and three in September), WMCC n = 15 (three in all months). A total of 44 final samples were aged: ASHFInt n = 12 (six in Week 4, and three in Weeks 8 and 12), ASHFExt n = 16 (six in Week 4, seven in Week 8, and three in Week 12), WMCC n = 16 (six in Week 4, seven in Week 8, and three in Week 12). (Table S10). A total of 63,273 OTU clusters were retained for ITS. The fungal Shannon alpha diversity in fresh guano was higher than in the aged guano from WMCC and ASHFExt, suggesting more diversity in fresh guano communities than aged (Fig. 4A; Table S11).

**Figure 4:**
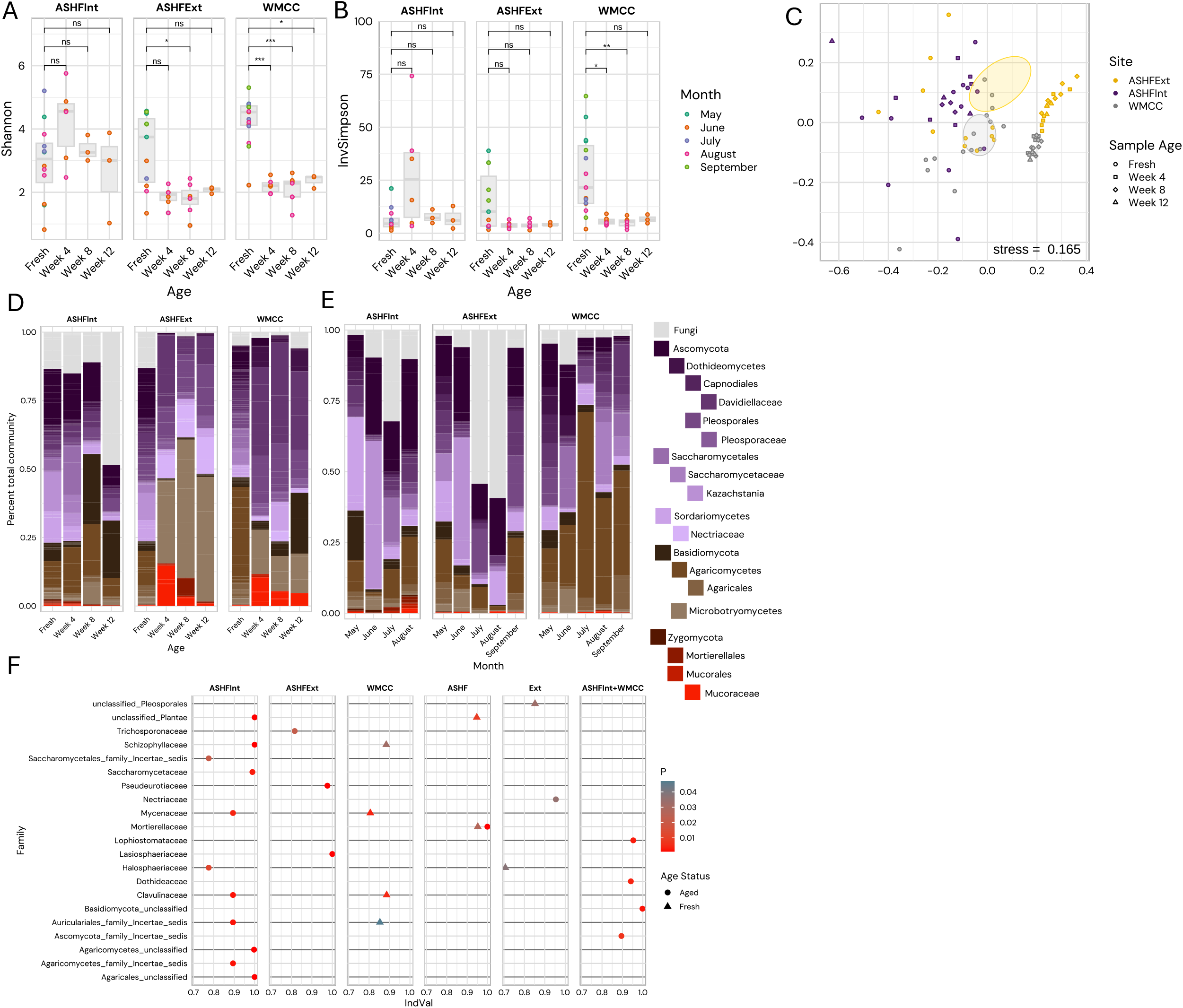
Fungal ITS Shannon diversity, inverse Simpson, Bray-Curtis dissimilarity and taxonomic distribution by site, month, and age. **(A)** Changes in Shannon diversity are shown by site and age (Fresh, Week 4, Week 8, and Week 12). Colored dots represent the month in which the guano was deposited. **(B)** Changes in inverse Simpson diversity are shown site and by age (Fresh, Week 4, Week 8, and Week 12). Color represents the month in which the guano was deposited. **(C)** The NMDS analysis of the fungal Bray-Curtis index is shown here to illustrate the similarity of the fungal compositions for both sites and aging. Colors represent the sites, while shapes indicate the age of the sample. **(D)** Fungal (ITS) taxonomic distribution and mean relative abundances across both sites and age (Fresh, Week 4, Week 8, Week 12). Represented taxa are organized by their taxonomic rank. The class, order, family, genus and species (if applicable) are represented as indented under their respective phyla. Gray bars indicate all taxa not represented on the plot. **(E)** Fungal (ITS) taxonomic distribution and mean relative abundances of fresh samples across both sites and month deposited. Represented taxa are organized by their taxonomic rank. The class, order, family, genus and species (if applicable) are indented under their respective phyla. Gray bars indicate all taxa not characterized. **(F)** Subset of significant fungal family site indicators, separated by Aged and Fresh samples. ASHF represents indicators of ASHFInt+ASHFExt and Ext represents indicators of ASHFExt+WMCC.

Among fresh samples, a two-factor ANOVA with site and month as factors detected significant differences in fresh sample fungal inverse Simpson diversity across sites (*F* = 16.170, *df* = 2, *p* ≤ 6.6e-5), across months (*F* = 12.073, *df* = 3, *p* ≤ 9.88e-5) but no significance in their interaction (*F* = 2.141, *df* = 6, *p* ≤ 0.0934; Table S11). Fungal Shannon and inverse Simpson diversity differed significantly by sample age (Shannon: *F* = 11.324, *df* = 3, *p* = 3.84e-6; Inverse Simpson: *F* = 4.531, *df* = 3, *p* = 0.006), site (Shannon: *F* = 7.018, *df* = 2, *p* = 0.002; Inverse Simpson: *F* = 2.679, *df* = 2, *p* = 0.076), and their interaction (Shannon: *F* = 6.476, *df* = 6, *p* = 1.82e-5; Inverse Simpson: *F* = 4.927, *df* = 6, *p* = 2.96e-4; Table S11). Tukey tests revealed significant declines in Shannon diversity from fresh to aged samples at WMCC (Week 4: *p* = 3.55e-04, Week 8: *p* = 4.15e-05, Week 12: *p =* 0.043) and ASHFExt (Week 4: *p* = 0.024), and in inverse Simpson diversity at WMCC (Week 4: *p* = 0.028, Week 8: *p =* 0.010; Table S11). In ASHFInt, Shannon diversity did not differ significantly between fresh and aged samples at any timepoint (Fig. 4; Table S11), although inverse Simpson diversity showed a strong increase from fresh to Week 4 (p = 0.031; Table S11). Inverse Simpson values in fresh guano at WMCC and ASHFInt reached 60–70, compared to aged samples falling below 15 (Fig. 4B). The diversity of fungal communities at WMCC started declining by Week 4 (the earliest aging time point), while ASHFExt showed a significant Shannon decline by Week 8. Following the decline both sites showed consistently low diversity of fungal communities.

The ITS Bray-Curtis NMDS shows two groups: ASHFInt (all ages) clusters with fresh ASHFExt and fresh WMCC. The other cluster consists of only aged (Week 4 to Week 12) ASHFxt and WMCC (Fig. 4C). The first group does not cluster very tightly, suggesting more diversity between sites. In contrast, the second group consisting of aged samples in ASHFExt and WMCC are much more clustered, indicating that over time the fungal communities become similar but still unique to their respective site. perMANOVA confirmed significant effects of site (Jaccard: *F* = 4.15, *df* = 2, *R*^2^ = 0.084, *p* = 0.001; Bray-Curtis: *F* = 7.76, *df* = 2, *R*^2^ = 0.135, *p* = 0.001), age (Jaccard: *F* = 3.16, *df* = 3, *R*^2^ = 0.096, *p* = 0.001; Bray-Curtis: *F* = 4.30, *df* = 3, *R*^2^ = 0.112, *p* = 0.001), and their interaction (Jaccard site:age: *F* = 1.88, *df* = 6, *R*^2^ = 0.114, *p* = 0.001; Bray-Curtis: *F* = 2.82, *df* = 6, *R*^2^ = 0.147, *p* = 0.001) on fungal community composition, with ASHFExt and WMCC showing more distinct compositional shifts than ASHFInt (Table S12).

Taxonomic relative abundance of the most abundant fungal taxa by family and genus was obtained across sites and between fresh and aged samples within each site (Fig. 4E; Table S13). Fresh samples from ASHFExt and ASHFInt were similar in fungal composition with *Saccharomycetaceae* (12.16% and 13.28% respective relative abundance) and unclassified *Ascomycota* being the most abundant taxa (10.24% and 15.46% relative abundances, respectively). WMCC’s most abundant family in fresh guano was *Peniophoraceae* (10.83%). Aged samples across all three testing sites were dominated by families within the phylum *Ascomycota* (53.65% at the ASHFInt site, 45.77% at the ASHFExt site, and 68.76% at WMCC) (Fig. 4D; Table S13). Aged ASHFInt samples show increasing relative abundance of the family *Trichocomaceae* across the aging series (Week 4 = 1.81%, Week 8 = 6.48%, Week 12 = 41.03%), a trend not observed at other sites. This trajectory was not formally tested at the family level since MaAsLin3 did not return significant fungal family:age associations (see below); the magnitude and site-specificity of the increase nonetheless warrant attention and are reflected in the family-level indicator analysis (Fig. 4F; Table S15). ASHFExt and WMCC exhibited similar aging profiles, with family *Davidiellaceae* remaining abundant across the decomposition process (WMCC: Week 4 = 32.31%, Week 8 = 36.79%, Week 12 = 29.13%; ASHFExt: Week 4 = 26.03%, Week 8 = 5.96%, Week 12 = 10.25%), although relative abundances were consistently higher at WMCC than at ASHFExt (Fig. 4D). In the exterior samples, the *Mucoromyceta* (*Mortierellales* and *Mucorales*) peaked at Week 4, then declined, but remained at low levels similar to fresh guano in ASHFInt (Fig. 4D).

Aged fungal samples contain many indicators of ASHFInt, but relatively few indicators of the other two sites individually, combined exterior sites, or combined ASHF sites. Aged fungal samples contained the only indicators of ASHFInt+WMCC. Fresh fungal samples contain no individual ASHF site indicators, only indicators of WMCC, ASHF combined, and exterior. *Mortierellaceae* is an indicator of the combined ASHF sites in both fresh and aged samples (fresh: IndVal stat = 0.952, *p* = 0.027; aged: stat = 0.999, *p* = 0.001), the only instance of shared fungal site indicators between fresh and aged samples. Six fungal families, including *Schizophyllaceae* (aged ASHFInt: stat = 1.000, *p* = 0.001; fresh WMCC: stat = 0.884, *p* = 0.032), serve as indicators for different sites in fresh versus aged samples (Fig. 4F; Table S15). Examination of the relationship between fungal family abundance and site characteristics did not reveal any significant associations in the MaAsLin3 analysis.

Guild composition varied across guano age status, site, and month (Fig. S3; Table S14). The composition of the five most abundant guilds differed between the age and fresh-month analyses. Across age status, fungal parasite-plant saprotroph was the dominant guild at ASHFExt, undefined saprotroph at ASHFInt, and animal pathogen-animal symbiotroph-endophyte-plant pathogen-undefined saprotroph at WMCC (Fig. S3A). Across the months of fresh guano, animal parasite-endophyte-undefined saprotroph-wood saprotroph was the dominant guild at both ASHF sites, whereas undefined saprotroph-wood saprotroph was most abundant at WMCC (Fig. S3B).

### CO1 Metabarcoding: OTU classification, diversity indices, and indicator taxa analysis

After demultiplexing, primer trimming, quality filtering, clustering at 95% similarity, medaka error correction, and chimera removal in PRONAME, samples meeting the minimum-replicate requirement (≥2 fresh samples per site per month, supporting site:month contrasts) were retained. The final analytical dataset comprised 31 fresh samples: ASHFInt n = 12 (three per month), ASHFExt n = 11 (missing one July replicate), and WMCC n = 8 (two per month). On average, 1,932 reads were generated per sample (median = 800, range 18–20,004). The ideal N50 for the 650 bp CO1 amplicon is 650–800 bp; the dataset average N50 was 606 bp (median = 665 bp, range 223–883 bp), indicating that most samples were suitable for downstream analysis (Table S16). Classification and relative abundance by genus, family, and order were indicated for each sample (Table S17-19).

Across the three sites, ASHFInt yielded the greatest number of arthropod orders unique to a single site, with five orders detected only at ASHFInt (the insect orders Megaloptera, Orthoptera, Neuroptera, and Odonata and the mite order Sarcoptiformes). WMCC contributed two unique orders (Psocodea, Plecoptera), and ASHFExt contributed none; every order at ASHFExt was also detected at one or both other sites (Fig. 5A; Fig. S4A). The same site ranking was observed at the family level (Fig. 5B; Fig. S4B): ASHFInt contained 25 unique families (including Apidae, Erebidae, Corydalidae, Tettigoniidae, Oripodidae, Hemerobiidae, and Libellulidae), WMCC contained seven (Aturidae, Buprestidae, Chloroperlidae, Crambidae, Hydroptilidae, Pallopteridae, Peripsocidae), and ASHFExt contained four (Caenidae, Clastopteridae, Gelechiidae, Nymphalidae). Sample size may contribute to the high uniqueness count at ASHFInt (n = 12 vs. n = 11 at ASHFExt and n = 8 at WMCC), but the gap is large enough to suggest a real difference.

**Figure 5:**
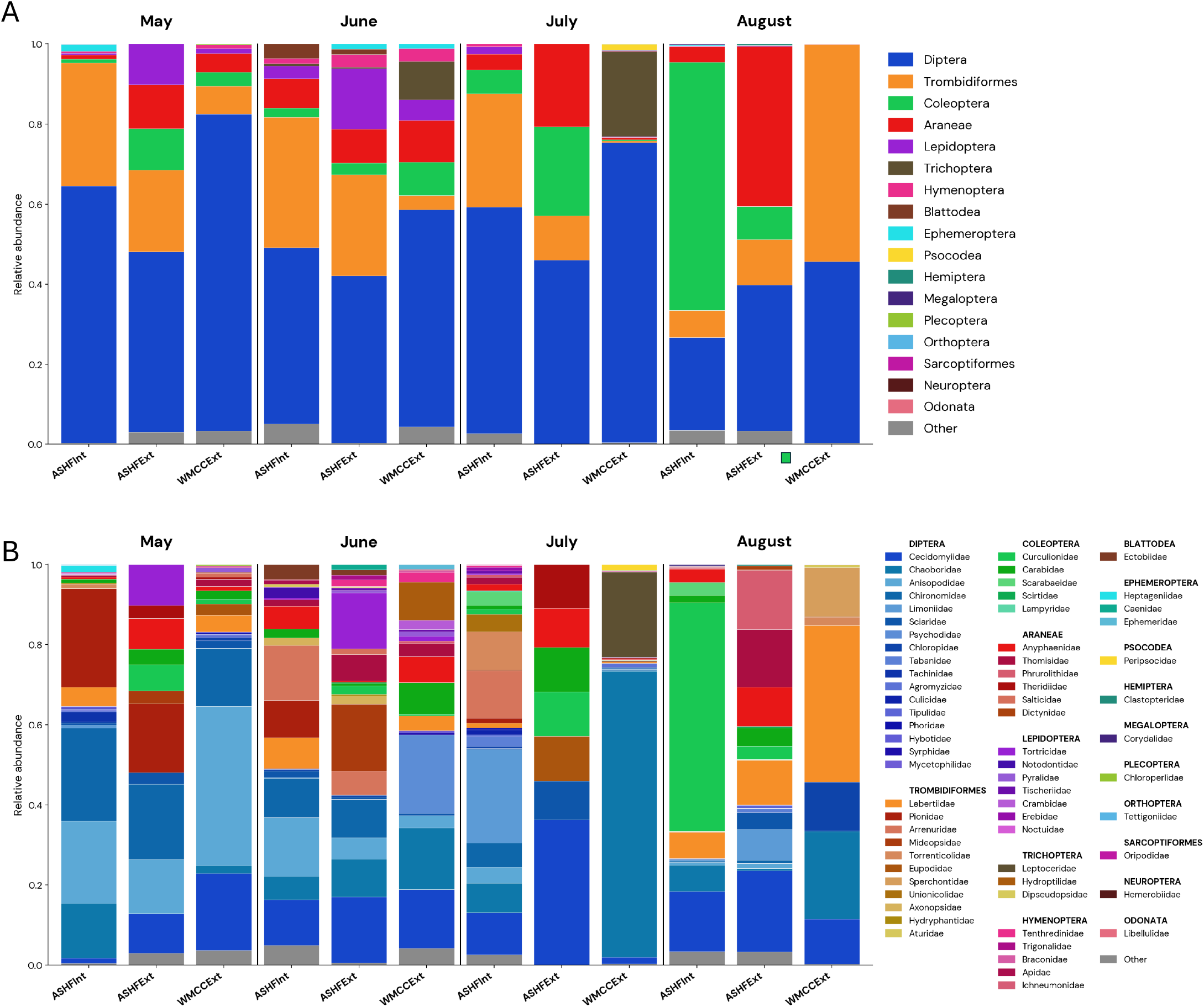
Order and family distributions by site and month. The relative abundance bar charts have the month on top, site on the bottom, and relative abundance on the left. **(A)** The top chart shows the relative abundance of the orders detected at each site from May to August. **(B)** The lower chart shows the relative abundance of the families detected at each site from May to August with the order that each family belongs to shown in bold.

Order-level richness varied across the May through August sampling window in site specific ways (Fig. S4B). At ASHFInt, richness was highest in May, declined to a low in July, and partially recovered in August. At ASHFExt, richness peaked in June, then declined through July to a minimum, before partially recovering in August. WMCC showed a strikingly different pattern: richness was similar in May and June, then peaked sharply in July before dropping again in August. The same site by month pattern was consistent at the family level (Fig. S4C), suggesting that the temporal differences are not artifacts of taxonomic resolution.

Diptera (flies) dominated relative dietary composition at all three sites in May, June, and July (Fig. 5A). In August, the dominant order was influenced by site: Coleoptera (beetles) at ASHFInt, Araneae (spiders) at ASHFExt, and Trombidiformes (one of the two mite orders) at WMCC. At the family level, the most abundant taxa included Chaoboridae (phantom midges) at WMCC in July, Curculionidae (weevils) at ASHFInt in August, and Lebertiidae (freshwater mites) at WMCC in August.

### CO1: Core and accessory diets

Dietary families were classified as core (detected in ≥80% of samples within a grouping) or accessory (detected in some but fewer than 80%) at three grouping levels: per site, per month, and per site:month cell (Fig. 6). Only one family (Cecidomyiidae, gall midges) was core at all three sites. ASHFInt had the largest core (five families: Anisopodidae, Anyphaenidae, Cecidomyiidae, Chaoboridae, Lebertiidae). WMCC had three (Cecidomyiidae, Chaoboridae, Lebertiidae), a subset of the ASHFInt core. ASHFExt had only Cecidomyiidae as core, with no private core families. The two aquatic-associated families shared between ASHFInt and WMCC (Chaoboridae, phantom midge larvae; Lebertiidae, freshwater mites) are obligate or near-obligate aquatic taxa, indicating that bats foraging from these two sites have reliable access to aquatic prey. The lack of core families observed in ASHFExt suggests potential differences in feeding behavior at this location.

**Figure 6:**
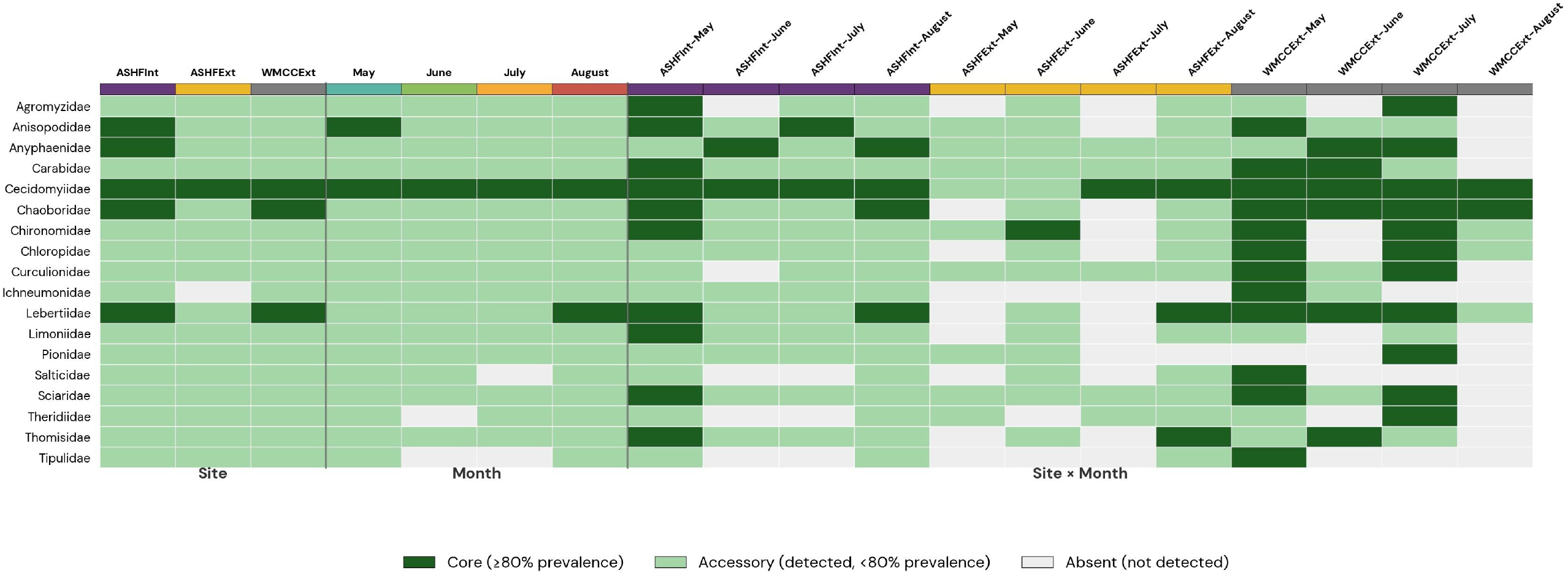
Core, accessory, and absent taxa across sites and months. Arthropod orders are represented on the y axis. Both site and month independently, and the interaction of site and month are represented on the x axis. Dark green, light green, and gray for each order indicate whether it is a core, accessory, or absent order respectively.

At the month level, only Cecidomyiidae was core in all four months. Anisopodidae was core in May only, and Lebertiidae only in August, the latter aligning with the late summer spike in trombidiform mites. Anyphaenidae and Chaoboridae were core at the site level but did not reach the ≥80% threshold within any single month (Fig. 6).

The mean per-sample relative abundance contributed by core families varied across sites. At WMCC the three-family core accounted for approximately 52% of mean per-sample relative abundance, at ASHFInt the five-family core contributed approximately 37%, and at ASHFExt the single-family core contributed only 20% (Fig. 6). This pattern is consistent with the within-site dispersion result from PERMDISP: ASHFExt has the broadest dietary heterogeneity of any site, with no small family core dominating individual samples.

Single-factor perMANOVA tested whether dietary composition differed among sites within each month, using both Bray-Curtis and Jaccard dissimilarities (Fig. S5A). Across the four months, only July showed statistically significant site differentiation, and only under Bray-Curtis (*F* = 2.31, *R*^2^ = 0.54, raw *p* = 0.010, BH-adjusted *q* = 0.039). May, June, and August did not differ by site under either metric. The July signal is consistent with the relative-abundance composition (Fig. 6B), in which July shows the most distinctive Chaoboridae signal at WMCC.

The complementary stratified analysis tested whether each site’s diet changed across the four months (Fig. S5B). WMCC showed clear temporal turnover under both metrics (Jaccard *F* = 1.42, *R*^2^ = 0.52, raw *p* = 0.016, *q* = 0.047 (passes *q* < 0.05); Bray–Curtis *F* = 1.73, *R*^2^ = 0.56, raw *p* = 0.019, *q* = 0.056 (passes *q* < 0.10)). The Jaccard signal indicates that the identity of taxa detected at WMCC shifted month to month (taxonomic turnover), while the Bray-Curtis signal indicates that the relative abundances of those taxa also shifted. Neither ASHFInt nor ASHFExt showed temporal turnover passing the FDR threshold under either metric, suggesting that the ASHF colony’s dietary composition is more stable across the sampling window than that at WMCC. Pairwise tests of WMCC’s temporal variation could not associate the difference to specific months.

PERMDISP testing identified heterogeneous within-group dispersion at the site:month level under both metrics (Bray-Curtis *p* = 0.009; Jaccard *p* = 0.005). At the site level, dispersion was homogeneous on Bray-Curtis (*p* = 0.673) and marginally heterogeneous on Jaccard (*p* = 0.043). ASHFExt had the widest Jaccard dispersion (site), consistent with the core diet finding that ASHFExt has no consistent dominant family across samples.

To identify families distinguishing the three sites, IndVal.g analysis was run on the 39 prevalence-filtered families under two contrasts. In the ASHF-only contrast (interior versus exterior, n = 23, WMCC excluded; Fig. S5C), two families showed raw *p* < 0.05, both favoring ASHFInt: Ichneumonidae (parasitoid wasps: IV.g = 0.71, raw *p* = 0.014, *q* = 0.534) and Chaoboridae (phantom midges: IV.g = 0.79, raw *p* = 0.037, *q* = 0.723). Neither family reached FDR significance.

In the across location contrast (ASHF combined versus WMCC, n = 31; Fig. S5D), two families showed raw *p* < 0.05, both favoring WMCC: Chaoboridae (IV.g = 0.91, *A* = 0.83, *B* = 1.00, raw *p* = 0.0013, *q* = 0.051) and Chloropidae (frit flies: IV.g = 0.77, raw *p* = 0.037, *q* = 0.716). The Chaoboridae signal at WMCC is significant: every WMCC sample contained Chaoboridae, and 83% of the family’s combined group mean relative abundance came from WMCC (specificity *A* = 0.83). Chaoboridae favors ASHFInt over ASHFExt and is the strongest indicator favoring WMCC over ASHF (combined); aquatic prey (Chaoboridae) are consistently shared by ASHFInt and WMCC, while ASHFExt is unique. This same pattern is visible in the core diet analysis, where Chaoboridae and Lebertiidae are core at ASHFInt and WMCC, but not at ASHFExt.

The same IndVal.g framework was applied to each month versus the other three months combined (Fig.s S5E, F). In May, two families showed raw *p* < 0.05, both favoring May: Anisopodidae (gnat-like flies: IV.g = 0.87, *A* = 0.87, *B* = 0.88, *p* = 0.0041, *q* = 0.161) and Chironomidae (non-biting midges: IV.g = 0.80, *p* = 0.033, *q* = 0.616). Neither reached the FDR threshold. In June, Ectobiidae (cockroaches) emerged as the leading indicator (IV.g = 0.71, *A* = 1.00, *B* = 0.50, *p* = 0.0022, *q* = 0.087), passing the *q* < 0.10 threshold. Specificity of 1.0 means every detection of Ectobiidae in the dataset came from a June sample, although fidelity of 0.50 indicates that Ectobiidae was detected in only half of the eight June samples. Axonopsidae (water mites) also showed a June signal (*p* = 0.042, *q* = 0.654). July had no families with raw *p* < 0.05. For August, three families showed raw *p* < 0.05: Lebertiidae (freshwater mites: IV.g = 0.87, *p* = 0.017, *q* = 0.352), Drosophilidae (vinegar flies: IV.g = 0.59, *p* = 0.022, *q* = 0.352), and Dictynidae (mesh-web spiders: IV.g = 0.60, *p* = 0.036, *q* = 0.352). Anisopodidae also showed a signal favoring May/June/July versus August (IV.g = 0.81, *p* = 0.032, *q* = 0.352), consistent with its early summer emergence (Fig. S5G). None passed the FDR threshold. The Lebertiidae result supports the core diet analysis, which identified this as the only family core to August samples.

### Targeted PCR assays for human, bat, and invasive forest threats and spore count investigation

Five organisms were targeted for qPCR: *Histoplasma* spp., *Blastomyces dermatitidis*, Pd, spotted lanternfly (*Lycorma delicatula*), and emerald ash borer (*Agrilus planipennis*). Presence was defined as detection in at least two of the three replicates for a sample.

*Pseudogymnoascus destructans* (Pd), the fungus causing white-nose syndrome, was detected in a high proportion of samples (31 in total) (Fig. 7A; Fig. S6A). Of these, 19 were freshly collected (45.2% of 42 total fresh samples), eight were collected after four weeks of aging (44.4% of 18 total four week aged samples), four were collected after eight weeks of aging (23.5% of 17 total eight week aged samples), and one was collected after 12 weeks (11.1% of nine total 12 week aged samples). Pd was most prevalent in ASHFInt samples (n = 11), followed by WMCC (n = 10) and ASHFExt samples (n = 10). Pd was most abundant in fresh samples collected in May (n = 8).

**Figure. 7:**
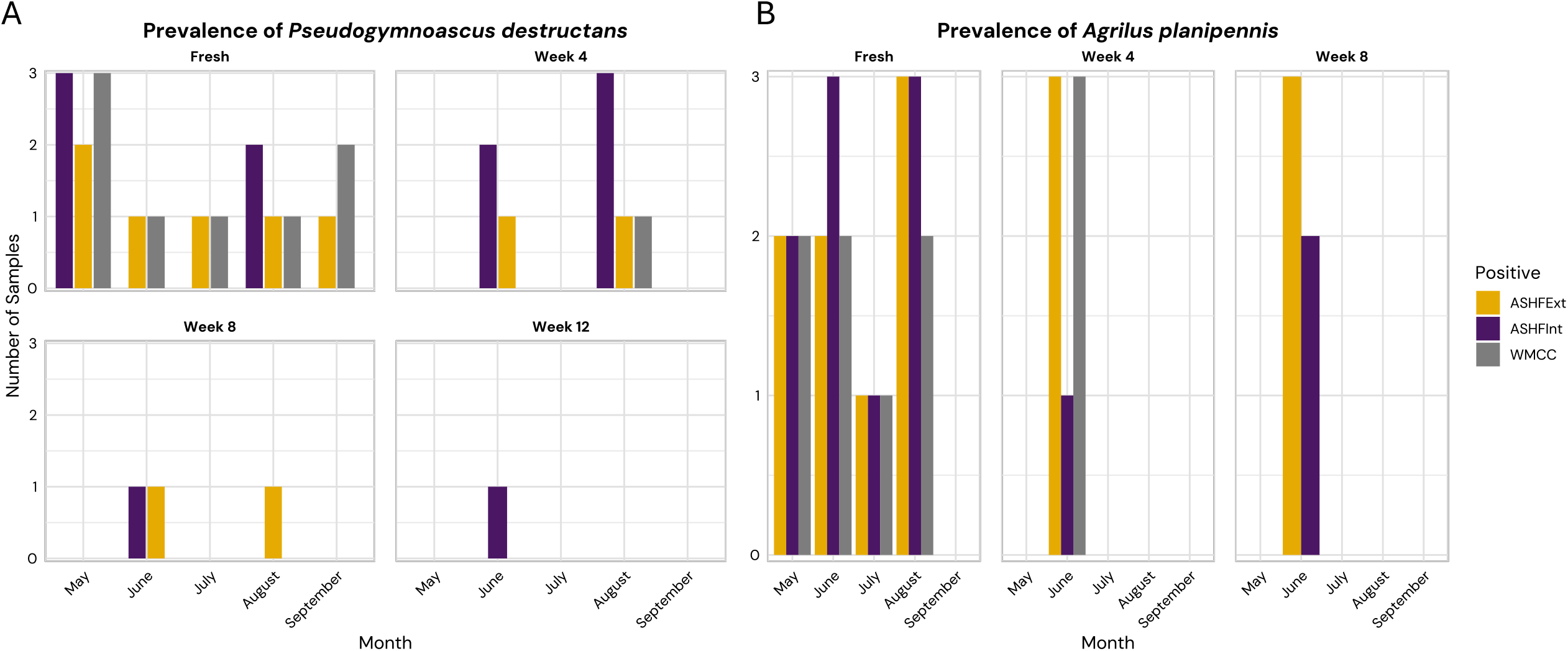
Prevalence of Pd and A. planipennis. **(A)** The presence of Pd, as defined by presence in two out of three triplicate copies of a sample, is displayed according to site, month and age. Color corresponds to the site at which a sample was collected. Week 4 and Week 8 samples were only collected in June and August, and Week 12 samples were collected in September. **(B)** The presence of *A. planipennis*, as defined by presence in two out of three triplicate copies of a sample, is displayed according to site, month and age. Color corresponds to the site at which a sample was collected. Week 4 and Week 8 samples were only collected in June and August. *A. planipennis* was not detected in Week 12 samples.

Results of 2024 spore counts at the sampling sites indicate the presence of Pd spores in nearly every swabbing site surveyed (Fig. S6B). Our findings suggest that Pd was present in most collected samples throughout the summer months, with highest counts present in ASHFExt bat houses in May and lowest counts found at all sites in August (Fig. S6B). Culturing attempts did not result in Pd growth; thus, viability of collected conidia was not confirmed.

Three species within the genus *Pseudomonas* have been previously identified as possible inhibitors of Pd growth (Li Z. et al. 2021). Although non-specific, the analysis of bacterial composition did identify a high abundance of bacteria within the order Pseudomonadales, especially within the Week 4 aged category (Fig. 2D). A total of 29 additional strains of bacteria with growth inhibiting effects on Pd were also identified within the Pseudomonadales genera *Acinetobacter* and *Pseudomonas* (Sun et al. 2025). The genus *Pseudomonas* was detected in samples at all three sample sites, with the highest abundance in Week 4 aged samples and lowest abundance in fresh samples. Additionally, this genus never increased above 0.01% relative abundance at the ASHFInt site, where Pd was detected in guano samples through Week 12 (Table S24). *Acinetobacter* was highly abundant in ASHFExt Week 4, 8, and 12 aged samples and present in WMCC Week 4, 8, and 12 aged samples, but absent from ASHFInt samples (Table S24). *Rhodococcus*, another bacterial genus which inhibits Pd growth *in vitro* (Jacewicz et al. 2026), was present in a similar pattern, with relatively higher abundances at ASHFExt and WMCC and low abundances at ASHFInt (Table S24).

### Classification of viral taxa from assembly and read level metagenomics

Aged samples had considerably higher DNA concentrations than fresh samples. Out of 86 samples, 48 samples were sequenced (those with acceptable DNA concentrations), and 40 of these were aged. The final dataset consisted of a total of 33 aged samples (n = 10 WMCC, n = 11 ASHFExt, and n = 11 ASHFInt samples). The average number of reads generated from aged samples (final and screen runs combined) was 3.6M (Table S20). The lowest-throughput sample was a Week 8 aged ASHFInt sample, which only had 733,896 reads, while the highest throughput sample was ASHFExt week 8 aged from August (9.1M reads). The average N50 was 1.8 kb, while the lowest was 434 bp. Sample replicates at this stage were combined, and host contamination varied greatly across combined samples, with the average alignment rate to the host being 20.25%, where the upper and lower limits were 0.46% and 61.86% respectively. Further, total reads removed as a result of host alignment ranged between 10k and 4.0M; on average 599k reads were removed (Table S20). A total of 12 metagenome assemblies were generated; for statistics, see Table S21.

Read level analysis of the shotgun metagenomic data resulted in the identification of 100 viral orders sorted by site and age. Of the detected species, 82 were classified as bacteriophages, while 18 non-bacteriophage viral orders were also identified. Among these non-bacteriophage viruses, six mastadenovirus strains were associated with bats, humans, canines, and simians. Those directly correlated to humans included Human mastadenovirus B, Human mastadenovirus C, and Human mastadenovirus E, with 8, 20, and 14 full-length read hits identified, respectively (Table S22). In addition, Bat mastadenovirus C was detected with a total of eight full-length read hits (Table S22). There were also three double-stranded DNA viruses that targeted protist or amoeba-like hosts, *Cotonvirus japonicum, Moumouvirus australiense*, and *Biavirus raunefjordenese*, each with two hits.

The metagenomic assembly level classification provided more resolution to detect non-bacteriophage viruses. Bacteriophages were still dominant and relatively stable across all sites and aging timepoints, particularly Tubulavirales (average 45% relative abundance), but several viruses with non-bacterial hosts were detected. Exterior sites show greater viral diversity overall, while ASHFInt predominately consists of a few key orders: Tubulavirales, Piccovirales, and Rowavirales in Week 4 samples and Algavirales in Week 8 samples. A comparison of the exterior sites shows that ASHFExt has an increased presence of viral orders compared to WMCC. Further, decomposition appears to have an effect on viral composition. At ASHFInt, aging at Week 8 results in the complete loss of all viral orders observed at Week 4 and is completely dominated by Algavirales (100%). Aging at ASHFExt shows the greatest turnover in viral composition: Priklausovirales (20%), Magrovirales (4.7%), and Pimascovirales (4.7%) are present only in Week 4. Aging from Week 8 to Week 12 introduces the presence of Petitvirales (13%), Orthopolintovira (4.7%), and Harevirales (9.2%), all of which are uniquely present at ASHFExt. No viral orders are unique to WMCC; however, a similar pattern of change in viral composition is observed with decomposition, most notably the complete loss of Lautamovirales (7.6%), and the new presence of Pimascovirales (6.6%) and Piccovirales (9.5%) from Week 4 to Week 8 (Fig. 8; Table S23). The viral composition remains stable from Week 8 to Week 12 at WMCC. No order level classifications were made for ASHFInt Week 12.

**Figure 8:**
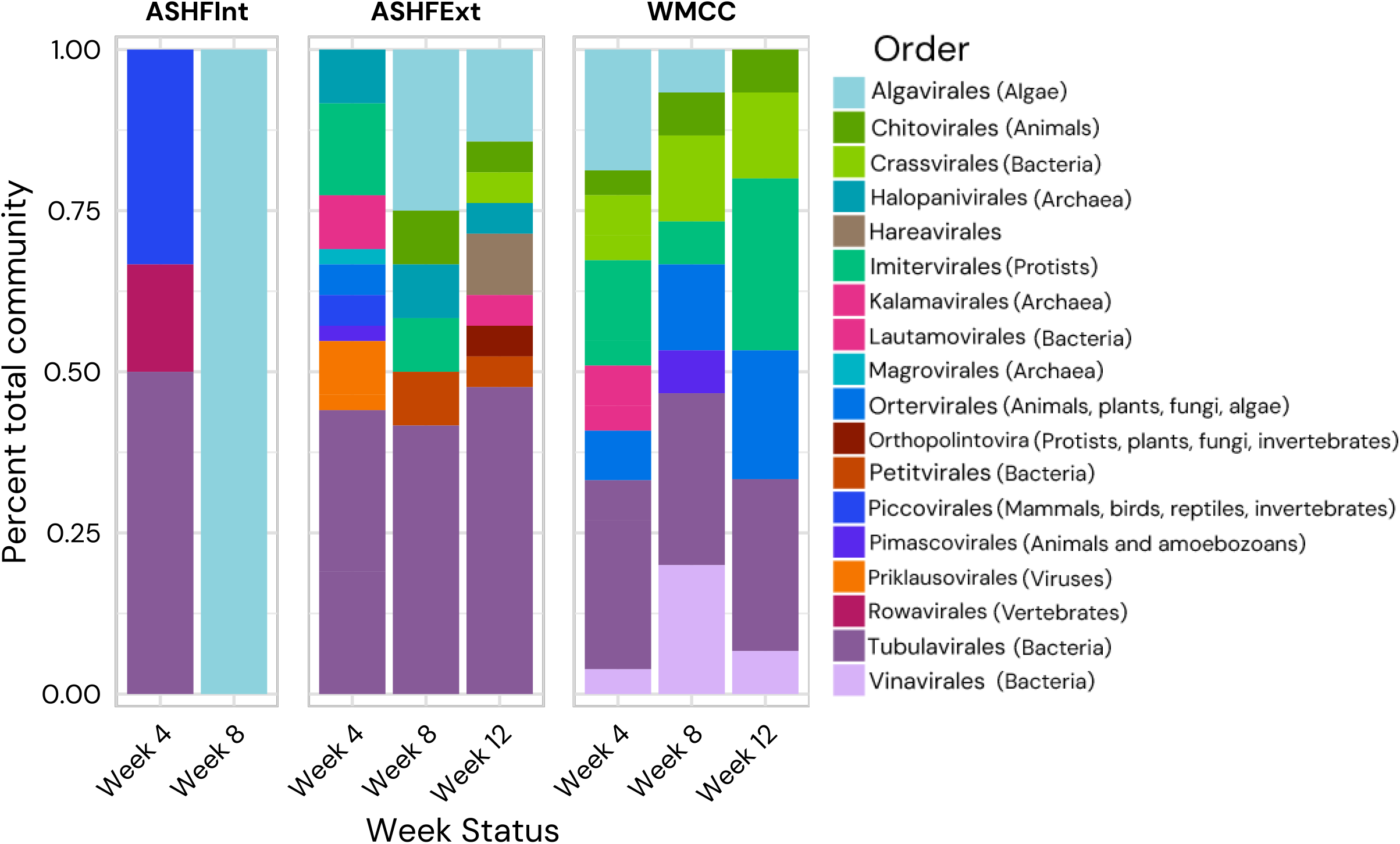
Relative abundance of viral orders detected in aged metagenomic assemblies. For each site across each timepoint, the relative abundance of viral orders is shown. Viral hosts are listed in parentheses next to each viral order in the legend.

## DISCUSSION

### Fresh guano retains a gut-associated signal whereas decomposition drives rapid environmental restructuring of microbial communities

Fresh guano retained a stronger gut-associated 16S signal than aged material and was more consistent across sites. Fresh samples were dominated by predominately anaerobic and facultative anaerobic taxa including Clostridiaceae, Enterobacteriaceae, and Enterococcaceae (Arboleya et al. 2012) (Fig. 2D). With decomposition, exterior samples shifted toward communities structured by environmental conditions, gaining taxa such as Pseudomonadaceae, Oxalobacteraceae, Sphingobacteriaceae, and Nocardiaceae (Fig. 2D). Microbial community source analysis, conducted in SourceTracker, confirmed that bacteria found in fresh samples contributed less than 5% of aged OTUs in nearly all exterior replicates (Fig. 2F). These shifts are consistent with prior work demonstrating that microbial communities in bat guano undergo rapid composition changes when exposed to aerobic environments (Fofanov et al. 2018). In contrast, guano communities at ASHFInt were more stable: fresh sample bacteria contributed 27–92% of aged OTUs, far higher than at any exterior site (Fig. 2F). The variation between interior and exterior guano communities suggests that the rapid change in exterior communities may be driven by exposure to soil microbiomes, whereas obligate anaerobes in interior guano likely faced less competition from environmental microbes.

### The guano microbiome changes temporally across months and during aging

The bacterial communities of bat guano are highly dynamic and shift during decomposition, a pattern documented across short and long timescales. Previous studies have demonstrated that across short timescales, bacterial diversity in bat guano declines upon aerobic exposure, with moderate disruption of diversity expected within as little as one hour of exposure (Fofanov et al. 2018). Across longer timescales diversity decreases with age; in guano accumulation over 120 years in a cave ecosystem, community composition transitioned from high diversity aerobic surface communities to low diversity communities dominated by *Actinomycetota* (McFarlane & Lundberg 2024). Most bat guano aging studies are conducted in cave environments where guano accumulates undisturbed, and comparatively little is known about how bacterial communities shift across seasonal timescales in non-cave environments. We found notable changes in diversity across the aging sequence, indicating that age of guano is an important variable when considering community composition.

Fresh guano alpha diversity was relatively stable across months, with no significant variation in alpha diversity for bacterial communities at either exterior site (Fig. 2A, B). Fresh bacterial composition was not significantly different across sites; however, it significantly changed through the months (Table S7). Fresh fungal composition was significantly different across sites and months (Table S12). Month-to-month variations in the microbiome of fresh guano may reflect seasonal fluctuations in prey consumed, with taxa turning over but maintaining similar proportions and therefore stable alpha diversity. Because of this difference in microbiome composition across months, aging series from June and August had distinct starting points, but all three sites started with similar guano bacteriomes. Despite these differences by month and similarities by site in fresh guano, aged communities diverged by site and converged between the two aging series, indicating that environmental exposure, even between ecologically similar sites, is a strong driver of community restructuring (Table S7; Table S12). Both the taxa present and the relative abundances of those taxa are changing across the aging series, a true example of community succession (Fig. 2C-E; Fig. 4C-E).

### Fungal diversity is highest in fresh guano and declines with environmental exposure

Fungal community composition differed across site and age, with fresh guano at external sites consistently exhibiting higher Shannon diversity than aged samples, most pronounced at WMCC where inverse Simpson diversity also declined significantly by Week 4. ASHFExt showed a significant decline in Shannon diversity by Week 8 (Fig. 4A), and both sites stabilizing at consistently low diversity in the following aging timepoints. This initial decline is consistent with environmental filtering: exposure to oxygen, desiccation, and different external microbial inputs selected for taxa capable of persisting under these conditions. Cave-associated bat guano shows similar patterns, with fungal assemblages reflecting substrate availability, environmental conditions, and external microbial communities rather than a stable host-associated community (Dimkić et al. 2021). In the interior environment, fungal diversity peaked in Week 4, then declined at Week 8 (Fig. 4A, B). The interior environment may have isolated the guano from immediate filtration by environmental factors, thus altering succession patterns.

### Exterior sites converge while the interior site diverges in bacterial and fungal succession

Small-scale spatial variation was the dominant contributor to bacterial and fungal composition in aged samples. In all tested beta diversity metrics, aged exterior sites cluster while the ASHFInt site aligns with fresh samples (Fig. 2C; Fig. 4C; Fig. S1C, D; Fig. S2C, D). Unsurprisingly, the most abundant bacterial and fungal taxa at all sites are considered cosmopolitan taxa that are common across many environments (Salgado-Salazar et al. 2013; Ozimek & Hanaka 2020; Telagathoti et al. 2021). The differences between the three sites suggest that shelter design and surrounding ecology jointly govern microbial richness and community composition.

Despite their small-scale differences in local habitat as indicated by differences in bat diet, the two exterior sites (ASHFExt and WMCC) exhibited similar shifts in both bacterial and fungal diversity over the aging period. One notable shift is the immediate loss of anaerobic bacterial families, such as Clostridiaceae (Fig. 2D). ASHFInt demonstrated the opposite pattern compared to exterior sites. Bacterial communities over time maintained anaerobic bacteria such as Clostridiaceae, and other taxa associated with exterior aged samples, such as *Actinobacteria*, remained low (Fig. 2D; Fig. 3A). Interior conditions, especially manmade roosts like bat houses and barns, may cultivate a unique guano microbiome by reducing weathering and colonization by competitors; both fungal and bacterial samples contain many ASHFInt site indicators but relatively few indicators of the exterior sites (Fig. 3A; Fig. 4F). In bacterial samples, Corynebacteriaceae, unclassified Corynebacteriales, and Bogoriellaceae were the only bacterial indicators of the combined ASHF sites that persisted across time, indicating that few bacterial taxa retain their site association through the decomposition process at ASHFExt (Fig. 3A). In fungal samples, the early peak in Mucoromyceta in both exterior sites (Fig. 4D) suggests an early successional role for these fungi in guano, while Basidiomycota were in higher abundances in Week 8 & 12. These patterns resemble the succession of fungal fruiting on dung in which Mucoromyceta fruited in the first week, with Basidiomycota such as *Coprinus* observed beginning in 2-3 weeks (Richardson, 2002).

Long-read CO1 metabarcoding reveals diverse arthropod prey with site and temporal signatures Bats are generalist foragers known to consume a wide array of arthropod taxa (Maslo et al. 2022). Insectivorous bat diets were historically characterized by visual assessment of insect taxa recovered from guano (O’Rourke et al. 2022; Webster & Whitaker 2005), but metabarcoding enables wider-scale assessment of diet and foraging habits with greater taxonomic resolution. Long-read Nanopore CO1 sequencing captures the full-length amplicon, enabling species-level identification and more accurate family assignments than short-read approaches. This approach has been used to characterize the diet of the brown long-eared bat (*Plecotus auritus*), and in soil mites recovered the known community more accurately than short-read Illumina (Varusk et al. 2025). While long reads improve resolution, database completeness and specificity remains the primary limitation of accurate CO1 metabarcoding. This was partly mitigated by subsetting the highly curated BOLD database to insects present in the study’s geographical region.

Insect consumption in *M. lucifugus* varies both temporally and spatially throughout the species’ range (O’Rourke et al. 2022; Clare et al. 2011), and similar spatial and temporal variability was observed here. Several orders observed were consistent with previous *M. lucifugus* guano CO1 studies (Diptera, Coleoptera, Lepidoptera, and Ephemeroptera; O’Rourke et al. 2022; Kunz & Whitaker Jr. 1983), though abundances differed. Diptera was the most abundant order across all months, and Coleoptera was abundant at ASHF, as expected. However, Ephemeroptera and Lepidoptera were rarely observed, contrasting with previous diet characterizations (Fig. 5A, B). Lepidoptera reached its greatest relative abundance in early spring (May–June), coinciding with emergence. Lepidopteran presence in *M. lucifugus* guano has been attributed to changes in relative abundance rather than preference (Wray et al. 2021), so the early-season peak and subsequent sharp decline after June at all sites likely reflect availability.

### Aquatic insect access drives site-level dietary differentiation

Several taxa were more common at WMCC than the two ASHF sites, potentially reflecting local habitat structure. Aquatic insects are more abundant at WMCC, which, despite similar ecology, has a larger body of water near the sample site than the smaller streams and ponds at ASHF (Fig. 1B, C). Two aquatic taxa were indicators of WMCC: phantom midges (Chaoboridae) and frit flies (Chloropidae) (Fig. 5C, D). Chloropidae are not strictly aquatic but often inhabit vegetative aquatic habitats (Merritt et al. 2009), matching the surrounding ecology of WMCC. Chaoboridae was not an exclusive indicator of WMCC, but its increased abundance and indicator status together support site-level differentiation driven by aquatic and semi-aquatic insect families. Increased phantom midge abundance at WMCC likely reflects greater availability of suitable larval and adult habitats, as these taxa require aquatic habitats to complete their life cycle. Chaoboridae was also an indicator of ASHFInt, differentiating the interior and exterior ASHF sites.

The differentiation of WMCC from combined ASHF sites reflects significant differences in both relative abundance and presence-absence of arthropod taxa (Fig. S5A). The unique presence of Chloroperlidae (stoneflies) and Aturidae (water mites), both obligate aquatic arthropods, further supports aquatic insect presence as the driver of WMCC’s divergence (Fig. S4C).

### Shifts in diet composition are driven by temporal signals and dietary preference

*M. lucifugus* diets shift temporally, and temporal factors are known to influence diet diversity more than spatial factors (Wray et al. 2021). Variation in arthropod diversity across the summer feeding season has been documented in Canadian *M. lucifugus* maternity colonies, where Diptera dominated early-season diets and shifted to Lepidoptera in the mid and late season, likely driven by temporal insect abundance and diversity (Clare et al. 2013). Here, the broad temporal shift was weaker, as Diptera remained dominant throughout the summer (Fig. 5), though Dipteran abundance decreased at the end of the season in August. This is consistent with reports that Diptera comprise up to 63% of the *M. lucifugus* diet, especially early in the maternity season, with abundance declining toward the end (Clare et al. 2011).

August showed the greatest shift in diet composition, characterized by increased Coleoptera, Araneae, and Trombidiformes (Fig. 5). Two dipteran families (Dictynidae and Drosophilidae) were indicators favoring August, but the strongest indicator was Lebertiidae (water mites, Trombidiformes). Lebertiidae are diverse and commonly inhabit temperate water bodies of constant and variable flow in deciduous-dominated regions (Gülle & Boyaci 2012); in August, they were core to both ASHF sites but accessory at WMCC, niches consistent with the land cover at each location (Fig. 1C). Larval lebertiids parasitize adult Diptera, particularly Chironomidae, attaching to the thorax (Martin & Gerecke 2009), while free living stages prey on a broader range of aquatic invertebrates, providing opportunities for consumption by bats. Water mites were observed attached to dipteran hosts captured during insect monitoring at WMCC (File S2). A late season increase in Curculionidae (weevils) in August provides further evidence of temporal shift (Fig. 5). Curculionidae have been reported as prey of several bat species, including the Indiana bat (*Myotis sodalis*) and the long-tailed bat (*Chalinolobus tuberculatus*) (Ling et al. 2023; Tuttle et al. 2006). In addition, Tuttle et al. (2006) found weevils emerged as a primary dietary component in August for *M. sodalis* in Indiana. This shift may indicate bat preference or changing prey availability at the end of the season.

*M. lucifugus* is known to show prey preferences and selective foraging (Wray et al. 2021). The early summer months (May and June) were dominated by Diptera and Trombidiformes, with a smaller contribution from Lepidoptera (Fig. 5). Within Diptera, non-biting midges (Chironomidae) and gnat-like flies (Anisopodidae) were associated with May (Fig. S5E), aligning with spring emergence patterns (Blackwood et al. 1995; Bouchard Jr. & Ferrington Jr. 2023) as foraging activity increases after hibernation. Because Chironomidae are a known preferred prey of *M. lucifugus*, their early-summer association could reflect prey preference, selective foraging, or simply emergence patterns. This signal was conserved across sites, indicating that seasonality contributes more to early-season dietary diversity than site-level differences in prey availability.

Axonopsidae and cockroaches (Ectobiidae) were associated with June (Fig. S5F). The Axonopsidae signal could reflect increased June abundance, as optimal temperatures and resource availability are known to affect abundance (Vasquez et al. 2020), who found that observations increased during transitional months such as June, October, and November with changes in temperature and precipitation. Given the overall low relative abundance of Axonopsidae, however, bats are more likely feeding preferentially on aquatic insects that host Axonopsidae.

### Mosquitoes are a minor component of the M. lucifugus diet

Bats are often cited as efficient pest suppressors in agricultural and human health contexts (Puig-Montserrat et al. 2020; Wetzler & Boyles 2017); however, the literature disagrees on the proportion of bat diets comprising mosquito prey (Gonsalves et al. 2013), and no bat species specializes on mosquitoes (Wetzler & Boyles 2017). Although *M. lucifugus* consumed some mosquito (Culicidae) prey at the Ashford interior and exterior sites, mosquitoes represented a small fraction of the diet (Fig. 5B). Mosquitoes are likely less worthwhile relative to the energy required to capture them: although Rydell et al. (2006) found a 92% success rate for *M. lucifugus* hunting mosquitoes in Alaska, at least 87 hours of foraging would be needed to meet a bat’s daily energetic needs (Wetzler & Boyles 2017), and a pregnant bat would need to consume over 90% of its body mass to subsist on mosquitoes alone, potentially impairing flight (Wetzler & Boyles 2017; Kurta et al. 1989). An earlier study of Indiana bats found mosquitoes were consumed most frequently during pregnancy, but Diptera generally comprise less than 25% of that species’ diet overall (Kurta & Whitaker Jr. 1998). Across the sampled months, Diptera were consumed most at WMCC, where Culicidae did not rank among the top 60 families, indicating that *M. lucifugus* prefer other prey even when mosquitoes are abundant (Fig. 5B).

### Consumed insects are a possible source of arthropod-associated fungi in guano

Insect diet is a potential source of the fungal families detected in the samples. Bats and their hibernacula harbor substantial mycological diversity, and these fungi can parasitize bats and their insect prey or serve as food for arthropods (Becker et al. 2023; de Groot et al. 2020; Vanderwolf et al. 2016). The fungal families detected here are therefore likely shaped by fungus-arthropod interactions in addition to fungus-bat interactions. For example, Mortierellaceae was an indicator of the ASHF sites (Fig. 4F), and *Mortierella* spp. have been shown to live on both spiders and bats in the same caves (Vanderwolf et al. 2016). Members of this family may have been associated with the spiders detected at ASHFInt through the summer (Fig. 5). Gall midges were prevalent across all three sites (Fig. 6) and are known to use fungi as a food source (Pyszko et al. 2025). They may have contributed fungi to the bat gut or been drawn to guano to feed.

Interactions between fungi and bat arthropod prey warrant further exploration. Many fungi could not be classified to species level, with classifications restricted to family and genus. Determining the source of microbiome components is difficult, but accounting for secondary consumption and the conditions enabling fungal growth and spread will help clarify the factors shaping the bat gut microbiome (Bowser et al. 2013).

### Guano enables non-invasive surveillance of invasive insects

Since 2002, the invasive beetle emerald ash borer (*Agrilus planipennis*, EAB) has devastated North American ash tree populations (Herms & McCullough 2014). Traditional surveillance has relied on visual monitoring and bark sifting (Jennings et al. 2018), while recent advances enable environmental DNA (eDNA) collection (Kyle et al. 2024). The detection of EAB in guano from every month except September, and in guano aged both four and eight weeks, validates bat guano as a potential resource for *A. planipennis* monitoring. Differential DNA degradation can bias biodiversity estimates from eDNA samples (Krehenwinkel et al. 2018), but the recovery of EAB borer from aged guano (Fig. 7B) indicates that this insect’s DNA remains viable in guano for up to eight weeks. Prior work has used bat guano to detect invasive insects, especially the spotted lanternfly (McHale et al. 2025), but that guano was extracted immediately after collection so the effect of degradation is unknown. No prior published study has used bat guano to detect EAB by molecular methods, and these results are promising for future conservation applications. The genus *Agrilus* was absent from the CO1 results (Table S17), though it is known to be difficult to identify by barcoding (Pentinsaari et al. 2014). Future guano surveillance for EAB should therefore prioritize qPCR over barcoding.

Evening bats (*Nycticeius humeralis*) are known to consume EAB (Münzer et al. 2016), though visual methods found them to be a very small dietary component. The detection of EAB in up to three of the 7 to 8 fresh samples per month suggests it may constitute a larger part of the diet than previously assumed. Emerald ash borer is primarily diurnal (Jennings et al. 2013), resting on trees at night when bats are most active. *M. lucifugus* exhibits both gleaning and hawking behaviors (Ratcliffe & Dawson 2003) and may show behavioral plasticity in response to abundant diurnal insects. The greater mouse-eared bat (*Myotis myotis*) adopts hawking strategies to capture comparatively risky large ground insects, favoring higher caloric reward over the lower reward, but more reliable, catch of smaller flying insects (Stidsholt et al. 2023).

### Pseudogymnoascus destructans persists in guano through the summer maternity season

White-nose syndrome, caused by Pd spores, has caused widespread mortality across many North American bat species, and guano sampling is an effective method of fungal detection for disease surveillance (Grider et al. 2021). Pd was detected in guano from every month sampled and in both fresh and aged guano (Fig. 7A), and spores were present in bat house and barn samples in every month at each site (Fig. S6B). Monitoring of Pd has focused primarily on hibernacula where white-nose syndrome infections are active, but these results indicate that the fungus persists in guano for at least 12 weeks following the return to maternity colonies, despite degrading quickly at high temperatures (Verant et al. 2012). Experimental inoculation found Pd detectable until 42 days after inoculation (Urbina et al. 2020), although that guano was held at temperatures lower than the average summer temperatures at both sample locations.

*Pseudogymnoascus destructans* was detected in Week 12 aged samples only at ASHFInt. Bat houses inside larger human structures provide shelter from UV exposure and direct precipitation, yet anecdotal observation indicated that interior temperatures at ASHFInt were consistently higher than outdoor ambient conditions, consistent with bat houses intended to retain heat (Fontaine et al. 2021). The Pd persistence pattern likely reflects the combined effects of physical shelter, including more stability of the interior microclimate, and the reduction of antifungal bacterial taxa (Fig. 7A). Spore counts from bat houses at ASHFInt were consistently moderate, while WMCC and ASHFExt showed larger accumulation depending on the month (Fig. S6B).

These results align with skin swab data from maternity colonies (Carpenter et al. 2016; Huebschman et al. 2019), though fungal transmission and risk within maternity colonies remain poorly understood. Residual detection may reflect conidia on skin, which can survive up to 60 days on bat fur at 30 °C and up to 180 days at 24 °C (Isidoro-Ayza et al. 2024; Campbell et al. 2019). The combination of residual conidia and grooming behavior may sustain the ongoing presence of Pd in guano observed here. Such residual material is unlikely to drive transmission at maternity colonies (Fischer et al. 2021), since, at least in European colonies, Pd strains do not show the loss of genetic differentiation expected under maternity colony transmission. Because fungal load from skin swabs also drops sharply over the summer, infection at maternity colonies is considered unlikely (Langwig et al. 2015). Dispersal to hibernacula, however, may pose a threat (Ballmann et al. 2017). Pd can survive and grow for up to four months on environmental substrates such as wood (Urbina et al. 2021), the material of both the bat boxes at WMCC and the boxes and barn at ASHF. Viable Pd has been cultured from wood surfaces of bat houses (Dobony & Johnson 2018), although culturing was performed at optimal temperatures for Pd growth, and whether this holds at higher temperatures is unclear. Bats returning to hibernacula experience lower temperatures ideal for psychrophilic fungi, so the presence of Pd in Week 12 aged guano is concerning.

Whether the presence of Pd in maternity colonies affects susceptibility to infection during hibernation is unclear. Bats do not appear to mount a sufficient antibody-mediated response to directly combat Pd infection (Lilley et al. 2017; Johnson et al. 2015) but do undergo a local inflammatory response (Lilley et al. 2017) that may be detrimental to hibernation survival. A cell-mediated response induced by a virally vectored vaccine increased survival in Pd-infected *M. lucifugus* (Rocke et al. 2019), demonstrating that some adaptive immunity is possible. It remains unclear whether summer exposure confers any immunological advantage to adult bats, or whether pups born in colonies with residual spores gain any advantage upon later exposure at hibernacula. Further study is needed to determine the effects of residual conidia on titer levels and immune responses following exposure in maternity colonies.

*Pseudogymnoascus destructans* has been shown to grow on autoclaved guano but not on fresh, non-autoclaved guano (Urbina et al. 2021), possibly due to bacterial communities involved in organic matter decomposition, including the highly abundant Enterobacteriaceae present at every site (Fig. 2C). A lack of antifungal bacteria, including the genera *Acinetobacter* and *Pseudomonas* (Sun et al. 2025), at ASHFInt may also have led to the detection of Pd through Week 12 (Table S24). Future work should investigate Pd growth on additional substrates during summer.

### Short-term aging is the primary driver of changes in viral diversity

Long-read shotgun metagenomics resolved 18 viral orders from aged guano. This resolution reflects the advantages of long reads, which span entire viral genomes or genes to improve downstream classification (Zaragoza-Solas et al. 2022) and produce more contiguous, complete genome assemblies (Warwick-Dugdale et al. 2019), and recent Nanopore virome work targeting bacteriophages reports the same gain in contiguity (Wirbel et al. 2025). Consistent with these advantages, most samples here recovered a substantial proportion of contigs exceeding 100 Kb, indicating that complete viral genomes were likely assembled (Table S21).

Bats are reservoirs of potentially zoonotic viruses and harbor the greatest viral diversity of any mammal (Y. Li et al. 2021), a pattern linked to communal roosting and migration (Hu et al. 2017; Drexler et al. 2014). Viral taxa commonly associated with bats include adeno-associated viruses, herpesviruses, astroviruses, and lyssaviruses (Dimkić et al. 2021). Diverse viruses have also been identified in *M. lucifugus* (Donaldson et al. 2010), including taxa that infect arthropod hosts as well as a large diversity of bacteriophages. However, few studies have characterized viruses from *M. lucifugus* guano, and fewer still from guano of North American populations in anthropic roosts. Work in this species has largely focused on cave environments, recovering SARS-related coronaviruses, arthropod and plant viruses, and bacteriophages (Donaldson et al. 2010). The virome of anthropic guano carries public health relevance given its proximity to human structures, yet few studies have evaluated viral diversity in truly anthropic guano. Soto-Lopez et al. (2025) examined cave guano near human hiking trails and found high abundances of Herpesvirales and other pathogenic viruses alongside the expected insect-infecting viruses (Tymoviridae and Piccovirales). Fewer studies still have assessed how guano age affects viral diversity. Viral composition, like microbial composition, may shift as nutrient and host availability change with decomposition. To our knowledge, this is the first study to assess the effects of in situ, short-term aging on the guano virome.

Guano typically hosts more environmental viruses and bacteriophages than host-specific pathogenic viruses, consistent with the present results (Roev et al. 2023). Aging had its greatest effect on non-bacteriophage composition: dominant bacteriophages such as Tubulavirales remained stable across sites and aging (Fig. 8). Site level viral diversity was broadly similar, with exceptions including Rowavirales, present only in ASHFInt Week 4 aged guano (Fig. 8). Previously detected in guano of Russian bat species (Roev et al. 2023), this order is notable because it infects mammalian hosts and includes the human adenoviruses also recovered through read-based identification (Table S22). Its restriction to early interior guano suggests it does not persist through decomposition. Because adenoviruses inactivate faster under warm conditions and UV exposure (Ghodshi et al. 2026), the reduced sunlight at ASHFInt may explain the elevated signal. Pathogenicity and transmission cannot be assessed here, though adenoviruses are host-specific and bat to human transmission is rare (Lee & Angiel et al. 2020).

A similar pattern was observed for Piccovirales, which infects a broad range of hosts, comprising most of the Week 4 interior virome but only a small portion at Week 4 ASHFExt (Fig. 8). Piccovirales contains a single family, Parvoviridae, whose members infect both vertebrates and invertebrates (Penzes et al. 2020). Parvoviruses in Californian roost guano have been attributed to rodent or canine urinary and fecal contamination (Y. Li et al. 2021), consistent with greater rodent presence in a barn environment such as ASHFInt. Parvoviridae also includes the insect-infecting Densovirinae, reported as a major component of the guano virome and likely derived from ingested arthropods (Li et al. 2010); Densovirinae was not detected here, nor were other insect-associated orders, possibly reflecting differences in sequencing coverage. Shifts in presence and abundance also occurred for Chitovirales (infecting animals), restricted to Week 8 and Week 12, and for the Imitervirales (protist infecting) (Fig. 8).

## Acknowledgments

The co-first authors are RaMP (Research and Mentoring for Postbaccalaureates) fellows at the University of Connecticut, supported with an award from the National Science Foundation (DBI-2217100 to ELJ, JLW, and RJO), which also supported the research. DJA received support through the National Science Foundation (DBI-2222328) and (DBI-1737778).

The authors acknowledge contributions of the HPC resources from the Computational Biology Core and sequencing resources from the Center for Genome Innovation, both within the Institute for Systems Genomics at the University of Connecticut. The authors also acknowledge the Ashford site property owner, Ruth Cutler.

## Data Archiving Statement

All code used in data analyses presented here is hosted at: https://gitlab.com/PlantGenomicsLab/myotis_lucifigus_metagenomics_2025_2026. Illumina and Nanopore sequence data for 16S, ITS, CO1, and metagenomic analysis are hosted at NCBI Bioproject PRJNA1482243.

## Conflict of Interest

The authors declare no conflicts of interest.

## Ethics Statement

This study used only non-invasively collected bat guano deposited beneath occupied roosts; no bats were captured, handled, or otherwise disturbed, and no animal procedures were performed. Because the work involved no live-animal handling or experimentation, it did not require Institutional Animal Care and Use Committee (IACUC) approval. Guano and spore samples were collected with the permission of the landowners and site managers at the Ashford, Connecticut residence and the White Memorial Conservation Center, and in coordination with the Connecticut Department of Energy and Environmental Protection. All sampling complied with relevant local and state wildlife and animal welfare regulations.

## Authorship Information

**Conceptualization:** K.P., N.P., R.J.O., E.L.J., J.L.W., J.F., A.M., D.F., H.R. **Methodology:** K.M., D.J.A., M.R., A.R., N.P., E.L.J., J.L.W., J.F. **Investigation:** J.D., L.E., A.G., D.J.H., D.L., R.L., R.M., K.S., K.A.S., J.T., S.C.V., K.P., K.M., D.J.A., M.R., A.R., A.M. **Formal analysis:** J.D., L.E., A.G., D.J.H., D.L., R.L., R.M., K.S., K.A.S., J.T., S.C.V., K.P., J.L.W., A.M. **Resources:** K.M., D.J.A., A.R., R.J.O., T.K., J.L.W., J.F. **Supervision:** K.M., K.C., R.J.O., E.L.J., J.L.W., A.M. **Project administration:** R.J.O., E.L.J., J.L.W., A.M., D.F., H.R. **Funding acquisition:** R.J.O., E.L.J., J.L.W. **Writing – original draft:** J.D., L.E., A.G., D.J.H., D.L., R.L., R.M., K.S., K.A.S., J.T., S.C.V., T.K., A.M. **Writing – review & editing:** all authors.

## MESH terms

- Chiroptera
- Microbiota
- Feces
- DNA Barcoding, Taxonomic
- High-Throughput Nucleotide Sequencing
- Fungi

## Notes

https://gitlab.com/PlantGenomicsLab/myotis_lucifigus_metagenomics_2025_2026

### Competing Interest Statement

The authors have declared no competing interest.

## References

Anderson, Marti J. (2001). A New Method for Non-Parametric Multivariate Analysis of Variance. Austral Ecology 26, no. 1 (2001): 32–46. 10.1111/j.1442-9993.2001.01070.pp.x.

Anderson, M. J., Ellingsen, K. E., & McArdle, B. H. (2006). Multivariate dispersion as a measure of beta diversity. Ecology Letters, 9(6), 683–693. 10.1111/j.1461-0248.2006.00926.x

Arboleya, S., Solís, G., Fernández, N., de los Reyes-Gavilán, C. G., & Gueimonde, M. (2012). Facultative to strict anaerobes ratio in the preterm infant microbiota: A target for intervention? Gut Microbes, 3(6), 583–588. 10.4161/gmic.21942

Ballmann, A. E., Torkelson, M. R., Bohuski, E. A., Russell, R. E., & Blehert, D. S. (2017). DISPERSAL HAZARDS OF PSEUDOGYMNOASCUS DESTRUCTANS BY BATS AND HUMAN ACTIVITY AT HIBERNACULA IN SUMMER. Journal of Wildlife Diseases, 53(4), 725. 10.7589/2016-09-206

Becker, P., van den Eynde, C., Baert, F., D’hooge, E., De Pauw, R., Normand, A. C., … Stubbe, D. (2023). Remarkable fungal biodiversity on northern Belgium bats and hibernacula. Mycologia, 115(4), 484–498. 10.1080/00275514.2023.2213138

Benjamini, Y., & Hochberg, Y. (1995). Controlling the False Discovery Rate: A Practical and Powerful Approach to Multiple Testing. Journal of the Royal Statistical Society. Series B (Methodological), 57(1), 289–300. https://www.jstor.org/stable/2346101

Blackwood, M. A., Hall, S. M., & Ferrington, L. C. (1995). Emergence of Chironomidae from Springs in the Central High Plains Region of the United States. JOURNAL OF THE KANSAS ENTOMOLOGICAL SOCIET, 68(2), 132–151.

Bourlat, S., Koch, M., Kirse, A., Langen, K., Espeland, M., Giebner, H., Decher, J., Ssymank, A., & Fonseca, V. (2023). Metabarcoding dietary analysis in the insectivorous bat Nyctalus leisleri and implications for conservation. Biodiversity Data Journal, 11, e111146. 10.3897/BDJ.11.e111146

Bowser, A. K., Diamond, A. W., & Addison, J. A. (2013). From Puffins to Plankton: A DNA-Based Analysis of a Seabird Food Chain in the Northern Gulf of Maine. PLOS ONE, 8(12), e83152. 10.1371/journal.pone.0083152

Breitwieser, F. P., & Salzberg, S. L. (2020). Pavian: Interactive analysis of metagenomics data for microbiome studies and pathogen identification. Bioinformatics, 36(4), 1303–1304. 10.1093/bioinformatics/btz715

Camargo, A. P., Roux, S., Schulz, F., Babinski, M., Xu, Y., Hu, B., Chain, P. S. G., Nayfach, S., & Kyrpides, N. C. (2024). Identification of mobile genetic elements with geNomad. Nature Biotechnology, 42(8), 1303–1312. 10.1038/s41587-023-01953-y

Campbell, L. J., Walsh, D. P., Blehert, D. S., & Lorch, J. M. (2019). LONG-TERM SURVIVAL OF PSEUDOGYMNOASCUS DESTRUCTANS AT ELEVATED TEMPERATURES. Journal of Wildlife Diseases, 56(2), 278. 10.7589/2019-04-106

Carpenter, G. M., Willcox, E. V., Bernard, R. F., & H. Stiver, W. (2016). Detection of Pseudogymnoascus destructans on Free-flying Male Bats Captured During Summer in the Southeastern USA. Journal of Wildlife Diseases, 52(4), 922–926. 10.7589/2016-02-041

Chaverri, P., & Chaverri, G. (2022). Fungal communities in feces of the frugivorous bat Ectophylla alba and its highly specialized Ficus colubrinae diet. Animal Microbiome, 4(1), 24. 10.1186/s42523-022-00169-w

Chuvochina, M., Gerken, J., Frentrup, M., Sandikci, Y., Goldmann, R., Freese, H. M., Göker, M., Sikorski, J., Yarza, P., Quast, C., Peplies, J., Glöckner, F. O., & Reimer, L. C. (2026). SILVA in 2026: A global core biodata resource for rRNA within the DSMZ digital diversity. Nucleic Acids Research, 54(D1), D334–D341. 10.1093/nar/gkaf1247

Clare, E. L., Barber, B. R., Sweeney, B. W., Hebert, P. D. N., & Fenton, M. B. (2011). Eating local: Influences of habitat on the diet of little brown bats (Myotis lucifugus). Molecular Ecology, 20(8), 1772–1780. 10.1111/j.1365-294X.2011.05040.x

Clare, E. L., Symondson, W. O. C., Broders, H., Fabianek, F., Fraser, E. E., MacKenzie, A., Boughen, A., Hamilton, R., Willis, C. K. R., Martinez-Nuñez, F., Menzies, A. K., Norquay, K. J. O., Brigham, M., Poissant, J., Rintoul, J., Barclay, R. M. R., & Reimer, J. P. (2013). The diet of Myotis lucifugus across Canada: Assessing foraging quality and diet variability. Molecular Ecology, 23(15), 3618–3632. 10.1111/mec.12542

Cunha, A. O. B., Bezerra, J. D. P., Oliveira, T. G. L., Barbier, E., Bernard, E., Machado, A. R., & Souza-Motta, C. M. (2020). Living in the dark: Bat caves as hotspots of fungal diversity. PLOS ONE, 15(12), e0243494. 10.1371/journal.pone.0243494

Danecek, P., Bonfield, J. K., Liddle, J., Marshall, J., Ohan, V., Pollard, M. O., Whitwham, A., Keane, T., McCarthy, S. A., Davies, R. M., & Li, H. (2021). Twelve years of SAMtools and BCFtools. GigaScience, 10(2), giab008. 10.1093/gigascience/giab008

De Cáceres, M., & Legendre, P. (2009). Associations between species and groups of sites: Indices and statistical inference. Ecology, 90(12), 3566–3574. 10.1890/08-1823.1

De Cáceres, M., Legendre, P., & Moretti, M. (2010). Improving indicator species analysis by combining groups of sites. Oikos, 119(10), 1674–1684. 10.1111/j.1600-0706.2010.18334.x

De Coster, W., & Rademakers, R. (2023). NanoPack2: Population-scale evaluation of long-read sequencing data. Bioinformatics, 39(5), btad311. 10.1093/bioinformatics/btad311

de Groot, M. D., Dumolein, I., Hiller, T., Sándor, A. D., Szentiványi, T., Schilthuizen, M., Aime, M. C., Verbeken, A., & Haelewaters, D. (2020). On the Fly: Tritrophic Associations of Bats, Bat Flies, and Fungi. Journal of Fungi, 6(4), 361. 10.3390/jof6040361

Dobony, C. A., & Johnson, J. B. (2018). Observed Resiliency of Little Brown Myotis to Long-Term White-Nose Syndrome Exposure. Journal of Fish and Wildlife Management, 9(1), 168–179. 10.3996/102017-JFWM-080

Donaldson, E. F., Haskew, A. N., Gates, J. E., Huynh, J., Moore, C. J., & Frieman, M. B. (2010). Metagenomic Analysis of the Viromes of Three North American Bat Species: Viral Diversity among Different Bat Species That Share a Common Habitat. Journal of Virology, 84(24), 13004–13018. 10.1128/JVI.01255-10

Drexler, J. F., Corman, V. M., & Drosten, C. (2014). Ecology, evolution and classification of bat coronaviruses in the aftermath of SARS. Antiviral Research, 101, 45–56. 10.1016/j.antiviral.2013.10.013

Dubois, B., Delitte, M., Lengrand, S., Bragard, C., Legrève, A., & Debode, F. (2024). PRONAME: A user-friendly pipeline to process long-read nanopore metabarcoding data by generating high-quality consensus sequences. Frontiers in Bioinformatics, 4, 1483255. 10.3389/fbinf.2024.1483255

Dufrêne, M., & Legendre, P. (1997). Species Assemblages and Indicator Species:the Need for a Flexible Asymmetrical Approach. Ecological Monographs, 67(3), 345–366. 10.1890/0012-9615(1997)067%255B0345:SAAIST%255D2.0.CO;2

Ekrem, T., Willassen, E., & Stur, E. (2007). A comprehensive DNA sequence library is essential for identification with DNA barcodes. Molecular Phylogenetics and Evolution, 43(2), 530–542. 10.1016/j.ympev.2006.11.021

Fischer, N. M., Altewischer, A., Ranpal, S., Dool, S., Kerth, G., & Puechmaille, S. J. (2022). Population genetics as a tool to elucidate pathogen reservoirs: Lessons from Pseudogymnoascus destructans, the causative agent of White-Nose disease in bats. Molecular Ecology, 31(2), 675–690. 10.1111/mec.16249

Fofanov, V. Y., Furstenau, T. N., Sanchez, D., Hepp, C. M., Cocking, J., Sobek, C., Pagel, N., Walker, F., & Chambers, C. L. (2018). Guano exposed: Impact of aerobic conditions on bat fecal microbiota. Ecology and Evolution, 8(11), 5563–5574. 10.1002/ece3.4084

Fontaine, A., Simard, A., Dubois, B., Dutel, J., & Elliott, K. H. (2021). Using mounting, orientation, and design to improve bat box thermodynamics in a northern temperate environment. Scientific Reports, 11(1), 7728. 10.1038/s41598-021-87327-3

Fu, L., Niu, B., Zhu, Z., Wu, S., & Li, W. (2012). CD-HIT: Accelerated for clustering the next-generation sequencing data. Bioinformatics, 28(23), 3150–3152. 10.1093/bioinformatics/bts565

Galan, M., Pons, J., Tournayre, O., Pierre, É., Leuchtmann, M., Pontier, D., & Charbonnel, N. (2018). Metabarcoding for the parallel identification of several hundred predators and their prey: Application to bat species diet analysis. Molecular Ecology Resources, 18(3), 474–489. 10.1111/1755-0998.12749

Gargas, A., Trest, M. T., Christensen, M., Volk, T. J., & Et, A. (2009). Geomyces destructans sp. Nov. Associated with bat white-nose syndrome. Mycotaxon, 108(1), 157–154. 10.5248/108.147

Gerbáčová, K., Maliničová, L., Kisková, J., Maslišová, V., Uhrin, M., & Pristaš, P. (2020). The Faecal Microbiome of Building-Dwelling Insectivorous Bats (Myotis myotis and Rhinolophus hipposideros) also Contains Antibiotic-Resistant Bacterial Representatives. Current Microbiology, 77(9), 2333–2344. 10.1007/s00284-020-02095-z

Ghodsi, S., Nikaeen, M., Gholipour, S., Rahmani, H. R., & Saderi, H. (2026). Viral persistence in soil: Insights from qPCR and cell culture detection of adenovirus. Journal of Environmental Chemical Engineering, 14(2), 121848. 10.1016/j.jece.2026.121848

Gonsalves, L., Bicknell, B., Law, B., Webb, C., & Monamy, V. (2013). Mosquito Consumption by Insectivorous Bats: Does Size Matter? PLoS ONE, 8(10), e77183. 10.1371/journal.pone.0077183

Grider, J. F., Russell, R. E., Ballmann, A. E., & Hefley, T. J. (2021). Long-term Pseudogymnoascus destructans surveillance data reveal factors contributing to pathogen presence. Ecosphere, 12(11), e03808. 10.1002/ecs2.3808

Gülle, P., & Boyaci, Y. (2012). Water mites of the genus Lebertia Neuman, 1880 (Acari, Hydrachnidia, Lebertiidae) from Turkey, with the description of one new species. ZooKeys, 238, 23–30. 10.3897/zookeys.238.3861

Heath, J. J., & Stireman III, J. O. (2010). Dissecting the association between a gall midge, Asteromyia carbonifera, and its symbiotic fungus, Botryosphaeria dothidea. Entomologia Experimentalis et Applicata, 137(1), 36–49. 10.1111/j.1570-7458.2010.01040.x

Herms, D. A., & McCullough, D. G. (2014). Emerald Ash Borer Invasion of North America: History, Biology, Ecology, Impacts, and Management. Annual Review of Entomology, 59, 13–30. 10.1146/annurev-ento-011613-162051

Hu, B., Zeng, L.-P., Yang, X.-L., Ge, X.-Y., Zhang, W., Li, B., Xie, J.-Z., Shen, X.-R., Zhang, Y.-Z., Wang, N., Luo, D.-S., Zheng, X.-S., Wang, M.-N., Daszak, P., Wang, L.-F., Cui, J., & Shi, Z.-L. (2017). Discovery of a rich gene pool of bat SARS-related coronaviruses provides new insights into the origin of SARS coronavirus. PLoS Pathogens, 13(11), e1006698. 10.1371/journal.ppat.1006698

Huebschman, J. J., Hoerner, S. A., White, J. P., Kaarakka, H. M., Parise, K. L., & Foster, J. T. (2019). Detection of Pseudogymnoascus destructans on Wisconsin Bats During Summer. Journal of Wildlife Diseases, 55(3), 673. 10.7589/2018-06-146

Ihrmark, K., Bödeker, I. T. M., Cruz-Martinez, K., Friberg, H., Kubartova, A., Schenck, J., Strid, Y., Stenlid, J., Brandström-Durling, M., Clemmensen, K. E., & Lindahl, B. D. (2012). New primers to amplify the fungal ITS2 region –evaluation by 454-sequencing of artificial and natural communities. FEMS Microbiology Ecology, 82(3), 666–677. 10.1111/j.1574-6941.2012.01437.x

Isidoro-Ayza, M., Lorch, J. M., & Klein, B. S. (2024). The skin I live in: Pathogenesis of white-nose syndrome of bats. PLOS Pathogens, 20(8), e1012342. 10.1371/journal.ppat.1012342

Jacewicz, M. J., Rogozynski, N. P., & Dixon, B. (2026). Strategies and limitations of the bat immune response to Pseudogymnoascus destructans: The causative agent of white-nose syndrome. Frontiers in Immunology, 16, 1736823. 10.3389/fimmu.2025.1736823

Jennings, D. E., Duan, J. J., & Shrewsbury, P. M. (2018). Comparing Methods for Monitoring Establishment of the Emerald Ash Borer (Agrilus planipennis, Coleoptera: Buprestidae) Egg Parasitoid Oobius agrili (Hymenoptera: Encyrtidae) in Maryland, USA. Forests, 9(10), 659. 10.3390/f9100659

Jennings, D. E., Taylor, P. B., & Duan, J. J. (2013). The mating and oviposition behavior of the invasive emerald ash borer (Agrilus planipennis), with reference to the influence of host tree condition. Journal of Pest Science, 87(1), 71–78. 10.1007/s10340-013-0539-1.

Johnson, J. S., Reeder, D. M., Lilley, T. M., Czirják, G.Á., Voigt, C. C., McMichael III, J. W., Meierhofer, M. B., Seery, C. W., Lumadue, S. S., Altmann, A. J., Toro, M. O., & Field, K. A. (2015). Antibodies to Pseudogymnoascus destructans are not sufficient for protection against white-nose syndrome. Ecology and Evolution, 5(11), 2203–2214. 10.1002/ece3.1502

Kim, D., Song, L., Breitwieser, F. P., & Salzberg, S. L. (2016). Centrifuge: Rapid and sensitive classification of metagenomic sequences. Genome Research, 26(12), 1721–1729. 10.1101/gr.210641.116

Knights, D., Kuczynski, J., Charlson, E. S., Zaneveld, J., Mozer, M. C., Collman, R. G., Bushman, F. D., Knight, R., & Kelley, S. T. (2011). Bayesian community-wide culture-independent microbial source tracking. Nature Methods, 8(9), 761–763. 10.1038/nmeth.1650

Kozich, J. J., Westcott, S. L., Baxter, N. T., Highlander, S. K., & Schloss, P. D. (2013). Development of a Dual-Index Sequencing Strategy and Curation Pipeline for Analyzing Amplicon Sequence Data on the MiSeq Illumina Sequencing Platform. Applied and Environmental Microbiology, 79(17), 5112–5120. 10.1128/AEM.01043-13

Krehenwinkel, H., Fong, M., Kennedy, S., Huang, E. G., Noriyuki, S., Cayetano, L., & Gillespie, R. (2018). The effect of DNA degradation bias in passive sampling devices on metabarcoding studies of arthropod communities and their associated microbiota. PLOS ONE, 13(1), e0189188. 10.1371/journal.pone.0189188

Kunz, T. H., & Whitaker Jr., J. O. (1983). An evaluation of fecal analysis for determining food habits of insectivorous bats. Canadian Journal of Zoology, 61(6), 1317–1321. 10.1139/z83-177

Kurta, A., Bell, G. P., Nagy, K. A., & Kunz, T. H. (1989). Energetics of Pregnancy and Lactation in Freeranging Little Brown Bats (Myotis lucifugus). Physiological Zoology, 62(3), 804–818. https://www.jstor.org/stable/30157928

Kurta, A., & Whitaker Jr, J. O. (1998). Diet of the Endangered Indiana Bat (Myotis sodalis) on the Northern Edge of Its Range. The American Midland Naturalist, 140(2), 280–286. 10.1674/0003-0031(1998)140%255B0280:DOTEIB%255D2.0.CO;2

Kyle, K. E., Allen, M. C., Siegert, N. W., Grabosky, J., & Lockwood, J. L. (2024). Design of an eDNA sampling method for detection of an endophagous forest pest. NeoBiota, 95, 149–164. 10.3897/neobiota.95.118267

Langwig, K. E., Frick, W. F., Reynolds, R., Parise, K. L., Drees, K. P., Hoyt, J. R., Cheng, T. L., Kunz, T. H., Foster, J. T., & Kilpatrick, A. M. (2015). Host and pathogen ecology drive the seasonal dynamics of a fungal disease, white-nose syndrome. Proceedings. Biological Sciences, 282(1799), 20142335. 10.1098/rspb.2014.2335

Lee, D. N., & Angiel, M. (2020). Two novel adenoviruses found in Cave Myotis bats (Myotis velifer) in Oklahoma. Virus Genes, 56(1), 99–103. 10.1007/s11262-019-01719-2

Li, H. (2021). New strategies to improve minimap2 alignment accuracy. Bioinformatics, 37(23), 4572–4574. 10.1093/bioinformatics/btab705

Li, L., Victoria, J. G., Wang, C., Jones, M., Fellers, G. M., Kunz, T. H., & Delwart, E. (2010). Bat Guano Virome: Predominance of Dietary Viruses from Insects and Plants plus Novel Mammalian Viruses. Journal of Virology, 84(14), 6955–6965. 10.1128/JVI.00501-10

Li, Y., Altan, E., Reyes, G., Halstead, B., Deng, X., & Delwart, E. (2021). Virome of Bat Guano from Nine Northern California Roosts. Journal of Virology, 95(3), e01713–20. 10.1128/JVI.01713-20

Li, Z., Li, A., Hoyt, J. R., Dai, W., Leng, H., Li, Y., Li, W., Liu, S., Jin, L., Sun, K., & Feng, J. (2021). Activity of bacteria isolated from bats against Pseudogymnoascus destructans in China. Microbial Biotechnology, 15(2), 469–481. 10.1111/1751-7915.13765

Lilley, T. M., Prokkola, J. M., Johnson, J. S., Rogers, E. J., Gronsky, S., Kurta, A., Reeder, D. M., & Field, K. A. (2017). Immune responses in hibernating little brown myotis (Myotis lucifugus) with white-nose syndrome. Proceedings of the Royal Society B: Biological Sciences, 284(1848), 20162232. 10.1098/rspb.2016.2232

Ling, N., Tempero, G. W., & Schamhart, T. (2023). Using faecal DNA metabarcoding to determine the diet of the long-tailed bat, Chalinolobus tuberculatus. New Zealand Journal of Zoology, 52(1), 55–62. 10.1080/03014223.2023.2240711

Lion, T. (2014). Adenovirus Infections in Immunocompetent and Immunocompromised Patients. Clinical Microbiology Reviews, 27(3), 441–462. 10.1128/CMR.00116-13

Lučan, R. K., Jor, T., Romportl, D., & Morelli, F. (2024). Use of synanthropic roosts by bats in Europe and North America. Mammal Review, 55(3), e12380. 10.1111/mam.12380

Lynch, J. P., & Kajon, A. E. (2016). Adenovirus: Epidemiology, Global Spread of Novel Serotypes, and Advances in Treatment and Prevention. Seminars in Respiratory and Critical Care Medicine, 37(4), 586–602. 10.1055/s-0036-1584923

Majidi, S., Rahiminejad, V., Razavi, E., & Nadimi, A. (2024). Heterostigmatic mites (Acari: Prostigmata) associated with mushroom-forming fungi (Basidiomycota: Agaricomycetes), with description of a new species of microdispid mite (Microdispidae). Biologia, 79(5), 1379–1389. 10.1007/s11756-024-01626-4

Marchiori, C. H. (2023). The characteristics of the families Anisopodidae and Mycetophilidae (Insecta: Diptera). International Journal of Life Science Research Archive, 5(2), 096–108. 10.53771/ijlsra.2023.5.2.0106

Martin, P., & Gerecke, R. (2009). Diptera as hosts of water mite larvae—An interesting relationship with many open questions. Lauterbornia, 68, 95–103.

Maslo, B., Mau, R. L., Kerwin, K., McDonough, R., McHale, E., & Foster, J. T. (2022). Bats provide a critical ecosystem service by consuming a large diversity of agricultural pest insects. Agriculture, Ecosystems & Environment, 324, 107722. 10.1016/j.agee.2021.107722

McFarlane, D. A., & Lundberg, J. (2024). Rates of diagenesis of tropical insectivorous bat guano accumulations: Implications for potential paleoenvironmental reconstruction. International Journal of Speleology, 53(1), 39–49. 10.5038/1827-806X.53.1.2494

McHale, E., Kwait, R., Kerwin, K., Kyle, K., Crosby, C., & Maslo, B. (2025). Detection of Spotted Lanternfly (Lycorma delicatula) by Bats: A qPCR Approach to Forest Pest Surveillance. Forests, 16(3), 443. 10.3390/f16030443

Merritt, R. W., Courtney, G. W., & Keiper, J. B. (2009). Chapter 76 - Diptera: (Flies, Mosquitoes, Midges, Gnats). In V. H. Resh & R. T. Cardé (Eds.), Encyclopedia of Insects (Second Edition) (pp. 284–297). Academic Press. 10.1016/B978-0-12-374144-8.00085-0

Münzer, O., Schaetz, B., & Kurta, A. (2016). Consumption of Insect Pests by the Evening Bat (Nycticeius humeralis) in Southeastern Michigan. The Great Lakes Entomologist, 49(1), 36–40. 10.22543/0090-0222.2520

Newman, M. M., Kloepper, L. N., Duncan, M., McInroy, J. A., & Kloepper, J. W. (2018). Variation in Bat Guano Bacterial Community Composition With Depth. Frontiers in Microbiology, 9, 914. 10.3389/fmicb.2018.00914

Nguyen, N. H., Song, Z., Bates, S. T., Branco, S., Tedersoo, L., Menke, J., Schilling, J. S., & Kennedy, P. G. (2016). FUNGuild: An open annotation tool for parsing fungal community datasets by ecological guild. Fungal Ecology, 20, 241–248. 10.1016/j.funeco.2015.06.006

Nickols, W. A., Kuntz, T., Shen, J., Maharjan, S., Mallick, H., Franzosa, E. A., Thompson, K. N., Nearing, J. T., & Huttenhower, C. (2026). MaAsLin 3: Refining and extending generalized multivariable linear models for meta-omic association discovery. Nature Methods, 23(3), 554–564. 10.1038/s41592-025-02923-9

Ogorek, R., Dylag, M., Kozak, B., Visnovska, Z., Tancinova, D., & Lejman, A. (2016). Fungi isolated and quantified from bat guano and air in Harmanecká and Driny Caves (Slovakia). Journal of Cave and Karst Studies, 78(1), 41–49. 10.4311/2015MB0108

O’Rourke, D., Rouillard, N. P., Parise, K. L., & Foster, J. T. (2022). Spatial and temporal variation in New Hampshire bat diets. Scientific Reports, 12(1), 14334. 10.1038/s41598-022-17631-z

Ozimek, E., & Hanaka, A. (2020). Mortierella Species as the Plant Growth-Promoting Fungi Present in the Agricultural Soils. Agriculture, 11(1), 7. 10.3390/agriculture11010007

Pedregosa, F., Pedregosa, F., Varoquaux, G., Varoquaux, G., Org, N., Gramfort, A., Gramfort, A., Michel, V., Michel, V., Fr, L., Thirion, B., Thirion, B., Grisel, O., Grisel, O., Blondel, M., Prettenhofer, P., Prettenhofer, P., Weiss, R., Dubourg, V., … Cournapeau, D. (2011). Scikit-learn: Machine Learning in Python. MACHINE LEARNING IN PYTHON. 10.5555/1953048.2078195

Pentinsaari, M., Mutanen, M., & Kaila, L. (2014). Cryptic diversity and signs of mitochondrial introgression in the Agrilus viridis species complex (Coleoptera: Buprestidae). European Journal of Entomology, 111(4), 475–486. 10.14411/eje.2014.072

Pénzes, J. J., Söderlund-Venermo, M., Canuti, M., Eis-Hübinger, A. M., Hughes, J., Cotmore, S. F., & Harrach, B. (2020). Reorganizing the family Parvoviridae: A revised taxonomy independent of the canonical approach based on host association. Archives of Virology, 165(9), 2133–2146. 10.1007/s00705-020-04632-4

Pike, N. (2011). Using false discovery rates for multiple comparisons in ecology and evolution. Methods in Ecology and Evolution, 2(3), 278–282. 10.1111/j.2041-210X.2010.00061.x

Portik, D. M., Brown, C. T., & Pierce-Ward, N. T. (2022). Evaluation of taxonomic classification and profiling methods for long-read shotgun metagenomic sequencing datasets. BMC Bioinformatics, 23(1), 541. 10.1186/s12859-022-05103-0

Puig-Montserrat, X., Flaquer, C., Gómez-Aguilera, N., Burgas, A., Mas, M., Tuneu, C., Marquès, E., & López-Baucells, A. (2020). Bats actively prey on mosquitoes and other deleterious insects in rice paddies: Potential impact on human health and agriculture. Pest Management Science, 76(11), 3759–3769. 10.1002/ps.5925

Pyszko, P., Šigutová, H., Ševčík, J., Drgová, M., Hařovská, D., & Drozd, P. (2025). Ambrosia gall midges (Diptera: Cecidomyiidae) and their microbial symbionts as a neglected model of fungus-farming evolution. FEMS Microbiology Reviews, 49, fuaf010. 10.1093/femsre/fuaf010

Ratcliffe, J. M., & Dawson, J. W. (2003). Behavioural flexibility: The little brown bat, Myotis lucifugus, and the northern long-eared bat,M.septentrionalis, both glean and hawk prey. Animal Behaviour, 66(5), 847–856. 10.1006/anbe.2003.2297

Ratnasingham, S., Wei, C., Chan, D., Agda, J., Agda, J., Ballesteros-Mejia, L., Boutou, H. A., El Bastami, Z. M., Ma, E., Manjunath, R., Rea, D., Ho, C., Telfer, A., McKeowan, J., Rahulan, M., Steinke, C., Dorsheimer, J., Milton, M., & Hebert, P. D. N. (2024). BOLD v4: A Centralized Bioinformatics Platform for DNA-Based Biodiversity Data. In R. DeSalle (Ed.), DNA Barcoding (Vol. 2744, pp. 403–441). Springer US. 10.1007/978-1-0716-3581-0_26

Richardson, M. (2002). The coprophilous succession. Fungal Diversity. 10. 101–111.

Rocke, T. E., Kingstad-Bakke, B., Wüthrich, M., Stading, B., Abbott, R. C., Isidoro-Ayza, M., Dobson, H. E., dos Santos Dias, L., Galles, K., Lankton, J. S., Falendysz, E. A., Lorch, J. M., Fites, J. S., Lopera-Madrid, J., White, J. P., Klein, B., & Osorio, J. E. (2019). Virally-vectored vaccine candidates against white-nose syndrome induce anti-fungal immune response in little brown bats (Myotis lucifugus). Scientific Reports, 9(1), 6788. 10.1038/s41598-019-43210-w

Roev, G. V., Borisova, N. I., Chistyakova, N. V., Agletdinov, M. R., Akimkin, V. G., & Khafizov, K. (2023). Unlocking the Viral Universe: Metagenomic Analysis of Bat Samples Using Next-Generation Sequencing. Microorganisms, 11(10), 2532. 10.3390/microorganisms11102532

Rydell, J., McNeill, D. P., & Eklöf, J. (2006). Capture success of little brown bats (Myotis lucifugus) feeding on mosquitoes. Journal of Zoology, 256(3), 379–381. 10.1017/S0952836902000419

Sæther, O. A. (2001). Phylogeny of the subfamilies of Chironomidae (Diptera). Systematic Entomology, 25(3), 393–403. 10.1046/j.1365-3113.2000.00111.x

Salgado-Salazar, C., Rossman, A. Y., & Chaverri, P. (2013). Not as Ubiquitous as We Thought: Taxonomic Crypsis, Hidden Diversity and Cryptic Speciation in the Cosmopolitan Fungus Thelonectria discophora (Nectriaceae, Hypocreales, Ascomycota). PLOS ONE, 8(10), e76737. 10.1371/journal.pone.0076737

Schloss, P. D., Westcott, S. L., Ryabin, T., Hall, J. R., Hartmann, M., Hollister, E. B., Lesniewski, R. A., Oakley, B. B., Parks, D. H., Robinson, C. J., Sahl, J. W., Stres, B., Thallinger, G. G., Van Horn, D. J., & Weber, C. F. (2009). Introducing mothur: Open-Source, Platform-Independent, Community-Supported Software for Describing and Comparing Microbial Communities. Applied and Environmental Microbiology, 75(23), 7537–7541. 10.1128/AEM.01541-09

Shaw, J., Marin, M. G., & Li, H. (2026). High-resolution metagenome assembly for modern long reads with myloasm. Nature Biotechnology. 10.1038/s41587-026-03053-z

Soto-López, J. D., García-Martín, J. M., Lizana-Ciudad, D., Lizana, M., Hernández-Tabernero, L., Fernández-Soto, P., Velásquez-González, O. E., Aragón, S. L., Belhassen-García, M., & Muro, A. (2025). Taxonomic and functional profiling of bat guano microbiota from hiking trail-associated tunnels: A potential risk for human health? Environmental Microbiome, 20(1), 123. 10.1186/s40793-025-00782-7

Stidsholt, L., Hubancheva, A., Greif, S., Goerlitz, H. R., Johnson, M., Yovel, Y., & Madsen, P. T. (2023). Echolocating bats prefer a high risk-high gain foraging strategy to increase prey profitability. eLife, 12, e84190. 10.7554/eLife.84190

Sun, S., Shan, M., Huang, Z., Lv, Y., Wei, Z., Shen, M., Sun, K., Li, Z., & Feng, J. (2025). Screening microbial inhibitors of Pseudogymnoascus destructans in Northern China. Microbiology Spectrum, 13(12), e01241–25. 10.1128/spectrum.01241-25

Telagathoti, A., Probst, M., & Peintner, U. (2021). Habitat, Snow-Cover and Soil pH, Affect the Distribution and Diversity of Mortierellaceae Species and Their Associations to Bacteria. Frontiers in Microbiology, 12. 10.3389/fmicb.2021.669784

Tuttle, N. M., Benson, D. P., & Sparks, D. W. (2006). Diet of the Myotis sodalis (Indiana Bat) at an Urban/Rural Interface. Northeastern Naturalist, 13(3), 435–442. 10.1656/1092-6194(2006)13%255B435:DOTMSI%255D2.0.CO;2

Urbina, J., Chestnut, T., Allen, J. M., & Levi, T. (2021). Pseudogymnoascus destructans growth in wood, soil and guano substrates. Scientific Reports, 11(1), 763. 10.1038/s41598-020-80707-1

Urbina, J., Chestnut, T., Schwalm, D., Allen, J., & Levi, T. (2020). Experimental evaluation of genomic DNA degradation rates for the pathogen Pseudogymnoascus destructans (Pd) in bat guano. PeerJ, 8, e8141. 10.7717/peerj.8141

Vanderwolf, K. J., Malloch, D., & McAlpine, D. F. (2016). Ectomycota Associated with Arthropods from Bat Hibernacula in Eastern Canada, with Particular Reference to Pseudogymnoascus destructans. Insects, 7(2), 16. 10.3390/insects7020016

Varusk, S., Sammet, K., Ariyan, M., Mubarak Alwutayd, K., & Anslan, S. (2025). DNA metabarcoding of mites from small soil samples: Limited agreement with morphological identifications but improved results from long-read sequencing. PeerJ, 13, e20205. 10.7717/peerj.20205

Vasquez, A. A., Carmona-Galindo, V., Qazazi, M. S., Walker, X. N., & Ram, J. L. (2020). Water mite assemblages reveal diverse genera, novel DNA barcodes and transitional periods of intermediate disturbance. Experimental and Applied Acarology, 80(4), 491–507. 10.1007/s10493-020-00476-4

Vazquez, J. M., Lauterbur, M. E., Mottaghinia, S., Gaucherand, L., Maesen, S., Singer, M., Villa, S., Bucci, M., Fraser, D., Gray-Sandoval, G., Haidar, Z. R., Han, M., Kohler, W., Lama, T. M., Le Corf, A., Loyer, C., McMillan, D., Li, S., Lo, J., … Sudmant, P. H. (2026). Extensive longevity and DNA virus-driven adaptation in nearctic Myotis bats. bioRxiv, 2024.10.10.617725. 10.1101/2024.10.10.617725

Verant, M. L., Boyles, J. G., Waldrep, W., Wibbelt, G., & Blehert, D. S. (2012). Temperature-Dependent Growth of Geomyces destructans, the Fungus That Causes Bat White-Nose Syndrome. PLoS ONE, 7(9), e46280. 10.1371/journal.pone.0046280

Verhoeven, K. J. F., Simonsen, K. L., & McIntyre, L. M. (2005). Implementing false discovery rate control: Increasing your power. Oikos, 108(3), 643–647. 10.1111/j.0030-1299.2005.13727.x

Virtanen, P., Gommers, R., Oliphant, T. E., Haberland, M., Reddy, T., Cournapeau, D., Burovski, E., Peterson, P., Weckesser, W., Bright, J., Van Der Walt, S. J., Brett, M., Wilson, J., Millman, K. J., Mayorov, N., Nelson, A. R. J., Jones, E., Kern, R., Larson, E., … Vázquez-Baeza, Y. (2020). SciPy 1.0: Fundamental algorithms for scientific computing in Python. Nature Methods, 17(3), 261–272. 10.1038/s41592-019-0686-2

Voigt, C. C., Phelps, K. L., Aguirre, L. F., Corrie Schoeman, M., Vanitharani, J., & Zubaid, A. (2015). Bats and Buildings: The Conservation of Synanthropic Bats. In C. C. Voigt & T. Kingston (Eds.), Bats in the Anthropocene: Conservation of Bats in a Changing World (pp. 427–462). Springer International Publishing. 10.1007/978-3-319-25220-9_14

Wang, Q., & Cole, J. R. (2024). Updated RDP taxonomy and RDP Classifier for more accurate taxonomic classification. Microbiology Resource Announcements, 13(4), e01063–23. 10.1128/mra.01063-23

Warwick-Dugdale, J., Solonenko, N., Moore, K., Chittick, L., Gregory, A. C., Allen, M. J., Sullivan, M. B., & Temperton, B. (2019). Long-read viral metagenomics captures abundant and microdiverse viral populations and their niche-defining genomic islands. PeerJ, 7, e6800. 10.7717/peerj.6800

Webster, J. M., & Whitaker, J. O. (2005). Study of Guano Communities of Big Brown Bat Colonies in Indiana and Neighboring Illinois Counties. Northeastern Naturalist, 12(2), 221–232. 10.1656/1092-6194(2005)012%255B0221:SOGCOB%255D2.0.CO;2

Westcott, S. L., & Schloss, P. D. (2017). OptiClust, an Improved Method for Assigning Amplicon-Based Sequence Data to Operational Taxonomic Units. mSphere, 2(2). 10.1128/mspheredirect.00073-17

Wetzler, G. C., & Boyles, J. G. (2017). The energetics of mosquito feeding by insectivorous bats. Canadian Journal of Zoology, 96(4), 373–377. 10.1139/cjz-2017-0162

Whitaker, J. O. (1995). Food of the Big Brown Bat Eptesicus fuscus from Maternity Colonies in Indiana and Illinois. The American Midland Naturalist, 134(2), 346–360. 10.2307/2426304

White, T. J., Bruns, T. D., Lee, S. B., & Taylor, J. W. (1990). Amplification and direct sequencing of fungal ribosomal RNA Genes for phylogenetics. PCR – Protocols and Applications – A Laboratory Manual. 315–322. https://www.researchgate.net/publication/223397588_White_T_J_T_D_Bruns_S_B_Lee_and_J_W_Taylor_Amplification_and_direct_sequencing_of_fungal_ribosomal_RNA_Genes_for_phylogenetics

Williams-Guillén, K., Olimpi, E., Maas, B., Taylor, P. J., & Arlettaz, R. (2015). Bats in the Anthropogenic Matrix: Challenges and Opportunities for the Conservation of Chiroptera and Their Ecosystem Services in Agricultural Landscapes. In C. C. Voigt & T. Kingston (Eds.), Bats in the Anthropocene: Conservation of Bats in a Changing World (pp. 151–186). Springer International Publishing. 10.1007/978-3-319-25220-9_6

Wirbel, J., Hickey, A. S., Chang, D., Enright, N. J., Dvorak, M., Chanin, R. B., Schmidtke, D. T., & Bhatt, A. S. (2025). Long-read metagenomics reveals phage dynamics in the human gut microbiome. Nature, 649(8098), 982–990. 10.1038/s41586-025-09786-2

Wray, A. K., Peery, M. Z., Jusino, M. A., Kochanski, J. M., Banik, M. T., Palmer, J. M., Lindner, D. L., & Gratton, C. (2021). Predator preferences shape the diets of arthropodivorous bats more than quantitative local prey abundance. Molecular Ecology, 30(3), 855–873. 10.1111/mec.15769

Zaragoza-Solas, A., Haro-Moreno, J. M., Rodriguez-Valera, F., & López-Pérez, M. (2022). Long-Read Metagenomics Improves the Recovery of Viral Diversity from Complex Natural Marine Samples. mSystems, 7(3), e00192–22. 10.1128/msystems.00192-22

