## Supplementary material for "Microbial, dietary insect, and pathogen communities in fresh and decomposing guano of anthropic little brown bat (Myotis lucifugus) maternity colonies": FileS2

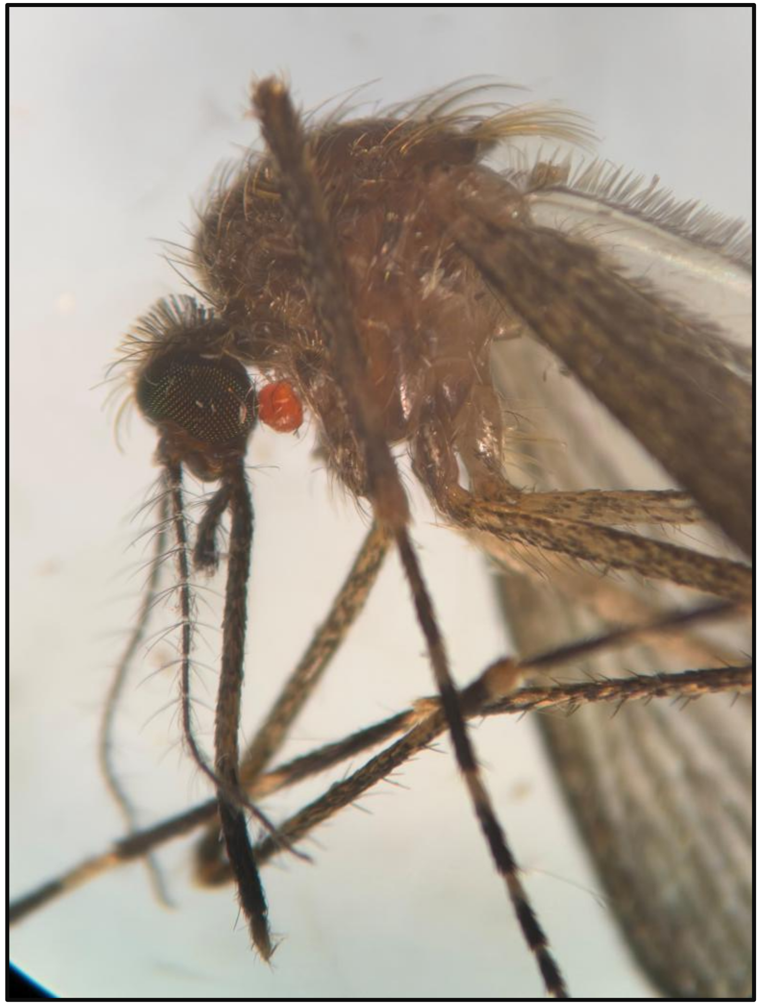

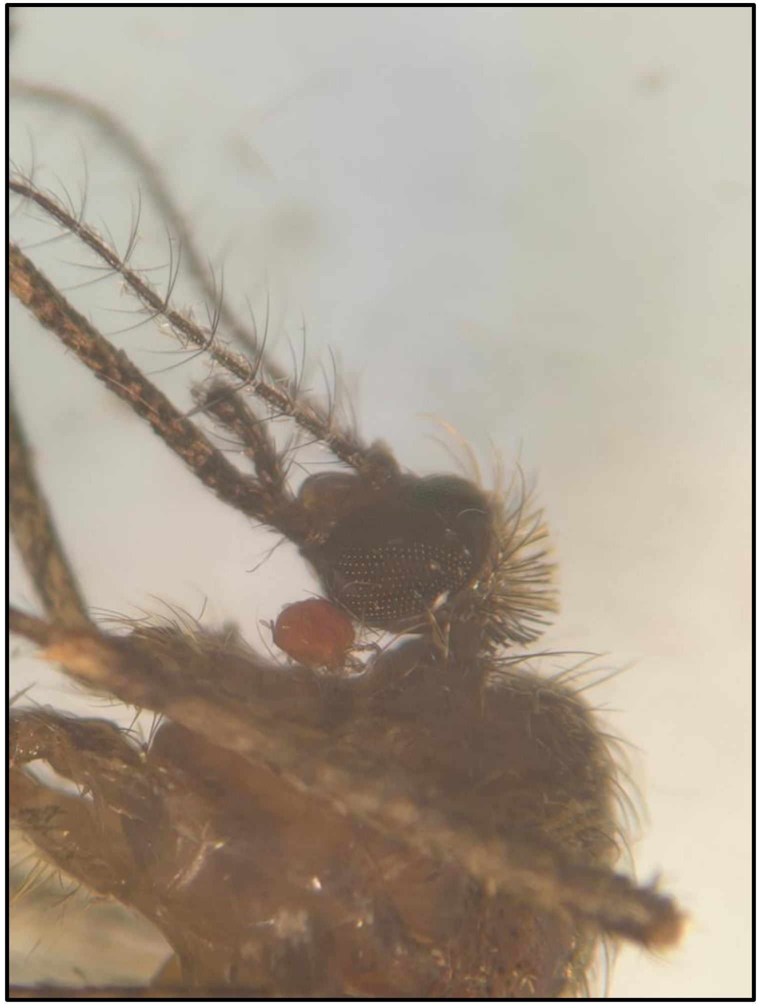

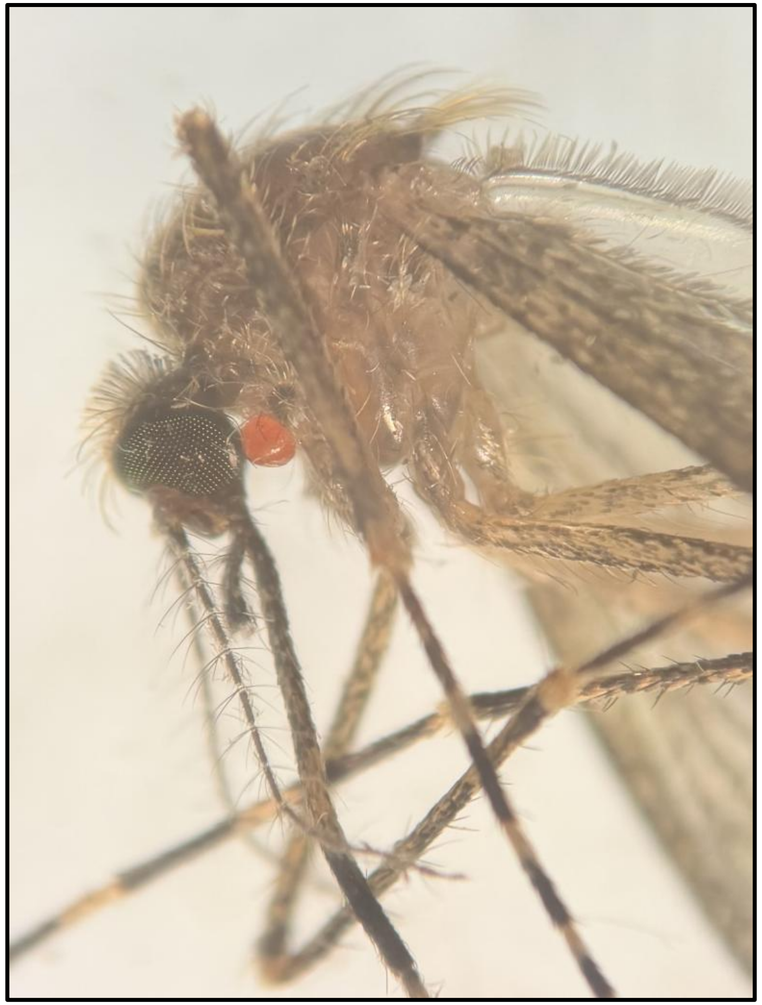

**Photographer**: Jacqui Barbieri, Seasonal Research Technician, The White Memorial Conservation Center, Inc., 80 Whitehall Rd., Litchfield, CT 06759. **Collection Date**: June 20, 2026. **Specimen Collector**: Craig Lehman, Wiederhold Bat Conservation Intern, The White Memorial Conservation Center, Inc., 80 Whitehall Rd., Litchfield, CT 06759. **Location**: 41.719295, -73.215028
**Site Description**: The site is located 160 meters north of Bantam Lake, 560 meters SSW of the White Memorial Conservation Center near the intersection of Lake (yellow blaze) and Windmill Hill Trails (green blaze). Wetland glade opening with 25% water surface and herbaceous vegetation consisting of sedges, ferns, skunk cabbage, bedstraw, and some snakeroot. Nearby tree canopy consisted of maples, hickory, oak, birch, and dense dogwood shrub.
