## Supplementary material for "Microbial, dietary insect, and pathogen communities in fresh and decomposing guano of anthropic little brown bat (Myotis lucifugus) maternity colonies": FileS1

### Supplemental Methods

#### *16S and ITS: PCR Methods*

Samples were amplified in triplicate 15 µL reactions using Go-Taq DNA polymerase (Promega) with the addition of bovine serum albumin (3.3 µg; New England BioLabs); triplicate PCR products from each biological sample were pooled prior to sequencing, such that reads from technical PCR replicates are combined at the bench level for both 16S and ITS workflows. Primers without indices or adapters (0.1 pmol) were added to the master mix to overcome inhibition from host DNA. The v4 16S PCR reaction was incubated at 95˚C for 2 minutes, then 30 cycles of 30 s at 95.0°C, 30 s at 50.0°C and 60 s at 72.0°C, followed by final extension as 72.0°C for 10 minutes. The ITS2 PCR reaction was incubated at 95˚C for 2 minutes, then 5 cycles of 30 s at 95.0°C, 60 s at 48.0°C and 60 s at 72.0°C, then 25 cycles of 30 s at 95.0°C, 60 s at 55.0°C and 60 s at 72.0°C followed by final extension as 72.0°C for 10 minutes.

#### *CO1 PCR Methods*

Made by Kendra Maas, David Askew, and Carlos Garcia-Robledo
Modifications by Kendra Maas, Kristiana Stover, and Ruiwen Lin

**Reagents**
GoTaq DNA Polymerase enzyme
Promega 5X polymerase buffer
ThermoFisher BSA (20mg/ml)
$MgSO_4$ (100mM)
Promega dNTPs (10mM)
LepF1NanoporeF primer (10µM)
LepR1NanoporeR primer (10µM)
LCO1490NanoporeF primer (10µM)
HCO2198RNanoporeR primer (10µM)

**Protocol**
For each sample, the master mix consisted of these following reagents and volumes: 10µL 5X buffer, 0.05µL $MgSO_4$, 1µL dNTP, 0.00625µL LepF1NanoporeF primer, 0.00625µL LepR1NanoporeR primer, 0.00625µL LCO1490NanoporeF primer, 0.00625µL HCO2198RNanoporeR, 0.4µL GoTaq, 30.45µL $H_2O$. Then add 5µL of sample, pipette mix, spin down, and start thermocycler reaction.

**Cycling Conditions**
The thermocycler was set to the following directions with a lid temperature of 102°C: 95°C for 2 minutes, 4 cycles of: 95°C for 30 seconds, 48°C for 40 seconds, 72°C for 1 minute; then 34 cycles of: 95°C for 30 seconds, 51°C for 40 seconds, 72°C for 1 minute; 72°C for 5 minutes, hold at 4°C indefinitely.

**KingFisher Robot Bead Cleaning**:
0.6 bead-to-sample ratio (20μL sample: 12μL bead + serapure mix)
200μL of 80% ethanol washed twice

| **npPCRbc12**<br>Lid Temp = 105°C | |
|---|---|
| **Step** | **Parameter** |
| 1 | Incubate at 95°C for 2 min |
| 2 | Incubate at 95°C for 30 sec |
| 3 | Incubate at 62°C for 1 min |
| 4 | Incubate at 72°C for 1 min |
| 5 | Repeat steps 2-4 for 12 cycles |
| 6 | Incubate at 72°C for 10 min |
| 7 | Hold at 12°C indefinitely |
| 8 | Finish |

| **Amplicon Sequencing Master Mix** | |
|---|---|
| **Reagent** | **Volume per sample (μL)** |
| 5X Buffer | 10 |
| MgSO4 | 0.05 |
| dNTP | 1 |
| HCO2198 primer | 0.0625 |
| LCO1490 primer | 0.0625 |
| lepF primer | 0.0625 |
| lepR primer | 0.0625 |
| GoTaq | 0.4 |
| H2O | 30.45 |

| | |
|---|---|
| Sample | 5 |

| CGMITES40x<br>Lid Temp = 102°C | |
|---|---|
| **Step** | **Parameter** |
| 1 | Incubate at 95°C for 2 min |
| 2 | Incubate at 95°C for 30 sec |
| 3 | Incubate at 48°C for 40 sec |
| 4 | Incubate at 72°C for 1 min |
| 5 | Repeat step 2-4 for 4 cycles |
| 6 | Incubate at 95°C for 30 sec |
| 7 | Incubate at 51°C for 40 sec |
| 8 | Incubate at 72°C for 1 min |
| 9 | Repeat steps 6-8 for 34 cycles |
| 10 | Incubate at 72°C for 5 min |
| 11 | Hold at 4°C indefinitely |
| 12 | Finish |

***<u>Shotgun Metagenomics Library Preparation Methods and Modifications</u>***

Shotgun metagenomic libraries were prepared with the Oxford Nanopore Native Barcoding Kit 96 V14 (SQK-NBD1114.6). The protocol was followed to the manufacturer's instructions with some modifications, which are detailed below.

***Barcode Ligation***

During the barcode ligation step, no nuclease water was added, and 3.75 uL of end-prepped DNA was added instead of 0.75 uL. The reaction was then incubated for 20 minutes, and 4uL of EDTA was added. For the "screen" metagenome run, only 1uL of barcoded DNA was added. During the high-throughput run, the amount of barcoded DNA added at this step depended on the sequencing success of the screen run. For specific inputs, see Table S1. Following addition of barcoded DNA, 0.6X beads were added opposed to 0.4X to retain longer fragments. Further, 200uL of long fragment buffer. The pellet was then suspended in 16uL of water. Incubation time was increased to 20 minutes.

***Adapter Ligation and Cleaning***

For the screen run, 15uL of the barcoded library was used for adapter ligation. For the high-throughput run, 35uL were used (according to manufacturer suggestion). During the adapter ligation reaction the following volumes were changed for the screen run only: 15uL of pooled barcoded sample, 2.5 uL of Native Adapter, 5 uL of NEB Quick Ligation Reaction Buffer (5X), and 2.5uL of Quick T4 Ligase.

During the cleaning step, 1X of of AMPure XP beads were added. The incubation at 37C was increased to 20 minutes. During library elution, 30uL was retained instead of 15uL. During sequencing, the final throughput run was sequenced on the PromthION, while the screen run was ran on the MinION, both on R10.4 flow cells. Each barcode set (1-24, and 1-24) were ran independently to avoid barcode redundancy.

***qPCR Methods***

Each sample was evaluated in triplicate on a 384 well plate. Each reaction contained 5 µL Luna® Universal Probe qPCR Master Mix, 1 µL primer-probe mix per target (see Table S4 for primer-probe concentrations), and 1 µL organismal DNA from the previous extraction, with water added to bring the total reaction volume to 10 µL. *Histoplasma* spp. and *Blastomyces dermatitidis* were run in one reaction and Pd, spotted lanternfly, and emerald ash borer in a second reaction. Bio-Rad CFX384 Touch Real-Time PCR Detection System was set to denaturation at 95 °C for 1 minute; 39 cycles of denaturation at 95 °C for 10 seconds and annealing at 58 °C for 30 seconds. The melt curve protocol consisted of 0.5 °C increments per cycle between 5 seconds each of 62 °C and 95 °C.

***Spore Count Methods***

In May through August of 2024, spores were collected from 11 bat houses across the ASHF little brown bat colony (BH4-BH14), as well as two little brown bat houses at WMCC (BH1, rocket roost) and a big brown bat colony in a barn at WMCC (High/low bat loft, floor edge/center, far floor, post A/B), using sterile collection swabs on locations within the barn and the inner mesh lining of the bat houses. The conidia were then suspended in a solution of 0.1% Tween surfactant solution, and a hemocytometer was used to quantify conidia and analyze summer variance of persisting Pd over the course of the sampling. Asymmetrical, curved spores visually resembling reference spores (Gargas et al. 2009) were identified as Pd. Culturing attempts were made using 50µl of spore solution at 12°C on potato dextrose agar.
