## Supplementary figures and images for "Microbial, dietary insect, and pathogen communities in fresh and decomposing guano of anthropic little brown bat (Myotis lucifugus) maternity colonies"

### FigureS1A

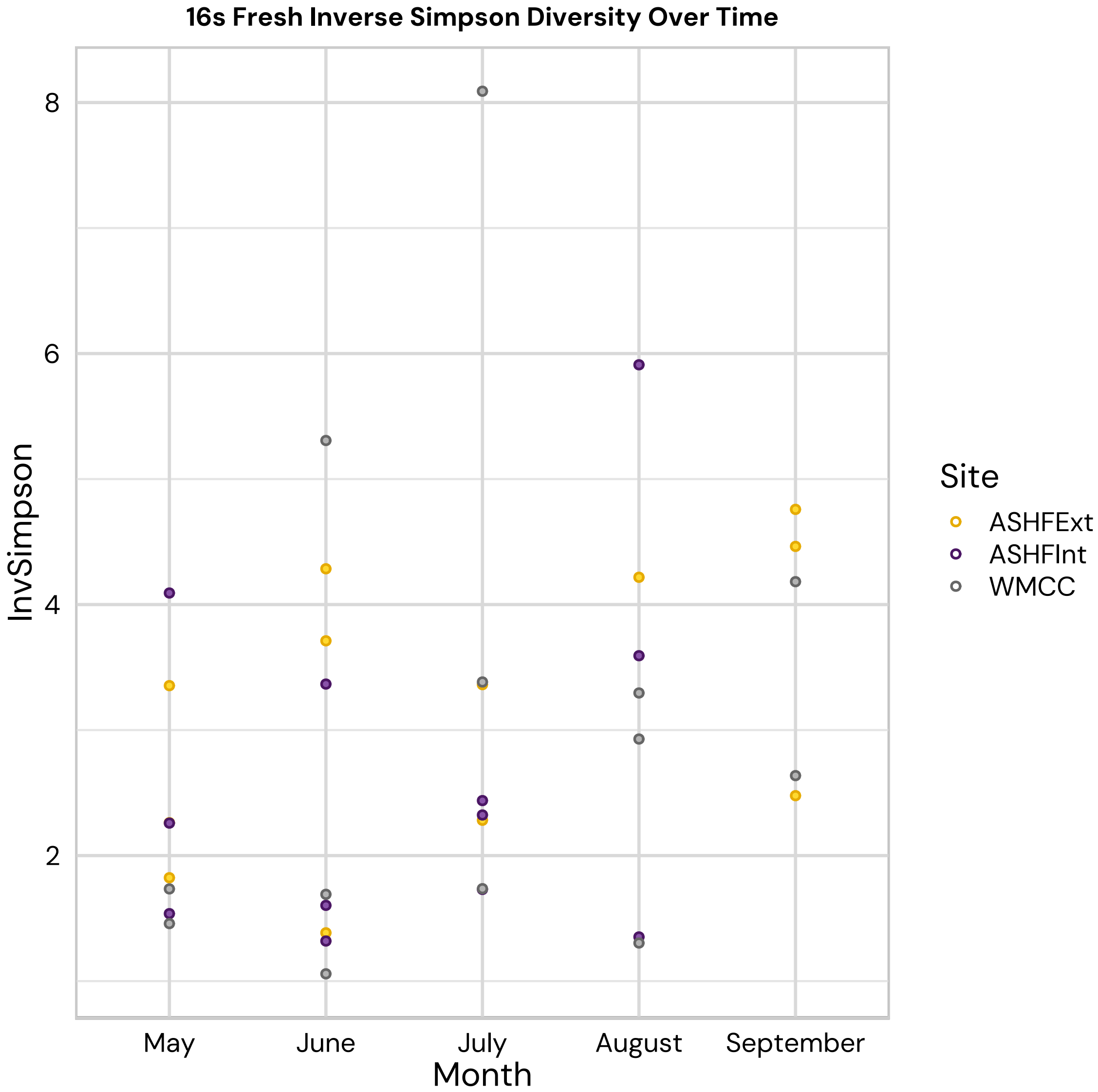

### FigureS1B

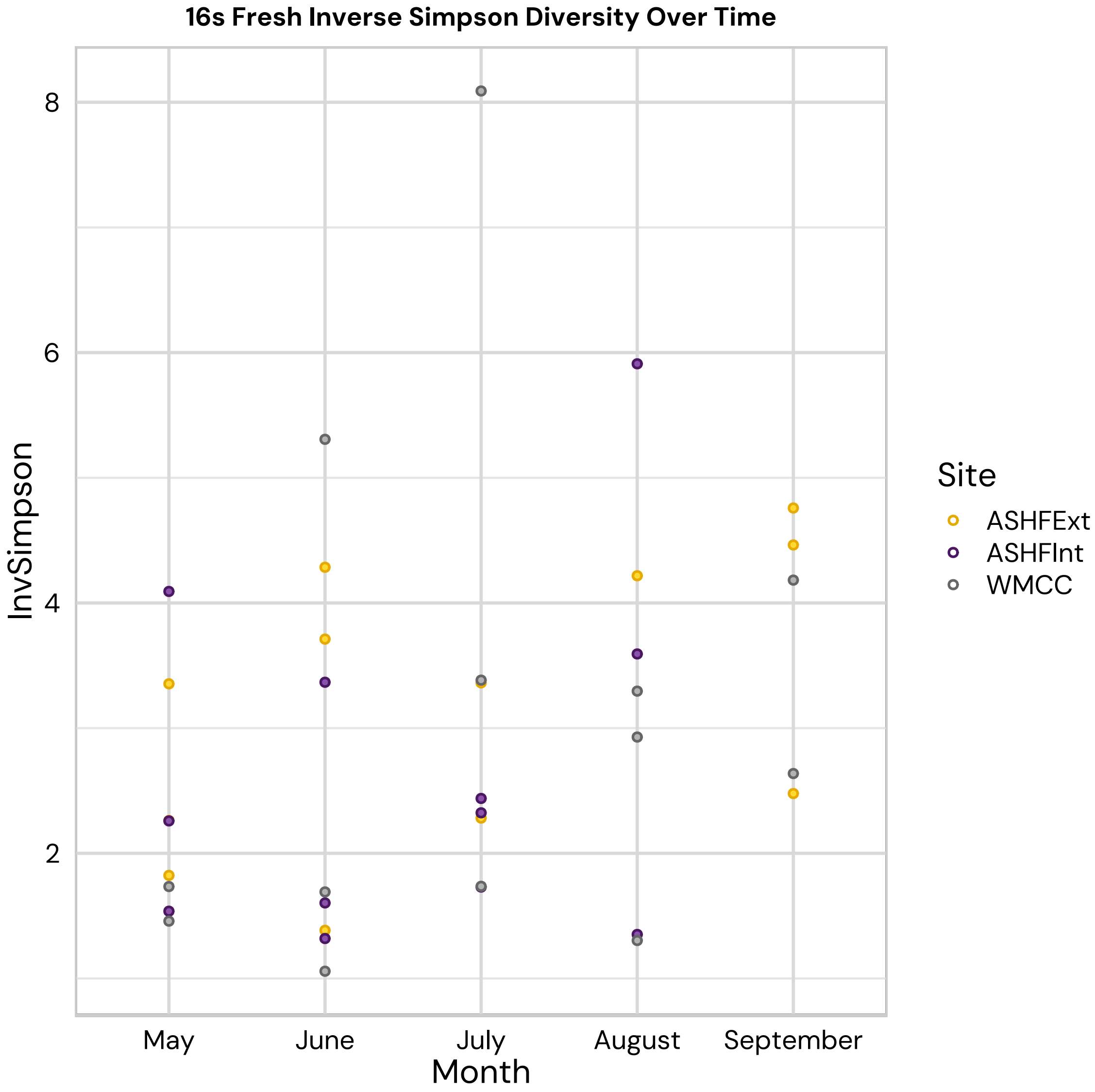

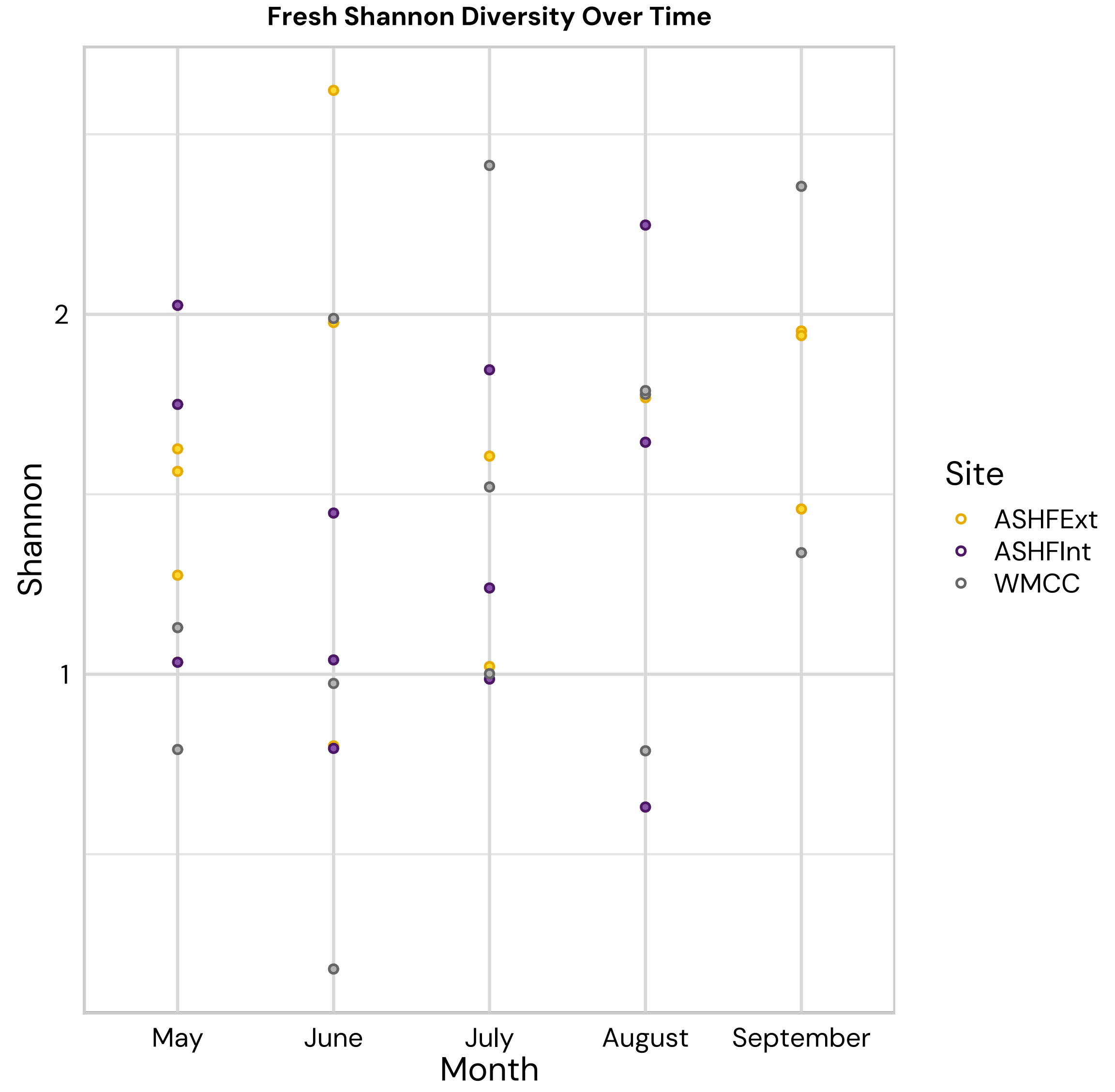

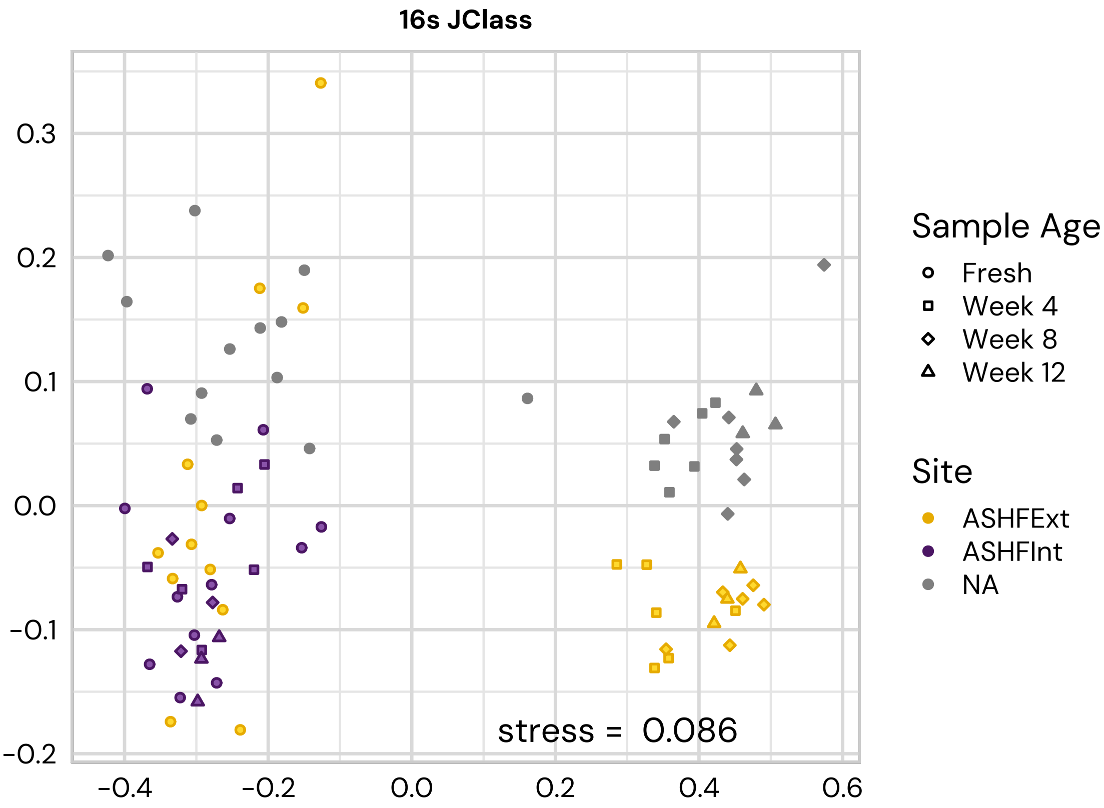

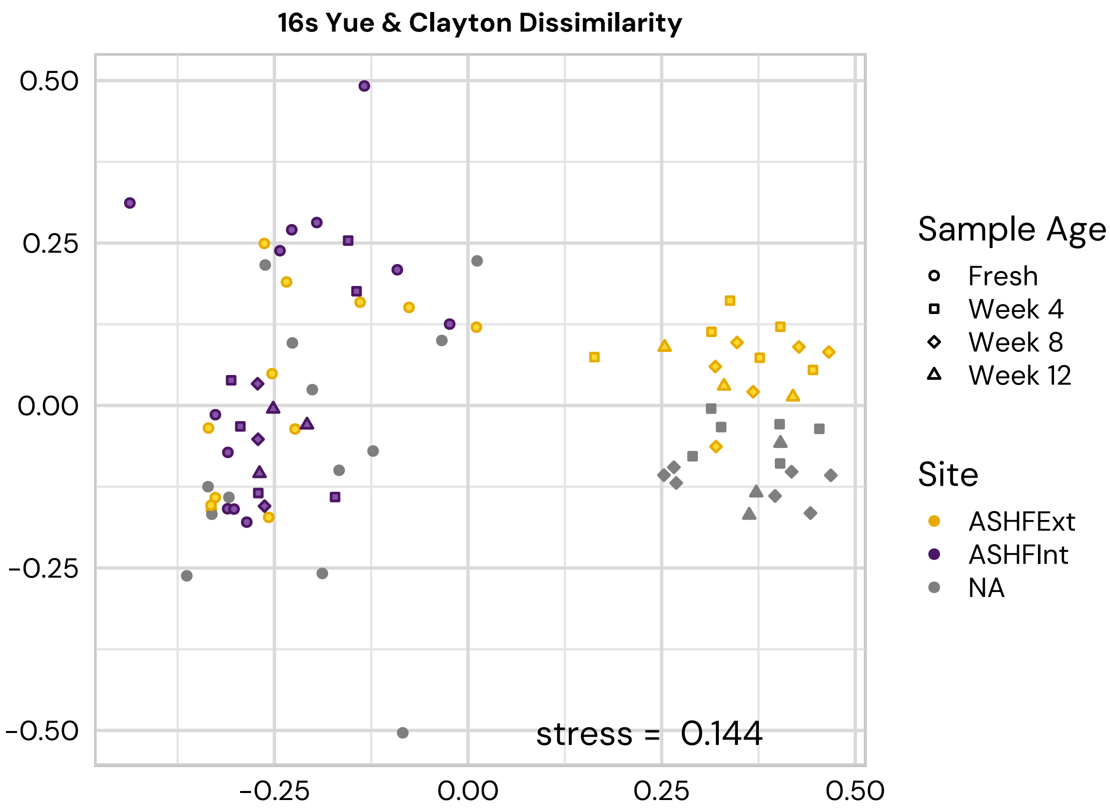

### FigureS1C

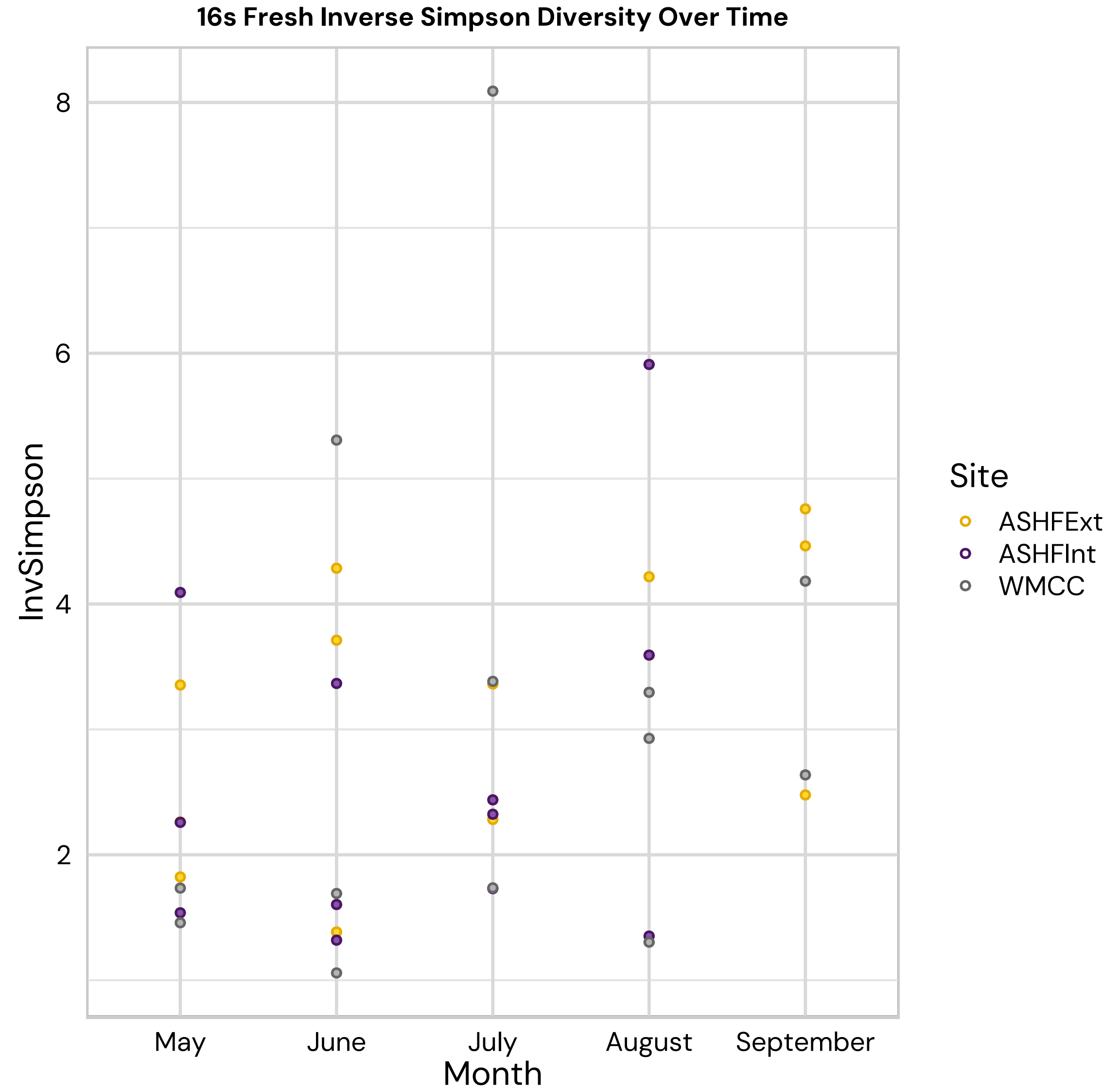

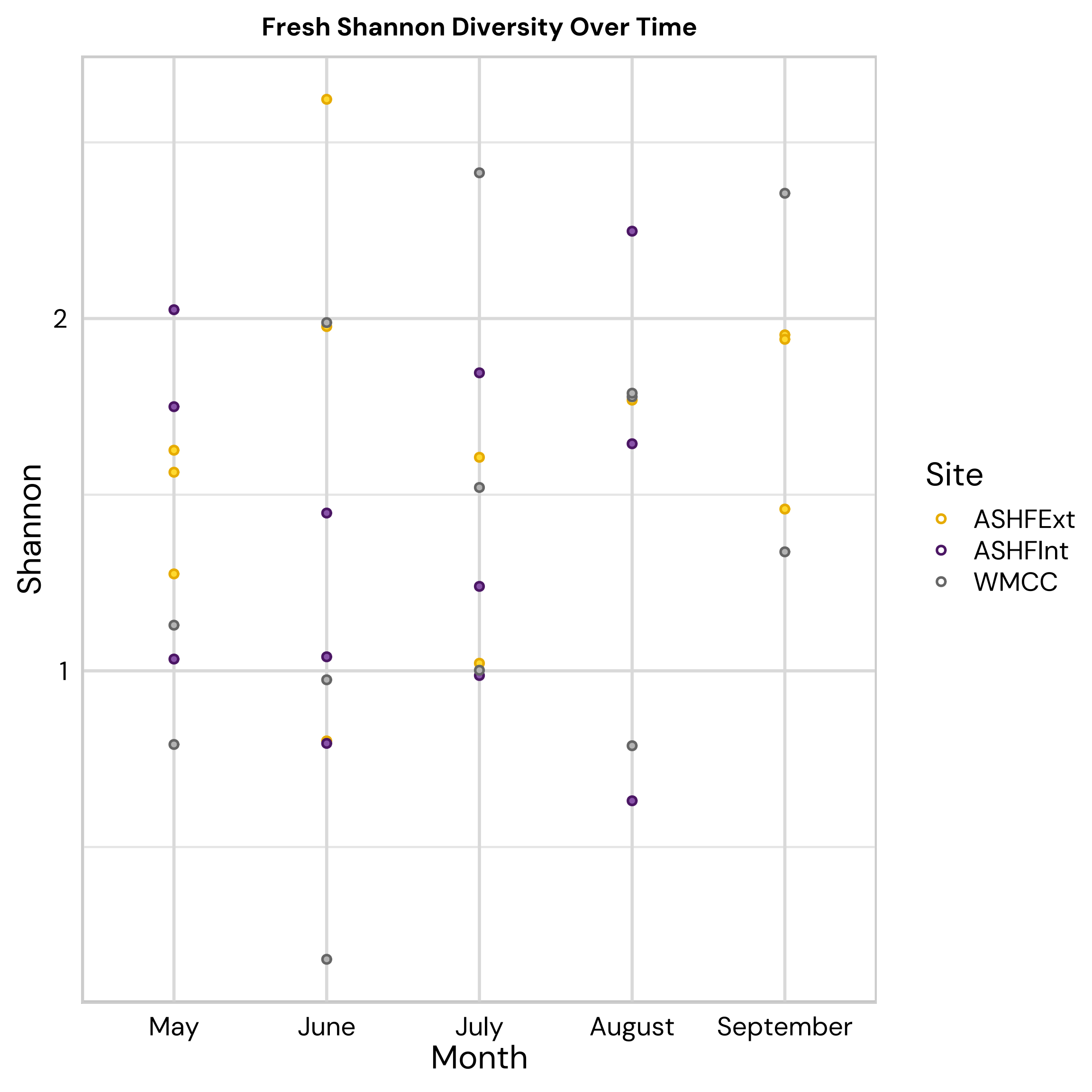

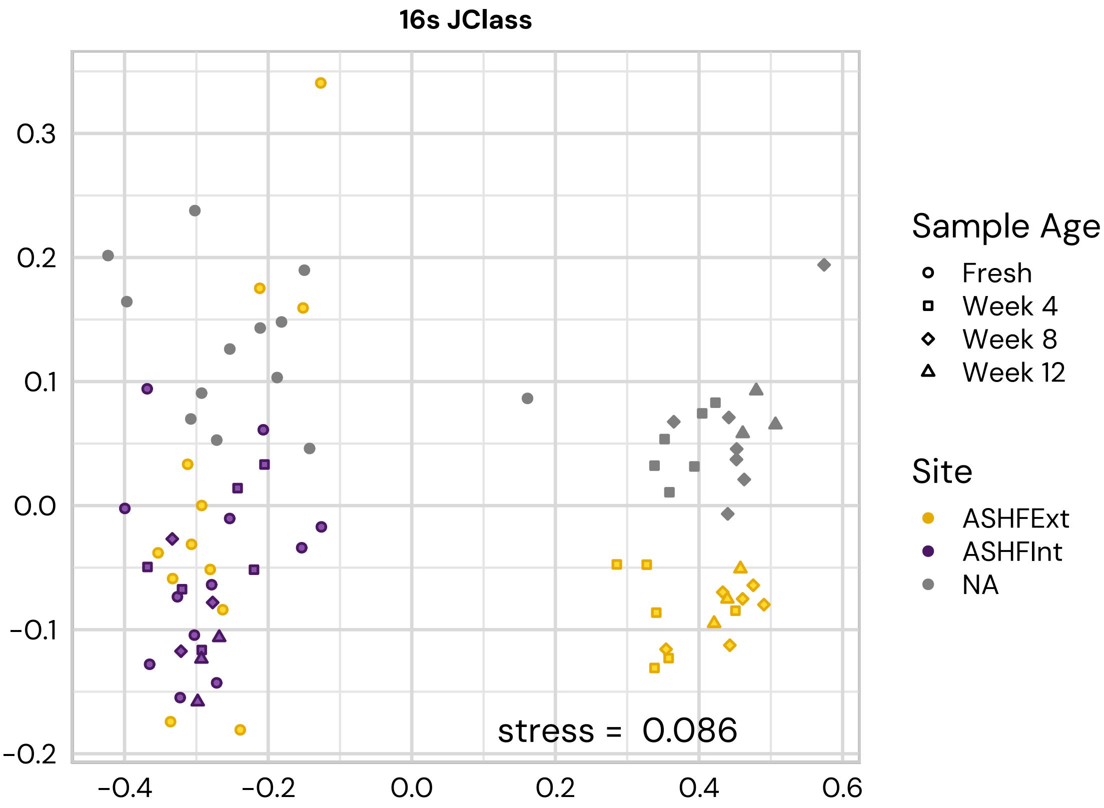

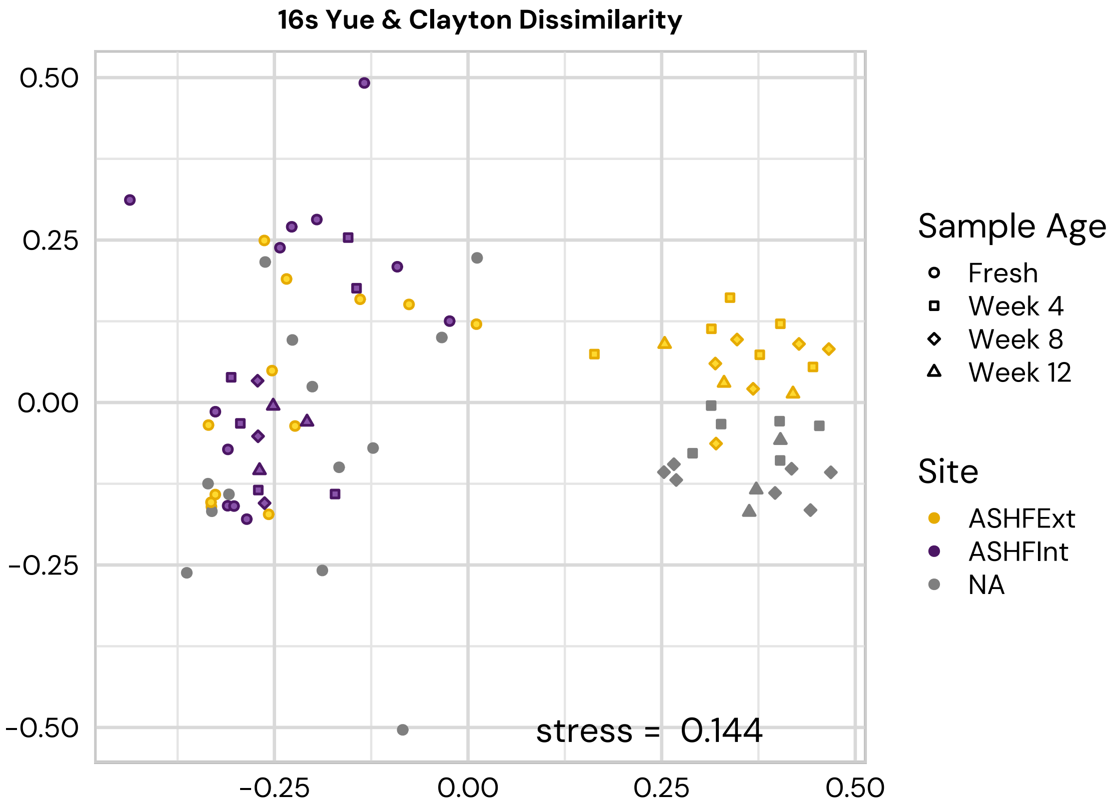

### FigureS1D

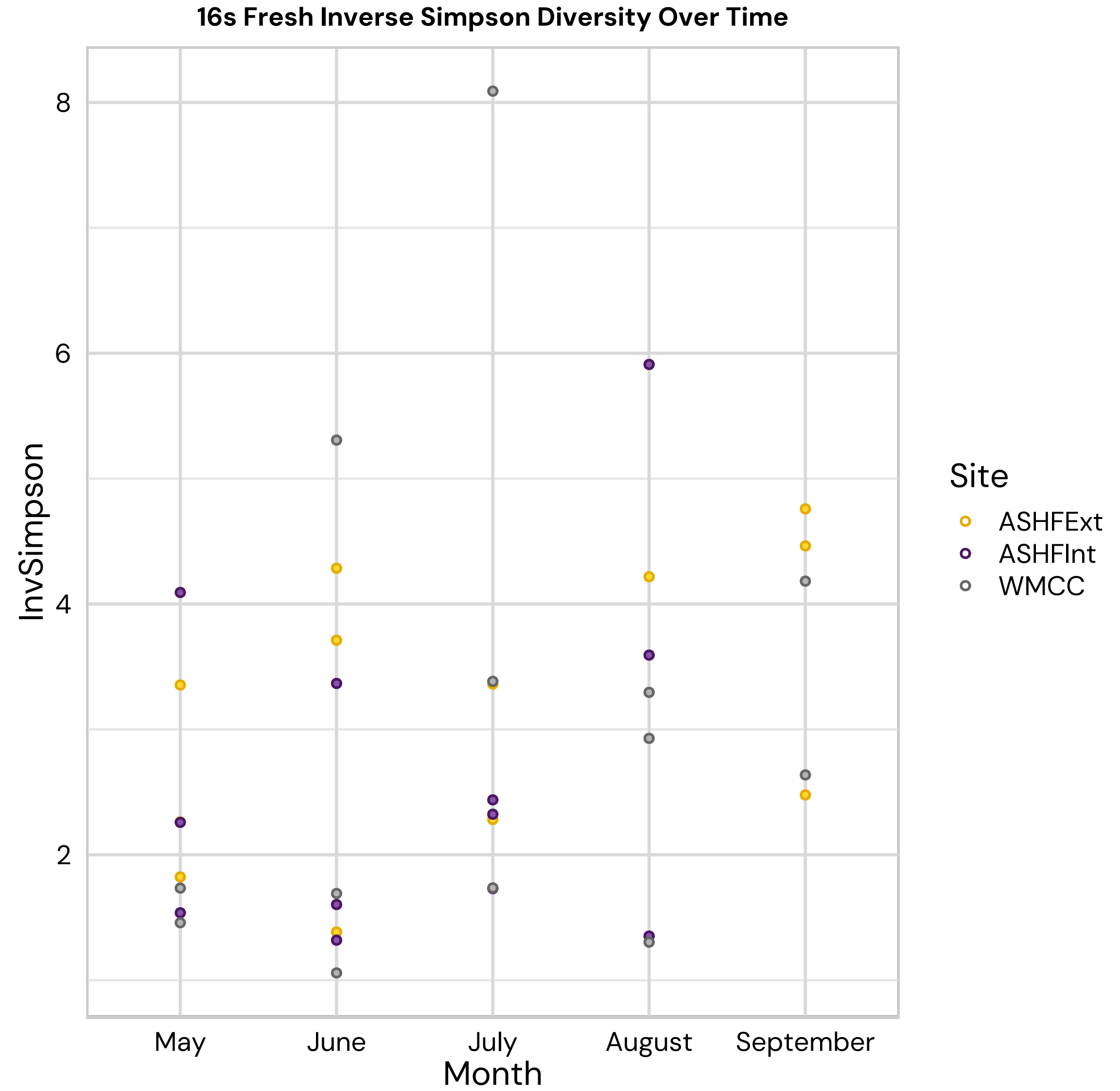

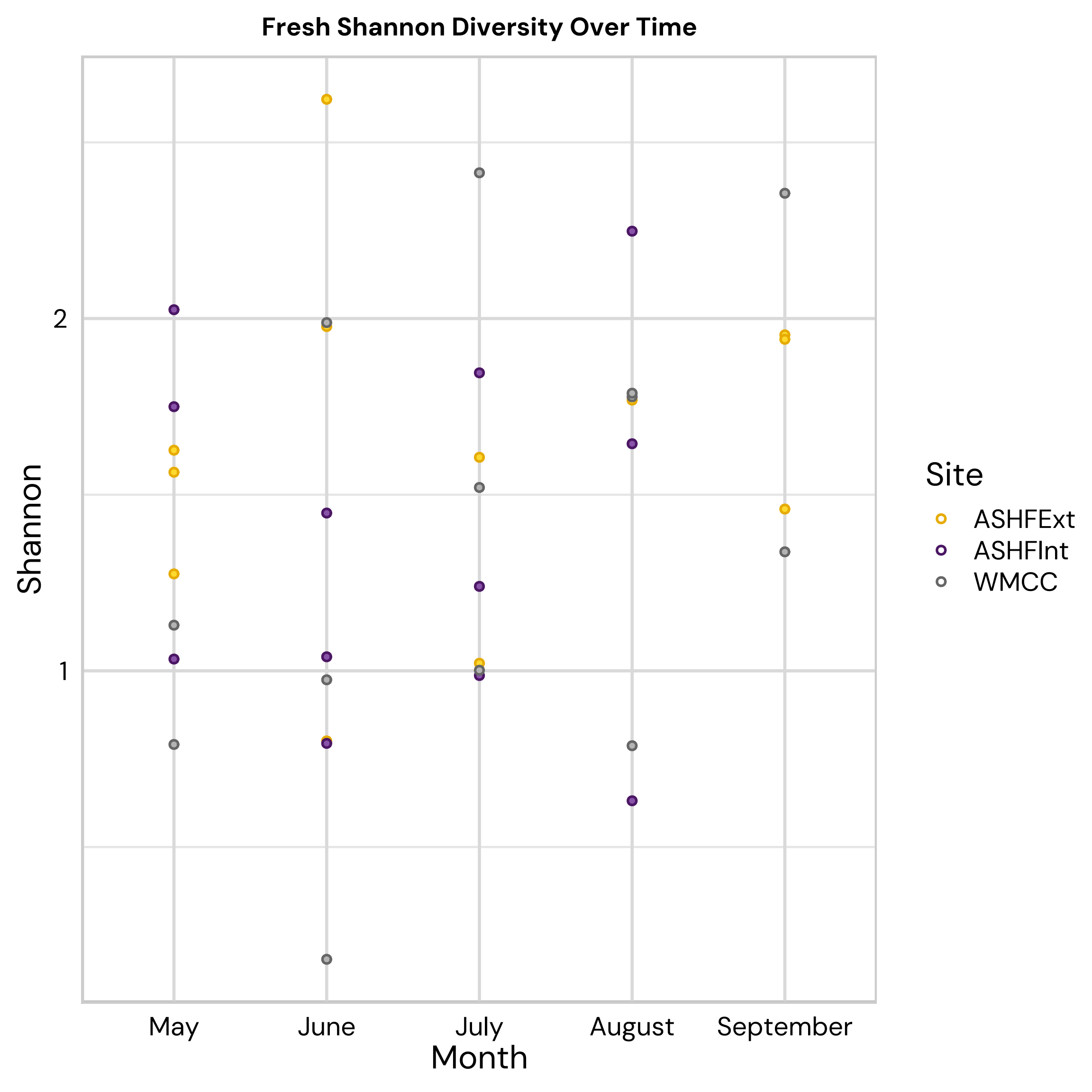

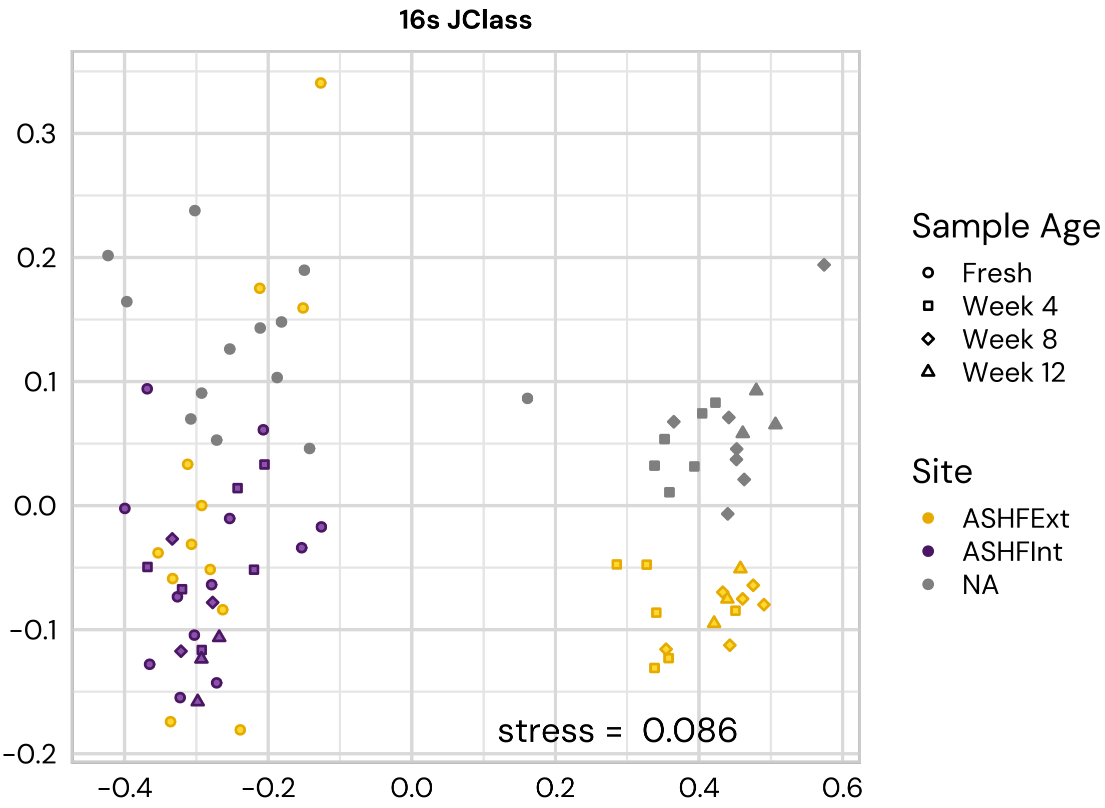

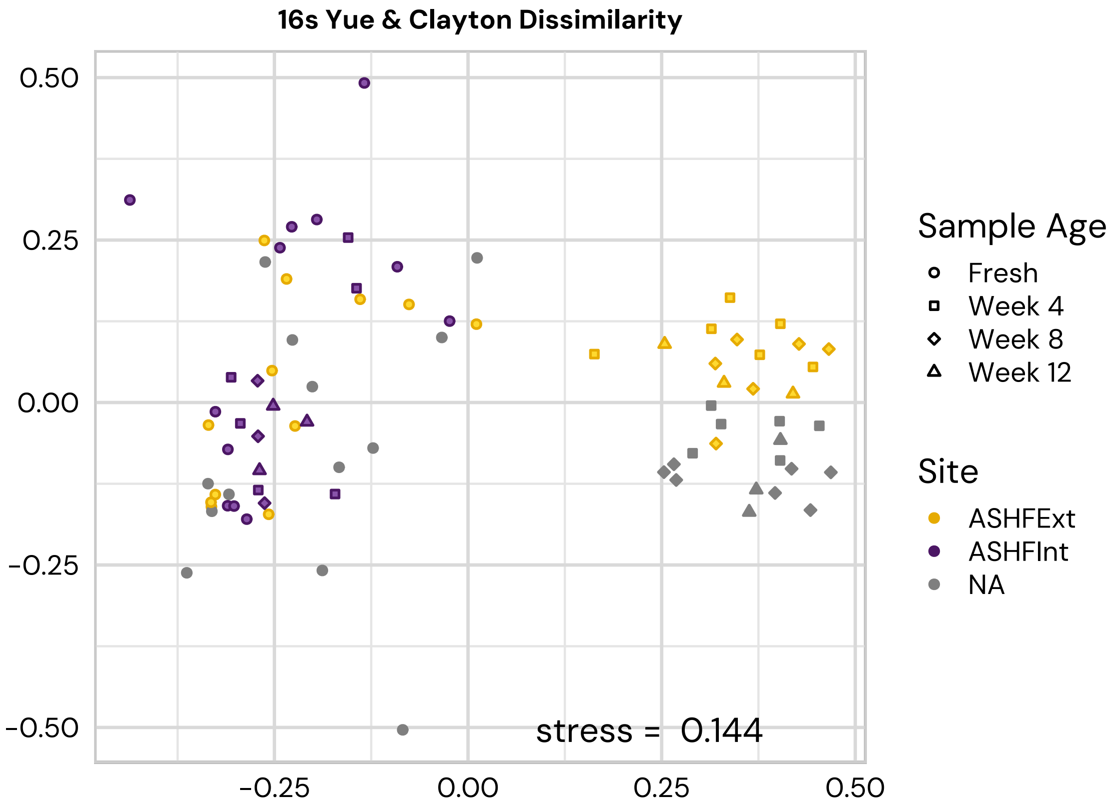

### FigureS2A

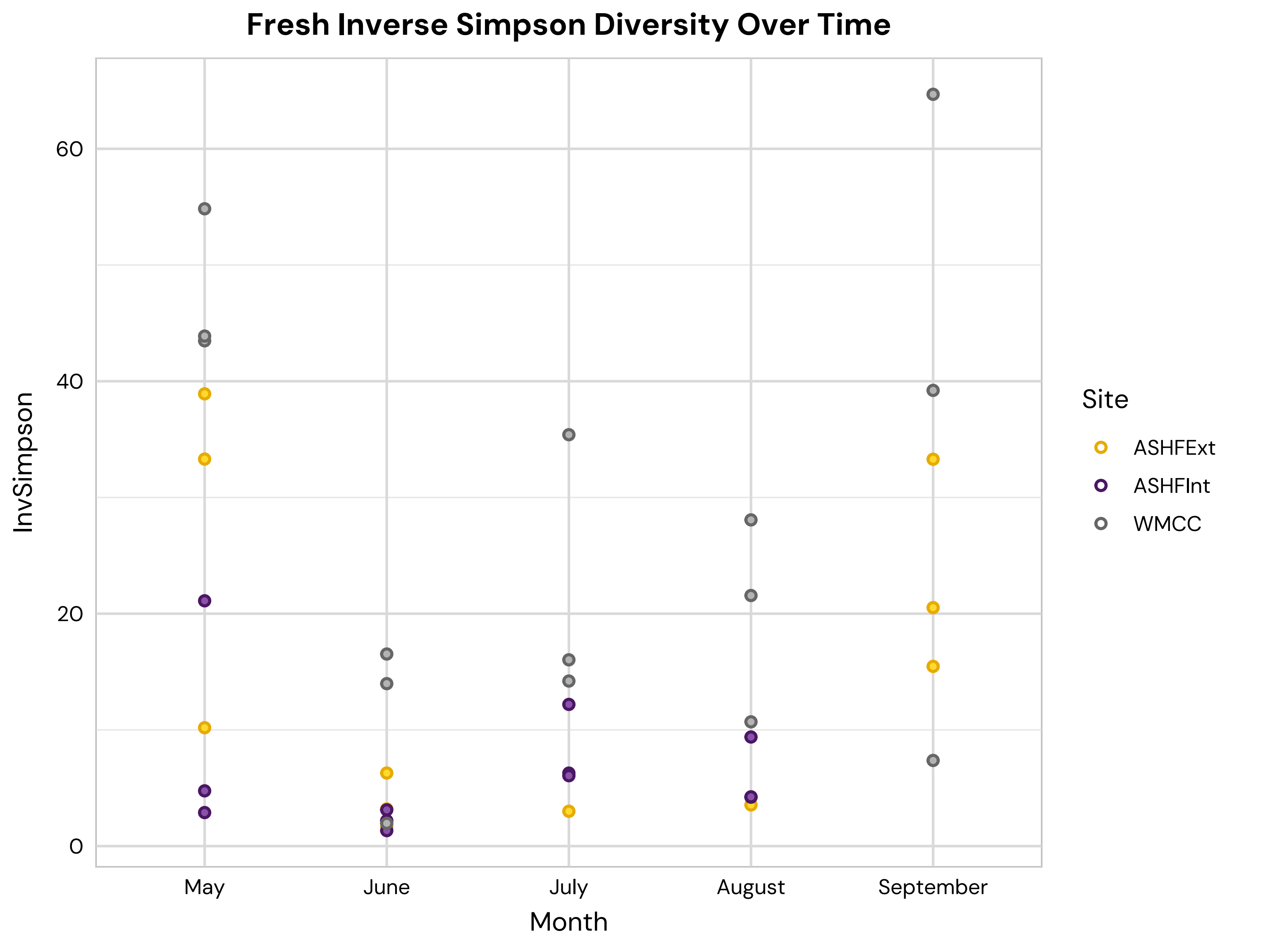

### FigureS2B

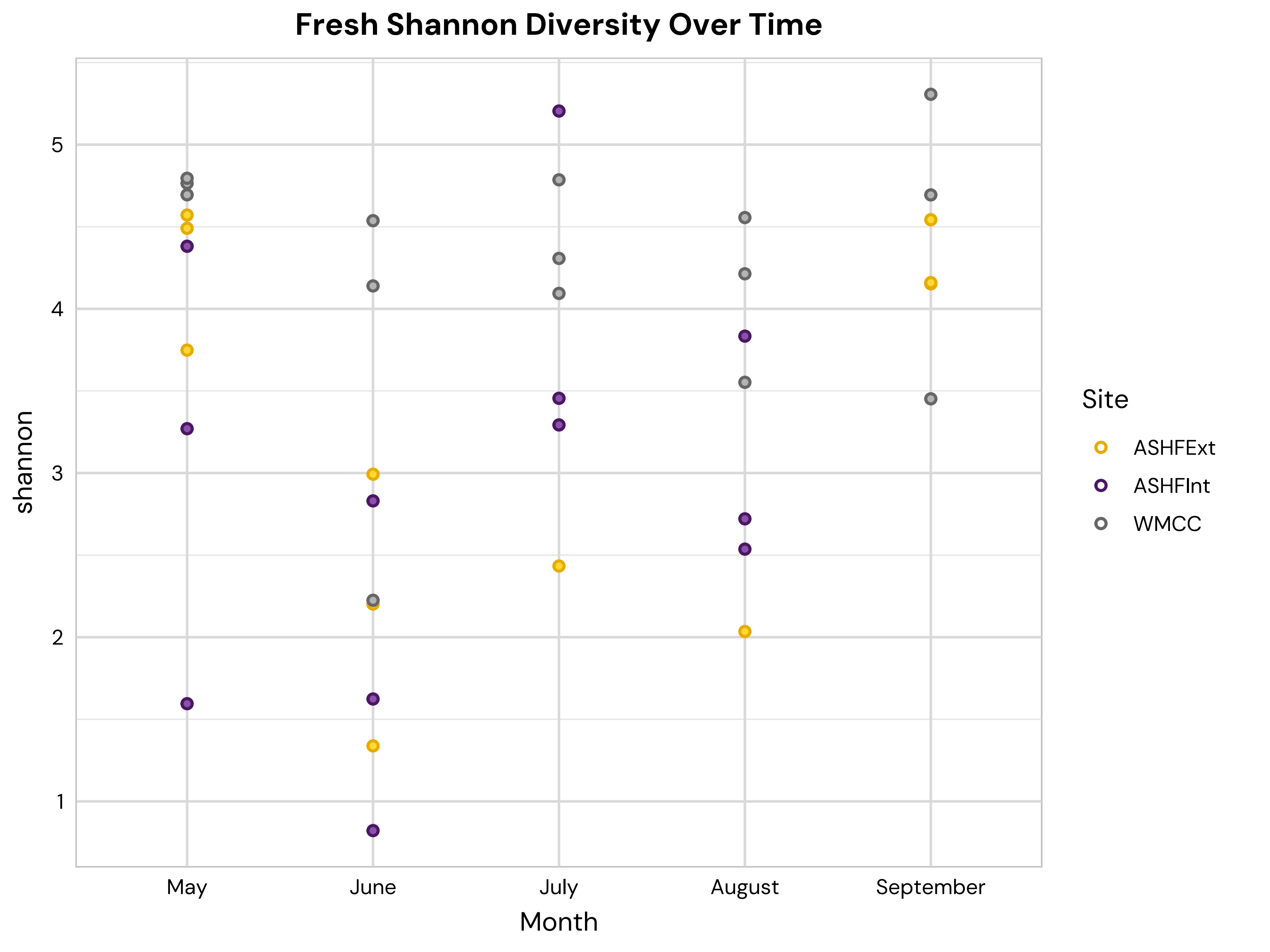

### FigureS2C

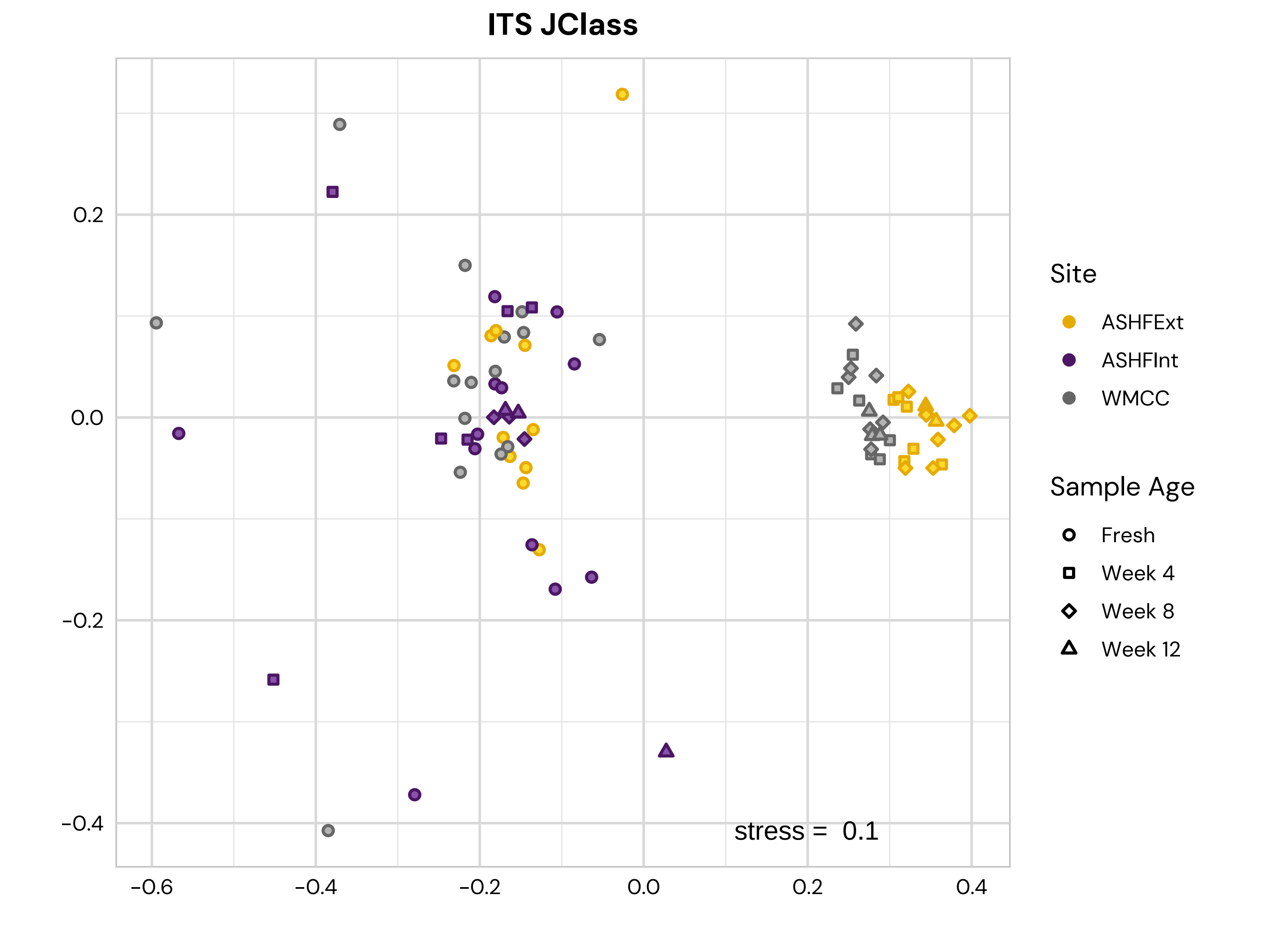

### FigureS2D

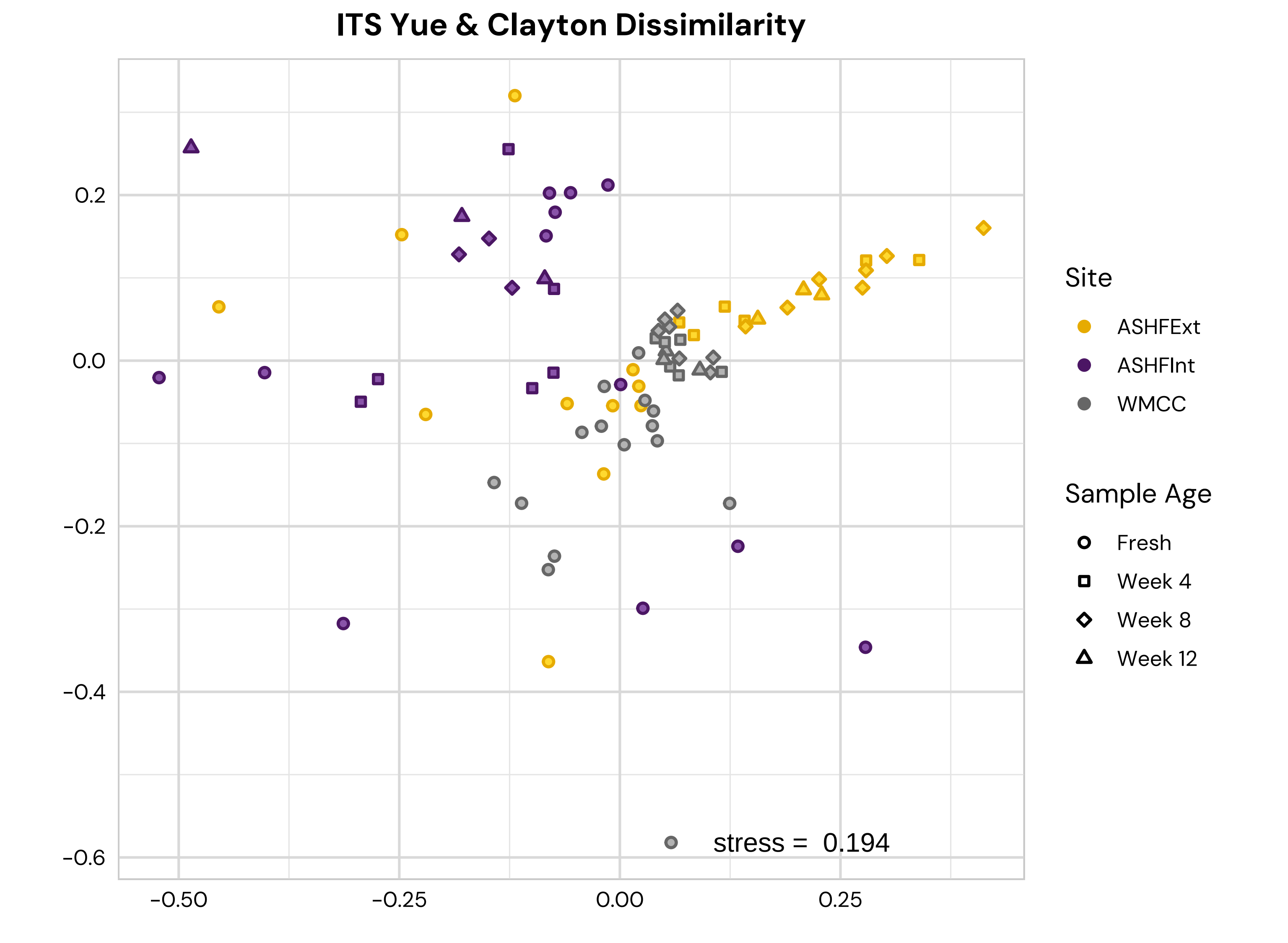

### FigureS3

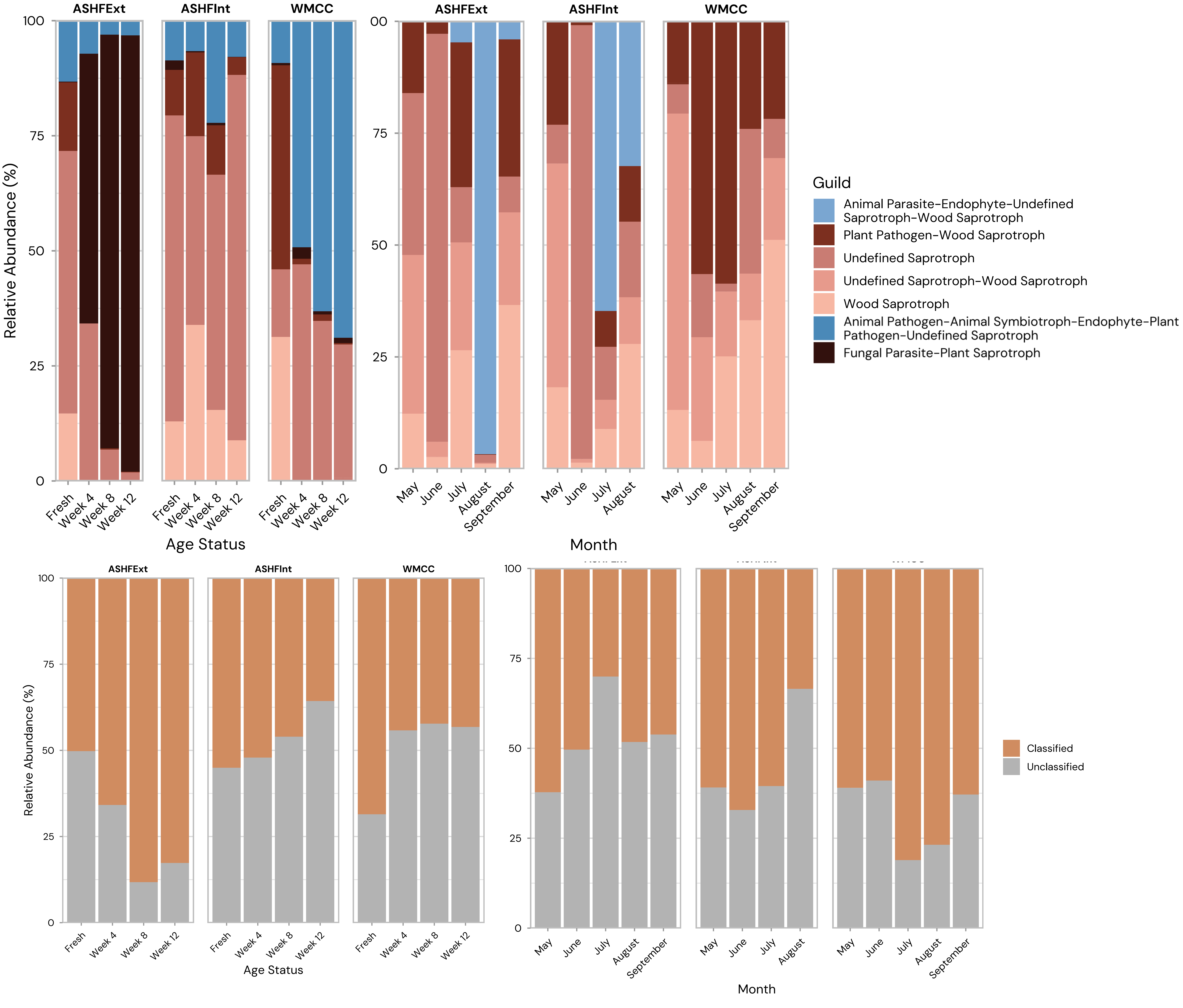

### FigureS4A

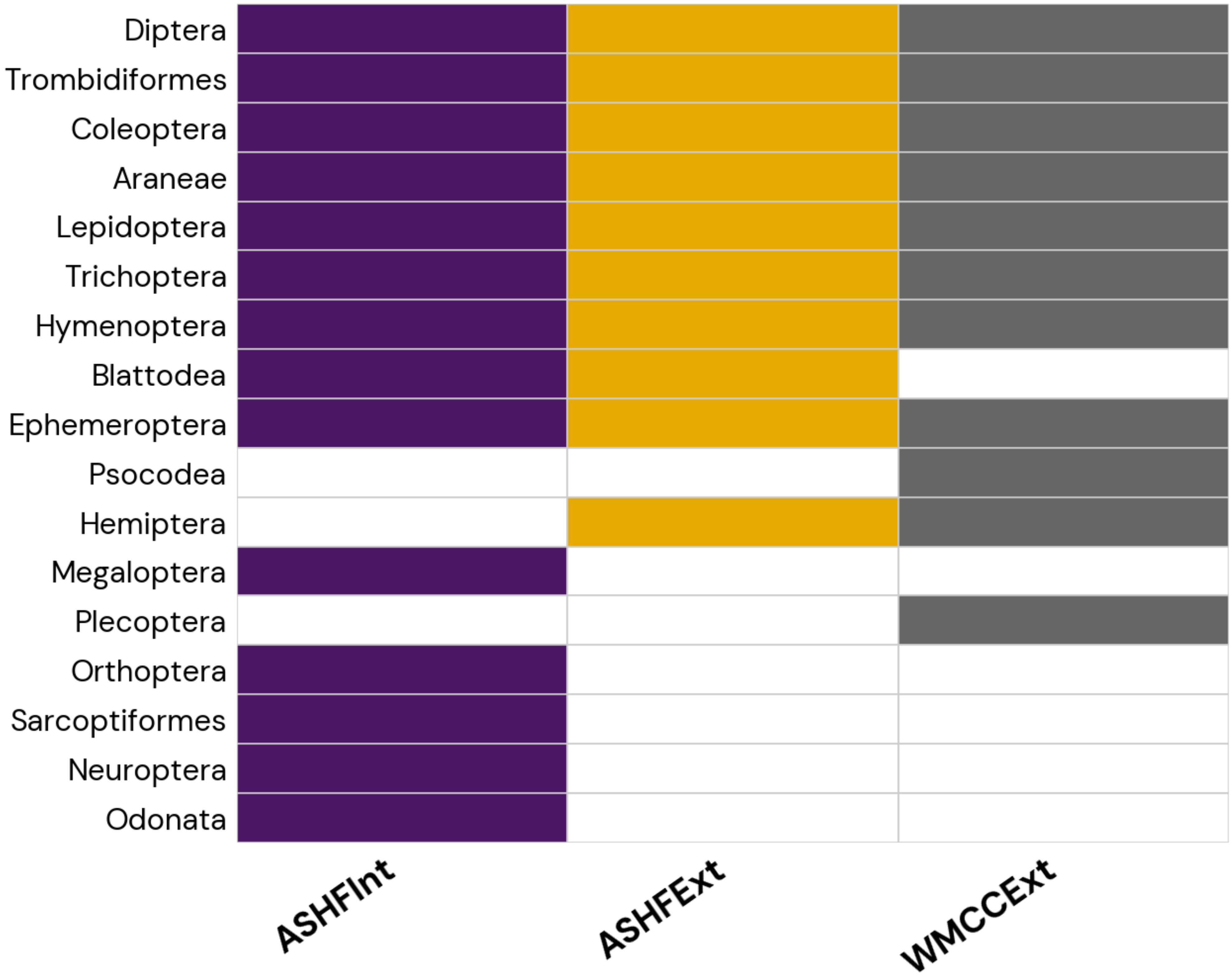

### FigureS4B

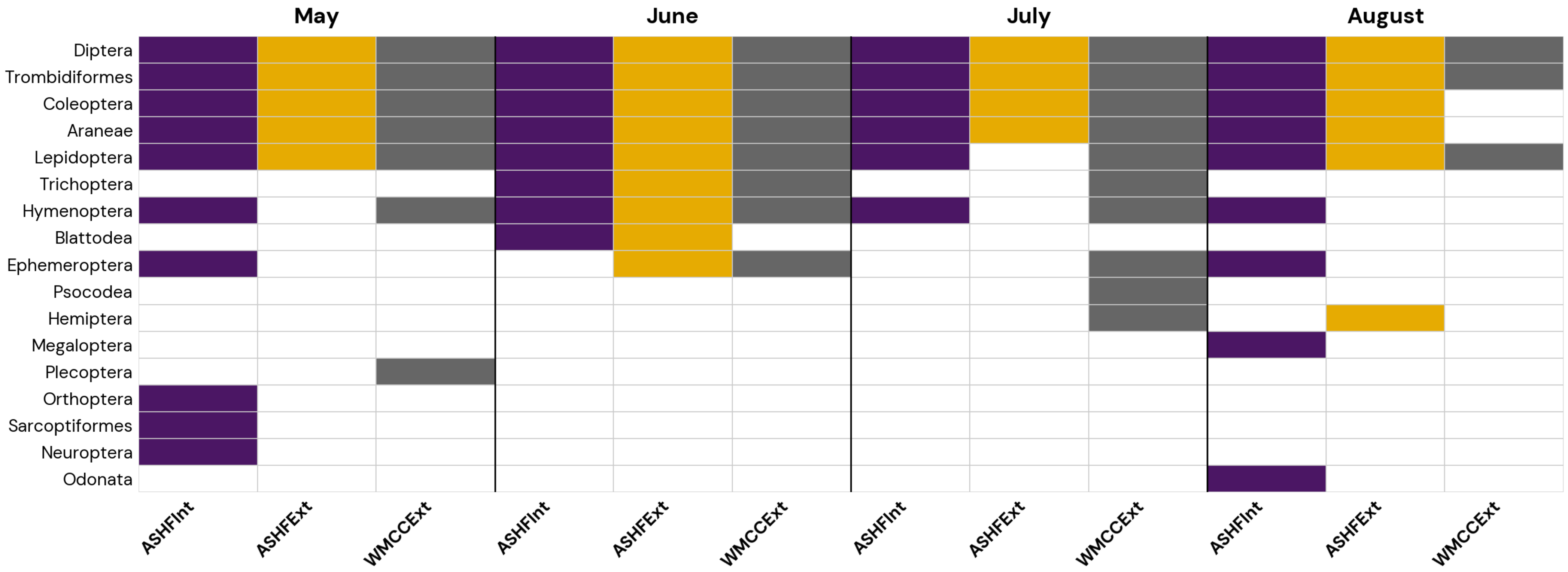

### FigureS4C

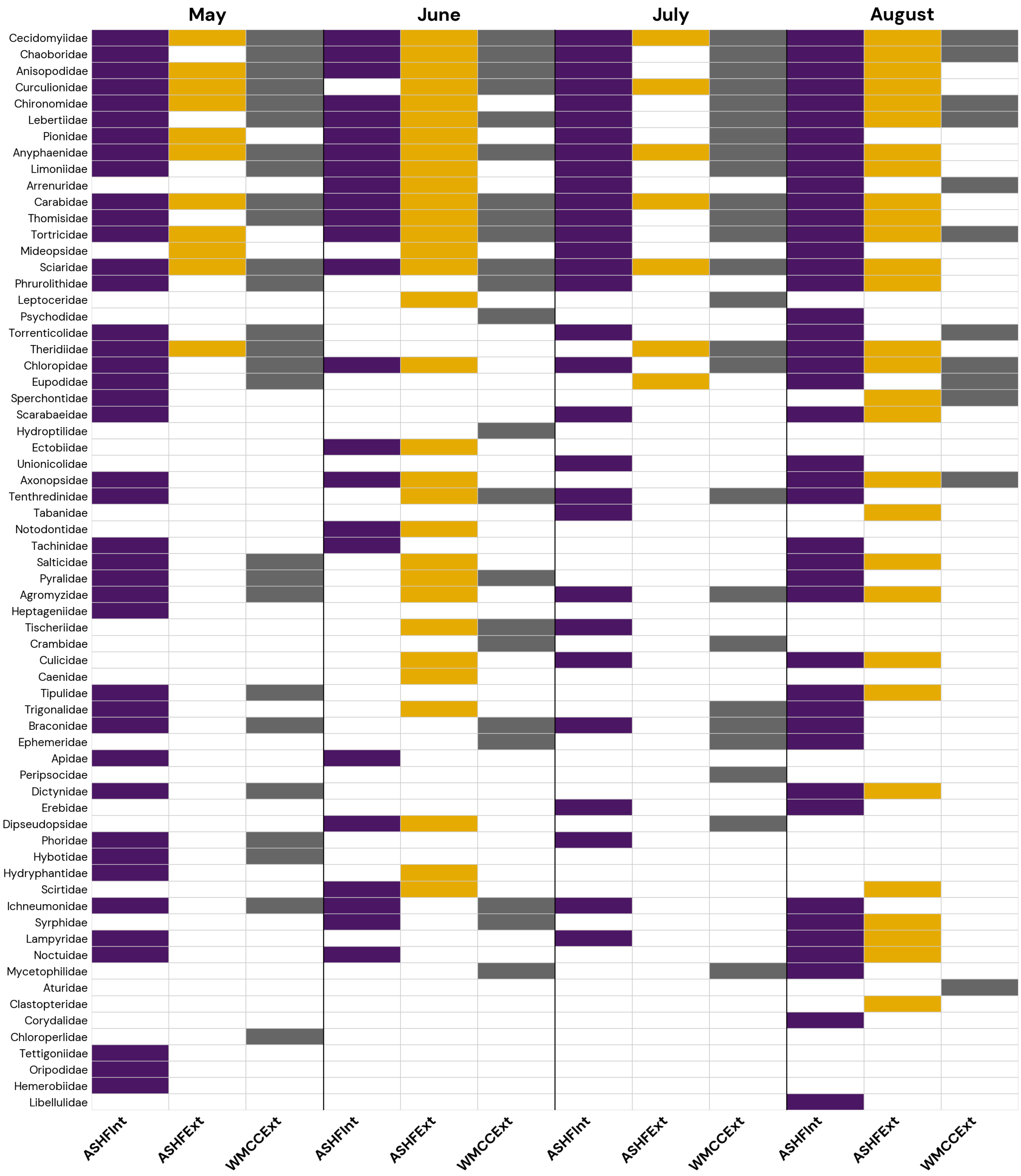

### FigureS5A

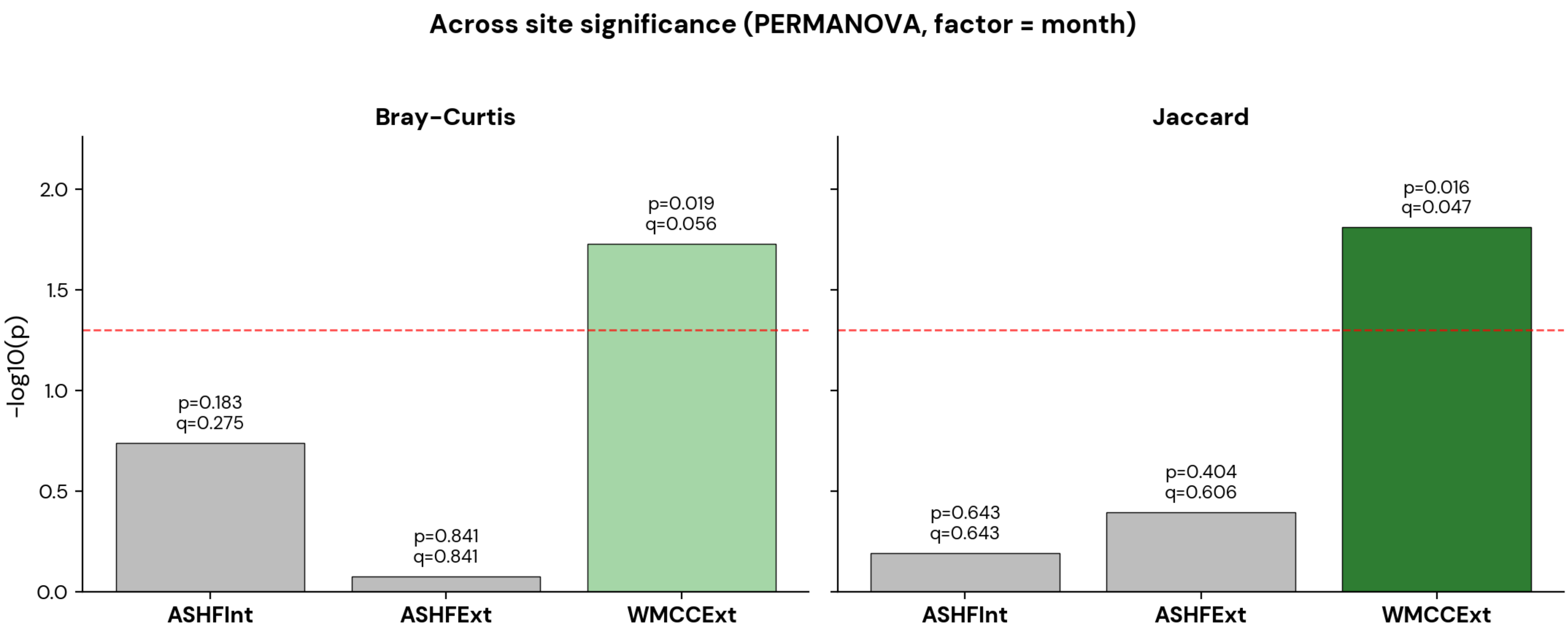

### FigureS5B

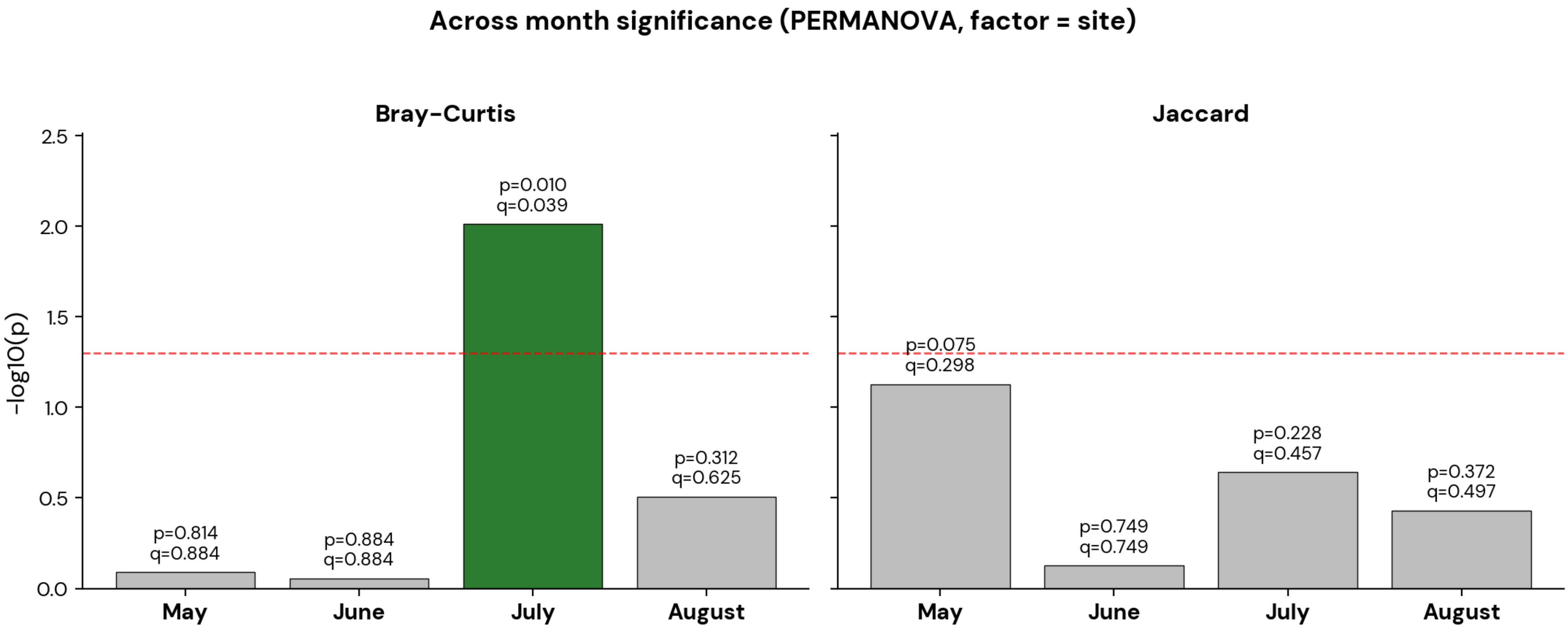

### FigureS5C

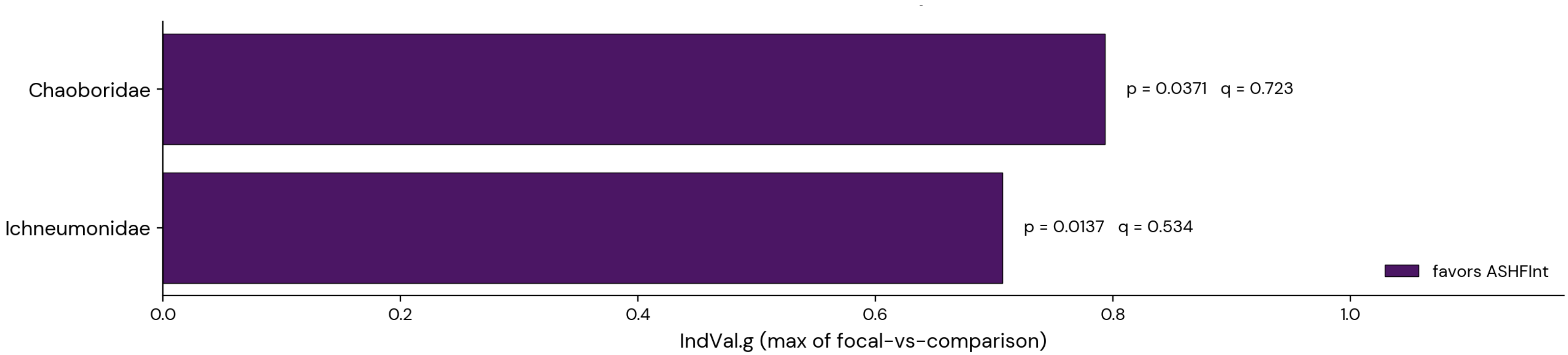

### FigureS5D

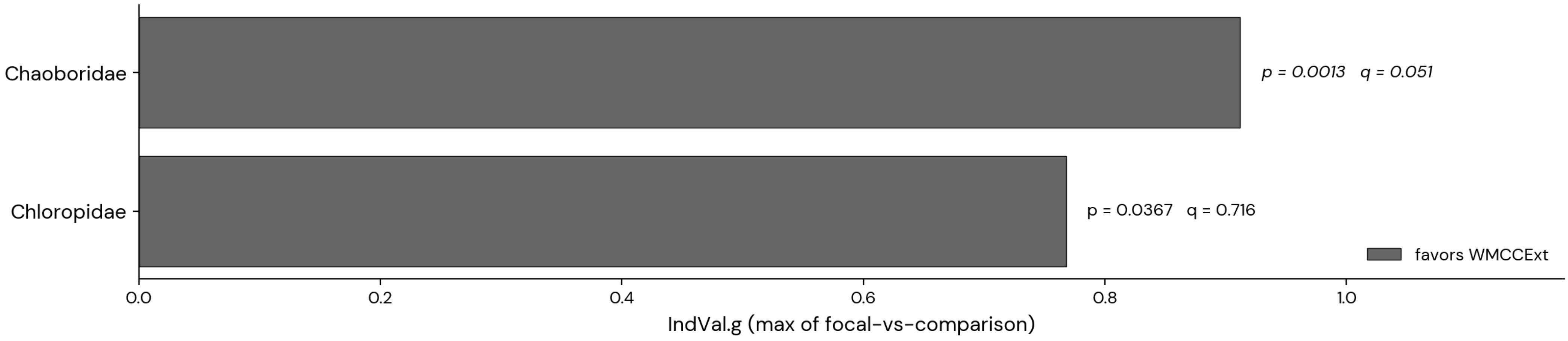

### FigureS5E

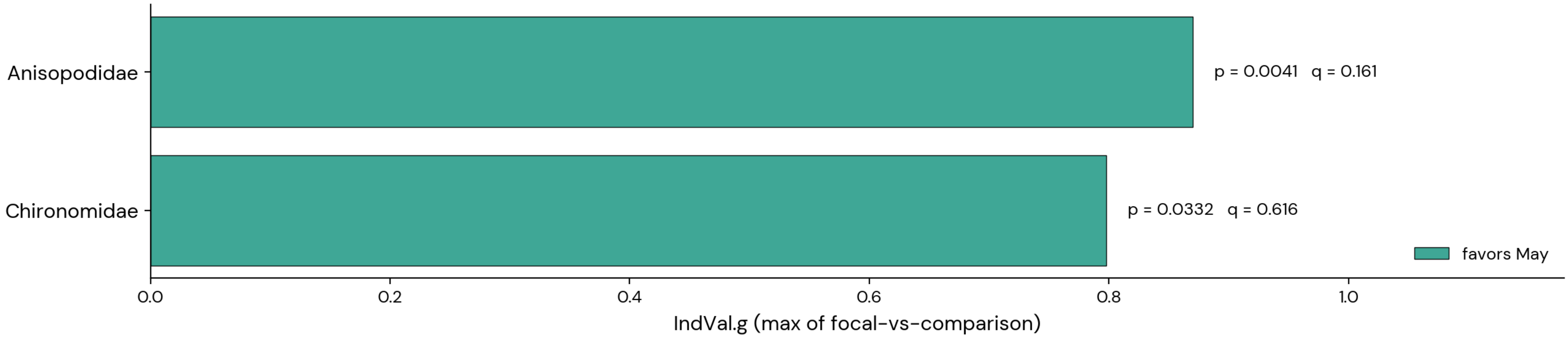

### FigureS5F

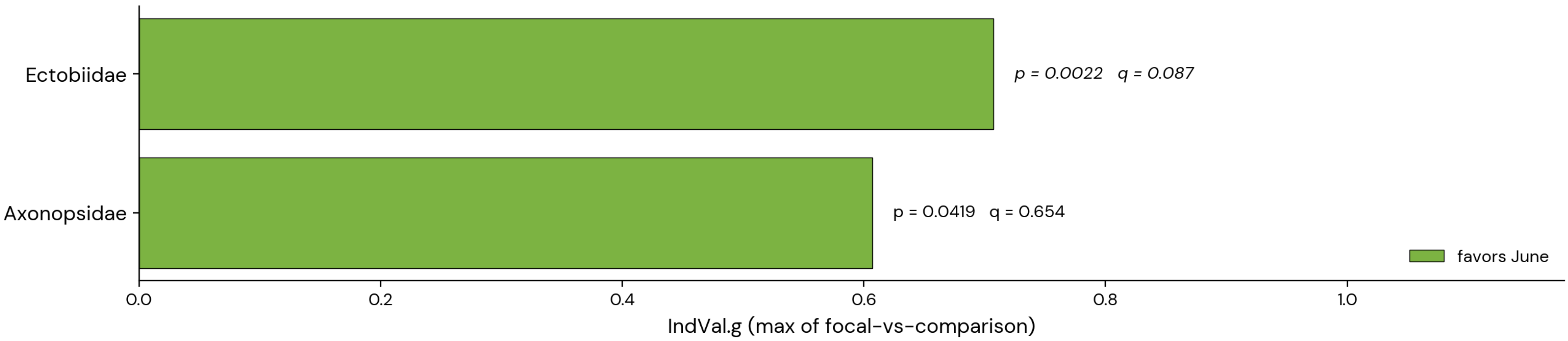

### FigureS5G

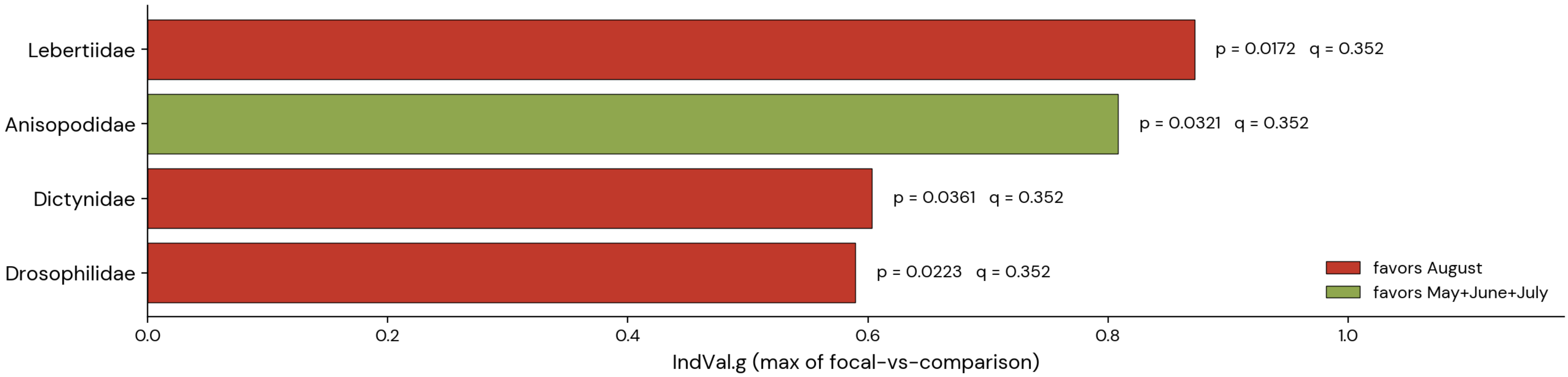

### FigureS6A

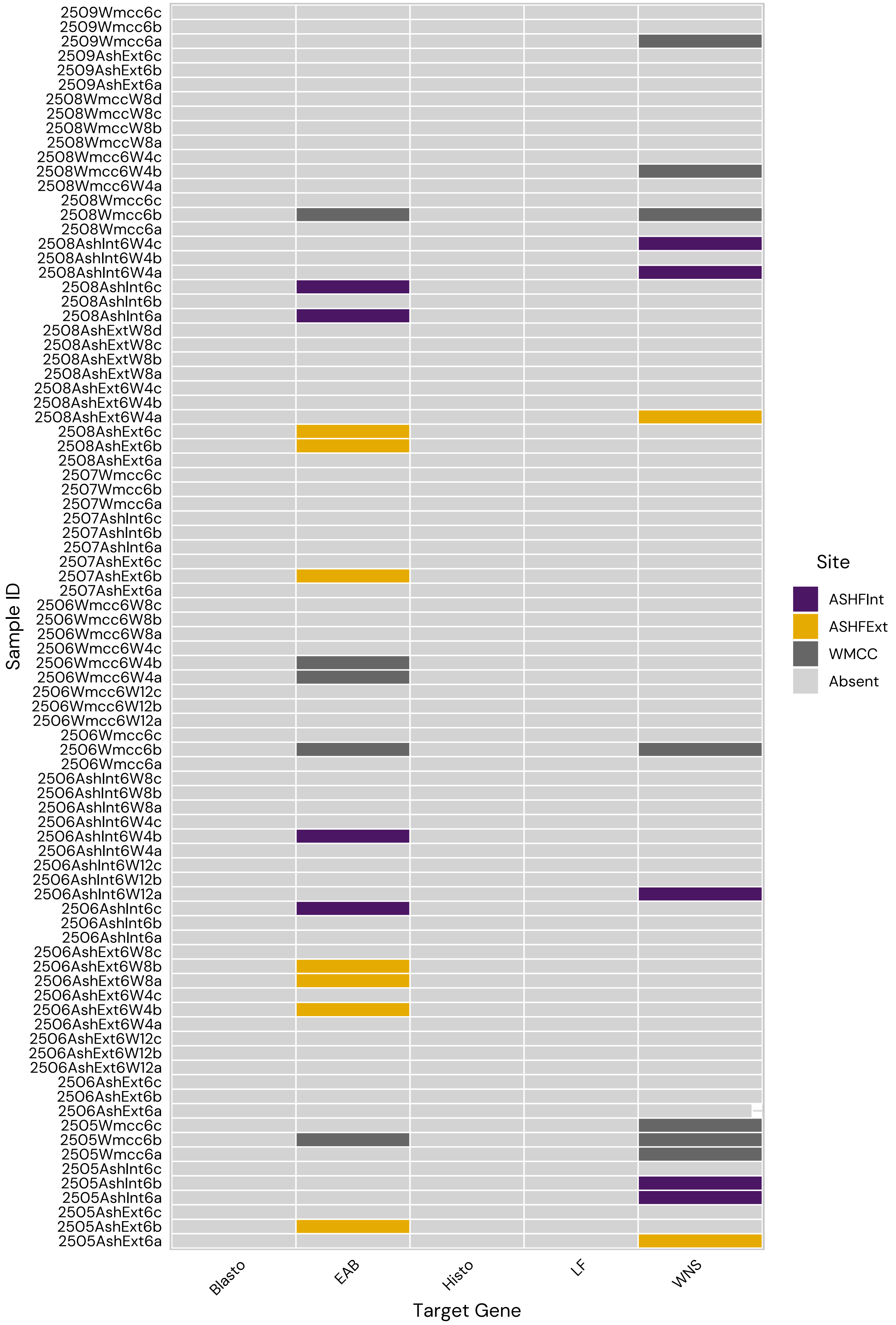

### FigureS6B

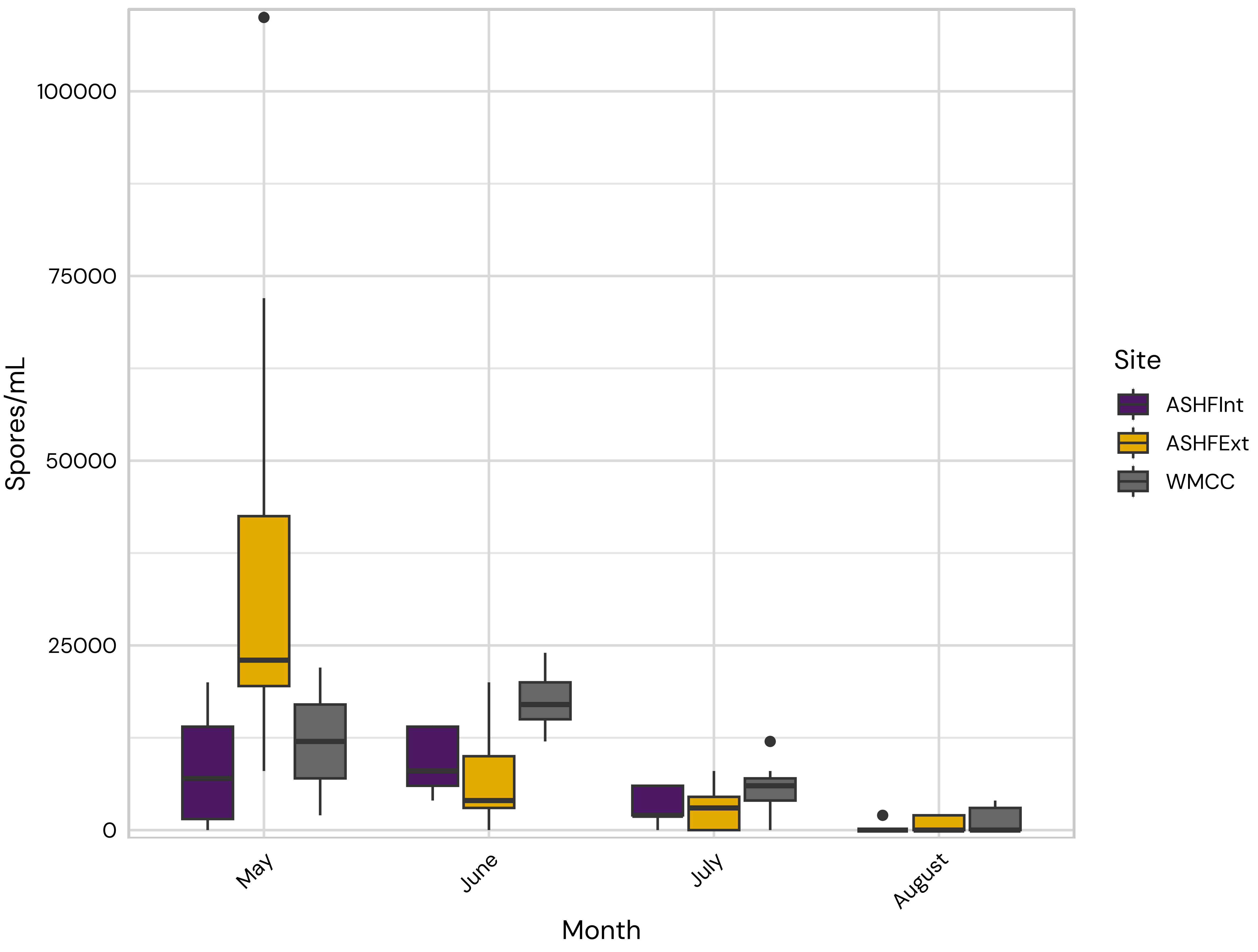
