## Supplementary material for "Microbial, dietary insect, and pathogen communities in fresh and decomposing guano of anthropic little brown bat (Myotis lucifugus) maternity colonies": Table S1

| Barcode | 1uL ng/uL | Total in ug | AVG length bp | pmol | fmol | fmol/ul | to pool (ul) | Added if different from pool volume |
|---|---|---|---|---|---|---|---|---|
| **A04** | | | | | | | | |
| barcode01 | 23.4 | 0.117 | 1,658.00 | 0.107 | 106.9 | 21.4 | 1.4 | |
| barcode02 | 12.88 | 0.06 | 849 | 0.115 | 114.9 | 23.0 | 1.3 | |
| barcode03 | 10.32 | 0.05 | 1,883 | 0.042 | 41.5 | 8.3 | 3.6 | |
| barcode04 | 7.24 | 0.04 | 1,974 | 0.028 | 27.8 | 5.6 | 5.4 | |
| barcode05 | 9 | 0.05 | 1,491.00 | 0.046 | 45.7 | 9.1 | 3.3 | |
| barcode06 | 9.88 | 0.05 | 2,074.00 | 0.036 | 36.1 | 7.2 | 4.2 | |
| barcode07 | 8.2 | 0.04 | 2,074.00 | 0.030 | 30.0 | 6.0 | 5.0 | |
| barcode08 | 8.36 | 0.04 | 2,361.00 | 0.027 | 26.8 | 5.4 | 5.6 | |
| barcode09 | 6.28 | 0.03 | 631 | 0.075 | 75.4 | 15.1 | 2.0 | |
| barcode10 | 6.2 | 0.03 | 535 | 0.088 | 87.8 | 17.6 | 1.7 | |
| barcode11 | 5 | 0.03 | 615 | 0.062 | 61.6 | 12.3 | 2.4 | |
| barcode12 | 3.38 | 0.02 | 2,057 | 0.012 | 12.4 | 2.5 | 12.0 | 5uL |
| barcode13 | 4.76 | 0.02 | 442 | 0.082 | 81.6 | 16.3 | 1.8 | |
| barcode14 | 4.76 | 0.02 | 668 | 0.054 | 54.0 | 10.8 | 2.8 | |
| barcode15 | 3.46 | 0.02 | 691 | 0.038 | 37.9 | 7.6 | 4.0 | |
| barcode16 | 4.16 | 0.02 | 607 | 0.052 | 51.9 | 10.4 | 2.9 | |
| barcode17 | 5 | 0.03 | 448 | 0.085 | 84.6 | 16.9 | 1.8 | |
| barcode18 | 2.32 | 0.01 | 439 | 0.040 | 40.0 | 8.0 | 3.7 | |
| barcode19 | 3.68 | 0.02 | 858 | 0.032 | 32.5 | 6.5 | 4.6 | |
| barcode20 | 4.24 | 0.02 | 482 | 0.067 | 66.6 | 13.3 | 2.3 | |
| barcode21 | 3.9 | 0.02 | 732 | 0.040 | 40.4 | 8.1 | 3.7 | |
| barcode22 | 3.5 | 0.02 | 631 | 0.042 | 42.0 | 8.4 | 3.6 | |
| barcode23 | 2.76 | 0.01 | 723 | 0.029 | 28.9 | 5.8 | 5.2 | |
| barcode24 | 3.12 | 0.02 | 1504 | 0.016 | 15.7 | 3.1 | 9.5 | |
| **A01** | | | | | | | | |
| barcode01 | 1.848 | 0.01 | 1,538.00 | 0.009 | 9.1 | 1.8 | 16.5 | Added all remaining volume |
| barcode02 | 10.72 | 0.05 | 2,450.00 | 0.033 | 33.1 | 6.6 | 4.5 | |
| barcode03 | 5.36 | 0.03 | 632 | 0.064 | 64.3 | 12.9 | 2.3 | |
| barcode04 | 4.36 | 0.02 | 685 | 0.048 | 48.2 | 9.6 | 3.1 | |
| barcode05 | 3.16 | 0.02 | 1,162.00 | 0.021 | 20.6 | 4.1 | 7.3 | Added all remaining volume |
| barcode06 | 2.9 | 0.01 | 1,362.00 | 0.016 | 16.1 | 3.2 | 9.3 | Added all remaining volume |
| barcode07 | 4.04 | 0.02 | 590 | 0.052 | 51.9 | 10.4 | 2.9 | |
| barcode08 | 40 | 0.20 | 3,098.00 | 0.098 | 97.8 | 19.6 | 1.5 | |
| barcode09 | 46.8 | 0.23 | 3,329.00 | 0.107 | 106.5 | 21.3 | 1.4 | |

| | | | | | | | | |
|---|---|---|---|---|---|---|---|---|
| barcode10 | 28.8 | 0.14 | 2,925.00 | 0.075 | 74.6 | 14.9 | 2.0 | |
| barcode11 | 40.4 | 0.20 | 1,459.00 | 0.210 | 209.8 | 42.0 | 0.7 | 1uL |
| barcode12 | 23.4 | 0.12 | 2,789.00 | 0.064 | 63.6 | 12.7 | 2.4 | |
| barcode13 | 28.4 | 0.14 | 2,816.00 | 0.076 | 76.4 | 15.3 | 2.0 | |
| barcode14 | 33 | 0.17 | 2,819.00 | 0.089 | 88.7 | 17.7 | 1.7 | |
| barcode15 | 26.8 | 0.13 | 2,797.00 | 0.073 | 72.6 | 14.5 | 2.1 | |
| barcode16 | 30.8 | 0.15 | 3,063.00 | 0.076 | 76.2 | 15.2 | 2.0 | |
| barcode17 | 16.52 | 0.08 | 2,799.00 | 0.045 | 44.7 | 8.9 | 3.4 | |
| barcode18 | 10.24 | 0.05 | 3,066.00 | 0.025 | 25.3 | 5.1 | 5.9 | |
| barcode19 | 11.2 | 0.06 | 3,248.00 | 0.026 | 26.1 | 5.2 | 5.7 | |
| barcode20 | 13.92 | 0.07 | 2,919.00 | 0.036 | 36.1 | 7.2 | 4.2 | |
| barcode21 | 9.52 | 0.05 | 2,612.00 | 0.028 | 27.6 | 5.5 | 5.4 | |
| barcode22 | 18.16 | 0.09 | 2,496.00 | 0.055 | 55.1 | 11.0 | 2.7 | |
| barcode23 | 7.64 | 0.04 | 1,798.00 | 0.032 | 32.2 | 6.4 | 4.7 | |
| barcode24 | 11.28 | 0.06 | 2,380.00 | 0.036 | 35.9 | 7.2 | 4.2 | |
