## Supplementary material for "Microbial, dietary insect, and pathogen communities in fresh and decomposing guano of anthropic little brown bat (Myotis lucifugus) maternity colonies": Table S2

| total otus | 8381 | | | | | |
|---|---|---|---|---|---|---|
| **Summary.seqs** | | | | | | |
| | Start | End | NBases | Ambigs | Polymer | Numseqs |
| Minimum | 1 | 246 | 246 | 0 | 3 | 1 |
| 2.5%-tile | 1 | 252 | 252 | 0 | 4 | 80736 |
| 25%-tile | 1 | 253 | 253 | 0 | 4 | 807352 |
| Median | 1 | 253 | 253 | 0 | 4 | 1614703 |
| 75%-tile | 1 | 253 | 253 | 0 | 5 | 2422054 |
| 97.5%-tile | 1 | 253 | 253 | 4 | 6 | 3148670 |
| Maximum | 1 | 502 | 502 | 219 | 232 | 3229405 |
| Mean | 1 | 253 | 253 | 0 | 4 | |
| # of unique seqs | 3229405 | | | | | |
| total # of seqs | 3229405 | | | | | |
| **screen.seqs** | | | | | | |
| | Start | End | NBases | Ambigs | Polymer | Numseqs |
| Minimum | 1 | 250 | 250 | 0 | 3 | 1 |
| 2.5%-tile | 1 | 252 | 252 | 0 | 4 | 69704 |
| 25%-tile | 1 | 253 | 253 | 0 | 4 | 697031 |
| Median | 1 | 253 | 253 | 0 | 4 | 1394062 |
| 75%-tile | 1 | 253 | 253 | 0 | 5 | 2091093 |
| 97.5%-tile | 1 | 253 | 253 | 0 | 6 | 2718420 |
| Maximum | 1 | 275 | 275 | 0 | 13 | 2788123 |
| Mean | 1 | 253 | 253 | 0 | 4 | |
| # of unique seqs | 2788123 | | | | | |
| total # of seqs | 2788123 | | | | | |
| **unique.seqs** | | | | | | |
| | Start | End | NBases | Ambigs | Polymer | Numseqs |
| Minimum | 1 | 250 | 250 | 0 | 3 | 1 |
| 2.5%-tile | 1 | 252 | 252 | 0 | 4 | 69704 |
| 25%-tile | 1 | 253 | 253 | 0 | 4 | 697031 |

| | | | | | | |
|---|---|---|---|---|---|---|
| Median | 1 | 253 | 253 | 0 | 4 | 1394062 |
| 75%-tile | 1 | 253 | 253 | 0 | 5 | 2091093 |
| 97.5%-tile | 1 | 253 | 253 | 0 | 6 | 2718420 |
| Maximum | 1 | 275 | 275 | 0 | 13 | 2788123 |
| Mean | 1 | 253 | 253 | 0 | 4 | |
| # of unique seqs | 374918 | | | | | |
| total # of seqs | 2788123 | | | | | |
| **align.seqs** | | | | | | |
| | Start | End | NBases | Ambigs | Polymer | Numseqs |
| Minimum | 1 | 10 | 1 | 0 | 1 | 1 |
| 2.5%-tile | 2 | 9584 | 252 | 0 | 4 | 69704 |
| 25%-tile | 2 | 9584 | 253 | 0 | 4 | 697031 |
| Median | 2 | 9584 | 253 | 0 | 4 | 1394062 |
| 75%-tile | 2 | 9584 | 253 | 0 | 5 | 2091093 |
| 97.5%-tile | 2 | 9584 | 253 | 0 | 6 | 2718420 |
| Maximum | 11433 | 11433 | 275 | 0 | 13 | 2788123 |
| Mean | 5 | 9585 | 252 | 0 | 4 | |
| # of unique seqs | 374918 | | | | | |
| total # of seqs | 2788123 | | | | | |
| **filter.seqs** | | | | | | |
| | Start | End | NBases | Ambigs | Polymer | Numseqs |
| Minimum | 1 | 861 | 224 | 0 | 3 | 1 |
| 2.5%-tile | 2 | 972 | 252 | 0 | 4 | 69667 |
| 25%-tile | 2 | 972 | 253 | 0 | 4 | 696662 |
| Median | 2 | 972 | 253 | 0 | 4 | 1393324 |
| 75%-tile | 2 | 972 | 253 | 0 | 5 | 2089986 |
| 97.5%-tile | 2 | 972 | 253 | 0 | 6 | 2716981 |
| Maximum | 2 | 1002 | 275 | 0 | 8 | 2786647 |
| Mean | 1 | 972 | 252 | 0 | 4 | |

| | | | | | | |
|---|---|---|---|---|---|---|
| # of unique seqs | 374120 | | | | | |
| total # of seqs | 2786647 | | | | | |
| **pre.cluster** | | | | | | |
| | Start | End | NBases | Ambigs | Polymer | Numseqs |
| Minimum | 1 | 861 | 224 | 0 | 3 | 1 |
| 2.5%-tile | 2 | 972 | 252 | 0 | 4 | 69667 |
| 25%-tile | 2 | 972 | 253 | 0 | 4 | 696662 |
| Median | 2 | 972 | 253 | 0 | 4 | 1393324 |
| 75%-tile | 2 | 972 | 253 | 0 | 5 | 2089986 |
| 97.5%-tile | 2 | 972 | 253 | 0 | 6 | 2716981 |
| Maximum | 2 | 1002 | 275 | 0 | 8 | 2786647 |
| Mean | 1 | 972 | 252 | 0 | 4 | |
| # of unique seqs | 144248 | | | | | |
| total # of seqs | 2786647 | | | | | |
| **chimera.vsearch** | | | | | | |
| | Start | End | NBases | Ambigs | Polymer | Numseqs |
| Minimum | 1 | 861 | 224 | 0 | 3 | 1 |
| 2.5%-tile | 2 | 972 | 252 | 0 | 4 | 63177 |
| 25%-tile | 2 | 972 | 253 | 0 | 4 | 631763 |
| Median | 2 | 972 | 253 | 0 | 4 | 1263526 |
| 75%-tile | 2 | 972 | 253 | 0 | 5 | 1895289 |
| 97.5%-tile | 2 | 972 | 253 | 0 | 6 | 2463875 |
| Maximum | 2 | 1002 | 275 | 0 | 8 | 2527051 |
| Mean | 1 | 972 | 253 | 0 | 4 | |
| # of unique seqs | 72777 | | | | | |
| total # of seqs | 2527051 | | | | | |
| **cluster** | | | | | | |
| | Start | End | NBases | Ambigs | Polymer | Numseqs |

| | | | | | | |
|---|---|---|---|---|---|---|
| Minimum | 1 | 861 | 243 | 0 | 3 | 1 |
| 2.5%-tile | 2 | 972 | 252 | 0 | 4 | 1779 |
| 25%-tile | 2 | 972 | 253 | 0 | 4 | 17785 |
| Median | 2 | 972 | 253 | 0 | 4 | 35569 |
| 75%-tile | 2 | 972 | 253 | 0 | 5 | 53353 |
| 97.5%-tile | 2 | 972 | 254 | 0 | 6 | 69359 |
| Maximum | 2 | 1002 | 271 | 0 | 8 | 71137 |
| Mean | 1 | 971 | 252 | 0 | 4 | |
| # of unique seqs | 971 | | | | | |
| total # of seqs | 71137 | | | | | |
