## Supplementary material for "Microbial, dietary insect, and pathogen communities in fresh and decomposing guano of anthropic little brown bat (Myotis lucifugus) maternity colonies": Table S3

| | | | | | | |
|---|---|---|---|---|---|---|
| Total OTUs | **63273** | | | | | |
| **summary.seqs** | | | | | | |
| | Start | End | Nbases | Ambigs | Polymer | Numseqs |
| Min | 1 | 244 | 244 | 0 | 3 | 1 |
| 2.5%-tile: | 1 | 252 | 252 | 0 | 4 | 77571 |
| 25%-tile: | 1 | 259 | 259 | 0 | 5 | 775702 |
| Median: | 1 | 264 | 264 | 0 | 6 | 1551403 |
| 75%-tile | 1 | 313 | 313 | 1 | 6 | 2327104 |
| 97.5%-tile | 1 | 477 | 477 | 14 | 14 | 3025234 |
| Max | 1 | 502 | 502 | 204 | 251 | 3102804 |
| Mean | 1 | 290 | 290 | 1 | 5 | |
| # Unique seqs | 3102804 | | | | | |
| total # seqs | 3102804 | | | | | |
| **screen.seqs** | | | | | | |
| | Start | End | Nbases | Ambigs | Polymer | Numseqs |
| Min | 1 | 249 | 249 | 0 | 3 | 1 |
| 2.5%-tile: | 1 | 252 | 252 | 0 | 4 | 66247 |
| 25%-tile: | 1 | 259 | 259 | 0 | 5 | 662461 |
| Median: | 1 | 262 | 262 | 0 | 6 | 1324921 |
| 75%-tile | 1 | 311 | 311 | 0 | 6 | 1987381 |
| 97.5%-tile | 1 | 370 | 370 | 2 | 9 | 2583595 |
| Max | 1 | 400 | 400 | 2 | 58 | 2649841 |
| Mean | 1 | 282 | 282 | 0 | 5 | |
| # Unique seqs | 2649841 | | | | | |
| total # seqs | 2649841 | | | | | |
| **unique.seqs** | | | | | | |
| | Start | End | Nbases | Ambigs | Polymer | Numseqs |
| Min | 1 | 249 | 249 | 0 | 3 | 1 |
| 2.5%-tile: | 1 | 252 | 252 | 0 | 4 | 66247 |
| 25%-tile: | 1 | 259 | 259 | 0 | 5 | 662461 |

| | | | | | | |
|---|---|---|---|---|---|---|
| Median: | 1 | 262 | 262 | 0 | 6 | 1324921 |
| 75%-tile | 1 | 311 | 311 | 0 | 6 | 1987381 |
| 97.5%-tile | 1 | 370 | 370 | 2 | 9 | 2583595 |
| Max | 1 | 400 | 400 | 2 | 58 | 2649841 |
| Mean | 1 | 282 | 282 | 0 | 5 | |
| # Unique seqs | 1001132 | | | | | |
| total # seqs | 2649841 | | | | | |
| **pre.cluster** | | | | | | |
| | Start | End | Nbases | Ambigs | Polymer | Numseqs |
| Min | 1 | 249 | 249 | 0 | 3 | 1 |
| 2.5%-tile: | 1 | 252 | 252 | 0 | 4 | 66247 |
| 25%-tile: | 1 | 259 | 259 | 0 | 5 | 662461 |
| Median: | 1 | 262 | 262 | 0 | 6 | 1324921 |
| 75%-tile | 1 | 311 | 311 | 0 | 6 | 1987381 |
| 97.5%-tile | 1 | 370 | 370 | 2 | 8 | 2583595 |
| Max | 1 | 400 | 400 | 2 | 58 | 2649841 |
| Mean | 1 | 282 | 282 | 0 | 5 | |
| # Unique seqs | 664993 | | | | | |
| total # seqs | 2649841 | | | | | |
| **chimera.vsearch** | | | | | | |
| | Start | End | Nbases | Ambigs | Polymer | Numseqs |
| Min | 1 | 249 | 249 | 0 | 3 | 1 |
| 2.5%-tile: | 1 | 252 | 252 | 0 | 4 | 65413 |
| 25%-tile: | 1 | 259 | 259 | 0 | 5 | 654129 |
| Median: | 1 | 262 | 262 | 0 | 6 | 1308257 |
| 75%-tile | 1 | 311 | 311 | 0 | 6 | 1962385 |
| 97.5%-tile | 1 | 370 | 370 | 2 | 9 | 2551100 |
| Max | 1 | 400 | 400 | 2 | 58 | 2616512 |
| Mean | 1 | 282 | 282 | 0 | 5 | |

| | | | | | | |
|---|---|---|---|---|---|---|
| # Unique seqs | 652155 | | | | | |
| total # seqs | 2616512 | | | | | |
| **cluster** | | | | | | |
| | Start | End | Nbases | Ambigs | Polymer | Numseqs |
| Min | 1 | 249 | 249 | 0 | 3 | 1 |
| 2.5%-tile: | 1 | 253 | 253 | 0 | 4 | 16296 |
| 25%-tile: | 1 | 260 | 260 | 0 | 5 | 162953 |
| Median: | 1 | 309 | 309 | 0 | 6 | 325906 |
| 75%-tile | 1 | 316 | 316 | 1 | 6 | 488858 |
| 97.5%-tile | 1 | 369 | 369 | 2 | 12 | 635515 |
| Max | 1 | 400 | 400 | 2 | 58 | 651810 |
| Mean | 1 | 293 | 293 | 0 | 5 | |
| # Unique seqs | | | | | | |
| total # seqs | 651810 | | | | | |
