## Supplementary material for "Microbial, dietary insect, and pathogen communities in fresh and decomposing guano of anthropic little brown bat (Myotis lucifugus) maternity colonies": Table S4

| Target | Primer/Probe | Sequence | Reference |
|---|---|---|---|
| *Histoplasma* | Histo_84R22_F | GCA GAR AAT TCG ACC YCA AGC C | López et al. (2017) |
| | Histo_15F23_R | GTA TCC CAC AGC ATC MYG GAG GT | |
| | Histo_45L23_HEX | /5HEX/CC TTC TTG C/ZEN/A ACT YCC YGC GTC T/3IABkFQ/ | |
| *Blastomyces* | Blasto1_V1741_F | TGT GCT GCT GCA CCT TGA TT | Sidamonidze et al. (2012) |
| | Blasto1_V1742_R | CAA CTT GGA TAA TGC GTG AAT ACA ATA | |
| | Blasto2_V1983_F | CTG TAC CTT GTA CCT TGA TT | |
| | Blasto2_V1984_R | CAA CTT GGA TAA TGC GTG AAT AAC ATA | |
| | Blasto_V1743_FAM | /56-FAM/AT TCA AGA C/ZEN/A ACA GAG CCC AGG CGA AAT /3IABkFQ/ | |
| *Pseudogymnoascus destructans* | WNS_nu-IGS-0169-59Gd_F | TGC CTC TCC GCC ATT AGT G | Muller et al. (2013) |
| | WNS_nu-IGS-0235-39Gd_R | ACC ACC GGC TCG CTA GGT | |
| | WNS_nu-IGS-0182/0204-Gd_TexRed | /5TEX615/CG TTA CAG CTT GCT CGG GCT GCC /3IAbRQSp/ | |
| Spotted lanternfly (*Lycorma delicatula*) | LF_LydeITS1F_F | AGC GTT TGA CAG CTG ACT CTT G | Valentin et al. (2020) |
| | LF_LydeITS1R_R | CGC CGA AGC GCA AAA A | |
| | LF_LydeITS1Tm_HEX | /5HEX/CC GCG GGA C/ZEN/C GGT A/3IABkFQ/ | |
| Emerald ash borer (*Agrilus planipennis*) | EAB_EABFOT_F | TCA AAG AAT GAT GTA TTT AAG TTT CGA TC | Kupper et al. (2025) |
| | EAB_EABROT_R | TAG CAA TTT TTA GAC TTC ATT TAG CTG G | |
| | EAB_EAB-RC-P1_FAM | /56-FAM/TA TGG TAA T/ZEN/T GCT CCC GCA AGA ACA GGT /3IABkFQ/ | |
| CO1 | CO1_F | RKTCAACMAATCATAAAGATATTGG | |
| | CO1_R | TAAACTTCWGGRTGWCCAAAAAWY | |
