## Supplementary material for "Microbial, dietary insect, and pathogen communities in fresh and decomposing guano of anthropic little brown bat (Myotis lucifugus) maternity colonies": Table S5

| Sample | Dups | GC | Seqs | SeqCount | | R1 avg | R2 avg |
|---|---|---|---|---|---|---|---|
| 2505AshExt6aBac.2505AshExt6aBac_S192_L001_R1_001 | 93.53753137 | 51 | 0.017532 | 17532 | | 35102.22826 | 35102.22826 |
| 2505AshExt6aBac.2505AshExt6aBac_S192_L001_R2_001 | 95.75633128 | 52 | 0.017532 | 17532 | | | |
| 2505AshExt6bBac.2505AshExt6bBac_S182_L001_R1_001 | 94.84381063 | 51 | 0.041232 | 41232 | | | |
| 2505AshExt6bBac.2505AshExt6bBac_S182_L001_R2_001 | 95.43558401 | 52 | 0.041232 | 41232 | | | |
| 2505AshExt6cBac.2505AshExt6cBac_S193_L001_R1_001 | 93.38238925 | 51 | 0.020249 | 20249 | | | |
| 2505AshExt6cBac.2505AshExt6cBac_S193_L001_R2_001 | 94.89851351 | 52 | 0.020249 | 20249 | | | |
| 2505AshInt6aBac.2505AshInt6aBac_S189_L001_R1_001 | 93.73354957 | 52 | 0.037613 | 37613 | | | |
| 2505AshInt6aBac.2505AshInt6aBac_S189_L001_R2_001 | 94.90601654 | 52 | 0.037613 | 37613 | | | |
| 2505AshInt6bBac.2505AshInt6bBac_S177_L001_R1_001 | 92.9194209 | 52 | 0.024659 | 24659 | | | |
| 2505AshInt6bBac.2505AshInt6bBac_S177_L001_R2_001 | 93.44255647 | 53 | 0.024659 | 24659 | | | |
| 2505AshInt6cBac.2505AshInt6cBac_S191_L001_R1_001 | 94.40194586 | 51 | 0.053858 | 53858 | | | |
| 2505AshInt6cBac.2505AshInt6cBac_S191_L001_R2_001 | 96.24011289 | 51 | 0.053858 | 53858 | | | |
| 2505Wmcc6aBac.2505Wmcc6aBac_S202_L001_R1_001 | 96.03422518 | 51 | 0.051541 | 51541 | | | |
| 2505Wmcc6aBac.2505Wmcc6aBac_S202_L001_R2_001 | 97.11879863 | 51 | 0.051541 | 51541 | | | |
| 2505Wmcc6bBac.2505Wmcc6bBac_S206_L001_R1_001 | 95.13576545 | 51 | 0.024012 | 24012 | | | |
| 2505Wmcc6bBac.2505Wmcc6bBac_S206_L001_R2_001 | 96.14775945 | 51 | 0.024012 | 24012 | | | |
| 2505Wmcc6cBac.2505Wmcc6cBac_S200_L001_R1_001 | 94.0513834 | 51 | 0.01012 | 10120 | | | |
| 2505Wmcc6cBac.2505Wmcc6cBac_S200_L001_R2_001 | 96.61067194 | 52 | 0.01012 | 10120 | | | |
| 2506AshExt6aBac.2506AshExt6aBac_S203_L001_R1_001 | 92.78828229 | 52 | 0.03755 | 37550 | | | |
| 2506AshExt6aBac.2506AshExt6aBac_S203_L001_R2_001 | 93.88015979 | 53 | 0.03755 | 37550 | | | |
| 2506AshExt6bBac.2506AshExt6bBac_S178_L001_R1_001 | 94.91904376 | 51 | 0.040701 | 40701 | | | |
| 2506AshExt6bBac.2506AshExt6bBac_S178_L001_R2_001 | 95.88216506 | 51 | 0.040701 | 40701 | | | |
| 2506AshExt6cBac.2506AshExt6cBac_S187_L001_R1_001 | 94.53602991 | 52 | 0.05884 | 58840 | | | |
| 2506AshExt6cBac.2506AshExt6cBac_S187_L001_R2_001 | 95.93473827 | 53 | 0.05884 | 58840 | | | |
| 2506AshInt6aBac.2506AshInt6aBac_S204_L001_R1_001 | 95.09280146 | 50 | 0.025269 | 25269 | | | |
| 2506AshInt6aBac.2506AshInt6aBac_S204_L001_R2_001 | 96.37896236 | 50 | 0.025269 | 25269 | | | |
| 2506AshInt6bBac.2506AshInt6bBac_S205_L001_R1_001 | 95.87739375 | 51 | 0.034517 | 34517 | | | |
| 2506AshInt6bBac.2506AshInt6bBac_S205_L001_R2_001 | 96.5466292 | 52 | 0.034517 | 34517 | | | |
| 2506AshInt6cBac.2506AshInt6cBac_S176_L001_R1_001 | 94.49811352 | 51 | 0.024119 | 24119 | | | |
| 2506AshInt6cBac.2506AshInt6cBac_S176_L001_R2_001 | 95.29416642 | 52 | 0.024119 | 24119 | | | |
| 2506Wmcc6aBac.2506Wmcc6aBac_S185_L001_R1_001 | 92.79314814 | 51 | 0.017747 | 17747 | | | |
| 2506Wmcc6aBac.2506Wmcc6aBac_S185_L001_R2_001 | 94.66388685 | 52 | 0.017747 | 17747 | | | |
| 2506Wmcc6bBac.2506Wmcc6bBac_S183_L001_R1_001 | 95.1172264 | 51 | 0.055235 | 55235 | | | |
| 2506Wmcc6bBac.2506Wmcc6bBac_S183_L001_R2_001 | 96.22159862 | 52 | 0.055235 | 55235 | | | |
| 2506Wmcc6cBac.2506Wmcc6cBac_S184_L001_R1_001 | 95.75542044 | 51 | 0.026151 | 26151 | | | |
| 2506Wmcc6cBac.2506Wmcc6cBac_S184_L001_R2_001 | 97.07085771 | 51 | 0.026151 | 26151 | | | |

| | | | | |
|---|---|---|---|---|
| 2507AshExt6W4aBac.2507AshExt6W4aBac_S181_L001_R1_001 | 93.13226059 | 52 | 0.037101 | 37101 |
| 2507AshExt6W4aBac.2507AshExt6W4aBac_S181_L001_R2_001 | 93.88965257 | 52 | 0.037101 | 37101 |
| 2507AshExt6W4bBac.2507AshExt6W4bBac_S195_L001_R1_001 | 93.98467489 | 52 | 0.067993 | 67993 |
| 2507AshExt6W4bBac.2507AshExt6W4bBac_S195_L001_R2_001 | 95.15097142 | 52 | 0.067993 | 67993 |
| 2507AshExt6W4cBac.2507AshExt6W4cBac_S198_L001_R1_001 | 93.99424023 | 52 | 0.050002 | 50002 |
| 2507AshExt6W4cBac.2507AshExt6W4cBac_S198_L001_R2_001 | 94.85020599 | 52 | 0.050002 | 50002 |
| 2507AshExt6aBac.2507AshExt6aBac_S167_L001_R1_001 | 94.32038486 | 49 | 0.027854 | 27854 |
| 2507AshExt6aBac.2507AshExt6aBac_S167_L001_R2_001 | 95.96826309 | 50 | 0.027854 | 27854 |
| 2507AshExt6bBac.2507AshExt6bBac_S173_L001_R1_001 | 94.52917488 | 49 | 0.023342 | 23342 |
| 2507AshExt6bBac.2507AshExt6bBac_S173_L001_R2_001 | 96.01148145 | 49 | 0.023342 | 23342 |
| 2507AshExt6cBac.2507AshExt6cBac_S174_L001_R1_001 | 78.93700787 | 47 | 0.000508 | 508 |
| 2507AshExt6cBac.2507AshExt6cBac_S174_L001_R2_001 | 83.66141732 | 48 | 0.000508 | 508 |
| 2507AshInt6W4aBac.2507AshInt6W4aBac_S197_L001_R1_001 | 95.55542154 | 52 | 0.049746 | 49746 |
| 2507AshInt6W4aBac.2507AshInt6W4aBac_S197_L001_R2_001 | 96.69521168 | 53 | 0.049746 | 49746 |
| 2507AshInt6W4bBac.2507AshInt6W4bBac_S196_L001_R1_001 | 95.05473542 | 51 | 0.034073 | 34073 |
| 2507AshInt6W4bBac.2507AshInt6W4bBac_S196_L001_R2_001 | 96.55739148 | 52 | 0.034073 | 34073 |
| 2507AshInt6W4cBac.2507AshInt6W4cBac_S157_L001_R1_001 | 94.85083134 | 52 | 0.025441 | 25441 |
| 2507AshInt6W4cBac.2507AshInt6W4cBac_S157_L001_R2_001 | 94.92551393 | 53 | 0.025441 | 25441 |
| 2507AshInt6aBac.2507AshInt6aBac_S199_L001_R1_001 | 94.98956159 | 52 | 0.04311 | 43110 |
| 2507AshInt6aBac.2507AshInt6aBac_S199_L001_R2_001 | 96.43934122 | 53 | 0.04311 | 43110 |
| 2507AshInt6bBac.2507AshInt6bBac_S165_L001_R1_001 | 94.32786468 | 48 | 0.023589 | 23589 |
| 2507AshInt6bBac.2507AshInt6bBac_S165_L001_R2_001 | 95.553012 | 49 | 0.023589 | 23589 |
| 2507AshInt6cBac.2507AshInt6cBac_S175_L001_R1_001 | 91.39280125 | 48 | 0.015336 | 15336 |
| 2507AshInt6cBac.2507AshInt6cBac_S175_L001_R2_001 | 94.18362024 | 49 | 0.015336 | 15336 |
| 2507Wmcc6W4aBac.2507Wmcc6W4aBac_S171_L001_R1_001 | 94.10433709 | 53 | 0.063199 | 63199 |
| 2507Wmcc6W4aBac.2507Wmcc6W4aBac_S171_L001_R2_001 | 94.95719869 | 54 | 0.063199 | 63199 |
| 2507Wmcc6W4bBac.2507Wmcc6W4bBac_S154_L001_R1_001 | 93.98943197 | 54 | 0.062074 | 62074 |
| 2507Wmcc6W4bBac.2507Wmcc6W4bBac_S154_L001_R2_001 | 94.51461159 | 55 | 0.062074 | 62074 |
| 2507Wmcc6W4cBac.2507Wmcc6W4cBac_S172_L001_R1_001 | 93.16860465 | 54 | 0.037152 | 37152 |
| 2507Wmcc6W4cBac.2507Wmcc6W4cBac_S172_L001_R2_001 | 94.12952196 | 54 | 0.037152 | 37152 |
| 2507Wmcc6aBac.2507Wmcc6aBac_S166_L001_R1_001 | 94.34877606 | 52 | 0.027003 | 27003 |
| 2507Wmcc6aBac.2507Wmcc6aBac_S166_L001_R2_001 | 95.32644521 | 53 | 0.027003 | 27003 |
| 2507Wmcc6bBac.2507Wmcc6bBac_S170_L001_R1_001 | 94.30469192 | 50 | 0.03538 | 35380 |
| 2507Wmcc6bBac.2507Wmcc6bBac_S170_L001_R2_001 | 96.19841718 | 51 | 0.03538 | 35380 |
| 2507Wmcc6cBac.2507Wmcc6cBac_S161_L001_R1_001 | 94.94308889 | 48 | 0.029959 | 29959 |
| 2507Wmcc6cBac.2507Wmcc6cBac_S161_L001_R2_001 | 96.08131113 | 48 | 0.029959 | 29959 |
| 2508AshExt6W8aBac.2508AshExt6W8aBac_S159_L001_R1_001 | 93.00461919 | 53 | 0.046545 | 46545 |

| | | | | |
|---|---|---|---|---|
| 2508AshExt6W8aBac.2508AshExt6W8aBac_S159_L001_R2_001 | 94.51713396 | 54 | 0.046545 | 46545 |
| 2508AshExt6W8bBac.2508AshExt6W8bBac_S152_L001_R1_001 | 89.27359131 | 53 | 0.011784 | 11784 |
| 2508AshExt6W8bBac.2508AshExt6W8bBac_S152_L001_R2_001 | 90.94534963 | 54 | 0.011784 | 11784 |
| 2508AshExt6W8cBac.2508AshExt6W8cBac_S169_L001_R1_001 | 92.09765642 | 53 | 0.022487 | 22487 |
| 2508AshExt6W8cBac.2508AshExt6W8cBac_S169_L001_R2_001 | 93.40063148 | 53 | 0.022487 | 22487 |
| 2508AshExt6aBac.2508AshExt6aBac_S160_L001_R1_001 | 76.85950413 | 48 | 0.000484 | 484 |
| 2508AshExt6aBac.2508AshExt6aBac_S160_L001_R2_001 | 80.5785124 | 48 | 0.000484 | 484 |
| 2508AshExt6bBac.2508AshExt6bBac_S168_L001_R1_001 | 0 | 45 | 0.000007 | 7 |
| 2508AshExt6bBac.2508AshExt6bBac_S168_L001_R2_001 | 14.28571429 | 45 | 0.000007 | 7 |
| 2508AshExt6cBac.2508AshExt6cBac_S158_L001_R1_001 | 95.63631261 | 53 | 0.049889 | 49889 |
| 2508AshExt6cBac.2508AshExt6cBac_S158_L001_R2_001 | 96.62651085 | 54 | 0.049889 | 49889 |
| 2508AshInt6W8aBac.2508AshInt6W8aBac_S153_L001_R1_001 | 94.48617444 | 51 | 0.030487 | 30487 |
| 2508AshInt6W8aBac.2508AshInt6W8aBac_S153_L001_R2_001 | 95.61124414 | 52 | 0.030487 | 30487 |
| 2508AshInt6W8bBac.2508AshInt6W8bBac_S164_L001_R1_001 | 95.40827429 | 51 | 0.041357 | 41357 |
| 2508AshInt6W8bBac.2508AshInt6W8bBac_S164_L001_R2_001 | 95.71535653 | 52 | 0.041357 | 41357 |
| 2508AshInt6W8cBac.2508AshInt6W8cBac_S163_L001_R1_001 | 94.67664544 | 51 | 0.033137 | 33137 |
| 2508AshInt6W8cBac.2508AshInt6W8cBac_S163_L001_R2_001 | 95.27416483 | 52 | 0.033137 | 33137 |
| 2508AshInt6aBac.2508AshInt6aBac_S155_L001_R1_001 | 95.56206824 | 51 | 0.038216 | 38216 |
| 2508AshInt6aBac.2508AshInt6aBac_S155_L001_R2_001 | 96.89920452 | 52 | 0.038216 | 38216 |
| 2508AshInt6bBac.2508AshInt6bBac_S156_L001_R1_001 | 95.39440963 | 53 | 0.030159 | 30159 |
| 2508AshInt6bBac.2508AshInt6bBac_S156_L001_R2_001 | 95.85530024 | 54 | 0.030159 | 30159 |
| 2508AshInt6cBac.2508AshInt6cBac_S162_L001_R1_001 | 92.70977847 | 51 | 0.022345 | 22345 |
| 2508AshInt6cBac.2508AshInt6cBac_S162_L001_R2_001 | 93.58245693 | 52 | 0.022345 | 22345 |
| 2508Wmcc6W8aBac.2508Wmcc6W8aBac_S194_L001_R1_001 | 92.62638053 | 54 | 0.055685 | 55685 |
| 2508Wmcc6W8aBac.2508Wmcc6W8aBac_S194_L001_R2_001 | 93.40576457 | 55 | 0.055685 | 55685 |
| 2508Wmcc6W8bBac.2508Wmcc6W8bBac_S190_L001_R1_001 | 92.38659574 | 54 | 0.044553 | 44553 |
| 2508Wmcc6W8bBac.2508Wmcc6W8bBac_S190_L001_R2_001 | 92.89834579 | 55 | 0.044553 | 44553 |
| 2508Wmcc6W8cBac.2508Wmcc6W8cBac_S188_L001_R1_001 | 93.18347202 | 54 | 0.063214 | 63214 |
| 2508Wmcc6W8cBac.2508Wmcc6W8cBac_S188_L001_R2_001 | 94.03138545 | 55 | 0.063214 | 63214 |
| 2508Wmcc6aBac.2508Wmcc6aBac_S179_L001_R1_001 | 93.32976568 | 50 | 0.026161 | 26161 |
| 2508Wmcc6aBac.2508Wmcc6aBac_S179_L001_R2_001 | 93.79228623 | 51 | 0.026161 | 26161 |
| 2508Wmcc6bBac.2508Wmcc6bBac_S186_L001_R1_001 | 94.58545939 | 51 | 0.030769 | 30769 |
| 2508Wmcc6bBac.2508Wmcc6bBac_S186_L001_R2_001 | 95.62871722 | 52 | 0.030769 | 30769 |
| 2508Wmcc6cBac.2508Wmcc6cBac_S180_L001_R1_001 | 94.61498574 | 53 | 0.03857 | 38570 |
| 2508Wmcc6cBac.2508Wmcc6cBac_S180_L001_R2_001 | 95.41094115 | 53 | 0.03857 | 38570 |
| 2509AshExt6W12aBac.2509AshExt6W12aBac_S286_L001_R1_0 | 95.95365373 | 52 | 0.039097 | 39097 |
| 2509AshExt6W12aBac.2509AshExt6W12aBac_S286_L001_R2_0 | 94.24764048 | 53 | 0.039097 | 39097 |

| | | | | |
|---|---|---|---|---|
| 2509AshExt6W12bBac.2509AshExt6W12bBac_S292_L001_R1_0 | 97.03427198 | 54 | 0.082779 | 82779 |
| 2509AshExt6W12bBac.2509AshExt6W12bBac_S292_L001_R2_0 | 94.98181906 | 55 | 0.082779 | 82779 |
| 2509AshExt6W12cBac.2509AshExt6W12cBac_S290_L001_R1_0 | 96.08230063 | 52 | 0.038007 | 38007 |
| 2509AshExt6W12cBac.2509AshExt6W12cBac_S290_L001_R2_0 | 94.3036809 | 53 | 0.038007 | 38007 |
| 2509AshExt6W4aBac.2509AshExt6W4aBac_S298_L001_R1_001 | 96.95672716 | 51 | 0.035981 | 35981 |
| 2509AshExt6W4aBac.2509AshExt6W4aBac_S298_L001_R2_001 | 95.58378033 | 52 | 0.035981 | 35981 |
| 2509AshExt6W4bBac.2509AshExt6W4bBac_S302_L001_R1_001 | 95.69476526 | 52 | 0.035933 | 35933 |
| 2509AshExt6W4bBac.2509AshExt6W4bBac_S302_L001_R2_001 | 94.00829321 | 53 | 0.035933 | 35933 |
| 2509AshExt6W4cBac.2509AshExt6W4cBac_S301_L001_R1_001 | 97.56868185 | 52 | 0.077859 | 77859 |
| 2509AshExt6W4cBac.2509AshExt6W4cBac_S301_L001_R2_001 | 96.26889634 | 53 | 0.077859 | 77859 |
| 2509AshExt6aBac.2509AshExt6aBac_S279_L001_R1_001 | 97.31859584 | 48 | 0.019542 | 19542 |
| 2509AshExt6aBac.2509AshExt6aBac_S279_L001_R2_001 | 96.47426057 | 49 | 0.019542 | 19542 |
| 2509AshExt6bBac.2509AshExt6bBac_S280_L001_R1_001 | 97.28541777 | 52 | 0.024571 | 24571 |
| 2509AshExt6bBac.2509AshExt6bBac_S280_L001_R2_001 | 95.12026373 | 53 | 0.024571 | 24571 |
| 2509AshExt6cBac.2509AshExt6cBac_S288_L001_R1_001 | 97.57853638 | 49 | 0.027917 | 27917 |
| 2509AshExt6cBac.2509AshExt6cBac_S288_L001_R2_001 | 96.09556901 | 50 | 0.027917 | 27917 |
| 2509AshInt6W12aBac.2509AshInt6W12aBac_S291_L001_R1_00 | 97.82865449 | 52 | 0.066042 | 66042 |
| 2509AshInt6W12aBac.2509AshInt6W12aBac_S291_L001_R2_00 | 96.86411677 | 53 | 0.066042 | 66042 |
| 2509AshInt6W12bBac.2509AshInt6W12bBac_S295_L001_R1_00 | 97.1677035 | 51 | 0.045652 | 45652 |
| 2509AshInt6W12bBac.2509AshInt6W12bBac_S295_L001_R2_00 | 95.95855603 | 52 | 0.045652 | 45652 |
| 2509AshInt6W12cBac.2509AshInt6W12cBac_S284_L001_R1_00 | 97.32442219 | 51 | 0.044738 | 44738 |
| 2509AshInt6W12cBac.2509AshInt6W12cBac_S284_L001_R2_00 | 96.68514462 | 52 | 0.044738 | 44738 |
| 2509AshInt6W4aBac.2509AshInt6W4aBac_S285_L001_R1_001 | 98.3847981 | 49 | 0.05473 | 54730 |
| 2509AshInt6W4aBac.2509AshInt6W4aBac_S285_L001_R2_001 | 97.72702357 | 49 | 0.05473 | 54730 |
| 2509AshInt6W4bBac.2509AshInt6W4bBac_S283_L001_R1_001 | 97.77399056 | 50 | 0.03814 | 38140 |
| 2509AshInt6W4bBac.2509AshInt6W4bBac_S283_L001_R2_001 | 97.25485055 | 51 | 0.03814 | 38140 |
| 2509AshInt6W4cBac.2509AshInt6W4cBac_S287_L001_R1_001 | 97.28734221 | 51 | 0.026063 | 26063 |
| 2509AshInt6W4cBac.2509AshInt6W4cBac_S287_L001_R2_001 | 96.57752369 | 51 | 0.026063 | 26063 |
| 2509Wmcc6W12aBac.2509Wmcc6W12aBac_S289_L001_R1_00 | 96.19449107 | 55 | 0.052424 | 52424 |
| 2509Wmcc6W12aBac.2509Wmcc6W12aBac_S289_L001_R2_00 | 94.7867389 | 56 | 0.052424 | 52424 |
| 2509Wmcc6W12bBac.2509Wmcc6W12bBac_S299_L001_R1_00 | 96.62284331 | 55 | 0.065262 | 65262 |
| 2509Wmcc6W12bBac.2509Wmcc6W12bBac_S299_L001_R2_00 | 94.99862094 | 56 | 0.065262 | 65262 |
| 2509Wmcc6W12cBac.2509Wmcc6W12cBac_S296_L001_R1_00 | 96.03271599 | 55 | 0.053185 | 53185 |
| 2509Wmcc6W12cBac.2509Wmcc6W12cBac_S296_L001_R2_00 | 92.69718906 | 56 | 0.053185 | 53185 |
| 2509Wmcc6W4aBac.2509Wmcc6W4aBac_S282_L001_R1_001 | 95.73984526 | 52 | 0.02068 | 20680 |
| 2509Wmcc6W4aBac.2509Wmcc6W4aBac_S282_L001_R2_001 | 93.68955513 | 53 | 0.02068 | 20680 |
| 2509Wmcc6W4bBac.2509Wmcc6W4bBac_S278_L001_R1_001 | 96.29050473 | 52 | 0.034533 | 34533 |

| | | | | |
|---|---|---|---|---|
| 2509Wmcc6W4bBac.2509Wmcc6W4bBac_S278_L001_R2_001 | 94.2692497 | 53 | 0.034533 | 34533 |
| 2509Wmcc6W4cBac.2509Wmcc6W4cBac_S277_L001_R1_001 | 96.85598656 | 54 | 0.036323 | 36323 |
| 2509Wmcc6W4cBac.2509Wmcc6W4cBac_S277_L001_R2_001 | 94.75263607 | 55 | 0.036323 | 36323 |
| 2509Wmcc6aBac.2509Wmcc6aBac_S281_L001_R1_001 | 97.65238343 | 52 | 0.030712 | 30712 |
| 2509Wmcc6aBac.2509Wmcc6aBac_S281_L001_R2_001 | 96.34344881 | 53 | 0.030712 | 30712 |
| 2509Wmcc6bBac.2509Wmcc6bBac_S297_L001_R1_001 | 92.37510955 | 50 | 0.003423 | 3423 |
| 2509Wmcc6bBac.2509Wmcc6bBac_S297_L001_R2_001 | 90.47619048 | 50 | 0.003423 | 3423 |
| 2509Wmcc6cBac.2509Wmcc6cBac_S300_L001_R1_001 | 96.32817173 | 53 | 0.030094 | 30094 |
| 2509Wmcc6cBac.2509Wmcc6cBac_S300_L001_R2_001 | 94.71323187 | 54 | 0.030094 | 30094 |
| 2510AshExtW8aBac.2510AshExtW8aBac_S308_L001_R1_001 | 95.75903362 | 53 | 0.057369 | 57369 |
| 2510AshExtW8aBac.2510AshExtW8aBac_S308_L001_R2_001 | 93.74749429 | 54 | 0.057369 | 57369 |
| 2510AshExtW8bBac.2510AshExtW8bBac_S303_L001_R1_001 | 93.89801014 | 53 | 0.026434 | 26434 |
| 2510AshExtW8bBac.2510AshExtW8bBac_S303_L001_R2_001 | 91.76817735 | 53 | 0.026434 | 26434 |
| 2510AshExtW8cBac.2510AshExtW8cBac_S307_L001_R1_001 | 95.89738945 | 53 | 0.044167 | 44167 |
| 2510AshExtW8cBac.2510AshExtW8cBac_S307_L001_R2_001 | 94.06344103 | 53 | 0.044167 | 44167 |
| 2510AshExtW8dBac.2510AshExtW8dBac_S304_L001_R1_001 | 96.23422809 | 53 | 0.082108 | 82108 |
| 2510AshExtW8dBac.2510AshExtW8dBac_S304_L001_R2_001 | 93.00189994 | 54 | 0.082108 | 82108 |
| 2510WmccW8aBac.2510WmccW8aBac_S294_L001_R1_001 | 95.30213792 | 53 | 0.043126 | 43126 |
| 2510WmccW8aBac.2510WmccW8aBac_S294_L001_R2_001 | 93.23378009 | 54 | 0.043126 | 43126 |
| 2510WmccW8bBac.2510WmccW8bBac_S293_L001_R1_001 | 96.81642149 | 54 | 0.067911 | 67911 |
| 2510WmccW8bBac.2510WmccW8bBac_S293_L001_R2_001 | 94.76373489 | 54 | 0.067911 | 67911 |
| 2510WmccW8cBac.2510WmccW8cBac_S306_L001_R1_001 | 95.73062073 | 53 | 0.026936 | 26936 |
| 2510WmccW8cBac.2510WmccW8cBac_S306_L001_R2_001 | 94.08226908 | 54 | 0.026936 | 26936 |
| 2510WmccW8dBac.2510WmccW8dBac_S305_L001_R1_001 | 94.39149192 | 53 | 0.049835 | 49835 |
| 2510WmccW8dBac.2510WmccW8dBac_S305_L001_R2_001 | 91.43573794 | 54 | 0.049835 | 49835 |
| EID2510BlankBac.EID2510BlankBac_S201_L001_R1_001 | 91.61258113 | 52 | 0.002003 | 2003 |
| EID2510BlankBac.EID2510BlankBac_S201_L001_R2_001 | 91.06340489 | 52 | 0.002003 | 2003 |
| EID2513BlankBac.EID2513BlankBac_S309_L001_R1_001 | 94.98189343 | 52 | 0.001933 | 1933 |
| EID2513BlankBac.EID2513BlankBac_S309_L001_R2_001 | 92.96430419 | 52 | 0.001933 | 1933 |
| RID001584Blank.RID001584Blank_S207_L001_R1_001 | 36.84210526 | 52 | 0.000038 | 38 |
| RID001584Blank.RID001584Blank_S207_L001_R2_001 | 47.36842105 | 53 | 0.000038 | 38 |
| RID001584Zymo.RID001584Zymo_S208_L001_R1_001 | 88.68513461 | 54 | 0.002563 | 2563 |
| RID001584Zymo.RID001584Zymo_S208_L001_R2_001 | 90.63597347 | 54 | 0.002563 | 2563 |
| RID001608Blank.RID001608Blank_S310_L001_R1_001 | 20 | 53 | 0.00001 | 10 |
| RID001608Blank.RID001608Blank_S310_L001_R2_001 | 20 | 55 | 0.00001 | 10 |
| RID001608Zymo.RID001608Zymo_S311_L001_R1_001 | 93.67710252 | 54 | 0.001629 | 1629 |
| RID001608Zymo.RID001608Zymo_S311_L001_R2_001 | 93.18600368 | 54 | 0.001629 | 1629 |
