## Supplementary material for "Microbial, dietary insect, and pathogen communities in fresh and decomposing guano of anthropic little brown bat (Myotis lucifugus) maternity colonies": Table S6

| Subset | ModelVariables | Df | SumSq | MeanSq | F Value | Pr(>F) |
|---|---|---|---|---|---|---|
| None | shannon~Age | 3 | 48.12 | 16.042 | 63.65 | <2.00E-16 |
| | shannon~Site | 2 | 13.43 | 6.717 | 26.65 | 2.84E-09 |
| | shannon~Age:Site | 6 | 10.92 | 1.82 | 7.22 | 5.50E-06 |
| | Residuals | 68 | 17.14 | 0.252 | | |
| | | **Tukey** | **Diff** | **Lower** | **Upper** | **p adj** |
| | | Week 12:ASHFExt-Fresh:ASHFExt | 1.885099 | 0.7884081 | 2.9817899 | 0.0000111 |
| | | Week 4:ASHFExt-Fresh:ASHFExt | 1.338019833 | 0.4885267 | 2.1875129 | 0.0000733 |
| | | Week 8:ASHFExt-Fresh:ASHFExt | 2.1588015 | 1.3093084 | 3.0082946 | 0 |
| | | Fresh:ASHFInt-Fresh:ASHFExt | -0.2441215 | -0.9377297 | 0.4494867 | 0.9879725 |
| | | Week 12:ASHFInt-Fresh:ASHFExt | 0.393556 | -0.7031349 | 1.4902469 | 0.9859786 |
| | | Week 4:ASHFInt-Fresh:ASHFExt | -0.013509167 | -0.8630023 | 0.8359839 | 1 |
| | | Week 8:ASHFInt-Fresh:ASHFExt | -0.093082 | -1.1897729 | 1.0036089 | 1 |
| | | Fresh:WMCC-Fresh:ASHFExt | -0.246366154 | -0.926505 | 0.4337727 | 0.9849141 |
| | | Week 12:WMCC-Fresh:ASHFExt | 1.928377333 | 0.8316864 | 3.0250682 | 0.0000065 |
| | | Week 4:WMCC-Fresh:ASHFExt | 1.36997 | 0.5204769 | 2.2194631 | 0.000045 |
| | | Week 8:WMCC-Fresh:ASHFExt | 2.056494714 | 1.2484656 | 2.8645238 | 0 |
| | | Week 4:ASHFExt-Week 12:ASHFExt | -0.547079167 | -1.7484438 | 0.6542855 | 0.9232481 |
| | | Week 8:ASHFExt-Week 12:ASHFExt | 0.2737025 | -0.9276622 | 1.4750672 | 0.999747 |
| | | Fresh:ASHFInt-Week 12:ASHFExt | -2.1292205 | -3.2259114 | -1.0325296 | 0.0000005 |
| | | Week 12:ASHFInt-Week 12:ASHFExt | -1.491543 | -2.8787594 | -0.1043266 | 0.0244436 |
| | | Week 4:ASHFInt-Week 12:ASHFExt | -1.898608167 | -3.0999728 | -0.6972435 | 0.0000684 |
| | | Week 8:ASHFInt-Week 12:ASHFExt | -1.978181 | -3.3653974 | -0.5909646 | 0.0004799 |
| | | Fresh:WMCC-Week 12:ASHFExt | -2.131465154 | -3.2196873 | -1.043243 | 0.0000004 |
| | | Week 12:WMCC-Week 12:ASHFExt | 0.043278333 | -1.3439381 | 1.4304948 | 1 |
| | | Week 4:WMCC-Week 12:ASHFExt | -0.515129 | -1.7164937 | 0.6862357 | 0.9482888 |
| | | Week 8:WMCC-Week 12:ASHFExt | 0.171395714 | -1.0010162 | 1.3438076 | 0.9999971 |
| | | Week 8:ASHFExt-Week 4:ASHFExt | 0.820781667 | -0.1601285 | 1.8016918 | 0.1903316 |
| | | Fresh:ASHFInt-Week 4:ASHFExt | -1.582141333 | -2.4316344 | -0.7326482 | 0.0000016 |
| | | Week 12:ASHFInt-Week 4:ASHFExt | -0.944463833 | -2.1458285 | 0.2569008 | 0.2683464 |
| | | Week 4:ASHFInt-Week 4:ASHFExt | -1.351529 | -2.3324392 | -0.3706188 | 0.000862 |
| | | Week 8:ASHFInt-Week 4:ASHFExt | -1.431101833 | -2.6324665 | -0.2297372 | 0.0073313 |
| | | Fresh:WMCC-Week 4:ASHFExt | -1.584385987 | -2.4229174 | -0.7458545 | 0.0000011 |
| | | Week 12:WMCC-Week 4:ASHFExt | 0.5903575 | -0.6110072 | 1.7917222 | 0.8781709 |
| | | Week 4:WMCC-Week 4:ASHFExt | 0.031950167 | -0.94896 | 1.0128603 | 1 |
| | | Week 8:WMCC-Week 4:ASHFExt | 0.718474881 | -0.2267538 | 1.6637036 | 0.3155744 |
| | | Fresh:ASHFInt-Week 8:ASHFExt | -2.402923 | -3.2524161 | -1.5534299 | 0 |
| | | Week 12:ASHFInt-Week 8:ASHFExt | -1.7652455 | -2.9666102 | -0.5638808 | 0.0002806 |
| | | Week 4:ASHFInt-Week 8:ASHFExt | -2.172310667 | -3.1532208 | -1.1914005 | 0 |

| | | | | | | |
|---|---|---|---|---|---|---|
| | | Week 8:ASHFInt-Week 8:ASHFExt | -2.2518835 | -3.4532482 | -1.0505188 | 0.0000013 |
| | | Fresh:WMCC-Week 8:ASHFExt | -2.405167654 | -3.2436991 | -1.5666362 | 0 |
| | | Week 12:WMCC-Week 8:ASHFExt | -0.230424167 | -1.4317888 | 0.9709405 | 0.9999533 |
| | | Week 4:WMCC-Week 8:ASHFExt | -0.7888315 | -1.7697417 | 0.1920787 | 0.2384494 |
| | | Week 8:WMCC-Week 8:ASHFExt | -0.102306786 | -1.0475355 | 0.8429219 | 0.9999999 |
| | | Week 12:ASHFInt-Fresh:ASHFInt | 0.6376775 | -0.4590134 | 1.7343684 | 0.7125955 |
| | | Week 4:ASHFInt-Fresh:ASHFInt | 0.230612333 | -0.6188808 | 1.0801054 | 0.9986952 |
| | | Week 8:ASHFInt-Fresh:ASHFInt | 0.1510395 | -0.9456514 | 1.2477304 | 0.9999984 |
| | | Fresh:WMCC-Fresh:ASHFInt | -0.002244654 | -0.6823835 | 0.6778942 | 1 |
| | | Week 12:WMCC-Fresh:ASHFInt | 2.172498833 | 1.0758079 | 3.2691897 | 0.0000003 |
| | | Week 4:WMCC-Fresh:ASHFInt | 1.6140915 | 0.7645984 | 2.4635846 | 0.0000009 |
| | | Week 8:WMCC-Fresh:ASHFInt | 2.300616214 | 1.4925871 | 3.1086453 | 0 |
| | | Week 4:ASHFInt-Week 12:ASHFInt | -0.407065167 | -1.6084298 | 0.7942995 | 0.9911425 |
| | | Week 8:ASHFInt-Week 12:ASHFInt | -0.486638 | -1.8738544 | 0.9005784 | 0.9882838 |
| | | Fresh:WMCC-Week 12:ASHFInt | -0.639922154 | -1.7281443 | 0.4483 | 0.6981664 |
| | | Week 12:WMCC-Week 12:ASHFInt | 1.534821333 | 0.1476049 | 2.9220378 | 0.0178814 |
| | | Week 4:WMCC-Week 12:ASHFInt | 0.976414 | -0.2249507 | 2.1777787 | 0.2250492 |
| | | Week 8:WMCC-Week 12:ASHFInt | 1.662938714 | 0.4905268 | 2.8353506 | 0.0005269 |
| | | Week 8:ASHFInt-Week 4:ASHFInt | -0.079572833 | -1.2809375 | 1.1217918 | 1 |
| | | Fresh:WMCC-Week 4:ASHFInt | -0.232856987 | -1.0713884 | 0.6056745 | 0.9983984 |
| | | Week 12:WMCC-Week 4:ASHFInt | 1.9418865 | 0.7405218 | 3.1432512 | 0.0000429 |
| | | Week 4:WMCC-Week 4:ASHFInt | 1.383479167 | 0.402569 | 2.3643893 | 0.0005809 |
| | | Week 8:WMCC-Week 4:ASHFInt | 2.070003881 | 1.1247752 | 3.0152326 | 0 |
| | | Fresh:WMCC-Week 8:ASHFInt | -0.153284154 | -1.2415063 | 0.934938 | 0.999998 |
| | | Week 12:WMCC-Week 8:ASHFInt | 2.021459333 | 0.6342429 | 3.4086758 | 0.0003265 |
| | | Week 4:WMCC-Week 8:ASHFInt | 1.463052 | 0.2616873 | 2.6644167 | 0.00548 |
| | | Week 8:WMCC-Week 8:ASHFInt | 2.149576714 | 0.9771648 | 3.3219886 | 0.0000024 |
| | | Week 12:WMCC-Fresh:WMCC | 2.174743487 | 1.0865214 | 3.2629656 | 0.0000002 |
| | | Week 4:WMCC-Fresh:WMCC | 1.616336154 | 0.7778047 | 2.4548676 | 0.0000006 |
| | | Week 8:WMCC-Fresh:WMCC | 2.302860868 | 1.5063638 | 3.0993579 | 0 |
| | | Week 4:WMCC-Week 12:WMCC | -0.558407333 | -1.759772 | 0.6429573 | 0.9127131 |
| | | Week 8:WMCC-Week 12:WMCC | 0.128117381 | -1.0442945 | 1.3005293 | 0.9999999 |
| | | Week 8:WMCC-Week 4:WMCC | 0.686524714 | -0.258704 | 1.6317534 | 0.3833791 |
| **Subset** | **ModelVariables** | **Df** | **SumSq** | **MeanSq** | **F Value** | **Pr(>F)** |
| None | invsimpson~Age | 3 | 2498.7 | 832.9 | 34.227 | 1.34E-13 |
| | invsimpson~Site | 2 | 616.9 | 308.4 | 12.675 | 2.10E-05 |
| | invsimpson~Age:Site | 6 | 750.5 | 125.1 | 5.141 | 2.09E-04 |
| | Residuals | 68 | 1654.7 | 24.3 | | |
| | | **Tukey** | **Diff** | **Lower** | **Upper** | **p adj** |

| | | | | | | |
|---|---|---|---|---|---|---|
| | | Week 12:ASHFExt-Fresh:ASHFExt | 12.60570342 | 1.82915141 | 23.38225543 | 0.0092399 |
| | | Week 4:ASHFExt-Fresh:ASHFExt | 6.85089808 | -1.49658321 | 15.19837938 | 0.2130768 |
| | | Week 8:ASHFExt-Fresh:ASHFExt | 20.85096708 | 12.50348579 | 29.19844838 | 0 |
| | | Fresh:ASHFInt-Fresh:ASHFExt | -0.571409 | -7.38709893 | 6.244280935 | 1 |
| | | Week 12:ASHFInt-Fresh:ASHFExt | 0.71276875 | -10.06378326 | 11.48932076 | 1 |
| | | Week 4:ASHFInt-Fresh:ASHFExt | -0.04385442 | -8.39133571 | 8.303626876 | 1 |
| | | Week 8:ASHFInt-Fresh:ASHFExt | -0.38287092 | -11.15942293 | 10.39368109 | 1 |
| | | Fresh:WMCC-Fresh:ASHFExt | -0.21332958 | -6.89666342 | 6.470004258 | 1 |
| | | Week 12:WMCC-Fresh:ASHFExt | 9.04208308 | -1.73446893 | 19.81863509 | 0.1872254 |
| | | Week 4:WMCC-Fresh:ASHFExt | 8.10200042 | -0.24548088 | 16.44948171 | 0.0651092 |
| | | Week 8:WMCC-Fresh:ASHFExt | 15.3044357 | 7.36439722 | 23.24447419 | 0.0000006 |
| | | Week 4:ASHFExt-Week 12:ASHFExt | -5.75480533 | -17.55992659 | 6.050315922 | 0.8837142 |
| | | Week 8:ASHFExt-Week 12:ASHFExt | 8.24526367 | -3.55985759 | 20.05038492 | 0.4439137 |
| | | Fresh:ASHFInt-Week 12:ASHFExt | -13.17711242 | -23.95366443 | -2.400560407 | 0.0051891 |
| | | Week 12:ASHFInt-Week 12:ASHFExt | -11.89293467 | -25.52431454 | 1.738445203 | 0.1461672 |
| | | Week 4:ASHFInt-Week 12:ASHFExt | -12.64955783 | -24.45467909 | -0.844436578 | 0.0253456 |
| | | Week 8:ASHFInt-Week 12:ASHFExt | -12.98857433 | -26.6199542 | 0.642805536 | 0.0760058 |
| | | Fresh:WMCC-Week 12:ASHFExt | -12.819033 | -23.51236715 | -2.125698854 | 0.0067529 |
| | | Week 12:WMCC-Week 12:ASHFExt | -3.56362033 | -17.1950002 | 10.06775954 | 0.9990756 |
| | | Week 4:WMCC-Week 12:ASHFExt | -4.503703 | -16.30882426 | 7.301418255 | 0.9775652 |
| | | Week 8:WMCC-Week 12:ASHFExt | 2.69873229 | -8.82188641 | 14.21935098 | 0.9996698 |
| | | Week 8:ASHFExt-Week 4:ASHFExt | 14.000069 | 4.36122786 | 23.63891014 | 0.0003463 |
| | | Fresh:ASHFInt-Week 4:ASHFExt | -7.42230708 | -15.76978838 | 0.925174209 | 0.1284438 |
| | | Week 12:ASHFInt-Week 4:ASHFExt | -6.13812933 | -17.94325059 | 5.666991922 | 0.8332475 |
| | | Week 4:ASHFInt-Week 4:ASHFExt | -6.8947525 | -16.53359364 | 2.744088643 | 0.4068346 |
| | | Week 8:ASHFInt-Week 4:ASHFExt | -7.233769 | -19.03889026 | 4.571352255 | 0.6423397 |
| | | Fresh:WMCC-Week 4:ASHFExt | -7.06422767 | -15.30399501 | 1.175539677 | 0.1638144 |
| | | Week 12:WMCC-Week 4:ASHFExt | 2.191185 | -9.61393626 | 13.99630626 | 0.9999664 |
| | | Week 4:WMCC-Week 4:ASHFExt | 1.25110233 | -8.38773881 | 10.88994348 | 0.9999992 |
| | | Week 8:WMCC-Week 4:ASHFExt | 8.45353762 | -0.83468211 | 17.74175735 | 0.1086539 |
| | | Fresh:ASHFInt-Week 8:ASHFExt | -21.42237608 | -29.76985738 | -13.07489479 | 0 |
| | | Week 12:ASHFInt-Week 8:ASHFExt | -20.13819833 | -31.94331959 | -8.333077078 | 0.0000132 |
| | | Week 4:ASHFInt-Week 8:ASHFExt | -20.8948215 | -30.53366264 | -11.25598036 | 0 |
| | | Week 8:ASHFInt-Week 8:ASHFExt | -21.233838 | -33.03895926 | -9.428716745 | 0.0000038 |
| | | Fresh:WMCC-Week 8:ASHFExt | -21.06429667 | -29.30406401 | -12.82452932 | 0 |
| | | Week 12:WMCC-Week 8:ASHFExt | -11.808884 | -23.61400526 | -0.003762745 | 0.0498546 |
| | | Week 4:WMCC-Week 8:ASHFExt | -12.74896667 | -22.38780781 | -3.110125524 | 0.00166 |
| | | Week 8:WMCC-Week 8:ASHFExt | -5.54653138 | -14.83475111 | 3.741688352 | 0.6778624 |
| | | Week 12:ASHFInt-Fresh:ASHFInt | 1.28417775 | -9.49237426 | 12.06072976 | 0.9999997 |

| | | | | | | |
|---|---|---|---|---|---|---|
| | | Week 4:ASHFInt-Fresh:ASHFInt | 0.52755458 | -7.81992671 | 8.875035876 | 1 |
| | | Week 8:ASHFInt-Fresh:ASHFInt | 0.18853808 | -10.58801393 | 10.96509009 | 1 |
| | | Fresh:WMCC-Fresh:ASHFInt | 0.35807942 | -6.32525442 | 7.041413258 | 1 |
| | | Week 12:WMCC-Fresh:ASHFInt | 9.61349208 | -1.16305993 | 20.39004409 | 0.1255393 |
| | | Week 4:WMCC-Fresh:ASHFInt | 8.67340942 | 0.32592812 | 17.02089071 | 0.0347241 |
| | | Week 8:WMCC-Fresh:ASHFInt | 15.8758447 | 7.93580622 | 23.81588319 | 0.0000002 |
| | | Week 4:ASHFInt-Week 12:ASHFInt | -0.75662317 | -12.56174442 | 11.04849809 | 1 |
| | | Week 8:ASHFInt-Week 12:ASHFInt | -1.09563967 | -14.72701954 | 12.5357402 | 1 |
| | | Fresh:WMCC-Week 12:ASHFInt | -0.92609833 | -11.61943248 | 9.767235813 | 1 |
| | | Week 12:WMCC-Week 12:ASHFInt | 8.32931433 | -5.30206554 | 21.9606942 | 0.646306 |
| | | Week 4:WMCC-Week 12:ASHFInt | 7.38923167 | -4.41588959 | 19.19435292 | 0.6118339 |
| | | Week 8:WMCC-Week 12:ASHFInt | 14.59166695 | 3.07104826 | 26.11228565 | 0.0031732 |
| | | Week 8:ASHFInt-Week 4:ASHFInt | -0.3390165 | -12.14413776 | 11.46610476 | 1 |
| | | Fresh:WMCC-Week 4:ASHFInt | -0.16947517 | -8.40924251 | 8.070292177 | 1 |
| | | Week 12:WMCC-Week 4:ASHFInt | 9.0859375 | -2.71918376 | 20.89105876 | 0.2977177 |
| | | Week 4:WMCC-Week 4:ASHFInt | 8.14585483 | -1.49298631 | 17.78469598 | 0.1792124 |
| | | Week 8:WMCC-Week 4:ASHFInt | 15.34829012 | 6.06007039 | 24.63650985 | 0.0000267 |
| | | Fresh:WMCC-Week 8:ASHFInt | 0.16954133 | -10.52379281 | 10.86287548 | 1 |
| | | Week 12:WMCC-Week 8:ASHFInt | 9.424954 | -4.20642587 | 23.05633387 | 0.4597216 |
| | | Week 4:WMCC-Week 8:ASHFInt | 8.48487133 | -3.32024992 | 20.28999259 | 0.39944 |
| | | Week 8:WMCC-Week 8:ASHFInt | 15.68730662 | 4.16668792 | 27.20792531 | 0.0010462 |
| | | Week 12:WMCC-Fresh:WMCC | 9.25541267 | -1.43792148 | 19.94874681 | 0.1540643 |
| | | Week 4:WMCC-Fresh:WMCC | 8.31533 | 0.07556266 | 16.55509734 | 0.0459611 |
| | | Week 8:WMCC-Fresh:WMCC | 15.51776529 | 7.69104611 | 23.34448447 | 0.0000003 |
| | | Week 4:WMCC-Week 12:WMCC | -0.94008267 | -12.74520392 | 10.86503859 | 1 |
| | | Week 8:WMCC-Week 12:WMCC | 6.26235262 | -5.25826608 | 17.78297131 | 0.7903613 |
| | | Week 8:WMCC-Week 4:WMCC | 7.20243529 | -2.08578445 | 16.49065502 | 0.2872191 |
| **Subset** | **ModelVariables** | **Df** | **SumSq** | **MeanSq** | **F Value** | **Pr(>F)** |
| Fresh | shannon~Month | 3 | 0.184 | 0.0615 | 0.158 | 9.23E-01 |
| *5 September samples removed | shannon~Site | 2 | 0.506 | 0.2531 | 0.65 | 5.33E-01 |
| | shannon~Month:Site | 6 | 1.347 | 0.2245 | 0.576 | 7.45E-01 |
| | Residuals | 20 | 7.792 | 0.3896 | | |
| **Subset** | **ModelVariables** | **Df** | **SumSq** | **MeanSq** | **F Value** | **Pr(>F)** |
| Fresh | invsimpson~Month | 3 | 4.48 | 1.4938 | 0.518 | 6.75E-01 |
| *5 September samples removed | invsimpson~Site | 2 | 1.13 | 0.5634 | 0.195 | 8.24E-01 |
| | invsimpson~Month:Site | 6 | 12.72 | 2.1194 | 0.735 | 6.28E-01 |
| | Residuals | 20 | 57.7 | 2.885 | | |
