## Supplementary material for "Microbial, dietary insect, and pathogen communities in fresh and decomposing guano of anthropic little brown bat (Myotis lucifugus) maternity colonies": Table S7

| Distance Matrix | Factor | Df | SumOfSqs | R2 | F | Pr(>F) |
|---|---|---|---|---|---|---|
| Jaccard | Site | 2 | 3.478 | 0.10612 | 5.3583 | 0.001 |
| Jaccard | Age | 3 | 3.332 | 0.10166 | 3.422 | 0.001 |
| Jaccard | Site* | 2 | 1.304 | 0.11223 | 2.0422 | 0.001 |
| Jaccard | Month* | 3 | 1.5855 | 0.13646 | 1.6554 | 0.001 |
| Jaccard | Site:Age | 6 | 3.894 | 0.11882 | 1.9997 | 0.001 |
| Jaccard | Site:Month* | 6 | 2.3441 | 0.20176 | 1.2238 | 0.003 |
| Bray-Curtis | Site | 2 | 4.1755 | 0.14100 | 8.7585 | 0.001 |
| Bray-Curtis | Age | 3 | 4.4404 | 0.14994 | 6.2095 | 0.001 |
| Bray-Curtis | Site* | 2 | 0.5538 | 0.05684 | 1.2528 | 0.224 |
| Bray-Curtis | Month* | 3 | 3.6881 | 0.37854 | 5.5623 | 0.001 |
| Bray-Curtis | Site:Age | 6 | 4.7895 | 0.16173 | 3.3489 | 0.001 |
| Bray-Curtis | Site:Month* | 6 | 1.0808 | 0.11093 | 0.815 | 0.765 |
| Yue & Clayton | Site | 2 | 4.0539 | 0.12871 | 7.4938 | 0.001 |
| Yue & Clayton | Age | 3 | 3.9721 | 0.12612 | 4.8952 | 0.001 |
| Yue & Clayton | Site* | 2 | 0.4948 | 0.04686 | 1.0447 | 0.361 |
| Yue & Clayton | Month* | 3 | 4.3729 | 0.41418 | 6.1555 | 0.001 |
| Yue & Clayton | Site:Age | 6 | 5.0768 | 0.16119 | 3.1282 | 0.001 |
| Yue & Clayton | Site:Month* | 6 | 0.9543 | 0.09038 | 0.6716 | 0.916 |

*Tests run on only fresh samples from May-August
