## Supplementary material for "Microbial, dietary insect, and pathogen communities in fresh and decomposing guano of anthropic little brown bat (Myotis lucifugus) maternity colonies": Table S8

| Site | Family | Status | Mean Relative Abundance |
|---|---|---|---|
| AshfExt | Clostridiaceae | Fresh | 47.83077 |
| AshfExt | Moraxellaceae | Aged | 30.31385 |
| AshfExt | Enterobacteriaceae | Fresh | 17.66462 |
| AshfExt | Oxalobacteraceae | Aged | 11.48846 |
| AshfExt | Sphingobacteriaceae | Aged | 6.773077 |
| AshfExt | Moraxellaceae | Fresh | 5.686154 |
| AshfExt | Weeksellaceae | Aged | 5.373077 |
| AshfExt | Enterobacteriaceae | Aged | 4.603077 |
| AshfExt | Pseudomonadaceae | Aged | 4.457692 |
| AshfExt | Micrococcaceae | Aged | 4.402308 |
| AshfExt | Nocardiaceae | Aged | 4.136923 |
| AshfExt | Carnobacteriaceae | Aged | 3.9 |
| AshfExt | Bacillaceae | Fresh | 3.593846 |
| AshfExt | Comamonadaceae | Aged | 2.909231 |
| AshfExt | Aeromonadaceae | Fresh | 2.355385 |
| AshfExt | Enterococcaceae | Fresh | 2.286923 |
| AshfExt | Nocardiaceae | Fresh | 2.186923 |
| AshfExt | Planococcaceae | Aged | 2.148462 |
| AshfExt | Hafniaceae | Fresh | 1.941538 |
| AshfExt | Enterobacterales_unclassified | Aged | 1.864615 |
| AshfExt | Dietziaceae | Aged | 1.524615 |
| AshfExt | Staphylococcaceae | Fresh | 1.31 |
| AshfExt | Oxalobacteraceae | Fresh | 1.181538 |
| AshfExt | Peptostreptococcaceae | Fresh | 1.15 |
| AshfExt | Chitinophagaceae | Aged | 1.123846 |
| AshfExt | Microbacteriaceae | Aged | 1.044615 |
| AshfExt | Weeksellaceae | Fresh | 0.987692 |
| AshfExt | Myxococcaceae | Aged | 0.974615 |
| AshfExt | Streptococcaceae | Fresh | 0.923846 |
| AshfExt | Diplorickettsiaceae | Fresh | 0.873846 |
| AshfExt | Micrococcaceae | Fresh | 0.857692 |

| | | | |
|---|---|---|---|
| AshfExt | Streptococcaceae | Aged | 0.845385 |
| AshfExt | Carnobacteriaceae | Fresh | 0.83 |
| AshfExt | Planococcaceae | Fresh | 0.783846 |
| AshfExt | Rhizobiaceae | Aged | 0.777692 |
| AshfExt | Deinococcaceae | Aged | 0.757692 |
| AshfExt | Enterobacterales_unclassified | Fresh | 0.712308 |
| AshfExt | Enterococcaceae | Aged | 0.689231 |
| AshfExt | Rhodobacteraceae | Aged | 0.676923 |
| AshfExt | Micrococcales_unclassified | Aged | 0.650769 |
| AshfExt | Lachnospiraceae | Fresh | 0.646154 |
| AshfExt | Hymenobacteraceae | Aged | 0.610769 |
| AshfExt | Corynebacteriaceae | Fresh | 0.581538 |
| AshfExt | Sphingomonadaceae | Aged | 0.51 |
| AshfExt | Paenibacillaceae | Aged | 0.508462 |
| AshfExt | Dietziaceae | Fresh | 0.482308 |
| AshfExt | Flavobacteriaceae | Aged | 0.45 |
| AshfExt | Fusobacteriaceae | Fresh | 0.449231 |
| AshfExt | Sphingobacteriaceae | Fresh | 0.439231 |
| AshfExt | Xanthomonadaceae | Aged | 0.439231 |
| AshfExt | Pseudomonadaceae | Fresh | 0.425385 |
| AshfExt | Devosiaceae | Aged | 0.418462 |
| AshfExt | Beijerinckiaceae | Aged | 0.386154 |
| AshfExt | Spirosomaceae | Aged | 0.380769 |
| AshfExt | Peptostreptococcaceae | Aged | 0.343846 |
| AshfExt | Bacteria_unclassified | Fresh | 0.321538 |
| AshfExt | Microbacteriaceae | Fresh | 0.316923 |
| AshfExt | Acetobacteraceae | Aged | 0.286154 |
| AshfExt | Caulobacteraceae | Aged | 0.286154 |
| AshfExt | Corynebacteriales_unclassified | Fresh | 0.249231 |
| AshfExt | Exiguobacteraceae | Aged | 0.246154 |
| AshfExt | Synergistaceae | Fresh | 0.24 |

| | | | |
|---|---|---|---|
| AshfExt | Nocardioidaceae | Aged | 0.236923 |
| AshfExt | Desulfovibrionaceae | Fresh | 0.213077 |
| AshfExt | Clostridiaceae | Aged | 0.207692 |
| AshfExt | Mycoplasmataceae | Fresh | 0.203846 |
| AshfExt | Alcaligenaceae | Aged | 0.202308 |
| AshfExt | Abditibacteriaceae | Aged | 0.191538 |
| AshfExt | Rickettsiaceae | Fresh | 0.191538 |
| AshfExt | Hafniaceae | Aged | 0.178462 |
| AshfExt | Geodermatophilaceae | Aged | 0.176923 |
| AshfExt | Salinisphaeraceae | Fresh | 0.171538 |
| AshfExt | Alcaligenaceae | Fresh | 0.163846 |
| AshfExt | Bacteroidia_unclassified | Fresh | 0.146923 |
| AshfExt | Chthoniobacteraceae | Aged | 0.141538 |
| AshfExt | JG30-KF-CM45 | Aged | 0.138462 |
| AshfExt | Dermabacteraceae | Aged | 0.137692 |
| AshfExt | Bacteriovoracaceae | Aged | 0.133077 |
| AshfExt | Kineosporiaceae | Aged | 0.128462 |
| AshfExt | Lactobacillaceae | Fresh | 0.123846 |
| AshfExt | Crocinitomicaceae | Aged | 0.122308 |
| AshfExt | Micromonosporaceae | Aged | 0.122308 |
| AshfExt | Comamonadaceae | Fresh | 0.113077 |
| AshfExt | Sericytochromatia_fa | Aged | 0.103846 |
| AshfExt | Micrococcales_unclassified | Fresh | 0.086923 |
| AshfExt | Bacillales_unclassified | Aged | 0.085385 |
| AshfExt | Gammaproteobacteria_unclassified | Aged | 0.077692 |
| AshfExt | Rhizobiaceae | Fresh | 0.075385 |
| AshfExt | Acetobacteraceae | Fresh | 0.071538 |
| AshfExt | Actinobacteria_unclassified | Aged | 0.06 |
| AshfExt | Xanthomonadaceae | Fresh | 0.059231 |
| AshfExt | Bacilli_unclassified | Aged | 0.056154 |
| AshfExt | Staphylococcaceae | Aged | 0.055385 |

| | | | |
|---|---|---|---|
| AshfExt | Mycobacteriaceae | Aged | 0.050769 |
| AshfExt | Helicobacteraceae | Fresh | 0.05 |
| AshfExt | Bdellovibrionaceae | Aged | 0.045385 |
| AshfExt | Dermabacteraceae | Fresh | 0.045385 |
| AshfExt | uncultured | Fresh | 0.043846 |
| AshfExt | Fimbriimonadaceae | Aged | 0.041538 |
| AshfExt | Azospirillaceae | Aged | 0.041538 |
| AshfExt | Bacteria_unclassified | Aged | 0.039231 |
| AshfExt | Beijerinckiaceae | Fresh | 0.039231 |
| AshfExt | Bacillales_unclassified | Fresh | 0.039231 |
| AshfExt | Solirubrobacteraceae | Aged | 0.038462 |
| AshfExt | Flavobacteriales_unclassified | Fresh | 0.037692 |
| AshfExt | Lactobacillales_unclassified | Aged | 0.037692 |
| AshfExt | Gemmatimonadaceae | Aged | 0.036154 |
| AshfExt | Morganellaceae | Fresh | 0.035385 |
| AshfExt | Orbaceae | Fresh | 0.035385 |
| AshfExt | Actinobacteria_unclassified | Fresh | 0.035385 |
| AshfExt | Burkholderiales_unclassified | Aged | 0.033077 |
| AshfExt | Bogoriellaceae | Aged | 0.033077 |
| AshfExt | Kineosporiaceae | Fresh | 0.033077 |
| AshfExt | Bacillaceae | Aged | 0.032308 |
| AshfExt | Pectobacteriaceae | Fresh | 0.032308 |
| AshfExt | Orbaceae | Aged | 0.031538 |
| AshfExt | Bogoriellaceae | Fresh | 0.030769 |
| AshfExt | Rhodobacteraceae | Fresh | 0.029231 |
| AshfExt | Deinococcaceae | Fresh | 0.029231 |
| AshfExt | Bacilli_unclassified | Fresh | 0.028462 |
| AshfExt | Firmicutes_unclassified | Fresh | 0.028462 |
| AshfExt | Paenibacillaceae | Fresh | 0.028462 |
| AshfExt | Verrucomicrobiaceae | Aged | 0.026923 |
| AshfExt | Phaselicystidaceae | Aged | 0.026154 |

| | | | |
|---|---|---|---|
| AshfExt | Cellvibrionaceae | Aged | 0.025385 |
| AshfExt | Ruminococcaceae | Fresh | 0.024615 |
| AshfExt | Iamiaceae | Aged | 0.023846 |
| AshfExt | Corynebacteriales_Incertae_Sedis | Fresh | 0.023846 |
| AshfExt | Leptotrichiaceae | Fresh | 0.023846 |
| AshfExt | Nocardioidaceae | Fresh | 0.023077 |
| AshfExt | WD2101_soil_group | Aged | 0.023077 |
| AshfExt | Flavobacteriaceae | Fresh | 0.022308 |
| AshfExt | Lactobacillales_unclassified | Fresh | 0.022308 |
| AshfExt | Oligoflexaceae | Aged | 0.022308 |
| AshfExt | Lactobacillaceae | Aged | 0.020769 |
| AshfExt | Exiguobacteraceae | Fresh | 0.020769 |
| AshfExt | Isosphaeraceae | Aged | 0.02 |
| AshfExt | Xanthobacteraceae | Aged | 0.02 |
| AshfExt | Corynebacteriaceae | Aged | 0.02 |
| AshfExt | Intrasporangiaceae | Aged | 0.019231 |
| AshfExt | Erysipelatoclostridiaceae | Fresh | 0.019231 |
| AshfExt | Gemmataceae | Aged | 0.019231 |
| AshfExt | Alphaproteobacteria_unclassified | Aged | 0.018462 |
| AshfExt | Corynebacteriales_unclassified | Aged | 0.017692 |
| AshfExt | Sphingomonadaceae | Fresh | 0.016923 |
| AshfExt | Brevibacteriaceae | Fresh | 0.016923 |
| AshfExt | Rhodothermaceae | Aged | 0.016923 |
| AshfExt | Gammaproteobacteria_unclassified | Fresh | 0.016154 |
| AshfExt | Hymenobacteraceae | Fresh | 0.015385 |
| AshfExt | Tepidisphaerales_unclassified | Aged | 0.014615 |
| AshfExt | Vagococcaceae | Fresh | 0.014615 |
| AshfExt | Bacteroidia_unclassified | Aged | 0.012308 |
| AshfExt | Sumerlaeaceae | Aged | 0.012308 |
| AshfExt | uncultured | Aged | 0.012308 |
| AshfExt | Brevibacteriaceae | Aged | 0.011538 |

| | | | |
|---|---|---|---|
| AshfExt | Chitinophagaceae | Fresh | 0.011538 |
| AshfExt | Geminicoccaceae | Aged | 0.011538 |
| AshfExt | Burkholderiaceae | Aged | 0.010769 |
| AshfExt | Cryptosporangiaceae | Aged | 0.010769 |
| AshfExt | Kaistiaceae | Aged | 0.010769 |
| AshfExt | Lachnospiraceae | Aged | 0.010769 |
| AshfExt | Rickettsiaceae | Aged | 0.010769 |
| AshfExt | Streptomycetaceae | Aged | 0.010769 |
| AshfExt | AKIW781 | Aged | 0.01 |
| AshfExt | Frankiaceae | Aged | 0.01 |
| AshfExt | JG30-KF-CM45 | Fresh | 0.01 |
| AshfExt | Absconditabacteriales_(SR1)_fa | Aged | 0.01 |
| AshfExt | Coriobacteriales_Incertae_Sedis | Fresh | 0.01 |
| AshfExt | Rhizobiales_Incertae_Sedis | Aged | 0.009231 |
| AshfExt | Propionibacteriales_unclassified | Fresh | 0.009231 |
| AshfExt | Spirosomaceae | Fresh | 0.008462 |
| AshfExt | Nakamurellaceae | Aged | 0.008462 |
| AshfExt | Saccharimonadales_unclassified | Aged | 0.008462 |
| AshfExt | 0319-6G20_fa | Aged | 0.007692 |
| AshfExt | Iamiaceae | Fresh | 0.007692 |
| AshfExt | Polyangiaceae | Aged | 0.007692 |
| AshfExt | Silvanigrellaceae | Aged | 0.007692 |
| AshfExt | 67-14 | Aged | 0.006923 |
| AshfExt | Budviciaceae | Fresh | 0.006923 |
| AshfExt | Dysgonomonadaceae | Fresh | 0.006154 |
| AshfExt | Blastocatellaceae | Aged | 0.006154 |
| AshfExt | Micromonosporaceae | Fresh | 0.006154 |
| AshfExt | Pseudonocardiaceae | Fresh | 0.006154 |
| AshfExt | Trueperaceae | Fresh | 0.006154 |
| AshfExt | Pseudonocardiaceae | Aged | 0.006154 |
| AshfExt | Brevinemataceae | Fresh | 0.005385 |

| | | | |
|---|---|---|---|
| AshfExt | Frankiales_unclassified | Aged | 0.005385 |
| AshfExt | Mycobacteriaceae | Fresh | 0.005385 |
| AshfExt | Rhizobiales_unclassified | Aged | 0.005385 |
| AshfExt | Geodermatophilaceae | Fresh | 0.004615 |
| AshfExt | Pseudomonadales_unclassified | Fresh | 0.004615 |
| AshfExt | Rubritaleaceae | Aged | 0.004615 |
| AshfExt | Ilumatobacteraceae | Aged | 0.004615 |
| AshfExt | Silvanigrellaceae | Fresh | 0.004615 |
| AshfExt | Steroidobacteraceae | Aged | 0.004615 |
| AshfExt | Tepidisphaeraceae | Aged | 0.004615 |
| AshfExt | mle1-27_fa | Aged | 0.004615 |
| AshfExt | Azospirillaceae | Fresh | 0.003846 |
| AshfExt | Burkholderiales_unclassified | Fresh | 0.003846 |
| AshfExt | Erysipelotrichaceae | Fresh | 0.003846 |
| AshfExt | Saprospiraceae | Fresh | 0.003846 |
| AshfExt | Solirubrobacteraceae | Fresh | 0.003846 |
| AshfExt | Yersiniaceae | Aged | 0.003846 |
| AshfExt | Sandaracinaceae | Aged | 0.003846 |
| AshfExt | Alphaproteobacteria_unclassified | Fresh | 0.003077 |
| AshfExt | Parcubacteria_unclassified | Aged | 0.003077 |
| AshfExt | Chlamydiales_unclassified | Aged | 0.003077 |
| AshfExt | Cyanobacteriia_unclassified | Fresh | 0.003077 |
| AshfExt | Dermacoccaceae | Aged | 0.003077 |
| AshfExt | Eubacteriaceae | Fresh | 0.003077 |
| AshfExt | Microtrichaceae | Aged | 0.003077 |
| AshfExt | WPS-2_fa | Fresh | 0.003077 |
| AshfExt | 67-14 | Fresh | 0.003077 |
| AshfExt | Diplorickettsiaceae | Aged | 0.003077 |
| AshfExt | Streptomycetaceae | Fresh | 0.003077 |
| AshfExt | Anaeromyxobacteraceae | Aged | 0.002308 |
| AshfExt | Anaplasmataceae | Fresh | 0.002308 |

| | | | |
|---|---|---|---|
| AshfExt | Bacteroidaceae | Aged | 0.002308 |
| AshfExt | Burkholderiaceae | Fresh | 0.002308 |
| AshfExt | Chlamydiaceae | Fresh | 0.002308 |
| AshfExt | Clostridia_unclassified | Fresh | 0.002308 |
| AshfExt | Isosphaeraceae | Fresh | 0.002308 |
| AshfExt | Oscillospirales_unclassified | Fresh | 0.002308 |
| AshfExt | Propionibacteriales_unclassified | Aged | 0.002308 |
| AshfExt | Rhodanobacteraceae | Aged | 0.002308 |
| AshfExt | Rhodospirillales_unclassified | Aged | 0.002308 |
| AshfExt | SM2D12 | Aged | 0.002308 |
| AshfExt | Xanthobacteraceae | Fresh | 0.002308 |
| AshfExt | Yersiniaceae | Fresh | 0.002308 |
| AshfExt | A0839 | Aged | 0.002308 |
| AshfExt | Cellulomonadaceae | Aged | 0.002308 |
| AshfExt | D05-2 | Aged | 0.002308 |
| AshfExt | Microgenomatia_fa | Aged | 0.002308 |
| AshfExt | Trueperaceae | Aged | 0.002308 |
| AshfExt | Xanthomonadales_unclassified | Fresh | 0.002308 |
| AshfExt | A4b | Aged | 0.001538 |
| AshfExt | Chitinibacteraceae | Fresh | 0.001538 |
| AshfExt | Chloroflexales_unclassified | Aged | 0.001538 |
| AshfExt | Entomoplasmataceae | Aged | 0.001538 |
| AshfExt | Firmicutes_unclassified | Aged | 0.001538 |
| AshfExt | Longimicrobiaceae | Aged | 0.001538 |
| AshfExt | Pirellulaceae | Aged | 0.001538 |
| AshfExt | Thermoleophilia_unclassified | Aged | 0.001538 |
| AshfExt | Anaerovoracaceae | Fresh | 0.001538 |
| AshfExt | Blattabacteriaceae | Fresh | 0.001538 |
| AshfExt | Eggerthellaceae | Fresh | 0.001538 |
| AshfExt | Family_XI | Fresh | 0.001538 |
| AshfExt | Halomonadaceae | Fresh | 0.001538 |

| | | | |
|---|---|---|---|
| AshfExt | Intrasporangiaceae | Fresh | 0.001538 |
| AshfExt | Nocardiopsaceae | Fresh | 0.001538 |
| AshfExt | Promicromonosporaceae | Aged | 0.001538 |
| AshfExt | RF39_fa | Fresh | 0.001538 |
| AshfExt | Salinisphaeraceae | Aged | 0.001538 |
| AshfExt | Bacteroidaceae | Fresh | 0.001538 |
| AshfExt | Erysipelatoclostridiaceae | Aged | 0.001538 |
| AshfExt | Herpetosiphonaceae | Aged | 0.001538 |
| AshfExt | Pseudomonadales_unclassified | Aged | 0.001538 |
| AshfExt | Saccharimonadaceae | Aged | 0.001538 |
| AshfExt | Thermoleophilia_unclassified | Fresh | 0.001538 |
| AshfExt | Acidimicrobiia_unclassified | Aged | 0.000769 |
| AshfExt | Acidobacteriales_unclassified | Aged | 0.000769 |
| AshfExt | Armatimonadales_fa | Aged | 0.000769 |
| AshfExt | Bacteroidales_unclassified | Fresh | 0.000769 |
| AshfExt | Chloroflexi_unclassified | Fresh | 0.000769 |
| AshfExt | Chroococcidiopsaceae | Aged | 0.000769 |
| AshfExt | Cyclobacteriaceae | Aged | 0.000769 |
| AshfExt | Hyphomicrobiaceae | Fresh | 0.000769 |
| AshfExt | Hyphomonadaceae | Aged | 0.000769 |
| AshfExt | LWQ8 | Aged | 0.000769 |
| AshfExt | Labraceae | Aged | 0.000769 |
| AshfExt | Magnetospirillaceae | Fresh | 0.000769 |
| AshfExt | Marinomonadaceae | Fresh | 0.000769 |
| AshfExt | Myxococcaceae | Fresh | 0.000769 |
| AshfExt | Nitrososphaeraceae | Aged | 0.000769 |
| AshfExt | Parachlamydiaceae | Aged | 0.000769 |
| AshfExt | Pasteurellaceae | Fresh | 0.000769 |
| AshfExt | Peptostreptococcales-Tissierellales_unclassified | Aged | 0.000769 |
| AshfExt | Rickettsiales_unclassified | Aged | 0.000769 |
| AshfExt | uncultured_fa | Aged | 0.000769 |

| | | | |
|---|---|---|---|
| AshfExt | Actinobacteriota_unclassified | Aged | 0.000769 |
| AshfExt | Anaerofustaceae | Fresh | 0.000769 |
| AshfExt | Anaplasmataceae | Aged | 0.000769 |
| AshfExt | Armatimonadales_unclassified | Aged | 0.000769 |
| AshfExt | Campylobacterales_unclassified | Fresh | 0.000769 |
| AshfExt | Caulobacteraceae | Fresh | 0.000769 |
| AshfExt | Chitinibacteraceae | Aged | 0.000769 |
| AshfExt | Christensenellaceae | Fresh | 0.000769 |
| AshfExt | D05-2 | Fresh | 0.000769 |
| AshfExt | Desulfobacterales_unclassified | Fresh | 0.000769 |
| AshfExt | Desulfobacterota_unclassified | Fresh | 0.000769 |
| AshfExt | Devosiaceae | Fresh | 0.000769 |
| AshfExt | Fusobacteriaceae | Aged | 0.000769 |
| AshfExt | Ilumatobacteraceae | Fresh | 0.000769 |
| AshfExt | Microtrichales_unclassified | Aged | 0.000769 |
| AshfExt | Microtrichales_unclassified | Fresh | 0.000769 |
| AshfExt | Morganellaceae | Aged | 0.000769 |
| AshfExt | Nakamurellaceae | Fresh | 0.000769 |
| AshfExt | Oscillospiraceae | Aged | 0.000769 |
| AshfExt | Oscillospiraceae | Fresh | 0.000769 |
| AshfExt | Proteobacteria_unclassified | Aged | 0.000769 |
| AshfExt | Puniceispirillales_Incertae_Sedis | Aged | 0.000769 |
| AshfExt | Rhodospirillales_unclassified | Fresh | 0.000769 |
| AshfExt | Roseiflexaceae | Aged | 0.000769 |
| AshfExt | Rs-K70_termite_group_fa | Fresh | 0.000769 |
| AshfExt | RsaHf231_fa | Fresh | 0.000769 |
| AshfExt | Rubritaleaceae | Fresh | 0.000769 |
| AshfExt | Saccharimonadaceae | Fresh | 0.000769 |
| AshfExt | Sanguibacteraceae | Aged | 0.000769 |
| AshfExt | Simkaniaceae | Fresh | 0.000769 |
| AshfExt | Solirubrobacterales_unclassified | Fresh | 0.000769 |

| | | | |
|---|---|---|---|
| AshfExt | Thermomonosporaceae | Aged | 0.000769 |
| AshfExt | Vampirovibrionales_unclassified | Aged | 0.000769 |
| AshfExt | vadinHA49_fa | Fresh | 0.000769 |
| AshfInt | Clostridiaceae | Aged | 55.83167 |
| AshfInt | Clostridiaceae | Fresh | 47.49 |
| AshfInt | Enterobacteriaceae | Aged | 19.505 |
| AshfInt | Enterobacteriaceae | Fresh | 17.75231 |
| AshfInt | Diplorickettsiaceae | Fresh | 6.769231 |
| AshfInt | Enterococcaceae | Fresh | 4.657692 |
| AshfInt | Enterococcaceae | Aged | 3.663333 |
| AshfInt | Hafniaceae | Aged | 3.625 |
| AshfInt | Peptostreptococcaceae | Aged | 3.495 |
| AshfInt | Bacillaceae | Fresh | 3.194615 |
| AshfInt | Fusobacteriaceae | Fresh | 2.996154 |
| AshfInt | Streptococcaceae | Fresh | 2.866154 |
| AshfInt | Mycoplasmataceae | Aged | 2.616667 |
| AshfInt | Salinisphaeraceae | Aged | 2.5725 |
| AshfInt | Hafniaceae | Fresh | 2.372308 |
| AshfInt | Simkaniaceae | Fresh | 2.36 |
| AshfInt | Aeromonadaceae | Aged | 1.6075 |
| AshfInt | Mycoplasmataceae | Fresh | 1.501538 |
| AshfInt | Aeromonadaceae | Fresh | 1.308462 |
| AshfInt | Morganellaceae | Aged | 1.281667 |
| AshfInt | Bacillaceae | Aged | 1.2625 |
| AshfInt | Staphylococcaceae | Fresh | 0.875385 |
| AshfInt | Staphylococcaceae | Aged | 0.810833 |
| AshfInt | Micrococcales_unclassified | Fresh | 0.630769 |
| AshfInt | Corynebacteriales_unclassified | Fresh | 0.559231 |
| AshfInt | Salinisphaeraceae | Fresh | 0.556154 |
| AshfInt | Fusobacteriaceae | Aged | 0.5375 |
| AshfInt | Streptococcaceae | Aged | 0.506667 |

| | | | |
|---|---|---|---|
| AshfInt | Enterobacterales_unclassified | Fresh | 0.434615 |
| AshfInt | Carnobacteriaceae | Fresh | 0.360769 |
| AshfInt | Helicobacteraceae | Fresh | 0.349231 |
| AshfInt | Bacteroidia_unclassified | Fresh | 0.333846 |
| AshfInt | Enterobacterales_unclassified | Aged | 0.330833 |
| AshfInt | Rickettsiaceae | Fresh | 0.321538 |
| AshfInt | Lachnospiraceae | Aged | 0.28 |
| AshfInt | Alcaligenaceae | Fresh | 0.262308 |
| AshfInt | Orbaceae | Aged | 0.258333 |
| AshfInt | Corynebacteriales_unclassified | Aged | 0.228333 |
| AshfInt | Peptostreptococcaceae | Fresh | 0.218462 |
| AshfInt | Rhizobiaceae | Fresh | 0.197692 |
| AshfInt | Ruminococcaceae | Fresh | 0.191538 |
| AshfInt | Corynebacteriaceae | Fresh | 0.176154 |
| AshfInt | Bacteroidia_unclassified | Aged | 0.158333 |
| AshfInt | Carnobacteriaceae | Aged | 0.1175 |
| AshfInt | Alcaligenaceae | Aged | 0.108333 |
| AshfInt | Synergistaceae | Aged | 0.105833 |
| AshfInt | Ruminococcaceae | Aged | 0.101667 |
| AshfInt | Lactobacillaceae | Fresh | 0.099231 |
| AshfInt | Acetobacteraceae | Fresh | 0.096923 |
| AshfInt | Firmicutes_unclassified | Aged | 0.095833 |
| AshfInt | Micrococcaceae | Fresh | 0.080769 |
| AshfInt | Microbacteriaceae | Fresh | 0.07 |
| AshfInt | Lachnospiraceae | Fresh | 0.069231 |
| AshfInt | Comamonadaceae | Fresh | 0.063846 |
| AshfInt | Leptotrichiaceae | Aged | 0.063333 |
| AshfInt | Morganellaceae | Fresh | 0.052308 |
| AshfInt | Lactobacillaceae | Aged | 0.051667 |
| AshfInt | Bogoriellaceae | Fresh | 0.048462 |
| AshfInt | Nocardioidaceae | Fresh | 0.041538 |

| | | | |
|---|---|---|---|
| AshfInt | Mycobacteriaceae | Fresh | 0.04 |
| AshfInt | Micrococcaceae | Aged | 0.039167 |
| AshfInt | Bacteria_unclassified | Fresh | 0.038462 |
| AshfInt | Flavobacteriaceae | Aged | 0.038333 |
| AshfInt | Coriobacteriales_Incertae_Sedis | Fresh | 0.036923 |
| AshfInt | Rhizobiaceae | Aged | 0.036667 |
| AshfInt | Lactobacillales_unclassified | Fresh | 0.035385 |
| AshfInt | Bogoriellaceae | Aged | 0.033333 |
| AshfInt | Lactobacillales_unclassified | Aged | 0.033333 |
| AshfInt | Bacilli_unclassified | Aged | 0.030833 |
| AshfInt | Mycobacteriaceae | Aged | 0.029167 |
| AshfInt | Corynebacteriales_Incertae_Sedis | Fresh | 0.027692 |
| AshfInt | Corynebacteriales_Incertae_Sedis | Aged | 0.0275 |
| AshfInt | Flavobacteriales_unclassified | Fresh | 0.026923 |
| AshfInt | Firmicutes_unclassified | Fresh | 0.025385 |
| AshfInt | Coriobacteriales_Incertae_Sedis | Aged | 0.024167 |
| AshfInt | Oscillospiraceae | Aged | 0.023333 |
| AshfInt | Flavobacteriales_unclassified | Aged | 0.0225 |
| AshfInt | Beijerinckiaceae | Fresh | 0.022308 |
| AshfInt | Corynebacteriaceae | Aged | 0.021667 |
| AshfInt | Moraxellaceae | Aged | 0.021667 |
| AshfInt | Bacilli_unclassified | Fresh | 0.020769 |
| AshfInt | RsaHf231_fa | Fresh | 0.02 |
| AshfInt | Simkaniaceae | Aged | 0.02 |
| AshfInt | JG30-KF-CM45 | Fresh | 0.019231 |
| AshfInt | Microbacteriaceae | Aged | 0.019167 |
| AshfInt | Pectobacteriaceae | Fresh | 0.017692 |
| AshfInt | Trueperaceae | Fresh | 0.017692 |
| AshfInt | Bacteria_unclassified | Aged | 0.0175 |
| AshfInt | Moraxellaceae | Fresh | 0.016923 |
| AshfInt | Planococcaceae | Aged | 0.016667 |

| | | | |
|---|---|---|---|
| AshfInt | Trueperaceae | Aged | 0.016667 |
| AshfInt | Bacillales_unclassified | Fresh | 0.016154 |
| AshfInt | Flavobacteriaceae | Fresh | 0.016154 |
| AshfInt | Budviciaceae | Aged | 0.015833 |
| AshfInt | Streptomycetaceae | Fresh | 0.015385 |
| AshfInt | Nocardiaceae | Fresh | 0.014615 |
| AshfInt | Rhodobacteraceae | Fresh | 0.014615 |
| AshfInt | uncultured | Fresh | 0.013846 |
| AshfInt | Micrococcales_unclassified | Aged | 0.013333 |
| AshfInt | Oscillospirales_unclassified | Aged | 0.013333 |
| AshfInt | Erysipelotrichaceae | Fresh | 0.013077 |
| AshfInt | Desulfovibrionaceae | Aged | 0.011667 |
| AshfInt | Nocardioidaceae | Aged | 0.011667 |
| AshfInt | Sphingobacteriaceae | Aged | 0.011667 |
| AshfInt | Iamiaceae | Fresh | 0.011538 |
| AshfInt | Comamonadaceae | Aged | 0.010833 |
| AshfInt | Iamiaceae | Aged | 0.010833 |
| AshfInt | Anaplasmataceae | Fresh | 0.010769 |
| AshfInt | Oscillospirales_unclassified | Fresh | 0.01 |
| AshfInt | JG30-KF-CM45 | Aged | 0.01 |
| AshfInt | Christensenellaceae | Aged | 0.01 |
| AshfInt | Sphingobacteriaceae | Fresh | 0.009231 |
| AshfInt | uncultured | Aged | 0.009167 |
| AshfInt | Clostridia_unclassified | Fresh | 0.008462 |
| AshfInt | Pseudomonadaceae | Aged | 0.008333 |
| AshfInt | Oscillospiraceae | Fresh | 0.007692 |
| AshfInt | Dysgonomonadaceae | Aged | 0.0075 |
| AshfInt | Erysipelatoclostridiaceae | Aged | 0.0075 |
| AshfInt | Rs-K70_termite_group_fa | Aged | 0.0075 |
| AshfInt | Saprospiraceae | Aged | 0.0075 |
| AshfInt | Chitinophagaceae | Aged | 0.006667 |

| | | | |
|---|---|---|---|
| AshfInt | Clostridia_unclassified | Aged | 0.006667 |
| AshfInt | Propionibacteriales_unclassified | Aged | 0.006667 |
| AshfInt | Pasteurellaceae | Aged | 0.006667 |
| AshfInt | Rhodobacteraceae | Aged | 0.006667 |
| AshfInt | Gammaproteobacteria_unclassified | Fresh | 0.006154 |
| AshfInt | Orbaceae | Fresh | 0.006154 |
| AshfInt | Propionibacteriales_unclassified | Fresh | 0.006154 |
| AshfInt | Anaerovoracaceae | Aged | 0.005833 |
| AshfInt | Brevinemataceae | Aged | 0.005 |
| AshfInt | Erysipelotrichaceae | Aged | 0.005 |
| AshfInt | Helicobacteraceae | Aged | 0.005 |
| AshfInt | Bacillales_unclassified | Aged | 0.004167 |
| AshfInt | Planococcaceae | Fresh | 0.003846 |
| AshfInt | Saprospiraceae | Fresh | 0.003846 |
| AshfInt | Peptostreptococcales-Tissierellales_unclassified | Aged | 0.003333 |
| AshfInt | Rickettsiaceae | Aged | 0.003333 |
| AshfInt | Streptomycetaceae | Aged | 0.003333 |
| AshfInt | Clostridiales_unclassified | Aged | 0.003333 |
| AshfInt | Gammaproteobacteria_unclassified | Aged | 0.003333 |
| AshfInt | Chitinophagaceae | Fresh | 0.003077 |
| AshfInt | Pasteurellaceae | Fresh | 0.003077 |
| AshfInt | Brevinemataceae | Fresh | 0.003077 |
| AshfInt | Christensenellaceae | Fresh | 0.003077 |
| AshfInt | Nocardiopsaceae | Fresh | 0.003077 |
| AshfInt | Vagococcaceae | Fresh | 0.003077 |
| AshfInt | Acetobacteraceae | Aged | 0.0025 |
| AshfInt | Family_XI | Aged | 0.0025 |
| AshfInt | Rhodanobacteraceae | Aged | 0.0025 |
| AshfInt | Acidaminococcaceae | Aged | 0.0025 |
| AshfInt | Paenibacillaceae | Aged | 0.0025 |
| AshfInt | Pseudonocardiaceae | Aged | 0.0025 |

| | | | |
|---|---|---|---|
| AshfInt | Beijerinckiaceae | Aged | 0.0025 |
| AshfInt | Nocardiaceae | Aged | 0.0025 |
| AshfInt | Pseudomonadales_unclassified | Aged | 0.0025 |
| AshfInt | Xanthomonadales_unclassified | Aged | 0.0025 |
| AshfInt | 67-14 | Fresh | 0.002308 |
| AshfInt | Actinobacteria_unclassified | Fresh | 0.002308 |
| AshfInt | Actinobacteriota_unclassified | Fresh | 0.002308 |
| AshfInt | Anaerovoracaceae | Fresh | 0.002308 |
| AshfInt | Brevibacteriaceae | Fresh | 0.002308 |
| AshfInt | RF39_fa | Fresh | 0.002308 |
| AshfInt | Weeksellaceae | Fresh | 0.002308 |
| AshfInt | Xanthomonadaceae | Fresh | 0.002308 |
| AshfInt | Actinobacteria_unclassified | Aged | 0.001667 |
| AshfInt | Cryomorphaceae | Aged | 0.001667 |
| AshfInt | Oxalobacteraceae | Aged | 0.001667 |
| AshfInt | RF39_fa | Aged | 0.001667 |
| AshfInt | WPS-2_fa | Aged | 0.001667 |
| AshfInt | Xanthomonadaceae | Aged | 0.001667 |
| AshfInt | Bacteriovoracaceae | Aged | 0.001667 |
| AshfInt | Bacteroidaceae | Aged | 0.001667 |
| AshfInt | Bradymonadaceae | Aged | 0.001667 |
| AshfInt | Cellulomonadaceae | Aged | 0.001667 |
| AshfInt | Cyclobacteriaceae | Aged | 0.001667 |
| AshfInt | Dietziaceae | Aged | 0.001667 |
| AshfInt | Diplorickettsiaceae | Aged | 0.001667 |
| AshfInt | Nitrososphaeraceae | Aged | 0.001667 |
| AshfInt | Weeksellaceae | Aged | 0.001667 |
| AshfInt | Chitinibacteraceae | Fresh | 0.001538 |
| AshfInt | Dermabacteraceae | Fresh | 0.001538 |
| AshfInt | Dietziaceae | Fresh | 0.001538 |
| AshfInt | Exiguobacteraceae | Fresh | 0.001538 |

| | | | |
|---|---|---|---|
| AshfInt | Frankiales_unclassified | Fresh | 0.001538 |
| AshfInt | Isosphaeraceae | Fresh | 0.001538 |
| AshfInt | Phormidiaceae | Fresh | 0.001538 |
| AshfInt | Spiroplasmataceae | Fresh | 0.001538 |
| AshfInt | Spirosomaceae | Fresh | 0.001538 |
| AshfInt | Budviciaceae | Fresh | 0.001538 |
| AshfInt | Catenulisporaceae | Fresh | 0.001538 |
| AshfInt | Desulfovibrionaceae | Fresh | 0.001538 |
| AshfInt | Erysipelatoclostridiaceae | Fresh | 0.001538 |
| AshfInt | Polyangiaceae | Fresh | 0.001538 |
| AshfInt | Pseudomonadaceae | Fresh | 0.001538 |
| AshfInt | Solirubrobacteraceae | Fresh | 0.001538 |
| AshfInt | Synergistaceae | Fresh | 0.001538 |
| AshfInt | Yersiniaceae | Fresh | 0.001538 |
| AshfInt | Family_XI | Fresh | 0.001538 |
| AshfInt | Nakamurellaceae | Fresh | 0.001538 |
| AshfInt | 67-14 | Aged | 0.000833 |
| AshfInt | Actinospicaceae | Aged | 0.000833 |
| AshfInt | Brevibacteriaceae | Aged | 0.000833 |
| AshfInt | Burkholderiaceae | Aged | 0.000833 |
| AshfInt | Burkholderiales_unclassified | Aged | 0.000833 |
| AshfInt | Catenulisporaceae | Aged | 0.000833 |
| AshfInt | Caulobacteraceae | Aged | 0.000833 |
| AshfInt | Crocinitomicaceae | Aged | 0.000833 |
| AshfInt | Dermabacteraceae | Aged | 0.000833 |
| AshfInt | Desulfobacterales_unclassified | Aged | 0.000833 |
| AshfInt | Devosiaceae | Aged | 0.000833 |
| AshfInt | Eubacteriaceae | Aged | 0.000833 |
| AshfInt | Gaiellaceae | Aged | 0.000833 |
| AshfInt | Gemmataceae | Aged | 0.000833 |
| AshfInt | Isosphaeraceae | Aged | 0.000833 |

| | | | |
|---|---|---|---|
| AshfInt | Methylococcaceae | Aged | 0.000833 |
| AshfInt | Micromonosporaceae | Aged | 0.000833 |
| AshfInt | Pectobacteriaceae | Aged | 0.000833 |
| AshfInt | Polyangiaceae | Aged | 0.000833 |
| AshfInt | Rhizobiales_Incertae_Sedis | Aged | 0.000833 |
| AshfInt | Sanguibacteraceae | Aged | 0.000833 |
| AshfInt | Solirubrobacteraceae | Aged | 0.000833 |
| AshfInt | Streptosporangiaceae | Aged | 0.000833 |
| AshfInt | Thermoleophilia_unclassified | Aged | 0.000833 |
| AshfInt | vadinHA49_fa | Aged | 0.000833 |
| AshfInt | Bacteroidaceae | Fresh | 0.000769 |
| AshfInt | Bradymonadaceae | Fresh | 0.000769 |
| AshfInt | Burkholderiaceae | Fresh | 0.000769 |
| AshfInt | Chromobacteriaceae | Fresh | 0.000769 |
| AshfInt | Desulfuromonadia_unclassified | Fresh | 0.000769 |
| AshfInt | Haloferacaceae | Fresh | 0.000769 |
| AshfInt | Halomicrobiaceae | Fresh | 0.000769 |
| AshfInt | Intrasporangiaceae | Fresh | 0.000769 |
| AshfInt | Microtrichaceae | Fresh | 0.000769 |
| AshfInt | Omnitrophaceae | Fresh | 0.000769 |
| AshfInt | Pseudonocardiaceae | Fresh | 0.000769 |
| AshfInt | Rhizobiales_Incertae_Sedis | Fresh | 0.000769 |
| AshfInt | SJA-15_fa | Fresh | 0.000769 |
| AshfInt | Saccharimonadaceae | Fresh | 0.000769 |
| AshfInt | Sphingomonadaceae | Fresh | 0.000769 |
| AshfInt | Thermoleophilia_unclassified | Fresh | 0.000769 |
| AshfInt | Tsukamurellaceae | Fresh | 0.000769 |
| AshfInt | Xanthobacteraceae | Fresh | 0.000769 |
| AshfInt | Burkholderiales_unclassified | Fresh | 0.000769 |
| AshfInt | Cyanobiaceae | Fresh | 0.000769 |
| AshfInt | Hyphomicrobiaceae | Fresh | 0.000769 |

| | | | |
|---|---|---|---|
| AshfInt | Legionellaceae | Fresh | 0.000769 |
| AshfInt | Microtrichales_unclassified | Fresh | 0.000769 |
| AshfInt | Nitrosomonadaceae | Fresh | 0.000769 |
| AshfInt | Peptostreptococcales-Tissierellales_fa | Fresh | 0.000769 |
| AshfInt | Peptostreptococcales-Tissierellales_unclassified | Fresh | 0.000769 |
| AshfInt | Pseudomonadales_unclassified | Fresh | 0.000769 |
| WMCC | Clostridiaceae | Fresh | 44.74692 |
| WMCC | Nocardiaceae | Aged | 19.94625 |
| WMCC | Oxalobacteraceae | Aged | 17.84813 |
| WMCC | Enterococcaceae | Fresh | 15.10385 |
| WMCC | Sphingobacteriaceae | Aged | 9.24875 |
| WMCC | Streptococcaceae | Fresh | 8.752308 |
| WMCC | Pseudomonadaceae | Aged | 7.6875 |
| WMCC | Enterobacteriaceae | Fresh | 7.145385 |
| WMCC | Aeromonadaceae | Fresh | 6.189231 |
| WMCC | Weeksellaceae | Aged | 5.289375 |
| WMCC | Simkaniaceae | Fresh | 4.384615 |
| WMCC | Enterobacteriaceae | Aged | 3.675 |
| WMCC | Xanthomonadaceae | Aged | 3.584375 |
| WMCC | Moraxellaceae | Aged | 3.31375 |
| WMCC | Rhizobiaceae | Aged | 3.093125 |
| WMCC | Micrococcaceae | Aged | 2.735625 |
| WMCC | Microbacteriaceae | Aged | 2.675625 |
| WMCC | Mycoplasmataceae | Fresh | 2.653846 |
| WMCC | Paenibacillaceae | Aged | 2.195625 |
| WMCC | Comamonadaceae | Aged | 2.17375 |
| WMCC | Desulfovibrionaceae | Fresh | 1.772308 |
| WMCC | Hafniaceae | Fresh | 1.646154 |
| WMCC | Enterobacterales_unclassified | Aged | 1.430625 |
| WMCC | Rubritaleaceae | Aged | 1.305625 |
| WMCC | Peptostreptococcaceae | Fresh | 1.269231 |

| | | | |
|---|---|---|---|
| WMCC | Sphingomonadaceae | Aged | 1.13 |
| WMCC | Morganellaceae | Fresh | 1.064615 |
| WMCC | Solirubrobacteraceae | Aged | 0.99 |
| WMCC | Burkholderiaceae | Aged | 0.919375 |
| WMCC | Beijerinckiaceae | Aged | 0.911875 |
| WMCC | Streptomycetaceae | Aged | 0.8025 |
| WMCC | Carnobacteriaceae | Aged | 0.775625 |
| WMCC | Chitinophagaceae | Aged | 0.744375 |
| WMCC | Diplorickettsiaceae | Fresh | 0.664615 |
| WMCC | Nocardioidaceae | Aged | 0.5625 |
| WMCC | Rhodanobacteraceae | Aged | 0.55875 |
| WMCC | Alcaligenaceae | Fresh | 0.505385 |
| WMCC | Staphylococcaceae | Fresh | 0.434615 |
| WMCC | Caulobacteraceae | Aged | 0.415625 |
| WMCC | Bacillaceae | Fresh | 0.406154 |
| WMCC | Xanthobacteraceae | Aged | 0.405 |
| WMCC | Enterobacterales_unclassified | Fresh | 0.404615 |
| WMCC | Alcaligenaceae | Aged | 0.343125 |
| WMCC | Salinisphaeraceae | Fresh | 0.338462 |
| WMCC | Lachnospiraceae | Fresh | 0.333077 |
| WMCC | Spirosomaceae | Aged | 0.321875 |
| WMCC | Flavobacteriaceae | Aged | 0.319375 |
| WMCC | Enterococcaceae | Aged | 0.316875 |
| WMCC | Chitinibacteraceae | Aged | 0.286875 |
| WMCC | Micromonosporaceae | Aged | 0.286875 |
| WMCC | Micrococcales_unclassified | Aged | 0.251875 |
| WMCC | Ruminococcaceae | Fresh | 0.219231 |
| WMCC | Devosiaceae | Aged | 0.200625 |
| WMCC | Staphylococcaceae | Aged | 0.18625 |
| WMCC | Fusobacteriaceae | Fresh | 0.167692 |
| WMCC | Abditibacteriaceae | Aged | 0.16625 |

| | | | |
|---|---|---|---|
| WMCC | Clostridiaceae | Aged | 0.158125 |
| WMCC | Actinobacteria_unclassified | Aged | 0.145625 |
| WMCC | Planococcaceae | Aged | 0.140625 |
| WMCC | Acetobacteraceae | Aged | 0.138125 |
| WMCC | Mycobacteriaceae | Aged | 0.12875 |
| WMCC | Azospirillaceae | Aged | 0.128125 |
| WMCC | Hymenobacteraceae | Aged | 0.119375 |
| WMCC | Comamonadaceae | Fresh | 0.11 |
| WMCC | Pseudomonadaceae | Fresh | 0.109231 |
| WMCC | Lactobacillaceae | Fresh | 0.100769 |
| WMCC | Chthoniobacteraceae | Aged | 0.095625 |
| WMCC | uncultured | Aged | 0.09125 |
| WMCC | Kaistiaceae | Aged | 0.086875 |
| WMCC | Dietziaceae | Aged | 0.084375 |
| WMCC | Bacteria_unclassified | Aged | 0.079375 |
| WMCC | Burkholderiales_unclassified | Aged | 0.076875 |
| WMCC | Lactobacillales_unclassified | Fresh | 0.076154 |
| WMCC | Crocinitomicaceae | Aged | 0.069375 |
| WMCC | Unknown_Family | Fresh | 0.066923 |
| WMCC | Oligoflexaceae | Aged | 0.06375 |
| WMCC | Nakamurellaceae | Aged | 0.06125 |
| WMCC | Nocardiaceae | Fresh | 0.059231 |
| WMCC | Orbaceae | Fresh | 0.053846 |
| WMCC | Sphingobacteriaceae | Fresh | 0.05 |
| WMCC | Gammaproteobacteria_unclassified | Aged | 0.049375 |
| WMCC | Microbacteriaceae | Fresh | 0.049231 |
| WMCC | Phormidiaceae | Fresh | 0.048462 |
| WMCC | Beijerinckiaceae | Fresh | 0.044615 |
| WMCC | Deinococcaceae | Aged | 0.044375 |
| WMCC | Helicobacteraceae | Fresh | 0.043846 |
| WMCC | Rhizobiaceae | Fresh | 0.043077 |

| | | | |
|---|---|---|---|
| WMCC | Kineosporiaceae | Aged | 0.041875 |
| WMCC | Phaselicystidaceae | Aged | 0.041875 |
| WMCC | Silvanigrellaceae | Fresh | 0.04 |
| WMCC | Microcystaceae | Fresh | 0.039231 |
| WMCC | Frankiaceae | Aged | 0.03875 |
| WMCC | Bdellovibrionaceae | Aged | 0.0375 |
| WMCC | Exiguobacteraceae | Aged | 0.03625 |
| WMCC | Bacillales_unclassified | Fresh | 0.036154 |
| WMCC | Oxalobacteraceae | Fresh | 0.035385 |
| WMCC | Solirubrobacteraceae | Fresh | 0.035385 |
| WMCC | Geodermatophilaceae | Aged | 0.031875 |
| WMCC | Firmicutes_unclassified | Aged | 0.03125 |
| WMCC | Firmicutes_unclassified | Fresh | 0.030769 |
| WMCC | Pseudonocardiaceae | Fresh | 0.029231 |
| WMCC | Xanthomonadaceae | Fresh | 0.029231 |
| WMCC | Fimbriimonadaceae | Aged | 0.028125 |
| WMCC | Rhizobiales_unclassified | Aged | 0.0275 |
| WMCC | Lachnospiraceae | Aged | 0.02625 |
| WMCC | Bacteriovoracaceae | Aged | 0.02625 |
| WMCC | 67-14 | Aged | 0.025 |
| WMCC | Rickettsiaceae | Fresh | 0.024615 |
| WMCC | Bacteria_unclassified | Fresh | 0.023846 |
| WMCC | Brevinemataceae | Fresh | 0.023846 |
| WMCC | IMCC26256_fa | Aged | 0.023125 |
| WMCC | WD2101_soil_group | Aged | 0.0225 |
| WMCC | Pasteurellaceae | Fresh | 0.021538 |
| WMCC | Corynebacteriales_unclassified | Aged | 0.02125 |
| WMCC | Cellvibrionaceae | Aged | 0.020625 |
| WMCC | Terrimicrobiaceae | Fresh | 0.019231 |
| WMCC | Acetobacteraceae | Fresh | 0.018462 |
| WMCC | Weeksellaceae | Fresh | 0.018462 |

| | | | |
|---|---|---|---|
| WMCC | Sphingomonadaceae | Fresh | 0.018462 |
| WMCC | Isosphaeraceae | Aged | 0.018125 |
| WMCC | Gaiellaceae | Aged | 0.018125 |
| WMCC | Pirellulaceae | Aged | 0.0175 |
| WMCC | Gemmataceae | Aged | 0.0175 |
| WMCC | Micrococcaceae | Fresh | 0.016923 |
| WMCC | Blastocatellaceae | Aged | 0.016875 |
| WMCC | Tepidisphaerales_unclassified | Aged | 0.016875 |
| WMCC | Pseudonocardiaceae | Aged | 0.016875 |
| WMCC | 0319-6G20_fa | Aged | 0.01625 |
| WMCC | Moraxellaceae | Fresh | 0.016154 |
| WMCC | Ilumatobacteraceae | Aged | 0.015625 |
| WMCC | Corynebacteriaceae | Fresh | 0.015385 |
| WMCC | Gammaproteobacteria_unclassified | Fresh | 0.015385 |
| WMCC | Peptostreptococcaceae | Aged | 0.015 |
| WMCC | KD4-96_fa | Aged | 0.015 |
| WMCC | Carnobacteriaceae | Fresh | 0.014615 |
| WMCC | uncultured | Fresh | 0.014615 |
| WMCC | Myxococcaceae | Aged | 0.014375 |
| WMCC | Polyangiaceae | Aged | 0.013125 |
| WMCC | Chitinibacteraceae | Fresh | 0.013077 |
| WMCC | Eggerthellaceae | Aged | 0.0125 |
| WMCC | Chthoniobacteraceae | Fresh | 0.012308 |
| WMCC | Streptomycetaceae | Fresh | 0.012308 |
| WMCC | Bacilli_unclassified | Fresh | 0.012308 |
| WMCC | Xanthobacteraceae | Fresh | 0.011538 |
| WMCC | Micromonosporaceae | Fresh | 0.011538 |
| WMCC | Vicinamibacteraceae | Aged | 0.01125 |
| WMCC | Caulobacteraceae | Fresh | 0.010769 |
| WMCC | Planococcaceae | Fresh | 0.010769 |
| WMCC | Frankiales_unclassified | Aged | 0.010625 |

| | | | |
|---|---|---|---|
| WMCC | Intrasporangiaceae | Aged | 0.010625 |
| WMCC | Saccharimonadaceae | Aged | 0.010625 |
| WMCC | Gaiellales_unclassified | Aged | 0.01 |
| WMCC | Saccharimonadales_unclassified | Aged | 0.01 |
| WMCC | Dysgonomonadaceae | Fresh | 0.01 |
| WMCC | Bacilli_unclassified | Aged | 0.009375 |
| WMCC | Silvanigrellaceae | Aged | 0.009375 |
| WMCC | Oscillospirales_unclassified | Fresh | 0.009231 |
| WMCC | Synergistaceae | Fresh | 0.009231 |
| WMCC | Gemmatimonadaceae | Aged | 0.00875 |
| WMCC | Acidobacteriaceae_(Subgroup_1) | Aged | 0.00875 |
| WMCC | Neisseriaceae | Fresh | 0.008462 |
| WMCC | Paenibacillaceae | Fresh | 0.008462 |
| WMCC | Pirellulaceae | Fresh | 0.008462 |
| WMCC | Rhodocyclaceae | Fresh | 0.008462 |
| WMCC | Alphaproteobacteria_unclassified | Aged | 0.008125 |
| WMCC | Sericytochromatia_fa | Aged | 0.008125 |
| WMCC | Burkholderiaceae | Fresh | 0.007692 |
| WMCC | Erysipelotrichaceae | Fresh | 0.007692 |
| WMCC | Nocardioidaceae | Fresh | 0.007692 |
| WMCC | Hyphomicrobiaceae | Aged | 0.0075 |
| WMCC | Halomonadaceae | Fresh | 0.006923 |
| WMCC | Rhizobiales_Incertae_Sedis | Fresh | 0.006923 |
| WMCC | Nitrososphaeraceae | Aged | 0.006875 |
| WMCC | Parachlamydiaceae | Aged | 0.006875 |
| WMCC | Bacillaceae | Aged | 0.00625 |
| WMCC | Bacteroidia_unclassified | Aged | 0.00625 |
| WMCC | Oscillospiraceae | Aged | 0.00625 |
| WMCC | Tepidisphaeraceae | Aged | 0.00625 |
| WMCC | Streptococcaceae | Aged | 0.00625 |
| WMCC | Micropepsaceae | Aged | 0.00625 |

| | | | |
|---|---|---|---|
| WMCC | Microtrichales_unclassified | Aged | 0.00625 |
| WMCC | SC-I-84 | Aged | 0.00625 |
| WMCC | TK10_fa | Aged | 0.00625 |
| WMCC | Rhodanobacteraceae | Fresh | 0.006154 |
| WMCC | Cyanobiaceae | Fresh | 0.006154 |
| WMCC | Erysipelatoclostridiaceae | Fresh | 0.006154 |
| WMCC | Spiroplasmataceae | Fresh | 0.006154 |
| WMCC | Reyranellaceae | Aged | 0.005625 |
| WMCC | Rhodobacteraceae | Aged | 0.005625 |
| WMCC | Legionellaceae | Aged | 0.005625 |
| WMCC | Coriobacteriales_Incertae_Sedis | Fresh | 0.005385 |
| WMCC | Alphaproteobacteria_unclassified | Fresh | 0.005385 |
| WMCC | Flavobacteriaceae | Fresh | 0.005385 |
| WMCC | Microtrichaceae | Fresh | 0.005385 |
| WMCC | Rubritaleaceae | Fresh | 0.005385 |
| WMCC | Chlamydiales_unclassified | Aged | 0.005 |
| WMCC | Cryptosporangiaceae | Aged | 0.005 |
| WMCC | Hafniaceae | Aged | 0.005 |
| WMCC | Vampirovibrionaceae | Aged | 0.005 |
| WMCC | Bacillales_unclassified | Aged | 0.005 |
| WMCC | JG30-KF-CM45 | Aged | 0.005 |
| WMCC | Nitrosomonadaceae | Aged | 0.005 |
| WMCC | Ilumatobacteraceae | Fresh | 0.004615 |
| WMCC | Cyanobacteriia_unclassified | Fresh | 0.004615 |
| WMCC | Leptotrichiaceae | Fresh | 0.004615 |
| WMCC | KD4-96_fa | Fresh | 0.004615 |
| WMCC | BIrii41 | Aged | 0.004375 |
| WMCC | Unknown_Family | Aged | 0.004375 |
| WMCC | Verrucomicrobiaceae | Aged | 0.004375 |
| WMCC | Corynebacteriales_Incertae_Sedis | Aged | 0.004375 |
| WMCC | Paracaedibacteraceae | Aged | 0.004375 |

| | | | |
|---|---|---|---|
| WMCC | Rhizobiales_Incertae_Sedis | Aged | 0.004375 |
| WMCC | Sanguibacteraceae | Aged | 0.004375 |
| WMCC | Corynebacteriales_Incertae_Sedis | Fresh | 0.003846 |
| WMCC | Frankiaceae | Fresh | 0.003846 |
| WMCC | Hungateiclostridiaceae | Fresh | 0.003846 |
| WMCC | Nakamurellaceae | Fresh | 0.003846 |
| WMCC | RsaHf231_fa | Fresh | 0.003846 |
| WMCC | Bacteriovoracaceae | Fresh | 0.003846 |
| WMCC | Burkholderiales_unclassified | Fresh | 0.003846 |
| WMCC | Chitinophagaceae | Fresh | 0.003846 |
| WMCC | Chromobacteriaceae | Fresh | 0.003846 |
| WMCC | Gemmataceae | Fresh | 0.003846 |
| WMCC | Bryobacteraceae | Aged | 0.00375 |
| WMCC | SM2D12 | Aged | 0.00375 |
| WMCC | Thermoleophilia_unclassified | Aged | 0.00375 |
| WMCC | Chloroflexi_unclassified | Aged | 0.00375 |
| WMCC | Xanthomonadales_unclassified | Aged | 0.00375 |
| WMCC | Cellulomonadaceae | Aged | 0.00375 |
| WMCC | MB-A2-108_fa | Aged | 0.00375 |
| WMCC | Pedosphaeraceae | Aged | 0.00375 |
| WMCC | Neisseriaceae | Aged | 0.003125 |
| WMCC | Acidimicrobiia_unclassified | Aged | 0.003125 |
| WMCC | Ruminococcaceae | Aged | 0.003125 |
| WMCC | env.OPS_17 | Aged | 0.003125 |
| WMCC | Koribacteraceae | Aged | 0.003125 |
| WMCC | Labraceae | Aged | 0.003125 |
| WMCC | Microtrichaceae | Aged | 0.003125 |
| WMCC | Pyrinomonadaceae | Aged | 0.003125 |
| WMCC | Roseiflexaceae | Aged | 0.003125 |
| WMCC | Sandaracinaceae | Aged | 0.003125 |
| WMCC | TRA3-20 | Aged | 0.003125 |

| | | | |
|---|---|---|---|
| WMCC | Kineosporiaceae | Fresh | 0.003077 |
| WMCC | Mycobacteriaceae | Fresh | 0.003077 |
| WMCC | Gaiellaceae | Fresh | 0.003077 |
| WMCC | Methanobacteriaceae | Fresh | 0.003077 |
| WMCC | Rhizobiales_unclassified | Fresh | 0.003077 |
| WMCC | Armatimonadales_fa | Aged | 0.0025 |
| WMCC | Promicromonosporaceae | Aged | 0.0025 |
| WMCC | Rickettsiaceae | Aged | 0.0025 |
| WMCC | Acidothermaceae | Aged | 0.0025 |
| WMCC | Dermabacteraceae | Aged | 0.0025 |
| WMCC | Diplorickettsiaceae | Aged | 0.0025 |
| WMCC | Lactobacillales_unclassified | Aged | 0.0025 |
| WMCC | Longimicrobiaceae | Aged | 0.0025 |
| WMCC | Opitutaceae | Aged | 0.0025 |
| WMCC | RF39_fa | Aged | 0.0025 |
| WMCC | Solibacteraceae | Aged | 0.0025 |
| WMCC | Actinobacteria_unclassified | Fresh | 0.002308 |
| WMCC | Anaerovoracaceae | Fresh | 0.002308 |
| WMCC | Clostridia_unclassified | Fresh | 0.002308 |
| WMCC | HOC36_fa | Fresh | 0.002308 |
| WMCC | Isosphaeraceae | Fresh | 0.002308 |
| WMCC | Nitrosomonadaceae | Fresh | 0.002308 |
| WMCC | 67-14 | Fresh | 0.002308 |
| WMCC | Frankiales_unclassified | Fresh | 0.002308 |
| WMCC | SJA-15_fa | Fresh | 0.002308 |
| WMCC | Sutterellaceae | Fresh | 0.002308 |
| WMCC | Eggerthellaceae | Fresh | 0.002308 |
| WMCC | AKIW781 | Aged | 0.001875 |
| WMCC | Propionibacteriales_unclassified | Aged | 0.001875 |
| WMCC | Sporichthyaceae | Aged | 0.001875 |
| WMCC | Subgroup_7_fa | Aged | 0.001875 |

| | | | |
|---|---|---|---|
| WMCC | uncultured_fa | Aged | 0.001875 |
| WMCC | Micavibrionaceae | Aged | 0.001875 |
| WMCC | Rhodomicrobiaceae | Aged | 0.001875 |
| WMCC | Steroidobacteraceae | Aged | 0.001875 |
| WMCC | Bacteroidia_unclassified | Fresh | 0.001538 |
| WMCC | MB-A2-108_fa | Fresh | 0.001538 |
| WMCC | Oscillospiraceae | Fresh | 0.001538 |
| WMCC | PeM15_fa | Fresh | 0.001538 |
| WMCC | Pectobacteriaceae | Fresh | 0.001538 |
| WMCC | SC-I-84 | Fresh | 0.001538 |
| WMCC | Spirosomaceae | Fresh | 0.001538 |
| WMCC | TK10_fa | Fresh | 0.001538 |
| WMCC | Bacteroidales_unclassified | Fresh | 0.001538 |
| WMCC | Geodermatophilaceae | Fresh | 0.001538 |
| WMCC | IMCC26256_fa | Fresh | 0.001538 |
| WMCC | JG30-KF-CM45 | Fresh | 0.001538 |
| WMCC | Oligoflexaceae | Fresh | 0.001538 |
| WMCC | Rhodobacteraceae | Fresh | 0.001538 |
| WMCC | Thermoleophilia_unclassified | Fresh | 0.001538 |
| WMCC | Legionellaceae | Fresh | 0.001538 |
| WMCC | Actinobacteriota_unclassified | Aged | 0.00125 |
| WMCC | Brevibacteriaceae | Aged | 0.00125 |
| WMCC | Chthoniobacterales_unclassified | Aged | 0.00125 |
| WMCC | Coriobacteriales_Incertae_Sedis | Aged | 0.00125 |
| WMCC | Cyanobacteriia_unclassified | Aged | 0.00125 |
| WMCC | Cytophagaceae | Aged | 0.00125 |
| WMCC | Gimesiaceae | Aged | 0.00125 |
| WMCC | Haliangiaceae | Aged | 0.00125 |
| WMCC | Hungateiclostridiaceae | Aged | 0.00125 |
| WMCC | Iamiaceae | Aged | 0.00125 |
| WMCC | LWQ8 | Aged | 0.00125 |

| | | | |
|---|---|---|---|
| WMCC | PLTA13_fa | Aged | 0.00125 |
| WMCC | Parcubacteria_unclassified | Aged | 0.00125 |
| WMCC | Proteobacteria_unclassified | Aged | 0.00125 |
| WMCC | Puniceicoccaceae | Aged | 0.00125 |
| WMCC | R7C24_fa | Aged | 0.00125 |
| WMCC | RBG-13-54-9_fa | Aged | 0.00125 |
| WMCC | S0134_terrestrial_group_fa | Aged | 0.00125 |
| WMCC | Solirubrobacterales_unclassified | Aged | 0.00125 |
| WMCC | Subgroup_2_fa | Aged | 0.00125 |
| WMCC | Thermoactinomycetaceae | Aged | 0.00125 |
| WMCC | Vicinamibacterales_unclassified | Aged | 0.00125 |
| WMCC | mle1-27_fa | Aged | 0.00125 |
| WMCC | 0319-6G20_fa | Fresh | 0.000769 |
| WMCC | A4b | Fresh | 0.000769 |
| WMCC | Alicyclobacillaceae | Fresh | 0.000769 |
| WMCC | Anaerolineaceae | Fresh | 0.000769 |
| WMCC | Anaerolineae_unclassified | Fresh | 0.000769 |
| WMCC | BIrii41 | Fresh | 0.000769 |
| WMCC | Bryobacteraceae | Fresh | 0.000769 |
| WMCC | C0119_fa | Fresh | 0.000769 |
| WMCC | CPR2_fa | Fresh | 0.000769 |
| WMCC | Caldilineaceae | Fresh | 0.000769 |
| WMCC | Caloramatoraceae | Fresh | 0.000769 |
| WMCC | Christensenellaceae | Fresh | 0.000769 |
| WMCC | DS-100_fa | Fresh | 0.000769 |
| WMCC | Desulfitobacteriaceae | Fresh | 0.000769 |
| WMCC | Devosiaceae | Fresh | 0.000769 |
| WMCC | Fimbriimonadaceae | Fresh | 0.000769 |
| WMCC | Gaiellales_unclassified | Fresh | 0.000769 |
| WMCC | Gemmatimonadaceae | Fresh | 0.000769 |
| WMCC | Geobacteraceae | Fresh | 0.000769 |

| | | | |
|---|---|---|---|
| WMCC | Halomicrobiaceae | Fresh | 0.000769 |
| WMCC | Hymenobacteraceae | Fresh | 0.000769 |
| WMCC | Hyphomicrobiaceae | Fresh | 0.000769 |
| WMCC | Iamiaceae | Fresh | 0.000769 |
| WMCC | Intrasporangiaceae | Fresh | 0.000769 |
| WMCC | Ktedonobacteraceae | Fresh | 0.000769 |
| WMCC | Methylococcaceae | Fresh | 0.000769 |
| WMCC | Methyloligellaceae | Fresh | 0.000769 |
| WMCC | Methylomonadaceae | Fresh | 0.000769 |
| WMCC | Micropepsaceae | Fresh | 0.000769 |
| WMCC | Nitrososphaeraceae | Fresh | 0.000769 |
| WMCC | Paracaedibacteraceae | Fresh | 0.000769 |
| WMCC | Polyangiaceae | Fresh | 0.000769 |
| WMCC | Pyrinomonadaceae | Fresh | 0.000769 |
| WMCC | Rice_Cluster_II | Fresh | 0.000769 |
| WMCC | Saccharimonadales_unclassified | Fresh | 0.000769 |
| WMCC | Sporomusaceae | Fresh | 0.000769 |
| WMCC | Subgroup_7_fa | Fresh | 0.000769 |
| WMCC | Syntrophorhabdaceae | Fresh | 0.000769 |
| WMCC | TRA3-20 | Fresh | 0.000769 |
| WMCC | Tepidisphaerales_unclassified | Fresh | 0.000769 |
| WMCC | Verrucomicrobiae_unclassified | Fresh | 0.000769 |
| WMCC | Vicinamibacteraceae | Fresh | 0.000769 |
| WMCC | Xiphinematobacteraceae | Fresh | 0.000769 |
| WMCC | Anaplasmataceae | Fresh | 0.000769 |
| WMCC | Bifidobacteriaceae | Fresh | 0.000769 |
| WMCC | Chlamydiaceae | Fresh | 0.000769 |
| WMCC | Corynebacteriales_unclassified | Fresh | 0.000769 |
| WMCC | Nostocaceae | Fresh | 0.000769 |
| WMCC | Thermoactinomycetaceae | Fresh | 0.000769 |
| WMCC | A4b | Aged | 0.000625 |

| | | | |
|---|---|---|---|
| WMCC | Actinomycetaceae | Aged | 0.000625 |
| WMCC | Aeromonadaceae | Aged | 0.000625 |
| WMCC | Alicyclobacillaceae | Aged | 0.000625 |
| WMCC | Anaeromyxobacteraceae | Aged | 0.000625 |
| WMCC | Babeliaceae | Aged | 0.000625 |
| WMCC | Babeliales_unclassified | Aged | 0.000625 |
| WMCC | Bacteroidaceae | Aged | 0.000625 |
| WMCC | Butyricicoccaceae | Aged | 0.000625 |
| WMCC | C0119_fa | Aged | 0.000625 |
| WMCC | CCD24_fa | Aged | 0.000625 |
| WMCC | Caldilineaceae | Aged | 0.000625 |
| WMCC | Clostridia_UCG-014_fa | Aged | 0.000625 |
| WMCC | Clostridia_unclassified | Aged | 0.000625 |
| WMCC | Coxiellaceae | Aged | 0.000625 |
| WMCC | DS-100_fa | Aged | 0.000625 |
| WMCC | Dermacoccaceae | Aged | 0.000625 |
| WMCC | Dongiaceae | Aged | 0.000625 |
| WMCC | Hyphomonadaceae | Aged | 0.000625 |
| WMCC | Kapabacteriales_fa | Aged | 0.000625 |
| WMCC | Lactobacillaceae | Aged | 0.000625 |
| WMCC | Methylacidiphilaceae | Aged | 0.000625 |
| WMCC | Microscillaceae | Aged | 0.000625 |
| WMCC | Morganellaceae | Aged | 0.000625 |
| WMCC | Nitrospiraceae | Aged | 0.000625 |
| WMCC | Polyangia_unclassified | Aged | 0.000625 |
| WMCC | Procabacteriaceae | Aged | 0.000625 |
| WMCC | Pseudomonadales_unclassified | Aged | 0.000625 |
| WMCC | Rhodospirillaceae | Aged | 0.000625 |
| WMCC | Rickettsiales_unclassified | Aged | 0.000625 |
| WMCC | Rubinisphaeraceae | Aged | 0.000625 |
| WMCC | Saccharimonadales_fa | Aged | 0.000625 |

| | | | |
|---|---|---|---|
| WMCC | Sporomusaceae | Aged | 0.000625 |
| WMCC | Subgroup_17_fa | Aged | 0.000625 |
| WMCC | Sumerlaeaceae | Aged | 0.000625 |
| WMCC | Thermomonosporaceae | Aged | 0.000625 |
| WMCC | Vermiphilaceae | Aged | 0.000625 |
| WMCC | Verrucomicrobiae_unclassified | Aged | 0.000625 |
| WMCC | WX65 | Aged | 0.000625 |
| WMCC | Xiphinematobacteraceae | Aged | 0.000625 |
