## Supplementary material for "Microbial, dietary insect, and pathogen communities in fresh and decomposing guano of anthropic little brown bat (Myotis lucifugus) maternity colonies": Table S9

| Age | Site | Family | Specificity | Fidelity | Stat | P |
|---|---|---|---|---|---|---|
| Aged | ASHFInt | Salinisphaeraceae | 0.9995 | 1 | 1 | 0.001 |
| Aged | ASHFInt | Clostridiaceae | 0.9936 | 1 | 0.997 | 0.001 |
| Aged | ASHFInt | Bacillaceae | 0.9711 | 1 | 0.985 | 0.001 |
| Aged | ASHFInt | Peptostreptococcaceae | 0.9178 | 1 | 0.958 | 0.001 |
| Aged | ASHFInt | Lachnospiraceae | 0.8854 | 1 | 0.941 | 0.001 |
| Aged | ASHFInt | Mycoplasmataceae | 0.9995 | 0.8333 | 0.913 | 0.001 |
| Aged | ASHFInt | Bacteroidia_unclassified | 0.9035 | 0.75 | 0.823 | 0.001 |
| Aged | ASHFInt | Fusobacteriaceae | 0.9988 | 0.6667 | 0.816 | 0.001 |
| Aged | ASHFInt | Morganellaceae | 0.9959 | 0.6667 | 0.815 | 0.001 |
| Aged | ASHFInt | Hafniaceae | 0.9578 | 0.6667 | 0.799 | 0.002 |
| Aged | ASHFInt | Corynebacteriales_Incertae_Sedis | 0.8627 | 0.5833 | 0.709 | 0.001 |
| Aged | ASHFInt | Leptotrichiaceae | 1 | 0.5 | 0.707 | 0.001 |
| Aged | ASHFInt | Aeromonadaceae | 0.9996 | 0.5 | 0.707 | 0.001 |
| Aged | ASHFInt | Firmicutes_unclassified | 0.7386 | 0.6667 | 0.702 | 0.027 |
| Aged | ASHFInt | Ruminococcaceae | 0.964 | 0.5 | 0.694 | 0.003 |
| Aged | ASHFInt | Oscillospiraceae | 0.7713 | 0.5833 | 0.671 | 0.003 |
| Aged | ASHFInt | Brevinemataceae | 1 | 0.4167 | 0.645 | 0.002 |
| Aged | ASHFInt | Christensenellaceae | 1 | 0.4167 | 0.645 | 0.002 |
| Aged | ASHFInt | Erysipelotrichaceae | 1 | 0.4167 | 0.645 | 0.003 |
| Aged | ASHFInt | Flavobacteriales_unclassified | 1 | 0.4167 | 0.645 | 0.001 |
| Aged | ASHFInt | Oscillospirales_unclassified | 1 | 0.4167 | 0.645 | 0.001 |
| Aged | ASHFInt | Synergistaceae | 1 | 0.4167 | 0.645 | 0.001 |
| Aged | ASHFInt | Coriobacteriales_Incertae_Sedis | 0.9508 | 0.4167 | 0.629 | 0.002 |
| Aged | ASHFInt | Clostridia_unclassified | 0.9143 | 0.4167 | 0.617 | 0.004 |
| Aged | ASHFInt | Trueperaceae | 0.8929 | 0.4167 | 0.61 | 0.01 |
| Aged | ASHFInt | Anaerovoracaceae | 1 | 0.3333 | 0.577 | 0.005 |
| Aged | ASHFInt | Budviciaceae | 1 | 0.3333 | 0.577 | 0.006 |
| Aged | ASHFInt | Simkaniaceae | 1 | 0.3333 | 0.577 | 0.004 |
| Aged | ASHFInt | Clostridiales_unclassified | 1 | 0.25 | 0.5 | 0.015 |
| Aged | ASHFInt | Desulfovibrionaceae | 1 | 0.25 | 0.5 | 0.023 |
| Aged | ASHFInt | Pasteurellaceae | 1 | 0.25 | 0.5 | 0.017 |

| | | | | | | |
|---|---|---|---|---|---|---|
| Aged | ASHFInt | Rs-K70_termite_group_fa | 1 | 0.25 | 0.5 | 0.023 |
| Aged | ASHFInt | Saprospiraceae | 1 | 0.25 | 0.5 | 0.022 |
| Aged | ASHFExt | Dietziaceae | 0.9528 | 1 | 0.976 | 0.001 |
| Aged | ASHFExt | Planococcaceae | 0.9415 | 1 | 0.97 | 0.001 |
| Aged | ASHFExt | Deinococcaceae | 0.9389 | 1 | 0.969 | 0.001 |
| Aged | ASHFExt | Rhodobacteraceae | 0.9799 | 0.9333 | 0.956 | 0.001 |
| Aged | ASHFExt | Dermabacteraceae | 0.9793 | 0.9333 | 0.956 | 0.001 |
| Aged | ASHFExt | Exiguobacteraceae | 0.8645 | 1 | 0.93 | 0.001 |
| Aged | ASHFExt | Bacillales_unclassified | 0.916 | 0.9333 | 0.925 | 0.001 |
| Aged | ASHFExt | Geodermatophilaceae | 0.8315 | 0.8 | 0.816 | 0.001 |
| Aged | ASHFExt | JG30-KF-CM45 | 0.89 | 0.7333 | 0.808 | 0.005 |
| Aged | ASHFExt | Myxococcaceae | 0.9833 | 0.5333 | 0.724 | 0.006 |
| Aged | ASHFExt | Brevibacteriaceae | 0.9057 | 0.5333 | 0.695 | 0.004 |
| Aged | ASHFExt | Gemmatimonadaceae | 0.7817 | 0.6 | 0.685 | 0.005 |
| Aged | ASHFExt | Sericytochromatia_fa | 0.9172 | 0.4667 | 0.654 | 0.009 |
| Aged | ASHFExt | Geminicoccaceae | 1 | 0.4 | 0.632 | 0.002 |
| Aged | ASHFExt | Absconditabacteriales_(SR1)_fa | 1 | 0.3333 | 0.577 | 0.005 |
| Aged | ASHFExt | Rhodothermaceae | 1 | 0.3333 | 0.577 | 0.007 |
| Aged | ASHFExt | Intrasporangiaceae | 0.6107 | 0.5333 | 0.571 | 0.041 |
| Aged | ASHFExt | D05-2 | 1 | 0.2667 | 0.516 | 0.014 |
| Aged | ASHFExt | Sumerlaeaceae | 0.9446 | 0.2667 | 0.502 | 0.044 |
| Aged | WMCC | Chitinibacteraceae | 0.9977 | 1 | 0.999 | 0.001 |
| Aged | WMCC | Rhodanobacteraceae | 0.992 | 1 | 0.996 | 0.001 |
| Aged | WMCC | Solirubrobacteraceae | 0.9654 | 1 | 0.983 | 0.001 |
| Aged | WMCC | Burkholderiaceae | 0.9876 | 0.9375 | 0.962 | 0.001 |
| Aged | WMCC | Xanthobacteraceae | 0.959 | 0.9375 | 0.948 | 0.001 |
| Aged | WMCC | Streptomycetaceae | 0.9845 | 0.8125 | 0.894 | 0.001 |
| Aged | WMCC | Kaistiaceae | 0.903 | 0.75 | 0.823 | 0.001 |
| Aged | WMCC | Rubritaleaceae | 0.9969 | 0.5625 | 0.749 | 0.004 |
| Aged | WMCC | Saccharimonadaceae | 0.8885 | 0.5 | 0.667 | 0.001 |
| Aged | WMCC | Rhizobiales_unclassified | 0.8549 | 0.5 | 0.654 | 0.018 |

| | | | | | | |
|---|---|---|---|---|---|---|
| Aged | WMCC | Paracaedibacteraceae | 1 | 0.375 | 0.612 | 0.002 |
| Aged | WMCC | Parachlamydiaceae | 0.9116 | 0.3125 | 0.534 | 0.02 |
| Aged | WMCC | Acidobacteriaceae_(Subgroup_1) | 1 | 0.25 | 0.5 | 0.027 |
| Aged | WMCC | Vampirovibrionaceae | 1 | 0.25 | 0.5 | 0.028 |
| Aged | ASHF | Corynebacteriales_unclassified | 0.9216 | 0.9259 | 0.924 | 0.008 |
| Aged | ASHF | Streptococcaceae | 0.995 | 0.7778 | 0.88 | 0.004 |
| Aged | ASHF | Lactobacillales_unclassified | 0.9642 | 0.7037 | 0.824 | 0.001 |
| Aged | ASHF | Bacilli_unclassified | 0.9071 | 0.7407 | 0.82 | 0.026 |
| Aged | ASHF | Bogoriellaceae | 1 | 0.6667 | 0.816 | 0.001 |
| Aged | ASHF | Corynebacteriaceae | 1 | 0.6296 | 0.793 | 0.002 |
| Aged | Ext | Hymenobacteraceae | 1 | 1 | 1 | 0.001 |
| Aged | Ext | Sphingomonadaceae | 1 | 1 | 1 | 0.001 |
| Aged | Ext | Spirosomaceae | 1 | 1 | 1 | 0.001 |
| Aged | Ext | Oxalobacteraceae | 0.9999 | 1 | 1 | 0.001 |
| Aged | Ext | Nocardiaceae | 0.9999 | 1 | 1 | 0.001 |
| Aged | Ext | Weeksellaceae | 0.9998 | 1 | 1 | 0.001 |
| Aged | Ext | Xanthomonadaceae | 0.9996 | 1 | 1 | 0.001 |
| Aged | Ext | Moraxellaceae | 0.9994 | 1 | 1 | 0.001 |
| Aged | Ext | Pseudomonadaceae | 0.9993 | 1 | 1 | 0.001 |
| Aged | Ext | Sphingobacteriaceae | 0.9992 | 1 | 1 | 0.001 |
| Aged | Ext | Paenibacillaceae | 0.9991 | 1 | 1 | 0.001 |
| Aged | Ext | Beijerinckiaceae | 0.998 | 1 | 0.999 | 0.001 |
| Aged | Ext | Comamonadaceae | 0.9977 | 1 | 0.999 | 0.001 |
| Aged | Ext | Microbacteriaceae | 0.995 | 1 | 0.998 | 0.001 |
| Aged | Ext | Rhizobiaceae | 0.9904 | 1 | 0.995 | 0.001 |
| Aged | Ext | Nocardioidaceae | 0.9853 | 1 | 0.993 | 0.001 |
| Aged | Ext | Micrococcales_unclassified | 0.9849 | 1 | 0.992 | 0.001 |
| Aged | Ext | Caulobacteraceae | 0.9987 | 0.9677 | 0.983 | 0.001 |
| Aged | Ext | Acetobacteraceae | 0.9941 | 0.9677 | 0.981 | 0.001 |
| Aged | Ext | Actinobacteria_unclassified | 0.9926 | 0.9355 | 0.964 | 0.001 |
| Aged | Ext | Burkholderiales_unclassified | 0.9924 | 0.9355 | 0.964 | 0.001 |

| | | | | | | |
|---|---|---|---|---|---|---|
| Aged | Ext | Gammaproteobacteria_unclassified | 0.975 | 0.9355 | 0.955 | 0.001 |
| Aged | Ext | Chitinophagaceae | 0.9961 | 0.9032 | 0.949 | 0.001 |
| Aged | Ext | Kineosporiaceae | 1 | 0.871 | 0.933 | 0.001 |
| Aged | Ext | Micromonosporaceae | 0.9979 | 0.871 | 0.932 | 0.001 |
| Aged | Ext | Devosiaceae | 0.9985 | 0.8065 | 0.897 | 0.001 |
| Aged | Ext | Abditibacteriaceae | 1 | 0.7097 | 0.842 | 0.001 |
| Aged | Ext | Flavobacteriaceae | 0.9491 | 0.7419 | 0.839 | 0.036 |
| Aged | Ext | Azospirillaceae | 1 | 0.6774 | 0.823 | 0.001 |
| Aged | Ext | Fimbriimonadaceae | 1 | 0.5161 | 0.718 | 0.002 |
| Aged | Ext | Oligoflexaceae | 1 | 0.4839 | 0.696 | 0.012 |
| Aged | Ext | Phaselicystidaceae | 1 | 0.4839 | 0.696 | 0.01 |
| Aged | Ext | Bdellovibrionaceae | 1 | 0.4516 | 0.672 | 0.014 |
| Aged | Ext | Chthoniobacteraceae | 1 | 0.4516 | 0.672 | 0.03 |
| Aged | Ext | Frankiaceae | 1 | 0.4516 | 0.672 | 0.021 |
| Aged | Ext | Nakamurellaceae | 1 | 0.4516 | 0.672 | 0.021 |
| Aged | Ext | Saccharimonadales_unclassified | 1 | 0.4516 | 0.672 | 0.01 |
| Aged | Ext | Alphaproteobacteria_unclassified | 1 | 0.4194 | 0.648 | 0.024 |
| Aged | Ext | Tepidisphaerales_unclassified | 1 | 0.4194 | 0.648 | 0.019 |
| Aged | Ext | 0319-6G20_fa | 1 | 0.3871 | 0.622 | 0.03 |
| Aged | Ext | Cryptosporangiaceae | 1 | 0.3871 | 0.622 | 0.031 |
| Aged | Ext | Silvanigrellaceae | 1 | 0.3871 | 0.622 | 0.02 |
| Fresh | ASHFInt | Micrococcales_unclassified | 0.9624 | 0.8333 | 0.896 | 0.001 |
| Fresh | WMCC | Cyanobiaceae | 0.8807 | 0.4615 | 0.638 | 0.011 |
| Fresh | WMCC | Pirellulaceae | 1 | 0.3846 | 0.62 | 0.009 |
| Fresh | WMCC | Phormidiaceae | 0.9668 | 0.3846 | 0.61 | 0.013 |
| Fresh | WMCC | Burkholderiaceae | 0.8219 | 0.3846 | 0.562 | 0.04 |
| Fresh | WMCC | Microcystaceae | 1 | 0.3077 | 0.555 | 0.026 |
| Fresh | ASHF | Carnobacteriaceae | 0.976 | 1 | 0.988 | 0.001 |
| Fresh | ASHF | Corynebacteriales_unclassified | 0.9991 | 0.8333 | 0.912 | 0.001 |
| Fresh | ASHF | Corynebacteriaceae | 0.9816 | 0.7917 | 0.882 | 0.029 |
| Fresh | ASHF | Bogoriellaceae | 1 | 0.625 | 0.791 | 0.004 |

| | | | | | | |
|---|---|---|---|---|---|---|
| Fresh | ASHF | Corynebacteriales_Incertae_Sedis | 0.9356 | 0.625 | 0.765 | 0.007 |
| Fresh | ASHF | Bacteroidia_unclassified | 0.9971 | 0.5833 | 0.763 | 0.006 |
| Fresh | Ext | Lachnospiraceae | 0.933 | 0.84 | 0.885 | 0.011 |
| Fresh | Ext | Aeromonadaceae | 0.9759 | 0.76 | 0.861 | 0.016 |
