## Supplementary material for "Microbial, dietary insect, and pathogen communities in fresh and decomposing guano of anthropic little brown bat (Myotis lucifugus) maternity colonies": Table S10

| Sample | Dups | GC | Seqs | SeqCount | | R1 Avg | R2 Avg |
|---|---|---|---|---|---|---|---|
| 2505AshExt6aFung_S308_L001_R1_001 | 88.68077001 | 52 | 0.034649 | 34649 | | 29548.40351 | 29548.40351 |
| 2505AshExt6aFung_S308_L001_R2_001 | 87.03570089 | 51 | 0.034649 | 34649 | | | |
| 2505AshExt6bFung_S298_L001_R1_001 | 86.67332446 | 54 | 0.009012 | 9012 | | | |
| 2505AshExt6bFung_S298_L001_R2_001 | 78.1846427 | 54 | 0.009012 | 9012 | | | |
| 2505AshExt6cFung_S309_L001_R1_001 | 79.25311203 | 51 | 0.00482 | 4820 | | | |
| 2505AshExt6cFung_S309_L001_R2_001 | 72.94605809 | 50 | 0.00482 | 4820 | | | |
| 2505AshInt6aFung_S305_L001_R1_001 | 86.5580766 | 44 | 0.019216 | 19216 | | | |
| 2505AshInt6aFung_S305_L001_R2_001 | 85.90237302 | 43 | 0.019216 | 19216 | | | |
| 2505AshInt6bFung_S293_L001_R1_001 | 85.40626583 | 53 | 0.013821 | 13821 | | | |
| 2505AshInt6bFung_S293_L001_R2_001 | 82.01287895 | 52 | 0.013821 | 13821 | | | |
| 2505AshInt6cFung_S307_L001_R1_001 | 92.32803472 | 53 | 0.104367 | 104367 | | | |
| 2505AshInt6cFung_S307_L001_R2_001 | 93.89845449 | 49 | 0.104367 | 104367 | | | |
| 2505Wmcc6aFung_S318_L001_R1_001 | 83.69579288 | 54 | 0.01545 | 15450 | | | |
| 2505Wmcc6aFung_S318_L001_R2_001 | 79.58576052 | 53 | 0.01545 | 15450 | | | |
| 2505Wmcc6bFung_S322_L001_R1_001 | 83.09133489 | 53 | 0.00854 | 8540 | | | |
| 2505Wmcc6bFung_S322_L001_R2_001 | 79.21545667 | 53 | 0.00854 | 8540 | | | |
| 2505Wmcc6cFung_S316_L001_R1_001 | 79.82897384 | 52 | 0.003976 | 3976 | | | |
| 2505Wmcc6cFung_S316_L001_R2_001 | 75.30181087 | 52 | 0.003976 | 3976 | | | |
| 2506AshExt6aFung_S319_L001_R1_001 | 89.93880657 | 58 | 0.032193 | 32193 | | | |
| 2506AshExt6aFung_S319_L001_R2_001 | 88.92305781 | 58 | 0.032193 | 32193 | | | |
| 2506AshExt6bFung_S294_L001_R1_001 | 86.39663182 | 58 | 0.013776 | 13776 | | | |
| 2506AshExt6bFung_S294_L001_R2_001 | 86.04094077 | 59 | 0.013776 | 13776 | | | |
| 2506AshExt6cFung_S303_L001_R1_001 | 92.21684341 | 49 | 0.053089 | 53089 | | | |
| 2506AshExt6cFung_S303_L001_R2_001 | 91.94560078 | 47 | 0.053089 | 53089 | | | |
| 2506AshInt6aFung_S320_L001_R1_001 | 93.62753639 | 45 | 0.063131 | 63131 | | | |
| 2506AshInt6aFung_S320_L001_R2_001 | 92.73415596 | 46 | 0.063131 | 63131 | | | |
| 2506AshInt6bFung_S321_L001_R1_001 | 91.69724915 | 49 | 0.031881 | 31881 | | | |
| 2506AshInt6bFung_S321_L001_R2_001 | 92.37790534 | 46 | 0.031881 | 31881 | | | |
| 2506AshInt6cFung_S292_L001_R1_001 | 93.85395704 | 48 | 0.048698 | 48698 | | | |
| 2506AshInt6cFung_S292_L001_R2_001 | 95.3016551 | 44 | 0.048698 | 48698 | | | |
| 2506Wmcc6aFung_S301_L001_R1_001 | 94.11770621 | 43 | 0.099451 | 99451 | | | |
| 2506Wmcc6aFung_S301_L001_R2_001 | 95.04077385 | 44 | 0.099451 | 99451 | | | |
| 2506Wmcc6bFung_S299_L001_R1_001 | 86.21873024 | 57 | 0.022146 | 22146 | | | |

| | | | | |
|---|---|---|---|---|
| 2506Wmcc6bFung_S299_L001_R2_001 | 81.19299196 | 56 | 0.022146 | 22146 |
| 2506Wmcc6cFung_S300_L001_R1_001 | 91.75957872 | 49 | 0.05507 | 55070 |
| 2506Wmcc6cFung_S300_L001_R2_001 | 91.81768658 | 50 | 0.05507 | 55070 |
| 2507AshExt6W4aFung_S297_L001_R1_001 | 92.98497479 | 54 | 0.061297 | 61297 |
| 2507AshExt6W4aFung_S297_L001_R2_001 | 94.62127021 | 53 | 0.061297 | 61297 |
| 2507AshExt6W4bFung_S311_L001_R1_001 | 93.23449614 | 49 | 0.098411 | 98411 |
| 2507AshExt6W4bFung_S311_L001_R2_001 | 94.08602697 | 46 | 0.098411 | 98411 |
| 2507AshExt6W4cFung_S314_L001_R1_001 | 92.48748535 | 54 | 0.053737 | 53737 |
| 2507AshExt6W4cFung_S314_L001_R2_001 | 93.06809089 | 53 | 0.053737 | 53737 |
| 2507AshExt6aFung_S283_L001_R1_001 | 71.51079137 | 51 | 0.00139 | 1390 |
| 2507AshExt6aFung_S283_L001_R2_001 | 66.83453237 | 51 | 0.00139 | 1390 |
| 2507AshExt6bFung_S289_L001_R1_001 | 87.21246417 | 47 | 0.013607 | 13607 |
| 2507AshExt6bFung_S289_L001_R2_001 | 87.98412582 | 48 | 0.013607 | 13607 |
| 2507AshExt6cFung_S290_L001_R1_001 | 56.08695652 | 43 | 0.00046 | 460 |
| 2507AshExt6cFung_S290_L001_R2_001 | 52.39130435 | 43 | 0.00046 | 460 |
| 2507AshInt6W4aFung_S313_L001_R1_001 | 85.81911466 | 49 | 0.016762 | 16762 |
| 2507AshInt6W4aFung_S313_L001_R2_001 | 83.42679871 | 49 | 0.016762 | 16762 |
| 2507AshInt6W4bFung_S312_L001_R1_001 | 92.17309324 | 46 | 0.053035 | 53035 |
| 2507AshInt6W4bFung_S312_L001_R2_001 | 91.66399547 | 47 | 0.053035 | 53035 |
| 2507AshInt6W4cFung_S273_L001_R1_001 | 87.13872189 | 51 | 0.022455 | 22455 |
| 2507AshInt6W4cFung_S273_L001_R2_001 | 85.46426186 | 51 | 0.022455 | 22455 |
| 2507AshInt6aFung_S315_L001_R1_001 | 88.48506028 | 54 | 0.051507 | 51507 |
| 2507AshInt6aFung_S315_L001_R2_001 | 87.35705826 | 54 | 0.051507 | 51507 |
| 2507AshInt6bFung_S281_L001_R1_001 | 86.75011276 | 53 | 0.017736 | 17736 |
| 2507AshInt6bFung_S281_L001_R2_001 | 83.82949932 | 53 | 0.017736 | 17736 |
| 2507AshInt6cFung_S291_L001_R1_001 | 86.51354131 | 48 | 0.014585 | 14585 |
| 2507AshInt6cFung_S291_L001_R2_001 | 85.72505999 | 49 | 0.014585 | 14585 |
| 2507Wmcc6W4aFung_S287_L001_R1_001 | 90.58655814 | 49 | 0.043491 | 43491 |
| 2507Wmcc6W4aFung_S287_L001_R2_001 | 92.1661953 | 48 | 0.043491 | 43491 |
| 2507Wmcc6W4bFung_S270_L001_R1_001 | 91.01831242 | 50 | 0.039536 | 39536 |
| 2507Wmcc6W4bFung_S270_L001_R2_001 | 91.8884055 | 49 | 0.039536 | 39536 |
| 2507Wmcc6W4cFung_S288_L001_R1_001 | 91.8058708 | 48 | 0.039211 | 39211 |
| 2507Wmcc6W4cFung_S288_L001_R2_001 | 92.83109332 | 47 | 0.039211 | 39211 |
| 2507Wmcc6aFung_S282_L001_R1_001 | 83.91068639 | 52 | 0.010883 | 10883 |

| | | | | |
|---|---|---|---|---|
| 2507Wmcc6aFung_S282_L001_R2_001 | 77.51539098 | 52 | 0.010883 | 10883 |
| 2507Wmcc6bFung_S286_L001_R1_001 | 86.06933281 | 52 | 0.020279 | 20279 |
| 2507Wmcc6bFung_S286_L001_R2_001 | 82.06025938 | 51 | 0.020279 | 20279 |
| 2507Wmcc6cFung_S277_L001_R1_001 | 86.12840286 | 53 | 0.010763 | 10763 |
| 2507Wmcc6cFung_S277_L001_R2_001 | 81.13908761 | 51 | 0.010763 | 10763 |
| 2508AshExt6W8aFung_S275_L001_R1_001 | 91.57645079 | 54 | 0.026141 | 26141 |
| 2508AshExt6W8aFung_S275_L001_R2_001 | 92.87708963 | 53 | 0.026141 | 26141 |
| 2508AshExt6W8bFung_S268_L001_R1_001 | 93.08022146 | 54 | 0.060869 | 60869 |
| 2508AshExt6W8bFung_S268_L001_R2_001 | 94.70502226 | 53 | 0.060869 | 60869 |
| 2508AshExt6W8cFung_S285_L001_R1_001 | 92.27148135 | 53 | 0.033564 | 33564 |
| 2508AshExt6W8cFung_S285_L001_R2_001 | 93.69860565 | 53 | 0.033564 | 33564 |
| 2508AshExt6aFung_S276_L001_R1_001 | 68.01470588 | 40 | 0.000272 | 272 |
| 2508AshExt6aFung_S276_L001_R2_001 | 60.66176471 | 44 | 0.000272 | 272 |
| 2508AshExt6bFung_S284_L001_R1_001 | 73.56746765 | 39 | 0.000541 | 541 |
| 2508AshExt6bFung_S284_L001_R2_001 | 70.42513863 | 43 | 0.000541 | 541 |
| 2508AshExt6cFung_S274_L001_R1_001 | 91.95337165 | 58 | 0.037059 | 37059 |
| 2508AshExt6cFung_S274_L001_R2_001 | 91.38940608 | 58 | 0.037059 | 37059 |
| 2508AshInt6W8aFung_S269_L001_R1_001 | 86.89618519 | 58 | 0.016934 | 16934 |
| 2508AshInt6W8aFung_S269_L001_R2_001 | 82.87468997 | 58 | 0.016934 | 16934 |
| 2508AshInt6W8bFung_S280_L001_R1_001 | 84.89693701 | 54 | 0.010382 | 10382 |
| 2508AshInt6W8bFung_S280_L001_R2_001 | 80.21575804 | 54 | 0.010382 | 10382 |
| 2508AshInt6W8cFung_S279_L001_R1_001 | 85.99235315 | 51 | 0.014385 | 14385 |
| 2508AshInt6W8cFung_S279_L001_R2_001 | 82.75286757 | 50 | 0.014385 | 14385 |
| 2508AshInt6aFung_S271_L001_R1_001 | 88.70013523 | 60 | 0.023664 | 23664 |
| 2508AshInt6aFung_S271_L001_R2_001 | 87.60141988 | 61 | 0.023664 | 23664 |
| 2508AshInt6bFung_S272_L001_R1_001 | 89.97398136 | 58 | 0.039587 | 39587 |
| 2508AshInt6bFung_S272_L001_R2_001 | 87.93543335 | 57 | 0.039587 | 39587 |
| 2508AshInt6cFung_S278_L001_R1_001 | 80.88262223 | 54 | 0.005461 | 5461 |
| 2508AshInt6cFung_S278_L001_R2_001 | 75.88353781 | 54 | 0.005461 | 5461 |
| 2508Wmcc6W8aFung_S310_L001_R1_001 | 90.63120205 | 49 | 0.048875 | 48875 |
| 2508Wmcc6W8aFung_S310_L001_R2_001 | 90.71713555 | 48 | 0.048875 | 48875 |
| 2508Wmcc6W8bFung_S306_L001_R1_001 | 92.55947629 | 50 | 0.06263 | 62630 |
| 2508Wmcc6W8bFung_S306_L001_R2_001 | 93.96136037 | 50 | 0.06263 | 62630 |
| 2508Wmcc6W8cFung_S304_L001_R1_001 | 92.01596455 | 49 | 0.056876 | 56876 |

| | | | | |
|---|---|---|---|---|
| 2508Wmcc6W8cFung_S304_L001_R2_001 | 93.34868837 | 48 | 0.056876 | 56876 |
| 2508Wmcc6aFung_S295_L001_R1_001 | 77.36389685 | 50 | 0.002792 | 2792 |
| 2508Wmcc6aFung_S295_L001_R2_001 | 72.70773639 | 49 | 0.002792 | 2792 |
| 2508Wmcc6bFung_S302_L001_R1_001 | 86.83700201 | 50 | 0.029393 | 29393 |
| 2508Wmcc6bFung_S302_L001_R2_001 | 83.50967918 | 49 | 0.029393 | 29393 |
| 2508Wmcc6cFung_S296_L001_R1_001 | 86.29080713 | 47 | 0.009986 | 9986 |
| 2508Wmcc6cFung_S296_L001_R2_001 | 84.32805928 | 45 | 0.009986 | 9986 |
| EID2510BlankFung_S317_L001_R1_001 | 63.32179931 | 41 | 0.000289 | 289 |
| EID2510BlankFung_S317_L001_R2_001 | 53.28719723 | 42 | 0.000289 | 289 |
| RID001585Blank_S323_L001_R1_001 | 56.61641541 | 41 | 0.000597 | 597 |
| RID001585Blank_S323_L001_R2_001 | 56.95142379 | 42 | 0.000597 | 597 |
| RID001585Zymo_S324_L001_R1_001 | 88.50424559 | 42 | 0.001531 | 1531 |
| RID001585Zymo_S324_L001_R2_001 | 89.15741346 | 40 | 0.001531 | 1531 |
