## Supplementary material for "Microbial, dietary insect, and pathogen communities in fresh and decomposing guano of anthropic little brown bat (Myotis lucifugus) maternity colonies": Table S11

| Subset | ModelVariables | Df | SumSq | MeanSq | F Value | Pr(>F) |
|---|---|---|---|---|---|---|
| None | shannon~Site | 2 | 10.78 | 5.389 | 7.018 | 1.67E-03 |
| | shannon~Age | 3 | 26.09 | 8.696 | 11.324 | 3.84E-06 |
| | shannon~Age:Site | 6 | 29.84 | 4.973 | 6.476 | 1.82E-05 |
| | Residuals | 70 | 53.76 | 0.768 | | |
| | | **Tukey** | **Diff** | **Lower** | **Upper** | **p adj** |
| | | Fresh:ASHFInt-Fresh:ASHFExt | -0.3693972 | -1.6061873 | 0.86739291 | 0.9969891 |
| | | Fresh:WMCC-Fresh:ASHFExt | 0.94124608 | -0.2349067 | 2.1173989 | 0.2454143 |
| | | Week 4:ASHFExt-Fresh:ASHFExt | -1.47091445 | -2.9746503 | 0.03282137 | 0.0609106 |
| | | Week 4:ASHFInt-Fresh:ASHFExt | 0.88315338 | -0.6205824 | 2.3868892 | 0.7011386 |
| | | Week 4:WMCC-Fresh:ASHFExt | -1.13185362 | -2.6355894 | 0.3718822 | 0.3307863 |
| | | Week 8:ASHFExt-Fresh:ASHFExt | -1.54058331 | -2.9731338 | -0.1080328 | 0.0243403 |
| | | Week 8:ASHFInt-Fresh:ASHFExt | 0.02596355 | -1.9038978 | 1.95582486 | 1 |
| | | Week 8:WMCC-Fresh:ASHFExt | -1.25025788 | -2.6828084 | 0.18229263 | 0.1461804 |
| | | Week 12:ASHFExt-Fresh:ASHFExt | -1.26371045 | -3.1935718 | 0.66615086 | 0.5455689 |
| | | Week 12:ASHFInt-Fresh:ASHFExt | -0.69565379 | -2.6255151 | 1.23420753 | 9.86E-01 |
| | | Week 12:WMCC-Fresh:ASHFExt | -0.96447412 | -2.8943354 | 0.9653872 | 0.8667792 |
| | | Fresh:WMCC-Fresh:ASHFInt | 1.31064328 | 0.1631116 | 2.45817496 | 0.0123119 |
| | | Week 4:ASHFExt-Fresh:ASHFInt | -1.10151725 | -2.5829743 | 0.37993978 | 0.3489452 |
| | | Week 4:ASHFInt-Fresh:ASHFInt | 1.25255058 | -0.2289064 | 2.73400761 | 0.1791685 |
| | | Week 4:WMCC-Fresh:ASHFInt | -0.76245642 | -2.2439134 | 0.71900061 | 0.8431885 |
| | | Week 8:ASHFExt-Fresh:ASHFInt | -1.17118611 | -2.5803328 | 0.2379606 | 0.1986146 |
| | | Week 8:ASHFInt-Fresh:ASHFInt | 0.39536075 | -1.517192 | 2.30791355 | 0.9999034 |
| | | Week 8:WMCC-Fresh:ASHFInt | -0.88086068 | -2.2900074 | 0.52828603 | 0.6151586 |
| | | Week 12:ASHFExt-Fresh:ASHFInt | -0.89431325 | -2.806866 | 1.01823955 | 0.9101489 |
| | | Week 12:ASHFInt-Fresh:ASHFInt | -0.32625658 | -2.2388094 | 1.58629621 | 0.999986 |
| | | Week 12:WMCC-Fresh:ASHFInt | -0.59507692 | -2.5076297 | 1.31747588 | 0.9957125 |
| | | Week 4:ASHFExt-Fresh:WMCC | -2.41216053 | -3.843384 | -0.98093707 | 0.0000165 |
| | | Week 4:ASHFInt-Fresh:WMCC | -0.0580927 | -1.4893162 | 1.37313076 | 1 |
| | | Week 4:WMCC-Fresh:WMCC | -2.0730997 | -3.5043232 | -0.64187624 | 3.55E-04 |
| | | Week 8:ASHFExt-Fresh:WMCC | -2.48182939 | -3.8380669 | -1.1255919 | 0.0000023 |
| | | Week 8:ASHFInt-Fresh:WMCC | -0.91528253 | -2.7891939 | 0.95862885 | 0.8832466 |
| | | Week 8:WMCC-Fresh:WMCC | -2.19150396 | -3.5477414 | -0.83526647 | 0.0000415 |
| | | Week 12:ASHFExt-Fresh:WMCC | -2.20495653 | -4.0788679 | -0.33104515 | 0.0085266 |
| | | Week 12:ASHFInt-Fresh:WMCC | -1.63689987 | -3.5108112 | 0.23701152 | 0.1453208 |

| | | | | | | |
|---|---|---|---|---|---|---|
| | | Week 12:WMCC-Fresh:WMCC | -1.9057202 | -3.7796316 | -0.03180882 | 0.0427341 |
| | | Week 4:ASHFInt-Week 4:ASHFExt | 2.35406783 | 0.6434286 | 4.06470706 | 0.0008643 |
| | | Week 4:WMCC-Week 4:ASHFExt | 0.33906083 | -1.3715784 | 2.04970006 | 0.9999363 |
| | | Week 8:ASHFExt-Week 4:ASHFExt | -0.06966886 | -1.7180821 | 1.57874435 | 1 |
| | | Week 8:ASHFInt-Week 4:ASHFExt | 1.496878 | -0.5982186 | 3.59197462 | 4.10E-01 |
| | | Week 8:WMCC-Week 4:ASHFExt | 0.22065657 | -1.4277566 | 1.86906977 | 0.9999989 |
| | | Week 12:ASHFExt-Week 4:ASHFExt | 0.207204 | -1.8878926 | 2.30230062 | 1 |
| | | Week 12:ASHFInt-Week 4:ASHFExt | 0.77526067 | -1.319836 | 2.87035728 | 0.9824101 |
| | | Week 12:WMCC-Week 4:ASHFExt | 0.50644033 | -1.5886563 | 2.60153695 | 0.9995621 |
| | | Week 4:WMCC-Week 4:ASHFInt | -2.015007 | -3.7256462 | -0.30436778 | 0.0084109 |
| | | Week 8:ASHFExt-Week 4:ASHFInt | -2.42373669 | -4.0721499 | -0.77532349 | 0.0002703 |
| | | Week 8:ASHFInt-Week 4:ASHFInt | -0.85718983 | -2.9522865 | 1.23790678 | 0.9630047 |
| | | Week 8:WMCC-Week 4:ASHFInt | -2.13341126 | -3.7818245 | -0.48499806 | 0.0022827 |
| | | Week 12:ASHFExt-Week 4:ASHFInt | -2.14686383 | -4.2419605 | -0.05176722 | 0.0397439 |
| | | Week 12:ASHFInt-Week 4:ASHFInt | -1.57880717 | -3.6739038 | 0.51628945 | 0.3290729 |
| | | Week 12:WMCC-Week 4:ASHFInt | -1.8476275 | -3.9427241 | 0.24746912 | 0.1362467 |
| | | Week 8:ASHFExt-Week 4:WMCC | -0.40872969 | -2.0571429 | 1.23968351 | 0.9994436 |
| | | Week 8:ASHFInt-Week 4:WMCC | 1.15781717 | -0.9372795 | 3.25291378 | 0.7739192 |
| | | Week 8:WMCC-Week 4:WMCC | -0.11840426 | -1.7668175 | 1.53000894 | 1 |
| | | Week 12:ASHFExt-Week 4:WMCC | -0.13185683 | -2.2269535 | 1.96323978 | 1 |
| | | Week 12:ASHFInt-Week 4:WMCC | 0.43619983 | -1.6588968 | 2.53129645 | 0.9998964 |
| | | Week 12:WMCC-Week 4:WMCC | 0.1673795 | -1.9277171 | 2.26247612 | 1 |
| | | Week 8:ASHFInt-Week 8:ASHFExt | 1.56654686 | -0.4780581 | 3.6111518 | 0.3050113 |
| | | Week 8:WMCC-Week 8:ASHFExt | 0.29032543 | -1.2934187 | 1.87406961 | 0.9999709 |
| | | Week 12:ASHFExt-Week 8:ASHFExt | 0.27687286 | -1.7677321 | 2.3214778 | 0.9999987 |
| | | Week 12:ASHFInt-Week 8:ASHFExt | 0.84492952 | -1.1996754 | 2.88953447 | 0.9603011 |
| | | Week 12:WMCC-Week 8:ASHFExt | 0.57610919 | -1.4684958 | 2.62071413 | 0.9982014 |
| | | Week 8:WMCC-Week 8:ASHFInt | -1.27622143 | -3.3208264 | 0.76838351 | 0.6172823 |
| | | Week 12:ASHFExt-Week 8:ASHFInt | -1.289674 | -3.7088832 | 1.12953519 | 0.8111184 |
| | | Week 12:ASHFInt-Week 8:ASHFInt | -0.72161733 | -3.1408265 | 1.69759186 | 0.9970231 |
| | | Week 12:WMCC-Week 8:ASHFInt | -0.99043767 | -3.4096469 | 1.42877153 | 0.9628347 |
| | | Week 12:ASHFExt-Week 8:WMCC | -0.01345257 | -2.0580575 | 2.03115237 | 1 |
| | | Week 12:ASHFInt-Week 8:WMCC | 0.5546041 | -1.4900008 | 2.59920904 | 0.998724 |
| | | Week 12:WMCC-Week 8:WMCC | 0.28578376 | -1.7588212 | 2.3303887 | 0.9999982 |
| | | Week 12:ASHFInt-Week 12:ASHFExt | 0.56805667 | -1.8511525 | 2.98726586 | 0.9996676 |

| | | | | | | |
|---|---|---|---|---|---|---|
| | | Week 12:WMCC-Week 12:ASHFExt | 0.29923633 | -2.1199729 | 2.71844553 | 0.9999995 |
| | | Week 12:WMCC-Week 12:ASHFInt | -0.26882033 | -2.6880295 | 2.15038886 | 0.9999998 |
| **Subset** | **ModelVariables** | **Df** | **SumSq** | **MeanSq** | **F Value** | **Pr(>F)** |
| None | invsimpson~Site | 2 | 835 | 417.5 | 2.679 | 7.57E-02 |
| | invsimpson~Age | 3 | 2118 | 706.1 | 4.531 | 5.83E-03 |
| | invsimpson~Age:Site | 6 | 4607 | 767.9 | 4.927 | 2.96E-04 |
| | Residuals | 70 | 10909 | 155.8 | | |
| | | **Tukey** | **Diff** | **Lower** | **Upper** | **p adj** |
| | | Fresh:ASHFInt-Fresh:ASHFExt | -8.9205667 | -26.538996 | 8.6978625 | 0.8566664 |
| | | Fresh:WMCC-Fresh:ASHFExt | 12.05903561 | -4.695598 | 28.8136693 | 0.3984184 |
| | | Week 4:ASHFExt-Fresh:ASHFExt | -11.55024362 | -32.9713909 | 9.8709037 | 0.7998378 |
| | | Week 4:ASHFInt-Fresh:ASHFExt | 13.27486938 | -8.1462779 | 34.6960167 | 0.6276892 |
| | | Week 4:WMCC-Fresh:ASHFExt | -9.60556845 | -31.0267157 | 11.8155788 | 0.9310661 |
| | | Week 8:ASHFExt-Fresh:ASHFExt | -11.59973674 | -32.0068288 | 8.8073554 | 0.7417355 |
| | | Week 8:ASHFInt-Fresh:ASHFExt | -7.64752012 | -35.138947 | 19.8439068 | 0.9983971 |
| | | Week 8:WMCC-Fresh:ASHFExt | -10.39818017 | -30.8052723 | 10.0089119 | 0.8515504 |
| | | Week 12:ASHFExt-Fresh:ASHFExt | -11.16397145 | -38.6553984 | 16.3274555 | 0.9649244 |
| | | Week 12:ASHFInt-Fresh:ASHFExt | -8.35096845 | -35.8423954 | 19.1404585 | 9.97E-01 |
| | | Week 12:WMCC-Fresh:ASHFExt | -8.75872279 | -36.2501497 | 18.7327041 | 0.9947562 |
| | | Fresh:WMCC-Fresh:ASHFInt | 20.97960232 | 4.6326851 | 37.3265196 | 0.0025874 |
| | | Week 4:ASHFExt-Fresh:ASHFInt | -2.62967692 | -23.7334563 | 18.4741025 | 0.9999995 |
| | | Week 4:ASHFInt-Fresh:ASHFInt | 22.19543608 | 1.0916567 | 43.2992155 | 0.030701 |
| | | Week 4:WMCC-Fresh:ASHFInt | -0.68500175 | -21.7887812 | 20.4187777 | 1 |
| | | Week 8:ASHFExt-Fresh:ASHFInt | -2.67917004 | -22.7528683 | 17.3945282 | 0.9999989 |
| | | Week 8:ASHFInt-Fresh:ASHFInt | 1.27304658 | -25.9718155 | 28.5179086 | 1 |
| | | Week 8:WMCC-Fresh:ASHFInt | -1.47761346 | -21.5513117 | 18.5960848 | 1 |
| | | Week 12:ASHFExt-Fresh:ASHFInt | -2.24340475 | -29.4882668 | 25.0014573 | 1 |
| | | Week 12:ASHFInt-Fresh:ASHFInt | 0.56959825 | -26.6752638 | 27.8144603 | 1.00E+00 |
| | | Week 12:WMCC-Fresh:ASHFInt | 0.16184392 | -27.0830181 | 27.406706 | 1 |
| | | Week 4:ASHFExt-Fresh:WMCC | -23.60927923 | -43.9974671 | -3.2210914 | 0.0104135 |
| | | Week 4:ASHFInt-Fresh:WMCC | 1.21583377 | -19.1723541 | 21.6040216 | 1 |
| | | Week 4:WMCC-Fresh:WMCC | -21.66460407 | -42.0527919 | -1.2764162 | 0.0276144 |
| | | Week 8:ASHFExt-Fresh:WMCC | -23.65877235 | -42.9787636 | -4.3387811 | 0.0050426 |
| | | Week 8:ASHFInt-Fresh:WMCC | -19.70655573 | -46.4009598 | 6.9878483 | 0.3597048 |
| | | Week 8:WMCC-Fresh:WMCC | -22.45721578 | -41.777207 | -3.1372245 | 0.0099374 |

| | | | | | | |
|---|---|---|---|---|---|---|
| | | Week 12:ASHFExt-Fresh:WMCC | -23.22300707 | -49.9174111 | 3.471397 | 0.1493267 |
| | | Week 12:ASHFInt-Fresh:WMCC | -20.41000407 | -47.1044081 | 6.2844 | 0.3080163 |
| | | Week 12:WMCC-Fresh:WMCC | -20.8177584 | -47.5121625 | 5.8766457 | 0.2801394 |
| | | Week 4:ASHFInt-Week 4:ASHFExt | 24.825113 | 0.4565676 | 49.1936584 | 0.0420362 |
| | | Week 4:WMCC-Week 4:ASHFExt | 1.94467517 | -22.4238703 | 26.3132206 | 1 |
| | | Week 8:ASHFExt-Week 4:ASHFExt | -0.04949312 | -23.5316111 | 23.4326248 | 1 |
| | | Week 8:ASHFInt-Week 4:ASHFExt | 3.9027235 | -25.9425276 | 33.7479746 | 0.9999991 |
| | | Week 8:WMCC-Week 4:ASHFExt | 1.15206345 | -22.3300545 | 24.6341814 | 1 |
| | | Week 12:ASHFExt-Week 4:ASHFExt | 0.38627217 | -29.4589789 | 30.2315232 | 1 |
| | | Week 12:ASHFInt-Week 4:ASHFExt | 3.19927517 | -26.6459759 | 33.0445262 | 0.9999999 |
| | | Week 12:WMCC-Week 4:ASHFExt | 2.79152083 | -27.0537302 | 32.6367719 | 1 |
| | | Week 4:WMCC-Week 4:ASHFInt | -22.88043783 | -47.2489833 | 1.4881076 | 0.0857015 |
| | | Week 8:ASHFExt-Week 4:ASHFInt | -24.87460612 | -48.3567241 | -1.3924882 | 0.0285215 |
| | | Week 8:ASHFInt-Week 4:ASHFInt | -20.9223895 | -50.7676406 | 8.9228616 | 0.4393609 |
| | | Week 8:WMCC-Week 4:ASHFInt | -23.67304955 | -47.1551675 | -0.1909316 | 0.0463946 |
| | | Week 12:ASHFExt-Week 4:ASHFInt | -24.43884083 | -54.2840919 | 5.4064102 | 0.2164178 |
| | | Week 12:ASHFInt-Week 4:ASHFInt | -21.62583783 | -51.4710889 | 8.2194132 | 0.3880701 |
| | | Week 12:WMCC-Week 4:ASHFInt | -22.03359217 | -51.8788432 | 7.8116589 | 0.3596369 |
| | | Week 8:ASHFExt-Week 4:WMCC | -1.99416829 | -25.4762862 | 21.4879497 | 1 |
| | | Week 8:ASHFInt-Week 4:WMCC | 1.95804833 | -27.8872027 | 31.8032994 | 1 |
| | | Week 8:WMCC-Week 4:WMCC | -0.79261171 | -24.2747297 | 22.6895062 | 1.00E+00 |
| | | Week 12:ASHFExt-Week 4:WMCC | -1.558403 | -31.4036541 | 28.2868481 | 1 |
| | | Week 12:ASHFInt-Week 4:WMCC | 1.2546 | -28.5906511 | 31.0998511 | 1 |
| | | Week 12:WMCC-Week 4:WMCC | 0.84684567 | -28.9984054 | 30.6920967 | 1 |
| | | Week 8:ASHFInt-Week 8:ASHFExt | 3.95221662 | -25.1737661 | 33.0781993 | 0.9999987 |
| | | Week 8:WMCC-Week 8:ASHFExt | 1.20155657 | -21.3593326 | 23.7624457 | 1 |
| | | Week 12:ASHFExt-Week 8:ASHFExt | 0.43576529 | -28.6902174 | 29.561748 | 1 |
| | | Week 12:ASHFInt-Week 8:ASHFExt | 3.24876829 | -25.8772144 | 32.374751 | 0.9999998 |
| | | Week 12:WMCC-Week 8:ASHFExt | 2.84101395 | -26.2849687 | 31.9669966 | 1 |
| | | Week 8:WMCC-Week 8:ASHFInt | -2.75066005 | -31.8766427 | 26.3753226 | 1 |
| | | Week 12:ASHFExt-Week 8:ASHFInt | -3.51645133 | -37.9787788 | 30.9458761 | 1.00E+00 |
| | | Week 12:ASHFInt-Week 8:ASHFInt | -0.70344833 | -35.1657758 | 33.7588791 | 1 |
| | | Week 12:WMCC-Week 8:ASHFInt | -1.11120267 | -35.5735301 | 33.3511248 | 1 |
| | | Week 12:ASHFExt-Week 8:WMCC | -0.76579129 | -29.891774 | 28.3601914 | 1 |
| | | Week 12:ASHFInt-Week 8:WMCC | 2.04721171 | -27.078771 | 31.1731944 | 1 |

| | | | | | | |
|---|---|---|---|---|---|---|
| | | Week 12:WMCC-Week 8:WMCC | 1.63945738 | -27.4865253 | 30.7654401 | 1 |
| | | Week 12:ASHFInt-Week 12:ASHFExt | 2.813003 | -31.6493245 | 37.2753305 | 1 |
| | | Week 12:WMCC-Week 12:ASHFExt | 2.40524867 | -32.0570788 | 36.8675761 | 1 |
| | | Week 12:WMCC-Week 12:ASHFInt | -0.40775433 | -34.8700818 | 34.0545731 | 1 |
| **Subset** | **ModelVariables** | **Df** | **SumSq** | **MeanSq** | **F Value** | **Pr(>F)** |
| Fresh | shannon~Month | 3 | 12.54 | 4.181 | 5.641 | 5.73E-03 |
| *5 September samples removed | shannon~Site | 2 | 11.24 | 5.618 | 7.58 | 3.55E-03 |
| | shannon~Month:Site | 6 | 5.66 | 0.943 | 1.273 | 3.14E-01 |
| | Residuals | 20 | 14.82 | 0.741 | | |
| **Subset** | **ModelVariables** | **Df** | **SumSq** | **MeanSq** | **F Value** | **Pr(>F)** |
| Fresh | invsimpson~Month | 3 | 2450.9 | 817 | 12.073 | 9.88E-05 |
| *5 September samples removed | invsimpson~Site | 2 | 2188.4 | 1094.2 | 16.17 | 6.64E-05 |
| | invsimpson~Month:Site | 6 | 869.2 | 144.9 | 2.141 | 9.34E-02 |
| | Residuals | 20 | 1353.4 | 67.7 | | |
