## Supplementary material for "Microbial, dietary insect, and pathogen communities in fresh and decomposing guano of anthropic little brown bat (Myotis lucifugus) maternity colonies": Table S12

| Distance Matrix | Factor | Df | SumOfSqs | R2 | F | Pr(>F) |
|---|---|---|---|---|---|---|
| Jaccard | Site | 2 | 2.888 | 0.08379 | 4.1496 | 0.001 |
| Jaccard | Age | 3 | 3.303 | 0.09584 | 3.1643 | 0.001 |
| Jaccard | Site* | 2 | 0.9951 | 0.07934 | 1.4196 | 0.001 |
| Jaccard | Month* | 3 | 2.0605 | 0.16429 | 1.9597 | 0.001 |
| Jaccard | Site:Age | 6 | 3.919 | 0.11368 | 1.8768 | 0.001 |
| Jaccard | Site:Month* | 6 | 2.4767 | 0.19747 | 1.1778 | 0.001 |
| Bray-Curtis | Site | 2 | 4.1841 | 0.13455 | 7.7605 | 0.001 |
| Bray-Curtis | Age | 3 | 3.4786 | 0.11187 | 4.3014 | 0.001 |
| Bray-Curtis | Site* | 2 | 1.1779 | 0.09938 | 2.2449 | 0.001 |
| Bray-Curtis | Month* | 3 | 2.8369 | 0.23935 | 3.6045 | 0.001 |
| Bray-Curtis | Site:Age | 6 | 4.5633 | 0.14675 | 2.8213 | 0.001 |
| Bray-Curtis | Site:Month* | 6 | 2.5907 | 0.21858 | 1.6458 | 0.001 |
| Yue & Clayton | Site | 2 | 4.600 | 0.13699 | 7.7695 | 0.001 |
| Yue & Clayton | Age | 3 | 3.138 | 0.09347 | 3.5341 | 0.001 |
| Yue & Clayton | Site* | 2 | 1.4719 | 0.10795 | 2.3444 | 0.002 |
| Yue & Clayton | Month* | 3 | 2.7028 | 0.19824 | 2.8700 | 0.001 |
| Yue & Clayton | Site:Age | 6 | 5.119 | 0.15245 | 2.8821 | 0.001 |
| Yue & Clayton | Site:Month* | 6 | 3.1814 | 0.23334 | 1.6891 | 0.001 |

*Month and Site:Month tests run on only fresh samples from May-August
