## Supplementary material for "Microbial, dietary insect, and pathogen communities in fresh and decomposing guano of anthropic little brown bat (Myotis lucifugus) maternity colonies": Table S13

| Site | Family | Age | Mean Relative Abundance |
|---|---|---|---|
| ASHFExt | Saccharomycetaceae | Fresh | 12.1636364 |
| ASHFExt | unclassified_Plantae | Fresh | 10.3363636 |
| ASHFExt | Ascomycota_unclassified | Fresh | 10.2363636 |
| ASHFExt | Fungi_unclassified | Fresh | 8.07272727 |
| ASHFExt | Davidiellaceae | Fresh | 7.34545455 |
| ASHFExt | Herpotrichiellaceae | Fresh | 3.95 |
| ASHFExt | Sporidiobolales_family_I | Fresh | 3.05909091 |
| ASHFExt | Pleosporales_family_Inc | Fresh | 2.97272727 |
| ASHFExt | Pleosporales_unclassified | Fresh | 2.23181818 |
| ASHFExt | Montagnulaceae | Fresh | 2.22727273 |
| ASHFExt | Strophariaceae | Fresh | 1.96363636 |
| ASHFExt | Peniophoraceae | Fresh | 1.93181818 |
| ASHFExt | Meruliaceae | Fresh | 1.80454546 |
| ASHFExt | Taphrinaceae | Fresh | 1.59545455 |
| ASHFExt | Mycosphaerellaceae | Fresh | 1.42272727 |
| ASHFExt | Sordariomycetes_unclass | Fresh | 1.32272727 |
| ASHFExt | Dothideaceae | Fresh | 1.03181818 |
| ASHFExt | Filobasidiaceae | Fresh | 0.92272727 |
| ASHFExt | Russulales_unclassified | Fresh | 0.89545455 |
| ASHFExt | Basidiomycota_unclassif | Fresh | 0.89090909 |
| ASHFExt | Polyporaceae | Fresh | 0.87272727 |
| ASHFExt | Schizoporaceae | Fresh | 0.87272727 |
| ASHFExt | Eurotiomycetes_unclassi | Fresh | 0.86363636 |
| ASHFExt | Capnodiales_unclassified | Fresh | 0.84090909 |
| ASHFExt | Valsaceae | Fresh | 0.79090909 |
| ASHFExt | Microascaceae | Fresh | 0.77272727 |
| ASHFExt | Amphisphaeriaceae | Fresh | 0.62272727 |
| ASHFExt | Ophiocordycipitaceae | Fresh | 0.61818182 |

| | | | |
|---|---|---|---|
| ASHFExt | Psathyrellaceae | Fresh | 0.54545455 |
| ASHFExt | Ganodermataceae | Fresh | 0.53636364 |
| ASHFExt | Gnomoniaceae | Fresh | 0.45454546 |
| ASHFExt | Xylariaceae | Fresh | 0.44545455 |
| ASHFExt | Hydnodontaceae | Fresh | 0.44090909 |
| ASHFExt | Capnodiales_family_Ince | Fresh | 0.43636364 |
| ASHFExt | unclassified_Corticiales | Fresh | 0.43181818 |
| ASHFExt | Cordycipitaceae | Fresh | 0.39545455 |
| ASHFExt | Dothioraceae | Fresh | 0.39545455 |
| ASHFExt | Stereaceae | Fresh | 0.36818182 |
| ASHFExt | Trichocomaceae | Fresh | 0.36818182 |
| ASHFExt | Agaricomycetes_unclass | Fresh | 0.36363636 |
| ASHFExt | Venturiaceae | Fresh | 0.36363636 |
| ASHFExt | Tremellales_family_Ince | Fresh | 0.36363636 |
| ASHFExt | Dothideomycetes_unclas | Fresh | 0.35454546 |
| ASHFExt | Meripilaceae | Fresh | 0.35454546 |
| ASHFExt | Botryosphaeriaceae | Fresh | 0.34545455 |
| ASHFExt | Diatrypaceae | Fresh | 0.34545455 |
| ASHFExt | Cucurbitariaceae | Fresh | 0.33181818 |
| ASHFExt | unclassified_Pleosporale | Fresh | 0.33181818 |
| ASHFExt | Eurotiomycetes_family_I | Fresh | 0.30454546 |
| ASHFExt | Thelebolaceae | Fresh | 0.26818182 |
| ASHFExt | Ascomycota_family_Inc | Fresh | 0.26363636 |
| ASHFExt | Ceratobasidiaceae | Fresh | 0.24545455 |
| ASHFExt | Mucoraceae | Fresh | 0.23636364 |
| ASHFExt | Phaeosphaeriaceae | Fresh | 0.23636364 |
| ASHFExt | Mycocaliciaceae | Fresh | 0.23181818 |
| ASHFExt | Phanerochaetaceae | Fresh | 0.22727273 |
| ASHFExt | Inocybaceae | Fresh | 0.22727273 |

| | | | |
|---|---|---|---|
| ASHFExt | Exobasidiaceae | Fresh | 0.21818182 |
| ASHFExt | Sclerotiniaceae | Fresh | 0.19545455 |
| ASHFExt | Hydnaceae | Fresh | 0.19090909 |
| ASHFExt | Pleosporaceae | Fresh | 0.19090909 |
| ASHFExt | Fomitopsidaceae | Fresh | 0.18636364 |
| ASHFExt | Diaporthales_family_Inc | Fresh | 0.18636364 |
| ASHFExt | Entolomataceae | Fresh | 0.18636364 |
| ASHFExt | Hymenochaetaceae | Fresh | 0.15909091 |
| ASHFExt | Hypocreales_unclassifiec | Fresh | 0.15909091 |
| ASHFExt | Leptosphaeriaceae | Fresh | 0.15454546 |
| ASHFExt | Chaetothyriales_unclassi | Fresh | 0.15 |
| ASHFExt | Marasmiaceae | Fresh | 0.13636364 |
| ASHFExt | Suillaceae | Fresh | 0.13636364 |
| ASHFExt | Pluteaceae | Fresh | 0.13181818 |
| ASHFExt | Agaricales_unclassified | Fresh | 0.12272727 |
| ASHFExt | Auriscalpiaceae | Fresh | 0.11818182 |
| ASHFExt | Glomerellaceae | Fresh | 0.11818182 |
| ASHFExt | Saccharomycetales_fami | Fresh | 0.11363636 |
| ASHFExt | Cantharellales_unclassifi | Fresh | 0.11363636 |
| ASHFExt | Lecanorales_unclassified | Fresh | 0.10909091 |
| ASHFExt | Helotiales_family_Incert | Fresh | 0.10454546 |
| ASHFExt | Lophiostomataceae | Fresh | 0.10454546 |
| ASHFExt | Pyronemataceae | Fresh | 0.10454546 |
| ASHFExt | Atheliaceae | Fresh | 0.1 |
| ASHFExt | Mortierellaceae | Fresh | 0.1 |
| ASHFExt | Hymenochaetales_family | Fresh | 0.1 |
| ASHFExt | Polyporales_unclassified | Fresh | 0.09545455 |
| ASHFExt | Clavicipitaceae | Fresh | 0.09545455 |
| ASHFExt | Nectriaceae | Fresh | 0.09545455 |

| | | | |
|---|---|---|---|
| ASHFExt | unclassified_Atheliales | Fresh | 0.09545455 |
| ASHFExt | Cortinariaceae | Fresh | 0.09090909 |
| ASHFExt | Metschnikowiaceae | Fresh | 0.09090909 |
| ASHFExt | Hypocreaceae | Fresh | 0.08636364 |
| ASHFExt | Helotiales_unclassified | Fresh | 0.08636364 |
| ASHFExt | Teratosphaeriaceae | Fresh | 0.08181818 |
| ASHFExt | Annulatascaceae | Fresh | 0.07727273 |
| ASHFExt | Polyporales_family_Ince | Fresh | 0.07272727 |
| ASHFExt | Agaricaceae | Fresh | 0.06818182 |
| ASHFExt | Tricholomataceae | Fresh | 0.06818182 |
| ASHFExt | Trichosphaeriales_family | Fresh | 0.05909091 |
| ASHFExt | Dothideomycetes_family | Fresh | 0.05909091 |
| ASHFExt | Hyaloscyphaceae | Fresh | 0.05909091 |
| ASHFExt | Agaricostilbaceae | Fresh | 0.05454546 |
| ASHFExt | Erythrobasidiales_family | Fresh | 0.05454546 |
| ASHFExt | Hypocreales_family_Inc | Fresh | 0.05454546 |
| ASHFExt | Teloschistales_unclassifi | Fresh | 0.05454546 |
| ASHFExt | Calosphaeriaceae | Fresh | 0.05 |
| ASHFExt | Graphostromataceae | Fresh | 0.05 |
| ASHFExt | Melampsoraceae | Fresh | 0.05 |
| ASHFExt | Sporormiaceae | Fresh | 0.05 |
| ASHFExt | Agaricostilbomycetes_ur | Fresh | 0.04545455 |
| ASHFExt | Umbelopsidaceae | Fresh | 0.04545455 |
| ASHFExt | Agaricomycetes_family_ | Fresh | 0.04090909 |
| ASHFExt | Microstromataceae | Fresh | 0.04090909 |
| ASHFExt | Taphrinales_unclassified | Fresh | 0.04090909 |
| ASHFExt | Xylariales_unclassified | Fresh | 0.04090909 |
| ASHFExt | Auriculariales_family_In | Fresh | 0.04090909 |
| ASHFExt | Bondarzewiaceae | Fresh | 0.04090909 |

| | | | |
|---|---|---|---|
| ASHFExt | Chaetothyriaceae | Fresh | 0.04090909 |
| ASHFExt | Botryobasidiaceae | Fresh | 0.04090909 |
| ASHFExt | Pezizaceae | Fresh | 0.04090909 |
| ASHFExt | Basidiobolaceae | Fresh | 0.04090909 |
| ASHFExt | Physciaceae | Fresh | 0.03636364 |
| ASHFExt | Diaporthales_unclassifie | Fresh | 0.03636364 |
| ASHFExt | Massarinaceae | Fresh | 0.03181818 |
| ASHFExt | Pleurotaceae | Fresh | 0.03181818 |
| ASHFExt | Ustilaginaceae | Fresh | 0.03181818 |
| ASHFExt | unclassified_Agaricostilt | Fresh | 0.03181818 |
| ASHFExt | Mucorales_unclassified | Fresh | 0.03181818 |
| ASHFExt | Hygrophoraceae | Fresh | 0.02727273 |
| ASHFExt | Lecanoromycetes_unclas | Fresh | 0.02727273 |
| ASHFExt | Halosphaeriaceae | Fresh | 0.02727273 |
| ASHFExt | Plectosphaerellaceae | Fresh | 0.02727273 |
| ASHFExt | Tremellomycetes_unclas | Fresh | 0.02272727 |
| ASHFExt | Chaetosphaeriaceae | Fresh | 0.02272727 |
| ASHFExt | Cystofilobasidiaceae | Fresh | 0.02272727 |
| ASHFExt | Elsinoaceae | Fresh | 0.02272727 |
| ASHFExt | Myxotrichaceae | Fresh | 0.02272727 |
| ASHFExt | Schizophyllaceae | Fresh | 0.02272727 |
| ASHFExt | Spizellomycetaceae | Fresh | 0.02272727 |
| ASHFExt | Tuberaceae | Fresh | 0.02272727 |
| ASHFExt | Bulgariaceae | Fresh | 0.02272727 |
| ASHFExt | Sarcosomataceae | Fresh | 0.02272727 |
| ASHFExt | unclassified_Myriangiale | Fresh | 0.02272727 |
| ASHFExt | Bionectriaceae | Fresh | 0.01818182 |
| ASHFExt | Bolbitiaceae | Fresh | 0.01818182 |
| ASHFExt | Chaetomiaceae | Fresh | 0.01818182 |

| | | | |
|---|---|---|---|
| ASHFExt | Clavulinaceae | Fresh | 0.01818182 |
| ASHFExt | Cystofilobasidiales_fami | Fresh | 0.01818182 |
| ASHFExt | Helotiaceae | Fresh | 0.01818182 |
| ASHFExt | Malasseziales_family_In | Fresh | 0.01818182 |
| ASHFExt | Mycenaceae | Fresh | 0.01818182 |
| ASHFExt | Phallaceae | Fresh | 0.01818182 |
| ASHFExt | Sebacinaceae | Fresh | 0.01818182 |
| ASHFExt | Serpulaceae | Fresh | 0.01818182 |
| ASHFExt | Tapinellaceae | Fresh | 0.01818182 |
| ASHFExt | Thelephoraceae | Fresh | 0.01818182 |
| ASHFExt | unclassified_Taphrinales | Fresh | 0.01818182 |
| ASHFExt | Microbotryaceae | Fresh | 0.01363636 |
| ASHFExt | Paxillaceae | Fresh | 0.01363636 |
| ASHFExt | Pucciniastraceae | Fresh | 0.01363636 |
| ASHFExt | Vibrisseaceae | Fresh | 0.01363636 |
| ASHFExt | Chaetothyriales_family_I | Fresh | 0.01363636 |
| ASHFExt | Russulaceae | Fresh | 0.01363636 |
| ASHFExt | Togniniaceae | Fresh | 0.01363636 |
| ASHFExt | unclassified_Cantharellal | Fresh | 0.01363636 |
| ASHFExt | Leotiomycetes_unclassif | Fresh | 0.00909091 |
| ASHFExt | Apiosporaceae | Fresh | 0.00909091 |
| ASHFExt | Ascosphaeraceae | Fresh | 0.00909091 |
| ASHFExt | Coniochaetaceae | Fresh | 0.00909091 |
| ASHFExt | Kickxellaceae | Fresh | 0.00909091 |
| ASHFExt | Lyophyllaceae | Fresh | 0.00909091 |
| ASHFExt | Parmeliaceae | Fresh | 0.00909091 |
| ASHFExt | Phragmidiaceae | Fresh | 0.00909091 |
| ASHFExt | Rhytismataceae | Fresh | 0.00909091 |
| ASHFExt | Septobasidiaceae | Fresh | 0.00909091 |

| | | | |
|---|---|---|---|
| ASHFExt | Sordariomycetes_family_ | Fresh | 0.00909091 |
| ASHFExt | Stictidaceae | Fresh | 0.00909091 |
| ASHFExt | Xenasmataceae | Fresh | 0.00909091 |
| ASHFExt | unclassified_Exobasidior | Fresh | 0.00909091 |
| ASHFExt | unclassified_Hysteriales | Fresh | 0.00909091 |
| ASHFExt | unclassified_Polyporales | Fresh | 0.00909091 |
| ASHFExt | unclassified_Trechispora | Fresh | 0.00909091 |
| ASHFExt | Rhytismatales_unclassifi | Fresh | 0.00909091 |
| ASHFExt | Pucciniomycetes_unclass | Fresh | 0.00454546 |
| ASHFExt | Agaricales_family_Incer | Fresh | 0.00454546 |
| ASHFExt | Anthracoideaceae | Fresh | 0.00454546 |
| ASHFExt | Cephalothecaceae | Fresh | 0.00454546 |
| ASHFExt | Ceratostomataceae | Fresh | 0.00454546 |
| ASHFExt | Chionosphaeraceae | Fresh | 0.00454546 |
| ASHFExt | Cladoniaceae | Fresh | 0.00454546 |
| ASHFExt | Corticiaceae | Fresh | 0.00454546 |
| ASHFExt | Cunninghamellaceae | Fresh | 0.00454546 |
| ASHFExt | Dacrymycetaceae | Fresh | 0.00454546 |
| ASHFExt | Dermateaceae | Fresh | 0.00454546 |
| ASHFExt | Diaporthaceae | Fresh | 0.00454546 |
| ASHFExt | Exobasidiomycetes_fami | Fresh | 0.00454546 |
| ASHFExt | Hebelomataceae | Fresh | 0.00454546 |
| ASHFExt | Hericiaceae | Fresh | 0.00454546 |
| ASHFExt | Leotiomycetes_family_I | Fresh | 0.00454546 |
| ASHFExt | Massariaceae | Fresh | 0.00454546 |
| ASHFExt | Melanconidaceae | Fresh | 0.00454546 |
| ASHFExt | Microbotryomycetes_fan | Fresh | 0.00454546 |
| ASHFExt | Pseudeurotiaceae | Fresh | 0.00454546 |
| ASHFExt | Pucciniaceae | Fresh | 0.00454546 |

| | | | |
|---|---|---|---|
| ASHFExt | Ramalinaceae | Fresh | 0.00454546 |
| ASHFExt | Rutstroemiaceae | Fresh | 0.00454546 |
| ASHFExt | Schizoparmaceae | Fresh | 0.00454546 |
| ASHFExt | Wallemiaceae | Fresh | 0.00454546 |
| ASHFExt | Xylariales_family_Incert | Fresh | 0.00454546 |
| ASHFExt | unclassified_Dothideales | Fresh | 0.00454546 |
| ASHFExt | Kickxellales_unclassifiec | Fresh | 0.00454546 |
| ASHFExt | Pucciniales_unclassified | Fresh | 0.00454546 |
| ASHFExt | Saccharomycetales_uncla | Fresh | 0.00454546 |
| ASHFExt | Sebacinales_unclassified | Fresh | 0.00454546 |
| ASHFExt | unknown_unclassified | Fresh | 0.00454546 |
| ASHFInt | Ascomycota_unclassifiec | Fresh | 15.4625 |
| ASHFInt | Saccharomycetaceae | Fresh | 13.2833333 |
| ASHFInt | unclassified_Plantae | Fresh | 10.25 |
| ASHFInt | Herpotrichiellaceae | Fresh | 8.125 |
| ASHFInt | Davidiellaceae | Fresh | 4.58333333 |
| ASHFInt | Pucciniaceae | Fresh | 3.975 |
| ASHFInt | Sordariomycetes_unclass | Fresh | 3.73333333 |
| ASHFInt | Hypocreales_unclassifiec | Fresh | 3.6875 |
| ASHFInt | Metschnikowiaceae | Fresh | 3.67083333 |
| ASHFInt | Sporidiobolales_family_I | Fresh | 2.46666667 |
| ASHFInt | Fungi_unclassified | Fresh | 2.42083333 |
| ASHFInt | Meruliaceae | Fresh | 1.79583333 |
| ASHFInt | Taphrinaceae | Fresh | 1.6875 |
| ASHFInt | Peniophoraceae | Fresh | 1.3625 |
| ASHFInt | Mycosphaerellaceae | Fresh | 1.30833333 |
| ASHFInt | Basidiomycota_unclassif | Fresh | 0.89166667 |
| ASHFInt | Pleosporales_family_Inc | Fresh | 0.76666667 |
| ASHFInt | Schizoporaceae | Fresh | 0.725 |

| | | | |
|---|---|---|---|
| ASHFInt | Stereaceae | Fresh | 0.72083333 |
| ASHFInt | Dothioraceae | Fresh | 0.71666667 |
| ASHFInt | Strophariaceae | Fresh | 0.69583333 |
| ASHFInt | Mortierellaceae | Fresh | 0.64583333 |
| ASHFInt | Venturiaceae | Fresh | 0.57916667 |
| ASHFInt | Polyporaceae | Fresh | 0.575 |
| ASHFInt | Eurotiomycetes_unclassi | Fresh | 0.55833333 |
| ASHFInt | Saccharomycetales_fami | Fresh | 0.55833333 |
| ASHFInt | Leotiomycetes_unclassif | Fresh | 0.52083333 |
| ASHFInt | Exobasidiaceae | Fresh | 0.50833333 |
| ASHFInt | Hypocreaceae | Fresh | 0.475 |
| ASHFInt | Polyporales_unclassified | Fresh | 0.41666667 |
| ASHFInt | Pleosporales_unclassifie | Fresh | 0.39583333 |
| ASHFInt | Amphisphaeriaceae | Fresh | 0.37916667 |
| ASHFInt | Pleurotaceae | Fresh | 0.375 |
| ASHFInt | Hydnodontaceae | Fresh | 0.36666667 |
| ASHFInt | Trichocomaceae | Fresh | 0.3625 |
| ASHFInt | Filobasidiaceae | Fresh | 0.35 |
| ASHFInt | Mucoraceae | Fresh | 0.35 |
| ASHFInt | Meripilaceae | Fresh | 0.34583333 |
| ASHFInt | Psathyrellaceae | Fresh | 0.34583333 |
| ASHFInt | Xylariaceae | Fresh | 0.34166667 |
| ASHFInt | Basidiobolaceae | Fresh | 0.33333333 |
| ASHFInt | Dothideaceae | Fresh | 0.275 |
| ASHFInt | Cordycipitaceae | Fresh | 0.27083333 |
| ASHFInt | Phanerochaetaceae | Fresh | 0.27083333 |
| ASHFInt | Ganodermataceae | Fresh | 0.26666667 |
| ASHFInt | Montagnulaceae | Fresh | 0.26666667 |
| ASHFInt | Tremellales_family_Ince | Fresh | 0.2625 |

| | | | |
|---|---|---|---|
| ASHFInt | Sclerotiniaceae | Fresh | 0.25416667 |
| ASHFInt | Debaryomycetaceae | Fresh | 0.24583333 |
| ASHFInt | Botryosphaeriaceae | Fresh | 0.2375 |
| ASHFInt | Ascomycota_family_Inc | Fresh | 0.2375 |
| ASHFInt | Agaricomycetes_unclass | Fresh | 0.22916667 |
| ASHFInt | Mucorales_unclassified | Fresh | 0.20416667 |
| ASHFInt | Capnodiales_unclassified | Fresh | 0.19583333 |
| ASHFInt | Inocybaceae | Fresh | 0.1875 |
| ASHFInt | Hypocreales_family_Inc | Fresh | 0.17083333 |
| ASHFInt | Suillaceae | Fresh | 0.1625 |
| ASHFInt | Ceratobasidiaceae | Fresh | 0.15833333 |
| ASHFInt | Diatrypaceae | Fresh | 0.15416667 |
| ASHFInt | Annulatascaceae | Fresh | 0.15 |
| ASHFInt | Dothideomycetes_unclas | Fresh | 0.14166667 |
| ASHFInt | Backusellaceae | Fresh | 0.1375 |
| ASHFInt | Gnomoniaceae | Fresh | 0.1375 |
| ASHFInt | Microascaceae | Fresh | 0.12083333 |
| ASHFInt | Botryobasidiaceae | Fresh | 0.1125 |
| ASHFInt | Umbelopsidaceae | Fresh | 0.10416667 |
| ASHFInt | Eurotiomycetes_family_I | Fresh | 0.10416667 |
| ASHFInt | Hymenochaetaceae | Fresh | 0.09583333 |
| ASHFInt | Cunninghamellaceae | Fresh | 0.09166667 |
| ASHFInt | Capnodiales_family_Inc | Fresh | 0.09166667 |
| ASHFInt | Hymenochaetales_family | Fresh | 0.09166667 |
| ASHFInt | Mycocaliciaceae | Fresh | 0.0875 |
| ASHFInt | Valsaceae | Fresh | 0.0875 |
| ASHFInt | Nectriaceae | Fresh | 0.08333333 |
| ASHFInt | Pleosporaceae | Fresh | 0.08333333 |
| ASHFInt | Hydnaceae | Fresh | 0.07916667 |

| | | | |
|---|---|---|---|
| ASHFInt | Lophiostomataceae | Fresh | 0.075 |
| ASHFInt | Agaricales_unclassified | Fresh | 0.075 |
| ASHFInt | Cantharellales_unclassifi | Fresh | 0.075 |
| ASHFInt | Graphostromataceae | Fresh | 0.06666667 |
| ASHFInt | Hyaloscyphaceae | Fresh | 0.06666667 |
| ASHFInt | Leptosphaeriaceae | Fresh | 0.0625 |
| ASHFInt | Massarinaceae | Fresh | 0.0625 |
| ASHFInt | Teratosphaeriaceae | Fresh | 0.0625 |
| ASHFInt | unclassified_Corticiales | Fresh | 0.0625 |
| ASHFInt | Diaporthales_family_Inc | Fresh | 0.0625 |
| ASHFInt | Myxotrichaceae | Fresh | 0.0625 |
| ASHFInt | Phaeosphaeriaceae | Fresh | 0.0625 |
| ASHFInt | Polyporales_family_Ince | Fresh | 0.0625 |
| ASHFInt | Pluteaceae | Fresh | 0.05833333 |
| ASHFInt | Trichosphaeriales_family | Fresh | 0.05833333 |
| ASHFInt | Chaetothyriales_unclassi | Fresh | 0.05833333 |
| ASHFInt | Glomerellaceae | Fresh | 0.05416667 |
| ASHFInt | Ophiostomataceae | Fresh | 0.05416667 |
| ASHFInt | Cucurbitariaceae | Fresh | 0.05 |
| ASHFInt | Fomitopsidaceae | Fresh | 0.05 |
| ASHFInt | Auriculariales_family_In | Fresh | 0.04583333 |
| ASHFInt | Helotiales_family_Incert | Fresh | 0.04583333 |
| ASHFInt | unclassified_Atheliales | Fresh | 0.04583333 |
| ASHFInt | Melampsoraceae | Fresh | 0.04583333 |
| ASHFInt | Lecanorales_unclassified | Fresh | 0.04583333 |
| ASHFInt | Clavicipitaceae | Fresh | 0.04166667 |
| ASHFInt | Trichosporonaceae | Fresh | 0.04166667 |
| ASHFInt | Auriscalpiaceae | Fresh | 0.04166667 |
| ASHFInt | unclassified_Pleosporale | Fresh | 0.04166667 |

| | | | |
|---|---|---|---|
| ASHFInt | Atheliaceae | Fresh | 0.0375 |
| ASHFInt | Cephalothecaceae | Fresh | 0.0375 |
| ASHFInt | Agaricaceae | Fresh | 0.03333333 |
| ASHFInt | Ophiocordycipitaceae | Fresh | 0.03333333 |
| ASHFInt | Vibrisseaceae | Fresh | 0.03333333 |
| ASHFInt | Tremellomycetes_unclas | Fresh | 0.03333333 |
| ASHFInt | Diaporthaceae | Fresh | 0.03333333 |
| ASHFInt | Kickxellaceae | Fresh | 0.03333333 |
| ASHFInt | Onygenaceae | Fresh | 0.03333333 |
| ASHFInt | Cystofilobasidiaceae | Fresh | 0.02916667 |
| ASHFInt | Dothideomycetes_family | Fresh | 0.02916667 |
| ASHFInt | Agaricostilbaceae | Fresh | 0.02916667 |
| ASHFInt | Entolomataceae | Fresh | 0.02916667 |
| ASHFInt | Schizophyllaceae | Fresh | 0.02916667 |
| ASHFInt | Boletaceae | Fresh | 0.025 |
| ASHFInt | Parmeliaceae | Fresh | 0.025 |
| ASHFInt | Physciaceae | Fresh | 0.025 |
| ASHFInt | Spizellomycetaceae | Fresh | 0.025 |
| ASHFInt | Ustilaginaceae | Fresh | 0.025 |
| ASHFInt | unclassified_Dothideales | Fresh | 0.025 |
| ASHFInt | Russulales_unclassified | Fresh | 0.025 |
| ASHFInt | Erythrobasidiales_family | Fresh | 0.025 |
| ASHFInt | unclassified_Saccharomy | Fresh | 0.02083333 |
| ASHFInt | Chaetosphaeriaceae | Fresh | 0.02083333 |
| ASHFInt | Dermateaceae | Fresh | 0.02083333 |
| ASHFInt | Pseudeurotiaceae | Fresh | 0.02083333 |
| ASHFInt | Sordariomycetes_family_ | Fresh | 0.02083333 |
| ASHFInt | Helotiales_unclassified | Fresh | 0.02083333 |
| ASHFInt | Sebacinales_unclassified | Fresh | 0.02083333 |

| | | | |
|---|---|---|---|
| ASHFInt | Agaricostilbomycetes_ur | Fresh | 0.01666667 |
| ASHFInt | Lecanoromycetes_unclas | Fresh | 0.01666667 |
| ASHFInt | Septobasidiaceae | Fresh | 0.01666667 |
| ASHFInt | Diaporthales_unclassifie | Fresh | 0.01666667 |
| ASHFInt | Bulgariaceae | Fresh | 0.01666667 |
| ASHFInt | unclassified_Trechispora | Fresh | 0.01666667 |
| ASHFInt | Pucciniales_unclassified | Fresh | 0.01666667 |
| ASHFInt | Boletinellaceae | Fresh | 0.01666667 |
| ASHFInt | Bondarzewiaceae | Fresh | 0.01666667 |
| ASHFInt | Cronartiaceae | Fresh | 0.01666667 |
| ASHFInt | unclassified_Hypocreale: | Fresh | 0.01666667 |
| ASHFInt | Boletales_unclassified | Fresh | 0.01666667 |
| ASHFInt | Agaricomycetes_family_ | Fresh | 0.0125 |
| ASHFInt | Calosphaeriaceae | Fresh | 0.0125 |
| ASHFInt | Marasmiaceae | Fresh | 0.0125 |
| ASHFInt | Plectosphaerellaceae | Fresh | 0.0125 |
| ASHFInt | Rutstroemiaceae | Fresh | 0.0125 |
| ASHFInt | Schizoparmaceae | Fresh | 0.0125 |
| ASHFInt | Wallemiaceae | Fresh | 0.0125 |
| ASHFInt | unclassified_Exobasidior | Fresh | 0.0125 |
| ASHFInt | unclassified_Taphrinales | Fresh | 0.0125 |
| ASHFInt | Pucciniomycetes_unclass | Fresh | 0.00833333 |
| ASHFInt | Candelariaceae | Fresh | 0.00833333 |
| ASHFInt | Corticiaceae | Fresh | 0.00833333 |
| ASHFInt | Cortinariaceae | Fresh | 0.00833333 |
| ASHFInt | Exobasidiomycetes_fami | Fresh | 0.00833333 |
| ASHFInt | Microstromataceae | Fresh | 0.00833333 |
| ASHFInt | Sarcosomataceae | Fresh | 0.00833333 |
| ASHFInt | Tubeufiaceae | Fresh | 0.00833333 |

| | | | |
|---|---|---|---|
| ASHFInt | unclassified_Agaricostilt | Fresh | 0.00833333 |
| ASHFInt | unclassified_Polyporales | Fresh | 0.00833333 |
| ASHFInt | Cystofilobasidiales_fami | Fresh | 0.00833333 |
| ASHFInt | Elaphomycetaceae | Fresh | 0.00833333 |
| ASHFInt | Pezizaceae | Fresh | 0.00833333 |
| ASHFInt | Pyronemataceae | Fresh | 0.00833333 |
| ASHFInt | unclassified_Atractiellale | Fresh | 0.00833333 |
| ASHFInt | unclassified_Cantharellal | Fresh | 0.00833333 |
| ASHFInt | Bolbitiaceae | Fresh | 0.00833333 |
| ASHFInt | Saccharomycopsidaceae | Fresh | 0.00833333 |
| ASHFInt | Agaricales_family_Incer | Fresh | 0.00416667 |
| ASHFInt | Ceratostomataceae | Fresh | 0.00416667 |
| ASHFInt | Clavulinaceae | Fresh | 0.00416667 |
| ASHFInt | Coniochaetales_family_I | Fresh | 0.00416667 |
| ASHFInt | Melanconidaceae | Fresh | 0.00416667 |
| ASHFInt | Mycenaceae | Fresh | 0.00416667 |
| ASHFInt | Stictidaceae | Fresh | 0.00416667 |
| ASHFInt | Thelephoraceae | Fresh | 0.00416667 |
| ASHFInt | Tricholomataceae | Fresh | 0.00416667 |
| ASHFInt | Tuberaceae | Fresh | 0.00416667 |
| ASHFInt | Xylariales_family_Incert | Fresh | 0.00416667 |
| ASHFInt | Kickxellales_unclassifiec | Fresh | 0.00416667 |
| ASHFInt | Myriangiales_unclassifie | Fresh | 0.00416667 |
| ASHFInt | Chytridiomycetes_unclas | Fresh | 0.00416667 |
| ASHFInt | Apiosporaceae | Fresh | 0.00416667 |
| ASHFInt | Chaetothyriaceae | Fresh | 0.00416667 |
| ASHFInt | Chaetothyriales_family_l | Fresh | 0.00416667 |
| ASHFInt | Chionosphaeraceae | Fresh | 0.00416667 |
| ASHFInt | Gyroporaceae | Fresh | 0.00416667 |

| | | | |
|---|---|---|---|
| ASHFInt | Holtermanniales_family_ | Fresh | 0.00416667 |
| ASHFInt | Niaceae | Fresh | 0.00416667 |
| ASHFInt | Orbiliaceae | Fresh | 0.00416667 |
| ASHFInt | Phragmidiaceae | Fresh | 0.00416667 |
| ASHFInt | Pichiaceae | Fresh | 0.00416667 |
| ASHFInt | Pucciniastraceae | Fresh | 0.00416667 |
| ASHFInt | Russulaceae | Fresh | 0.00416667 |
| ASHFInt | Sporormiaceae | Fresh | 0.00416667 |
| ASHFInt | Xenasmataceae | Fresh | 0.00416667 |
| ASHFInt | unclassified_Agaricomy | Fresh | 0.00416667 |
| ASHFInt | unclassified_Eurotiomyc | Fresh | 0.00416667 |
| ASHFInt | unclassified_Fungi | Fresh | 0.00416667 |
| ASHFInt | unclassified_Leotiomyce | Fresh | 0.00416667 |
| ASHFInt | Xylariales_unclassified | Fresh | 0.00416667 |
| WMCCExt | Peniophoraceae | Fresh | 10.8333333 |
| WMCCExt | Davidiellaceae | Fresh | 7.35666667 |
| WMCCExt | Ascomycota_unclassified | Fresh | 6.62 |
| WMCCExt | Saccharomycetaceae | Fresh | 6.02333333 |
| WMCCExt | Metschnikowiaceae | Fresh | 5.53666667 |
| WMCCExt | Meruliaceae | Fresh | 5.47666667 |
| WMCCExt | Sporidiobolales_family_ | Fresh | 3.22333333 |
| WMCCExt | Schizoporaceae | Fresh | 2.68 |
| WMCCExt | Mycosphaerellaceae | Fresh | 2.37666667 |
| WMCCExt | Dothideaceae | Fresh | 1.92666667 |
| WMCCExt | Pleosporales_family_Inc | Fresh | 1.92 |
| WMCCExt | Polyporaceae | Fresh | 1.91666667 |
| WMCCExt | Ganodermataceae | Fresh | 1.56666667 |
| WMCCExt | unclassified_Plantae | Fresh | 1.5 |
| WMCCExt | Meripilaceae | Fresh | 1.46 |

| | | | |
|---|---|---|---|
| WMCCExt | Fungi_unclassified | Fresh | 1.45333333 |
| WMCCExt | Strophariaceae | Fresh | 1.44333333 |
| WMCCExt | Dothioraceae | Fresh | 1.38333333 |
| WMCCExt | Pleosporales_unclassifie | Fresh | 1.37 |
| WMCCExt | Basidiomycota_unclassif | Fresh | 1.19666667 |
| WMCCExt | Hymenochaetales_family | Fresh | 1.18333333 |
| WMCCExt | Psathyrellaceae | Fresh | 1.16 |
| WMCCExt | Agaricomycetes_unclass | Fresh | 1.14 |
| WMCCExt | Eurotiomycetes_unclassi | Fresh | 1.10333333 |
| WMCCExt | Saccharomycetales_fami | Fresh | 1.10333333 |
| WMCCExt | Taphrinaceae | Fresh | 1.08666667 |
| WMCCExt | Montagnulaceae | Fresh | 0.90333333 |
| WMCCExt | Phanerochaetaceae | Fresh | 0.78 |
| WMCCExt | Polyporales_unclassified | Fresh | 0.75666667 |
| WMCCExt | Inocybaceae | Fresh | 0.72 |
| WMCCExt | Sordariomycetes_unclass | Fresh | 0.71 |
| WMCCExt | Hydnodontaceae | Fresh | 0.68333333 |
| WMCCExt | unclassified_Corticiales | Fresh | 0.66666667 |
| WMCCExt | Agaricales_unclassified | Fresh | 0.64 |
| WMCCExt | Phaeosphaeriaceae | Fresh | 0.63 |
| WMCCExt | Suillaceae | Fresh | 0.57666667 |
| WMCCExt | Sclerotiniaceae | Fresh | 0.57 |
| WMCCExt | Tremellales_family_Ince | Fresh | 0.56 |
| WMCCExt | Pleosporaceae | Fresh | 0.49333333 |
| WMCCExt | Ceratobasidiaceae | Fresh | 0.48 |
| WMCCExt | Xylariaceae | Fresh | 0.45666667 |
| WMCCExt | Fomitopsidaceae | Fresh | 0.45333333 |
| WMCCExt | Amphisphaeriaceae | Fresh | 0.44666667 |
| WMCCExt | Capnodiales_unclassifiec | Fresh | 0.43333333 |

| | | | |
|---|---|---|---|
| WMCCExt | Cantharellales_unclassifi | Fresh | 0.42666667 |
| WMCCExt | Diatrypaceae | Fresh | 0.42666667 |
| WMCCExt | Capnodiales_family_Ince | Fresh | 0.42 |
| WMCCExt | Stereaceae | Fresh | 0.41333333 |
| WMCCExt | Hymenochaetaceae | Fresh | 0.40333333 |
| WMCCExt | Trichocomaceae | Fresh | 0.39666667 |
| WMCCExt | Physciaceae | Fresh | 0.37333333 |
| WMCCExt | Hydnaceae | Fresh | 0.34 |
| WMCCExt | Gnomoniaceae | Fresh | 0.32666667 |
| WMCCExt | Mucoraceae | Fresh | 0.31 |
| WMCCExt | Venturiaceae | Fresh | 0.30333333 |
| WMCCExt | Cucurbitariaceae | Fresh | 0.28333333 |
| WMCCExt | Exobasidiaceae | Fresh | 0.27 |
| WMCCExt | Schizophyllaceae | Fresh | 0.27 |
| WMCCExt | Auriculariales_family_In | Fresh | 0.26666667 |
| WMCCExt | Polyporales_family_Ince | Fresh | 0.26333333 |
| WMCCExt | Lophiostomataceae | Fresh | 0.26 |
| WMCCExt | Herpotrichiellaceae | Fresh | 0.24333333 |
| WMCCExt | Helotiales_family_Incert | Fresh | 0.24333333 |
| WMCCExt | Clavulinaceae | Fresh | 0.23666667 |
| WMCCExt | Entolomataceae | Fresh | 0.23666667 |
| WMCCExt | Auriscalpiaceae | Fresh | 0.19666667 |
| WMCCExt | Botryosphaeriaceae | Fresh | 0.19333333 |
| WMCCExt | Valsaceae | Fresh | 0.19 |
| WMCCExt | Hypocreales_family_Ince | Fresh | 0.18666667 |
| WMCCExt | Capnodiaceae | Fresh | 0.17333333 |
| WMCCExt | Sporormiaceae | Fresh | 0.17 |
| WMCCExt | Hypocreales_unclassified | Fresh | 0.16666667 |
| WMCCExt | Tricholomataceae | Fresh | 0.15666667 |

| | | | |
|---|---|---|---|
| WMCCExt | Ascomycota_family_Inc | Fresh | 0.15333333 |
| WMCCExt | Filobasidiaceae | Fresh | 0.15333333 |
| WMCCExt | Leptosphaeriaceae | Fresh | 0.15 |
| WMCCExt | Nectriaceae | Fresh | 0.14666667 |
| WMCCExt | Cordycipitaceae | Fresh | 0.14333333 |
| WMCCExt | Pluteaceae | Fresh | 0.14333333 |
| WMCCExt | Teratosphaeriaceae | Fresh | 0.14333333 |
| WMCCExt | Diaporthales_family_Inc | Fresh | 0.14 |
| WMCCExt | Marasmiaceae | Fresh | 0.13666667 |
| WMCCExt | Mortierellaceae | Fresh | 0.13666667 |
| WMCCExt | Atheliaceae | Fresh | 0.13333333 |
| WMCCExt | Didymosphaeriaceae | Fresh | 0.13333333 |
| WMCCExt | Lecanorales_unclassified | Fresh | 0.13333333 |
| WMCCExt | Chaetomiaceae | Fresh | 0.13 |
| WMCCExt | Chaetothyriaceae | Fresh | 0.12666667 |
| WMCCExt | Russulales_unclassified | Fresh | 0.12333333 |
| WMCCExt | Dothideomycetes_unclas | Fresh | 0.12333333 |
| WMCCExt | unclassified_Pleosporale | Fresh | 0.12333333 |
| WMCCExt | Botryobasidiaceae | Fresh | 0.11333333 |
| WMCCExt | Massarinaceae | Fresh | 0.10666667 |
| WMCCExt | Agaricostilbaceae | Fresh | 0.10333333 |
| WMCCExt | Melampsoraceae | Fresh | 0.1 |
| WMCCExt | Glomerellaceae | Fresh | 0.09666667 |
| WMCCExt | Trichosphaeriales_family | Fresh | 0.09 |
| WMCCExt | Hypocreaceae | Fresh | 0.09 |
| WMCCExt | Hebelomataceae | Fresh | 0.08666667 |
| WMCCExt | Mycenaceae | Fresh | 0.08666667 |
| WMCCExt | Mycocaliciaceae | Fresh | 0.08 |
| WMCCExt | Umbelopsidaceae | Fresh | 0.08 |

| | | | |
|---|---|---|---|
| WMCCExt | Eurotiomycetes_family_I | Fresh | 0.07666667 |
| WMCCExt | Helotiales_unclassified | Fresh | 0.07666667 |
| WMCCExt | unclassified_Atheliales | Fresh | 0.07333333 |
| WMCCExt | Pleurotaceae | Fresh | 0.07 |
| WMCCExt | Ustilaginaceae | Fresh | 0.06666667 |
| WMCCExt | Dothideomycetes_family | Fresh | 0.06666667 |
| WMCCExt | Phallaceae | Fresh | 0.06666667 |
| WMCCExt | Xenasmataceae | Fresh | 0.06333333 |
| WMCCExt | Corticiaceae | Fresh | 0.06 |
| WMCCExt | Annulatascaceae | Fresh | 0.06 |
| WMCCExt | Chaetothyriales_unclassi | Fresh | 0.06 |
| WMCCExt | Tremellomycetes_unclas | Fresh | 0.05666667 |
| WMCCExt | Thelephoraceae | Fresh | 0.05666667 |
| WMCCExt | unclassified_Exobasidiomy | Fresh | 0.05666667 |
| WMCCExt | unclassified_Polyporales | Fresh | 0.05333333 |
| WMCCExt | Cortinariaceae | Fresh | 0.05333333 |
| WMCCExt | Cystofilobasidiaceae | Fresh | 0.05333333 |
| WMCCExt | Exobasidiomycetes_fami | Fresh | 0.05333333 |
| WMCCExt | Agaricaceae | Fresh | 0.05 |
| WMCCExt | Agaricomycetes_family_ | Fresh | 0.05 |
| WMCCExt | Clavicipitaceae | Fresh | 0.05 |
| WMCCExt | Cephalothecaceae | Fresh | 0.05 |
| WMCCExt | Erythrobasidiales_family | Fresh | 0.04666667 |
| WMCCExt | Halosphaeriaceae | Fresh | 0.04333333 |
| WMCCExt | Leotiomycetes_family_In | Fresh | 0.04333333 |
| WMCCExt | Sordariomycetes_family_ | Fresh | 0.04333333 |
| WMCCExt | Xylariales_unclassified | Fresh | 0.04333333 |
| WMCCExt | Teloschistales_unclassifi | Fresh | 0.04 |
| WMCCExt | Leotiomycetes_unclassif | Fresh | 0.04 |

| | | | |
|---|---|---|---|
| WMCCExt | Vibrisseaceae | Fresh | 0.04 |
| WMCCExt | Pucciniomycetes_unclass | Fresh | 0.03666667 |
| WMCCExt | Plectosphaerellaceae | Fresh | 0.03666667 |
| WMCCExt | Diaporthaceae | Fresh | 0.03666667 |
| WMCCExt | Graphostromataceae | Fresh | 0.03666667 |
| WMCCExt | Atheliales_unclassified | Fresh | 0.03666667 |
| WMCCExt | Lachnocladiaceae | Fresh | 0.03666667 |
| WMCCExt | unclassified_Agaricostilb | Fresh | 0.03333333 |
| WMCCExt | Verrucariales_family_Inc | Fresh | 0.03333333 |
| WMCCExt | Boletales_unclassified | Fresh | 0.03333333 |
| WMCCExt | Boletaceae | Fresh | 0.03333333 |
| WMCCExt | Spizellomycetaceae | Fresh | 0.03333333 |
| WMCCExt | Bondarzewiaceae | Fresh | 0.03 |
| WMCCExt | Parmeliaceae | Fresh | 0.03 |
| WMCCExt | Togniniaceae | Fresh | 0.03 |
| WMCCExt | Gomphaceae | Fresh | 0.03 |
| WMCCExt | Myriangiales_family_Inc | Fresh | 0.03 |
| WMCCExt | unclassified_Trechispora | Fresh | 0.02666667 |
| WMCCExt | Chaetosphaeriaceae | Fresh | 0.02666667 |
| WMCCExt | Lyophyllaceae | Fresh | 0.02666667 |
| WMCCExt | Microascaceae | Fresh | 0.02666667 |
| WMCCExt | Serpulaceae | Fresh | 0.02666667 |
| WMCCExt | Saccharomycetales_uncla | Fresh | 0.02333333 |
| WMCCExt | Hyaloscyphaceae | Fresh | 0.02333333 |
| WMCCExt | Russulaceae | Fresh | 0.02333333 |
| WMCCExt | Xylariales_family_Incert | Fresh | 0.02333333 |
| WMCCExt | unclassified_Atractiellal | Fresh | 0.02333333 |
| WMCCExt | Lecanoromycetes_unclas | Fresh | 0.02 |
| WMCCExt | Gymnoascaceae | Fresh | 0.02 |

| | | | |
|---|---|---|---|
| WMCCExt | Wallemiaceae | Fresh | 0.02 |
| WMCCExt | Sebacinales_unclassified | Fresh | 0.02 |
| WMCCExt | Rhytismataceae | Fresh | 0.02 |
| WMCCExt | Diaporthales_unclassifie | Fresh | 0.02 |
| WMCCExt | Agaricales_family_Incer | Fresh | 0.01666667 |
| WMCCExt | Microstromataceae | Fresh | 0.01666667 |
| WMCCExt | Bolbitiaceae | Fresh | 0.01666667 |
| WMCCExt | Elsinoaceae | Fresh | 0.01666667 |
| WMCCExt | Geastraceae | Fresh | 0.01666667 |
| WMCCExt | Phragmidiaceae | Fresh | 0.01666667 |
| WMCCExt | Pseudeurotiaceae | Fresh | 0.01666667 |
| WMCCExt | Pucciniastraceae | Fresh | 0.01666667 |
| WMCCExt | Tuberaceae | Fresh | 0.01666667 |
| WMCCExt | Mucorales_unclassified | Fresh | 0.01666667 |
| WMCCExt | Hygrophoraceae | Fresh | 0.01666667 |
| WMCCExt | unclassified_Tremellales | Fresh | 0.01666667 |
| WMCCExt | Sordariales_unclassified | Fresh | 0.01666667 |
| WMCCExt | Septobasidiaceae | Fresh | 0.01333333 |
| WMCCExt | Apiosporaceae | Fresh | 0.01333333 |
| WMCCExt | Calosphaeriaceae | Fresh | 0.01333333 |
| WMCCExt | Lecanoraceae | Fresh | 0.01333333 |
| WMCCExt | Myxotrichaceae | Fresh | 0.01333333 |
| WMCCExt | Physalacriaceae | Fresh | 0.01333333 |
| WMCCExt | unclassified_Dothideales | Fresh | 0.01333333 |
| WMCCExt | Pucciniales_unclassified | Fresh | 0.01333333 |
| WMCCExt | Taphrinales_unclassified | Fresh | 0.01333333 |
| WMCCExt | Chytridiomycetes_unclas | Fresh | 0.01333333 |
| WMCCExt | Cystofilobasidiales_fami | Fresh | 0.01333333 |
| WMCCExt | unclassified_Taphrinales | Fresh | 0.01333333 |

| | | | |
|---|---|---|---|
| WMCCExt | Lulworthiaceae | Fresh | 0.01 |
| WMCCExt | Cladoniaceae | Fresh | 0.01 |
| WMCCExt | Eurotiales_family_Incert | Fresh | 0.01 |
| WMCCExt | Melanconidaceae | Fresh | 0.01 |
| WMCCExt | Ophiocordycipitaceae | Fresh | 0.01 |
| WMCCExt | Quambalariaceae | Fresh | 0.01 |
| WMCCExt | Sordariaceae | Fresh | 0.01 |
| WMCCExt | Typhulaceae | Fresh | 0.01 |
| WMCCExt | unclassified_Cantharellal | Fresh | 0.01 |
| WMCCExt | Thelephorales_unclassifi | Fresh | 0.01 |
| WMCCExt | Agaricostilbomycetes_ur | Fresh | 0.00666667 |
| WMCCExt | Bankeraceae | Fresh | 0.00666667 |
| WMCCExt | Basidiobolaceae | Fresh | 0.00666667 |
| WMCCExt | Basidiomycota_family_I | Fresh | 0.00666667 |
| WMCCExt | Boliniaceae | Fresh | 0.00666667 |
| WMCCExt | Chaetothyriales_family_ | Fresh | 0.00666667 |
| WMCCExt | Chionosphaeraceae | Fresh | 0.00666667 |
| WMCCExt | Coniochaetaceae | Fresh | 0.00666667 |
| WMCCExt | Cronartiaceae | Fresh | 0.00666667 |
| WMCCExt | Hygrophoropsidaceae | Fresh | 0.00666667 |
| WMCCExt | Lasiosphaeriaceae | Fresh | 0.00666667 |
| WMCCExt | Niaceae | Fresh | 0.00666667 |
| WMCCExt | Onygenaceae | Fresh | 0.00666667 |
| WMCCExt | Paxillaceae | Fresh | 0.00666667 |
| WMCCExt | Pterulaceae | Fresh | 0.00666667 |
| WMCCExt | Saccharomycopsidaceae | Fresh | 0.00666667 |
| WMCCExt | Schizoparmaceae | Fresh | 0.00666667 |
| WMCCExt | Sebacinaceae | Fresh | 0.00666667 |
| WMCCExt | Sirobasidiaceae | Fresh | 0.00666667 |

| | | | |
|---|---|---|---|
| WMCCExt | Stephanosporaceae | Fresh | 0.00666667 |
| WMCCExt | Stictidaceae | Fresh | 0.00666667 |
| WMCCExt | Tilletiaceae | Fresh | 0.00666667 |
| WMCCExt | Tubeufiaceae | Fresh | 0.00666667 |
| WMCCExt | unclassified_Pezizales | Fresh | 0.00666667 |
| WMCCExt | Arachnomycetaceae | Fresh | 0.00666667 |
| WMCCExt | Entomophthoraceae | Fresh | 0.00666667 |
| WMCCExt | Glomeraceae | Fresh | 0.00666667 |
| WMCCExt | Pucciniaceae | Fresh | 0.00666667 |
| WMCCExt | Cystofilobasidiales_uncla | Fresh | 0.00666667 |
| WMCCExt | Acarosporaceae | Fresh | 0.00333333 |
| WMCCExt | Arthrodermataceae | Fresh | 0.00333333 |
| WMCCExt | Bionectriaceae | Fresh | 0.00333333 |
| WMCCExt | Caliciaceae | Fresh | 0.00333333 |
| WMCCExt | Candelariaceae | Fresh | 0.00333333 |
| WMCCExt | Cantharellales_family_In | Fresh | 0.00333333 |
| WMCCExt | Coniophoraceae | Fresh | 0.00333333 |
| WMCCExt | Eocronartiaceae | Fresh | 0.00333333 |
| WMCCExt | Gyroporaceae | Fresh | 0.00333333 |
| WMCCExt | Hericiaceae | Fresh | 0.00333333 |
| WMCCExt | Lentitheciaceae | Fresh | 0.00333333 |
| WMCCExt | Pleomassariaceae | Fresh | 0.00333333 |
| WMCCExt | Ramalinaceae | Fresh | 0.00333333 |
| WMCCExt | Sarcosomataceae | Fresh | 0.00333333 |
| WMCCExt | Tapinellaceae | Fresh | 0.00333333 |
| WMCCExt | unclassified_Agaricomyc | Fresh | 0.00333333 |
| WMCCExt | unclassified_Cystofiloba | Fresh | 0.00333333 |
| WMCCExt | unclassified_Entylomatal | Fresh | 0.00333333 |
| WMCCExt | unclassified_Fungi | Fresh | 0.00333333 |

| | | | |
|---|---|---|---|
| WMCCExt | unclassified_Hypocreales | Fresh | 0.00333333 |
| WMCCExt | unclassified_Hysteriales | Fresh | 0.00333333 |
| WMCCExt | unclassified_Leotiomyce | Fresh | 0.00333333 |
| WMCCExt | unclassified_Saccharomy | Fresh | 0.00333333 |
| WMCCExt | Corticiales_unclassified | Fresh | 0.00333333 |
| WMCCExt | Myriangiales_unclassifie | Fresh | 0.00333333 |
| WMCCExt | Rhizophydiales_unclassi | Fresh | 0.00333333 |
| WMCCExt | Discinaceae | Fresh | 0.00333333 |
| WMCCExt | Helotiaceae | Fresh | 0.00333333 |
| WMCCExt | Microbotryomycetes_fan | Fresh | 0.00333333 |
| WMCCExt | Rutstroemiaceae | Fresh | 0.00333333 |
| WMCCExt | Stereocaulaceae | Fresh | 0.00333333 |
| WMCCExt | Tulasnellaceae | Fresh | 0.00333333 |
| ASHFExt | Sporidiobolales_family_ | W12 | 45.25 |
| ASHFExt | Pleosporales_family_Inc | W12 | 15.8833333 |
| ASHFExt | Nectriaceae | W12 | 13.3666667 |
| ASHFExt | Davidiellaceae | W12 | 10.25 |
| ASHFExt | Pleosporaceae | W12 | 4.28333333 |
| ASHFExt | Lasiosphaeriaceae | W12 | 2.88333333 |
| ASHFExt | Phaeosphaeriaceae | W12 | 1.18333333 |
| ASHFExt | Pseudeurotiaceae | W12 | 0.95 |
| ASHFExt | Filobasidiaceae | W12 | 0.88333333 |
| ASHFExt | Mucoraceae | W12 | 0.81666667 |
| ASHFExt | Pleosporales_unclassified | W12 | 0.76666667 |
| ASHFExt | Mortierellaceae | W12 | 0.56666667 |
| ASHFExt | Amphisphaeriaceae | W12 | 0.55 |
| ASHFExt | Tremellales_family_Ince | W12 | 0.3 |
| ASHFExt | Montagnulaceae | W12 | 0.28333333 |
| ASHFExt | unclassified_Pleosporales | W12 | 0.28333333 |

| | | | |
|---|---|---|---|
| ASHFExt | Ascomycota_unclassified | W12 | 0.25 |
| ASHFExt | Holtermanniales_family_ | W12 | 0.23333333 |
| ASHFExt | Fungi_unclassified | W12 | 0.15 |
| ASHFExt | Rhizophlyctidaceae | W12 | 0.11666667 |
| ASHFExt | Chaetomiaceae | W12 | 0.1 |
| ASHFExt | Leptosphaeriaceae | W12 | 0.06666667 |
| ASHFExt | Dothioraceae | W12 | 0.05 |
| ASHFExt | Glomerellaceae | W12 | 0.05 |
| ASHFExt | Helotiaceae | W12 | 0.05 |
| ASHFExt | Hypocreales_family_Inc | W12 | 0.05 |
| ASHFExt | Ceratobasidiaceae | W12 | 0.03333333 |
| ASHFExt | Umbelopsidaceae | W12 | 0.03333333 |
| ASHFExt | Mucorales_unclassified | W12 | 0.03333333 |
| ASHFExt | Sporormiaceae | W12 | 0.03333333 |
| ASHFExt | unclassified_Fungi | W12 | 0.03333333 |
| ASHFExt | Tremellomycetes_unclas | W12 | 0.01666667 |
| ASHFExt | Ascomycota_family_Inc | W12 | 0.01666667 |
| ASHFExt | Cystofilobasidiaceae | W12 | 0.01666667 |
| ASHFExt | Diaporthaceae | W12 | 0.01666667 |
| ASHFExt | Didymosphaeriaceae | W12 | 0.01666667 |
| ASHFExt | Eremomycetaceae | W12 | 0.01666667 |
| ASHFExt | Herpotrichiellaceae | W12 | 0.01666667 |
| ASHFExt | Leucosporidiaceae | W12 | 0.01666667 |
| ASHFExt | Lophiostomataceae | W12 | 0.01666667 |
| ASHFExt | Magnaporthaceae | W12 | 0.01666667 |
| ASHFExt | Spizellomycetaceae | W12 | 0.01666667 |
| ASHFExt | Xylariales_family_Incert | W12 | 0.01666667 |
| ASHFExt | Basidiomycota_unclassif | W12 | 0.01666667 |
| ASHFInt | Trichocomaceae | W12 | 41.0333333 |

| | | | |
|---|---|---|---|
| ASHFInt | Wallemiaceae | W12 | 18.7 |
| ASHFInt | Ascomycota_unclassified | W12 | 8.03333333 |
| ASHFInt | unclassified_Plantae | W12 | 5.76666667 |
| ASHFInt | Pleosporales_family_Inc | W12 | 3.2 |
| ASHFInt | Davidiellaceae | W12 | 2.08333333 |
| ASHFInt | Sporidiobolales_family_I | W12 | 2.06666667 |
| ASHFInt | Fungi_unclassified | W12 | 2.03333333 |
| ASHFInt | Saccharomycetaceae | W12 | 1.55 |
| ASHFInt | Meruliaceae | W12 | 1.43333333 |
| ASHFInt | Taphrinaceae | W12 | 1.15 |
| ASHFInt | Polyporaceae | W12 | 0.68333333 |
| ASHFInt | Dothideaceae | W12 | 0.66666667 |
| ASHFInt | Agaricomycetes_unclassi | W12 | 0.6 |
| ASHFInt | Peniophoraceae | W12 | 0.46666667 |
| ASHFInt | Pleosporales_unclassified | W12 | 0.46666667 |
| ASHFInt | Stereaceae | W12 | 0.41666667 |
| ASHFInt | Tremellales_family_Ince | W12 | 0.4 |
| ASHFInt | Basidiomycota_unclassif | W12 | 0.38333333 |
| ASHFInt | Schizoporaceae | W12 | 0.35 |
| ASHFInt | Dothioraceae | W12 | 0.35 |
| ASHFInt | Hydnodontaceae | W12 | 0.35 |
| ASHFInt | Hymenochaetales_family | W12 | 0.35 |
| ASHFInt | Leptosphaeriaceae | W12 | 0.3 |
| ASHFInt | Ophiostomataceae | W12 | 0.3 |
| ASHFInt | Phanerochaetaceae | W12 | 0.25 |
| ASHFInt | Microascaceae | W12 | 0.23333333 |
| ASHFInt | Pleosporaceae | W12 | 0.23333333 |
| ASHFInt | Venturiaceae | W12 | 0.23333333 |
| ASHFInt | Eurotiomycetes_unclassi | W12 | 0.21666667 |

| | | | |
|---|---|---|---|
| ASHFInt | Hymenochaetaceae | W12 | 0.21666667 |
| ASHFInt | Mucoraceae | W12 | 0.21666667 |
| ASHFInt | Meripilaceae | W12 | 0.2 |
| ASHFInt | Marasmiaceae | W12 | 0.18333333 |
| ASHFInt | Strophariaceae | W12 | 0.18333333 |
| ASHFInt | Filobasidiaceae | W12 | 0.16666667 |
| ASHFInt | Mycosphaerellaceae | W12 | 0.16666667 |
| ASHFInt | Cystofilobasidiaceae | W12 | 0.16666667 |
| ASHFInt | Exobasidiaceae | W12 | 0.15 |
| ASHFInt | Glomerellaceae | W12 | 0.13333333 |
| ASHFInt | Herpotrichiellaceae | W12 | 0.13333333 |
| ASHFInt | Fomitopsidaceae | W12 | 0.11666667 |
| ASHFInt | Psathyrellaceae | W12 | 0.11666667 |
| ASHFInt | Sclerotiniaceae | W12 | 0.11666667 |
| ASHFInt | Agaricomycetes_family_ | W12 | 0.1 |
| ASHFInt | Amphisphaeriaceae | W12 | 0.1 |
| ASHFInt | Ascomycota_family_Inc | W12 | 0.1 |
| ASHFInt | Capnodiales_family_Inc | W12 | 0.1 |
| ASHFInt | Cordycipitaceae | W12 | 0.1 |
| ASHFInt | Erysiphaceae | W12 | 0.1 |
| ASHFInt | Melampsoraceae | W12 | 0.1 |
| ASHFInt | Montagnulaceae | W12 | 0.1 |
| ASHFInt | Pleurotaceae | W12 | 0.1 |
| ASHFInt | Capnodiales_unclassified | W12 | 0.1 |
| ASHFInt | Schizophyllaceae | W12 | 0.08333333 |
| ASHFInt | Erythrobasidiales_family | W12 | 0.08333333 |
| ASHFInt | Ganodermataceae | W12 | 0.08333333 |
| ASHFInt | Xylariaceae | W12 | 0.08333333 |
| ASHFInt | Cantharellales_unclassifi | W12 | 0.08333333 |

| | | | |
|---|---|---|---|
| ASHFInt | Clavicipitaceae | W12 | 0.06666667 |
| ASHFInt | Hydnaceae | W12 | 0.06666667 |
| ASHFInt | Inocybaceae | W12 | 0.06666667 |
| ASHFInt | Pluteaceae | W12 | 0.06666667 |
| ASHFInt | Polyporales_family_Ince | W12 | 0.06666667 |
| ASHFInt | unclassified_Corticiales | W12 | 0.06666667 |
| ASHFInt | Agaricales_unclassified | W12 | 0.06666667 |
| ASHFInt | Sordariomycetes_unclass | W12 | 0.05 |
| ASHFInt | Atheliaceae | W12 | 0.05 |
| ASHFInt | Basidiobolaceae | W12 | 0.05 |
| ASHFInt | Botryobasidiaceae | W12 | 0.05 |
| ASHFInt | Mortierellaceae | W12 | 0.05 |
| ASHFInt | Polyporales_unclassified | W12 | 0.05 |
| ASHFInt | Dothideomycetes_unclas | W12 | 0.03333333 |
| ASHFInt | Agaricostilbaceae | W12 | 0.03333333 |
| ASHFInt | Auriculariales_family_In | W12 | 0.03333333 |
| ASHFInt | Calosphaeriaceae | W12 | 0.03333333 |
| ASHFInt | Clavulinaceae | W12 | 0.03333333 |
| ASHFInt | Diaporthales_family_Inc | W12 | 0.03333333 |
| ASHFInt | Elsinoaceae | W12 | 0.03333333 |
| ASHFInt | Massarinaceae | W12 | 0.03333333 |
| ASHFInt | Mycenaceae | W12 | 0.03333333 |
| ASHFInt | Pyronemataceae | W12 | 0.03333333 |
| ASHFInt | unclassified_Pleosporale | W12 | 0.03333333 |
| ASHFInt | unclassified_Trechispora | W12 | 0.03333333 |
| ASHFInt | Chaetothyriales_unclassi | W12 | 0.03333333 |
| ASHFInt | Leotiomycetes_unclassif | W12 | 0.03333333 |
| ASHFInt | Cucurbitariaceae | W12 | 0.03333333 |
| ASHFInt | Hypocreales_family_Inc | W12 | 0.03333333 |

| | | | |
|---|---|---|---|
| ASHFInt | Pseudeurotiaceae | W12 | 0.03333333 |
| ASHFInt | Ustilaginaceae | W12 | 0.03333333 |
| ASHFInt | Hypocreales_unclassifiec | W12 | 0.03333333 |
| ASHFInt | Tremellomycetes_unclas | W12 | 0.01666667 |
| ASHFInt | Amanitaceae | W12 | 0.01666667 |
| ASHFInt | Auriscalpiaceae | W12 | 0.01666667 |
| ASHFInt | Boletaceae | W12 | 0.01666667 |
| ASHFInt | Ceratobasidiaceae | W12 | 0.01666667 |
| ASHFInt | Chaetomiaceae | W12 | 0.01666667 |
| ASHFInt | Chaetothyriaceae | W12 | 0.01666667 |
| ASHFInt | Clavariaceae | W12 | 0.01666667 |
| ASHFInt | Coniochaetaceae | W12 | 0.01666667 |
| ASHFInt | Corticiaceae | W12 | 0.01666667 |
| ASHFInt | Diatrypaceae | W12 | 0.01666667 |
| ASHFInt | Dothideomycetes_family | W12 | 0.01666667 |
| ASHFInt | Entolomataceae | W12 | 0.01666667 |
| ASHFInt | Eurotiomycetes_family_] | W12 | 0.01666667 |
| ASHFInt | Helotiales_family_Incert | W12 | 0.01666667 |
| ASHFInt | Hypocreaceae | W12 | 0.01666667 |
| ASHFInt | Lachnocladiaceae | W12 | 0.01666667 |
| ASHFInt | Lyophyllaceae | W12 | 0.01666667 |
| ASHFInt | Malasseziales_family_In | W12 | 0.01666667 |
| ASHFInt | Pezizaceae | W12 | 0.01666667 |
| ASHFInt | Phaeosphaeriaceae | W12 | 0.01666667 |
| ASHFInt | Physalacriaceae | W12 | 0.01666667 |
| ASHFInt | Plectosphaerellaceae | W12 | 0.01666667 |
| ASHFInt | Rhytismataceae | W12 | 0.01666667 |
| ASHFInt | Russulaceae | W12 | 0.01666667 |
| ASHFInt | Rutstroemiaceae | W12 | 0.01666667 |

| | | | |
|---|---|---|---|
| ASHFInt | Schizoparmaceae | W12 | 0.01666667 |
| ASHFInt | Sporormiaceae | W12 | 0.01666667 |
| ASHFInt | Tricholomataceae | W12 | 0.01666667 |
| ASHFInt | Trichosphaeriales_family | W12 | 0.01666667 |
| ASHFInt | Tulasnellaceae | W12 | 0.01666667 |
| ASHFInt | unclassified_Atheliales | W12 | 0.01666667 |
| ASHFInt | unclassified_Taphrinales | W12 | 0.01666667 |
| ASHFInt | Botryosphaeriales_uncla: | W12 | 0.01666667 |
| ASHFInt | Helotiales_unclassified | W12 | 0.01666667 |
| ASHFInt | Mucorales_unclassified | W12 | 0.01666667 |
| ASHFInt | Pucciniales_unclassified | W12 | 0.01666667 |
| WMCCExt | Davidiellaceae | W12 | 29.1333333 |
| WMCCExt | Tremellales_family_Ince | W12 | 21.0833333 |
| WMCCExt | Sporidiobolales_family_] | W12 | 15.15 |
| WMCCExt | Dothioraceae | W12 | 7.56666667 |
| WMCCExt | Pleosporales_family_Inc | W12 | 5.15 |
| WMCCExt | Mucoraceae | W12 | 5.03333333 |
| WMCCExt | Spizellomycetaceae | W12 | 4.58333333 |
| WMCCExt | Pleosporaceae | W12 | 3.61666667 |
| WMCCExt | Pleosporales_unclassifiec | W12 | 1.91666667 |
| WMCCExt | Phaeosphaeriaceae | W12 | 1.45 |
| WMCCExt | Nectriaceae | W12 | 1.41666667 |
| WMCCExt | Fungi_unclassified | W12 | 0.81666667 |
| WMCCExt | Ascomycota_unclassifiec | W12 | 0.48333333 |
| WMCCExt | Tremellomycetes_unclas | W12 | 0.38333333 |
| WMCCExt | Sporormiaceae | W12 | 0.25 |
| WMCCExt | Basidiomycota_unclassif | W12 | 0.21666667 |
| WMCCExt | Lophiostomataceae | W12 | 0.2 |
| WMCCExt | Sordariaceae | W12 | 0.18333333 |

| | | | |
|---|---|---|---|
| WMCCExt | Chaetothyriaceae | W12 | 0.15 |
| WMCCExt | Helotiaceae | W12 | 0.15 |
| WMCCExt | Venturiaceae | W12 | 0.15 |
| WMCCExt | unclassified_Chytridiomy | W12 | 0.1 |
| WMCCExt | Herpotrichiellaceae | W12 | 0.08333333 |
| WMCCExt | Ascomycota_family_Inc | W12 | 0.06666667 |
| WMCCExt | Hypocreales_family_Inc | W12 | 0.06666667 |
| WMCCExt | Helotiales_unclassified | W12 | 0.06666667 |
| WMCCExt | Leotiomycetes_unclassif | W12 | 0.05 |
| WMCCExt | Cucurbitariaceae | W12 | 0.05 |
| WMCCExt | Leptosphaeriaceae | W12 | 0.05 |
| WMCCExt | Montagnulaceae | W12 | 0.05 |
| WMCCExt | Coniochaetaceae | W12 | 0.03333333 |
| WMCCExt | Tubeufiaceae | W12 | 0.03333333 |
| WMCCExt | unclassified_Pleosporale | W12 | 0.03333333 |
| WMCCExt | Polyporales_unclassified | W12 | 0.03333333 |
| WMCCExt | Sordariomycetes_unclass | W12 | 0.01666667 |
| WMCCExt | Amphisphaeriaceae | W12 | 0.01666667 |
| WMCCExt | Cystobasidiaceae | W12 | 0.01666667 |
| WMCCExt | Dothideaceae | W12 | 0.01666667 |
| WMCCExt | Filobasidiaceae | W12 | 0.01666667 |
| WMCCExt | Helotiales_family_Incert | W12 | 0.01666667 |
| WMCCExt | Meruliaceae | W12 | 0.01666667 |
| WMCCExt | Pluteaceae | W12 | 0.01666667 |
| WMCCExt | Taphrinaceae | W12 | 0.01666667 |
| WMCCExt | Vibrisseaceae | W12 | 0.01666667 |
| WMCCExt | unclassified_Chytridiomy | W12 | 0.01666667 |
| WMCCExt | unclassified_Exobasidior | W12 | 0.01666667 |
| ASHFExt | Sporidiobolales_family_ | W4 | 30.325 |

| | | | |
|---|---|---|---|
| ASHFExt | Davidiellaceae | W4 | 26.0333333 |
| ASHFExt | Mucoraceae | W4 | 14.7 |
| ASHFExt | Pleosporales_family_Inc | W4 | 9.85833333 |
| ASHFExt | Nectriaceae | W4 | 8.58333333 |
| ASHFExt | Pleosporaceae | W4 | 3.125 |
| ASHFExt | Lasiosphaeriaceae | W4 | 1.64166667 |
| ASHFExt | Pleosporales_unclassified | W4 | 1.38333333 |
| ASHFExt | Filobasidiaceae | W4 | 0.70833333 |
| ASHFExt | Umbelopsidaceae | W4 | 0.61666667 |
| ASHFExt | Sporormiaceae | W4 | 0.38333333 |
| ASHFExt | Amphisphaeriaceae | W4 | 0.30833333 |
| ASHFExt | Leotiomycetes_unclassif | W4 | 0.24166667 |
| ASHFExt | Pseudeurotiaceae | W4 | 0.24166667 |
| ASHFExt | Mortierellaceae | W4 | 0.225 |
| ASHFExt | Tremellales_family_Ince | W4 | 0.2 |
| ASHFExt | Phaeosphaeriaceae | W4 | 0.175 |
| ASHFExt | Fungi_unclassified | W4 | 0.125 |
| ASHFExt | Saccharomycetaceae | W4 | 0.11666667 |
| ASHFExt | Rhizophlyctidaceae | W4 | 0.10833333 |
| ASHFExt | Montagnulaceae | W4 | 0.09166667 |
| ASHFExt | Dothioraceae | W4 | 0.075 |
| ASHFExt | Backusellaceae | W4 | 0.06666667 |
| ASHFExt | Holtermanniales_family_ | W4 | 0.05833333 |
| ASHFExt | Ascomycota_unclassified | W4 | 0.05 |
| ASHFExt | Mycosphaerellaceae | W4 | 0.04166667 |
| ASHFExt | Metschnikowiaceae | W4 | 0.04166667 |
| ASHFExt | Chaetomiaceae | W4 | 0.03333333 |
| ASHFExt | Hypocreales_family_Inc | W4 | 0.03333333 |
| ASHFExt | Spizellomycetaceae | W4 | 0.03333333 |

| | | | |
|---|---|---|---|
| ASHFExt | Eurotiomycetes_unclassi | W4 | 0.025 |
| ASHFExt | Didymosphaeriaceae | W4 | 0.025 |
| ASHFExt | Dothideaceae | W4 | 0.025 |
| ASHFExt | Glomerellaceae | W4 | 0.025 |
| ASHFExt | Helotiales_family_Incert | W4 | 0.025 |
| ASHFExt | Magnaporthaceae | W4 | 0.025 |
| ASHFExt | unclassified_Plantae | W4 | 0.025 |
| ASHFExt | Ceratobasidiaceae | W4 | 0.01666667 |
| ASHFExt | Cucurbitariaceae | W4 | 0.01666667 |
| ASHFExt | Sclerotiniaceae | W4 | 0.01666667 |
| ASHFExt | Trichocomaceae | W4 | 0.01666667 |
| ASHFExt | unclassified_Pleosporale: | W4 | 0.01666667 |
| ASHFExt | Chaetothyriales_unclassi | W4 | 0.01666667 |
| ASHFExt | Dothideomycetes_unclas | W4 | 0.00833333 |
| ASHFExt | Tremellomycetes_unclas | W4 | 0.00833333 |
| ASHFExt | Chaetothyriaceae | W4 | 0.00833333 |
| ASHFExt | Lophiostomataceae | W4 | 0.00833333 |
| ASHFExt | Massarinaceae | W4 | 0.00833333 |
| ASHFExt | Rutstroemiaceae | W4 | 0.00833333 |
| ASHFExt | Taphrinaceae | W4 | 0.00833333 |
| ASHFExt | Capnodiales_unclassifiec | W4 | 0.00833333 |
| ASHFExt | Agaricomycetes_unclass: | W4 | 0.00833333 |
| ASHFExt | Capnodiales_family_Ince | W4 | 0.00833333 |
| ASHFExt | Stereaceae | W4 | 0.00833333 |
| ASHFExt | Basidiomycota_unclassif | W4 | 0.00833333 |
| ASHFInt | Fungi_unclassified | W4 | 16.2916667 |
| ASHFInt | Saccharomycetales_fami | W4 | 11.8083333 |
| ASHFInt | unclassified_Plantae | W4 | 10.625 |
| ASHFInt | Saccharomycetaceae | W4 | 10.4416667 |

| | | | |
|---|---|---|---|
| ASHFInt | Ascomycota_unclassifie | W4 | 5.73333333 |
| ASHFInt | Metschnikowiaceae | W4 | 4.99166667 |
| ASHFInt | Davidiellaceae | W4 | 3.725 |
| ASHFInt | Meruliaceae | W4 | 3.28333333 |
| ASHFInt | Trichocomaceae | W4 | 1.80833333 |
| ASHFInt | Polyporaceae | W4 | 1.44166667 |
| ASHFInt | Ganodermataceae | W4 | 1.4 |
| ASHFInt | Microascaceae | W4 | 1.25 |
| ASHFInt | Sporidiobolales_family_I | W4 | 1.25 |
| ASHFInt | Peniophoraceae | W4 | 1.075 |
| ASHFInt | Pleosporales_family_Inc | W4 | 1 |
| ASHFInt | Taphrinaceae | W4 | 0.96666667 |
| ASHFInt | Schizoporaceae | W4 | 0.95833333 |
| ASHFInt | Xylariaceae | W4 | 0.94166667 |
| ASHFInt | unclassified_Eurotiomyc | W4 | 0.94166667 |
| ASHFInt | Meripilaceae | W4 | 0.89166667 |
| ASHFInt | Polyporales_unclassified | W4 | 0.81666667 |
| ASHFInt | Mycocaliciaceae | W4 | 0.78333333 |
| ASHFInt | Hydnodontaceae | W4 | 0.75 |
| ASHFInt | Myxotrichaceae | W4 | 0.75 |
| ASHFInt | Suillaceae | W4 | 0.725 |
| ASHFInt | Mortierellaceae | W4 | 0.69166667 |
| ASHFInt | Pleosporales_unclassifie | W4 | 0.64166667 |
| ASHFInt | Hypocreaceae | W4 | 0.55 |
| ASHFInt | Basidiomycota_unclassif | W4 | 0.53333333 |
| ASHFInt | Chaetomiaceae | W4 | 0.51666667 |
| ASHFInt | Agaricomycetes_unclass | W4 | 0.50833333 |
| ASHFInt | Mucoraceae | W4 | 0.45 |
| ASHFInt | Amphisphaeriaceae | W4 | 0.39166667 |

| | | | |
|---|---|---|---|
| ASHFInt | Phanerochaetaceae | W4 | 0.38333333 |
| ASHFInt | Stereaceae | W4 | 0.36666667 |
| ASHFInt | Wallemiaceae | W4 | 0.36666667 |
| ASHFInt | Psathyrellaceae | W4 | 0.34166667 |
| ASHFInt | Hymenochaetales_family | W4 | 0.3 |
| ASHFInt | Exobasidiaceae | W4 | 0.29166667 |
| ASHFInt | Mycosphaerellaceae | W4 | 0.28333333 |
| ASHFInt | Venturiaceae | W4 | 0.28333333 |
| ASHFInt | Inocybaceae | W4 | 0.25833333 |
| ASHFInt | Capnodiales_family_Ince | W4 | 0.24166667 |
| ASHFInt | Eurotiomycetes_unclassi | W4 | 0.23333333 |
| ASHFInt | Hymenochaetaceae | W4 | 0.23333333 |
| ASHFInt | Cantharellales_unclassifi | W4 | 0.23333333 |
| ASHFInt | Botryosphaeriaceae | W4 | 0.21666667 |
| ASHFInt | Dothideaceae | W4 | 0.18333333 |
| ASHFInt | Fomitopsidaceae | W4 | 0.18333333 |
| ASHFInt | Nectriaceae | W4 | 0.18333333 |
| ASHFInt | Tremellales_family_Ince | W4 | 0.18333333 |
| ASHFInt | Leptosphaeriaceae | W4 | 0.175 |
| ASHFInt | Sordariomycetes_unclass | W4 | 0.16666667 |
| ASHFInt | Pleosporaceae | W4 | 0.16666667 |
| ASHFInt | Umbelopsidaceae | W4 | 0.16666667 |
| ASHFInt | Capnodiales_unclassified | W4 | 0.16666667 |
| ASHFInt | Strophariaceae | W4 | 0.15833333 |
| ASHFInt | Sclerotiniaceae | W4 | 0.15833333 |
| ASHFInt | unclassified_Corticiales | W4 | 0.15833333 |
| ASHFInt | Lophiostomataceae | W4 | 0.15 |
| ASHFInt | Backusellaceae | W4 | 0.14166667 |
| ASHFInt | Agaricales_unclassified | W4 | 0.14166667 |

| | | | |
|---|---|---|---|
| ASHFInt | Herpotrichiellaceae | W4 | 0.13333333 |
| ASHFInt | Thelephoraceae | W4 | 0.13333333 |
| ASHFInt | Auriculariales_family_In | W4 | 0.13333333 |
| ASHFInt | Botryobasidiaceae | W4 | 0.11666667 |
| ASHFInt | Filobasidiaceae | W4 | 0.11666667 |
| ASHFInt | Dothioraceae | W4 | 0.1 |
| ASHFInt | Hypocreales_family_Inc | W4 | 0.09166667 |
| ASHFInt | Agaricomycetes_family_ | W4 | 0.08333333 |
| ASHFInt | Diaporthales_family_Inc | W4 | 0.08333333 |
| ASHFInt | Hydnaceae | W4 | 0.08333333 |
| ASHFInt | Pluteaceae | W4 | 0.08333333 |
| ASHFInt | Annulatascaceae | W4 | 0.08333333 |
| ASHFInt | Boletaceae | W4 | 0.08333333 |
| ASHFInt | Trichosphaeriales_family | W4 | 0.08333333 |
| ASHFInt | Russulales_unclassified | W4 | 0.08333333 |
| ASHFInt | Ascomycota_family_Inc | W4 | 0.075 |
| ASHFInt | Ceratobasidiaceae | W4 | 0.075 |
| ASHFInt | Gymnoascaceae | W4 | 0.075 |
| ASHFInt | Montagnulaceae | W4 | 0.075 |
| ASHFInt | Onygenaceae | W4 | 0.075 |
| ASHFInt | Polyporales_family_Ince | W4 | 0.075 |
| ASHFInt | Saccharomycetales_uncl | W4 | 0.075 |
| ASHFInt | Entolomataceae | W4 | 0.06666667 |
| ASHFInt | Mycenaceae | W4 | 0.06666667 |
| ASHFInt | unclassified_Trechispora | W4 | 0.06666667 |
| ASHFInt | Cordycipitaceae | W4 | 0.05833333 |
| ASHFInt | Cunninghamellaceae | W4 | 0.05833333 |
| ASHFInt | Dothideomycetes_family | W4 | 0.05833333 |
| ASHFInt | Phaeosphaeriaceae | W4 | 0.05833333 |

| | | | |
|---|---|---|---|
| ASHFInt | Melampsoraceae | W4 | 0.05833333 |
| ASHFInt | Dothideomycetes_unclas | W4 | 0.05 |
| ASHFInt | Russulaceae | W4 | 0.05 |
| ASHFInt | Sporormiaceae | W4 | 0.05 |
| ASHFInt | Vibrisseaceae | W4 | 0.05 |
| ASHFInt | Helotiales_unclassified | W4 | 0.05 |
| ASHFInt | Erysiphaceae | W4 | 0.04166667 |
| ASHFInt | Graphostromataceae | W4 | 0.04166667 |
| ASHFInt | Xenasmataceae | W4 | 0.04166667 |
| ASHFInt | Atheliaceae | W4 | 0.04166667 |
| ASHFInt | Auriscalpiaceae | W4 | 0.04166667 |
| ASHFInt | Bondarzewiaceae | W4 | 0.04166667 |
| ASHFInt | Cystofilobasidiaceae | W4 | 0.04166667 |
| ASHFInt | Eurotiomycetes_family_I | W4 | 0.04166667 |
| ASHFInt | Helotiales_family_Incert | W4 | 0.04166667 |
| ASHFInt | Hypocreales_unclassified | W4 | 0.04166667 |
| ASHFInt | Microstromataceae | W4 | 0.03333333 |
| ASHFInt | Ophiostomataceae | W4 | 0.03333333 |
| ASHFInt | Sordariomycetes_family_ | W4 | 0.03333333 |
| ASHFInt | Lachnocladiaceae | W4 | 0.03333333 |
| ASHFInt | Leotiomycetes_family_In | W4 | 0.03333333 |
| ASHFInt | Marasmiaceae | W4 | 0.03333333 |
| ASHFInt | unclassified_Exobasidion | W4 | 0.03333333 |
| ASHFInt | unclassified_Polyporales | W4 | 0.03333333 |
| ASHFInt | Pucciniales_unclassified | W4 | 0.03333333 |
| ASHFInt | Agaricaceae | W4 | 0.025 |
| ASHFInt | Caliciaceae | W4 | 0.025 |
| ASHFInt | Chaetosphaeriaceae | W4 | 0.025 |
| ASHFInt | Clavicipitaceae | W4 | 0.025 |

| | | | |
|---|---|---|---|
| ASHFInt | Cucurbitariaceae | W4 | 0.025 |
| ASHFInt | Diaporthaceae | W4 | 0.025 |
| ASHFInt | Erythrobasidiales_family | W4 | 0.025 |
| ASHFInt | Gnomoniaceae | W4 | 0.025 |
| ASHFInt | Massarinaceae | W4 | 0.025 |
| ASHFInt | Physciaceae | W4 | 0.025 |
| ASHFInt | Plectosphaerellaceae | W4 | 0.025 |
| ASHFInt | Pleurotaceae | W4 | 0.025 |
| ASHFInt | Rutstroemiaceae | W4 | 0.025 |
| ASHFInt | Sarcosomataceae | W4 | 0.025 |
| ASHFInt | Teratosphaeriaceae | W4 | 0.025 |
| ASHFInt | unclassified_Agaricomy( | W4 | 0.025 |
| ASHFInt | unclassified_Agaricostilt | W4 | 0.025 |
| ASHFInt | unclassified_Pleosporale: | W4 | 0.025 |
| ASHFInt | unclassified_Taphrinales | W4 | 0.025 |
| ASHFInt | Mucorales_unclassified | W4 | 0.025 |
| ASHFInt | Taphrinales_unclassified | W4 | 0.025 |
| ASHFInt | Leotiomycetes_unclassif | W4 | 0.01666667 |
| ASHFInt | Bulgariaceae | W4 | 0.01666667 |
| ASHFInt | Chorioactidaceae | W4 | 0.01666667 |
| ASHFInt | Clavulinaceae | W4 | 0.01666667 |
| ASHFInt | Exobasidiomycetes_fami | W4 | 0.01666667 |
| ASHFInt | Hyaloscyphaceae | W4 | 0.01666667 |
| ASHFInt | Pucciniaceae | W4 | 0.01666667 |
| ASHFInt | Schizophyllaceae | W4 | 0.01666667 |
| ASHFInt | Ustilaginaceae | W4 | 0.01666667 |
| ASHFInt | Eurotiales_unclassified | W4 | 0.01666667 |
| ASHFInt | Pezizales_unclassified | W4 | 0.01666667 |
| ASHFInt | Sordariales_unclassified | W4 | 0.01666667 |

| | | | |
|---|---|---|---|
| ASHFInt | Debaryomycetaceae | W4 | 0.01666667 |
| ASHFInt | Agaricostilbomycetes_ur | W4 | 0.00833333 |
| ASHFInt | Pucciniomycetes_unclass | W4 | 0.00833333 |
| ASHFInt | Ajellomycetaceae | W4 | 0.00833333 |
| ASHFInt | Ascosphaeraceae | W4 | 0.00833333 |
| ASHFInt | Boletinellaceae | W4 | 0.00833333 |
| ASHFInt | Cephalothecaceae | W4 | 0.00833333 |
| ASHFInt | Cladoniaceae | W4 | 0.00833333 |
| ASHFInt | Entomophthoraceae | W4 | 0.00833333 |
| ASHFInt | Halosphaeriaceae | W4 | 0.00833333 |
| ASHFInt | Lentamycetaceae | W4 | 0.00833333 |
| ASHFInt | Melanconidaceae | W4 | 0.00833333 |
| ASHFInt | Ophiocordycipitaceae | W4 | 0.00833333 |
| ASHFInt | Phallaceae | W4 | 0.00833333 |
| ASHFInt | Pucciniastraceae | W4 | 0.00833333 |
| ASHFInt | Septobasidiaceae | W4 | 0.00833333 |
| ASHFInt | Tapinellaceae | W4 | 0.00833333 |
| ASHFInt | Tilletiaceae | W4 | 0.00833333 |
| ASHFInt | Trichosporonaceae | W4 | 0.00833333 |
| ASHFInt | Tubeufiaceae | W4 | 0.00833333 |
| ASHFInt | Valsaceae | W4 | 0.00833333 |
| ASHFInt | Verrucariaceae | W4 | 0.00833333 |
| ASHFInt | Xylariales_family_Incert | W4 | 0.00833333 |
| ASHFInt | unclassified_Hypocreales | W4 | 0.00833333 |
| ASHFInt | Gomphales_unclassified | W4 | 0.00833333 |
| ASHFInt | Lecanorales_unclassified | W4 | 0.00833333 |
| ASHFInt | Cronartiaceae | W4 | 0.00833333 |
| ASHFInt | Diatrypaceae | W4 | 0.00833333 |
| ASHFInt | Glomerellaceae | W4 | 0.00833333 |

| | | | |
|---|---|---|---|
| ASHFInt | Lecanoraceae | W4 | 0.00833333 |
| ASHFInt | Orbiliaceae | W4 | 0.00833333 |
| ASHFInt | Pseudeurotiaceae | W4 | 0.00833333 |
| ASHFInt | Pterulaceae | W4 | 0.00833333 |
| ASHFInt | Pyronemataceae | W4 | 0.00833333 |
| ASHFInt | Ramalinaceae | W4 | 0.00833333 |
| ASHFInt | unclassified_Auricularial | W4 | 0.00833333 |
| ASHFInt | unclassified_Saccharomy | W4 | 0.00833333 |
| ASHFInt | Boletales_unclassified | W4 | 0.00833333 |
| ASHFInt | Chaetothyriales_unclassi | W4 | 0.00833333 |
| ASHFInt | Trechisporales_unclassif | W4 | 0.00833333 |
| ASHFInt | Xylariales_unclassified | W4 | 0.00833333 |
| WMCCExt | Davidiellaceae | W4 | 32.3083333 |
| WMCCExt | Sporidiobolales_family_I | W4 | 15.775 |
| WMCCExt | Pleosporales_family_Inc | W4 | 10.825 |
| WMCCExt | Mucoraceae | W4 | 10.25 |
| WMCCExt | Dothioraceae | W4 | 7.53333333 |
| WMCCExt | Pleosporaceae | W4 | 4.86666667 |
| WMCCExt | Pyronemataceae | W4 | 2.55 |
| WMCCExt | Sporormiaceae | W4 | 2.375 |
| WMCCExt | Phaeosphaeriaceae | W4 | 1.725 |
| WMCCExt | Pleosporales_unclassifie | W4 | 1.49166667 |
| WMCCExt | Tremellales_family_Ince | W4 | 1.45833333 |
| WMCCExt | Trichocomaceae | W4 | 1.275 |
| WMCCExt | Fungi_unclassified | W4 | 1.15 |
| WMCCExt | Mucorales_unclassified | W4 | 1.06666667 |
| WMCCExt | Nectriaceae | W4 | 0.84166667 |
| WMCCExt | Tremellomycetes_unclas | W4 | 0.83333333 |
| WMCCExt | Lophiostomataceae | W4 | 0.625 |

| | | | |
|---|---|---|---|
| WMCCExt | unclassified_Exobasidior | W4 | 0.60833333 |
| WMCCExt | Montagnulaceae | W4 | 0.58333333 |
| WMCCExt | Leptosphaeriaceae | W4 | 0.31666667 |
| WMCCExt | Ascomycota_unclassifiec | W4 | 0.3 |
| WMCCExt | Umbelopsidaceae | W4 | 0.19166667 |
| WMCCExt | Hypocreales_family_Inc | W4 | 0.175 |
| WMCCExt | Basidiomycota_unclassif | W4 | 0.125 |
| WMCCExt | Chaetomiaceae | W4 | 0.09166667 |
| WMCCExt | Coniochaetaceae | W4 | 0.09166667 |
| WMCCExt | Dothideaceae | W4 | 0.075 |
| WMCCExt | Cucurbitariaceae | W4 | 0.05833333 |
| WMCCExt | Amphisphaeriaceae | W4 | 0.04166667 |
| WMCCExt | Cystofilobasidiaceae | W4 | 0.04166667 |
| WMCCExt | Ascomycota_family_Inc | W4 | 0.025 |
| WMCCExt | Cordycipitaceae | W4 | 0.025 |
| WMCCExt | Holtermanniales_family_ | W4 | 0.025 |
| WMCCExt | Mycosphaerellaceae | W4 | 0.025 |
| WMCCExt | Diaporthaceae | W4 | 0.01666667 |
| WMCCExt | Microascaceae | W4 | 0.01666667 |
| WMCCExt | unclassified_Chytridiomy | W4 | 0.01666667 |
| WMCCExt | Agaricomycetes_unclass | W4 | 0.01666667 |
| WMCCExt | Helotiaceae | W4 | 0.01666667 |
| WMCCExt | Meruliaceae | W4 | 0.01666667 |
| WMCCExt | Trichosphaeriales_family | W4 | 0.01666667 |
| WMCCExt | Ustilaginales_family_Inc | W4 | 0.01666667 |
| WMCCExt | Bionectriaceae | W4 | 0.00833333 |
| WMCCExt | Chaetothyriaceae | W4 | 0.00833333 |
| WMCCExt | Exobasidiomycetes_fami | W4 | 0.00833333 |
| WMCCExt | Gnomoniaceae | W4 | 0.00833333 |

| | | | |
|---|---|---|---|
| WMCCExt | Magnaporthaceae | W4 | 0.00833333 |
| WMCCExt | Metschnikowiaceae | W4 | 0.00833333 |
| WMCCExt | Ophiocordycipitaceae | W4 | 0.00833333 |
| WMCCExt | Peniophoraceae | W4 | 0.00833333 |
| WMCCExt | Plectosphaerellaceae | W4 | 0.00833333 |
| WMCCExt | Ustilaginaceae | W4 | 0.00833333 |
| WMCCExt | Helotiales_unclassified | W4 | 0.00833333 |
| WMCCExt | Glomerellaceae | W4 | 0.00833333 |
| WMCCExt | Hydnaceae | W4 | 0.00833333 |
| WMCCExt | Vibrisseaceae | W4 | 0.00833333 |
| ASHFExt | Sporidiobolales_family_I | W8 | 49.5214286 |
| ASHFExt | Pleosporales_family_Inc | W8 | 12.7428571 |
| ASHFExt | Nectriaceae | W8 | 11.6142857 |
| ASHFExt | Mortierellaceae | W8 | 6.95714286 |
| ASHFExt | Davidiellaceae | W8 | 5.95714286 |
| ASHFExt | Mucoraceae | W8 | 2.98571429 |
| ASHFExt | Pleosporaceae | W8 | 2.75 |
| ASHFExt | Lasiosphaeriaceae | W8 | 1.82142857 |
| ASHFExt | Rhizophlyctidaceae | W8 | 0.7 |
| ASHFExt | Filobasidiaceae | W8 | 0.6 |
| ASHFExt | Fungi_unclassified | W8 | 0.56428571 |
| ASHFExt | Pseudeurotiaceae | W8 | 0.52142857 |
| ASHFExt | Umbelopsidaceae | W8 | 0.49285714 |
| ASHFExt | Amphisphaeriaceae | W8 | 0.45 |
| ASHFExt | Pleosporales_unclassified | W8 | 0.41428571 |
| ASHFExt | Phaeosphaeriaceae | W8 | 0.25714286 |
| ASHFExt | Montagnulaceae | W8 | 0.20714286 |
| ASHFExt | Spizellomycetaceae | W8 | 0.19285714 |
| ASHFExt | Holtermanniales_family_ | W8 | 0.17142857 |

| | | | |
|---|---|---|---|
| ASHFExt | Tremellales_family_Ince | W8 | 0.12857143 |
| ASHFExt | Trichocomaceae | W8 | 0.1 |
| ASHFExt | Corticiaceae | W8 | 0.09285714 |
| ASHFExt | Backusellaceae | W8 | 0.06428571 |
| ASHFExt | Hypocreales_family_Inc | W8 | 0.06428571 |
| ASHFExt | Chaetomiaceae | W8 | 0.05714286 |
| ASHFExt | Ceratobasidiaceae | W8 | 0.05 |
| ASHFExt | unclassified_Pleosporale | W8 | 0.05 |
| ASHFExt | Herpotrichiellaceae | W8 | 0.03571429 |
| ASHFExt | Leptosphaeriaceae | W8 | 0.03571429 |
| ASHFExt | unclassified_Pezizomyce | W8 | 0.03571429 |
| ASHFExt | Magnaporthaceae | W8 | 0.02857143 |
| ASHFExt | Orbiliaceae | W8 | 0.02857143 |
| ASHFExt | Wallemiaceae | W8 | 0.02857143 |
| ASHFExt | unclassified_Fungi | W8 | 0.02857143 |
| ASHFExt | Helotiaceae | W8 | 0.02142857 |
| ASHFExt | Ascomycota_unclassified | W8 | 0.02142857 |
| ASHFExt | Leotiomycetes_unclassif | W8 | 0.01428571 |
| ASHFExt | Meruliaceae | W8 | 0.01428571 |
| ASHFExt | Mycosphaerellaceae | W8 | 0.01428571 |
| ASHFExt | Chaetothyriales_unclassi | W8 | 0.01428571 |
| ASHFExt | Tremellomycetes_unclas | W8 | 0.01428571 |
| ASHFExt | Cystofilobasidiaceae | W8 | 0.01428571 |
| ASHFExt | Onygenaceae | W8 | 0.01428571 |
| ASHFExt | Hypocreales_unclassified | W8 | 0.01428571 |
| ASHFExt | Eurotiomycetes_unclassi | W8 | 0.00714286 |
| ASHFExt | Sordariomycetes_unclass | W8 | 0.00714286 |
| ASHFExt | Dothideaceae | W8 | 0.00714286 |
| ASHFExt | Ganodermataceae | W8 | 0.00714286 |

| | | | |
|---|---|---|---|
| ASHFExt | Meripilaceae | W8 | 0.00714286 |
| ASHFExt | Microascaceae | W8 | 0.00714286 |
| ASHFExt | Peniophoraceae | W8 | 0.00714286 |
| ASHFExt | Psathyrellaceae | W8 | 0.00714286 |
| ASHFExt | Ustilaginaceae | W8 | 0.00714286 |
| ASHFExt | unclassified_Plantae | W8 | 0.00714286 |
| ASHFExt | Onygenales_unclassified | W8 | 0.00714286 |
| ASHFExt | Sordariales_unclassified | W8 | 0.00714286 |
| ASHFExt | Basidiomycota_unclassif | W8 | 0.00714286 |
| ASHFInt | unclassified_Plantae | W8 | 32.6333333 |
| ASHFInt | Wallemiaceae | W8 | 13.0166667 |
| ASHFInt | Trichocomaceae | W8 | 6.48333333 |
| ASHFInt | Sporidiobolales_family_I | W8 | 6.45 |
| ASHFInt | Davidiellaceae | W8 | 4.43333333 |
| ASHFInt | Ascomycota_unclassified | W8 | 3.75 |
| ASHFInt | Taphrinaceae | W8 | 3.63333333 |
| ASHFInt | Meruliaceae | W8 | 3.23333333 |
| ASHFInt | Peniophoraceae | W8 | 1.71666667 |
| ASHFInt | Pleosporales_family_Inc | W8 | 1.55 |
| ASHFInt | Schizoporaceae | W8 | 1.48333333 |
| ASHFInt | Basidiomycota_unclassif | W8 | 1.01666667 |
| ASHFInt | Fungi_unclassified | W8 | 0.93333333 |
| ASHFInt | Dothideaceae | W8 | 0.93333333 |
| ASHFInt | Polyporaceae | W8 | 0.88333333 |
| ASHFInt | Agaricomycetes_unclass | W8 | 0.8 |
| ASHFInt | Saccharomycetaceae | W8 | 0.76666667 |
| ASHFInt | Hymenochaetales_family | W8 | 0.68333333 |
| ASHFInt | Mycosphaerellaceae | W8 | 0.6 |
| ASHFInt | Stereaceae | W8 | 0.58333333 |

| | | | |
|---|---|---|---|
| ASHFInt | Filobasidiaceae | W8 | 0.48333333 |
| ASHFInt | Pleosporales_unclassified | W8 | 0.48333333 |
| ASHFInt | Hydnodontaceae | W8 | 0.46666667 |
| ASHFInt | Melampsoraceae | W8 | 0.41666667 |
| ASHFInt | Pleurotaceae | W8 | 0.41666667 |
| ASHFInt | Ganodermataceae | W8 | 0.4 |
| ASHFInt | Lophiostomataceae | W8 | 0.4 |
| ASHFInt | Psathyrellaceae | W8 | 0.4 |
| ASHFInt | Sordariomycetes_unclass | W8 | 0.36666667 |
| ASHFInt | Exobasidiaceae | W8 | 0.36666667 |
| ASHFInt | Phanerochaetaceae | W8 | 0.33333333 |
| ASHFInt | Dothioraceae | W8 | 0.31666667 |
| ASHFInt | Hymenochaetaceae | W8 | 0.28333333 |
| ASHFInt | Sclerotiniaceae | W8 | 0.28333333 |
| ASHFInt | Tremellales_family_Ince | W8 | 0.28333333 |
| ASHFInt | Eurotiomycetes_unclassi | W8 | 0.26666667 |
| ASHFInt | Annulatascaceae | W8 | 0.25 |
| ASHFInt | Strophariaceae | W8 | 0.25 |
| ASHFInt | Xylariaceae | W8 | 0.25 |
| ASHFInt | Pleosporaceae | W8 | 0.23333333 |
| ASHFInt | Sporormiaceae | W8 | 0.23333333 |
| ASHFInt | Atheliaceae | W8 | 0.21666667 |
| ASHFInt | Meripilaceae | W8 | 0.21666667 |
| ASHFInt | Venturiaceae | W8 | 0.2 |
| ASHFInt | Microascaceae | W8 | 0.2 |
| ASHFInt | Polyporales_unclassified | W8 | 0.2 |
| ASHFInt | Agaricomycetes_family_ | W8 | 0.18333333 |
| ASHFInt | Amphisphaeriaceae | W8 | 0.18333333 |
| ASHFInt | Cantharellales_unclassifi | W8 | 0.18333333 |

| | | | |
|---|---|---|---|
| ASHFInt | Capnodiales_unclassified | W8 | 0.18333333 |
| ASHFInt | Hydnaceae | W8 | 0.16666667 |
| ASHFInt | Mortierellaceae | W8 | 0.16666667 |
| ASHFInt | Umbelopsidaceae | W8 | 0.16666667 |
| ASHFInt | Physciaceae | W8 | 0.16666667 |
| ASHFInt | Schizophyllaceae | W8 | 0.15 |
| ASHFInt | unclassified_Agaricostilt | W8 | 0.15 |
| ASHFInt | Gnomoniaceae | W8 | 0.13333333 |
| ASHFInt | Montagnulaceae | W8 | 0.13333333 |
| ASHFInt | Capnodiales_family_Ince | W8 | 0.11666667 |
| ASHFInt | Hyaloscyphaceae | W8 | 0.11666667 |
| ASHFInt | Nectriaceae | W8 | 0.11666667 |
| ASHFInt | Vibrisseaceae | W8 | 0.11666667 |
| ASHFInt | Agaricales_unclassified | W8 | 0.11666667 |
| ASHFInt | Taphrinales_unclassified | W8 | 0.11666667 |
| ASHFInt | Botryobasidiaceae | W8 | 0.1 |
| ASHFInt | Fomitopsidaceae | W8 | 0.1 |
| ASHFInt | Helotiales_family_Incert | W8 | 0.1 |
| ASHFInt | Herpotrichiellaceae | W8 | 0.1 |
| ASHFInt | Marasmiaceae | W8 | 0.1 |
| ASHFInt | Mycenaceae | W8 | 0.1 |
| ASHFInt | Trichosphaeriales_family | W8 | 0.1 |
| ASHFInt | Inocybaceae | W8 | 0.08333333 |
| ASHFInt | Erythrobasidiales_family | W8 | 0.08333333 |
| ASHFInt | Ophiostomataceae | W8 | 0.08333333 |
| ASHFInt | Basidiobolaceae | W8 | 0.06666667 |
| ASHFInt | Candelariaceae | W8 | 0.06666667 |
| ASHFInt | Phycomycetaceae | W8 | 0.06666667 |
| ASHFInt | Suillaceae | W8 | 0.06666667 |

| | | | |
|---|---|---|---|
| ASHFInt | Auriculariales_family_In | W8 | 0.06666667 |
| ASHFInt | Ceratobasidiaceae | W8 | 0.06666667 |
| ASHFInt | Diaporthales_family_Inc | W8 | 0.06666667 |
| ASHFInt | Microstromataceae | W8 | 0.06666667 |
| ASHFInt | Mucoraceae | W8 | 0.06666667 |
| ASHFInt | Pluteaceae | W8 | 0.06666667 |
| ASHFInt | unclassified_Corticiales | W8 | 0.06666667 |
| ASHFInt | Russulales_unclassified | W8 | 0.06666667 |
| ASHFInt | Auriscalpiaceae | W8 | 0.05 |
| ASHFInt | Cordycipitaceae | W8 | 0.05 |
| ASHFInt | Cucurbitariaceae | W8 | 0.05 |
| ASHFInt | Diatrypaceae | W8 | 0.05 |
| ASHFInt | Leptosphaeriaceae | W8 | 0.05 |
| ASHFInt | Parmeliaceae | W8 | 0.05 |
| ASHFInt | Plectosphaerellaceae | W8 | 0.05 |
| ASHFInt | Saccharomycetales_fami | W8 | 0.05 |
| ASHFInt | Teratosphaeriaceae | W8 | 0.05 |
| ASHFInt | Tricholomataceae | W8 | 0.05 |
| ASHFInt | Agaricostilbaceae | W8 | 0.03333333 |
| ASHFInt | Botryosphaeriaceae | W8 | 0.03333333 |
| ASHFInt | Clavicipitaceae | W8 | 0.03333333 |
| ASHFInt | Clavulinaceae | W8 | 0.03333333 |
| ASHFInt | Diaporthaceae | W8 | 0.03333333 |
| ASHFInt | Halosphaeriaceae | W8 | 0.03333333 |
| ASHFInt | Ustilaginaceae | W8 | 0.03333333 |
| ASHFInt | unclassified_Taphrinales | W8 | 0.03333333 |
| ASHFInt | Diaporthales_unclassifie | W8 | 0.03333333 |
| ASHFInt | Helotiales_unclassified | W8 | 0.03333333 |
| ASHFInt | Dermateaceae | W8 | 0.03333333 |

| | | | |
|---|---|---|---|
| ASHFInt | Hypocreaceae | W8 | 0.03333333 |
| ASHFInt | Massarinaceae | W8 | 0.03333333 |
| ASHFInt | Mycocaliciaceae | W8 | 0.03333333 |
| ASHFInt | Polyporales_family_Ince | W8 | 0.03333333 |
| ASHFInt | Pseudeurotiaceae | W8 | 0.03333333 |
| ASHFInt | Thelephoraceae | W8 | 0.03333333 |
| ASHFInt | unclassified_Pleosporale | W8 | 0.03333333 |
| ASHFInt | Atheliales_unclassified | W8 | 0.03333333 |
| ASHFInt | Dothideomycetes_unclas | W8 | 0.01666667 |
| ASHFInt | Leotiomycetes_unclassif | W8 | 0.01666667 |
| ASHFInt | Pucciniomycetes_unclass | W8 | 0.01666667 |
| ASHFInt | Tremellomycetes_unclas | W8 | 0.01666667 |
| ASHFInt | Arachnomycetaceae | W8 | 0.01666667 |
| ASHFInt | Ascomycota_family_Inc | W8 | 0.01666667 |
| ASHFInt | Ascosphaeraceae | W8 | 0.01666667 |
| ASHFInt | Bolbitiaceae | W8 | 0.01666667 |
| ASHFInt | Boliniaceae | W8 | 0.01666667 |
| ASHFInt | Bondarzewiaceae | W8 | 0.01666667 |
| ASHFInt | Calosphaeriaceae | W8 | 0.01666667 |
| ASHFInt | Cephalothecaceae | W8 | 0.01666667 |
| ASHFInt | Ceratostomataceae | W8 | 0.01666667 |
| ASHFInt | Cystofilobasidiaceae | W8 | 0.01666667 |
| ASHFInt | Didymosphaeriaceae | W8 | 0.01666667 |
| ASHFInt | Elsinoaceae | W8 | 0.01666667 |
| ASHFInt | Gomphaceae | W8 | 0.01666667 |
| ASHFInt | Hypocreales_family_Inc | W8 | 0.01666667 |
| ASHFInt | Magnaporthaceae | W8 | 0.01666667 |
| ASHFInt | Melanconidaceae | W8 | 0.01666667 |
| ASHFInt | Micropeltidaceae | W8 | 0.01666667 |

| | | | |
|---|---|---|---|
| ASHFInt | Pezizaceae | W8 | 0.01666667 |
| ASHFInt | Phaeosphaeriaceae | W8 | 0.01666667 |
| ASHFInt | Quambalariaceae | W8 | 0.01666667 |
| ASHFInt | Schizoparmaceae | W8 | 0.01666667 |
| ASHFInt | Sebacinaceae | W8 | 0.01666667 |
| ASHFInt | Sordariomycetes_family_ | W8 | 0.01666667 |
| ASHFInt | Spizellomycetaceae | W8 | 0.01666667 |
| ASHFInt | Tubeufiaceae | W8 | 0.01666667 |
| ASHFInt | Xylariales_family_Incert | W8 | 0.01666667 |
| ASHFInt | unclassified_Atheliales | W8 | 0.01666667 |
| ASHFInt | unclassified_Atractiellale | W8 | 0.01666667 |
| ASHFInt | unclassified_Polyporales | W8 | 0.01666667 |
| ASHFInt | unclassified_Saccharomy | W8 | 0.01666667 |
| ASHFInt | Boletales_unclassified | W8 | 0.01666667 |
| ASHFInt | Eurotiales_unclassified | W8 | 0.01666667 |
| ASHFInt | Hypocreales_unclassifiec | W8 | 0.01666667 |
| ASHFInt | Lecanorales_unclassified | W8 | 0.01666667 |
| ASHFInt | Sebacinales_unclassified | W8 | 0.01666667 |
| WMCCExt | Davidiellaceae | W8 | 36.7928571 |
| WMCCExt | Sporidiobolales_family_ | W8 | 11.7642857 |
| WMCCExt | Pleosporales_family_Inc | W8 | 7.74285714 |
| WMCCExt | Mucoraceae | W8 | 5.59285714 |
| WMCCExt | Hypocreales_family_Inc | W8 | 4.90714286 |
| WMCCExt | Chaetomiaceae | W8 | 4.58571429 |
| WMCCExt | Pleosporaceae | W8 | 4.32142857 |
| WMCCExt | Tremellales_family_Ince | W8 | 4.19285714 |
| WMCCExt | Nectriaceae | W8 | 4.11428571 |
| WMCCExt | Lophiostomataceae | W8 | 3.43571429 |
| WMCCExt | Pleosporales_unclassifiec | W8 | 2.85 |

| | | | |
|---|---|---|---|
| WMCCExt | Dothioraceae | W8 | 1.97857143 |
| WMCCExt | Phaeosphaeriaceae | W8 | 1.63571429 |
| WMCCExt | Sporormiaceae | W8 | 1.44285714 |
| WMCCExt | Fungi_unclassified | W8 | 0.98571429 |
| WMCCExt | Tremellomycetes_unclas | W8 | 0.62142857 |
| WMCCExt | Montagnulaceae | W8 | 0.31428571 |
| WMCCExt | Ascomycota_unclassifie | W8 | 0.27142857 |
| WMCCExt | Leptosphaeriaceae | W8 | 0.22142857 |
| WMCCExt | Cucurbitariaceae | W8 | 0.2 |
| WMCCExt | Trichocomaceae | W8 | 0.17857143 |
| WMCCExt | Umbelopsidaceae | W8 | 0.17142857 |
| WMCCExt | Cordycipitaceae | W8 | 0.16428571 |
| WMCCExt | Helotiaceae | W8 | 0.14285714 |
| WMCCExt | Ophiocordycipitaceae | W8 | 0.12857143 |
| WMCCExt | Agaricomycetes_unclass | W8 | 0.1 |
| WMCCExt | Ascomycota_family_Inc | W8 | 0.09285714 |
| WMCCExt | Clavicipitaceae | W8 | 0.08571429 |
| WMCCExt | Spizellomycetaceae | W8 | 0.07857143 |
| WMCCExt | unclassified_Exobasidio | W8 | 0.07857143 |
| WMCCExt | Basidiomycota_unclassif | W8 | 0.07142857 |
| WMCCExt | Mucorales_unclassified | W8 | 0.07142857 |
| WMCCExt | Filobasidiaceae | W8 | 0.05 |
| WMCCExt | Chaetothyriales_family_ | W8 | 0.04285714 |
| WMCCExt | Xylariales_family_Incert | W8 | 0.04285714 |
| WMCCExt | Helotiales_unclassified | W8 | 0.04285714 |
| WMCCExt | Sordariaceae | W8 | 0.03571429 |
| WMCCExt | unclassified_Chytridiom | W8 | 0.03571429 |
| WMCCExt | Ascobolaceae | W8 | 0.03571429 |
| WMCCExt | Dothideaceae | W8 | 0.02857143 |

| | | | |
|---|---|---|---|
| WMCCExt | Helotiales_family_Incert | W8 | 0.02857143 |
| WMCCExt | Mycosphaerellaceae | W8 | 0.02857143 |
| WMCCExt | Leotiomycetes_unclassif | W8 | 0.02142857 |
| WMCCExt | Holtermanniales_family_ | W8 | 0.02142857 |
| WMCCExt | Ustilaginaceae | W8 | 0.02142857 |
| WMCCExt | Capnodiales_family_Ince | W8 | 0.01428571 |
| WMCCExt | Didymosphaeriaceae | W8 | 0.01428571 |
| WMCCExt | Leucosporidiaceae | W8 | 0.01428571 |
| WMCCExt | unclassified_Pleosporale | W8 | 0.01428571 |
| WMCCExt | Plectosphaerellaceae | W8 | 0.01428571 |
| WMCCExt | Pseudeurotiaceae | W8 | 0.01428571 |
| WMCCExt | Ustilaginales_family_Inc | W8 | 0.01428571 |
| WMCCExt | Chaetothyriales_unclassi | W8 | 0.01428571 |
| WMCCExt | Agaricostilbaceae | W8 | 0.00714286 |
| WMCCExt | Coniochaetaceae | W8 | 0.00714286 |
| WMCCExt | Cystofilobasidiaceae | W8 | 0.00714286 |
| WMCCExt | Dothideomycetes_family | W8 | 0.00714286 |
| WMCCExt | Erythrobasidiales_family | W8 | 0.00714286 |
| WMCCExt | Herpotrichiellaceae | W8 | 0.00714286 |
| WMCCExt | Hymenochaetaceae | W8 | 0.00714286 |
| WMCCExt | Lasiosphaeriaceae | W8 | 0.00714286 |
| WMCCExt | Leotiaceae | W8 | 0.00714286 |
| WMCCExt | Magnaporthaceae | W8 | 0.00714286 |
| WMCCExt | Meruliaceae | W8 | 0.00714286 |
| WMCCExt | Mortierellaceae | W8 | 0.00714286 |
| WMCCExt | Orbiliaceae | W8 | 0.00714286 |
| WMCCExt | Peniophoraceae | W8 | 0.00714286 |
| WMCCExt | Saccharomycetaceae | W8 | 0.00714286 |
| WMCCExt | Sordariales_unclassified | W8 | 0.00714286 |
