## Supplementary material for "Microbial, dietary insect, and pathogen communities in fresh and decomposing guano of anthropic little brown bat (Myotis lucifugus) maternity colonies": Table S15

| Age | Site | Family | Specificity | Fidelity | Stat | P |
|---|---|---|---|---|---|---|
| Aged | ASHFInt | Atheliaceae | 1 | 1 | 1 | 0.001 |
| Aged | ASHFInt | Cordycipitaceae | 1 | 1 | 1 | 0.001 |
| Aged | ASHFInt | DiaporthalesfamilyIncertaesedis | 1 | 1 | 1 | 0.001 |
| Aged | ASHFInt | Exobasidiaceae | 1 | 1 | 1 | 0.001 |
| Aged | ASHFInt | Fomitopsidaceae | 1 | 1 | 1 | 0.001 |
| Aged | ASHFInt | Hyaloscyphaceae | 1 | 1 | 1 | 0.001 |
| Aged | ASHFInt | Inocybaceae | 1 | 1 | 1 | 0.001 |
| Aged | ASHFInt | Marasmiaceae | 1 | 1 | 1 | 0.001 |
| Aged | ASHFInt | Massarinaceae | 1 | 1 | 1 | 0.001 |
| Aged | ASHFInt | Melampsoraceae | 1 | 1 | 1 | 0.001 |
| Aged | ASHFInt | Microascaceae | 1 | 1 | 1 | 0.001 |
| Aged | ASHFInt | Microstromataceae | 1 | 1 | 1 | 0.001 |
| Aged | ASHFInt | Pluteaceae | 1 | 1 | 1 | 0.001 |
| Aged | ASHFInt | Schizophyllaceae | 1 | 1 | 1 | 0.001 |
| Aged | ASHFInt | Schizoporaceae | 1 | 1 | 1 | 0.001 |
| Aged | ASHFInt | Strophariaceae | 1 | 1 | 1 | 0.001 |
| Aged | ASHFInt | Wallemiaceae | 1 | 1 | 1 | 0.001 |
| Aged | ASHFInt | Agaricalesunclassified | 1 | 1 | 1 | 0.001 |
| Aged | ASHFInt | Capnodialesunclassified | 1 | 1 | 1 | 0.001 |
| Aged | ASHFInt | unclassifiedPlantae | 0.9998 | 1 | 1 | 0.001 |
| Aged | ASHFInt | Taphrinaceae | 0.9987 | 1 | 0.999 | 0.002 |
| Aged | ASHFInt | Trichocomaceae | 0.9976 | 1 | 0.999 | 0.001 |
| Aged | ASHFInt | Stereaceae | 0.9968 | 1 | 0.998 | 0.001 |
| Aged | ASHFInt | HymenochaetalesfamilyIncertaesedis | 0.9964 | 1 | 0.998 | 0.001 |
| Aged | ASHFInt | Hydnodontaceae | 0.9959 | 1 | 0.998 | 0.001 |
| Aged | ASHFInt | Meruliaceae | 0.9958 | 1 | 0.998 | 0.001 |
| Aged | ASHFInt | Polyporaceae | 0.9944 | 1 | 0.997 | 0.002 |
| Aged | ASHFInt | Psathyrellaceae | 0.9942 | 1 | 0.997 | 0.001 |
| Aged | ASHFInt | CapnodialesfamilyIncertaesedis | 0.9935 | 1 | 0.997 | 0.001 |
| Aged | ASHFInt | Agaricomycetesunclassified | 0.9934 | 1 | 0.997 | 0.001 |
| Aged | ASHFInt | Xylariaceae | 0.9933 | 1 | 0.997 | 0.001 |

| | | | | | | |
|---|---|---|---|---|---|---|
| Aged | ASHFInt | Sordariomycetesunclassified | 0.9917 | 1 | 0.996 | 0.002 |
| Aged | ASHFInt | Sclerotiniaceae | 0.9916 | 1 | 0.996 | 0.002 |
| Aged | ASHFInt | Cantharellalesunclassified | 0.9914 | 1 | 0.996 | 0.001 |
| Aged | ASHFInt | Hypocreaceae | 0.9909 | 1 | 0.995 | 0.001 |
| Aged | ASHFInt | Ganodermataceae | 0.9906 | 1 | 0.995 | 0.001 |
| Aged | ASHFInt | Phanerochaetaceae | 0.9895 | 1 | 0.995 | 0.001 |
| Aged | ASHFInt | Hydnaceae | 0.9865 | 1 | 0.993 | 0.001 |
| Aged | ASHFInt | Peniophoraceae | 0.9848 | 1 | 0.992 | 0.001 |
| Aged | ASHFInt | Mycosphaerellaceae | 0.9813 | 1 | 0.991 | 0.002 |
| Aged | ASHFInt | Saccharomycetaceae | 0.9789 | 1 | 0.989 | 0.003 |
| Aged | ASHFInt | Hymenochaetaceae | 0.9738 | 1 | 0.987 | 0.001 |
| Aged | ASHFInt | Meripilaceae | 0.9735 | 1 | 0.987 | 0.001 |
| Aged | ASHFInt | Venturiaceae | 0.9706 | 1 | 0.985 | 0.001 |
| Aged | ASHFInt | Eurotiomycetesunclassified | 0.9664 | 1 | 0.983 | 0.001 |
| Aged | ASHFInt | Gnomoniaceae | 0.9627 | 1 | 0.981 | 0.002 |
| Aged | ASHFInt | Ceratobasidiaceae | 0.9194 | 1 | 0.959 | 0.001 |
| Aged | ASHFInt | Vibrisseaceae | 0.9083 | 1 | 0.953 | 0.001 |
| Aged | ASHFInt | Herpotrichiellaceae | 0.8239 | 1 | 0.908 | 0.01 |
| Aged | ASHFInt | Dothideomycetesunclassified | 1 | 0.8 | 0.894 | 0.005 |
| Aged | ASHFInt | AgaricomycetesfamilyIncertaesedis | 1 | 0.8 | 0.894 | 0.002 |
| Aged | ASHFInt | AuricularialesfamilyIncertaesedis | 1 | 0.8 | 0.894 | 0.002 |
| Aged | ASHFInt | Botryobasidiaceae | 1 | 0.8 | 0.894 | 0.002 |
| Aged | ASHFInt | Clavicipitaceae | 1 | 0.8 | 0.894 | 0.004 |
| Aged | ASHFInt | Clavulinaceae | 1 | 0.8 | 0.894 | 0.002 |
| Aged | ASHFInt | HelotialesfamilyIncertaesedis | 1 | 0.8 | 0.894 | 0.002 |
| Aged | ASHFInt | Mycenaceae | 1 | 0.8 | 0.894 | 0.004 |
| Aged | ASHFInt | Mycocaliciaceae | 1 | 0.8 | 0.894 | 0.004 |
| Aged | ASHFInt | Pleurotaceae | 1 | 0.8 | 0.894 | 0.004 |
| Aged | ASHFInt | PolyporalesfamilyIncertaesedis | 1 | 0.8 | 0.894 | 0.002 |
| Aged | ASHFInt | unclassifiedCorticiales | 1 | 0.8 | 0.894 | 0.002 |
| Aged | ASHFInt | Helotialesunclassified | 1 | 0.8 | 0.894 | 0.002 |

| | | | | | | |
|---|---|---|---|---|---|---|
| Aged | ASHFInt | Hypocrealesunclassified | 1 | 0.8 | 0.894 | 0.002 |
| Aged | ASHFInt | Lecanoralesunclassified | 1 | 0.8 | 0.894 | 0.002 |
| Aged | ASHFInt | Polyporalesunclassified | 1 | 0.8 | 0.894 | 0.002 |
| Aged | ASHFInt | Russulalesunclassified | 1 | 0.8 | 0.894 | 0.002 |
| Aged | ASHFInt | Ustilaginaceae | 0.7934 | 1 | 0.891 | 0.003 |
| Aged | ASHFInt | Diatrypaceae | 0.973 | 0.8 | 0.882 | 0.003 |
| Aged | ASHFInt | DothideomycetesfamilyIncertaesedis | 0.9231 | 0.8 | 0.859 | 0.013 |
| Aged | ASHFInt | Leptosphaeriaceae | 0.8485 | 0.8 | 0.824 | 0.015 |
| Aged | ASHFInt | Agaricostilbomycetesunclassified | 1 | 0.6 | 0.775 | 0.013 |
| Aged | ASHFInt | Auriscalpiaceae | 1 | 0.6 | 0.775 | 0.012 |
| Aged | ASHFInt | Cephalothecaceae | 1 | 0.6 | 0.775 | 0.019 |
| Aged | ASHFInt | Chaetosphaeriaceae | 1 | 0.6 | 0.775 | 0.017 |
| Aged | ASHFInt | Diaporthaceae | 1 | 0.6 | 0.775 | 0.012 |
| Aged | ASHFInt | Entolomataceae | 1 | 0.6 | 0.775 | 0.016 |
| Aged | ASHFInt | Erysiphaceae | 1 | 0.6 | 0.775 | 0.013 |
| Aged | ASHFInt | EurotiomycetesfamilyIncertaesedis | 1 | 0.6 | 0.775 | 0.016 |
| Aged | ASHFInt | Halosphaeriaceae | 1 | 0.6 | 0.775 | 0.012 |
| Aged | ASHFInt | Lachnocladiaceae | 1 | 0.6 | 0.775 | 0.013 |
| Aged | ASHFInt | Lecanoraceae | 1 | 0.6 | 0.775 | 0.017 |
| Aged | ASHFInt | SaccharomycetalesfamilyIncertaesedis | 1 | 0.6 | 0.775 | 0.021 |
| Aged | ASHFInt | SordariomycetesfamilyIncertaesedis | 1 | 0.6 | 0.775 | 0.012 |
| Aged | ASHFInt | Stephanosporaceae | 1 | 0.6 | 0.775 | 0.019 |
| Aged | ASHFInt | unclassifiedAgaricostilbomycetes | 1 | 0.6 | 0.775 | 0.017 |
| Aged | ASHFInt | unclassifiedTaphrinales | 1 | 0.6 | 0.775 | 0.016 |
| Aged | ASHFInt | unclassifiedTrechisporales | 1 | 0.6 | 0.775 | 0.019 |
| Aged | ASHFInt | Taphrinalesunclassified | 1 | 0.6 | 0.775 | 0.017 |
| Aged | ASHFInt | Physciaceae | 0.9777 | 0.6 | 0.766 | 0.049 |
| Aged | ASHFInt | Annulatascaceae | 0.9754 | 0.6 | 0.765 | 0.027 |
| Aged | ASHFInt | Plectosphaerellaceae | 0.9194 | 0.6 | 0.743 | 0.027 |
| Aged | ASHFInt | Puccinialesunclassified | 0.8438 | 0.6 | 0.712 | 0.037 |
| Aged | ASHFInt | Botryosphaeriaceae | 0.8276 | 0.6 | 0.705 | 0.039 |

| | | | | | | |
|---|---|---|---|---|---|---|
| Aged | ASHFExt | Lasiosphaeriaceae | 1 | 1 | 1 | 0.001 |
| Aged | ASHFExt | Pseudeurotiaceae | 0.9531 | 1 | 0.976 | 0.001 |
| Aged | ASHFExt | Trichosporonaceae | 1 | 0.6667 | 0.816 | 0.022 |
| Aged | WMCC | Tremellomycetesunclassified | 0.9833 | 1 | 0.992 | 0.001 |
| Aged | WMCC | unclassifiedChytridiomycetes | 1 | 0.6667 | 0.816 | 0.018 |
| Aged | WMCC | Chaetothyriaceae | 0.8333 | 0.6667 | 0.745 | 0.029 |
| Aged | ASHF | Mortierellaceae | 0.997 | 1 | 0.999 | 0.001 |
| Aged | ASHF | Filobasidiaceae | 0.9718 | 1 | 0.986 | 0.005 |
| Aged | ASHFInt+WMCC | Basidiomycotaunclassified | 0.9951 | 1 | 0.998 | 0.002 |
| Aged | ASHFInt+WMCC | Lophiostomataceae | 1 | 0.9091 | 0.953 | 0.003 |
| Aged | ASHFInt+WMCC | Dothideaceae | 0.9738 | 0.9091 | 0.941 | 0.004 |
| Aged | ASHFInt+WMCC | AscomycotafamilyIncertaesedis | 0.9795 | 0.8182 | 0.895 | 0.007 |
| Aged | Ext | Nectriaceae | 0.9939 | 0.9167 | 0.954 | 0.035 |
| Aged | Ext | HoltermannialesfamilyIncertaesedis | 1 | 0.8333 | 0.913 | 0.006 |
| Fresh | WMCC | Clavulinaceae | 0.9415 | 0.8333 | 0.886 | 0.005 |
| Fresh | WMCC | Schizophyllaceae | 0.9369 | 0.8333 | 0.884 | 0.032 |
| Fresh | WMCC | AuricularialesfamilyIncertaesedis | 0.8724 | 0.8333 | 0.853 | 0.047 |
| Fresh | WMCC | LeotiomycetesfamilyIncertaesedis | 0.7237 | 1 | 0.851 | 0.021 |
| Fresh | WMCC | Didymosphaeriaceae | 0.9969 | 0.6667 | 0.815 | 0.01 |
| Fresh | WMCC | Mycenaceae | 0.9735 | 0.6667 | 0.806 | 0.004 |
| Fresh | WMCC | Niaceae | 0.8696 | 0.6667 | 0.761 | 0.015 |
| Fresh | WMCC | unclassifiedExobasidiomycetes | 0.8505 | 0.6667 | 0.753 | 0.036 |
| Fresh | WMCC | unclassifiedAtheliales | 0.7506 | 0.6667 | 0.707 | 0.029 |
| Fresh | WMCC | Coniochaetaceae | 0.75 | 0.6667 | 0.707 | 0.03 |
| Fresh | WMCC | Lulworthiaceae | 1 | 0.5 | 0.707 | 0.026 |
| Fresh | WMCC | Physalacriaceae | 1 | 0.5 | 0.707 | 0.026 |
| Fresh | WMCC | Tulasnellaceae | 1 | 0.5 | 0.707 | 0.037 |
| Fresh | WMCC | Auricularialesunclassified | 1 | 0.5 | 0.707 | 0.024 |
| Fresh | WMCC | Pucciniomycetesunclassified | 0.9701 | 0.5 | 0.696 | 0.014 |

| | | | | | | |
|---|---|---|---|---|---|---|
| Fresh | WMCC | Bionectriaceae | 0.9091 | 0.5 | 0.674 | 0.031 |
| Fresh | ASHF | Mortierellaceae | 0.9675 | 0.9375 | 0.952 | 0.027 |
| Fresh | ASHF | unclassifiedPlantae | 0.9574 | 0.9375 | 0.947 | 0.009 |
| Fresh | ASHF | Umbelopsidaceae | 0.8291 | 0.875 | 0.852 | 0.049 |
| Fresh | Ext | unclassifiedPleosporales | 0.9673 | 0.75 | 0.852 | 0.033 |
| Fresh | Ext | Halosphaeriaceae | 1 | 0.5 | 0.707 | 0.038 |
