## Supplementary material for "Microbial, dietary insect, and pathogen communities in fresh and decomposing guano of anthropic little brown bat (Myotis lucifugus) maternity colonies": Table S16

| Barcode | Rename Sample ID | Original Read Count | Final Read Count (FRESH ONLY) - All Resequenced | Read N50 |
|---|---|---|---|---|
| barcode04 | 2508AshInt6a | 1,566 | 2,345.00 | 882 |
| barcode05 | 2508AshInt6b | 19,349 | 20,004.00 | 881 |
| barcode07 | 2508AshExt6c | 3,629 | 3,629.00 | 879 |
| barcode09 | 2508AshExt6a | 1,430 | 1,430.00 | 223 |
| barcode10 | 2507Wmcc6c | 4,148 | 4,763.00 | 881 |
| barcode11 | 2508AshInt6c | 178 | 339 | 257 |
| barcode14 | 2507AshInt6b | 933.00 | 1,761.00 | 315 |
| barcode15 | 2507Wmcc6a | 124 | 533 | 635 |
| barcode16 | 2507AshExt6a | 226 | 226 | 248 |
| barcode17 | 2508AshExt6b | 33 | 565 | 239 |
| barcode19 | 2507Wmcc6b | 871 | 985 | 874 |
| barcode22 | 2507AshExt6b | 3,681 | 3,955.00 | 233 |
| barcode23 | 2507AshExt6c | 14 | 130 | 258 |
| barcode24 | 2507AshInt6c | 2,348 | 2,406 | 275 |
| barcode25 | 2506AshInt6c | 422 | 428 | 873 |
| barcode26 | 2505AshInt6b | 800 | 1,239.00 | 876 |
| barcode27 | 2506AshExt6b | 383 | 387 | 611 |
| barcode28 | 2508Wmcc6a | 16 | 18 | 280 |
| barcode29 | 2508Wmcc6c | 1,016 | 1,031 | 874 |
| barcode31 | 2505AshExt6b | 117 | 119 | 665 |
| barcode32 | 2506Wmcc6b | 3,540 | 3,796 | 265 |
| barcode33 | 2506Wmcc6c | 87 | 273 | 665 |
| barcode34 | 2506Wmcc6a | 1 | 165 | 251 |
| barcode35 | 2508Wmcc6b | 115 | 121 | 423 |
| barcode36 | 2506AshExt6c | 257 | 617 | 257 |
| barcode38 | 2505AshInt6a | 10,191 | 10,223.00 | 881 |
| barcode40 | 2505AshInt6c | 783 | 800 | 881 |
| barcode41 | 2505AshExt6a | 219 | 342 | 247 |
| barcode42 | 2505AshExt6c | 44 | 62 | 524 |
| barcode48 | 2507AshInt6a | 65 | 74 | 855 |
| barcode49 | 2505Wmcc6c | 1,242 | 3,269.00 | 258 |
| barcode51 | 2505Wmcc6a | 1,097 | 1,176.00 | 880 |
| barcode52 | 2506AshExt6a | 1,634 | 1,861.00 | 883 |

| | | | | |
|---|---|---|---|---|
| barcode53 | 2506AshInt6a | 28 | 501 | 241 |
| barcode54 | 2506AshInt6b | 886 | 1,016 | 855 |
| barcode55 | 2505Wmcc6b | 94 | 188 | 869 |
| barcode67 | 2509AshExt6a | 90 | 99 | 604 |
| barcode68 | 2509AshExt6b | 2,952 | 2,965.00 | 878 |
| barcode69 | 2509Wmcc6a | 1,008 | 1,075.00 | 881 |
| barcode76 | 2509AshExt6c | 17 | 24 | 856 |
| barcode85 | 2509Wmcc6b | 89 | 462 | 871 |
| barcode88 | 2509Wmcc6c | 1,593 | 5,762.00 | 881 |
| | | | 1,932.48 | |
| | | | 20,004.00 | |
| | | | 18.00 | 606.3095238 |
