## Supplementary material for "Microbial, dietary insect, and pathogen communities in fresh and decomposing guano of anthropic little brown bat (Myotis lucifugus) maternity colonies": Table S17

| Sample ID | Site | Month | Order | Family | Genus |
|---|---|---|---|---|---|
| 2505AshInt6a | AshInt | May | Araneae | Agelenidae | Coras |
| 2505AshInt6a | AshInt | May | Araneae | Amaurobiidae | Callobius |
| 2505AshInt6a | AshInt | May | Araneae | Anyphaenidae | Anyphaena |
| 2505AshInt6a | AshInt | May | Araneae | Cicurinidae | Cicurina |
| 2505AshInt6a | AshInt | May | Araneae | Dictynidae | Dictyna |
| 2505AshInt6a | AshInt | May | Araneae | Gnaphosidae | Gnaphosa |
| 2505AshInt6a | AshInt | May | Araneae | Linyphiidae | Grammonota |
| 2505AshInt6a | AshInt | May | Araneae | Lycosidae | Rabidosa |
| 2505AshInt6a | AshInt | May | Araneae | Lycosidae | Schizocosa |
| 2505AshInt6a | AshInt | May | Araneae | Phrurolithidae | Scotinella |
| 2505AshInt6a | AshInt | May | Araneae | Salticidae | Maevia |
| 2505AshInt6a | AshInt | May | Araneae | Salticidae | Sibianor |
| 2505AshInt6a | AshInt | May | Araneae | Theridiidae | Canalidion |
| 2505AshInt6a | AshInt | May | Araneae | Theridiidae | Rugathodes |
| 2505AshInt6a | AshInt | May | Araneae | Theridiidae | Spintharus |
| 2505AshInt6a | AshInt | May | Araneae | Thomisidae | Mecaphesa |
| 2505AshInt6a | AshInt | May | Blattodea | Ectobiidae | Parcoblatta |
| 2505AshInt6a | AshInt | May | Coleoptera | Buprestidae | Pachyschelus |
| 2505AshInt6a | AshInt | May | Coleoptera | Carabidae | Apenes |
| 2505AshInt6a | AshInt | May | Coleoptera | Carabidae | Harpalus |
| 2505AshInt6a | AshInt | May | Coleoptera | Carabidae | Other |
| 2505AshInt6a | AshInt | May | Coleoptera | Carabidae | Polyderis |
| 2505AshInt6a | AshInt | May | Coleoptera | Curculionidae | Conotrachelus |
| 2505AshInt6a | AshInt | May | Coleoptera | Curculionidae | Cyrtepistomus |
| 2505AshInt6a | AshInt | May | Coleoptera | Curculionidae | Listronotus |
| 2505AshInt6a | AshInt | May | Coleoptera | Dytiscidae | Acilius |
| 2505AshInt6a | AshInt | May | Coleoptera | Lampyridae | Photinus |
| 2505AshInt6a | AshInt | May | Coleoptera | Scarabaeidae | Dialytes |
| 2505AshInt6a | AshInt | May | Coleoptera | Scarabaeidae | Melanocanthon |
| 2505AshInt6a | AshInt | May | Coleoptera | Scirtidae | Contacyphon |
| 2505AshInt6a | AshInt | May | Diptera | Agromyzidae | Calycomyza |

| | | | | | |
|---|---|---|---|---|---|
| 2505AshInt6a | AshInt | May | Diptera | Agromyzidae | Cerodontha |
| 2505AshInt6a | AshInt | May | Diptera | Agromyzidae | Japanagromyza |
| 2505AshInt6a | AshInt | May | Diptera | Agromyzidae | Liriomyza |
| 2505AshInt6a | AshInt | May | Diptera | Agromyzidae | Nemorimyza |
| 2505AshInt6a | AshInt | May | Diptera | Agromyzidae | Phytomyza |
| 2505AshInt6a | AshInt | May | Diptera | Anisopodidae | Mycetobia |
| 2505AshInt6a | AshInt | May | Diptera | Calliphoridae | Lucilia |
| 2505AshInt6a | AshInt | May | Diptera | Cecidomyiidae | Other |
| 2505AshInt6a | AshInt | May | Diptera | Chaoboridae | Chaoborus |
| 2505AshInt6a | AshInt | May | Diptera | Chaoboridae | Other |
| 2505AshInt6a | AshInt | May | Diptera | Chironomidae | Ablabesmyia |
| 2505AshInt6a | AshInt | May | Diptera | Chironomidae | Chironomus |
| 2505AshInt6a | AshInt | May | Diptera | Chironomidae | Dicrotendipes |
| 2505AshInt6a | AshInt | May | Diptera | Chironomidae | Endochironomus |
| 2505AshInt6a | AshInt | May | Diptera | Chironomidae | Glyptotendipes |
| 2505AshInt6a | AshInt | May | Diptera | Chironomidae | Microtendipes |
| 2505AshInt6a | AshInt | May | Diptera | Chironomidae | Orthocladius |
| 2505AshInt6a | AshInt | May | Diptera | Chironomidae | Other |
| 2505AshInt6a | AshInt | May | Diptera | Chironomidae | Other |
| 2505AshInt6a | AshInt | May | Diptera | Chironomidae | Other |
| 2505AshInt6a | AshInt | May | Diptera | Chironomidae | Other |
| 2505AshInt6a | AshInt | May | Diptera | Chironomidae | Parachironomus |
| 2505AshInt6a | AshInt | May | Diptera | Chironomidae | Polypedilum |
| 2505AshInt6a | AshInt | May | Diptera | Chironomidae | Pseudochironomus |
| 2505AshInt6a | AshInt | May | Diptera | Chironomidae | Tanytarsus |
| 2505AshInt6a | AshInt | May | Diptera | Chironomidae | Xenochironomus |
| 2505AshInt6a | AshInt | May | Diptera | Chloropidae | Apotropina |
| 2505AshInt6a | AshInt | May | Diptera | Chloropidae | Elachiptera |
| 2505AshInt6a | AshInt | May | Diptera | Chloropidae | Liohippelates |
| 2505AshInt6a | AshInt | May | Diptera | Chloropidae | Oscinella |
| 2505AshInt6a | AshInt | May | Diptera | Culicidae | Coquillettidia |

| | | | | | |
|---|---|---|---|---|---|
| 2505AshInt6a | AshInt | May | Diptera | Culicidae | Culex |
| 2505AshInt6a | AshInt | May | Diptera | Culicidae | Uranotaenia |
| 2505AshInt6a | AshInt | May | Diptera | Drosophilidae | Drosophila |
| 2505AshInt6a | AshInt | May | Diptera | Empididae | Other |
| 2505AshInt6a | AshInt | May | Diptera | Hybotidae | Platypalpus |
| 2505AshInt6a | AshInt | May | Diptera | Hybotidae | Syndyas |
| 2505AshInt6a | AshInt | May | Diptera | Keroplatidae | Other |
| 2505AshInt6a | AshInt | May | Diptera | Limoniidae | Dicranomyia |
| 2505AshInt6a | AshInt | May | Diptera | Limoniidae | Dicranophragma |
| 2505AshInt6a | AshInt | May | Diptera | Limoniidae | Epiphragma |
| 2505AshInt6a | AshInt | May | Diptera | Limoniidae | Erioptera |
| 2505AshInt6a | AshInt | May | Diptera | Limoniidae | Metalimnobia |
| 2505AshInt6a | AshInt | May | Diptera | Limoniidae | Ormosia |
| 2505AshInt6a | AshInt | May | Diptera | Limoniidae | Other |
| 2505AshInt6a | AshInt | May | Diptera | Limoniidae | Other |
| 2505AshInt6a | AshInt | May | Diptera | Limoniidae | Pseudolimnophila |
| 2505AshInt6a | AshInt | May | Diptera | Muscidae | Mydaea |
| 2505AshInt6a | AshInt | May | Diptera | Mycetophilidae | Mycomya |
| 2505AshInt6a | AshInt | May | Diptera | Mycetophilidae | Neoempheria |
| 2505AshInt6a | AshInt | May | Diptera | Opomyzidae | Geomyza |
| 2505AshInt6a | AshInt | May | Diptera | Pallopteridae | Toxonevra |
| 2505AshInt6a | AshInt | May | Diptera | Phoridae | Diplonevra |
| 2505AshInt6a | AshInt | May | Diptera | Polleniidae | Pollenia |
| 2505AshInt6a | AshInt | May | Diptera | Psychodidae | Clogmia |
| 2505AshInt6a | AshInt | May | Diptera | Psychodidae | Psychoda |
| 2505AshInt6a | AshInt | May | Diptera | Sarcophagidae | Ravinia |
| 2505AshInt6a | AshInt | May | Diptera | Sciaridae | Claustropyga |
| 2505AshInt6a | AshInt | May | Diptera | Sciaridae | Corynoptera |
| 2505AshInt6a | AshInt | May | Diptera | Sciaridae | Epidapus |
| 2505AshInt6a | AshInt | May | Diptera | Sciaridae | Other |
| 2505AshInt6a | AshInt | May | Diptera | Sphaeroceridae | Leptocera |

| | | | | | |
|---|---|---|---|---|---|
| 2505AshInt6a | AshInt | May | Diptera | Syrphidae | Chalcosyrphus |
| 2505AshInt6a | AshInt | May | Diptera | Syrphidae | Palpada |
| 2505AshInt6a | AshInt | May | Diptera | Tabanidae | Hybomitra |
| 2505AshInt6a | AshInt | May | Diptera | Tabanidae | Other |
| 2505AshInt6a | AshInt | May | Diptera | Tachinidae | Cryptomeigenia |
| 2505AshInt6a | AshInt | May | Diptera | Tachinidae | Cyzenis |
| 2505AshInt6a | AshInt | May | Diptera | Tachinidae | Zaira |
| 2505AshInt6a | AshInt | May | Diptera | Tipulidae | Dolichopeza |
| 2505AshInt6a | AshInt | May | Diptera | Tipulidae | Tipula |
| 2505AshInt6a | AshInt | May | Ephemeroptera | Caenidae | Caenis |
| 2505AshInt6a | AshInt | May | Ephemeroptera | Ephemeridae | Hexagenia |
| 2505AshInt6a | AshInt | May | Ephemeroptera | Heptageniidae | Epeorus |
| 2505AshInt6a | AshInt | May | Hemiptera | Clastopteridae | Clastoptera |
| 2505AshInt6a | AshInt | May | Hemiptera | Lygaeidae | Kleidocerys |
| 2505AshInt6a | AshInt | May | Hymenoptera | Apidae | Triepeolus |
| 2505AshInt6a | AshInt | May | Hymenoptera | Braconidae | Bracon |
| 2505AshInt6a | AshInt | May | Hymenoptera | Braconidae | Chorebus |
| 2505AshInt6a | AshInt | May | Hymenoptera | Ichneumonidae | Megastylus |
| 2505AshInt6a | AshInt | May | Hymenoptera | Ichneumonidae | Polyblastus |
| 2505AshInt6a | AshInt | May | Hymenoptera | Other | Other |
| 2505AshInt6a | AshInt | May | Hymenoptera | Tenthredinidae | Dolerus |
| 2505AshInt6a | AshInt | May | Hymenoptera | Tenthredinidae | Nematus |
| 2505AshInt6a | AshInt | May | Hymenoptera | Trigonalidae | Orthogonalys |
| 2505AshInt6a | AshInt | May | Lepidoptera | Crambidae | Blepharomastix |
| 2505AshInt6a | AshInt | May | Lepidoptera | Crambidae | Parapoynx |
| 2505AshInt6a | AshInt | May | Lepidoptera | Elachistidae | Elachista |
| 2505AshInt6a | AshInt | May | Lepidoptera | Erebidae | Panopoda |
| 2505AshInt6a | AshInt | May | Lepidoptera | Gelechiidae | Chionodes |
| 2505AshInt6a | AshInt | May | Lepidoptera | Noctuidae | Condica |
| 2505AshInt6a | AshInt | May | Lepidoptera | Noctuidae | Leuconycta |
| 2505AshInt6a | AshInt | May | Lepidoptera | Noctuidae | Pyreferra |

| | | | | | |
|---|---|---|---|---|---|
| 2505AshInt6a | AshInt | May | Lepidoptera | Notodontidae | Heterocampa |
| 2505AshInt6a | AshInt | May | Lepidoptera | Notodontidae | Hyparpax |
| 2505AshInt6a | AshInt | May | Lepidoptera | Nymphalidae | Oeneis |
| 2505AshInt6a | AshInt | May | Lepidoptera | Pyralidae | Atheloca |
| 2505AshInt6a | AshInt | May | Lepidoptera | Pyralidae | Elasmopalpus |
| 2505AshInt6a | AshInt | May | Lepidoptera | Tischeriidae | Coptotriche |
| 2505AshInt6a | AshInt | May | Lepidoptera | Tortricidae | Acleris |
| 2505AshInt6a | AshInt | May | Lepidoptera | Tortricidae | Argyrotaenia |
| 2505AshInt6a | AshInt | May | Lepidoptera | Tortricidae | Rhyacionia |
| 2505AshInt6a | AshInt | May | Megaloptera | Corydalidae | Corydalus |
| 2505AshInt6a | AshInt | May | Neuroptera | Hemerobiidae | Hemerobius |
| 2505AshInt6a | AshInt | May | Odonata | Libellulidae | Brachymesia |
| 2505AshInt6a | AshInt | May | Orthoptera | Tettigoniidae | Conocephalus |
| 2505AshInt6a | AshInt | May | Other | Other | Other |
| 2505AshInt6a | AshInt | May | Other | Other | Other |
| 2505AshInt6a | AshInt | May | Plecoptera | Chloroperlidae | Sweltsa |
| 2505AshInt6a | AshInt | May | Psocodea | Peripsocidae | Peripsocus |
| 2505AshInt6a | AshInt | May | Sarcoptiformes | Oripodidae | Other |
| 2505AshInt6a | AshInt | May | Trichoptera | Dipseudopsidae | Phylocentropus |
| 2505AshInt6a | AshInt | May | Trichoptera | Hydroptilidae | Orthotrichia |
| 2505AshInt6a | AshInt | May | Trichoptera | Leptoceridae | Oecetis |
| 2505AshInt6a | AshInt | May | Trombidiformes | Arrenuridae | Arrenurus |
| 2505AshInt6a | AshInt | May | Trombidiformes | Aturidae | Aturus |
| 2505AshInt6a | AshInt | May | Trombidiformes | Axonopsidae | Brachypoda |
| 2505AshInt6a | AshInt | May | Trombidiformes | Eupodidae | Other |
| 2505AshInt6a | AshInt | May | Trombidiformes | Feltriidae | Feltria |
| 2505AshInt6a | AshInt | May | Trombidiformes | Hydryphantidae | Hydryphantes |
| 2505AshInt6a | AshInt | May | Trombidiformes | Hydryphantidae | Parathyas |
| 2505AshInt6a | AshInt | May | Trombidiformes | Lebertiidae | Lebertia |
| 2505AshInt6a | AshInt | May | Trombidiformes | Mideopsidae | Mideopsis |
| 2505AshInt6a | AshInt | May | Trombidiformes | Pionidae | Piona |

| | | | | | |
|---|---|---|---|---|---|
| 2505AshInt6a | AshInt | May | Trombidiformes | Pionidae | Tiphys |
| 2505AshInt6a | AshInt | May | Trombidiformes | Sperchontidae | Sperchon |
| 2505AshInt6a | AshInt | May | Trombidiformes | Tarsonemidae | Phytonemus |
| 2505AshInt6a | AshInt | May | Trombidiformes | Torrenticolidae | Testudacarus |
| 2505AshInt6a | AshInt | May | Trombidiformes | Torrenticolidae | Torrenticola |
| 2505AshInt6a | AshInt | May | Trombidiformes | Unionicolidae | Koenikea |
| 2505AshInt6b | AshInt | May | Araneae | Agelenidae | Coras |
| 2505AshInt6b | AshInt | May | Araneae | Amaurobiidae | Callobius |
| 2505AshInt6b | AshInt | May | Araneae | Anyphaenidae | Anyphaena |
| 2505AshInt6b | AshInt | May | Araneae | Cicurinidae | Cicurina |
| 2505AshInt6b | AshInt | May | Araneae | Dictynidae | Dictyna |
| 2505AshInt6b | AshInt | May | Araneae | Gnaphosidae | Gnaphosa |
| 2505AshInt6b | AshInt | May | Araneae | Linyphiidae | Grammonota |
| 2505AshInt6b | AshInt | May | Araneae | Lycosidae | Rabidosa |
| 2505AshInt6b | AshInt | May | Araneae | Lycosidae | Schizocosa |
| 2505AshInt6b | AshInt | May | Araneae | Phrurolithidae | Scotinella |
| 2505AshInt6b | AshInt | May | Araneae | Salticidae | Maevia |
| 2505AshInt6b | AshInt | May | Araneae | Salticidae | Sibianor |
| 2505AshInt6b | AshInt | May | Araneae | Theridiidae | Canalidion |
| 2505AshInt6b | AshInt | May | Araneae | Theridiidae | Rugathodes |
| 2505AshInt6b | AshInt | May | Araneae | Theridiidae | Spintharus |
| 2505AshInt6b | AshInt | May | Araneae | Thomisidae | Mecaphesa |
| 2505AshInt6b | AshInt | May | Blattodea | Ectobiidae | Parcoblatta |
| 2505AshInt6b | AshInt | May | Coleoptera | Buprestidae | Pachyschelus |
| 2505AshInt6b | AshInt | May | Coleoptera | Carabidae | Apenes |
| 2505AshInt6b | AshInt | May | Coleoptera | Carabidae | Harpalus |
| 2505AshInt6b | AshInt | May | Coleoptera | Carabidae | Other |
| 2505AshInt6b | AshInt | May | Coleoptera | Carabidae | Polyderis |
| 2505AshInt6b | AshInt | May | Coleoptera | Curculionidae | Conotrachelus |
| 2505AshInt6b | AshInt | May | Coleoptera | Curculionidae | Cyrtepistomus |
| 2505AshInt6b | AshInt | May | Coleoptera | Curculionidae | Listronotus |

| | | | | | |
|---|---|---|---|---|---|
| 2505AshInt6b | AshInt | May | Coleoptera | Dytiscidae | Acilius |
| 2505AshInt6b | AshInt | May | Coleoptera | Lampyridae | Photinus |
| 2505AshInt6b | AshInt | May | Coleoptera | Scarabaeidae | Dialytes |
| 2505AshInt6b | AshInt | May | Coleoptera | Scarabaeidae | Melanocanthon |
| 2505AshInt6b | AshInt | May | Coleoptera | Scirtidae | Contacyphon |
| 2505AshInt6b | AshInt | May | Diptera | Agromyzidae | Calycomyza |
| 2505AshInt6b | AshInt | May | Diptera | Agromyzidae | Cerodontha |
| 2505AshInt6b | AshInt | May | Diptera | Agromyzidae | Japanagromyza |
| 2505AshInt6b | AshInt | May | Diptera | Agromyzidae | Liriomyza |
| 2505AshInt6b | AshInt | May | Diptera | Agromyzidae | Nemorimyza |
| 2505AshInt6b | AshInt | May | Diptera | Agromyzidae | Phytomyza |
| 2505AshInt6b | AshInt | May | Diptera | Anisopodidae | Mycetobia |
| 2505AshInt6b | AshInt | May | Diptera | Calliphoridae | Lucilia |
| 2505AshInt6b | AshInt | May | Diptera | Cecidomyiidae | Other |
| 2505AshInt6b | AshInt | May | Diptera | Chaoboridae | Chaoborus |
| 2505AshInt6b | AshInt | May | Diptera | Chaoboridae | Other |
| 2505AshInt6b | AshInt | May | Diptera | Chironomidae | Ablabesmyia |
| 2505AshInt6b | AshInt | May | Diptera | Chironomidae | Chironomus |
| 2505AshInt6b | AshInt | May | Diptera | Chironomidae | Dicrotendipes |
| 2505AshInt6b | AshInt | May | Diptera | Chironomidae | Endochironomus |
| 2505AshInt6b | AshInt | May | Diptera | Chironomidae | Glyptotendipes |
| 2505AshInt6b | AshInt | May | Diptera | Chironomidae | Microtendipes |
| 2505AshInt6b | AshInt | May | Diptera | Chironomidae | Orthocladius |
| 2505AshInt6b | AshInt | May | Diptera | Chironomidae | Other |
| 2505AshInt6b | AshInt | May | Diptera | Chironomidae | Other |
| 2505AshInt6b | AshInt | May | Diptera | Chironomidae | Other |
| 2505AshInt6b | AshInt | May | Diptera | Chironomidae | Other |
| 2505AshInt6b | AshInt | May | Diptera | Chironomidae | Parachironomus |
| 2505AshInt6b | AshInt | May | Diptera | Chironomidae | Polypedilum |
| 2505AshInt6b | AshInt | May | Diptera | Chironomidae | Pseudochironomus |
| 2505AshInt6b | AshInt | May | Diptera | Chironomidae | Tanytarsus |

| | | | | | |
|---|---|---|---|---|---|
| 2505AshInt6b | AshInt | May | Diptera | Chironomidae | Xenochironomus |
| 2505AshInt6b | AshInt | May | Diptera | Chloropidae | Apotropina |
| 2505AshInt6b | AshInt | May | Diptera | Chloropidae | Elachiptera |
| 2505AshInt6b | AshInt | May | Diptera | Chloropidae | Liohippelates |
| 2505AshInt6b | AshInt | May | Diptera | Chloropidae | Oscinella |
| 2505AshInt6b | AshInt | May | Diptera | Culicidae | Coquillettidia |
| 2505AshInt6b | AshInt | May | Diptera | Culicidae | Culex |
| 2505AshInt6b | AshInt | May | Diptera | Culicidae | Uranotaenia |
| 2505AshInt6b | AshInt | May | Diptera | Drosophilidae | Drosophila |
| 2505AshInt6b | AshInt | May | Diptera | Empididae | Other |
| 2505AshInt6b | AshInt | May | Diptera | Hybotidae | Platypalpus |
| 2505AshInt6b | AshInt | May | Diptera | Hybotidae | Syndyas |
| 2505AshInt6b | AshInt | May | Diptera | Keroplatidae | Other |
| 2505AshInt6b | AshInt | May | Diptera | Limoniidae | Dicranomyia |
| 2505AshInt6b | AshInt | May | Diptera | Limoniidae | Dicranophragma |
| 2505AshInt6b | AshInt | May | Diptera | Limoniidae | Epiphragma |
| 2505AshInt6b | AshInt | May | Diptera | Limoniidae | Erioptera |
| 2505AshInt6b | AshInt | May | Diptera | Limoniidae | Metalimnobia |
| 2505AshInt6b | AshInt | May | Diptera | Limoniidae | Ormosia |
| 2505AshInt6b | AshInt | May | Diptera | Limoniidae | Other |
| 2505AshInt6b | AshInt | May | Diptera | Limoniidae | Other |
| 2505AshInt6b | AshInt | May | Diptera | Limoniidae | Pseudolimnophila |
| 2505AshInt6b | AshInt | May | Diptera | Muscidae | Mydaea |
| 2505AshInt6b | AshInt | May | Diptera | Mycetophilidae | Mycomya |
| 2505AshInt6b | AshInt | May | Diptera | Mycetophilidae | Neoempheria |
| 2505AshInt6b | AshInt | May | Diptera | Opomyzidae | Geomyza |
| 2505AshInt6b | AshInt | May | Diptera | Pallopteridae | Toxonevra |
| 2505AshInt6b | AshInt | May | Diptera | Phoridae | Diplonevra |
| 2505AshInt6b | AshInt | May | Diptera | Polleniidae | Pollenia |
| 2505AshInt6b | AshInt | May | Diptera | Psychodidae | Clogmia |
| 2505AshInt6b | AshInt | May | Diptera | Psychodidae | Psychoda |

| | | | | | |
|---|---|---|---|---|---|
| 2505AshInt6b | AshInt | May | Diptera | Sarcophagidae | Ravinia |
| 2505AshInt6b | AshInt | May | Diptera | Sciaridae | Claustropyga |
| 2505AshInt6b | AshInt | May | Diptera | Sciaridae | Corynoptera |
| 2505AshInt6b | AshInt | May | Diptera | Sciaridae | Epidapus |
| 2505AshInt6b | AshInt | May | Diptera | Sciaridae | Other |
| 2505AshInt6b | AshInt | May | Diptera | Sphaeroceridae | Leptocera |
| 2505AshInt6b | AshInt | May | Diptera | Syrphidae | Chalcosyrphus |
| 2505AshInt6b | AshInt | May | Diptera | Syrphidae | Palpada |
| 2505AshInt6b | AshInt | May | Diptera | Tabanidae | Hybomitra |
| 2505AshInt6b | AshInt | May | Diptera | Tabanidae | Other |
| 2505AshInt6b | AshInt | May | Diptera | Tachinidae | Cryptomeigenia |
| 2505AshInt6b | AshInt | May | Diptera | Tachinidae | Cyzenis |
| 2505AshInt6b | AshInt | May | Diptera | Tachinidae | Zaira |
| 2505AshInt6b | AshInt | May | Diptera | Tipulidae | Dolichopeza |
| 2505AshInt6b | AshInt | May | Diptera | Tipulidae | Tipula |
| 2505AshInt6b | AshInt | May | Ephemeroptera | Caenidae | Caenis |
| 2505AshInt6b | AshInt | May | Ephemeroptera | Ephemeridae | Hexagenia |
| 2505AshInt6b | AshInt | May | Ephemeroptera | Heptageniidae | Epeorus |
| 2505AshInt6b | AshInt | May | Hemiptera | Clastopteridae | Clastoptera |
| 2505AshInt6b | AshInt | May | Hemiptera | Lygaeidae | Kleidocerys |
| 2505AshInt6b | AshInt | May | Hymenoptera | Apidae | Triepeolus |
| 2505AshInt6b | AshInt | May | Hymenoptera | Braconidae | Bracon |
| 2505AshInt6b | AshInt | May | Hymenoptera | Braconidae | Chorebus |
| 2505AshInt6b | AshInt | May | Hymenoptera | Ichneumonidae | Megastylus |
| 2505AshInt6b | AshInt | May | Hymenoptera | Ichneumonidae | Polyblastus |
| 2505AshInt6b | AshInt | May | Hymenoptera | Other | Other |
| 2505AshInt6b | AshInt | May | Hymenoptera | Tenthredinidae | Dolerus |
| 2505AshInt6b | AshInt | May | Hymenoptera | Tenthredinidae | Nematus |
| 2505AshInt6b | AshInt | May | Hymenoptera | Trigonalidae | Orthogonalys |
| 2505AshInt6b | AshInt | May | Lepidoptera | Crambidae | Blepharomastix |
| 2505AshInt6b | AshInt | May | Lepidoptera | Crambidae | Parapoynx |

| | | | | | |
|---|---|---|---|---|---|
| 2505AshInt6b | AshInt | May | Lepidoptera | Elachistidae | Elachista |
| 2505AshInt6b | AshInt | May | Lepidoptera | Erebidae | Panopoda |
| 2505AshInt6b | AshInt | May | Lepidoptera | Gelechiidae | Chionodes |
| 2505AshInt6b | AshInt | May | Lepidoptera | Noctuidae | Condica |
| 2505AshInt6b | AshInt | May | Lepidoptera | Noctuidae | Leuconycta |
| 2505AshInt6b | AshInt | May | Lepidoptera | Noctuidae | Pyreferra |
| 2505AshInt6b | AshInt | May | Lepidoptera | Notodontidae | Heterocampa |
| 2505AshInt6b | AshInt | May | Lepidoptera | Notodontidae | Hyparpax |
| 2505AshInt6b | AshInt | May | Lepidoptera | Nymphalidae | Oeneis |
| 2505AshInt6b | AshInt | May | Lepidoptera | Pyralidae | Atheloca |
| 2505AshInt6b | AshInt | May | Lepidoptera | Pyralidae | Elasmopalpus |
| 2505AshInt6b | AshInt | May | Lepidoptera | Tischeriidae | Coptotriche |
| 2505AshInt6b | AshInt | May | Lepidoptera | Tortricidae | Acleris |
| 2505AshInt6b | AshInt | May | Lepidoptera | Tortricidae | Argyrotaenia |
| 2505AshInt6b | AshInt | May | Lepidoptera | Tortricidae | Rhyacionia |
| 2505AshInt6b | AshInt | May | Megaloptera | Corydalidae | Corydalus |
| 2505AshInt6b | AshInt | May | Neuroptera | Hemerobiidae | Hemerobius |
| 2505AshInt6b | AshInt | May | Odonata | Libellulidae | Brachymesia |
| 2505AshInt6b | AshInt | May | Orthoptera | Tettigoniidae | Conocephalus |
| 2505AshInt6b | AshInt | May | Other | Other | Other |
| 2505AshInt6b | AshInt | May | Other | Other | Other |
| 2505AshInt6b | AshInt | May | Plecoptera | Chloroperlidae | Sweltsa |
| 2505AshInt6b | AshInt | May | Psocodea | Peripsocidae | Peripsocus |
| 2505AshInt6b | AshInt | May | Sarcoptiformes | Oripodidae | Other |
| 2505AshInt6b | AshInt | May | Trichoptera | Dipseudopsidae | Phylocentropus |
| 2505AshInt6b | AshInt | May | Trichoptera | Hydroptilidae | Orthotrichia |
| 2505AshInt6b | AshInt | May | Trichoptera | Leptoceridae | Oecetis |
| 2505AshInt6b | AshInt | May | Trombidiformes | Arrenuridae | Arrenurus |
| 2505AshInt6b | AshInt | May | Trombidiformes | Aturidae | Aturus |
| 2505AshInt6b | AshInt | May | Trombidiformes | Axonopsidae | Brachypoda |
| 2505AshInt6b | AshInt | May | Trombidiformes | Eupodidae | Other |

| | | | | | |
|---|---|---|---|---|---|
| 2505AshInt6b | AshInt | May | Trombidiformes | Feltriidae | Feltria |
| 2505AshInt6b | AshInt | May | Trombidiformes | Hydryphantidae | Hydryphantes |
| 2505AshInt6b | AshInt | May | Trombidiformes | Hydryphantidae | Parathyas |
| 2505AshInt6b | AshInt | May | Trombidiformes | Lebertiidae | Lebertia |
| 2505AshInt6b | AshInt | May | Trombidiformes | Mideopsidae | Mideopsis |
| 2505AshInt6b | AshInt | May | Trombidiformes | Pionidae | Piona |
| 2505AshInt6b | AshInt | May | Trombidiformes | Pionidae | Tiphys |
| 2505AshInt6b | AshInt | May | Trombidiformes | Sperchontidae | Sperchon |
| 2505AshInt6b | AshInt | May | Trombidiformes | Tarsonemidae | Phytonemus |
| 2505AshInt6b | AshInt | May | Trombidiformes | Torrenticolidae | Testudacarus |
| 2505AshInt6b | AshInt | May | Trombidiformes | Torrenticolidae | Torrenticola |
| 2505AshInt6b | AshInt | May | Trombidiformes | Unionicolidae | Koenikea |
| 2505AshInt6c | AshInt | May | Araneae | Agelenidae | Coras |
| 2505AshInt6c | AshInt | May | Araneae | Amaurobiidae | Callobius |
| 2505AshInt6c | AshInt | May | Araneae | Anyphaenidae | Anyphaena |
| 2505AshInt6c | AshInt | May | Araneae | Cicurinidae | Cicurina |
| 2505AshInt6c | AshInt | May | Araneae | Dictynidae | Dictyna |
| 2505AshInt6c | AshInt | May | Araneae | Gnaphosidae | Gnaphosa |
| 2505AshInt6c | AshInt | May | Araneae | Linyphiidae | Grammonota |
| 2505AshInt6c | AshInt | May | Araneae | Lycosidae | Rabidosa |
| 2505AshInt6c | AshInt | May | Araneae | Lycosidae | Schizocosa |
| 2505AshInt6c | AshInt | May | Araneae | Phrurolithidae | Scotinella |
| 2505AshInt6c | AshInt | May | Araneae | Salticidae | Maevia |
| 2505AshInt6c | AshInt | May | Araneae | Salticidae | Sibianor |
| 2505AshInt6c | AshInt | May | Araneae | Theridiidae | Canalidion |
| 2505AshInt6c | AshInt | May | Araneae | Theridiidae | Rugathodes |
| 2505AshInt6c | AshInt | May | Araneae | Theridiidae | Spintharus |
| 2505AshInt6c | AshInt | May | Araneae | Thomisidae | Mecaphesa |
| 2505AshInt6c | AshInt | May | Blattodea | Ectobiidae | Parcoblatta |
| 2505AshInt6c | AshInt | May | Coleoptera | Buprestidae | Pachyschelus |
| 2505AshInt6c | AshInt | May | Coleoptera | Carabidae | Apenes |

| | | | | | |
|---|---|---|---|---|---|
| 2505AshInt6c | AshInt | May | Coleoptera | Carabidae | Harpalus |
| 2505AshInt6c | AshInt | May | Coleoptera | Carabidae | Other |
| 2505AshInt6c | AshInt | May | Coleoptera | Carabidae | Polyderis |
| 2505AshInt6c | AshInt | May | Coleoptera | Curculionidae | Conotrachelus |
| 2505AshInt6c | AshInt | May | Coleoptera | Curculionidae | Cyrtepistomus |
| 2505AshInt6c | AshInt | May | Coleoptera | Curculionidae | Listronotus |
| 2505AshInt6c | AshInt | May | Coleoptera | Dytiscidae | Acilius |
| 2505AshInt6c | AshInt | May | Coleoptera | Lampyridae | Photinus |
| 2505AshInt6c | AshInt | May | Coleoptera | Scarabaeidae | Dialytes |
| 2505AshInt6c | AshInt | May | Coleoptera | Scarabaeidae | Melanocanthon |
| 2505AshInt6c | AshInt | May | Coleoptera | Scirtidae | Contacyphon |
| 2505AshInt6c | AshInt | May | Diptera | Agromyzidae | Calycomyza |
| 2505AshInt6c | AshInt | May | Diptera | Agromyzidae | Cerodontha |
| 2505AshInt6c | AshInt | May | Diptera | Agromyzidae | Japanagromyza |
| 2505AshInt6c | AshInt | May | Diptera | Agromyzidae | Liriomyza |
| 2505AshInt6c | AshInt | May | Diptera | Agromyzidae | Nemorimyza |
| 2505AshInt6c | AshInt | May | Diptera | Agromyzidae | Phytomyza |
| 2505AshInt6c | AshInt | May | Diptera | Anisopodidae | Mycetobia |
| 2505AshInt6c | AshInt | May | Diptera | Calliphoridae | Lucilia |
| 2505AshInt6c | AshInt | May | Diptera | Cecidomyiidae | Other |
| 2505AshInt6c | AshInt | May | Diptera | Chaoboridae | Chaoborus |
| 2505AshInt6c | AshInt | May | Diptera | Chaoboridae | Other |
| 2505AshInt6c | AshInt | May | Diptera | Chironomidae | Ablabesmyia |
| 2505AshInt6c | AshInt | May | Diptera | Chironomidae | Chironomus |
| 2505AshInt6c | AshInt | May | Diptera | Chironomidae | Dicrotendipes |
| 2505AshInt6c | AshInt | May | Diptera | Chironomidae | Endochironomus |
| 2505AshInt6c | AshInt | May | Diptera | Chironomidae | Glyptotendipes |
| 2505AshInt6c | AshInt | May | Diptera | Chironomidae | Microtendipes |
| 2505AshInt6c | AshInt | May | Diptera | Chironomidae | Orthocladius |
| 2505AshInt6c | AshInt | May | Diptera | Chironomidae | Other |
| 2505AshInt6c | AshInt | May | Diptera | Chironomidae | Other |

| | | | | | |
|---|---|---|---|---|---|
| 2505AshInt6c | AshInt | May | Diptera | Chironomidae | Other |
| 2505AshInt6c | AshInt | May | Diptera | Chironomidae | Other |
| 2505AshInt6c | AshInt | May | Diptera | Chironomidae | Parachironomus |
| 2505AshInt6c | AshInt | May | Diptera | Chironomidae | Polypedilum |
| 2505AshInt6c | AshInt | May | Diptera | Chironomidae | Pseudochironomus |
| 2505AshInt6c | AshInt | May | Diptera | Chironomidae | Tanytarsus |
| 2505AshInt6c | AshInt | May | Diptera | Chironomidae | Xenochironomus |
| 2505AshInt6c | AshInt | May | Diptera | Chloropidae | Apotropina |
| 2505AshInt6c | AshInt | May | Diptera | Chloropidae | Elachiptera |
| 2505AshInt6c | AshInt | May | Diptera | Chloropidae | Liohippelates |
| 2505AshInt6c | AshInt | May | Diptera | Chloropidae | Oscinella |
| 2505AshInt6c | AshInt | May | Diptera | Culicidae | Coquillettidia |
| 2505AshInt6c | AshInt | May | Diptera | Culicidae | Culex |
| 2505AshInt6c | AshInt | May | Diptera | Culicidae | Uranotaenia |
| 2505AshInt6c | AshInt | May | Diptera | Drosophilidae | Drosophila |
| 2505AshInt6c | AshInt | May | Diptera | Empididae | Other |
| 2505AshInt6c | AshInt | May | Diptera | Hybotidae | Platypalpus |
| 2505AshInt6c | AshInt | May | Diptera | Hybotidae | Syndyas |
| 2505AshInt6c | AshInt | May | Diptera | Keroplatidae | Other |
| 2505AshInt6c | AshInt | May | Diptera | Limoniidae | Dicranomyia |
| 2505AshInt6c | AshInt | May | Diptera | Limoniidae | Dicranophragma |
| 2505AshInt6c | AshInt | May | Diptera | Limoniidae | Epiphragma |
| 2505AshInt6c | AshInt | May | Diptera | Limoniidae | Erioptera |
| 2505AshInt6c | AshInt | May | Diptera | Limoniidae | Metalimnobia |
| 2505AshInt6c | AshInt | May | Diptera | Limoniidae | Ormosia |
| 2505AshInt6c | AshInt | May | Diptera | Limoniidae | Other |
| 2505AshInt6c | AshInt | May | Diptera | Limoniidae | Other |
| 2505AshInt6c | AshInt | May | Diptera | Limoniidae | Pseudolimnophila |
| 2505AshInt6c | AshInt | May | Diptera | Muscidae | Mydaea |
| 2505AshInt6c | AshInt | May | Diptera | Mycetophilidae | Mycomya |
| 2505AshInt6c | AshInt | May | Diptera | Mycetophilidae | Neoempheria |

| | | | | | |
|---|---|---|---|---|---|
| 2505AshInt6c | AshInt | May | Diptera | Opomyzidae | Geomyza |
| 2505AshInt6c | AshInt | May | Diptera | Pallopteridae | Toxonevra |
| 2505AshInt6c | AshInt | May | Diptera | Phoridae | Diplonevra |
| 2505AshInt6c | AshInt | May | Diptera | Polleniidae | Pollenia |
| 2505AshInt6c | AshInt | May | Diptera | Psychodidae | Clogmia |
| 2505AshInt6c | AshInt | May | Diptera | Psychodidae | Psychoda |
| 2505AshInt6c | AshInt | May | Diptera | Sarcophagidae | Ravinia |
| 2505AshInt6c | AshInt | May | Diptera | Sciaridae | Claustropyga |
| 2505AshInt6c | AshInt | May | Diptera | Sciaridae | Corynoptera |
| 2505AshInt6c | AshInt | May | Diptera | Sciaridae | Epidapus |
| 2505AshInt6c | AshInt | May | Diptera | Sciaridae | Other |
| 2505AshInt6c | AshInt | May | Diptera | Sphaeroceridae | Leptocera |
| 2505AshInt6c | AshInt | May | Diptera | Syrphidae | Chalcosyrphus |
| 2505AshInt6c | AshInt | May | Diptera | Syrphidae | Palpada |
| 2505AshInt6c | AshInt | May | Diptera | Tabanidae | Hybomitra |
| 2505AshInt6c | AshInt | May | Diptera | Tabanidae | Other |
| 2505AshInt6c | AshInt | May | Diptera | Tachinidae | Cryptomeigenia |
| 2505AshInt6c | AshInt | May | Diptera | Tachinidae | Cyzenis |
| 2505AshInt6c | AshInt | May | Diptera | Tachinidae | Zaira |
| 2505AshInt6c | AshInt | May | Diptera | Tipulidae | Dolichopeza |
| 2505AshInt6c | AshInt | May | Diptera | Tipulidae | Tipula |
| 2505AshInt6c | AshInt | May | Ephemeroptera | Caenidae | Caenis |
| 2505AshInt6c | AshInt | May | Ephemeroptera | Ephemeridae | Hexagenia |
| 2505AshInt6c | AshInt | May | Ephemeroptera | Heptageniidae | Epeorus |
| 2505AshInt6c | AshInt | May | Hemiptera | Clastopteridae | Clastoptera |
| 2505AshInt6c | AshInt | May | Hemiptera | Lygaeidae | Kleidocerys |
| 2505AshInt6c | AshInt | May | Hymenoptera | Apidae | Triepeolus |
| 2505AshInt6c | AshInt | May | Hymenoptera | Braconidae | Bracon |
| 2505AshInt6c | AshInt | May | Hymenoptera | Braconidae | Chorebus |
| 2505AshInt6c | AshInt | May | Hymenoptera | Ichneumonidae | Megastylus |
| 2505AshInt6c | AshInt | May | Hymenoptera | Ichneumonidae | Polyblastus |

| | | | | | |
|---|---|---|---|---|---|
| 2505AshInt6c | AshInt | May | Hymenoptera | Other | Other |
| 2505AshInt6c | AshInt | May | Hymenoptera | Tenthredinidae | Dolerus |
| 2505AshInt6c | AshInt | May | Hymenoptera | Tenthredinidae | Nematus |
| 2505AshInt6c | AshInt | May | Hymenoptera | Trigonalidae | Orthogonalys |
| 2505AshInt6c | AshInt | May | Lepidoptera | Crambidae | Blepharomastix |
| 2505AshInt6c | AshInt | May | Lepidoptera | Crambidae | Parapoynx |
| 2505AshInt6c | AshInt | May | Lepidoptera | Elachistidae | Elachista |
| 2505AshInt6c | AshInt | May | Lepidoptera | Erebidae | Panopoda |
| 2505AshInt6c | AshInt | May | Lepidoptera | Gelechiidae | Chionodes |
| 2505AshInt6c | AshInt | May | Lepidoptera | Noctuidae | Condica |
| 2505AshInt6c | AshInt | May | Lepidoptera | Noctuidae | Leuconycta |
| 2505AshInt6c | AshInt | May | Lepidoptera | Noctuidae | Pyreferra |
| 2505AshInt6c | AshInt | May | Lepidoptera | Notodontidae | Heterocampa |
| 2505AshInt6c | AshInt | May | Lepidoptera | Notodontidae | Hyparpax |
| 2505AshInt6c | AshInt | May | Lepidoptera | Nymphalidae | Oeneis |
| 2505AshInt6c | AshInt | May | Lepidoptera | Pyralidae | Atheloca |
| 2505AshInt6c | AshInt | May | Lepidoptera | Pyralidae | Elasmopalpus |
| 2505AshInt6c | AshInt | May | Lepidoptera | Tischeriidae | Coptotriche |
| 2505AshInt6c | AshInt | May | Lepidoptera | Tortricidae | Acleris |
| 2505AshInt6c | AshInt | May | Lepidoptera | Tortricidae | Argyrotaenia |
| 2505AshInt6c | AshInt | May | Lepidoptera | Tortricidae | Rhyacionia |
| 2505AshInt6c | AshInt | May | Megaloptera | Corydalidae | Corydalus |
| 2505AshInt6c | AshInt | May | Neuroptera | Hemerobiidae | Hemerobius |
| 2505AshInt6c | AshInt | May | Odonata | Libellulidae | Brachymesia |
| 2505AshInt6c | AshInt | May | Orthoptera | Tettigoniidae | Conocephalus |
| 2505AshInt6c | AshInt | May | Other | Other | Other |
| 2505AshInt6c | AshInt | May | Other | Other | Other |
| 2505AshInt6c | AshInt | May | Plecoptera | Chloroperlidae | Sweltsa |
| 2505AshInt6c | AshInt | May | Psocodea | Peripsocidae | Peripsocus |
| 2505AshInt6c | AshInt | May | Sarcoptiformes | Oripodidae | Other |
| 2505AshInt6c | AshInt | May | Trichoptera | Dipseudopsidae | Phylocentropus |

| | | | | | |
|---|---|---|---|---|---|
| 2505AshInt6c | AshInt | May | Trichoptera | Hydroptilidae | Orthotrichia |
| 2505AshInt6c | AshInt | May | Trichoptera | Leptoceridae | Oecetis |
| 2505AshInt6c | AshInt | May | Trombidiformes | Arrenuridae | Arrenurus |
| 2505AshInt6c | AshInt | May | Trombidiformes | Aturidae | Aturus |
| 2505AshInt6c | AshInt | May | Trombidiformes | Axonopsidae | Brachypoda |
| 2505AshInt6c | AshInt | May | Trombidiformes | Eupodidae | Other |
| 2505AshInt6c | AshInt | May | Trombidiformes | Feltriidae | Feltria |
| 2505AshInt6c | AshInt | May | Trombidiformes | Hydryphantidae | Hydryphantes |
| 2505AshInt6c | AshInt | May | Trombidiformes | Hydryphantidae | Parathyas |
| 2505AshInt6c | AshInt | May | Trombidiformes | Lebertiidae | Lebertia |
| 2505AshInt6c | AshInt | May | Trombidiformes | Mideopsidae | Mideopsis |
| 2505AshInt6c | AshInt | May | Trombidiformes | Pionidae | Piona |
| 2505AshInt6c | AshInt | May | Trombidiformes | Pionidae | Tiphys |
| 2505AshInt6c | AshInt | May | Trombidiformes | Sperchontidae | Sperchon |
| 2505AshInt6c | AshInt | May | Trombidiformes | Tarsonemidae | Phytonemus |
| 2505AshInt6c | AshInt | May | Trombidiformes | Torrenticolidae | Testudacarus |
| 2505AshInt6c | AshInt | May | Trombidiformes | Torrenticolidae | Torrenticola |
| 2505AshInt6c | AshInt | May | Trombidiformes | Unionicolidae | Koenikea |
| 2506AshInt6a | AshInt | June | Araneae | Agelenidae | Coras |
| 2506AshInt6a | AshInt | June | Araneae | Amaurobiidae | Callobius |
| 2506AshInt6a | AshInt | June | Araneae | Anyphaenidae | Anyphaena |
| 2506AshInt6a | AshInt | June | Araneae | Cicurinidae | Cicurina |
| 2506AshInt6a | AshInt | June | Araneae | Dictynidae | Dictyna |
| 2506AshInt6a | AshInt | June | Araneae | Gnaphosidae | Gnaphosa |
| 2506AshInt6a | AshInt | June | Araneae | Linyphiidae | Grammonota |
| 2506AshInt6a | AshInt | June | Araneae | Lycosidae | Rabidosa |
| 2506AshInt6a | AshInt | June | Araneae | Lycosidae | Schizocosa |
| 2506AshInt6a | AshInt | June | Araneae | Phrurolithidae | Scotinella |
| 2506AshInt6a | AshInt | June | Araneae | Salticidae | Maevia |
| 2506AshInt6a | AshInt | June | Araneae | Salticidae | Sibianor |
| 2506AshInt6a | AshInt | June | Araneae | Theridiidae | Canalidion |

| | | | | | |
|---|---|---|---|---|---|
| 2506AshInt6a | AshInt | June | Araneae | Theridiidae | Rugathodes |
| 2506AshInt6a | AshInt | June | Araneae | Theridiidae | Spintharus |
| 2506AshInt6a | AshInt | June | Araneae | Thomisidae | Mecaphesa |
| 2506AshInt6a | AshInt | June | Blattodea | Ectobiidae | Parcoblatta |
| 2506AshInt6a | AshInt | June | Coleoptera | Buprestidae | Pachyschelus |
| 2506AshInt6a | AshInt | June | Coleoptera | Carabidae | Apenes |
| 2506AshInt6a | AshInt | June | Coleoptera | Carabidae | Harpalus |
| 2506AshInt6a | AshInt | June | Coleoptera | Carabidae | Other |
| 2506AshInt6a | AshInt | June | Coleoptera | Carabidae | Polyderis |
| 2506AshInt6a | AshInt | June | Coleoptera | Curculionidae | Conotrachelus |
| 2506AshInt6a | AshInt | June | Coleoptera | Curculionidae | Cyrtepistomus |
| 2506AshInt6a | AshInt | June | Coleoptera | Curculionidae | Listronotus |
| 2506AshInt6a | AshInt | June | Coleoptera | Dytiscidae | Acilius |
| 2506AshInt6a | AshInt | June | Coleoptera | Lampyridae | Photinus |
| 2506AshInt6a | AshInt | June | Coleoptera | Scarabaeidae | Dialytes |
| 2506AshInt6a | AshInt | June | Coleoptera | Scarabaeidae | Melanocanthon |
| 2506AshInt6a | AshInt | June | Coleoptera | Scirtidae | Contacyphon |
| 2506AshInt6a | AshInt | June | Diptera | Agromyzidae | Calycomyza |
| 2506AshInt6a | AshInt | June | Diptera | Agromyzidae | Cerodontha |
| 2506AshInt6a | AshInt | June | Diptera | Agromyzidae | Japanagromyza |
| 2506AshInt6a | AshInt | June | Diptera | Agromyzidae | Liriomyza |
| 2506AshInt6a | AshInt | June | Diptera | Agromyzidae | Nemorimyza |
| 2506AshInt6a | AshInt | June | Diptera | Agromyzidae | Phytomyza |
| 2506AshInt6a | AshInt | June | Diptera | Anisopodidae | Mycetobia |
| 2506AshInt6a | AshInt | June | Diptera | Calliphoridae | Lucilia |
| 2506AshInt6a | AshInt | June | Diptera | Cecidomyiidae | Other |
| 2506AshInt6a | AshInt | June | Diptera | Chaoboridae | Chaoborus |
| 2506AshInt6a | AshInt | June | Diptera | Chaoboridae | Other |
| 2506AshInt6a | AshInt | June | Diptera | Chironomidae | Ablabesmyia |
| 2506AshInt6a | AshInt | June | Diptera | Chironomidae | Chironomus |
| 2506AshInt6a | AshInt | June | Diptera | Chironomidae | Dicrotendipes |

| | | | | | |
|---|---|---|---|---|---|
| 2506AshInt6a | AshInt | June | Diptera | Chironomidae | Endochironomus |
| 2506AshInt6a | AshInt | June | Diptera | Chironomidae | Glyptotendipes |
| 2506AshInt6a | AshInt | June | Diptera | Chironomidae | Microtendipes |
| 2506AshInt6a | AshInt | June | Diptera | Chironomidae | Orthocladius |
| 2506AshInt6a | AshInt | June | Diptera | Chironomidae | Other |
| 2506AshInt6a | AshInt | June | Diptera | Chironomidae | Other |
| 2506AshInt6a | AshInt | June | Diptera | Chironomidae | Other |
| 2506AshInt6a | AshInt | June | Diptera | Chironomidae | Other |
| 2506AshInt6a | AshInt | June | Diptera | Chironomidae | Parachironomus |
| 2506AshInt6a | AshInt | June | Diptera | Chironomidae | Polypedilum |
| 2506AshInt6a | AshInt | June | Diptera | Chironomidae | Pseudochironomus |
| 2506AshInt6a | AshInt | June | Diptera | Chironomidae | Tanytarsus |
| 2506AshInt6a | AshInt | June | Diptera | Chironomidae | Xenochironomus |
| 2506AshInt6a | AshInt | June | Diptera | Chloropidae | Apotropina |
| 2506AshInt6a | AshInt | June | Diptera | Chloropidae | Elachiptera |
| 2506AshInt6a | AshInt | June | Diptera | Chloropidae | Liohippelates |
| 2506AshInt6a | AshInt | June | Diptera | Chloropidae | Oscinella |
| 2506AshInt6a | AshInt | June | Diptera | Culicidae | Coquillettidia |
| 2506AshInt6a | AshInt | June | Diptera | Culicidae | Culex |
| 2506AshInt6a | AshInt | June | Diptera | Culicidae | Uranotaenia |
| 2506AshInt6a | AshInt | June | Diptera | Drosophilidae | Drosophila |
| 2506AshInt6a | AshInt | June | Diptera | Empididae | Other |
| 2506AshInt6a | AshInt | June | Diptera | Hybotidae | Platypalpus |
| 2506AshInt6a | AshInt | June | Diptera | Hybotidae | Syndyas |
| 2506AshInt6a | AshInt | June | Diptera | Keroplatidae | Other |
| 2506AshInt6a | AshInt | June | Diptera | Limoniidae | Dicranomyia |
| 2506AshInt6a | AshInt | June | Diptera | Limoniidae | Dicranophragma |
| 2506AshInt6a | AshInt | June | Diptera | Limoniidae | Epiphragma |
| 2506AshInt6a | AshInt | June | Diptera | Limoniidae | Erioptera |
| 2506AshInt6a | AshInt | June | Diptera | Limoniidae | Metalimnobia |
| 2506AshInt6a | AshInt | June | Diptera | Limoniidae | Ormosia |

| | | | | | |
|---|---|---|---|---|---|
| 2506AshInt6a | AshInt | June | Diptera | Limoniidae | Other |
| 2506AshInt6a | AshInt | June | Diptera | Limoniidae | Other |
| 2506AshInt6a | AshInt | June | Diptera | Limoniidae | Pseudolimnophila |
| 2506AshInt6a | AshInt | June | Diptera | Muscidae | Mydaea |
| 2506AshInt6a | AshInt | June | Diptera | Mycetophilidae | Mycomya |
| 2506AshInt6a | AshInt | June | Diptera | Mycetophilidae | Neoempheria |
| 2506AshInt6a | AshInt | June | Diptera | Opomyzidae | Geomyza |
| 2506AshInt6a | AshInt | June | Diptera | Pallopteridae | Toxonevra |
| 2506AshInt6a | AshInt | June | Diptera | Phoridae | Diplonevra |
| 2506AshInt6a | AshInt | June | Diptera | Polleniidae | Pollenia |
| 2506AshInt6a | AshInt | June | Diptera | Psychodidae | Clogmia |
| 2506AshInt6a | AshInt | June | Diptera | Psychodidae | Psychoda |
| 2506AshInt6a | AshInt | June | Diptera | Sarcophagidae | Ravinia |
| 2506AshInt6a | AshInt | June | Diptera | Sciaridae | Claustropyga |
| 2506AshInt6a | AshInt | June | Diptera | Sciaridae | Corynoptera |
| 2506AshInt6a | AshInt | June | Diptera | Sciaridae | Epidapus |
| 2506AshInt6a | AshInt | June | Diptera | Sciaridae | Other |
| 2506AshInt6a | AshInt | June | Diptera | Sphaeroceridae | Leptocera |
| 2506AshInt6a | AshInt | June | Diptera | Syrphidae | Chalcosyrphus |
| 2506AshInt6a | AshInt | June | Diptera | Syrphidae | Palpada |
| 2506AshInt6a | AshInt | June | Diptera | Tabanidae | Hybomitra |
| 2506AshInt6a | AshInt | June | Diptera | Tabanidae | Other |
| 2506AshInt6a | AshInt | June | Diptera | Tachinidae | Cryptomeigenia |
| 2506AshInt6a | AshInt | June | Diptera | Tachinidae | Cyzenis |
| 2506AshInt6a | AshInt | June | Diptera | Tachinidae | Zaira |
| 2506AshInt6a | AshInt | June | Diptera | Tipulidae | Dolichopeza |
| 2506AshInt6a | AshInt | June | Diptera | Tipulidae | Tipula |
| 2506AshInt6a | AshInt | June | Ephemeroptera | Caenidae | Caenis |
| 2506AshInt6a | AshInt | June | Ephemeroptera | Ephemeridae | Hexagenia |
| 2506AshInt6a | AshInt | June | Ephemeroptera | Heptageniidae | Epeorus |
| 2506AshInt6a | AshInt | June | Hemiptera | Clastopteridae | Clastoptera |

| | | | | | |
|---|---|---|---|---|---|
| 2506AshInt6a | AshInt | June | Hemiptera | Lygaeidae | Kleidocerys |
| 2506AshInt6a | AshInt | June | Hymenoptera | Apidae | Triepeolus |
| 2506AshInt6a | AshInt | June | Hymenoptera | Braconidae | Bracon |
| 2506AshInt6a | AshInt | June | Hymenoptera | Braconidae | Chorebus |
| 2506AshInt6a | AshInt | June | Hymenoptera | Ichneumonidae | Megastylus |
| 2506AshInt6a | AshInt | June | Hymenoptera | Ichneumonidae | Polyblastus |
| 2506AshInt6a | AshInt | June | Hymenoptera | Other | Other |
| 2506AshInt6a | AshInt | June | Hymenoptera | Tenthredinidae | Dolerus |
| 2506AshInt6a | AshInt | June | Hymenoptera | Tenthredinidae | Nematus |
| 2506AshInt6a | AshInt | June | Hymenoptera | Trigonalidae | Orthogonalys |
| 2506AshInt6a | AshInt | June | Lepidoptera | Crambidae | Blepharomastix |
| 2506AshInt6a | AshInt | June | Lepidoptera | Crambidae | Parapoynx |
| 2506AshInt6a | AshInt | June | Lepidoptera | Elachistidae | Elachista |
| 2506AshInt6a | AshInt | June | Lepidoptera | Erebidae | Panopoda |
| 2506AshInt6a | AshInt | June | Lepidoptera | Gelechiidae | Chionodes |
| 2506AshInt6a | AshInt | June | Lepidoptera | Noctuidae | Condica |
| 2506AshInt6a | AshInt | June | Lepidoptera | Noctuidae | Leuconycta |
| 2506AshInt6a | AshInt | June | Lepidoptera | Noctuidae | Pyreferra |
| 2506AshInt6a | AshInt | June | Lepidoptera | Notodontidae | Heterocampa |
| 2506AshInt6a | AshInt | June | Lepidoptera | Notodontidae | Hyparpax |
| 2506AshInt6a | AshInt | June | Lepidoptera | Nymphalidae | Oeneis |
| 2506AshInt6a | AshInt | June | Lepidoptera | Pyralidae | Atheloca |
| 2506AshInt6a | AshInt | June | Lepidoptera | Pyralidae | Elasmopalpus |
| 2506AshInt6a | AshInt | June | Lepidoptera | Tischeriidae | Coptotriche |
| 2506AshInt6a | AshInt | June | Lepidoptera | Tortricidae | Acleris |
| 2506AshInt6a | AshInt | June | Lepidoptera | Tortricidae | Argyrotaenia |
| 2506AshInt6a | AshInt | June | Lepidoptera | Tortricidae | Rhyacionia |
| 2506AshInt6a | AshInt | June | Megaloptera | Corydalidae | Corydalus |
| 2506AshInt6a | AshInt | June | Neuroptera | Hemerobiidae | Hemerobius |
| 2506AshInt6a | AshInt | June | Odonata | Libellulidae | Brachymesia |
| 2506AshInt6a | AshInt | June | Orthoptera | Tettigoniidae | Conocephalus |

| | | | | | |
|---|---|---|---|---|---|
| 2506AshInt6a | AshInt | June | Other | Other | Other |
| 2506AshInt6a | AshInt | June | Other | Other | Other |
| 2506AshInt6a | AshInt | June | Plecoptera | Chloroperlidae | Sweltsa |
| 2506AshInt6a | AshInt | June | Psocodea | Peripsocidae | Peripsocus |
| 2506AshInt6a | AshInt | June | Sarcoptiformes | Oripodidae | Other |
| 2506AshInt6a | AshInt | June | Trichoptera | Dipseudopsidae | Phylocentropus |
| 2506AshInt6a | AshInt | June | Trichoptera | Hydroptilidae | Orthotrichia |
| 2506AshInt6a | AshInt | June | Trichoptera | Leptoceridae | Oecetis |
| 2506AshInt6a | AshInt | June | Trombidiformes | Arrenuridae | Arrenurus |
| 2506AshInt6a | AshInt | June | Trombidiformes | Aturidae | Aturus |
| 2506AshInt6a | AshInt | June | Trombidiformes | Axonopsidae | Brachypoda |
| 2506AshInt6a | AshInt | June | Trombidiformes | Eupodidae | Other |
| 2506AshInt6a | AshInt | June | Trombidiformes | Feltriidae | Feltria |
| 2506AshInt6a | AshInt | June | Trombidiformes | Hydryphantidae | Hydryphantes |
| 2506AshInt6a | AshInt | June | Trombidiformes | Hydryphantidae | Parathyas |
| 2506AshInt6a | AshInt | June | Trombidiformes | Lebertiidae | Lebertia |
| 2506AshInt6a | AshInt | June | Trombidiformes | Mideopsidae | Mideopsis |
| 2506AshInt6a | AshInt | June | Trombidiformes | Pionidae | Piona |
| 2506AshInt6a | AshInt | June | Trombidiformes | Pionidae | Tiphys |
| 2506AshInt6a | AshInt | June | Trombidiformes | Sperchontidae | Sperchon |
| 2506AshInt6a | AshInt | June | Trombidiformes | Tarsonemidae | Phytonemus |
| 2506AshInt6a | AshInt | June | Trombidiformes | Torrenticolidae | Testudacarus |
| 2506AshInt6a | AshInt | June | Trombidiformes | Torrenticolidae | Torrenticola |
| 2506AshInt6a | AshInt | June | Trombidiformes | Unionicolidae | Koenikea |
| 2506AshInt6b | AshInt | June | Araneae | Agelenidae | Coras |
| 2506AshInt6b | AshInt | June | Araneae | Amaurobiidae | Callobius |
| 2506AshInt6b | AshInt | June | Araneae | Anyphaenidae | Anyphaena |
| 2506AshInt6b | AshInt | June | Araneae | Cicurinidae | Cicurina |
| 2506AshInt6b | AshInt | June | Araneae | Dictynidae | Dictyna |
| 2506AshInt6b | AshInt | June | Araneae | Gnaphosidae | Gnaphosa |
| 2506AshInt6b | AshInt | June | Araneae | Linyphiidae | Grammonota |

| | | | | | |
|---|---|---|---|---|---|
| 2506AshInt6b | AshInt | June | Araneae | Lycosidae | Rabidosa |
| 2506AshInt6b | AshInt | June | Araneae | Lycosidae | Schizocosa |
| 2506AshInt6b | AshInt | June | Araneae | Phrurolithidae | Scotinella |
| 2506AshInt6b | AshInt | June | Araneae | Salticidae | Maevia |
| 2506AshInt6b | AshInt | June | Araneae | Salticidae | Sibianor |
| 2506AshInt6b | AshInt | June | Araneae | Theridiidae | Canalidion |
| 2506AshInt6b | AshInt | June | Araneae | Theridiidae | Rugathodes |
| 2506AshInt6b | AshInt | June | Araneae | Theridiidae | Spintharus |
| 2506AshInt6b | AshInt | June | Araneae | Thomisidae | Mecaphesa |
| 2506AshInt6b | AshInt | June | Blattodea | Ectobiidae | Parcoblatta |
| 2506AshInt6b | AshInt | June | Coleoptera | Buprestidae | Pachyschelus |
| 2506AshInt6b | AshInt | June | Coleoptera | Carabidae | Apenes |
| 2506AshInt6b | AshInt | June | Coleoptera | Carabidae | Harpalus |
| 2506AshInt6b | AshInt | June | Coleoptera | Carabidae | Other |
| 2506AshInt6b | AshInt | June | Coleoptera | Carabidae | Polyderis |
| 2506AshInt6b | AshInt | June | Coleoptera | Curculionidae | Conotrachelus |
| 2506AshInt6b | AshInt | June | Coleoptera | Curculionidae | Cyrtepistomus |
| 2506AshInt6b | AshInt | June | Coleoptera | Curculionidae | Listronotus |
| 2506AshInt6b | AshInt | June | Coleoptera | Dytiscidae | Acilius |
| 2506AshInt6b | AshInt | June | Coleoptera | Lampyridae | Photinus |
| 2506AshInt6b | AshInt | June | Coleoptera | Scarabaeidae | Dialytes |
| 2506AshInt6b | AshInt | June | Coleoptera | Scarabaeidae | Melanocanthon |
| 2506AshInt6b | AshInt | June | Coleoptera | Scirtidae | Contacyphon |
| 2506AshInt6b | AshInt | June | Diptera | Agromyzidae | Calycomyza |
| 2506AshInt6b | AshInt | June | Diptera | Agromyzidae | Cerodontha |
| 2506AshInt6b | AshInt | June | Diptera | Agromyzidae | Japanagromyza |
| 2506AshInt6b | AshInt | June | Diptera | Agromyzidae | Liriomyza |
| 2506AshInt6b | AshInt | June | Diptera | Agromyzidae | Nemorimyza |
| 2506AshInt6b | AshInt | June | Diptera | Agromyzidae | Phytomyza |
| 2506AshInt6b | AshInt | June | Diptera | Anisopodidae | Mycetobia |
| 2506AshInt6b | AshInt | June | Diptera | Calliphoridae | Lucilia |

| | | | | | |
|---|---|---|---|---|---|
| 2506AshInt6b | AshInt | June | Diptera | Cecidomyiidae | Other |
| 2506AshInt6b | AshInt | June | Diptera | Chaoboridae | Chaoborus |
| 2506AshInt6b | AshInt | June | Diptera | Chaoboridae | Other |
| 2506AshInt6b | AshInt | June | Diptera | Chironomidae | Ablabesmyia |
| 2506AshInt6b | AshInt | June | Diptera | Chironomidae | Chironomus |
| 2506AshInt6b | AshInt | June | Diptera | Chironomidae | Dicrotendipes |
| 2506AshInt6b | AshInt | June | Diptera | Chironomidae | Endochironomus |
| 2506AshInt6b | AshInt | June | Diptera | Chironomidae | Glyptotendipes |
| 2506AshInt6b | AshInt | June | Diptera | Chironomidae | Microtendipes |
| 2506AshInt6b | AshInt | June | Diptera | Chironomidae | Orthocladius |
| 2506AshInt6b | AshInt | June | Diptera | Chironomidae | Other |
| 2506AshInt6b | AshInt | June | Diptera | Chironomidae | Other |
| 2506AshInt6b | AshInt | June | Diptera | Chironomidae | Other |
| 2506AshInt6b | AshInt | June | Diptera | Chironomidae | Other |
| 2506AshInt6b | AshInt | June | Diptera | Chironomidae | Parachironomus |
| 2506AshInt6b | AshInt | June | Diptera | Chironomidae | Polypedilum |
| 2506AshInt6b | AshInt | June | Diptera | Chironomidae | Pseudochironomus |
| 2506AshInt6b | AshInt | June | Diptera | Chironomidae | Tanytarsus |
| 2506AshInt6b | AshInt | June | Diptera | Chironomidae | Xenochironomus |
| 2506AshInt6b | AshInt | June | Diptera | Chloropidae | Apotropina |
| 2506AshInt6b | AshInt | June | Diptera | Chloropidae | Elachiptera |
| 2506AshInt6b | AshInt | June | Diptera | Chloropidae | Liohippelates |
| 2506AshInt6b | AshInt | June | Diptera | Chloropidae | Oscinella |
| 2506AshInt6b | AshInt | June | Diptera | Culicidae | Coquillettidia |
| 2506AshInt6b | AshInt | June | Diptera | Culicidae | Culex |
| 2506AshInt6b | AshInt | June | Diptera | Culicidae | Uranotaenia |
| 2506AshInt6b | AshInt | June | Diptera | Drosophilidae | Drosophila |
| 2506AshInt6b | AshInt | June | Diptera | Empididae | Other |
| 2506AshInt6b | AshInt | June | Diptera | Hybotidae | Platypalpus |
| 2506AshInt6b | AshInt | June | Diptera | Hybotidae | Syndyas |
| 2506AshInt6b | AshInt | June | Diptera | Keroplatidae | Other |

| | | | | | |
|---|---|---|---|---|---|
| 2506AshInt6b | AshInt | June | Diptera | Limoniidae | Dicranomyia |
| 2506AshInt6b | AshInt | June | Diptera | Limoniidae | Dicranophragma |
| 2506AshInt6b | AshInt | June | Diptera | Limoniidae | Epiphragma |
| 2506AshInt6b | AshInt | June | Diptera | Limoniidae | Erioptera |
| 2506AshInt6b | AshInt | June | Diptera | Limoniidae | Metalimnobia |
| 2506AshInt6b | AshInt | June | Diptera | Limoniidae | Ormosia |
| 2506AshInt6b | AshInt | June | Diptera | Limoniidae | Other |
| 2506AshInt6b | AshInt | June | Diptera | Limoniidae | Other |
| 2506AshInt6b | AshInt | June | Diptera | Limoniidae | Pseudolimnophila |
| 2506AshInt6b | AshInt | June | Diptera | Muscidae | Mydaea |
| 2506AshInt6b | AshInt | June | Diptera | Mycetophilidae | Mycomya |
| 2506AshInt6b | AshInt | June | Diptera | Mycetophilidae | Neoempheria |
| 2506AshInt6b | AshInt | June | Diptera | Opomyzidae | Geomyza |
| 2506AshInt6b | AshInt | June | Diptera | Pallopteridae | Toxonevra |
| 2506AshInt6b | AshInt | June | Diptera | Phoridae | Diplonevra |
| 2506AshInt6b | AshInt | June | Diptera | Polleniidae | Pollenia |
| 2506AshInt6b | AshInt | June | Diptera | Psychodidae | Clogmia |
| 2506AshInt6b | AshInt | June | Diptera | Psychodidae | Psychoda |
| 2506AshInt6b | AshInt | June | Diptera | Sarcophagidae | Ravinia |
| 2506AshInt6b | AshInt | June | Diptera | Sciaridae | Claustropyga |
| 2506AshInt6b | AshInt | June | Diptera | Sciaridae | Corynoptera |
| 2506AshInt6b | AshInt | June | Diptera | Sciaridae | Epidapus |
| 2506AshInt6b | AshInt | June | Diptera | Sciaridae | Other |
| 2506AshInt6b | AshInt | June | Diptera | Sphaeroceridae | Leptocera |
| 2506AshInt6b | AshInt | June | Diptera | Syrphidae | Chalcosyrphus |
| 2506AshInt6b | AshInt | June | Diptera | Syrphidae | Palpada |
| 2506AshInt6b | AshInt | June | Diptera | Tabanidae | Hybomitra |
| 2506AshInt6b | AshInt | June | Diptera | Tabanidae | Other |
| 2506AshInt6b | AshInt | June | Diptera | Tachinidae | Cryptomeigenia |
| 2506AshInt6b | AshInt | June | Diptera | Tachinidae | Cyzenis |
| 2506AshInt6b | AshInt | June | Diptera | Tachinidae | Zaira |

| | | | | | |
|---|---|---|---|---|---|
| 2506AshInt6b | AshInt | June | Diptera | Tipulidae | Dolichopeza |
| 2506AshInt6b | AshInt | June | Diptera | Tipulidae | Tipula |
| 2506AshInt6b | AshInt | June | Ephemeroptera | Caenidae | Caenis |
| 2506AshInt6b | AshInt | June | Ephemeroptera | Ephemeridae | Hexagenia |
| 2506AshInt6b | AshInt | June | Ephemeroptera | Heptageniidae | Epeorus |
| 2506AshInt6b | AshInt | June | Hemiptera | Clastopteridae | Clastoptera |
| 2506AshInt6b | AshInt | June | Hemiptera | Lygaeidae | Kleidocerys |
| 2506AshInt6b | AshInt | June | Hymenoptera | Apidae | Triepeolus |
| 2506AshInt6b | AshInt | June | Hymenoptera | Braconidae | Bracon |
| 2506AshInt6b | AshInt | June | Hymenoptera | Braconidae | Chorebus |
| 2506AshInt6b | AshInt | June | Hymenoptera | Ichneumonidae | Megastylus |
| 2506AshInt6b | AshInt | June | Hymenoptera | Ichneumonidae | Polyblastus |
| 2506AshInt6b | AshInt | June | Hymenoptera | Other | Other |
| 2506AshInt6b | AshInt | June | Hymenoptera | Tenthredinidae | Dolerus |
| 2506AshInt6b | AshInt | June | Hymenoptera | Tenthredinidae | Nematus |
| 2506AshInt6b | AshInt | June | Hymenoptera | Trigonalidae | Orthogonalys |
| 2506AshInt6b | AshInt | June | Lepidoptera | Crambidae | Blepharomastix |
| 2506AshInt6b | AshInt | June | Lepidoptera | Crambidae | Parapoynx |
| 2506AshInt6b | AshInt | June | Lepidoptera | Elachistidae | Elachista |
| 2506AshInt6b | AshInt | June | Lepidoptera | Erebidae | Panopoda |
| 2506AshInt6b | AshInt | June | Lepidoptera | Gelechiidae | Chionodes |
| 2506AshInt6b | AshInt | June | Lepidoptera | Noctuidae | Condica |
| 2506AshInt6b | AshInt | June | Lepidoptera | Noctuidae | Leuconycta |
| 2506AshInt6b | AshInt | June | Lepidoptera | Noctuidae | Pyreferra |
| 2506AshInt6b | AshInt | June | Lepidoptera | Notodontidae | Heterocampa |
| 2506AshInt6b | AshInt | June | Lepidoptera | Notodontidae | Hyparpax |
| 2506AshInt6b | AshInt | June | Lepidoptera | Nymphalidae | Oeneis |
| 2506AshInt6b | AshInt | June | Lepidoptera | Pyralidae | Atheloca |
| 2506AshInt6b | AshInt | June | Lepidoptera | Pyralidae | Elasmopalpus |
| 2506AshInt6b | AshInt | June | Lepidoptera | Tischeriidae | Coptotriche |
| 2506AshInt6b | AshInt | June | Lepidoptera | Tortricidae | Acleris |

| | | | | | |
|---|---|---|---|---|---|
| 2506AshInt6b | AshInt | June | Lepidoptera | Tortricidae | Argyrotaenia |
| 2506AshInt6b | AshInt | June | Lepidoptera | Tortricidae | Rhyacionia |
| 2506AshInt6b | AshInt | June | Megaloptera | Corydalidae | Corydalus |
| 2506AshInt6b | AshInt | June | Neuroptera | Hemerobiidae | Hemerobius |
| 2506AshInt6b | AshInt | June | Odonata | Libellulidae | Brachymesia |
| 2506AshInt6b | AshInt | June | Orthoptera | Tettigoniidae | Conocephalus |
| 2506AshInt6b | AshInt | June | Other | Other | Other |
| 2506AshInt6b | AshInt | June | Other | Other | Other |
| 2506AshInt6b | AshInt | June | Plecoptera | Chloroperlidae | Sweltsa |
| 2506AshInt6b | AshInt | June | Psocodea | Peripsocidae | Peripsocus |
| 2506AshInt6b | AshInt | June | Sarcoptiformes | Oripodidae | Other |
| 2506AshInt6b | AshInt | June | Trichoptera | Dipseudopsidae | Phylocentropus |
| 2506AshInt6b | AshInt | June | Trichoptera | Hydroptilidae | Orthotrichia |
| 2506AshInt6b | AshInt | June | Trichoptera | Leptoceridae | Oecetis |
| 2506AshInt6b | AshInt | June | Trombidiformes | Arrenuridae | Arrenurus |
| 2506AshInt6b | AshInt | June | Trombidiformes | Aturidae | Aturus |
| 2506AshInt6b | AshInt | June | Trombidiformes | Axonopsidae | Brachypoda |
| 2506AshInt6b | AshInt | June | Trombidiformes | Eupodidae | Other |
| 2506AshInt6b | AshInt | June | Trombidiformes | Feltriidae | Feltria |
| 2506AshInt6b | AshInt | June | Trombidiformes | Hydryphantidae | Hydryphantes |
| 2506AshInt6b | AshInt | June | Trombidiformes | Hydryphantidae | Parathyas |
| 2506AshInt6b | AshInt | June | Trombidiformes | Lebertiidae | Lebertia |
| 2506AshInt6b | AshInt | June | Trombidiformes | Mideopsidae | Mideopsis |
| 2506AshInt6b | AshInt | June | Trombidiformes | Pionidae | Piona |
| 2506AshInt6b | AshInt | June | Trombidiformes | Pionidae | Tiphys |
| 2506AshInt6b | AshInt | June | Trombidiformes | Sperchontidae | Sperchon |
| 2506AshInt6b | AshInt | June | Trombidiformes | Tarsonemidae | Phytonemus |
| 2506AshInt6b | AshInt | June | Trombidiformes | Torrenticolidae | Testudacarus |
| 2506AshInt6b | AshInt | June | Trombidiformes | Torrenticolidae | Torrenticola |
| 2506AshInt6b | AshInt | June | Trombidiformes | Unionicolidae | Koenikea |
| 2506AshInt6c | AshInt | June | Araneae | Agelenidae | Coras |

| | | | | | |
|---|---|---|---|---|---|
| 2506AshInt6c | AshInt | June | Araneae | Amaurobiidae | Callobius |
| 2506AshInt6c | AshInt | June | Araneae | Anyphaenidae | Anyphaena |
| 2506AshInt6c | AshInt | June | Araneae | Cicurinidae | Cicurina |
| 2506AshInt6c | AshInt | June | Araneae | Dictynidae | Dictyna |
| 2506AshInt6c | AshInt | June | Araneae | Gnaphosidae | Gnaphosa |
| 2506AshInt6c | AshInt | June | Araneae | Linyphiidae | Grammonota |
| 2506AshInt6c | AshInt | June | Araneae | Lycosidae | Rabidosa |
| 2506AshInt6c | AshInt | June | Araneae | Lycosidae | Schizocosa |
| 2506AshInt6c | AshInt | June | Araneae | Phrurolithidae | Scotinella |
| 2506AshInt6c | AshInt | June | Araneae | Salticidae | Maevia |
| 2506AshInt6c | AshInt | June | Araneae | Salticidae | Sibianor |
| 2506AshInt6c | AshInt | June | Araneae | Theridiidae | Canalidion |
| 2506AshInt6c | AshInt | June | Araneae | Theridiidae | Rugathodes |
| 2506AshInt6c | AshInt | June | Araneae | Theridiidae | Spintharus |
| 2506AshInt6c | AshInt | June | Araneae | Thomisidae | Mecaphesa |
| 2506AshInt6c | AshInt | June | Blattodea | Ectobiidae | Parcoblatta |
| 2506AshInt6c | AshInt | June | Coleoptera | Buprestidae | Pachyschelus |
| 2506AshInt6c | AshInt | June | Coleoptera | Carabidae | Apenes |
| 2506AshInt6c | AshInt | June | Coleoptera | Carabidae | Harpalus |
| 2506AshInt6c | AshInt | June | Coleoptera | Carabidae | Other |
| 2506AshInt6c | AshInt | June | Coleoptera | Carabidae | Polyderis |
| 2506AshInt6c | AshInt | June | Coleoptera | Curculionidae | Conotrachelus |
| 2506AshInt6c | AshInt | June | Coleoptera | Curculionidae | Cyrtepistomus |
| 2506AshInt6c | AshInt | June | Coleoptera | Curculionidae | Listronotus |
| 2506AshInt6c | AshInt | June | Coleoptera | Dytiscidae | Acilius |
| 2506AshInt6c | AshInt | June | Coleoptera | Lampyridae | Photinus |
| 2506AshInt6c | AshInt | June | Coleoptera | Scarabaeidae | Dialytes |
| 2506AshInt6c | AshInt | June | Coleoptera | Scarabaeidae | Melanocanthon |
| 2506AshInt6c | AshInt | June | Coleoptera | Scirtidae | Contacyphon |
| 2506AshInt6c | AshInt | June | Diptera | Agromyzidae | Calycomyza |
| 2506AshInt6c | AshInt | June | Diptera | Agromyzidae | Cerodontha |

| | | | | | |
|---|---|---|---|---|---|
| 2506AshInt6c | AshInt | June | Diptera | Agromyzidae | Japanagromyza |
| 2506AshInt6c | AshInt | June | Diptera | Agromyzidae | Liriomyza |
| 2506AshInt6c | AshInt | June | Diptera | Agromyzidae | Nemorimyza |
| 2506AshInt6c | AshInt | June | Diptera | Agromyzidae | Phytomyza |
| 2506AshInt6c | AshInt | June | Diptera | Anisopodidae | Mycetobia |
| 2506AshInt6c | AshInt | June | Diptera | Calliphoridae | Lucilia |
| 2506AshInt6c | AshInt | June | Diptera | Cecidomyiidae | Other |
| 2506AshInt6c | AshInt | June | Diptera | Chaoboridae | Chaoborus |
| 2506AshInt6c | AshInt | June | Diptera | Chaoboridae | Other |
| 2506AshInt6c | AshInt | June | Diptera | Chironomidae | Ablabesmyia |
| 2506AshInt6c | AshInt | June | Diptera | Chironomidae | Chironomus |
| 2506AshInt6c | AshInt | June | Diptera | Chironomidae | Dicrotendipes |
| 2506AshInt6c | AshInt | June | Diptera | Chironomidae | Endochironomus |
| 2506AshInt6c | AshInt | June | Diptera | Chironomidae | Glyptotendipes |
| 2506AshInt6c | AshInt | June | Diptera | Chironomidae | Microtendipes |
| 2506AshInt6c | AshInt | June | Diptera | Chironomidae | Orthocladius |
| 2506AshInt6c | AshInt | June | Diptera | Chironomidae | Other |
| 2506AshInt6c | AshInt | June | Diptera | Chironomidae | Other |
| 2506AshInt6c | AshInt | June | Diptera | Chironomidae | Other |
| 2506AshInt6c | AshInt | June | Diptera | Chironomidae | Other |
| 2506AshInt6c | AshInt | June | Diptera | Chironomidae | Parachironomus |
| 2506AshInt6c | AshInt | June | Diptera | Chironomidae | Polypedilum |
| 2506AshInt6c | AshInt | June | Diptera | Chironomidae | Pseudochironomus |
| 2506AshInt6c | AshInt | June | Diptera | Chironomidae | Tanytarsus |
| 2506AshInt6c | AshInt | June | Diptera | Chironomidae | Xenochironomus |
| 2506AshInt6c | AshInt | June | Diptera | Chloropidae | Apotropina |
| 2506AshInt6c | AshInt | June | Diptera | Chloropidae | Elachiptera |
| 2506AshInt6c | AshInt | June | Diptera | Chloropidae | Liohippelates |
| 2506AshInt6c | AshInt | June | Diptera | Chloropidae | Oscinella |
| 2506AshInt6c | AshInt | June | Diptera | Culicidae | Coquillettidia |
| 2506AshInt6c | AshInt | June | Diptera | Culicidae | Culex |

| | | | | | |
|---|---|---|---|---|---|
| 2506AshInt6c | AshInt | June | Diptera | Culicidae | Uranotaenia |
| 2506AshInt6c | AshInt | June | Diptera | Drosophilidae | Drosophila |
| 2506AshInt6c | AshInt | June | Diptera | Empididae | Other |
| 2506AshInt6c | AshInt | June | Diptera | Hybotidae | Platypalpus |
| 2506AshInt6c | AshInt | June | Diptera | Hybotidae | Syndyas |
| 2506AshInt6c | AshInt | June | Diptera | Keroplatidae | Other |
| 2506AshInt6c | AshInt | June | Diptera | Limoniidae | Dicranomyia |
| 2506AshInt6c | AshInt | June | Diptera | Limoniidae | Dicranophragma |
| 2506AshInt6c | AshInt | June | Diptera | Limoniidae | Epiphragma |
| 2506AshInt6c | AshInt | June | Diptera | Limoniidae | Erioptera |
| 2506AshInt6c | AshInt | June | Diptera | Limoniidae | Metalimnobia |
| 2506AshInt6c | AshInt | June | Diptera | Limoniidae | Ormosia |
| 2506AshInt6c | AshInt | June | Diptera | Limoniidae | Other |
| 2506AshInt6c | AshInt | June | Diptera | Limoniidae | Other |
| 2506AshInt6c | AshInt | June | Diptera | Limoniidae | Pseudolimnophila |
| 2506AshInt6c | AshInt | June | Diptera | Muscidae | Mydaea |
| 2506AshInt6c | AshInt | June | Diptera | Mycetophilidae | Mycomya |
| 2506AshInt6c | AshInt | June | Diptera | Mycetophilidae | Neoempheria |
| 2506AshInt6c | AshInt | June | Diptera | Opomyzidae | Geomyza |
| 2506AshInt6c | AshInt | June | Diptera | Pallopteridae | Toxonevra |
| 2506AshInt6c | AshInt | June | Diptera | Phoridae | Diplonevra |
| 2506AshInt6c | AshInt | June | Diptera | Polleniidae | Pollenia |
| 2506AshInt6c | AshInt | June | Diptera | Psychodidae | Clogmia |
| 2506AshInt6c | AshInt | June | Diptera | Psychodidae | Psychoda |
| 2506AshInt6c | AshInt | June | Diptera | Sarcophagidae | Ravinia |
| 2506AshInt6c | AshInt | June | Diptera | Sciaridae | Claustropyga |
| 2506AshInt6c | AshInt | June | Diptera | Sciaridae | Corynoptera |
| 2506AshInt6c | AshInt | June | Diptera | Sciaridae | Epidapus |
| 2506AshInt6c | AshInt | June | Diptera | Sciaridae | Other |
| 2506AshInt6c | AshInt | June | Diptera | Sphaeroceridae | Leptocera |
| 2506AshInt6c | AshInt | June | Diptera | Syrphidae | Chalcosyrphus |

| | | | | | |
|---|---|---|---|---|---|
| 2506AshInt6c | AshInt | June | Diptera | Syrphidae | Palpada |
| 2506AshInt6c | AshInt | June | Diptera | Tabanidae | Hybomitra |
| 2506AshInt6c | AshInt | June | Diptera | Tabanidae | Other |
| 2506AshInt6c | AshInt | June | Diptera | Tachinidae | Cryptomeigenia |
| 2506AshInt6c | AshInt | June | Diptera | Tachinidae | Cyzenis |
| 2506AshInt6c | AshInt | June | Diptera | Tachinidae | Zaira |
| 2506AshInt6c | AshInt | June | Diptera | Tipulidae | Dolichopeza |
| 2506AshInt6c | AshInt | June | Diptera | Tipulidae | Tipula |
| 2506AshInt6c | AshInt | June | Ephemeroptera | Caenidae | Caenis |
| 2506AshInt6c | AshInt | June | Ephemeroptera | Ephemeridae | Hexagenia |
| 2506AshInt6c | AshInt | June | Ephemeroptera | Heptageniidae | Epeorus |
| 2506AshInt6c | AshInt | June | Hemiptera | Clastopteridae | Clastoptera |
| 2506AshInt6c | AshInt | June | Hemiptera | Lygaeidae | Kleidocerys |
| 2506AshInt6c | AshInt | June | Hymenoptera | Apidae | Triepeolus |
| 2506AshInt6c | AshInt | June | Hymenoptera | Braconidae | Bracon |
| 2506AshInt6c | AshInt | June | Hymenoptera | Braconidae | Chorebus |
| 2506AshInt6c | AshInt | June | Hymenoptera | Ichneumonidae | Megastylus |
| 2506AshInt6c | AshInt | June | Hymenoptera | Ichneumonidae | Polyblastus |
| 2506AshInt6c | AshInt | June | Hymenoptera | Other | Other |
| 2506AshInt6c | AshInt | June | Hymenoptera | Tenthredinidae | Dolerus |
| 2506AshInt6c | AshInt | June | Hymenoptera | Tenthredinidae | Nematus |
| 2506AshInt6c | AshInt | June | Hymenoptera | Trigonalidae | Orthogonalys |
| 2506AshInt6c | AshInt | June | Lepidoptera | Crambidae | Blepharomastix |
| 2506AshInt6c | AshInt | June | Lepidoptera | Crambidae | Parapoynx |
| 2506AshInt6c | AshInt | June | Lepidoptera | Elachistidae | Elachista |
| 2506AshInt6c | AshInt | June | Lepidoptera | Erebidae | Panopoda |
| 2506AshInt6c | AshInt | June | Lepidoptera | Gelechiidae | Chionodes |
| 2506AshInt6c | AshInt | June | Lepidoptera | Noctuidae | Condica |
| 2506AshInt6c | AshInt | June | Lepidoptera | Noctuidae | Leuconycta |
| 2506AshInt6c | AshInt | June | Lepidoptera | Noctuidae | Pyreferra |
| 2506AshInt6c | AshInt | June | Lepidoptera | Notodontidae | Heterocampa |

| | | | | | |
|---|---|---|---|---|---|
| 2506AshInt6c | AshInt | June | Lepidoptera | Notodontidae | Hyparpax |
| 2506AshInt6c | AshInt | June | Lepidoptera | Nymphalidae | Oeneis |
| 2506AshInt6c | AshInt | June | Lepidoptera | Pyralidae | Atheloca |
| 2506AshInt6c | AshInt | June | Lepidoptera | Pyralidae | Elasmopalpus |
| 2506AshInt6c | AshInt | June | Lepidoptera | Tischeriidae | Coptotriche |
| 2506AshInt6c | AshInt | June | Lepidoptera | Tortricidae | Acleris |
| 2506AshInt6c | AshInt | June | Lepidoptera | Tortricidae | Argyrotaenia |
| 2506AshInt6c | AshInt | June | Lepidoptera | Tortricidae | Rhyacionia |
| 2506AshInt6c | AshInt | June | Megaloptera | Corydalidae | Corydalus |
| 2506AshInt6c | AshInt | June | Neuroptera | Hemerobiidae | Hemerobius |
| 2506AshInt6c | AshInt | June | Odonata | Libellulidae | Brachymesia |
| 2506AshInt6c | AshInt | June | Orthoptera | Tettigoniidae | Conocephalus |
| 2506AshInt6c | AshInt | June | Other | Other | Other |
| 2506AshInt6c | AshInt | June | Other | Other | Other |
| 2506AshInt6c | AshInt | June | Plecoptera | Chloroperlidae | Sweltsa |
| 2506AshInt6c | AshInt | June | Psocodea | Peripsocidae | Peripsocus |
| 2506AshInt6c | AshInt | June | Sarcoptiformes | Oripodidae | Other |
| 2506AshInt6c | AshInt | June | Trichoptera | Dipseudopsidae | Phylocentropus |
| 2506AshInt6c | AshInt | June | Trichoptera | Hydroptilidae | Orthotrichia |
| 2506AshInt6c | AshInt | June | Trichoptera | Leptoceridae | Oecetis |
| 2506AshInt6c | AshInt | June | Trombidiformes | Arrenuridae | Arrenurus |
| 2506AshInt6c | AshInt | June | Trombidiformes | Aturidae | Aturus |
| 2506AshInt6c | AshInt | June | Trombidiformes | Axonopsidae | Brachypoda |
| 2506AshInt6c | AshInt | June | Trombidiformes | Eupodidae | Other |
| 2506AshInt6c | AshInt | June | Trombidiformes | Feltriidae | Feltria |
| 2506AshInt6c | AshInt | June | Trombidiformes | Hydryphantidae | Hydryphantes |
| 2506AshInt6c | AshInt | June | Trombidiformes | Hydryphantidae | Parathyas |
| 2506AshInt6c | AshInt | June | Trombidiformes | Lebertiidae | Lebertia |
| 2506AshInt6c | AshInt | June | Trombidiformes | Mideopsidae | Mideopsis |
| 2506AshInt6c | AshInt | June | Trombidiformes | Pionidae | Piona |
| 2506AshInt6c | AshInt | June | Trombidiformes | Pionidae | Tiphys |

| | | | | | |
|---|---|---|---|---|---|
| 2506AshInt6c | AshInt | June | Trombidiformes | Sperchontidae | Sperchon |
| 2506AshInt6c | AshInt | June | Trombidiformes | Tarsonemidae | Phytonemus |
| 2506AshInt6c | AshInt | June | Trombidiformes | Torrenticolidae | Testudacarus |
| 2506AshInt6c | AshInt | June | Trombidiformes | Torrenticolidae | Torrenticola |
| 2506AshInt6c | AshInt | June | Trombidiformes | Unionicolidae | Koenikea |
| 2507AshInt6a | AshInt | July | Araneae | Agelenidae | Coras |
| 2507AshInt6a | AshInt | July | Araneae | Amaurobiidae | Callobius |
| 2507AshInt6a | AshInt | July | Araneae | Anyphaenidae | Anyphaena |
| 2507AshInt6a | AshInt | July | Araneae | Cicurinidae | Cicurina |
| 2507AshInt6a | AshInt | July | Araneae | Dictynidae | Dictyna |
| 2507AshInt6a | AshInt | July | Araneae | Gnaphosidae | Gnaphosa |
| 2507AshInt6a | AshInt | July | Araneae | Linyphiidae | Grammonota |
| 2507AshInt6a | AshInt | July | Araneae | Lycosidae | Rabidosa |
| 2507AshInt6a | AshInt | July | Araneae | Lycosidae | Schizocosa |
| 2507AshInt6a | AshInt | July | Araneae | Phrurolithidae | Scotinella |
| 2507AshInt6a | AshInt | July | Araneae | Salticidae | Maevia |
| 2507AshInt6a | AshInt | July | Araneae | Salticidae | Sibianor |
| 2507AshInt6a | AshInt | July | Araneae | Theridiidae | Canalidion |
| 2507AshInt6a | AshInt | July | Araneae | Theridiidae | Rugathodes |
| 2507AshInt6a | AshInt | July | Araneae | Theridiidae | Spintharus |
| 2507AshInt6a | AshInt | July | Araneae | Thomisidae | Mecaphesa |
| 2507AshInt6a | AshInt | July | Blattodea | Ectobiidae | Parcoblatta |
| 2507AshInt6a | AshInt | July | Coleoptera | Buprestidae | Pachyschelus |
| 2507AshInt6a | AshInt | July | Coleoptera | Carabidae | Apenes |
| 2507AshInt6a | AshInt | July | Coleoptera | Carabidae | Harpalus |
| 2507AshInt6a | AshInt | July | Coleoptera | Carabidae | Other |
| 2507AshInt6a | AshInt | July | Coleoptera | Carabidae | Polyderis |
| 2507AshInt6a | AshInt | July | Coleoptera | Curculionidae | Conotrachelus |
| 2507AshInt6a | AshInt | July | Coleoptera | Curculionidae | Cyrtepistomus |
| 2507AshInt6a | AshInt | July | Coleoptera | Curculionidae | Listronotus |
| 2507AshInt6a | AshInt | July | Coleoptera | Dytiscidae | Acilius |

| | | | | | |
|---|---|---|---|---|---|
| 2507AshInt6a | AshInt | July | Coleoptera | Lampyridae | Photinus |
| 2507AshInt6a | AshInt | July | Coleoptera | Scarabaeidae | Dialytes |
| 2507AshInt6a | AshInt | July | Coleoptera | Scarabaeidae | Melanocanthon |
| 2507AshInt6a | AshInt | July | Coleoptera | Scirtidae | Contacyphon |
| 2507AshInt6a | AshInt | July | Diptera | Agromyzidae | Calycomyza |
| 2507AshInt6a | AshInt | July | Diptera | Agromyzidae | Cerodontha |
| 2507AshInt6a | AshInt | July | Diptera | Agromyzidae | Japanagromyza |
| 2507AshInt6a | AshInt | July | Diptera | Agromyzidae | Liriomyza |
| 2507AshInt6a | AshInt | July | Diptera | Agromyzidae | Nemorimyza |
| 2507AshInt6a | AshInt | July | Diptera | Agromyzidae | Phytomyza |
| 2507AshInt6a | AshInt | July | Diptera | Anisopodidae | Mycetobia |
| 2507AshInt6a | AshInt | July | Diptera | Calliphoridae | Lucilia |
| 2507AshInt6a | AshInt | July | Diptera | Cecidomyiidae | Other |
| 2507AshInt6a | AshInt | July | Diptera | Chaoboridae | Chaoborus |
| 2507AshInt6a | AshInt | July | Diptera | Chaoboridae | Other |
| 2507AshInt6a | AshInt | July | Diptera | Chironomidae | Ablabesmyia |
| 2507AshInt6a | AshInt | July | Diptera | Chironomidae | Chironomus |
| 2507AshInt6a | AshInt | July | Diptera | Chironomidae | Dicrotendipes |
| 2507AshInt6a | AshInt | July | Diptera | Chironomidae | Endochironomus |
| 2507AshInt6a | AshInt | July | Diptera | Chironomidae | Glyptotendipes |
| 2507AshInt6a | AshInt | July | Diptera | Chironomidae | Microtendipes |
| 2507AshInt6a | AshInt | July | Diptera | Chironomidae | Orthocladius |
| 2507AshInt6a | AshInt | July | Diptera | Chironomidae | Other |
| 2507AshInt6a | AshInt | July | Diptera | Chironomidae | Other |
| 2507AshInt6a | AshInt | July | Diptera | Chironomidae | Other |
| 2507AshInt6a | AshInt | July | Diptera | Chironomidae | Other |
| 2507AshInt6a | AshInt | July | Diptera | Chironomidae | Parachironomus |
| 2507AshInt6a | AshInt | July | Diptera | Chironomidae | Polypedilum |
| 2507AshInt6a | AshInt | July | Diptera | Chironomidae | Pseudochironomus |
| 2507AshInt6a | AshInt | July | Diptera | Chironomidae | Tanytarsus |
| 2507AshInt6a | AshInt | July | Diptera | Chironomidae | Xenochironomus |

| | | | | | |
|---|---|---|---|---|---|
| 2507AshInt6a | AshInt | July | Diptera | Chloropidae | Apotropina |
| 2507AshInt6a | AshInt | July | Diptera | Chloropidae | Elachiptera |
| 2507AshInt6a | AshInt | July | Diptera | Chloropidae | Liohippelates |
| 2507AshInt6a | AshInt | July | Diptera | Chloropidae | Oscinella |
| 2507AshInt6a | AshInt | July | Diptera | Culicidae | Coquillettidia |
| 2507AshInt6a | AshInt | July | Diptera | Culicidae | Culex |
| 2507AshInt6a | AshInt | July | Diptera | Culicidae | Uranotaenia |
| 2507AshInt6a | AshInt | July | Diptera | Drosophilidae | Drosophila |
| 2507AshInt6a | AshInt | July | Diptera | Empididae | Other |
| 2507AshInt6a | AshInt | July | Diptera | Hybotidae | Platypalpus |
| 2507AshInt6a | AshInt | July | Diptera | Hybotidae | Syndyas |
| 2507AshInt6a | AshInt | July | Diptera | Keroplatidae | Other |
| 2507AshInt6a | AshInt | July | Diptera | Limoniidae | Dicranomyia |
| 2507AshInt6a | AshInt | July | Diptera | Limoniidae | Dicranophragma |
| 2507AshInt6a | AshInt | July | Diptera | Limoniidae | Epiphragma |
| 2507AshInt6a | AshInt | July | Diptera | Limoniidae | Erioptera |
| 2507AshInt6a | AshInt | July | Diptera | Limoniidae | Metalimnobia |
| 2507AshInt6a | AshInt | July | Diptera | Limoniidae | Ormosia |
| 2507AshInt6a | AshInt | July | Diptera | Limoniidae | Other |
| 2507AshInt6a | AshInt | July | Diptera | Limoniidae | Other |
| 2507AshInt6a | AshInt | July | Diptera | Limoniidae | Pseudolimnophila |
| 2507AshInt6a | AshInt | July | Diptera | Muscidae | Mydaea |
| 2507AshInt6a | AshInt | July | Diptera | Mycetophilidae | Mycomya |
| 2507AshInt6a | AshInt | July | Diptera | Mycetophilidae | Neoempheria |
| 2507AshInt6a | AshInt | July | Diptera | Opomyzidae | Geomyza |
| 2507AshInt6a | AshInt | July | Diptera | Pallopteridae | Toxonevra |
| 2507AshInt6a | AshInt | July | Diptera | Phoridae | Diplonevra |
| 2507AshInt6a | AshInt | July | Diptera | Polleniidae | Pollenia |
| 2507AshInt6a | AshInt | July | Diptera | Psychodidae | Clogmia |
| 2507AshInt6a | AshInt | July | Diptera | Psychodidae | Psychoda |
| 2507AshInt6a | AshInt | July | Diptera | Sarcophagidae | Ravinia |

| | | | | | |
|---|---|---|---|---|---|
| 2507AshInt6a | AshInt | July | Diptera | Sciaridae | Claustropyga |
| 2507AshInt6a | AshInt | July | Diptera | Sciaridae | Corynoptera |
| 2507AshInt6a | AshInt | July | Diptera | Sciaridae | Epidapus |
| 2507AshInt6a | AshInt | July | Diptera | Sciaridae | Other |
| 2507AshInt6a | AshInt | July | Diptera | Sphaeroceridae | Leptocera |
| 2507AshInt6a | AshInt | July | Diptera | Syrphidae | Chalcosyrphus |
| 2507AshInt6a | AshInt | July | Diptera | Syrphidae | Palpada |
| 2507AshInt6a | AshInt | July | Diptera | Tabanidae | Hybomitra |
| 2507AshInt6a | AshInt | July | Diptera | Tabanidae | Other |
| 2507AshInt6a | AshInt | July | Diptera | Tachinidae | Cryptomeigenia |
| 2507AshInt6a | AshInt | July | Diptera | Tachinidae | Cyzenis |
| 2507AshInt6a | AshInt | July | Diptera | Tachinidae | Zaira |
| 2507AshInt6a | AshInt | July | Diptera | Tipulidae | Dolichopeza |
| 2507AshInt6a | AshInt | July | Diptera | Tipulidae | Tipula |
| 2507AshInt6a | AshInt | July | Ephemeroptera | Caenidae | Caenis |
| 2507AshInt6a | AshInt | July | Ephemeroptera | Ephemeridae | Hexagenia |
| 2507AshInt6a | AshInt | July | Ephemeroptera | Heptageniidae | Epeorus |
| 2507AshInt6a | AshInt | July | Hemiptera | Clastopteridae | Clastoptera |
| 2507AshInt6a | AshInt | July | Hemiptera | Lygaeidae | Kleidocerys |
| 2507AshInt6a | AshInt | July | Hymenoptera | Apidae | Triepeolus |
| 2507AshInt6a | AshInt | July | Hymenoptera | Braconidae | Bracon |
| 2507AshInt6a | AshInt | July | Hymenoptera | Braconidae | Chorebus |
| 2507AshInt6a | AshInt | July | Hymenoptera | Ichneumonidae | Megastylus |
| 2507AshInt6a | AshInt | July | Hymenoptera | Ichneumonidae | Polyblastus |
| 2507AshInt6a | AshInt | July | Hymenoptera | Other | Other |
| 2507AshInt6a | AshInt | July | Hymenoptera | Tenthredinidae | Dolerus |
| 2507AshInt6a | AshInt | July | Hymenoptera | Tenthredinidae | Nematus |
| 2507AshInt6a | AshInt | July | Hymenoptera | Trigonalidae | Orthogonalys |
| 2507AshInt6a | AshInt | July | Lepidoptera | Crambidae | Blepharomastix |
| 2507AshInt6a | AshInt | July | Lepidoptera | Crambidae | Parapoynx |
| 2507AshInt6a | AshInt | July | Lepidoptera | Elachistidae | Elachista |

| | | | | | |
|---|---|---|---|---|---|
| 2507AshInt6a | AshInt | July | Lepidoptera | Erebidae | Panopoda |
| 2507AshInt6a | AshInt | July | Lepidoptera | Gelechiidae | Chionodes |
| 2507AshInt6a | AshInt | July | Lepidoptera | Noctuidae | Condica |
| 2507AshInt6a | AshInt | July | Lepidoptera | Noctuidae | Leuconycta |
| 2507AshInt6a | AshInt | July | Lepidoptera | Noctuidae | Pyreferra |
| 2507AshInt6a | AshInt | July | Lepidoptera | Notodontidae | Heterocampa |
| 2507AshInt6a | AshInt | July | Lepidoptera | Notodontidae | Hyparpax |
| 2507AshInt6a | AshInt | July | Lepidoptera | Nymphalidae | Oeneis |
| 2507AshInt6a | AshInt | July | Lepidoptera | Pyralidae | Atheloca |
| 2507AshInt6a | AshInt | July | Lepidoptera | Pyralidae | Elasmopalpus |
| 2507AshInt6a | AshInt | July | Lepidoptera | Tischeriidae | Coptotriche |
| 2507AshInt6a | AshInt | July | Lepidoptera | Tortricidae | Acleris |
| 2507AshInt6a | AshInt | July | Lepidoptera | Tortricidae | Argyrotaenia |
| 2507AshInt6a | AshInt | July | Lepidoptera | Tortricidae | Rhyacionia |
| 2507AshInt6a | AshInt | July | Megaloptera | Corydalidae | Corydalus |
| 2507AshInt6a | AshInt | July | Neuroptera | Hemerobiidae | Hemerobius |
| 2507AshInt6a | AshInt | July | Odonata | Libellulidae | Brachymesia |
| 2507AshInt6a | AshInt | July | Orthoptera | Tettigoniidae | Conocephalus |
| 2507AshInt6a | AshInt | July | Other | Other | Other |
| 2507AshInt6a | AshInt | July | Other | Other | Other |
| 2507AshInt6a | AshInt | July | Plecoptera | Chloroperlidae | Sweltsa |
| 2507AshInt6a | AshInt | July | Psocodea | Peripsocidae | Peripsocus |
| 2507AshInt6a | AshInt | July | Sarcoptiformes | Oripodidae | Other |
| 2507AshInt6a | AshInt | July | Trichoptera | Dipseudopsidae | Phylocentropus |
| 2507AshInt6a | AshInt | July | Trichoptera | Hydroptilidae | Orthotrichia |
| 2507AshInt6a | AshInt | July | Trichoptera | Leptoceridae | Oecetis |
| 2507AshInt6a | AshInt | July | Trombidiformes | Arrenuridae | Arrenurus |
| 2507AshInt6a | AshInt | July | Trombidiformes | Aturidae | Aturus |
| 2507AshInt6a | AshInt | July | Trombidiformes | Axonopsidae | Brachypoda |
| 2507AshInt6a | AshInt | July | Trombidiformes | Eupodidae | Other |
| 2507AshInt6a | AshInt | July | Trombidiformes | Feltriidae | Feltria |

| | | | | | |
|---|---|---|---|---|---|
| 2507AshInt6a | AshInt | July | Trombidiformes | Hydryphantidae | Hydryphantes |
| 2507AshInt6a | AshInt | July | Trombidiformes | Hydryphantidae | Parathyas |
| 2507AshInt6a | AshInt | July | Trombidiformes | Lebertiidae | Lebertia |
| 2507AshInt6a | AshInt | July | Trombidiformes | Mideopsidae | Mideopsis |
| 2507AshInt6a | AshInt | July | Trombidiformes | Pionidae | Piona |
| 2507AshInt6a | AshInt | July | Trombidiformes | Pionidae | Tiphys |
| 2507AshInt6a | AshInt | July | Trombidiformes | Sperchontidae | Sperchon |
| 2507AshInt6a | AshInt | July | Trombidiformes | Tarsonemidae | Phytonemus |
| 2507AshInt6a | AshInt | July | Trombidiformes | Torrenticolidae | Testudacarus |
| 2507AshInt6a | AshInt | July | Trombidiformes | Torrenticolidae | Torrenticola |
| 2507AshInt6a | AshInt | July | Trombidiformes | Unionicolidae | Koenikea |
| 2507AshInt6b | AshInt | July | Araneae | Agelenidae | Coras |
| 2507AshInt6b | AshInt | July | Araneae | Amaurobiidae | Callobius |
| 2507AshInt6b | AshInt | July | Araneae | Anyphaenidae | Anyphaena |
| 2507AshInt6b | AshInt | July | Araneae | Cicurinidae | Cicurina |
| 2507AshInt6b | AshInt | July | Araneae | Dictynidae | Dictyna |
| 2507AshInt6b | AshInt | July | Araneae | Gnaphosidae | Gnaphosa |
| 2507AshInt6b | AshInt | July | Araneae | Linyphiidae | Grammonota |
| 2507AshInt6b | AshInt | July | Araneae | Lycosidae | Rabidosa |
| 2507AshInt6b | AshInt | July | Araneae | Lycosidae | Schizocosa |
| 2507AshInt6b | AshInt | July | Araneae | Phrurolithidae | Scotinella |
| 2507AshInt6b | AshInt | July | Araneae | Salticidae | Maevia |
| 2507AshInt6b | AshInt | July | Araneae | Salticidae | Sibianor |
| 2507AshInt6b | AshInt | July | Araneae | Theridiidae | Canalidion |
| 2507AshInt6b | AshInt | July | Araneae | Theridiidae | Rugathodes |
| 2507AshInt6b | AshInt | July | Araneae | Theridiidae | Spintharus |
| 2507AshInt6b | AshInt | July | Araneae | Thomisidae | Mecaphesa |
| 2507AshInt6b | AshInt | July | Blattodea | Ectobiidae | Parcoblatta |
| 2507AshInt6b | AshInt | July | Coleoptera | Buprestidae | Pachyschelus |
| 2507AshInt6b | AshInt | July | Coleoptera | Carabidae | Apenes |
| 2507AshInt6b | AshInt | July | Coleoptera | Carabidae | Harpalus |

| | | | | | |
|---|---|---|---|---|---|
| 2507AshInt6b | AshInt | July | Coleoptera | Carabidae | Other |
| 2507AshInt6b | AshInt | July | Coleoptera | Carabidae | Polyderis |
| 2507AshInt6b | AshInt | July | Coleoptera | Curculionidae | Conotrachelus |
| 2507AshInt6b | AshInt | July | Coleoptera | Curculionidae | Cyrtepistomus |
| 2507AshInt6b | AshInt | July | Coleoptera | Curculionidae | Listronotus |
| 2507AshInt6b | AshInt | July | Coleoptera | Dytiscidae | Acilius |
| 2507AshInt6b | AshInt | July | Coleoptera | Lampyridae | Photinus |
| 2507AshInt6b | AshInt | July | Coleoptera | Scarabaeidae | Dialytes |
| 2507AshInt6b | AshInt | July | Coleoptera | Scarabaeidae | Melanocanthon |
| 2507AshInt6b | AshInt | July | Coleoptera | Scirtidae | Contacyphon |
| 2507AshInt6b | AshInt | July | Diptera | Agromyzidae | Calycomyza |
| 2507AshInt6b | AshInt | July | Diptera | Agromyzidae | Cerodontha |
| 2507AshInt6b | AshInt | July | Diptera | Agromyzidae | Japanagromyza |
| 2507AshInt6b | AshInt | July | Diptera | Agromyzidae | Liriomyza |
| 2507AshInt6b | AshInt | July | Diptera | Agromyzidae | Nemorimyza |
| 2507AshInt6b | AshInt | July | Diptera | Agromyzidae | Phytomyza |
| 2507AshInt6b | AshInt | July | Diptera | Anisopodidae | Mycetobia |
| 2507AshInt6b | AshInt | July | Diptera | Calliphoridae | Lucilia |
| 2507AshInt6b | AshInt | July | Diptera | Cecidomyiidae | Other |
| 2507AshInt6b | AshInt | July | Diptera | Chaoboridae | Chaoborus |
| 2507AshInt6b | AshInt | July | Diptera | Chaoboridae | Other |
| 2507AshInt6b | AshInt | July | Diptera | Chironomidae | Ablabesmyia |
| 2507AshInt6b | AshInt | July | Diptera | Chironomidae | Chironomus |
| 2507AshInt6b | AshInt | July | Diptera | Chironomidae | Dicrotendipes |
| 2507AshInt6b | AshInt | July | Diptera | Chironomidae | Endochironomus |
| 2507AshInt6b | AshInt | July | Diptera | Chironomidae | Glyptotendipes |
| 2507AshInt6b | AshInt | July | Diptera | Chironomidae | Microtendipes |
| 2507AshInt6b | AshInt | July | Diptera | Chironomidae | Orthocladius |
| 2507AshInt6b | AshInt | July | Diptera | Chironomidae | Other |
| 2507AshInt6b | AshInt | July | Diptera | Chironomidae | Other |
| 2507AshInt6b | AshInt | July | Diptera | Chironomidae | Other |

| | | | | | |
|---|---|---|---|---|---|
| 2507AshInt6b | AshInt | July | Diptera | Chironomidae | Other |
| 2507AshInt6b | AshInt | July | Diptera | Chironomidae | Parachironomus |
| 2507AshInt6b | AshInt | July | Diptera | Chironomidae | Polypedilum |
| 2507AshInt6b | AshInt | July | Diptera | Chironomidae | Pseudochironomus |
| 2507AshInt6b | AshInt | July | Diptera | Chironomidae | Tanytarsus |
| 2507AshInt6b | AshInt | July | Diptera | Chironomidae | Xenochironomus |
| 2507AshInt6b | AshInt | July | Diptera | Chloropidae | Apotropina |
| 2507AshInt6b | AshInt | July | Diptera | Chloropidae | Elachiptera |
| 2507AshInt6b | AshInt | July | Diptera | Chloropidae | Liohippelates |
| 2507AshInt6b | AshInt | July | Diptera | Chloropidae | Oscinella |
| 2507AshInt6b | AshInt | July | Diptera | Culicidae | Coquillettidia |
| 2507AshInt6b | AshInt | July | Diptera | Culicidae | Culex |
| 2507AshInt6b | AshInt | July | Diptera | Culicidae | Uranotaenia |
| 2507AshInt6b | AshInt | July | Diptera | Drosophilidae | Drosophila |
| 2507AshInt6b | AshInt | July | Diptera | Empididae | Other |
| 2507AshInt6b | AshInt | July | Diptera | Hybotidae | Platypalpus |
| 2507AshInt6b | AshInt | July | Diptera | Hybotidae | Syndyas |
| 2507AshInt6b | AshInt | July | Diptera | Keroplatidae | Other |
| 2507AshInt6b | AshInt | July | Diptera | Limoniidae | Dicranomyia |
| 2507AshInt6b | AshInt | July | Diptera | Limoniidae | Dicranophragma |
| 2507AshInt6b | AshInt | July | Diptera | Limoniidae | Epiphragma |
| 2507AshInt6b | AshInt | July | Diptera | Limoniidae | Erioptera |
| 2507AshInt6b | AshInt | July | Diptera | Limoniidae | Metalimnobia |
| 2507AshInt6b | AshInt | July | Diptera | Limoniidae | Ormosia |
| 2507AshInt6b | AshInt | July | Diptera | Limoniidae | Other |
| 2507AshInt6b | AshInt | July | Diptera | Limoniidae | Other |
| 2507AshInt6b | AshInt | July | Diptera | Limoniidae | Pseudolimnophila |
| 2507AshInt6b | AshInt | July | Diptera | Muscidae | Mydaea |
| 2507AshInt6b | AshInt | July | Diptera | Mycetophilidae | Mycomya |
| 2507AshInt6b | AshInt | July | Diptera | Mycetophilidae | Neoempheria |
| 2507AshInt6b | AshInt | July | Diptera | Opomyzidae | Geomyza |

| | | | | | |
|---|---|---|---|---|---|
| 2507AshInt6b | AshInt | July | Diptera | Pallopteridae | Toxonevra |
| 2507AshInt6b | AshInt | July | Diptera | Phoridae | Diplonevra |
| 2507AshInt6b | AshInt | July | Diptera | Polleniidae | Pollenia |
| 2507AshInt6b | AshInt | July | Diptera | Psychodidae | Clogmia |
| 2507AshInt6b | AshInt | July | Diptera | Psychodidae | Psychoda |
| 2507AshInt6b | AshInt | July | Diptera | Sarcophagidae | Ravinia |
| 2507AshInt6b | AshInt | July | Diptera | Sciaridae | Claustropyga |
| 2507AshInt6b | AshInt | July | Diptera | Sciaridae | Corynoptera |
| 2507AshInt6b | AshInt | July | Diptera | Sciaridae | Epidapus |
| 2507AshInt6b | AshInt | July | Diptera | Sciaridae | Other |
| 2507AshInt6b | AshInt | July | Diptera | Sphaeroceridae | Leptocera |
| 2507AshInt6b | AshInt | July | Diptera | Syrphidae | Chalcosyrphus |
| 2507AshInt6b | AshInt | July | Diptera | Syrphidae | Palpada |
| 2507AshInt6b | AshInt | July | Diptera | Tabanidae | Hybomitra |
| 2507AshInt6b | AshInt | July | Diptera | Tabanidae | Other |
| 2507AshInt6b | AshInt | July | Diptera | Tachinidae | Cryptomeigenia |
| 2507AshInt6b | AshInt | July | Diptera | Tachinidae | Cyzenis |
| 2507AshInt6b | AshInt | July | Diptera | Tachinidae | Zaira |
| 2507AshInt6b | AshInt | July | Diptera | Tipulidae | Dolichopeza |
| 2507AshInt6b | AshInt | July | Diptera | Tipulidae | Tipula |
| 2507AshInt6b | AshInt | July | Ephemeroptera | Caenidae | Caenis |
| 2507AshInt6b | AshInt | July | Ephemeroptera | Ephemeridae | Hexagenia |
| 2507AshInt6b | AshInt | July | Ephemeroptera | Heptageniidae | Epeorus |
| 2507AshInt6b | AshInt | July | Hemiptera | Clastopteridae | Clastoptera |
| 2507AshInt6b | AshInt | July | Hemiptera | Lygaeidae | Kleidocerys |
| 2507AshInt6b | AshInt | July | Hymenoptera | Apidae | Triepeolus |
| 2507AshInt6b | AshInt | July | Hymenoptera | Braconidae | Bracon |
| 2507AshInt6b | AshInt | July | Hymenoptera | Braconidae | Chorebus |
| 2507AshInt6b | AshInt | July | Hymenoptera | Ichneumonidae | Megastylus |
| 2507AshInt6b | AshInt | July | Hymenoptera | Ichneumonidae | Polyblastus |
| 2507AshInt6b | AshInt | July | Hymenoptera | Other | Other |

| | | | | | |
|---|---|---|---|---|---|
| 2507AshInt6b | AshInt | July | Hymenoptera | Tenthredinidae | Dolerus |
| 2507AshInt6b | AshInt | July | Hymenoptera | Tenthredinidae | Nematus |
| 2507AshInt6b | AshInt | July | Hymenoptera | Trigonalidae | Orthogonalys |
| 2507AshInt6b | AshInt | July | Lepidoptera | Crambidae | Blepharomastix |
| 2507AshInt6b | AshInt | July | Lepidoptera | Crambidae | Parapoynx |
| 2507AshInt6b | AshInt | July | Lepidoptera | Elachistidae | Elachista |
| 2507AshInt6b | AshInt | July | Lepidoptera | Erebidae | Panopoda |
| 2507AshInt6b | AshInt | July | Lepidoptera | Gelechiidae | Chionodes |
| 2507AshInt6b | AshInt | July | Lepidoptera | Noctuidae | Condica |
| 2507AshInt6b | AshInt | July | Lepidoptera | Noctuidae | Leuconycta |
| 2507AshInt6b | AshInt | July | Lepidoptera | Noctuidae | Pyreferra |
| 2507AshInt6b | AshInt | July | Lepidoptera | Notodontidae | Heterocampa |
| 2507AshInt6b | AshInt | July | Lepidoptera | Notodontidae | Hyparpax |
| 2507AshInt6b | AshInt | July | Lepidoptera | Nymphalidae | Oeneis |
| 2507AshInt6b | AshInt | July | Lepidoptera | Pyralidae | Atheloca |
| 2507AshInt6b | AshInt | July | Lepidoptera | Pyralidae | Elasmopalpus |
| 2507AshInt6b | AshInt | July | Lepidoptera | Tischeriidae | Coptotriche |
| 2507AshInt6b | AshInt | July | Lepidoptera | Tortricidae | Acleris |
| 2507AshInt6b | AshInt | July | Lepidoptera | Tortricidae | Argyrotaenia |
| 2507AshInt6b | AshInt | July | Lepidoptera | Tortricidae | Rhyacionia |
| 2507AshInt6b | AshInt | July | Megaloptera | Corydalidae | Corydalus |
| 2507AshInt6b | AshInt | July | Neuroptera | Hemerobiidae | Hemerobius |
| 2507AshInt6b | AshInt | July | Odonata | Libellulidae | Brachymesia |
| 2507AshInt6b | AshInt | July | Orthoptera | Tettigoniidae | Conocephalus |
| 2507AshInt6b | AshInt | July | Other | Other | Other |
| 2507AshInt6b | AshInt | July | Other | Other | Other |
| 2507AshInt6b | AshInt | July | Plecoptera | Chloroperlidae | Sweltsa |
| 2507AshInt6b | AshInt | July | Psocodea | Peripsocidae | Peripsocus |
| 2507AshInt6b | AshInt | July | Sarcoptiformes | Oripodidae | Other |
| 2507AshInt6b | AshInt | July | Trichoptera | Dipseudopsidae | Phylocentropus |
| 2507AshInt6b | AshInt | July | Trichoptera | Hydroptilidae | Orthotrichia |

| | | | | | |
|---|---|---|---|---|---|
| 2507AshInt6b | AshInt | July | Trichoptera | Leptoceridae | Oecetis |
| 2507AshInt6b | AshInt | July | Trombidiformes | Arrenuridae | Arrenurus |
| 2507AshInt6b | AshInt | July | Trombidiformes | Aturidae | Aturus |
| 2507AshInt6b | AshInt | July | Trombidiformes | Axonopsidae | Brachypoda |
| 2507AshInt6b | AshInt | July | Trombidiformes | Eupodidae | Other |
| 2507AshInt6b | AshInt | July | Trombidiformes | Feltriidae | Feltria |
| 2507AshInt6b | AshInt | July | Trombidiformes | Hydryphantidae | Hydryphantes |
| 2507AshInt6b | AshInt | July | Trombidiformes | Hydryphantidae | Parathyas |
| 2507AshInt6b | AshInt | July | Trombidiformes | Lebertiidae | Lebertia |
| 2507AshInt6b | AshInt | July | Trombidiformes | Mideopsidae | Mideopsis |
| 2507AshInt6b | AshInt | July | Trombidiformes | Pionidae | Piona |
| 2507AshInt6b | AshInt | July | Trombidiformes | Pionidae | Tiphys |
| 2507AshInt6b | AshInt | July | Trombidiformes | Sperchontidae | Sperchon |
| 2507AshInt6b | AshInt | July | Trombidiformes | Tarsonemidae | Phytonemus |
| 2507AshInt6b | AshInt | July | Trombidiformes | Torrenticolidae | Testudacarus |
| 2507AshInt6b | AshInt | July | Trombidiformes | Torrenticolidae | Torrenticola |
| 2507AshInt6b | AshInt | July | Trombidiformes | Unionicolidae | Koenikea |
| 2507AshInt6c | AshInt | July | Araneae | Agelenidae | Coras |
| 2507AshInt6c | AshInt | July | Araneae | Amaurobiidae | Callobius |
| 2507AshInt6c | AshInt | July | Araneae | Anyphaenidae | Anyphaena |
| 2507AshInt6c | AshInt | July | Araneae | Cicurinidae | Cicurina |
| 2507AshInt6c | AshInt | July | Araneae | Dictynidae | Dictyna |
| 2507AshInt6c | AshInt | July | Araneae | Gnaphosidae | Gnaphosa |
| 2507AshInt6c | AshInt | July | Araneae | Linyphiidae | Grammonota |
| 2507AshInt6c | AshInt | July | Araneae | Lycosidae | Rabidosa |
| 2507AshInt6c | AshInt | July | Araneae | Lycosidae | Schizocosa |
| 2507AshInt6c | AshInt | July | Araneae | Phrurolithidae | Scotinella |
| 2507AshInt6c | AshInt | July | Araneae | Salticidae | Maevia |
| 2507AshInt6c | AshInt | July | Araneae | Salticidae | Sibianor |
| 2507AshInt6c | AshInt | July | Araneae | Theridiidae | Canalidion |
| 2507AshInt6c | AshInt | July | Araneae | Theridiidae | Rugathodes |

| | | | | | |
|---|---|---|---|---|---|
| 2507AshInt6c | AshInt | July | Araneae | Theridiidae | Spintharus |
| 2507AshInt6c | AshInt | July | Araneae | Thomisidae | Mecaphesa |
| 2507AshInt6c | AshInt | July | Blattodea | Ectobiidae | Parcoblatta |
| 2507AshInt6c | AshInt | July | Coleoptera | Buprestidae | Pachyschelus |
| 2507AshInt6c | AshInt | July | Coleoptera | Carabidae | Apenes |
| 2507AshInt6c | AshInt | July | Coleoptera | Carabidae | Harpalus |
| 2507AshInt6c | AshInt | July | Coleoptera | Carabidae | Other |
| 2507AshInt6c | AshInt | July | Coleoptera | Carabidae | Polyderis |
| 2507AshInt6c | AshInt | July | Coleoptera | Curculionidae | Conotrachelus |
| 2507AshInt6c | AshInt | July | Coleoptera | Curculionidae | Cyrtepistomus |
| 2507AshInt6c | AshInt | July | Coleoptera | Curculionidae | Listronotus |
| 2507AshInt6c | AshInt | July | Coleoptera | Dytiscidae | Acilius |
| 2507AshInt6c | AshInt | July | Coleoptera | Lampyridae | Photinus |
| 2507AshInt6c | AshInt | July | Coleoptera | Scarabaeidae | Dialytes |
| 2507AshInt6c | AshInt | July | Coleoptera | Scarabaeidae | Melanocanthon |
| 2507AshInt6c | AshInt | July | Coleoptera | Scirtidae | Contacyphon |
| 2507AshInt6c | AshInt | July | Diptera | Agromyzidae | Calycomyza |
| 2507AshInt6c | AshInt | July | Diptera | Agromyzidae | Cerodontha |
| 2507AshInt6c | AshInt | July | Diptera | Agromyzidae | Japanagromyza |
| 2507AshInt6c | AshInt | July | Diptera | Agromyzidae | Liriomyza |
| 2507AshInt6c | AshInt | July | Diptera | Agromyzidae | Nemorimyza |
| 2507AshInt6c | AshInt | July | Diptera | Agromyzidae | Phytomyza |
| 2507AshInt6c | AshInt | July | Diptera | Anisopodidae | Mycetobia |
| 2507AshInt6c | AshInt | July | Diptera | Calliphoridae | Lucilia |
| 2507AshInt6c | AshInt | July | Diptera | Cecidomyiidae | Other |
| 2507AshInt6c | AshInt | July | Diptera | Chaoboridae | Chaoborus |
| 2507AshInt6c | AshInt | July | Diptera | Chaoboridae | Other |
| 2507AshInt6c | AshInt | July | Diptera | Chironomidae | Ablabesmyia |
| 2507AshInt6c | AshInt | July | Diptera | Chironomidae | Chironomus |
| 2507AshInt6c | AshInt | July | Diptera | Chironomidae | Dicrotendipes |
| 2507AshInt6c | AshInt | July | Diptera | Chironomidae | Endochironomus |

| | | | | | |
|---|---|---|---|---|---|
| 2507AshInt6c | AshInt | July | Diptera | Chironomidae | Glyptotendipes |
| 2507AshInt6c | AshInt | July | Diptera | Chironomidae | Microtendipes |
| 2507AshInt6c | AshInt | July | Diptera | Chironomidae | Orthocladius |
| 2507AshInt6c | AshInt | July | Diptera | Chironomidae | Other |
| 2507AshInt6c | AshInt | July | Diptera | Chironomidae | Other |
| 2507AshInt6c | AshInt | July | Diptera | Chironomidae | Other |
| 2507AshInt6c | AshInt | July | Diptera | Chironomidae | Other |
| 2507AshInt6c | AshInt | July | Diptera | Chironomidae | Parachironomus |
| 2507AshInt6c | AshInt | July | Diptera | Chironomidae | Polypedilum |
| 2507AshInt6c | AshInt | July | Diptera | Chironomidae | Pseudochironomus |
| 2507AshInt6c | AshInt | July | Diptera | Chironomidae | Tanytarsus |
| 2507AshInt6c | AshInt | July | Diptera | Chironomidae | Xenochironomus |
| 2507AshInt6c | AshInt | July | Diptera | Chloropidae | Apotropina |
| 2507AshInt6c | AshInt | July | Diptera | Chloropidae | Elachiptera |
| 2507AshInt6c | AshInt | July | Diptera | Chloropidae | Liohippelates |
| 2507AshInt6c | AshInt | July | Diptera | Chloropidae | Oscinella |
| 2507AshInt6c | AshInt | July | Diptera | Culicidae | Coquillettidia |
| 2507AshInt6c | AshInt | July | Diptera | Culicidae | Culex |
| 2507AshInt6c | AshInt | July | Diptera | Culicidae | Uranotaenia |
| 2507AshInt6c | AshInt | July | Diptera | Drosophilidae | Drosophila |
| 2507AshInt6c | AshInt | July | Diptera | Empididae | Other |
| 2507AshInt6c | AshInt | July | Diptera | Hybotidae | Platypalpus |
| 2507AshInt6c | AshInt | July | Diptera | Hybotidae | Syndyas |
| 2507AshInt6c | AshInt | July | Diptera | Keroplatidae | Other |
| 2507AshInt6c | AshInt | July | Diptera | Limoniidae | Dicranomyia |
| 2507AshInt6c | AshInt | July | Diptera | Limoniidae | Dicranophragma |
| 2507AshInt6c | AshInt | July | Diptera | Limoniidae | Epiphragma |
| 2507AshInt6c | AshInt | July | Diptera | Limoniidae | Erioptera |
| 2507AshInt6c | AshInt | July | Diptera | Limoniidae | Metalimnobia |
| 2507AshInt6c | AshInt | July | Diptera | Limoniidae | Ormosia |
| 2507AshInt6c | AshInt | July | Diptera | Limoniidae | Other |

| | | | | | |
|---|---|---|---|---|---|
| 2507AshInt6c | AshInt | July | Diptera | Limoniidae | Other |
| 2507AshInt6c | AshInt | July | Diptera | Limoniidae | Pseudolimnophila |
| 2507AshInt6c | AshInt | July | Diptera | Muscidae | Mydaea |
| 2507AshInt6c | AshInt | July | Diptera | Mycetophilidae | Mycomya |
| 2507AshInt6c | AshInt | July | Diptera | Mycetophilidae | Neoempheria |
| 2507AshInt6c | AshInt | July | Diptera | Opomyzidae | Geomyza |
| 2507AshInt6c | AshInt | July | Diptera | Pallopteridae | Toxonevra |
| 2507AshInt6c | AshInt | July | Diptera | Phoridae | Diplonevra |
| 2507AshInt6c | AshInt | July | Diptera | Polleniidae | Pollenia |
| 2507AshInt6c | AshInt | July | Diptera | Psychodidae | Clogmia |
| 2507AshInt6c | AshInt | July | Diptera | Psychodidae | Psychoda |
| 2507AshInt6c | AshInt | July | Diptera | Sarcophagidae | Ravinia |
| 2507AshInt6c | AshInt | July | Diptera | Sciaridae | Claustropyga |
| 2507AshInt6c | AshInt | July | Diptera | Sciaridae | Corynoptera |
| 2507AshInt6c | AshInt | July | Diptera | Sciaridae | Epidapus |
| 2507AshInt6c | AshInt | July | Diptera | Sciaridae | Other |
| 2507AshInt6c | AshInt | July | Diptera | Sphaeroceridae | Leptocera |
| 2507AshInt6c | AshInt | July | Diptera | Syrphidae | Chalcosyrphus |
| 2507AshInt6c | AshInt | July | Diptera | Syrphidae | Palpada |
| 2507AshInt6c | AshInt | July | Diptera | Tabanidae | Hybomitra |
| 2507AshInt6c | AshInt | July | Diptera | Tabanidae | Other |
| 2507AshInt6c | AshInt | July | Diptera | Tachinidae | Cryptomeigenia |
| 2507AshInt6c | AshInt | July | Diptera | Tachinidae | Cyzenis |
| 2507AshInt6c | AshInt | July | Diptera | Tachinidae | Zaira |
| 2507AshInt6c | AshInt | July | Diptera | Tipulidae | Dolichopeza |
| 2507AshInt6c | AshInt | July | Diptera | Tipulidae | Tipula |
| 2507AshInt6c | AshInt | July | Ephemeroptera | Caenidae | Caenis |
| 2507AshInt6c | AshInt | July | Ephemeroptera | Ephemeridae | Hexagenia |
| 2507AshInt6c | AshInt | July | Ephemeroptera | Heptageniidae | Epeorus |
| 2507AshInt6c | AshInt | July | Hemiptera | Clastopteridae | Clastoptera |
| 2507AshInt6c | AshInt | July | Hemiptera | Lygaeidae | Kleidocerys |

| | | | | | |
|---|---|---|---|---|---|
| 2507AshInt6c | AshInt | July | Hymenoptera | Apidae | Triepeolus |
| 2507AshInt6c | AshInt | July | Hymenoptera | Braconidae | Bracon |
| 2507AshInt6c | AshInt | July | Hymenoptera | Braconidae | Chorebus |
| 2507AshInt6c | AshInt | July | Hymenoptera | Ichneumonidae | Megastylus |
| 2507AshInt6c | AshInt | July | Hymenoptera | Ichneumonidae | Polyblastus |
| 2507AshInt6c | AshInt | July | Hymenoptera | Other | Other |
| 2507AshInt6c | AshInt | July | Hymenoptera | Tenthredinidae | Dolerus |
| 2507AshInt6c | AshInt | July | Hymenoptera | Tenthredinidae | Nematus |
| 2507AshInt6c | AshInt | July | Hymenoptera | Trigonalidae | Orthogonalys |
| 2507AshInt6c | AshInt | July | Lepidoptera | Crambidae | Blepharomastix |
| 2507AshInt6c | AshInt | July | Lepidoptera | Crambidae | Parapoynx |
| 2507AshInt6c | AshInt | July | Lepidoptera | Elachistidae | Elachista |
| 2507AshInt6c | AshInt | July | Lepidoptera | Erebidae | Panopoda |
| 2507AshInt6c | AshInt | July | Lepidoptera | Gelechiidae | Chionodes |
| 2507AshInt6c | AshInt | July | Lepidoptera | Noctuidae | Condica |
| 2507AshInt6c | AshInt | July | Lepidoptera | Noctuidae | Leuconycta |
| 2507AshInt6c | AshInt | July | Lepidoptera | Noctuidae | Pyreferra |
| 2507AshInt6c | AshInt | July | Lepidoptera | Notodontidae | Heterocampa |
| 2507AshInt6c | AshInt | July | Lepidoptera | Notodontidae | Hyparpax |
| 2507AshInt6c | AshInt | July | Lepidoptera | Nymphalidae | Oeneis |
| 2507AshInt6c | AshInt | July | Lepidoptera | Pyralidae | Atheloca |
| 2507AshInt6c | AshInt | July | Lepidoptera | Pyralidae | Elasmopalpus |
| 2507AshInt6c | AshInt | July | Lepidoptera | Tischeriidae | Coptotriche |
| 2507AshInt6c | AshInt | July | Lepidoptera | Tortricidae | Acleris |
| 2507AshInt6c | AshInt | July | Lepidoptera | Tortricidae | Argyrotaenia |
| 2507AshInt6c | AshInt | July | Lepidoptera | Tortricidae | Rhyacionia |
| 2507AshInt6c | AshInt | July | Megaloptera | Corydalidae | Corydalus |
| 2507AshInt6c | AshInt | July | Neuroptera | Hemerobiidae | Hemerobius |
| 2507AshInt6c | AshInt | July | Odonata | Libellulidae | Brachymesia |
| 2507AshInt6c | AshInt | July | Orthoptera | Tettigoniidae | Conocephalus |
| 2507AshInt6c | AshInt | July | Other | Other | Other |

| | | | | | |
|---|---|---|---|---|---|
| 2507AshInt6c | AshInt | July | Other | Other | Other |
| 2507AshInt6c | AshInt | July | Plecoptera | Chloroperlidae | Sweltsa |
| 2507AshInt6c | AshInt | July | Psocodea | Peripsocidae | Peripsocus |
| 2507AshInt6c | AshInt | July | Sarcoptiformes | Oripodidae | Other |
| 2507AshInt6c | AshInt | July | Trichoptera | Dipseudopsidae | Phylocentropus |
| 2507AshInt6c | AshInt | July | Trichoptera | Hydroptilidae | Orthotrichia |
| 2507AshInt6c | AshInt | July | Trichoptera | Leptoceridae | Oecetis |
| 2507AshInt6c | AshInt | July | Trombidiformes | Arrenuridae | Arrenurus |
| 2507AshInt6c | AshInt | July | Trombidiformes | Aturidae | Aturus |
| 2507AshInt6c | AshInt | July | Trombidiformes | Axonopsidae | Brachypoda |
| 2507AshInt6c | AshInt | July | Trombidiformes | Eupodidae | Other |
| 2507AshInt6c | AshInt | July | Trombidiformes | Feltriidae | Feltria |
| 2507AshInt6c | AshInt | July | Trombidiformes | Hydryphantidae | Hydryphantes |
| 2507AshInt6c | AshInt | July | Trombidiformes | Hydryphantidae | Parathyas |
| 2507AshInt6c | AshInt | July | Trombidiformes | Lebertiidae | Lebertia |
| 2507AshInt6c | AshInt | July | Trombidiformes | Mideopsidae | Mideopsis |
| 2507AshInt6c | AshInt | July | Trombidiformes | Pionidae | Piona |
| 2507AshInt6c | AshInt | July | Trombidiformes | Pionidae | Tiphys |
| 2507AshInt6c | AshInt | July | Trombidiformes | Sperchontidae | Sperchon |
| 2507AshInt6c | AshInt | July | Trombidiformes | Tarsonemidae | Phytonemus |
| 2507AshInt6c | AshInt | July | Trombidiformes | Torrenticolidae | Testudacarus |
| 2507AshInt6c | AshInt | July | Trombidiformes | Torrenticolidae | Torrenticola |
| 2507AshInt6c | AshInt | July | Trombidiformes | Unionicolidae | Koenikea |
| 2508AshInt6a | AshInt | August | Araneae | Agelenidae | Coras |
| 2508AshInt6a | AshInt | August | Araneae | Amaurobiidae | Callobius |
| 2508AshInt6a | AshInt | August | Araneae | Anyphaenidae | Anyphaena |
| 2508AshInt6a | AshInt | August | Araneae | Cicurinidae | Cicurina |
| 2508AshInt6a | AshInt | August | Araneae | Dictynidae | Dictyna |
| 2508AshInt6a | AshInt | August | Araneae | Gnaphosidae | Gnaphosa |
| 2508AshInt6a | AshInt | August | Araneae | Linyphiidae | Grammonota |
| 2508AshInt6a | AshInt | August | Araneae | Lycosidae | Rabidosa |

| | | | | | |
|---|---|---|---|---|---|
| 2508AshInt6a | AshInt | August | Araneae | Lycosidae | Schizocosa |
| 2508AshInt6a | AshInt | August | Araneae | Phrurolithidae | Scotinella |
| 2508AshInt6a | AshInt | August | Araneae | Salticidae | Maevia |
| 2508AshInt6a | AshInt | August | Araneae | Salticidae | Sibianor |
| 2508AshInt6a | AshInt | August | Araneae | Theridiidae | Canalidion |
| 2508AshInt6a | AshInt | August | Araneae | Theridiidae | Rugathodes |
| 2508AshInt6a | AshInt | August | Araneae | Theridiidae | Spintharus |
| 2508AshInt6a | AshInt | August | Araneae | Thomisidae | Mecaphesa |
| 2508AshInt6a | AshInt | August | Blattodea | Ectobiidae | Parcoblatta |
| 2508AshInt6a | AshInt | August | Coleoptera | Buprestidae | Pachyschelus |
| 2508AshInt6a | AshInt | August | Coleoptera | Carabidae | Apenes |
| 2508AshInt6a | AshInt | August | Coleoptera | Carabidae | Harpalus |
| 2508AshInt6a | AshInt | August | Coleoptera | Carabidae | Other |
| 2508AshInt6a | AshInt | August | Coleoptera | Carabidae | Polyderis |
| 2508AshInt6a | AshInt | August | Coleoptera | Curculionidae | Conotrachelus |
| 2508AshInt6a | AshInt | August | Coleoptera | Curculionidae | Cyrtepistomus |
| 2508AshInt6a | AshInt | August | Coleoptera | Curculionidae | Listronotus |
| 2508AshInt6a | AshInt | August | Coleoptera | Dytiscidae | Acilius |
| 2508AshInt6a | AshInt | August | Coleoptera | Lampyridae | Photinus |
| 2508AshInt6a | AshInt | August | Coleoptera | Scarabaeidae | Dialytes |
| 2508AshInt6a | AshInt | August | Coleoptera | Scarabaeidae | Melanocanthon |
| 2508AshInt6a | AshInt | August | Coleoptera | Scirtidae | Contacyphon |
| 2508AshInt6a | AshInt | August | Diptera | Agromyzidae | Calycomyza |
| 2508AshInt6a | AshInt | August | Diptera | Agromyzidae | Cerodontha |
| 2508AshInt6a | AshInt | August | Diptera | Agromyzidae | Japanagromyza |
| 2508AshInt6a | AshInt | August | Diptera | Agromyzidae | Liriomyza |
| 2508AshInt6a | AshInt | August | Diptera | Agromyzidae | Nemorimyza |
| 2508AshInt6a | AshInt | August | Diptera | Agromyzidae | Phytomyza |
| 2508AshInt6a | AshInt | August | Diptera | Anisopodidae | Mycetobia |
| 2508AshInt6a | AshInt | August | Diptera | Calliphoridae | Lucilia |
| 2508AshInt6a | AshInt | August | Diptera | Cecidomyiidae | Other |

| | | | | | |
|---|---|---|---|---|---|
| 2508AshInt6a | AshInt | August | Diptera | Chaoboridae | Chaoborus |
| 2508AshInt6a | AshInt | August | Diptera | Chaoboridae | Other |
| 2508AshInt6a | AshInt | August | Diptera | Chironomidae | Ablabesmyia |
| 2508AshInt6a | AshInt | August | Diptera | Chironomidae | Chironomus |
| 2508AshInt6a | AshInt | August | Diptera | Chironomidae | Dicrotendipes |
| 2508AshInt6a | AshInt | August | Diptera | Chironomidae | Endochironomus |
| 2508AshInt6a | AshInt | August | Diptera | Chironomidae | Glyptotendipes |
| 2508AshInt6a | AshInt | August | Diptera | Chironomidae | Microtendipes |
| 2508AshInt6a | AshInt | August | Diptera | Chironomidae | Orthocladius |
| 2508AshInt6a | AshInt | August | Diptera | Chironomidae | Other |
| 2508AshInt6a | AshInt | August | Diptera | Chironomidae | Other |
| 2508AshInt6a | AshInt | August | Diptera | Chironomidae | Other |
| 2508AshInt6a | AshInt | August | Diptera | Chironomidae | Other |
| 2508AshInt6a | AshInt | August | Diptera | Chironomidae | Parachironomus |
| 2508AshInt6a | AshInt | August | Diptera | Chironomidae | Polypedilum |
| 2508AshInt6a | AshInt | August | Diptera | Chironomidae | Pseudochironomus |
| 2508AshInt6a | AshInt | August | Diptera | Chironomidae | Tanytarsus |
| 2508AshInt6a | AshInt | August | Diptera | Chironomidae | Xenochironomus |
| 2508AshInt6a | AshInt | August | Diptera | Chloropidae | Apotropina |
| 2508AshInt6a | AshInt | August | Diptera | Chloropidae | Elachiptera |
| 2508AshInt6a | AshInt | August | Diptera | Chloropidae | Liohippelates |
| 2508AshInt6a | AshInt | August | Diptera | Chloropidae | Oscinella |
| 2508AshInt6a | AshInt | August | Diptera | Culicidae | Coquillettidia |
| 2508AshInt6a | AshInt | August | Diptera | Culicidae | Culex |
| 2508AshInt6a | AshInt | August | Diptera | Culicidae | Uranotaenia |
| 2508AshInt6a | AshInt | August | Diptera | Drosophilidae | Drosophila |
| 2508AshInt6a | AshInt | August | Diptera | Empididae | Other |
| 2508AshInt6a | AshInt | August | Diptera | Hybotidae | Platypalpus |
| 2508AshInt6a | AshInt | August | Diptera | Hybotidae | Syndyas |
| 2508AshInt6a | AshInt | August | Diptera | Keroplatidae | Other |
| 2508AshInt6a | AshInt | August | Diptera | Limoniidae | Dicranomyia |

| | | | | | |
|---|---|---|---|---|---|
| 2508AshInt6a | AshInt | August | Diptera | Limoniidae | Dicranophragma |
| 2508AshInt6a | AshInt | August | Diptera | Limoniidae | Epiphragma |
| 2508AshInt6a | AshInt | August | Diptera | Limoniidae | Erioptera |
| 2508AshInt6a | AshInt | August | Diptera | Limoniidae | Metalimnobia |
| 2508AshInt6a | AshInt | August | Diptera | Limoniidae | Ormosia |
| 2508AshInt6a | AshInt | August | Diptera | Limoniidae | Other |
| 2508AshInt6a | AshInt | August | Diptera | Limoniidae | Other |
| 2508AshInt6a | AshInt | August | Diptera | Limoniidae | Pseudolimnophila |
| 2508AshInt6a | AshInt | August | Diptera | Muscidae | Mydaea |
| 2508AshInt6a | AshInt | August | Diptera | Mycetophilidae | Mycomya |
| 2508AshInt6a | AshInt | August | Diptera | Mycetophilidae | Neoempheria |
| 2508AshInt6a | AshInt | August | Diptera | Opomyzidae | Geomyza |
| 2508AshInt6a | AshInt | August | Diptera | Pallopteridae | Toxonevra |
| 2508AshInt6a | AshInt | August | Diptera | Phoridae | Diplonevra |
| 2508AshInt6a | AshInt | August | Diptera | Polleniidae | Pollenia |
| 2508AshInt6a | AshInt | August | Diptera | Psychodidae | Clogmia |
| 2508AshInt6a | AshInt | August | Diptera | Psychodidae | Psychoda |
| 2508AshInt6a | AshInt | August | Diptera | Sarcophagidae | Ravinia |
| 2508AshInt6a | AshInt | August | Diptera | Sciaridae | Claustropyga |
| 2508AshInt6a | AshInt | August | Diptera | Sciaridae | Corynoptera |
| 2508AshInt6a | AshInt | August | Diptera | Sciaridae | Epidapus |
| 2508AshInt6a | AshInt | August | Diptera | Sciaridae | Other |
| 2508AshInt6a | AshInt | August | Diptera | Sphaeroceridae | Leptocera |
| 2508AshInt6a | AshInt | August | Diptera | Syrphidae | Chalcosyrphus |
| 2508AshInt6a | AshInt | August | Diptera | Syrphidae | Palpada |
| 2508AshInt6a | AshInt | August | Diptera | Tabanidae | Hybomitra |
| 2508AshInt6a | AshInt | August | Diptera | Tabanidae | Other |
| 2508AshInt6a | AshInt | August | Diptera | Tachinidae | Cryptomeigenia |
| 2508AshInt6a | AshInt | August | Diptera | Tachinidae | Cyzenis |
| 2508AshInt6a | AshInt | August | Diptera | Tachinidae | Zaira |
| 2508AshInt6a | AshInt | August | Diptera | Tipulidae | Dolichopeza |

| | | | | | |
|---|---|---|---|---|---|
| 2508AshInt6a | AshInt | August | Diptera | Tipulidae | Tipula |
| 2508AshInt6a | AshInt | August | Ephemeroptera | Caenidae | Caenis |
| 2508AshInt6a | AshInt | August | Ephemeroptera | Ephemeridae | Hexagenia |
| 2508AshInt6a | AshInt | August | Ephemeroptera | Heptageniidae | Epeorus |
| 2508AshInt6a | AshInt | August | Hemiptera | Clastopteridae | Clastoptera |
| 2508AshInt6a | AshInt | August | Hemiptera | Lygaeidae | Kleidocerys |
| 2508AshInt6a | AshInt | August | Hymenoptera | Apidae | Triepeolus |
| 2508AshInt6a | AshInt | August | Hymenoptera | Braconidae | Bracon |
| 2508AshInt6a | AshInt | August | Hymenoptera | Braconidae | Chorebus |
| 2508AshInt6a | AshInt | August | Hymenoptera | Ichneumonidae | Megastylus |
| 2508AshInt6a | AshInt | August | Hymenoptera | Ichneumonidae | Polyblastus |
| 2508AshInt6a | AshInt | August | Hymenoptera | Other | Other |
| 2508AshInt6a | AshInt | August | Hymenoptera | Tenthredinidae | Dolerus |
| 2508AshInt6a | AshInt | August | Hymenoptera | Tenthredinidae | Nematus |
| 2508AshInt6a | AshInt | August | Hymenoptera | Trigonalidae | Orthogonalys |
| 2508AshInt6a | AshInt | August | Lepidoptera | Crambidae | Blepharomastix |
| 2508AshInt6a | AshInt | August | Lepidoptera | Crambidae | Parapoynx |
| 2508AshInt6a | AshInt | August | Lepidoptera | Elachistidae | Elachista |
| 2508AshInt6a | AshInt | August | Lepidoptera | Erebidae | Panopoda |
| 2508AshInt6a | AshInt | August | Lepidoptera | Gelechiidae | Chionodes |
| 2508AshInt6a | AshInt | August | Lepidoptera | Noctuidae | Condica |
| 2508AshInt6a | AshInt | August | Lepidoptera | Noctuidae | Leuconycta |
| 2508AshInt6a | AshInt | August | Lepidoptera | Noctuidae | Pyreferra |
| 2508AshInt6a | AshInt | August | Lepidoptera | Notodontidae | Heterocampa |
| 2508AshInt6a | AshInt | August | Lepidoptera | Notodontidae | Hyparpax |
| 2508AshInt6a | AshInt | August | Lepidoptera | Nymphalidae | Oeneis |
| 2508AshInt6a | AshInt | August | Lepidoptera | Pyralidae | Atheloca |
| 2508AshInt6a | AshInt | August | Lepidoptera | Pyralidae | Elasmopalpus |
| 2508AshInt6a | AshInt | August | Lepidoptera | Tischeriidae | Coptotriche |
| 2508AshInt6a | AshInt | August | Lepidoptera | Tortricidae | Acleris |
| 2508AshInt6a | AshInt | August | Lepidoptera | Tortricidae | Argyrotaenia |

| | | | | | |
|---|---|---|---|---|---|
| 2508AshInt6a | AshInt | August | Lepidoptera | Tortricidae | Rhyacionia |
| 2508AshInt6a | AshInt | August | Megaloptera | Corydalidae | Corydalus |
| 2508AshInt6a | AshInt | August | Neuroptera | Hemerobiidae | Hemerobius |
| 2508AshInt6a | AshInt | August | Odonata | Libellulidae | Brachymesia |
| 2508AshInt6a | AshInt | August | Orthoptera | Tettigoniidae | Conocephalus |
| 2508AshInt6a | AshInt | August | Other | Other | Other |
| 2508AshInt6a | AshInt | August | Other | Other | Other |
| 2508AshInt6a | AshInt | August | Plecoptera | Chloroperlidae | Sweltsa |
| 2508AshInt6a | AshInt | August | Psocodea | Peripsocidae | Peripsocus |
| 2508AshInt6a | AshInt | August | Sarcoptiformes | Oripodidae | Other |
| 2508AshInt6a | AshInt | August | Trichoptera | Dipseudopsidae | Phylocentropus |
| 2508AshInt6a | AshInt | August | Trichoptera | Hydroptilidae | Orthotrichia |
| 2508AshInt6a | AshInt | August | Trichoptera | Leptoceridae | Oecetis |
| 2508AshInt6a | AshInt | August | Trombidiformes | Arrenuridae | Arrenurus |
| 2508AshInt6a | AshInt | August | Trombidiformes | Aturidae | Aturus |
| 2508AshInt6a | AshInt | August | Trombidiformes | Axonopsidae | Brachypoda |
| 2508AshInt6a | AshInt | August | Trombidiformes | Eupodidae | Other |
| 2508AshInt6a | AshInt | August | Trombidiformes | Feltriidae | Feltria |
| 2508AshInt6a | AshInt | August | Trombidiformes | Hydryphantidae | Hydryphantes |
| 2508AshInt6a | AshInt | August | Trombidiformes | Hydryphantidae | Parathyas |
| 2508AshInt6a | AshInt | August | Trombidiformes | Lebertiidae | Lebertia |
| 2508AshInt6a | AshInt | August | Trombidiformes | Mideopsidae | Mideopsis |
| 2508AshInt6a | AshInt | August | Trombidiformes | Pionidae | Piona |
| 2508AshInt6a | AshInt | August | Trombidiformes | Pionidae | Tiphys |
| 2508AshInt6a | AshInt | August | Trombidiformes | Sperchontidae | Sperchon |
| 2508AshInt6a | AshInt | August | Trombidiformes | Tarsonemidae | Phytonemus |
| 2508AshInt6a | AshInt | August | Trombidiformes | Torrenticolidae | Testudacarus |
| 2508AshInt6a | AshInt | August | Trombidiformes | Torrenticolidae | Torrenticola |
| 2508AshInt6a | AshInt | August | Trombidiformes | Unionicolidae | Koenikea |
| 2508AshInt6b | AshInt | August | Araneae | Agelenidae | Coras |
| 2508AshInt6b | AshInt | August | Araneae | Amaurobiidae | Callobius |

| | | | | | |
|---|---|---|---|---|---|
| 2508AshInt6b | AshInt | August | Araneae | Anyphaenidae | Anyphaena |
| 2508AshInt6b | AshInt | August | Araneae | Cicurinidae | Cicurina |
| 2508AshInt6b | AshInt | August | Araneae | Dictynidae | Dictyna |
| 2508AshInt6b | AshInt | August | Araneae | Gnaphosidae | Gnaphosa |
| 2508AshInt6b | AshInt | August | Araneae | Linyphiidae | Grammonota |
| 2508AshInt6b | AshInt | August | Araneae | Lycosidae | Rabidosa |
| 2508AshInt6b | AshInt | August | Araneae | Lycosidae | Schizocosa |
| 2508AshInt6b | AshInt | August | Araneae | Phrurolithidae | Scotinella |
| 2508AshInt6b | AshInt | August | Araneae | Salticidae | Maevia |
| 2508AshInt6b | AshInt | August | Araneae | Salticidae | Sibianor |
| 2508AshInt6b | AshInt | August | Araneae | Theridiidae | Canalidion |
| 2508AshInt6b | AshInt | August | Araneae | Theridiidae | Rugathodes |
| 2508AshInt6b | AshInt | August | Araneae | Theridiidae | Spintharus |
| 2508AshInt6b | AshInt | August | Araneae | Thomisidae | Mecaphesa |
| 2508AshInt6b | AshInt | August | Blattodea | Ectobiidae | Parcoblatta |
| 2508AshInt6b | AshInt | August | Coleoptera | Buprestidae | Pachyschelus |
| 2508AshInt6b | AshInt | August | Coleoptera | Carabidae | Apenes |
| 2508AshInt6b | AshInt | August | Coleoptera | Carabidae | Harpalus |
| 2508AshInt6b | AshInt | August | Coleoptera | Carabidae | Other |
| 2508AshInt6b | AshInt | August | Coleoptera | Carabidae | Polyderis |
| 2508AshInt6b | AshInt | August | Coleoptera | Curculionidae | Conotrachelus |
| 2508AshInt6b | AshInt | August | Coleoptera | Curculionidae | Cyrtepistomus |
| 2508AshInt6b | AshInt | August | Coleoptera | Curculionidae | Listronotus |
| 2508AshInt6b | AshInt | August | Coleoptera | Dytiscidae | Acilius |
| 2508AshInt6b | AshInt | August | Coleoptera | Lampyridae | Photinus |
| 2508AshInt6b | AshInt | August | Coleoptera | Scarabaeidae | Dialytes |
| 2508AshInt6b | AshInt | August | Coleoptera | Scarabaeidae | Melanocanthon |
| 2508AshInt6b | AshInt | August | Coleoptera | Scirtidae | Contacyphon |
| 2508AshInt6b | AshInt | August | Diptera | Agromyzidae | Calycomyza |
| 2508AshInt6b | AshInt | August | Diptera | Agromyzidae | Cerodontha |
| 2508AshInt6b | AshInt | August | Diptera | Agromyzidae | Japanagromyza |

| | | | | | |
|---|---|---|---|---|---|
| 2508AshInt6b | AshInt | August | Diptera | Agromyzidae | Liriomyza |
| 2508AshInt6b | AshInt | August | Diptera | Agromyzidae | Nemorimyza |
| 2508AshInt6b | AshInt | August | Diptera | Agromyzidae | Phytomyza |
| 2508AshInt6b | AshInt | August | Diptera | Anisopodidae | Mycetobia |
| 2508AshInt6b | AshInt | August | Diptera | Calliphoridae | Lucilia |
| 2508AshInt6b | AshInt | August | Diptera | Cecidomyiidae | Other |
| 2508AshInt6b | AshInt | August | Diptera | Chaoboridae | Chaoborus |
| 2508AshInt6b | AshInt | August | Diptera | Chaoboridae | Other |
| 2508AshInt6b | AshInt | August | Diptera | Chironomidae | Ablabesmyia |
| 2508AshInt6b | AshInt | August | Diptera | Chironomidae | Chironomus |
| 2508AshInt6b | AshInt | August | Diptera | Chironomidae | Dicrotendipes |
| 2508AshInt6b | AshInt | August | Diptera | Chironomidae | Endochironomus |
| 2508AshInt6b | AshInt | August | Diptera | Chironomidae | Glyptotendipes |
| 2508AshInt6b | AshInt | August | Diptera | Chironomidae | Microtendipes |
| 2508AshInt6b | AshInt | August | Diptera | Chironomidae | Orthocladius |
| 2508AshInt6b | AshInt | August | Diptera | Chironomidae | Other |
| 2508AshInt6b | AshInt | August | Diptera | Chironomidae | Other |
| 2508AshInt6b | AshInt | August | Diptera | Chironomidae | Other |
| 2508AshInt6b | AshInt | August | Diptera | Chironomidae | Other |
| 2508AshInt6b | AshInt | August | Diptera | Chironomidae | Parachironomus |
| 2508AshInt6b | AshInt | August | Diptera | Chironomidae | Polypedilum |
| 2508AshInt6b | AshInt | August | Diptera | Chironomidae | Pseudochironomus |
| 2508AshInt6b | AshInt | August | Diptera | Chironomidae | Tanytarsus |
| 2508AshInt6b | AshInt | August | Diptera | Chironomidae | Xenochironomus |
| 2508AshInt6b | AshInt | August | Diptera | Chloropidae | Apotropina |
| 2508AshInt6b | AshInt | August | Diptera | Chloropidae | Elachiptera |
| 2508AshInt6b | AshInt | August | Diptera | Chloropidae | Liohippelates |
| 2508AshInt6b | AshInt | August | Diptera | Chloropidae | Oscinella |
| 2508AshInt6b | AshInt | August | Diptera | Culicidae | Coquillettidia |
| 2508AshInt6b | AshInt | August | Diptera | Culicidae | Culex |
| 2508AshInt6b | AshInt | August | Diptera | Culicidae | Uranotaenia |

| | | | | | |
|---|---|---|---|---|---|
| 2508AshInt6b | AshInt | August | Diptera | Drosophilidae | Drosophila |
| 2508AshInt6b | AshInt | August | Diptera | Empididae | Other |
| 2508AshInt6b | AshInt | August | Diptera | Hybotidae | Platypalpus |
| 2508AshInt6b | AshInt | August | Diptera | Hybotidae | Syndyas |
| 2508AshInt6b | AshInt | August | Diptera | Keroplatidae | Other |
| 2508AshInt6b | AshInt | August | Diptera | Limoniidae | Dicranomyia |
| 2508AshInt6b | AshInt | August | Diptera | Limoniidae | Dicranophragma |
| 2508AshInt6b | AshInt | August | Diptera | Limoniidae | Epiphragma |
| 2508AshInt6b | AshInt | August | Diptera | Limoniidae | Erioptera |
| 2508AshInt6b | AshInt | August | Diptera | Limoniidae | Metalimnobia |
| 2508AshInt6b | AshInt | August | Diptera | Limoniidae | Ormosia |
| 2508AshInt6b | AshInt | August | Diptera | Limoniidae | Other |
| 2508AshInt6b | AshInt | August | Diptera | Limoniidae | Other |
| 2508AshInt6b | AshInt | August | Diptera | Limoniidae | Pseudolimnophila |
| 2508AshInt6b | AshInt | August | Diptera | Muscidae | Mydaea |
| 2508AshInt6b | AshInt | August | Diptera | Mycetophilidae | Mycomya |
| 2508AshInt6b | AshInt | August | Diptera | Mycetophilidae | Neoempheria |
| 2508AshInt6b | AshInt | August | Diptera | Opomyzidae | Geomyza |
| 2508AshInt6b | AshInt | August | Diptera | Pallopteridae | Toxonevra |
| 2508AshInt6b | AshInt | August | Diptera | Phoridae | Diplonevra |
| 2508AshInt6b | AshInt | August | Diptera | Polleniidae | Pollenia |
| 2508AshInt6b | AshInt | August | Diptera | Psychodidae | Clogmia |
| 2508AshInt6b | AshInt | August | Diptera | Psychodidae | Psychoda |
| 2508AshInt6b | AshInt | August | Diptera | Sarcophagidae | Ravinia |
| 2508AshInt6b | AshInt | August | Diptera | Sciaridae | Claustropyga |
| 2508AshInt6b | AshInt | August | Diptera | Sciaridae | Corynoptera |
| 2508AshInt6b | AshInt | August | Diptera | Sciaridae | Epidapus |
| 2508AshInt6b | AshInt | August | Diptera | Sciaridae | Other |
| 2508AshInt6b | AshInt | August | Diptera | Sphaeroceridae | Leptocera |
| 2508AshInt6b | AshInt | August | Diptera | Syrphidae | Chalcosyrphus |
| 2508AshInt6b | AshInt | August | Diptera | Syrphidae | Palpada |

| | | | | | |
|---|---|---|---|---|---|
| 2508AshInt6b | AshInt | August | Diptera | Tabanidae | Hybomitra |
| 2508AshInt6b | AshInt | August | Diptera | Tabanidae | Other |
| 2508AshInt6b | AshInt | August | Diptera | Tachinidae | Cryptomeigenia |
| 2508AshInt6b | AshInt | August | Diptera | Tachinidae | Cyzenis |
| 2508AshInt6b | AshInt | August | Diptera | Tachinidae | Zaira |
| 2508AshInt6b | AshInt | August | Diptera | Tipulidae | Dolichopeza |
| 2508AshInt6b | AshInt | August | Diptera | Tipulidae | Tipula |
| 2508AshInt6b | AshInt | August | Ephemeroptera | Caenidae | Caenis |
| 2508AshInt6b | AshInt | August | Ephemeroptera | Ephemeridae | Hexagenia |
| 2508AshInt6b | AshInt | August | Ephemeroptera | Heptageniidae | Epeorus |
| 2508AshInt6b | AshInt | August | Hemiptera | Clastopteridae | Clastoptera |
| 2508AshInt6b | AshInt | August | Hemiptera | Lygaeidae | Kleidocerys |
| 2508AshInt6b | AshInt | August | Hymenoptera | Apidae | Triepeolus |
| 2508AshInt6b | AshInt | August | Hymenoptera | Braconidae | Bracon |
| 2508AshInt6b | AshInt | August | Hymenoptera | Braconidae | Chorebus |
| 2508AshInt6b | AshInt | August | Hymenoptera | Ichneumonidae | Megastylus |
| 2508AshInt6b | AshInt | August | Hymenoptera | Ichneumonidae | Polyblastus |
| 2508AshInt6b | AshInt | August | Hymenoptera | Other | Other |
| 2508AshInt6b | AshInt | August | Hymenoptera | Tenthredinidae | Dolerus |
| 2508AshInt6b | AshInt | August | Hymenoptera | Tenthredinidae | Nematus |
| 2508AshInt6b | AshInt | August | Hymenoptera | Trigonalidae | Orthogonalys |
| 2508AshInt6b | AshInt | August | Lepidoptera | Crambidae | Blepharomastix |
| 2508AshInt6b | AshInt | August | Lepidoptera | Crambidae | Parapoynx |
| 2508AshInt6b | AshInt | August | Lepidoptera | Elachistidae | Elachista |
| 2508AshInt6b | AshInt | August | Lepidoptera | Erebidae | Panopoda |
| 2508AshInt6b | AshInt | August | Lepidoptera | Gelechiidae | Chionodes |
| 2508AshInt6b | AshInt | August | Lepidoptera | Noctuidae | Condica |
| 2508AshInt6b | AshInt | August | Lepidoptera | Noctuidae | Leuconycta |
| 2508AshInt6b | AshInt | August | Lepidoptera | Noctuidae | Pyreferra |
| 2508AshInt6b | AshInt | August | Lepidoptera | Notodontidae | Heterocampa |
| 2508AshInt6b | AshInt | August | Lepidoptera | Notodontidae | Hyparpax |

| | | | | | |
|---|---|---|---|---|---|
| 2508AshInt6b | AshInt | August | Lepidoptera | Nymphalidae | Oeneis |
| 2508AshInt6b | AshInt | August | Lepidoptera | Pyralidae | Atheloca |
| 2508AshInt6b | AshInt | August | Lepidoptera | Pyralidae | Elasmopalpus |
| 2508AshInt6b | AshInt | August | Lepidoptera | Tischeriidae | Coptotriche |
| 2508AshInt6b | AshInt | August | Lepidoptera | Tortricidae | Acleris |
| 2508AshInt6b | AshInt | August | Lepidoptera | Tortricidae | Argyrotaenia |
| 2508AshInt6b | AshInt | August | Lepidoptera | Tortricidae | Rhyacionia |
| 2508AshInt6b | AshInt | August | Megaloptera | Corydalidae | Corydalus |
| 2508AshInt6b | AshInt | August | Neuroptera | Hemerobiidae | Hemerobius |
| 2508AshInt6b | AshInt | August | Odonata | Libellulidae | Brachymesia |
| 2508AshInt6b | AshInt | August | Orthoptera | Tettigoniidae | Conocephalus |
| 2508AshInt6b | AshInt | August | Other | Other | Other |
| 2508AshInt6b | AshInt | August | Other | Other | Other |
| 2508AshInt6b | AshInt | August | Plecoptera | Chloroperlidae | Sweltsa |
| 2508AshInt6b | AshInt | August | Psocodea | Peripsocidae | Peripsocus |
| 2508AshInt6b | AshInt | August | Sarcoptiformes | Oripodidae | Other |
| 2508AshInt6b | AshInt | August | Trichoptera | Dipseudopsidae | Phylocentropus |
| 2508AshInt6b | AshInt | August | Trichoptera | Hydroptilidae | Orthotrichia |
| 2508AshInt6b | AshInt | August | Trichoptera | Leptoceridae | Oecetis |
| 2508AshInt6b | AshInt | August | Trombidiformes | Arrenuridae | Arrenurus |
| 2508AshInt6b | AshInt | August | Trombidiformes | Aturidae | Aturus |
| 2508AshInt6b | AshInt | August | Trombidiformes | Axonopsidae | Brachypoda |
| 2508AshInt6b | AshInt | August | Trombidiformes | Eupodidae | Other |
| 2508AshInt6b | AshInt | August | Trombidiformes | Feltriidae | Feltria |
| 2508AshInt6b | AshInt | August | Trombidiformes | Hydryphantidae | Hydryphantes |
| 2508AshInt6b | AshInt | August | Trombidiformes | Hydryphantidae | Parathyas |
| 2508AshInt6b | AshInt | August | Trombidiformes | Lebertiidae | Lebertia |
| 2508AshInt6b | AshInt | August | Trombidiformes | Mideopsidae | Mideopsis |
| 2508AshInt6b | AshInt | August | Trombidiformes | Pionidae | Piona |
| 2508AshInt6b | AshInt | August | Trombidiformes | Pionidae | Tiphys |
| 2508AshInt6b | AshInt | August | Trombidiformes | Sperchontidae | Sperchon |

| | | | | | |
|---|---|---|---|---|---|
| 2508AshInt6b | AshInt | August | Trombidiformes | Tarsonemidae | Phytonemus |
| 2508AshInt6b | AshInt | August | Trombidiformes | Torrenticolidae | Testudacarus |
| 2508AshInt6b | AshInt | August | Trombidiformes | Torrenticolidae | Torrenticola |
| 2508AshInt6b | AshInt | August | Trombidiformes | Unionicolidae | Koenikea |
| 2508AshInt6c | AshInt | August | Araneae | Agelenidae | Coras |
| 2508AshInt6c | AshInt | August | Araneae | Amaurobiidae | Callobius |
| 2508AshInt6c | AshInt | August | Araneae | Anyphaenidae | Anyphaena |
| 2508AshInt6c | AshInt | August | Araneae | Cicurinidae | Cicurina |
| 2508AshInt6c | AshInt | August | Araneae | Dictynidae | Dictyna |
| 2508AshInt6c | AshInt | August | Araneae | Gnaphosidae | Gnaphosa |
| 2508AshInt6c | AshInt | August | Araneae | Linyphiidae | Grammonota |
| 2508AshInt6c | AshInt | August | Araneae | Lycosidae | Rabidosa |
| 2508AshInt6c | AshInt | August | Araneae | Lycosidae | Schizocosa |
| 2508AshInt6c | AshInt | August | Araneae | Phrurolithidae | Scotinella |
| 2508AshInt6c | AshInt | August | Araneae | Salticidae | Maevia |
| 2508AshInt6c | AshInt | August | Araneae | Salticidae | Sibianor |
| 2508AshInt6c | AshInt | August | Araneae | Theridiidae | Canalidion |
| 2508AshInt6c | AshInt | August | Araneae | Theridiidae | Rugathodes |
| 2508AshInt6c | AshInt | August | Araneae | Theridiidae | Spintharus |
| 2508AshInt6c | AshInt | August | Araneae | Thomisidae | Mecaphesa |
| 2508AshInt6c | AshInt | August | Blattodea | Ectobiidae | Parcoblatta |
| 2508AshInt6c | AshInt | August | Coleoptera | Buprestidae | Pachyschelus |
| 2508AshInt6c | AshInt | August | Coleoptera | Carabidae | Apenes |
| 2508AshInt6c | AshInt | August | Coleoptera | Carabidae | Harpalus |
| 2508AshInt6c | AshInt | August | Coleoptera | Carabidae | Other |
| 2508AshInt6c | AshInt | August | Coleoptera | Carabidae | Polyderis |
| 2508AshInt6c | AshInt | August | Coleoptera | Curculionidae | Conotrachelus |
| 2508AshInt6c | AshInt | August | Coleoptera | Curculionidae | Cyrtepistomus |
| 2508AshInt6c | AshInt | August | Coleoptera | Curculionidae | Listronotus |
| 2508AshInt6c | AshInt | August | Coleoptera | Dytiscidae | Acilius |
| 2508AshInt6c | AshInt | August | Coleoptera | Lampyridae | Photinus |

| | | | | | |
|---|---|---|---|---|---|
| 2508AshInt6c | AshInt | August | Coleoptera | Scarabaeidae | Dialytes |
| 2508AshInt6c | AshInt | August | Coleoptera | Scarabaeidae | Melanocanthon |
| 2508AshInt6c | AshInt | August | Coleoptera | Scirtidae | Contacyphon |
| 2508AshInt6c | AshInt | August | Diptera | Agromyzidae | Calycomyza |
| 2508AshInt6c | AshInt | August | Diptera | Agromyzidae | Cerodontha |
| 2508AshInt6c | AshInt | August | Diptera | Agromyzidae | Japanagromyza |
| 2508AshInt6c | AshInt | August | Diptera | Agromyzidae | Liriomyza |
| 2508AshInt6c | AshInt | August | Diptera | Agromyzidae | Nemorimyza |
| 2508AshInt6c | AshInt | August | Diptera | Agromyzidae | Phytomyza |
| 2508AshInt6c | AshInt | August | Diptera | Anisopodidae | Mycetobia |
| 2508AshInt6c | AshInt | August | Diptera | Calliphoridae | Lucilia |
| 2508AshInt6c | AshInt | August | Diptera | Cecidomyiidae | Other |
| 2508AshInt6c | AshInt | August | Diptera | Chaoboridae | Chaoborus |
| 2508AshInt6c | AshInt | August | Diptera | Chaoboridae | Other |
| 2508AshInt6c | AshInt | August | Diptera | Chironomidae | Ablabesmyia |
| 2508AshInt6c | AshInt | August | Diptera | Chironomidae | Chironomus |
| 2508AshInt6c | AshInt | August | Diptera | Chironomidae | Dicrotendipes |
| 2508AshInt6c | AshInt | August | Diptera | Chironomidae | Endochironomus |
| 2508AshInt6c | AshInt | August | Diptera | Chironomidae | Glyptotendipes |
| 2508AshInt6c | AshInt | August | Diptera | Chironomidae | Microtendipes |
| 2508AshInt6c | AshInt | August | Diptera | Chironomidae | Orthocladius |
| 2508AshInt6c | AshInt | August | Diptera | Chironomidae | Other |
| 2508AshInt6c | AshInt | August | Diptera | Chironomidae | Other |
| 2508AshInt6c | AshInt | August | Diptera | Chironomidae | Other |
| 2508AshInt6c | AshInt | August | Diptera | Chironomidae | Other |
| 2508AshInt6c | AshInt | August | Diptera | Chironomidae | Parachironomus |
| 2508AshInt6c | AshInt | August | Diptera | Chironomidae | Polypedilum |
| 2508AshInt6c | AshInt | August | Diptera | Chironomidae | Pseudochironomus |
| 2508AshInt6c | AshInt | August | Diptera | Chironomidae | Tanytarsus |
| 2508AshInt6c | AshInt | August | Diptera | Chironomidae | Xenochironomus |
| 2508AshInt6c | AshInt | August | Diptera | Chloropidae | Apotropina |

| | | | | | |
|---|---|---|---|---|---|
| 2508AshInt6c | AshInt | August | Diptera | Chloropidae | Elachiptera |
| 2508AshInt6c | AshInt | August | Diptera | Chloropidae | Liohippelates |
| 2508AshInt6c | AshInt | August | Diptera | Chloropidae | Oscinella |
| 2508AshInt6c | AshInt | August | Diptera | Culicidae | Coquillettidia |
| 2508AshInt6c | AshInt | August | Diptera | Culicidae | Culex |
| 2508AshInt6c | AshInt | August | Diptera | Culicidae | Uranotaenia |
| 2508AshInt6c | AshInt | August | Diptera | Drosophilidae | Drosophila |
| 2508AshInt6c | AshInt | August | Diptera | Empididae | Other |
| 2508AshInt6c | AshInt | August | Diptera | Hybotidae | Platypalpus |
| 2508AshInt6c | AshInt | August | Diptera | Hybotidae | Syndyas |
| 2508AshInt6c | AshInt | August | Diptera | Keroplatidae | Other |
| 2508AshInt6c | AshInt | August | Diptera | Limoniidae | Dicranomyia |
| 2508AshInt6c | AshInt | August | Diptera | Limoniidae | Dicranophragma |
| 2508AshInt6c | AshInt | August | Diptera | Limoniidae | Epiphragma |
| 2508AshInt6c | AshInt | August | Diptera | Limoniidae | Erioptera |
| 2508AshInt6c | AshInt | August | Diptera | Limoniidae | Metalimnobia |
| 2508AshInt6c | AshInt | August | Diptera | Limoniidae | Ormosia |
| 2508AshInt6c | AshInt | August | Diptera | Limoniidae | Other |
| 2508AshInt6c | AshInt | August | Diptera | Limoniidae | Other |
| 2508AshInt6c | AshInt | August | Diptera | Limoniidae | Pseudolimnophila |
| 2508AshInt6c | AshInt | August | Diptera | Muscidae | Mydaea |
| 2508AshInt6c | AshInt | August | Diptera | Mycetophilidae | Mycomya |
| 2508AshInt6c | AshInt | August | Diptera | Mycetophilidae | Neoempheria |
| 2508AshInt6c | AshInt | August | Diptera | Opomyzidae | Geomyza |
| 2508AshInt6c | AshInt | August | Diptera | Pallopteridae | Toxonevra |
| 2508AshInt6c | AshInt | August | Diptera | Phoridae | Diplonevra |
| 2508AshInt6c | AshInt | August | Diptera | Polleniidae | Pollenia |
| 2508AshInt6c | AshInt | August | Diptera | Psychodidae | Clogmia |
| 2508AshInt6c | AshInt | August | Diptera | Psychodidae | Psychoda |
| 2508AshInt6c | AshInt | August | Diptera | Sarcophagidae | Ravinia |
| 2508AshInt6c | AshInt | August | Diptera | Sciaridae | Claustropyga |

| | | | | | |
|---|---|---|---|---|---|
| 2508AshInt6c | AshInt | August | Diptera | Sciaridae | Corynoptera |
| 2508AshInt6c | AshInt | August | Diptera | Sciaridae | Epidapus |
| 2508AshInt6c | AshInt | August | Diptera | Sciaridae | Other |
| 2508AshInt6c | AshInt | August | Diptera | Sphaeroceridae | Leptocera |
| 2508AshInt6c | AshInt | August | Diptera | Syrphidae | Chalcosyrphus |
| 2508AshInt6c | AshInt | August | Diptera | Syrphidae | Palpada |
| 2508AshInt6c | AshInt | August | Diptera | Tabanidae | Hybomitra |
| 2508AshInt6c | AshInt | August | Diptera | Tabanidae | Other |
| 2508AshInt6c | AshInt | August | Diptera | Tachinidae | Cryptomeigenia |
| 2508AshInt6c | AshInt | August | Diptera | Tachinidae | Cyzenis |
| 2508AshInt6c | AshInt | August | Diptera | Tachinidae | Zaira |
| 2508AshInt6c | AshInt | August | Diptera | Tipulidae | Dolichopeza |
| 2508AshInt6c | AshInt | August | Diptera | Tipulidae | Tipula |
| 2508AshInt6c | AshInt | August | Ephemeroptera | Caenidae | Caenis |
| 2508AshInt6c | AshInt | August | Ephemeroptera | Ephemeridae | Hexagenia |
| 2508AshInt6c | AshInt | August | Ephemeroptera | Heptageniidae | Epeorus |
| 2508AshInt6c | AshInt | August | Hemiptera | Clastopteridae | Clastoptera |
| 2508AshInt6c | AshInt | August | Hemiptera | Lygaeidae | Kleidocerys |
| 2508AshInt6c | AshInt | August | Hymenoptera | Apidae | Triepeolus |
| 2508AshInt6c | AshInt | August | Hymenoptera | Braconidae | Bracon |
| 2508AshInt6c | AshInt | August | Hymenoptera | Braconidae | Chorebus |
| 2508AshInt6c | AshInt | August | Hymenoptera | Ichneumonidae | Megastylus |
| 2508AshInt6c | AshInt | August | Hymenoptera | Ichneumonidae | Polyblastus |
| 2508AshInt6c | AshInt | August | Hymenoptera | Other | Other |
| 2508AshInt6c | AshInt | August | Hymenoptera | Tenthredinidae | Dolerus |
| 2508AshInt6c | AshInt | August | Hymenoptera | Tenthredinidae | Nematus |
| 2508AshInt6c | AshInt | August | Hymenoptera | Trigonalidae | Orthogonalys |
| 2508AshInt6c | AshInt | August | Lepidoptera | Crambidae | Blepharomastix |
| 2508AshInt6c | AshInt | August | Lepidoptera | Crambidae | Parapoynx |
| 2508AshInt6c | AshInt | August | Lepidoptera | Elachistidae | Elachista |
| 2508AshInt6c | AshInt | August | Lepidoptera | Erebidae | Panopoda |

| | | | | | |
|---|---|---|---|---|---|
| 2508AshInt6c | AshInt | August | Lepidoptera | Gelechiidae | Chionodes |
| 2508AshInt6c | AshInt | August | Lepidoptera | Noctuidae | Condica |
| 2508AshInt6c | AshInt | August | Lepidoptera | Noctuidae | Leuconycta |
| 2508AshInt6c | AshInt | August | Lepidoptera | Noctuidae | Pyreferra |
| 2508AshInt6c | AshInt | August | Lepidoptera | Notodontidae | Heterocampa |
| 2508AshInt6c | AshInt | August | Lepidoptera | Notodontidae | Hyparpax |
| 2508AshInt6c | AshInt | August | Lepidoptera | Nymphalidae | Oeneis |
| 2508AshInt6c | AshInt | August | Lepidoptera | Pyralidae | Atheloca |
| 2508AshInt6c | AshInt | August | Lepidoptera | Pyralidae | Elasmopalpus |
| 2508AshInt6c | AshInt | August | Lepidoptera | Tischeriidae | Coptotriche |
| 2508AshInt6c | AshInt | August | Lepidoptera | Tortricidae | Acleris |
| 2508AshInt6c | AshInt | August | Lepidoptera | Tortricidae | Argyrotaenia |
| 2508AshInt6c | AshInt | August | Lepidoptera | Tortricidae | Rhyacionia |
| 2508AshInt6c | AshInt | August | Megaloptera | Corydalidae | Corydalus |
| 2508AshInt6c | AshInt | August | Neuroptera | Hemerobiidae | Hemerobius |
| 2508AshInt6c | AshInt | August | Odonata | Libellulidae | Brachymesia |
| 2508AshInt6c | AshInt | August | Orthoptera | Tettigoniidae | Conocephalus |
| 2508AshInt6c | AshInt | August | Other | Other | Other |
| 2508AshInt6c | AshInt | August | Other | Other | Other |
| 2508AshInt6c | AshInt | August | Plecoptera | Chloroperlidae | Sweltsa |
| 2508AshInt6c | AshInt | August | Psocodea | Peripsocidae | Peripsocus |
| 2508AshInt6c | AshInt | August | Sarcoptiformes | Oripodidae | Other |
| 2508AshInt6c | AshInt | August | Trichoptera | Dipseudopsidae | Phylocentropus |
| 2508AshInt6c | AshInt | August | Trichoptera | Hydroptilidae | Orthotrichia |
| 2508AshInt6c | AshInt | August | Trichoptera | Leptoceridae | Oecetis |
| 2508AshInt6c | AshInt | August | Trombidiformes | Arrenuridae | Arrenurus |
| 2508AshInt6c | AshInt | August | Trombidiformes | Aturidae | Aturus |
| 2508AshInt6c | AshInt | August | Trombidiformes | Axonopsidae | Brachypoda |
| 2508AshInt6c | AshInt | August | Trombidiformes | Eupodidae | Other |
| 2508AshInt6c | AshInt | August | Trombidiformes | Feltriidae | Feltria |
| 2508AshInt6c | AshInt | August | Trombidiformes | Hydryphantidae | Hydryphantes |

| | | | | | |
|---|---|---|---|---|---|
| 2508AshInt6c | AshInt | August | Trombidiformes | Hydryphantidae | Parathyas |
| 2508AshInt6c | AshInt | August | Trombidiformes | Lebertiidae | Lebertia |
| 2508AshInt6c | AshInt | August | Trombidiformes | Mideopsidae | Mideopsis |
| 2508AshInt6c | AshInt | August | Trombidiformes | Pionidae | Piona |
| 2508AshInt6c | AshInt | August | Trombidiformes | Pionidae | Tiphys |
| 2508AshInt6c | AshInt | August | Trombidiformes | Sperchontidae | Sperchon |
| 2508AshInt6c | AshInt | August | Trombidiformes | Tarsonemidae | Phytonemus |
| 2508AshInt6c | AshInt | August | Trombidiformes | Torrenticolidae | Testudacarus |
| 2508AshInt6c | AshInt | August | Trombidiformes | Torrenticolidae | Torrenticola |
| 2508AshInt6c | AshInt | August | Trombidiformes | Unionicolidae | Koenikea |
| 2505AshExt6a | AshExt | May | Araneae | Agelenidae | Coras |
| 2505AshExt6a | AshExt | May | Araneae | Amaurobiidae | Callobius |
| 2505AshExt6a | AshExt | May | Araneae | Anyphaenidae | Anyphaena |
| 2505AshExt6a | AshExt | May | Araneae | Cicurinidae | Cicurina |
| 2505AshExt6a | AshExt | May | Araneae | Dictynidae | Dictyna |
| 2505AshExt6a | AshExt | May | Araneae | Gnaphosidae | Gnaphosa |
| 2505AshExt6a | AshExt | May | Araneae | Linyphiidae | Grammonota |
| 2505AshExt6a | AshExt | May | Araneae | Lycosidae | Rabidosa |
| 2505AshExt6a | AshExt | May | Araneae | Lycosidae | Schizocosa |
| 2505AshExt6a | AshExt | May | Araneae | Phrurolithidae | Scotinella |
| 2505AshExt6a | AshExt | May | Araneae | Salticidae | Maevia |
| 2505AshExt6a | AshExt | May | Araneae | Salticidae | Sibianor |
| 2505AshExt6a | AshExt | May | Araneae | Theridiidae | Canalidion |
| 2505AshExt6a | AshExt | May | Araneae | Theridiidae | Rugathodes |
| 2505AshExt6a | AshExt | May | Araneae | Theridiidae | Spintharus |
| 2505AshExt6a | AshExt | May | Araneae | Thomisidae | Mecaphesa |
| 2505AshExt6a | AshExt | May | Blattodea | Ectobiidae | Parcoblatta |
| 2505AshExt6a | AshExt | May | Coleoptera | Buprestidae | Pachyschelus |
| 2505AshExt6a | AshExt | May | Coleoptera | Carabidae | Apenes |
| 2505AshExt6a | AshExt | May | Coleoptera | Carabidae | Harpalus |
| 2505AshExt6a | AshExt | May | Coleoptera | Carabidae | Other |

| | | | | | |
|---|---|---|---|---|---|
| 2505AshExt6a | AshExt | May | Coleoptera | Carabidae | Polyderis |
| 2505AshExt6a | AshExt | May | Coleoptera | Curculionidae | Conotrachelus |
| 2505AshExt6a | AshExt | May | Coleoptera | Curculionidae | Cyrtepistomus |
| 2505AshExt6a | AshExt | May | Coleoptera | Curculionidae | Listronotus |
| 2505AshExt6a | AshExt | May | Coleoptera | Dytiscidae | Acilius |
| 2505AshExt6a | AshExt | May | Coleoptera | Lampyridae | Photinus |
| 2505AshExt6a | AshExt | May | Coleoptera | Scarabaeidae | Dialytes |
| 2505AshExt6a | AshExt | May | Coleoptera | Scarabaeidae | Melanocanthon |
| 2505AshExt6a | AshExt | May | Coleoptera | Scirtidae | Contacyphon |
| 2505AshExt6a | AshExt | May | Diptera | Agromyzidae | Calycomyza |
| 2505AshExt6a | AshExt | May | Diptera | Agromyzidae | Cerodontha |
| 2505AshExt6a | AshExt | May | Diptera | Agromyzidae | Japanagromyza |
| 2505AshExt6a | AshExt | May | Diptera | Agromyzidae | Liriomyza |
| 2505AshExt6a | AshExt | May | Diptera | Agromyzidae | Nemorimyza |
| 2505AshExt6a | AshExt | May | Diptera | Agromyzidae | Phytomyza |
| 2505AshExt6a | AshExt | May | Diptera | Anisopodidae | Mycetobia |
| 2505AshExt6a | AshExt | May | Diptera | Calliphoridae | Lucilia |
| 2505AshExt6a | AshExt | May | Diptera | Cecidomyiidae | Other |
| 2505AshExt6a | AshExt | May | Diptera | Chaoboridae | Chaoborus |
| 2505AshExt6a | AshExt | May | Diptera | Chaoboridae | Other |
| 2505AshExt6a | AshExt | May | Diptera | Chironomidae | Ablabesmyia |
| 2505AshExt6a | AshExt | May | Diptera | Chironomidae | Chironomus |
| 2505AshExt6a | AshExt | May | Diptera | Chironomidae | Dicrotendipes |
| 2505AshExt6a | AshExt | May | Diptera | Chironomidae | Endochironomus |
| 2505AshExt6a | AshExt | May | Diptera | Chironomidae | Glyptotendipes |
| 2505AshExt6a | AshExt | May | Diptera | Chironomidae | Microtendipes |
| 2505AshExt6a | AshExt | May | Diptera | Chironomidae | Orthocladius |
| 2505AshExt6a | AshExt | May | Diptera | Chironomidae | Other |
| 2505AshExt6a | AshExt | May | Diptera | Chironomidae | Other |
| 2505AshExt6a | AshExt | May | Diptera | Chironomidae | Other |
| 2505AshExt6a | AshExt | May | Diptera | Chironomidae | Other |

| | | | | | |
|---|---|---|---|---|---|
| 2505AshExt6a | AshExt | May | Diptera | Chironomidae | Parachironomus |
| 2505AshExt6a | AshExt | May | Diptera | Chironomidae | Polypedilum |
| 2505AshExt6a | AshExt | May | Diptera | Chironomidae | Pseudochironomus |
| 2505AshExt6a | AshExt | May | Diptera | Chironomidae | Tanytarsus |
| 2505AshExt6a | AshExt | May | Diptera | Chironomidae | Xenochironomus |
| 2505AshExt6a | AshExt | May | Diptera | Chloropidae | Apotropina |
| 2505AshExt6a | AshExt | May | Diptera | Chloropidae | Elachiptera |
| 2505AshExt6a | AshExt | May | Diptera | Chloropidae | Liohippelates |
| 2505AshExt6a | AshExt | May | Diptera | Chloropidae | Oscinella |
| 2505AshExt6a | AshExt | May | Diptera | Culicidae | Coquillettidia |
| 2505AshExt6a | AshExt | May | Diptera | Culicidae | Culex |
| 2505AshExt6a | AshExt | May | Diptera | Culicidae | Uranotaenia |
| 2505AshExt6a | AshExt | May | Diptera | Drosophilidae | Drosophila |
| 2505AshExt6a | AshExt | May | Diptera | Empididae | Other |
| 2505AshExt6a | AshExt | May | Diptera | Hybotidae | Platypalpus |
| 2505AshExt6a | AshExt | May | Diptera | Hybotidae | Syndyas |
| 2505AshExt6a | AshExt | May | Diptera | Keroplatidae | Other |
| 2505AshExt6a | AshExt | May | Diptera | Limoniidae | Dicranomyia |
| 2505AshExt6a | AshExt | May | Diptera | Limoniidae | Dicranophragma |
| 2505AshExt6a | AshExt | May | Diptera | Limoniidae | Epiphragma |
| 2505AshExt6a | AshExt | May | Diptera | Limoniidae | Erioptera |
| 2505AshExt6a | AshExt | May | Diptera | Limoniidae | Metalimnobia |
| 2505AshExt6a | AshExt | May | Diptera | Limoniidae | Ormosia |
| 2505AshExt6a | AshExt | May | Diptera | Limoniidae | Other |
| 2505AshExt6a | AshExt | May | Diptera | Limoniidae | Other |
| 2505AshExt6a | AshExt | May | Diptera | Limoniidae | Pseudolimnophila |
| 2505AshExt6a | AshExt | May | Diptera | Muscidae | Mydaea |
| 2505AshExt6a | AshExt | May | Diptera | Mycetophilidae | Mycomya |
| 2505AshExt6a | AshExt | May | Diptera | Mycetophilidae | Neoempheria |
| 2505AshExt6a | AshExt | May | Diptera | Opomyzidae | Geomyza |
| 2505AshExt6a | AshExt | May | Diptera | Pallopteridae | Toxonevra |

| | | | | | |
|---|---|---|---|---|---|
| 2505AshExt6a | AshExt | May | Diptera | Phoridae | Diplonevra |
| 2505AshExt6a | AshExt | May | Diptera | Polleniidae | Pollenia |
| 2505AshExt6a | AshExt | May | Diptera | Psychodidae | Clogmia |
| 2505AshExt6a | AshExt | May | Diptera | Psychodidae | Psychoda |
| 2505AshExt6a | AshExt | May | Diptera | Sarcophagidae | Ravinia |
| 2505AshExt6a | AshExt | May | Diptera | Sciaridae | Claustropyga |
| 2505AshExt6a | AshExt | May | Diptera | Sciaridae | Corynoptera |
| 2505AshExt6a | AshExt | May | Diptera | Sciaridae | Epidapus |
| 2505AshExt6a | AshExt | May | Diptera | Sciaridae | Other |
| 2505AshExt6a | AshExt | May | Diptera | Sphaeroceridae | Leptocera |
| 2505AshExt6a | AshExt | May | Diptera | Syrphidae | Chalcosyrphus |
| 2505AshExt6a | AshExt | May | Diptera | Syrphidae | Palpada |
| 2505AshExt6a | AshExt | May | Diptera | Tabanidae | Hybomitra |
| 2505AshExt6a | AshExt | May | Diptera | Tabanidae | Other |
| 2505AshExt6a | AshExt | May | Diptera | Tachinidae | Cryptomeigenia |
| 2505AshExt6a | AshExt | May | Diptera | Tachinidae | Cyzenis |
| 2505AshExt6a | AshExt | May | Diptera | Tachinidae | Zaira |
| 2505AshExt6a | AshExt | May | Diptera | Tipulidae | Dolichopeza |
| 2505AshExt6a | AshExt | May | Diptera | Tipulidae | Tipula |
| 2505AshExt6a | AshExt | May | Ephemeroptera | Caenidae | Caenis |
| 2505AshExt6a | AshExt | May | Ephemeroptera | Ephemeridae | Hexagenia |
| 2505AshExt6a | AshExt | May | Ephemeroptera | Heptageniidae | Epeorus |
| 2505AshExt6a | AshExt | May | Hemiptera | Clastopteridae | Clastoptera |
| 2505AshExt6a | AshExt | May | Hemiptera | Lygaeidae | Kleidocerys |
| 2505AshExt6a | AshExt | May | Hymenoptera | Apidae | Triepeolus |
| 2505AshExt6a | AshExt | May | Hymenoptera | Braconidae | Bracon |
| 2505AshExt6a | AshExt | May | Hymenoptera | Braconidae | Chorebus |
| 2505AshExt6a | AshExt | May | Hymenoptera | Ichneumonidae | Megastylus |
| 2505AshExt6a | AshExt | May | Hymenoptera | Ichneumonidae | Polyblastus |
| 2505AshExt6a | AshExt | May | Hymenoptera | Other | Other |
| 2505AshExt6a | AshExt | May | Hymenoptera | Tenthredinidae | Dolerus |

| | | | | | |
|---|---|---|---|---|---|
| 2505AshExt6a | AshExt | May | Hymenoptera | Tenthredinidae | Nematus |
| 2505AshExt6a | AshExt | May | Hymenoptera | Trigonalidae | Orthogonalys |
| 2505AshExt6a | AshExt | May | Lepidoptera | Crambidae | Blepharomastix |
| 2505AshExt6a | AshExt | May | Lepidoptera | Crambidae | Parapoynx |
| 2505AshExt6a | AshExt | May | Lepidoptera | Elachistidae | Elachista |
| 2505AshExt6a | AshExt | May | Lepidoptera | Erebidae | Panopoda |
| 2505AshExt6a | AshExt | May | Lepidoptera | Gelechiidae | Chionodes |
| 2505AshExt6a | AshExt | May | Lepidoptera | Noctuidae | Condica |
| 2505AshExt6a | AshExt | May | Lepidoptera | Noctuidae | Leuconycta |
| 2505AshExt6a | AshExt | May | Lepidoptera | Noctuidae | Pyreferra |
| 2505AshExt6a | AshExt | May | Lepidoptera | Notodontidae | Heterocampa |
| 2505AshExt6a | AshExt | May | Lepidoptera | Notodontidae | Hyparpax |
| 2505AshExt6a | AshExt | May | Lepidoptera | Nymphalidae | Oeneis |
| 2505AshExt6a | AshExt | May | Lepidoptera | Pyralidae | Atheloca |
| 2505AshExt6a | AshExt | May | Lepidoptera | Pyralidae | Elasmopalpus |
| 2505AshExt6a | AshExt | May | Lepidoptera | Tischeriidae | Coptotriche |
| 2505AshExt6a | AshExt | May | Lepidoptera | Tortricidae | Acleris |
| 2505AshExt6a | AshExt | May | Lepidoptera | Tortricidae | Argyrotaenia |
| 2505AshExt6a | AshExt | May | Lepidoptera | Tortricidae | Rhyacionia |
| 2505AshExt6a | AshExt | May | Megaloptera | Corydalidae | Corydalus |
| 2505AshExt6a | AshExt | May | Neuroptera | Hemerobiidae | Hemerobius |
| 2505AshExt6a | AshExt | May | Odonata | Libellulidae | Brachymesia |
| 2505AshExt6a | AshExt | May | Orthoptera | Tettigoniidae | Conocephalus |
| 2505AshExt6a | AshExt | May | Other | Other | Other |
| 2505AshExt6a | AshExt | May | Other | Other | Other |
| 2505AshExt6a | AshExt | May | Plecoptera | Chloroperlidae | Sweltsa |
| 2505AshExt6a | AshExt | May | Psocodea | Peripsocidae | Peripsocus |
| 2505AshExt6a | AshExt | May | Sarcoptiformes | Oripodidae | Other |
| 2505AshExt6a | AshExt | May | Trichoptera | Dipseudopsidae | Phylocentropus |
| 2505AshExt6a | AshExt | May | Trichoptera | Hydroptilidae | Orthotrichia |
| 2505AshExt6a | AshExt | May | Trichoptera | Leptoceridae | Oecetis |

| | | | | | |
|---|---|---|---|---|---|
| 2505AshExt6a | AshExt | May | Trombidiformes | Arrenuridae | Arrenurus |
| 2505AshExt6a | AshExt | May | Trombidiformes | Aturidae | Aturus |
| 2505AshExt6a | AshExt | May | Trombidiformes | Axonopsidae | Brachypoda |
| 2505AshExt6a | AshExt | May | Trombidiformes | Eupodidae | Other |
| 2505AshExt6a | AshExt | May | Trombidiformes | Feltriidae | Feltria |
| 2505AshExt6a | AshExt | May | Trombidiformes | Hydryphantidae | Hydryphantes |
| 2505AshExt6a | AshExt | May | Trombidiformes | Hydryphantidae | Parathyas |
| 2505AshExt6a | AshExt | May | Trombidiformes | Lebertiidae | Lebertia |
| 2505AshExt6a | AshExt | May | Trombidiformes | Mideopsidae | Mideopsis |
| 2505AshExt6a | AshExt | May | Trombidiformes | Pionidae | Piona |
| 2505AshExt6a | AshExt | May | Trombidiformes | Pionidae | Tiphys |
| 2505AshExt6a | AshExt | May | Trombidiformes | Sperchontidae | Sperchon |
| 2505AshExt6a | AshExt | May | Trombidiformes | Tarsonemidae | Phytonemus |
| 2505AshExt6a | AshExt | May | Trombidiformes | Torrenticolidae | Testudacarus |
| 2505AshExt6a | AshExt | May | Trombidiformes | Torrenticolidae | Torrenticola |
| 2505AshExt6a | AshExt | May | Trombidiformes | Unionicolidae | Koenikea |
| 2505AshExt6b | AshExt | May | Araneae | Agelenidae | Coras |
| 2505AshExt6b | AshExt | May | Araneae | Amaurobiidae | Callobius |
| 2505AshExt6b | AshExt | May | Araneae | Anyphaenidae | Anyphaena |
| 2505AshExt6b | AshExt | May | Araneae | Cicurinidae | Cicurina |
| 2505AshExt6b | AshExt | May | Araneae | Dictynidae | Dictyna |
| 2505AshExt6b | AshExt | May | Araneae | Gnaphosidae | Gnaphosa |
| 2505AshExt6b | AshExt | May | Araneae | Linyphiidae | Grammonota |
| 2505AshExt6b | AshExt | May | Araneae | Lycosidae | Rabidosa |
| 2505AshExt6b | AshExt | May | Araneae | Lycosidae | Schizocosa |
| 2505AshExt6b | AshExt | May | Araneae | Phrurolithidae | Scotinella |
| 2505AshExt6b | AshExt | May | Araneae | Salticidae | Maevia |
| 2505AshExt6b | AshExt | May | Araneae | Salticidae | Sibianor |
| 2505AshExt6b | AshExt | May | Araneae | Theridiidae | Canalidion |
| 2505AshExt6b | AshExt | May | Araneae | Theridiidae | Rugathodes |
| 2505AshExt6b | AshExt | May | Araneae | Theridiidae | Spintharus |

| | | | | | |
|---|---|---|---|---|---|
| 2505AshExt6b | AshExt | May | Araneae | Thomisidae | Mecaphesa |
| 2505AshExt6b | AshExt | May | Blattodea | Ectobiidae | Parcoblatta |
| 2505AshExt6b | AshExt | May | Coleoptera | Buprestidae | Pachyschelus |
| 2505AshExt6b | AshExt | May | Coleoptera | Carabidae | Apenes |
| 2505AshExt6b | AshExt | May | Coleoptera | Carabidae | Harpalus |
| 2505AshExt6b | AshExt | May | Coleoptera | Carabidae | Other |
| 2505AshExt6b | AshExt | May | Coleoptera | Carabidae | Polyderis |
| 2505AshExt6b | AshExt | May | Coleoptera | Curculionidae | Conotrachelus |
| 2505AshExt6b | AshExt | May | Coleoptera | Curculionidae | Cyrtepistomus |
| 2505AshExt6b | AshExt | May | Coleoptera | Curculionidae | Listronotus |
| 2505AshExt6b | AshExt | May | Coleoptera | Dytiscidae | Acilius |
| 2505AshExt6b | AshExt | May | Coleoptera | Lampyridae | Photinus |
| 2505AshExt6b | AshExt | May | Coleoptera | Scarabaeidae | Dialytes |
| 2505AshExt6b | AshExt | May | Coleoptera | Scarabaeidae | Melanocanthon |
| 2505AshExt6b | AshExt | May | Coleoptera | Scirtidae | Contacyphon |
| 2505AshExt6b | AshExt | May | Diptera | Agromyzidae | Calycomyza |
| 2505AshExt6b | AshExt | May | Diptera | Agromyzidae | Cerodontha |
| 2505AshExt6b | AshExt | May | Diptera | Agromyzidae | Japanagromyza |
| 2505AshExt6b | AshExt | May | Diptera | Agromyzidae | Liriomyza |
| 2505AshExt6b | AshExt | May | Diptera | Agromyzidae | Nemorimyza |
| 2505AshExt6b | AshExt | May | Diptera | Agromyzidae | Phytomyza |
| 2505AshExt6b | AshExt | May | Diptera | Anisopodidae | Mycetobia |
| 2505AshExt6b | AshExt | May | Diptera | Calliphoridae | Lucilia |
| 2505AshExt6b | AshExt | May | Diptera | Cecidomyiidae | Other |
| 2505AshExt6b | AshExt | May | Diptera | Chaoboridae | Chaoborus |
| 2505AshExt6b | AshExt | May | Diptera | Chaoboridae | Other |
| 2505AshExt6b | AshExt | May | Diptera | Chironomidae | Ablabesmyia |
| 2505AshExt6b | AshExt | May | Diptera | Chironomidae | Chironomus |
| 2505AshExt6b | AshExt | May | Diptera | Chironomidae | Dicrotendipes |
| 2505AshExt6b | AshExt | May | Diptera | Chironomidae | Endochironomus |
| 2505AshExt6b | AshExt | May | Diptera | Chironomidae | Glyptotendipes |

| | | | | | |
|---|---|---|---|---|---|
| 2505AshExt6b | AshExt | May | Diptera | Chironomidae | Microtendipes |
| 2505AshExt6b | AshExt | May | Diptera | Chironomidae | Orthocladius |
| 2505AshExt6b | AshExt | May | Diptera | Chironomidae | Other |
| 2505AshExt6b | AshExt | May | Diptera | Chironomidae | Other |
| 2505AshExt6b | AshExt | May | Diptera | Chironomidae | Other |
| 2505AshExt6b | AshExt | May | Diptera | Chironomidae | Other |
| 2505AshExt6b | AshExt | May | Diptera | Chironomidae | Parachironomus |
| 2505AshExt6b | AshExt | May | Diptera | Chironomidae | Polypedilum |
| 2505AshExt6b | AshExt | May | Diptera | Chironomidae | Pseudochironomus |
| 2505AshExt6b | AshExt | May | Diptera | Chironomidae | Tanytarsus |
| 2505AshExt6b | AshExt | May | Diptera | Chironomidae | Xenochironomus |
| 2505AshExt6b | AshExt | May | Diptera | Chloropidae | Apotropina |
| 2505AshExt6b | AshExt | May | Diptera | Chloropidae | Elachiptera |
| 2505AshExt6b | AshExt | May | Diptera | Chloropidae | Liohippelates |
| 2505AshExt6b | AshExt | May | Diptera | Chloropidae | Oscinella |
| 2505AshExt6b | AshExt | May | Diptera | Culicidae | Coquillettidia |
| 2505AshExt6b | AshExt | May | Diptera | Culicidae | Culex |
| 2505AshExt6b | AshExt | May | Diptera | Culicidae | Uranotaenia |
| 2505AshExt6b | AshExt | May | Diptera | Drosophilidae | Drosophila |
| 2505AshExt6b | AshExt | May | Diptera | Empididae | Other |
| 2505AshExt6b | AshExt | May | Diptera | Hybotidae | Platypalpus |
| 2505AshExt6b | AshExt | May | Diptera | Hybotidae | Syndyas |
| 2505AshExt6b | AshExt | May | Diptera | Keroplatidae | Other |
| 2505AshExt6b | AshExt | May | Diptera | Limoniidae | Dicranomyia |
| 2505AshExt6b | AshExt | May | Diptera | Limoniidae | Dicranophragma |
| 2505AshExt6b | AshExt | May | Diptera | Limoniidae | Epiphragma |
| 2505AshExt6b | AshExt | May | Diptera | Limoniidae | Erioptera |
| 2505AshExt6b | AshExt | May | Diptera | Limoniidae | Metalimnobia |
| 2505AshExt6b | AshExt | May | Diptera | Limoniidae | Ormosia |
| 2505AshExt6b | AshExt | May | Diptera | Limoniidae | Other |
| 2505AshExt6b | AshExt | May | Diptera | Limoniidae | Other |

| | | | | | |
|---|---|---|---|---|---|
| 2505AshExt6b | AshExt | May | Diptera | Limoniidae | Pseudolimnophila |
| 2505AshExt6b | AshExt | May | Diptera | Muscidae | Mydaea |
| 2505AshExt6b | AshExt | May | Diptera | Mycetophilidae | Mycomya |
| 2505AshExt6b | AshExt | May | Diptera | Mycetophilidae | Neoempheria |
| 2505AshExt6b | AshExt | May | Diptera | Opomyzidae | Geomyza |
| 2505AshExt6b | AshExt | May | Diptera | Pallopteridae | Toxonevra |
| 2505AshExt6b | AshExt | May | Diptera | Phoridae | Diplonevra |
| 2505AshExt6b | AshExt | May | Diptera | Polleniidae | Pollenia |
| 2505AshExt6b | AshExt | May | Diptera | Psychodidae | Clogmia |
| 2505AshExt6b | AshExt | May | Diptera | Psychodidae | Psychoda |
| 2505AshExt6b | AshExt | May | Diptera | Sarcophagidae | Ravinia |
| 2505AshExt6b | AshExt | May | Diptera | Sciaridae | Claustropyga |
| 2505AshExt6b | AshExt | May | Diptera | Sciaridae | Corynoptera |
| 2505AshExt6b | AshExt | May | Diptera | Sciaridae | Epidapus |
| 2505AshExt6b | AshExt | May | Diptera | Sciaridae | Other |
| 2505AshExt6b | AshExt | May | Diptera | Sphaeroceridae | Leptocera |
| 2505AshExt6b | AshExt | May | Diptera | Syrphidae | Chalcosyrphus |
| 2505AshExt6b | AshExt | May | Diptera | Syrphidae | Palpada |
| 2505AshExt6b | AshExt | May | Diptera | Tabanidae | Hybomitra |
| 2505AshExt6b | AshExt | May | Diptera | Tabanidae | Other |
| 2505AshExt6b | AshExt | May | Diptera | Tachinidae | Cryptomeigenia |
| 2505AshExt6b | AshExt | May | Diptera | Tachinidae | Cyzenis |
| 2505AshExt6b | AshExt | May | Diptera | Tachinidae | Zaira |
| 2505AshExt6b | AshExt | May | Diptera | Tipulidae | Dolichopeza |
| 2505AshExt6b | AshExt | May | Diptera | Tipulidae | Tipula |
| 2505AshExt6b | AshExt | May | Ephemeroptera | Caenidae | Caenis |
| 2505AshExt6b | AshExt | May | Ephemeroptera | Ephemeridae | Hexagenia |
| 2505AshExt6b | AshExt | May | Ephemeroptera | Heptageniidae | Epeorus |
| 2505AshExt6b | AshExt | May | Hemiptera | Clastopteridae | Clastoptera |
| 2505AshExt6b | AshExt | May | Hemiptera | Lygaeidae | Kleidocerys |
| 2505AshExt6b | AshExt | May | Hymenoptera | Apidae | Triepeolus |

| | | | | | |
|---|---|---|---|---|---|
| 2505AshExt6b | AshExt | May | Hymenoptera | Braconidae | Bracon |
| 2505AshExt6b | AshExt | May | Hymenoptera | Braconidae | Chorebus |
| 2505AshExt6b | AshExt | May | Hymenoptera | Ichneumonidae | Megastylus |
| 2505AshExt6b | AshExt | May | Hymenoptera | Ichneumonidae | Polyblastus |
| 2505AshExt6b | AshExt | May | Hymenoptera | Other | Other |
| 2505AshExt6b | AshExt | May | Hymenoptera | Tenthredinidae | Dolerus |
| 2505AshExt6b | AshExt | May | Hymenoptera | Tenthredinidae | Nematus |
| 2505AshExt6b | AshExt | May | Hymenoptera | Trigonalidae | Orthogonalys |
| 2505AshExt6b | AshExt | May | Lepidoptera | Crambidae | Blepharomastix |
| 2505AshExt6b | AshExt | May | Lepidoptera | Crambidae | Parapoynx |
| 2505AshExt6b | AshExt | May | Lepidoptera | Elachistidae | Elachista |
| 2505AshExt6b | AshExt | May | Lepidoptera | Erebidae | Panopoda |
| 2505AshExt6b | AshExt | May | Lepidoptera | Gelechiidae | Chionodes |
| 2505AshExt6b | AshExt | May | Lepidoptera | Noctuidae | Condica |
| 2505AshExt6b | AshExt | May | Lepidoptera | Noctuidae | Leuconycta |
| 2505AshExt6b | AshExt | May | Lepidoptera | Noctuidae | Pyreferra |
| 2505AshExt6b | AshExt | May | Lepidoptera | Notodontidae | Heterocampa |
| 2505AshExt6b | AshExt | May | Lepidoptera | Notodontidae | Hyparpax |
| 2505AshExt6b | AshExt | May | Lepidoptera | Nymphalidae | Oeneis |
| 2505AshExt6b | AshExt | May | Lepidoptera | Pyralidae | Atheloca |
| 2505AshExt6b | AshExt | May | Lepidoptera | Pyralidae | Elasmopalpus |
| 2505AshExt6b | AshExt | May | Lepidoptera | Tischeriidae | Coptotriche |
| 2505AshExt6b | AshExt | May | Lepidoptera | Tortricidae | Acleris |
| 2505AshExt6b | AshExt | May | Lepidoptera | Tortricidae | Argyrotaenia |
| 2505AshExt6b | AshExt | May | Lepidoptera | Tortricidae | Rhyacionia |
| 2505AshExt6b | AshExt | May | Megaloptera | Corydalidae | Corydalus |
| 2505AshExt6b | AshExt | May | Neuroptera | Hemerobiidae | Hemerobius |
| 2505AshExt6b | AshExt | May | Odonata | Libellulidae | Brachymesia |
| 2505AshExt6b | AshExt | May | Orthoptera | Tettigoniidae | Conocephalus |
| 2505AshExt6b | AshExt | May | Other | Other | Other |
| 2505AshExt6b | AshExt | May | Other | Other | Other |

| | | | | | |
|---|---|---|---|---|---|
| 2505AshExt6b | AshExt | May | Plecoptera | Chloroperlidae | Sweltsa |
| 2505AshExt6b | AshExt | May | Psocodea | Peripsocidae | Peripsocus |
| 2505AshExt6b | AshExt | May | Sarcoptiformes | Oripodidae | Other |
| 2505AshExt6b | AshExt | May | Trichoptera | Dipseudopsidae | Phylocentropus |
| 2505AshExt6b | AshExt | May | Trichoptera | Hydroptilidae | Orthotrichia |
| 2505AshExt6b | AshExt | May | Trichoptera | Leptoceridae | Oecetis |
| 2505AshExt6b | AshExt | May | Trombidiformes | Arrenuridae | Arrenurus |
| 2505AshExt6b | AshExt | May | Trombidiformes | Aturidae | Aturus |
| 2505AshExt6b | AshExt | May | Trombidiformes | Axonopsidae | Brachypoda |
| 2505AshExt6b | AshExt | May | Trombidiformes | Eupodidae | Other |
| 2505AshExt6b | AshExt | May | Trombidiformes | Feltriidae | Feltria |
| 2505AshExt6b | AshExt | May | Trombidiformes | Hydryphantidae | Hydryphantes |
| 2505AshExt6b | AshExt | May | Trombidiformes | Hydryphantidae | Parathyas |
| 2505AshExt6b | AshExt | May | Trombidiformes | Lebertiidae | Lebertia |
| 2505AshExt6b | AshExt | May | Trombidiformes | Mideopsidae | Mideopsis |
| 2505AshExt6b | AshExt | May | Trombidiformes | Pionidae | Piona |
| 2505AshExt6b | AshExt | May | Trombidiformes | Pionidae | Tiphys |
| 2505AshExt6b | AshExt | May | Trombidiformes | Sperchontidae | Sperchon |
| 2505AshExt6b | AshExt | May | Trombidiformes | Tarsonemidae | Phytonemus |
| 2505AshExt6b | AshExt | May | Trombidiformes | Torrenticolidae | Testudacarus |
| 2505AshExt6b | AshExt | May | Trombidiformes | Torrenticolidae | Torrenticola |
| 2505AshExt6b | AshExt | May | Trombidiformes | Unionicolidae | Koenikea |
| 2505AshExt6c | AshExt | May | Araneae | Agelenidae | Coras |
| 2505AshExt6c | AshExt | May | Araneae | Amaurobiidae | Callobius |
| 2505AshExt6c | AshExt | May | Araneae | Anyphaenidae | Anyphaena |
| 2505AshExt6c | AshExt | May | Araneae | Cicurinidae | Cicurina |
| 2505AshExt6c | AshExt | May | Araneae | Dictynidae | Dictyna |
| 2505AshExt6c | AshExt | May | Araneae | Gnaphosidae | Gnaphosa |
| 2505AshExt6c | AshExt | May | Araneae | Linyphiidae | Grammonota |
| 2505AshExt6c | AshExt | May | Araneae | Lycosidae | Rabidosa |
| 2505AshExt6c | AshExt | May | Araneae | Lycosidae | Schizocosa |

| | | | | | |
|---|---|---|---|---|---|
| 2505AshExt6c | AshExt | May | Araneae | Phrurolithidae | Scotinella |
| 2505AshExt6c | AshExt | May | Araneae | Salticidae | Maevia |
| 2505AshExt6c | AshExt | May | Araneae | Salticidae | Sibianor |
| 2505AshExt6c | AshExt | May | Araneae | Theridiidae | Canalidion |
| 2505AshExt6c | AshExt | May | Araneae | Theridiidae | Rugathodes |
| 2505AshExt6c | AshExt | May | Araneae | Theridiidae | Spintharus |
| 2505AshExt6c | AshExt | May | Araneae | Thomisidae | Mecaphesa |
| 2505AshExt6c | AshExt | May | Blattodea | Ectobiidae | Parcoblatta |
| 2505AshExt6c | AshExt | May | Coleoptera | Buprestidae | Pachyschelus |
| 2505AshExt6c | AshExt | May | Coleoptera | Carabidae | Apenes |
| 2505AshExt6c | AshExt | May | Coleoptera | Carabidae | Harpalus |
| 2505AshExt6c | AshExt | May | Coleoptera | Carabidae | Other |
| 2505AshExt6c | AshExt | May | Coleoptera | Carabidae | Polyderis |
| 2505AshExt6c | AshExt | May | Coleoptera | Curculionidae | Conotrachelus |
| 2505AshExt6c | AshExt | May | Coleoptera | Curculionidae | Cyrtepistomus |
| 2505AshExt6c | AshExt | May | Coleoptera | Curculionidae | Listronotus |
| 2505AshExt6c | AshExt | May | Coleoptera | Dytiscidae | Acilius |
| 2505AshExt6c | AshExt | May | Coleoptera | Lampyridae | Photinus |
| 2505AshExt6c | AshExt | May | Coleoptera | Scarabaeidae | Dialytes |
| 2505AshExt6c | AshExt | May | Coleoptera | Scarabaeidae | Melanocanthon |
| 2505AshExt6c | AshExt | May | Coleoptera | Scirtidae | Contacyphon |
| 2505AshExt6c | AshExt | May | Diptera | Agromyzidae | Calycomyza |
| 2505AshExt6c | AshExt | May | Diptera | Agromyzidae | Cerodontha |
| 2505AshExt6c | AshExt | May | Diptera | Agromyzidae | Japanagromyza |
| 2505AshExt6c | AshExt | May | Diptera | Agromyzidae | Liriomyza |
| 2505AshExt6c | AshExt | May | Diptera | Agromyzidae | Nemorimyza |
| 2505AshExt6c | AshExt | May | Diptera | Agromyzidae | Phytomyza |
| 2505AshExt6c | AshExt | May | Diptera | Anisopodidae | Mycetobia |
| 2505AshExt6c | AshExt | May | Diptera | Calliphoridae | Lucilia |
| 2505AshExt6c | AshExt | May | Diptera | Cecidomyiidae | Other |
| 2505AshExt6c | AshExt | May | Diptera | Chaoboridae | Chaoborus |

| | | | | | |
|---|---|---|---|---|---|
| 2505AshExt6c | AshExt | May | Diptera | Chaoboridae | Other |
| 2505AshExt6c | AshExt | May | Diptera | Chironomidae | Ablabesmyia |
| 2505AshExt6c | AshExt | May | Diptera | Chironomidae | Chironomus |
| 2505AshExt6c | AshExt | May | Diptera | Chironomidae | Dicrotendipes |
| 2505AshExt6c | AshExt | May | Diptera | Chironomidae | Endochironomus |
| 2505AshExt6c | AshExt | May | Diptera | Chironomidae | Glyptotendipes |
| 2505AshExt6c | AshExt | May | Diptera | Chironomidae | Microtendipes |
| 2505AshExt6c | AshExt | May | Diptera | Chironomidae | Orthocladius |
| 2505AshExt6c | AshExt | May | Diptera | Chironomidae | Other |
| 2505AshExt6c | AshExt | May | Diptera | Chironomidae | Other |
| 2505AshExt6c | AshExt | May | Diptera | Chironomidae | Other |
| 2505AshExt6c | AshExt | May | Diptera | Chironomidae | Other |
| 2505AshExt6c | AshExt | May | Diptera | Chironomidae | Parachironomus |
| 2505AshExt6c | AshExt | May | Diptera | Chironomidae | Polypedilum |
| 2505AshExt6c | AshExt | May | Diptera | Chironomidae | Pseudochironomus |
| 2505AshExt6c | AshExt | May | Diptera | Chironomidae | Tanytarsus |
| 2505AshExt6c | AshExt | May | Diptera | Chironomidae | Xenochironomus |
| 2505AshExt6c | AshExt | May | Diptera | Chloropidae | Apotropina |
| 2505AshExt6c | AshExt | May | Diptera | Chloropidae | Elachiptera |
| 2505AshExt6c | AshExt | May | Diptera | Chloropidae | Liohippelates |
| 2505AshExt6c | AshExt | May | Diptera | Chloropidae | Oscinella |
| 2505AshExt6c | AshExt | May | Diptera | Culicidae | Coquillettidia |
| 2505AshExt6c | AshExt | May | Diptera | Culicidae | Culex |
| 2505AshExt6c | AshExt | May | Diptera | Culicidae | Uranotaenia |
| 2505AshExt6c | AshExt | May | Diptera | Drosophilidae | Drosophila |
| 2505AshExt6c | AshExt | May | Diptera | Empididae | Other |
| 2505AshExt6c | AshExt | May | Diptera | Hybotidae | Platypalpus |
| 2505AshExt6c | AshExt | May | Diptera | Hybotidae | Syndyas |
| 2505AshExt6c | AshExt | May | Diptera | Keroplatidae | Other |
| 2505AshExt6c | AshExt | May | Diptera | Limoniidae | Dicranomyia |
| 2505AshExt6c | AshExt | May | Diptera | Limoniidae | Dicranophragma |

| | | | | | |
|---|---|---|---|---|---|
| 2505AshExt6c | AshExt | May | Diptera | Limoniidae | Epiphragma |
| 2505AshExt6c | AshExt | May | Diptera | Limoniidae | Erioptera |
| 2505AshExt6c | AshExt | May | Diptera | Limoniidae | Metalimnobia |
| 2505AshExt6c | AshExt | May | Diptera | Limoniidae | Ormosia |
| 2505AshExt6c | AshExt | May | Diptera | Limoniidae | Other |
| 2505AshExt6c | AshExt | May | Diptera | Limoniidae | Other |
| 2505AshExt6c | AshExt | May | Diptera | Limoniidae | Pseudolimnophila |
| 2505AshExt6c | AshExt | May | Diptera | Muscidae | Mydaea |
| 2505AshExt6c | AshExt | May | Diptera | Mycetophilidae | Mycomya |
| 2505AshExt6c | AshExt | May | Diptera | Mycetophilidae | Neoempheria |
| 2505AshExt6c | AshExt | May | Diptera | Opomyzidae | Geomyza |
| 2505AshExt6c | AshExt | May | Diptera | Pallopteridae | Toxonevra |
| 2505AshExt6c | AshExt | May | Diptera | Phoridae | Diplonevra |
| 2505AshExt6c | AshExt | May | Diptera | Polleniidae | Pollenia |
| 2505AshExt6c | AshExt | May | Diptera | Psychodidae | Clogmia |
| 2505AshExt6c | AshExt | May | Diptera | Psychodidae | Psychoda |
| 2505AshExt6c | AshExt | May | Diptera | Sarcophagidae | Ravinia |
| 2505AshExt6c | AshExt | May | Diptera | Sciaridae | Claustropyga |
| 2505AshExt6c | AshExt | May | Diptera | Sciaridae | Corynoptera |
| 2505AshExt6c | AshExt | May | Diptera | Sciaridae | Epidapus |
| 2505AshExt6c | AshExt | May | Diptera | Sciaridae | Other |
| 2505AshExt6c | AshExt | May | Diptera | Sphaeroceridae | Leptocera |
| 2505AshExt6c | AshExt | May | Diptera | Syrphidae | Chalcosyrphus |
| 2505AshExt6c | AshExt | May | Diptera | Syrphidae | Palpada |
| 2505AshExt6c | AshExt | May | Diptera | Tabanidae | Hybomitra |
| 2505AshExt6c | AshExt | May | Diptera | Tabanidae | Other |
| 2505AshExt6c | AshExt | May | Diptera | Tachinidae | Cryptomeigenia |
| 2505AshExt6c | AshExt | May | Diptera | Tachinidae | Cyzenis |
| 2505AshExt6c | AshExt | May | Diptera | Tachinidae | Zaira |
| 2505AshExt6c | AshExt | May | Diptera | Tipulidae | Dolichopeza |
| 2505AshExt6c | AshExt | May | Diptera | Tipulidae | Tipula |

| | | | | | |
|---|---|---|---|---|---|
| 2505AshExt6c | AshExt | May | Ephemeroptera | Caenidae | Caenis |
| 2505AshExt6c | AshExt | May | Ephemeroptera | Ephemeridae | Hexagenia |
| 2505AshExt6c | AshExt | May | Ephemeroptera | Heptageniidae | Epeorus |
| 2505AshExt6c | AshExt | May | Hemiptera | Clastopteridae | Clastoptera |
| 2505AshExt6c | AshExt | May | Hemiptera | Lygaeidae | Kleidocerys |
| 2505AshExt6c | AshExt | May | Hymenoptera | Apidae | Triepeolus |
| 2505AshExt6c | AshExt | May | Hymenoptera | Braconidae | Bracon |
| 2505AshExt6c | AshExt | May | Hymenoptera | Braconidae | Chorebus |
| 2505AshExt6c | AshExt | May | Hymenoptera | Ichneumonidae | Megastylus |
| 2505AshExt6c | AshExt | May | Hymenoptera | Ichneumonidae | Polyblastus |
| 2505AshExt6c | AshExt | May | Hymenoptera | Other | Other |
| 2505AshExt6c | AshExt | May | Hymenoptera | Tenthredinidae | Dolerus |
| 2505AshExt6c | AshExt | May | Hymenoptera | Tenthredinidae | Nematus |
| 2505AshExt6c | AshExt | May | Hymenoptera | Trigonalidae | Orthogonalys |
| 2505AshExt6c | AshExt | May | Lepidoptera | Crambidae | Blepharomastix |
| 2505AshExt6c | AshExt | May | Lepidoptera | Crambidae | Parapoynx |
| 2505AshExt6c | AshExt | May | Lepidoptera | Elachistidae | Elachista |
| 2505AshExt6c | AshExt | May | Lepidoptera | Erebidae | Panopoda |
| 2505AshExt6c | AshExt | May | Lepidoptera | Gelechiidae | Chionodes |
| 2505AshExt6c | AshExt | May | Lepidoptera | Noctuidae | Condica |
| 2505AshExt6c | AshExt | May | Lepidoptera | Noctuidae | Leuconycta |
| 2505AshExt6c | AshExt | May | Lepidoptera | Noctuidae | Pyreferra |
| 2505AshExt6c | AshExt | May | Lepidoptera | Notodontidae | Heterocampa |
| 2505AshExt6c | AshExt | May | Lepidoptera | Notodontidae | Hyparpax |
| 2505AshExt6c | AshExt | May | Lepidoptera | Nymphalidae | Oeneis |
| 2505AshExt6c | AshExt | May | Lepidoptera | Pyralidae | Atheloca |
| 2505AshExt6c | AshExt | May | Lepidoptera | Pyralidae | Elasmopalpus |
| 2505AshExt6c | AshExt | May | Lepidoptera | Tischeriidae | Coptotriche |
| 2505AshExt6c | AshExt | May | Lepidoptera | Tortricidae | Acleris |
| 2505AshExt6c | AshExt | May | Lepidoptera | Tortricidae | Argyrotaenia |
| 2505AshExt6c | AshExt | May | Lepidoptera | Tortricidae | Rhyacionia |

| | | | | | |
|---|---|---|---|---|---|
| 2505AshExt6c | AshExt | May | Megaloptera | Corydalidae | Corydalus |
| 2505AshExt6c | AshExt | May | Neuroptera | Hemerobiidae | Hemerobius |
| 2505AshExt6c | AshExt | May | Odonata | Libellulidae | Brachymesia |
| 2505AshExt6c | AshExt | May | Orthoptera | Tettigoniidae | Conocephalus |
| 2505AshExt6c | AshExt | May | Other | Other | Other |
| 2505AshExt6c | AshExt | May | Other | Other | Other |
| 2505AshExt6c | AshExt | May | Plecoptera | Chloroperlidae | Sweltsa |
| 2505AshExt6c | AshExt | May | Psocodea | Peripsocidae | Peripsocus |
| 2505AshExt6c | AshExt | May | Sarcoptiformes | Oripodidae | Other |
| 2505AshExt6c | AshExt | May | Trichoptera | Dipseudopsidae | Phylocentropus |
| 2505AshExt6c | AshExt | May | Trichoptera | Hydroptilidae | Orthotrichia |
| 2505AshExt6c | AshExt | May | Trichoptera | Leptoceridae | Oecetis |
| 2505AshExt6c | AshExt | May | Trombidiformes | Arrenuridae | Arrenurus |
| 2505AshExt6c | AshExt | May | Trombidiformes | Aturidae | Aturus |
| 2505AshExt6c | AshExt | May | Trombidiformes | Axonopsidae | Brachypoda |
| 2505AshExt6c | AshExt | May | Trombidiformes | Eupodidae | Other |
| 2505AshExt6c | AshExt | May | Trombidiformes | Feltriidae | Feltria |
| 2505AshExt6c | AshExt | May | Trombidiformes | Hydryphantidae | Hydryphantes |
| 2505AshExt6c | AshExt | May | Trombidiformes | Hydryphantidae | Parathyas |
| 2505AshExt6c | AshExt | May | Trombidiformes | Lebertiidae | Lebertia |
| 2505AshExt6c | AshExt | May | Trombidiformes | Mideopsidae | Mideopsis |
| 2505AshExt6c | AshExt | May | Trombidiformes | Pionidae | Piona |
| 2505AshExt6c | AshExt | May | Trombidiformes | Pionidae | Tiphys |
| 2505AshExt6c | AshExt | May | Trombidiformes | Sperchontidae | Sperchon |
| 2505AshExt6c | AshExt | May | Trombidiformes | Tarsonemidae | Phytonemus |
| 2505AshExt6c | AshExt | May | Trombidiformes | Torrenticolidae | Testudacarus |
| 2505AshExt6c | AshExt | May | Trombidiformes | Torrenticolidae | Torrenticola |
| 2505AshExt6c | AshExt | May | Trombidiformes | Unionicolidae | Koenikea |
| 2506AshExt6a | AshExt | June | Araneae | Agelenidae | Coras |
| 2506AshExt6a | AshExt | June | Araneae | Amaurobiidae | Callobius |
| 2506AshExt6a | AshExt | June | Araneae | Anyphaenidae | Anyphaena |

| | | | | | |
|---|---|---|---|---|---|
| 2506AshExt6a | AshExt | June | Araneae | Cicurinidae | Cicurina |
| 2506AshExt6a | AshExt | June | Araneae | Dictynidae | Dictyna |
| 2506AshExt6a | AshExt | June | Araneae | Gnaphosidae | Gnaphosa |
| 2506AshExt6a | AshExt | June | Araneae | Linyphiidae | Grammonota |
| 2506AshExt6a | AshExt | June | Araneae | Lycosidae | Rabidosa |
| 2506AshExt6a | AshExt | June | Araneae | Lycosidae | Schizocosa |
| 2506AshExt6a | AshExt | June | Araneae | Phrurolithidae | Scotinella |
| 2506AshExt6a | AshExt | June | Araneae | Salticidae | Maevia |
| 2506AshExt6a | AshExt | June | Araneae | Salticidae | Sibianor |
| 2506AshExt6a | AshExt | June | Araneae | Theridiidae | Canalidion |
| 2506AshExt6a | AshExt | June | Araneae | Theridiidae | Rugathodes |
| 2506AshExt6a | AshExt | June | Araneae | Theridiidae | Spintharus |
| 2506AshExt6a | AshExt | June | Araneae | Thomisidae | Mecaphesa |
| 2506AshExt6a | AshExt | June | Blattodea | Ectobiidae | Parcoblatta |
| 2506AshExt6a | AshExt | June | Coleoptera | Buprestidae | Pachyschelus |
| 2506AshExt6a | AshExt | June | Coleoptera | Carabidae | Apenes |
| 2506AshExt6a | AshExt | June | Coleoptera | Carabidae | Harpalus |
| 2506AshExt6a | AshExt | June | Coleoptera | Carabidae | Other |
| 2506AshExt6a | AshExt | June | Coleoptera | Carabidae | Polyderis |
| 2506AshExt6a | AshExt | June | Coleoptera | Curculionidae | Conotrachelus |
| 2506AshExt6a | AshExt | June | Coleoptera | Curculionidae | Cyrtepistomus |
| 2506AshExt6a | AshExt | June | Coleoptera | Curculionidae | Listronotus |
| 2506AshExt6a | AshExt | June | Coleoptera | Dytiscidae | Acilius |
| 2506AshExt6a | AshExt | June | Coleoptera | Lampyridae | Photinus |
| 2506AshExt6a | AshExt | June | Coleoptera | Scarabaeidae | Dialytes |
| 2506AshExt6a | AshExt | June | Coleoptera | Scarabaeidae | Melanocanthon |
| 2506AshExt6a | AshExt | June | Coleoptera | Scirtidae | Contacyphon |
| 2506AshExt6a | AshExt | June | Diptera | Agromyzidae | Calycomyza |
| 2506AshExt6a | AshExt | June | Diptera | Agromyzidae | Cerodontha |
| 2506AshExt6a | AshExt | June | Diptera | Agromyzidae | Japanagromyza |
| 2506AshExt6a | AshExt | June | Diptera | Agromyzidae | Liriomyza |

| | | | | | |
|---|---|---|---|---|---|
| 2506AshExt6a | AshExt | June | Diptera | Agromyzidae | Nemorimyza |
| 2506AshExt6a | AshExt | June | Diptera | Agromyzidae | Phytomyza |
| 2506AshExt6a | AshExt | June | Diptera | Anisopodidae | Mycetobia |
| 2506AshExt6a | AshExt | June | Diptera | Calliphoridae | Lucilia |
| 2506AshExt6a | AshExt | June | Diptera | Cecidomyiidae | Other |
| 2506AshExt6a | AshExt | June | Diptera | Chaoboridae | Chaoborus |
| 2506AshExt6a | AshExt | June | Diptera | Chaoboridae | Other |
| 2506AshExt6a | AshExt | June | Diptera | Chironomidae | Ablabesmyia |
| 2506AshExt6a | AshExt | June | Diptera | Chironomidae | Chironomus |
| 2506AshExt6a | AshExt | June | Diptera | Chironomidae | Dicrotendipes |
| 2506AshExt6a | AshExt | June | Diptera | Chironomidae | Endochironomus |
| 2506AshExt6a | AshExt | June | Diptera | Chironomidae | Glyptotendipes |
| 2506AshExt6a | AshExt | June | Diptera | Chironomidae | Microtendipes |
| 2506AshExt6a | AshExt | June | Diptera | Chironomidae | Orthocladius |
| 2506AshExt6a | AshExt | June | Diptera | Chironomidae | Other |
| 2506AshExt6a | AshExt | June | Diptera | Chironomidae | Other |
| 2506AshExt6a | AshExt | June | Diptera | Chironomidae | Other |
| 2506AshExt6a | AshExt | June | Diptera | Chironomidae | Other |
| 2506AshExt6a | AshExt | June | Diptera | Chironomidae | Parachironomus |
| 2506AshExt6a | AshExt | June | Diptera | Chironomidae | Polypedilum |
| 2506AshExt6a | AshExt | June | Diptera | Chironomidae | Pseudochironomus |
| 2506AshExt6a | AshExt | June | Diptera | Chironomidae | Tanytarsus |
| 2506AshExt6a | AshExt | June | Diptera | Chironomidae | Xenochironomus |
| 2506AshExt6a | AshExt | June | Diptera | Chloropidae | Apotropina |
| 2506AshExt6a | AshExt | June | Diptera | Chloropidae | Elachiptera |
| 2506AshExt6a | AshExt | June | Diptera | Chloropidae | Liohippelates |
| 2506AshExt6a | AshExt | June | Diptera | Chloropidae | Oscinella |
| 2506AshExt6a | AshExt | June | Diptera | Culicidae | Coquillettidia |
| 2506AshExt6a | AshExt | June | Diptera | Culicidae | Culex |
| 2506AshExt6a | AshExt | June | Diptera | Culicidae | Uranotaenia |
| 2506AshExt6a | AshExt | June | Diptera | Drosophilidae | Drosophila |

| | | | | | |
|---|---|---|---|---|---|
| 2506AshExt6a | AshExt | June | Diptera | Empididae | Other |
| 2506AshExt6a | AshExt | June | Diptera | Hybotidae | Platypalpus |
| 2506AshExt6a | AshExt | June | Diptera | Hybotidae | Syndyas |
| 2506AshExt6a | AshExt | June | Diptera | Keroplatidae | Other |
| 2506AshExt6a | AshExt | June | Diptera | Limoniidae | Dicranomyia |
| 2506AshExt6a | AshExt | June | Diptera | Limoniidae | Dicranophragma |
| 2506AshExt6a | AshExt | June | Diptera | Limoniidae | Epiphragma |
| 2506AshExt6a | AshExt | June | Diptera | Limoniidae | Erioptera |
| 2506AshExt6a | AshExt | June | Diptera | Limoniidae | Metalimnobia |
| 2506AshExt6a | AshExt | June | Diptera | Limoniidae | Ormosia |
| 2506AshExt6a | AshExt | June | Diptera | Limoniidae | Other |
| 2506AshExt6a | AshExt | June | Diptera | Limoniidae | Other |
| 2506AshExt6a | AshExt | June | Diptera | Limoniidae | Pseudolimnophila |
| 2506AshExt6a | AshExt | June | Diptera | Muscidae | Mydaea |
| 2506AshExt6a | AshExt | June | Diptera | Mycetophilidae | Mycomya |
| 2506AshExt6a | AshExt | June | Diptera | Mycetophilidae | Neoempheria |
| 2506AshExt6a | AshExt | June | Diptera | Opomyzidae | Geomyza |
| 2506AshExt6a | AshExt | June | Diptera | Pallopteridae | Toxonevra |
| 2506AshExt6a | AshExt | June | Diptera | Phoridae | Diplonevra |
| 2506AshExt6a | AshExt | June | Diptera | Polleniidae | Pollenia |
| 2506AshExt6a | AshExt | June | Diptera | Psychodidae | Clogmia |
| 2506AshExt6a | AshExt | June | Diptera | Psychodidae | Psychoda |
| 2506AshExt6a | AshExt | June | Diptera | Sarcophagidae | Ravinia |
| 2506AshExt6a | AshExt | June | Diptera | Sciaridae | Claustropyga |
| 2506AshExt6a | AshExt | June | Diptera | Sciaridae | Corynoptera |
| 2506AshExt6a | AshExt | June | Diptera | Sciaridae | Epidapus |
| 2506AshExt6a | AshExt | June | Diptera | Sciaridae | Other |
| 2506AshExt6a | AshExt | June | Diptera | Sphaeroceridae | Leptocera |
| 2506AshExt6a | AshExt | June | Diptera | Syrphidae | Chalcosyrphus |
| 2506AshExt6a | AshExt | June | Diptera | Syrphidae | Palpada |
| 2506AshExt6a | AshExt | June | Diptera | Tabanidae | Hybomitra |

| | | | | | |
|---|---|---|---|---|---|
| 2506AshExt6a | AshExt | June | Diptera | Tabanidae | Other |
| 2506AshExt6a | AshExt | June | Diptera | Tachinidae | Cryptomeigenia |
| 2506AshExt6a | AshExt | June | Diptera | Tachinidae | Cyzenis |
| 2506AshExt6a | AshExt | June | Diptera | Tachinidae | Zaira |
| 2506AshExt6a | AshExt | June | Diptera | Tipulidae | Dolichopeza |
| 2506AshExt6a | AshExt | June | Diptera | Tipulidae | Tipula |
| 2506AshExt6a | AshExt | June | Ephemeroptera | Caenidae | Caenis |
| 2506AshExt6a | AshExt | June | Ephemeroptera | Ephemeridae | Hexagenia |
| 2506AshExt6a | AshExt | June | Ephemeroptera | Heptageniidae | Epeorus |
| 2506AshExt6a | AshExt | June | Hemiptera | Clastopteridae | Clastoptera |
| 2506AshExt6a | AshExt | June | Hemiptera | Lygaeidae | Kleidocerys |
| 2506AshExt6a | AshExt | June | Hymenoptera | Apidae | Triepeolus |
| 2506AshExt6a | AshExt | June | Hymenoptera | Braconidae | Bracon |
| 2506AshExt6a | AshExt | June | Hymenoptera | Braconidae | Chorebus |
| 2506AshExt6a | AshExt | June | Hymenoptera | Ichneumonidae | Megastylus |
| 2506AshExt6a | AshExt | June | Hymenoptera | Ichneumonidae | Polyblastus |
| 2506AshExt6a | AshExt | June | Hymenoptera | Other | Other |
| 2506AshExt6a | AshExt | June | Hymenoptera | Tenthredinidae | Dolerus |
| 2506AshExt6a | AshExt | June | Hymenoptera | Tenthredinidae | Nematus |
| 2506AshExt6a | AshExt | June | Hymenoptera | Trigonalidae | Orthogonalys |
| 2506AshExt6a | AshExt | June | Lepidoptera | Crambidae | Blepharomastix |
| 2506AshExt6a | AshExt | June | Lepidoptera | Crambidae | Parapoynx |
| 2506AshExt6a | AshExt | June | Lepidoptera | Elachistidae | Elachista |
| 2506AshExt6a | AshExt | June | Lepidoptera | Erebidae | Panopoda |
| 2506AshExt6a | AshExt | June | Lepidoptera | Gelechiidae | Chionodes |
| 2506AshExt6a | AshExt | June | Lepidoptera | Noctuidae | Condica |
| 2506AshExt6a | AshExt | June | Lepidoptera | Noctuidae | Leuconycta |
| 2506AshExt6a | AshExt | June | Lepidoptera | Noctuidae | Pyreferra |
| 2506AshExt6a | AshExt | June | Lepidoptera | Notodontidae | Heterocampa |
| 2506AshExt6a | AshExt | June | Lepidoptera | Notodontidae | Hyparpax |
| 2506AshExt6a | AshExt | June | Lepidoptera | Nymphalidae | Oeneis |

| | | | | | |
|---|---|---|---|---|---|
| 2506AshExt6a | AshExt | June | Lepidoptera | Pyralidae | Atheloca |
| 2506AshExt6a | AshExt | June | Lepidoptera | Pyralidae | Elasmopalpus |
| 2506AshExt6a | AshExt | June | Lepidoptera | Tischeriidae | Coptotriche |
| 2506AshExt6a | AshExt | June | Lepidoptera | Tortricidae | Acleris |
| 2506AshExt6a | AshExt | June | Lepidoptera | Tortricidae | Argyrotaenia |
| 2506AshExt6a | AshExt | June | Lepidoptera | Tortricidae | Rhyacionia |
| 2506AshExt6a | AshExt | June | Megaloptera | Corydalidae | Corydalus |
| 2506AshExt6a | AshExt | June | Neuroptera | Hemerobiidae | Hemerobius |
| 2506AshExt6a | AshExt | June | Odonata | Libellulidae | Brachymesia |
| 2506AshExt6a | AshExt | June | Orthoptera | Tettigoniidae | Conocephalus |
| 2506AshExt6a | AshExt | June | Other | Other | Other |
| 2506AshExt6a | AshExt | June | Other | Other | Other |
| 2506AshExt6a | AshExt | June | Plecoptera | Chloroperlidae | Sweltsa |
| 2506AshExt6a | AshExt | June | Psocodea | Peripsocidae | Peripsocus |
| 2506AshExt6a | AshExt | June | Sarcoptiformes | Oripodidae | Other |
| 2506AshExt6a | AshExt | June | Trichoptera | Dipseudopsidae | Phylocentropus |
| 2506AshExt6a | AshExt | June | Trichoptera | Hydroptilidae | Orthotrichia |
| 2506AshExt6a | AshExt | June | Trichoptera | Leptoceridae | Oecetis |
| 2506AshExt6a | AshExt | June | Trombidiformes | Arrenuridae | Arrenurus |
| 2506AshExt6a | AshExt | June | Trombidiformes | Aturidae | Aturus |
| 2506AshExt6a | AshExt | June | Trombidiformes | Axonopsidae | Brachypoda |
| 2506AshExt6a | AshExt | June | Trombidiformes | Eupodidae | Other |
| 2506AshExt6a | AshExt | June | Trombidiformes | Feltriidae | Feltria |
| 2506AshExt6a | AshExt | June | Trombidiformes | Hydryphantidae | Hydryphantes |
| 2506AshExt6a | AshExt | June | Trombidiformes | Hydryphantidae | Parathyas |
| 2506AshExt6a | AshExt | June | Trombidiformes | Lebertiidae | Lebertia |
| 2506AshExt6a | AshExt | June | Trombidiformes | Mideopsidae | Mideopsis |
| 2506AshExt6a | AshExt | June | Trombidiformes | Pionidae | Piona |
| 2506AshExt6a | AshExt | June | Trombidiformes | Pionidae | Tiphys |
| 2506AshExt6a | AshExt | June | Trombidiformes | Sperchontidae | Sperchon |
| 2506AshExt6a | AshExt | June | Trombidiformes | Tarsonemidae | Phytonemus |

| | | | | | |
|---|---|---|---|---|---|
| 2506AshExt6a | AshExt | June | Trombidiformes | Torrenticolidae | Testudacarus |
| 2506AshExt6a | AshExt | June | Trombidiformes | Torrenticolidae | Torrenticola |
| 2506AshExt6a | AshExt | June | Trombidiformes | Unionicolidae | Koenikea |
| 2506AshExt6b | AshExt | June | Araneae | Agelenidae | Coras |
| 2506AshExt6b | AshExt | June | Araneae | Amaurobiidae | Callobius |
| 2506AshExt6b | AshExt | June | Araneae | Anyphaenidae | Anyphaena |
| 2506AshExt6b | AshExt | June | Araneae | Cicurinidae | Cicurina |
| 2506AshExt6b | AshExt | June | Araneae | Dictynidae | Dictyna |
| 2506AshExt6b | AshExt | June | Araneae | Gnaphosidae | Gnaphosa |
| 2506AshExt6b | AshExt | June | Araneae | Linyphiidae | Grammonota |
| 2506AshExt6b | AshExt | June | Araneae | Lycosidae | Rabidosa |
| 2506AshExt6b | AshExt | June | Araneae | Lycosidae | Schizocosa |
| 2506AshExt6b | AshExt | June | Araneae | Phrurolithidae | Scotinella |
| 2506AshExt6b | AshExt | June | Araneae | Salticidae | Maevia |
| 2506AshExt6b | AshExt | June | Araneae | Salticidae | Sibianor |
| 2506AshExt6b | AshExt | June | Araneae | Theridiidae | Canalidion |
| 2506AshExt6b | AshExt | June | Araneae | Theridiidae | Rugathodes |
| 2506AshExt6b | AshExt | June | Araneae | Theridiidae | Spintharus |
| 2506AshExt6b | AshExt | June | Araneae | Thomisidae | Mecaphesa |
| 2506AshExt6b | AshExt | June | Blattodea | Ectobiidae | Parcoblatta |
| 2506AshExt6b | AshExt | June | Coleoptera | Buprestidae | Pachyschelus |
| 2506AshExt6b | AshExt | June | Coleoptera | Carabidae | Apenes |
| 2506AshExt6b | AshExt | June | Coleoptera | Carabidae | Harpalus |
| 2506AshExt6b | AshExt | June | Coleoptera | Carabidae | Other |
| 2506AshExt6b | AshExt | June | Coleoptera | Carabidae | Polyderis |
| 2506AshExt6b | AshExt | June | Coleoptera | Curculionidae | Conotrachelus |
| 2506AshExt6b | AshExt | June | Coleoptera | Curculionidae | Cyrtepistomus |
| 2506AshExt6b | AshExt | June | Coleoptera | Curculionidae | Listronotus |
| 2506AshExt6b | AshExt | June | Coleoptera | Dytiscidae | Acilius |
| 2506AshExt6b | AshExt | June | Coleoptera | Lampyridae | Photinus |
| 2506AshExt6b | AshExt | June | Coleoptera | Scarabaeidae | Dialytes |

| | | | | | |
|---|---|---|---|---|---|
| 2506AshExt6b | AshExt | June | Coleoptera | Scarabaeidae | Melanocanthon |
| 2506AshExt6b | AshExt | June | Coleoptera | Scirtidae | Contacyphon |
| 2506AshExt6b | AshExt | June | Diptera | Agromyzidae | Calycomyza |
| 2506AshExt6b | AshExt | June | Diptera | Agromyzidae | Cerodontha |
| 2506AshExt6b | AshExt | June | Diptera | Agromyzidae | Japanagromyza |
| 2506AshExt6b | AshExt | June | Diptera | Agromyzidae | Liriomyza |
| 2506AshExt6b | AshExt | June | Diptera | Agromyzidae | Nemorimyza |
| 2506AshExt6b | AshExt | June | Diptera | Agromyzidae | Phytomyza |
| 2506AshExt6b | AshExt | June | Diptera | Anisopodidae | Mycetobia |
| 2506AshExt6b | AshExt | June | Diptera | Calliphoridae | Lucilia |
| 2506AshExt6b | AshExt | June | Diptera | Cecidomyiidae | Other |
| 2506AshExt6b | AshExt | June | Diptera | Chaoboridae | Chaoborus |
| 2506AshExt6b | AshExt | June | Diptera | Chaoboridae | Other |
| 2506AshExt6b | AshExt | June | Diptera | Chironomidae | Ablabesmyia |
| 2506AshExt6b | AshExt | June | Diptera | Chironomidae | Chironomus |
| 2506AshExt6b | AshExt | June | Diptera | Chironomidae | Dicrotendipes |
| 2506AshExt6b | AshExt | June | Diptera | Chironomidae | Endochironomus |
| 2506AshExt6b | AshExt | June | Diptera | Chironomidae | Glyptotendipes |
| 2506AshExt6b | AshExt | June | Diptera | Chironomidae | Microtendipes |
| 2506AshExt6b | AshExt | June | Diptera | Chironomidae | Orthocladius |
| 2506AshExt6b | AshExt | June | Diptera | Chironomidae | Other |
| 2506AshExt6b | AshExt | June | Diptera | Chironomidae | Other |
| 2506AshExt6b | AshExt | June | Diptera | Chironomidae | Other |
| 2506AshExt6b | AshExt | June | Diptera | Chironomidae | Other |
| 2506AshExt6b | AshExt | June | Diptera | Chironomidae | Parachironomus |
| 2506AshExt6b | AshExt | June | Diptera | Chironomidae | Polypedilum |
| 2506AshExt6b | AshExt | June | Diptera | Chironomidae | Pseudochironomus |
| 2506AshExt6b | AshExt | June | Diptera | Chironomidae | Tanytarsus |
| 2506AshExt6b | AshExt | June | Diptera | Chironomidae | Xenochironomus |
| 2506AshExt6b | AshExt | June | Diptera | Chloropidae | Apotropina |
| 2506AshExt6b | AshExt | June | Diptera | Chloropidae | Elachiptera |

| | | | | | |
|---|---|---|---|---|---|
| 2506AshExt6b | AshExt | June | Diptera | Chloropidae | Liohippelates |
| 2506AshExt6b | AshExt | June | Diptera | Chloropidae | Oscinella |
| 2506AshExt6b | AshExt | June | Diptera | Culicidae | Coquillettidia |
| 2506AshExt6b | AshExt | June | Diptera | Culicidae | Culex |
| 2506AshExt6b | AshExt | June | Diptera | Culicidae | Uranotaenia |
| 2506AshExt6b | AshExt | June | Diptera | Drosophilidae | Drosophila |
| 2506AshExt6b | AshExt | June | Diptera | Empididae | Other |
| 2506AshExt6b | AshExt | June | Diptera | Hybotidae | Platypalpus |
| 2506AshExt6b | AshExt | June | Diptera | Hybotidae | Syndyas |
| 2506AshExt6b | AshExt | June | Diptera | Keroplatidae | Other |
| 2506AshExt6b | AshExt | June | Diptera | Limoniidae | Dicranomyia |
| 2506AshExt6b | AshExt | June | Diptera | Limoniidae | Dicranophragma |
| 2506AshExt6b | AshExt | June | Diptera | Limoniidae | Epiphragma |
| 2506AshExt6b | AshExt | June | Diptera | Limoniidae | Erioptera |
| 2506AshExt6b | AshExt | June | Diptera | Limoniidae | Metalimnobia |
| 2506AshExt6b | AshExt | June | Diptera | Limoniidae | Ormosia |
| 2506AshExt6b | AshExt | June | Diptera | Limoniidae | Other |
| 2506AshExt6b | AshExt | June | Diptera | Limoniidae | Other |
| 2506AshExt6b | AshExt | June | Diptera | Limoniidae | Pseudolimnophila |
| 2506AshExt6b | AshExt | June | Diptera | Muscidae | Mydaea |
| 2506AshExt6b | AshExt | June | Diptera | Mycetophilidae | Mycomya |
| 2506AshExt6b | AshExt | June | Diptera | Mycetophilidae | Neoempheria |
| 2506AshExt6b | AshExt | June | Diptera | Opomyzidae | Geomyza |
| 2506AshExt6b | AshExt | June | Diptera | Pallopteridae | Toxonevra |
| 2506AshExt6b | AshExt | June | Diptera | Phoridae | Diplonevra |
| 2506AshExt6b | AshExt | June | Diptera | Polleniidae | Pollenia |
| 2506AshExt6b | AshExt | June | Diptera | Psychodidae | Clogmia |
| 2506AshExt6b | AshExt | June | Diptera | Psychodidae | Psychoda |
| 2506AshExt6b | AshExt | June | Diptera | Sarcophagidae | Ravinia |
| 2506AshExt6b | AshExt | June | Diptera | Sciaridae | Claustropyga |
| 2506AshExt6b | AshExt | June | Diptera | Sciaridae | Corynoptera |

| | | | | | |
|---|---|---|---|---|---|
| 2506AshExt6b | AshExt | June | Diptera | Sciaridae | Epidapus |
| 2506AshExt6b | AshExt | June | Diptera | Sciaridae | Other |
| 2506AshExt6b | AshExt | June | Diptera | Sphaeroceridae | Leptocera |
| 2506AshExt6b | AshExt | June | Diptera | Syrphidae | Chalcosyrphus |
| 2506AshExt6b | AshExt | June | Diptera | Syrphidae | Palpada |
| 2506AshExt6b | AshExt | June | Diptera | Tabanidae | Hybomitra |
| 2506AshExt6b | AshExt | June | Diptera | Tabanidae | Other |
| 2506AshExt6b | AshExt | June | Diptera | Tachinidae | Cryptomeigenia |
| 2506AshExt6b | AshExt | June | Diptera | Tachinidae | Cyzenis |
| 2506AshExt6b | AshExt | June | Diptera | Tachinidae | Zaira |
| 2506AshExt6b | AshExt | June | Diptera | Tipulidae | Dolichopeza |
| 2506AshExt6b | AshExt | June | Diptera | Tipulidae | Tipula |
| 2506AshExt6b | AshExt | June | Ephemeroptera | Caenidae | Caenis |
| 2506AshExt6b | AshExt | June | Ephemeroptera | Ephemeridae | Hexagenia |
| 2506AshExt6b | AshExt | June | Ephemeroptera | Heptageniidae | Epeorus |
| 2506AshExt6b | AshExt | June | Hemiptera | Clastopteridae | Clastoptera |
| 2506AshExt6b | AshExt | June | Hemiptera | Lygaeidae | Kleidocerys |
| 2506AshExt6b | AshExt | June | Hymenoptera | Apidae | Triepeolus |
| 2506AshExt6b | AshExt | June | Hymenoptera | Braconidae | Bracon |
| 2506AshExt6b | AshExt | June | Hymenoptera | Braconidae | Chorebus |
| 2506AshExt6b | AshExt | June | Hymenoptera | Ichneumonidae | Megastylus |
| 2506AshExt6b | AshExt | June | Hymenoptera | Ichneumonidae | Polyblastus |
| 2506AshExt6b | AshExt | June | Hymenoptera | Other | Other |
| 2506AshExt6b | AshExt | June | Hymenoptera | Tenthredinidae | Dolerus |
| 2506AshExt6b | AshExt | June | Hymenoptera | Tenthredinidae | Nematus |
| 2506AshExt6b | AshExt | June | Hymenoptera | Trigonalidae | Orthogonalys |
| 2506AshExt6b | AshExt | June | Lepidoptera | Crambidae | Blepharomastix |
| 2506AshExt6b | AshExt | June | Lepidoptera | Crambidae | Parapoynx |
| 2506AshExt6b | AshExt | June | Lepidoptera | Elachistidae | Elachista |
| 2506AshExt6b | AshExt | June | Lepidoptera | Erebidae | Panopoda |
| 2506AshExt6b | AshExt | June | Lepidoptera | Gelechiidae | Chionodes |

| | | | | | |
|---|---|---|---|---|---|
| 2506AshExt6b | AshExt | June | Lepidoptera | Noctuidae | Condica |
| 2506AshExt6b | AshExt | June | Lepidoptera | Noctuidae | Leuconycta |
| 2506AshExt6b | AshExt | June | Lepidoptera | Noctuidae | Pyreferra |
| 2506AshExt6b | AshExt | June | Lepidoptera | Notodontidae | Heterocampa |
| 2506AshExt6b | AshExt | June | Lepidoptera | Notodontidae | Hyparpax |
| 2506AshExt6b | AshExt | June | Lepidoptera | Nymphalidae | Oeneis |
| 2506AshExt6b | AshExt | June | Lepidoptera | Pyralidae | Atheloca |
| 2506AshExt6b | AshExt | June | Lepidoptera | Pyralidae | Elasmopalpus |
| 2506AshExt6b | AshExt | June | Lepidoptera | Tischeriidae | Coptotriche |
| 2506AshExt6b | AshExt | June | Lepidoptera | Tortricidae | Acleris |
| 2506AshExt6b | AshExt | June | Lepidoptera | Tortricidae | Argyrotaenia |
| 2506AshExt6b | AshExt | June | Lepidoptera | Tortricidae | Rhyacionia |
| 2506AshExt6b | AshExt | June | Megaloptera | Corydalidae | Corydalus |
| 2506AshExt6b | AshExt | June | Neuroptera | Hemerobiidae | Hemerobius |
| 2506AshExt6b | AshExt | June | Odonata | Libellulidae | Brachymesia |
| 2506AshExt6b | AshExt | June | Orthoptera | Tettigoniidae | Conocephalus |
| 2506AshExt6b | AshExt | June | Other | Other | Other |
| 2506AshExt6b | AshExt | June | Other | Other | Other |
| 2506AshExt6b | AshExt | June | Plecoptera | Chloroperlidae | Sweltsa |
| 2506AshExt6b | AshExt | June | Psocodea | Peripsocidae | Peripsocus |
| 2506AshExt6b | AshExt | June | Sarcoptiformes | Oripodidae | Other |
| 2506AshExt6b | AshExt | June | Trichoptera | Dipseudopsidae | Phylocentropus |
| 2506AshExt6b | AshExt | June | Trichoptera | Hydroptilidae | Orthotrichia |
| 2506AshExt6b | AshExt | June | Trichoptera | Leptoceridae | Oecetis |
| 2506AshExt6b | AshExt | June | Trombidiformes | Arrenuridae | Arrenurus |
| 2506AshExt6b | AshExt | June | Trombidiformes | Aturidae | Aturus |
| 2506AshExt6b | AshExt | June | Trombidiformes | Axonopsidae | Brachypoda |
| 2506AshExt6b | AshExt | June | Trombidiformes | Eupodidae | Other |
| 2506AshExt6b | AshExt | June | Trombidiformes | Feltriidae | Feltria |
| 2506AshExt6b | AshExt | June | Trombidiformes | Hydryphantidae | Hydryphantes |
| 2506AshExt6b | AshExt | June | Trombidiformes | Hydryphantidae | Parathyas |

| | | | | | |
|---|---|---|---|---|---|
| 2506AshExt6b | AshExt | June | Trombidiformes | Lebertiidae | Lebertia |
| 2506AshExt6b | AshExt | June | Trombidiformes | Mideopsidae | Mideopsis |
| 2506AshExt6b | AshExt | June | Trombidiformes | Pionidae | Piona |
| 2506AshExt6b | AshExt | June | Trombidiformes | Pionidae | Tiphys |
| 2506AshExt6b | AshExt | June | Trombidiformes | Sperchontidae | Sperchon |
| 2506AshExt6b | AshExt | June | Trombidiformes | Tarsonemidae | Phytonemus |
| 2506AshExt6b | AshExt | June | Trombidiformes | Torrenticolidae | Testudacarus |
| 2506AshExt6b | AshExt | June | Trombidiformes | Torrenticolidae | Torrenticola |
| 2506AshExt6b | AshExt | June | Trombidiformes | Unionicolidae | Koenikea |
| 2506AshExt6c | AshExt | June | Araneae | Agelenidae | Coras |
| 2506AshExt6c | AshExt | June | Araneae | Amaurobiidae | Callobius |
| 2506AshExt6c | AshExt | June | Araneae | Anyphaenidae | Anyphaena |
| 2506AshExt6c | AshExt | June | Araneae | Cicurinidae | Cicurina |
| 2506AshExt6c | AshExt | June | Araneae | Dictynidae | Dictyna |
| 2506AshExt6c | AshExt | June | Araneae | Gnaphosidae | Gnaphosa |
| 2506AshExt6c | AshExt | June | Araneae | Linyphiidae | Grammonota |
| 2506AshExt6c | AshExt | June | Araneae | Lycosidae | Rabidosa |
| 2506AshExt6c | AshExt | June | Araneae | Lycosidae | Schizocosa |
| 2506AshExt6c | AshExt | June | Araneae | Phrurolithidae | Scotinella |
| 2506AshExt6c | AshExt | June | Araneae | Salticidae | Maevia |
| 2506AshExt6c | AshExt | June | Araneae | Salticidae | Sibianor |
| 2506AshExt6c | AshExt | June | Araneae | Theridiidae | Canalidion |
| 2506AshExt6c | AshExt | June | Araneae | Theridiidae | Rugathodes |
| 2506AshExt6c | AshExt | June | Araneae | Theridiidae | Spintharus |
| 2506AshExt6c | AshExt | June | Araneae | Thomisidae | Mecaphesa |
| 2506AshExt6c | AshExt | June | Blattodea | Ectobiidae | Parcoblatta |
| 2506AshExt6c | AshExt | June | Coleoptera | Buprestidae | Pachyschelus |
| 2506AshExt6c | AshExt | June | Coleoptera | Carabidae | Apenes |
| 2506AshExt6c | AshExt | June | Coleoptera | Carabidae | Harpalus |
| 2506AshExt6c | AshExt | June | Coleoptera | Carabidae | Other |
| 2506AshExt6c | AshExt | June | Coleoptera | Carabidae | Polyderis |

| | | | | | |
|---|---|---|---|---|---|
| 2506AshExt6c | AshExt | June | Coleoptera | Curculionidae | Conotrachelus |
| 2506AshExt6c | AshExt | June | Coleoptera | Curculionidae | Cyrtepistomus |
| 2506AshExt6c | AshExt | June | Coleoptera | Curculionidae | Listronotus |
| 2506AshExt6c | AshExt | June | Coleoptera | Dytiscidae | Acilius |
| 2506AshExt6c | AshExt | June | Coleoptera | Lampyridae | Photinus |
| 2506AshExt6c | AshExt | June | Coleoptera | Scarabaeidae | Dialytes |
| 2506AshExt6c | AshExt | June | Coleoptera | Scarabaeidae | Melanocanthon |
| 2506AshExt6c | AshExt | June | Coleoptera | Scirtidae | Contacyphon |
| 2506AshExt6c | AshExt | June | Diptera | Agromyzidae | Calycomyza |
| 2506AshExt6c | AshExt | June | Diptera | Agromyzidae | Cerodontha |
| 2506AshExt6c | AshExt | June | Diptera | Agromyzidae | Japanagromyza |
| 2506AshExt6c | AshExt | June | Diptera | Agromyzidae | Liriomyza |
| 2506AshExt6c | AshExt | June | Diptera | Agromyzidae | Nemorimyza |
| 2506AshExt6c | AshExt | June | Diptera | Agromyzidae | Phytomyza |
| 2506AshExt6c | AshExt | June | Diptera | Anisopodidae | Mycetobia |
| 2506AshExt6c | AshExt | June | Diptera | Calliphoridae | Lucilia |
| 2506AshExt6c | AshExt | June | Diptera | Cecidomyiidae | Other |
| 2506AshExt6c | AshExt | June | Diptera | Chaoboridae | Chaoborus |
| 2506AshExt6c | AshExt | June | Diptera | Chaoboridae | Other |
| 2506AshExt6c | AshExt | June | Diptera | Chironomidae | Ablabesmyia |
| 2506AshExt6c | AshExt | June | Diptera | Chironomidae | Chironomus |
| 2506AshExt6c | AshExt | June | Diptera | Chironomidae | Dicrotendipes |
| 2506AshExt6c | AshExt | June | Diptera | Chironomidae | Endochironomus |
| 2506AshExt6c | AshExt | June | Diptera | Chironomidae | Glyptotendipes |
| 2506AshExt6c | AshExt | June | Diptera | Chironomidae | Microtendipes |
| 2506AshExt6c | AshExt | June | Diptera | Chironomidae | Orthocladius |
| 2506AshExt6c | AshExt | June | Diptera | Chironomidae | Other |
| 2506AshExt6c | AshExt | June | Diptera | Chironomidae | Other |
| 2506AshExt6c | AshExt | June | Diptera | Chironomidae | Other |
| 2506AshExt6c | AshExt | June | Diptera | Chironomidae | Other |
| 2506AshExt6c | AshExt | June | Diptera | Chironomidae | Parachironomus |

| | | | | | |
|---|---|---|---|---|---|
| 2506AshExt6c | AshExt | June | Diptera | Chironomidae | Polypedilum |
| 2506AshExt6c | AshExt | June | Diptera | Chironomidae | Pseudochironomus |
| 2506AshExt6c | AshExt | June | Diptera | Chironomidae | Tanytarsus |
| 2506AshExt6c | AshExt | June | Diptera | Chironomidae | Xenochironomus |
| 2506AshExt6c | AshExt | June | Diptera | Chloropidae | Apotropina |
| 2506AshExt6c | AshExt | June | Diptera | Chloropidae | Elachiptera |
| 2506AshExt6c | AshExt | June | Diptera | Chloropidae | Liohippelates |
| 2506AshExt6c | AshExt | June | Diptera | Chloropidae | Oscinella |
| 2506AshExt6c | AshExt | June | Diptera | Culicidae | Coquillettidia |
| 2506AshExt6c | AshExt | June | Diptera | Culicidae | Culex |
| 2506AshExt6c | AshExt | June | Diptera | Culicidae | Uranotaenia |
| 2506AshExt6c | AshExt | June | Diptera | Drosophilidae | Drosophila |
| 2506AshExt6c | AshExt | June | Diptera | Empididae | Other |
| 2506AshExt6c | AshExt | June | Diptera | Hybotidae | Platypalpus |
| 2506AshExt6c | AshExt | June | Diptera | Hybotidae | Syndyas |
| 2506AshExt6c | AshExt | June | Diptera | Keroplatidae | Other |
| 2506AshExt6c | AshExt | June | Diptera | Limoniidae | Dicranomyia |
| 2506AshExt6c | AshExt | June | Diptera | Limoniidae | Dicranophragma |
| 2506AshExt6c | AshExt | June | Diptera | Limoniidae | Epiphragma |
| 2506AshExt6c | AshExt | June | Diptera | Limoniidae | Erioptera |
| 2506AshExt6c | AshExt | June | Diptera | Limoniidae | Metalimnobia |
| 2506AshExt6c | AshExt | June | Diptera | Limoniidae | Ormosia |
| 2506AshExt6c | AshExt | June | Diptera | Limoniidae | Other |
| 2506AshExt6c | AshExt | June | Diptera | Limoniidae | Other |
| 2506AshExt6c | AshExt | June | Diptera | Limoniidae | Pseudolimnophila |
| 2506AshExt6c | AshExt | June | Diptera | Muscidae | Mydaea |
| 2506AshExt6c | AshExt | June | Diptera | Mycetophilidae | Mycomya |
| 2506AshExt6c | AshExt | June | Diptera | Mycetophilidae | Neoempheria |
| 2506AshExt6c | AshExt | June | Diptera | Opomyzidae | Geomyza |
| 2506AshExt6c | AshExt | June | Diptera | Pallopteridae | Toxonevra |
| 2506AshExt6c | AshExt | June | Diptera | Phoridae | Diplonevra |

| | | | | | |
|---|---|---|---|---|---|
| 2506AshExt6c | AshExt | June | Diptera | Polleniidae | Pollenia |
| 2506AshExt6c | AshExt | June | Diptera | Psychodidae | Clogmia |
| 2506AshExt6c | AshExt | June | Diptera | Psychodidae | Psychoda |
| 2506AshExt6c | AshExt | June | Diptera | Sarcophagidae | Ravinia |
| 2506AshExt6c | AshExt | June | Diptera | Sciaridae | Claustropyga |
| 2506AshExt6c | AshExt | June | Diptera | Sciaridae | Corynoptera |
| 2506AshExt6c | AshExt | June | Diptera | Sciaridae | Epidapus |
| 2506AshExt6c | AshExt | June | Diptera | Sciaridae | Other |
| 2506AshExt6c | AshExt | June | Diptera | Sphaeroceridae | Leptocera |
| 2506AshExt6c | AshExt | June | Diptera | Syrphidae | Chalcosyrphus |
| 2506AshExt6c | AshExt | June | Diptera | Syrphidae | Palpada |
| 2506AshExt6c | AshExt | June | Diptera | Tabanidae | Hybomitra |
| 2506AshExt6c | AshExt | June | Diptera | Tabanidae | Other |
| 2506AshExt6c | AshExt | June | Diptera | Tachinidae | Cryptomeigenia |
| 2506AshExt6c | AshExt | June | Diptera | Tachinidae | Cyzenis |
| 2506AshExt6c | AshExt | June | Diptera | Tachinidae | Zaira |
| 2506AshExt6c | AshExt | June | Diptera | Tipulidae | Dolichopeza |
| 2506AshExt6c | AshExt | June | Diptera | Tipulidae | Tipula |
| 2506AshExt6c | AshExt | June | Ephemeroptera | Caenidae | Caenis |
| 2506AshExt6c | AshExt | June | Ephemeroptera | Ephemeridae | Hexagenia |
| 2506AshExt6c | AshExt | June | Ephemeroptera | Heptageniidae | Epeorus |
| 2506AshExt6c | AshExt | June | Hemiptera | Clastopteridae | Clastoptera |
| 2506AshExt6c | AshExt | June | Hemiptera | Lygaeidae | Kleidocerys |
| 2506AshExt6c | AshExt | June | Hymenoptera | Apidae | Triepeolus |
| 2506AshExt6c | AshExt | June | Hymenoptera | Braconidae | Bracon |
| 2506AshExt6c | AshExt | June | Hymenoptera | Braconidae | Chorebus |
| 2506AshExt6c | AshExt | June | Hymenoptera | Ichneumonidae | Megastylus |
| 2506AshExt6c | AshExt | June | Hymenoptera | Ichneumonidae | Polyblastus |
| 2506AshExt6c | AshExt | June | Hymenoptera | Other | Other |
| 2506AshExt6c | AshExt | June | Hymenoptera | Tenthredinidae | Dolerus |
| 2506AshExt6c | AshExt | June | Hymenoptera | Tenthredinidae | Nematus |

| | | | | | |
|---|---|---|---|---|---|
| 2506AshExt6c | AshExt | June | Hymenoptera | Trigonalidae | Orthogonalys |
| 2506AshExt6c | AshExt | June | Lepidoptera | Crambidae | Blepharomastix |
| 2506AshExt6c | AshExt | June | Lepidoptera | Crambidae | Parapoynx |
| 2506AshExt6c | AshExt | June | Lepidoptera | Elachistidae | Elachista |
| 2506AshExt6c | AshExt | June | Lepidoptera | Erebidae | Panopoda |
| 2506AshExt6c | AshExt | June | Lepidoptera | Gelechiidae | Chionodes |
| 2506AshExt6c | AshExt | June | Lepidoptera | Noctuidae | Condica |
| 2506AshExt6c | AshExt | June | Lepidoptera | Noctuidae | Leuconycta |
| 2506AshExt6c | AshExt | June | Lepidoptera | Noctuidae | Pyreferra |
| 2506AshExt6c | AshExt | June | Lepidoptera | Notodontidae | Heterocampa |
| 2506AshExt6c | AshExt | June | Lepidoptera | Notodontidae | Hyparpax |
| 2506AshExt6c | AshExt | June | Lepidoptera | Nymphalidae | Oeneis |
| 2506AshExt6c | AshExt | June | Lepidoptera | Pyralidae | Atheloca |
| 2506AshExt6c | AshExt | June | Lepidoptera | Pyralidae | Elasmopalpus |
| 2506AshExt6c | AshExt | June | Lepidoptera | Tischeriidae | Coptotriche |
| 2506AshExt6c | AshExt | June | Lepidoptera | Tortricidae | Acleris |
| 2506AshExt6c | AshExt | June | Lepidoptera | Tortricidae | Argyrotaenia |
| 2506AshExt6c | AshExt | June | Lepidoptera | Tortricidae | Rhyacionia |
| 2506AshExt6c | AshExt | June | Megaloptera | Corydalidae | Corydalus |
| 2506AshExt6c | AshExt | June | Neuroptera | Hemerobiidae | Hemerobius |
| 2506AshExt6c | AshExt | June | Odonata | Libellulidae | Brachymesia |
| 2506AshExt6c | AshExt | June | Orthoptera | Tettigoniidae | Conocephalus |
| 2506AshExt6c | AshExt | June | Other | Other | Other |
| 2506AshExt6c | AshExt | June | Other | Other | Other |
| 2506AshExt6c | AshExt | June | Plecoptera | Chloroperlidae | Sweltsa |
| 2506AshExt6c | AshExt | June | Psocodea | Peripsocidae | Peripsocus |
| 2506AshExt6c | AshExt | June | Sarcoptiformes | Oripodidae | Other |
| 2506AshExt6c | AshExt | June | Trichoptera | Dipseudopsidae | Phylocentropus |
| 2506AshExt6c | AshExt | June | Trichoptera | Hydroptilidae | Orthotrichia |
| 2506AshExt6c | AshExt | June | Trichoptera | Leptoceridae | Oecetis |
| 2506AshExt6c | AshExt | June | Trombidiformes | Arrenuridae | Arrenurus |

| | | | | | |
|---|---|---|---|---|---|
| 2506AshExt6c | AshExt | June | Trombidiformes | Aturidae | Aturus |
| 2506AshExt6c | AshExt | June | Trombidiformes | Axonopsidae | Brachypoda |
| 2506AshExt6c | AshExt | June | Trombidiformes | Eupodidae | Other |
| 2506AshExt6c | AshExt | June | Trombidiformes | Feltriidae | Feltria |
| 2506AshExt6c | AshExt | June | Trombidiformes | Hydryphantidae | Hydryphantes |
| 2506AshExt6c | AshExt | June | Trombidiformes | Hydryphantidae | Parathyas |
| 2506AshExt6c | AshExt | June | Trombidiformes | Lebertiidae | Lebertia |
| 2506AshExt6c | AshExt | June | Trombidiformes | Mideopsidae | Mideopsis |
| 2506AshExt6c | AshExt | June | Trombidiformes | Pionidae | Piona |
| 2506AshExt6c | AshExt | June | Trombidiformes | Pionidae | Tiphys |
| 2506AshExt6c | AshExt | June | Trombidiformes | Sperchontidae | Sperchon |
| 2506AshExt6c | AshExt | June | Trombidiformes | Tarsonemidae | Phytonemus |
| 2506AshExt6c | AshExt | June | Trombidiformes | Torrenticolidae | Testudacarus |
| 2506AshExt6c | AshExt | June | Trombidiformes | Torrenticolidae | Torrenticola |
| 2506AshExt6c | AshExt | June | Trombidiformes | Unionicolidae | Koenikea |
| 2507AshExt6a | AshExt | July | Araneae | Agelenidae | Coras |
| 2507AshExt6a | AshExt | July | Araneae | Amaurobiidae | Callobius |
| 2507AshExt6a | AshExt | July | Araneae | Anyphaenidae | Anyphaena |
| 2507AshExt6a | AshExt | July | Araneae | Cicurinidae | Cicurina |
| 2507AshExt6a | AshExt | July | Araneae | Dictynidae | Dictyna |
| 2507AshExt6a | AshExt | July | Araneae | Gnaphosidae | Gnaphosa |
| 2507AshExt6a | AshExt | July | Araneae | Linyphiidae | Grammonota |
| 2507AshExt6a | AshExt | July | Araneae | Lycosidae | Rabidosa |
| 2507AshExt6a | AshExt | July | Araneae | Lycosidae | Schizocosa |
| 2507AshExt6a | AshExt | July | Araneae | Phrurolithidae | Scotinella |
| 2507AshExt6a | AshExt | July | Araneae | Salticidae | Maevia |
| 2507AshExt6a | AshExt | July | Araneae | Salticidae | Sibianor |
| 2507AshExt6a | AshExt | July | Araneae | Theridiidae | Canalidion |
| 2507AshExt6a | AshExt | July | Araneae | Theridiidae | Rugathodes |
| 2507AshExt6a | AshExt | July | Araneae | Theridiidae | Spintharus |
| 2507AshExt6a | AshExt | July | Araneae | Thomisidae | Mecaphesa |

| | | | | | |
|---|---|---|---|---|---|
| 2507AshExt6a | AshExt | July | Blattodea | Ectobiidae | Parcoblatta |
| 2507AshExt6a | AshExt | July | Coleoptera | Buprestidae | Pachyschelus |
| 2507AshExt6a | AshExt | July | Coleoptera | Carabidae | Apenes |
| 2507AshExt6a | AshExt | July | Coleoptera | Carabidae | Harpalus |
| 2507AshExt6a | AshExt | July | Coleoptera | Carabidae | Other |
| 2507AshExt6a | AshExt | July | Coleoptera | Carabidae | Polyderis |
| 2507AshExt6a | AshExt | July | Coleoptera | Curculionidae | Conotrachelus |
| 2507AshExt6a | AshExt | July | Coleoptera | Curculionidae | Cyrtepistomus |
| 2507AshExt6a | AshExt | July | Coleoptera | Curculionidae | Listronotus |
| 2507AshExt6a | AshExt | July | Coleoptera | Dytiscidae | Acilius |
| 2507AshExt6a | AshExt | July | Coleoptera | Lampyridae | Photinus |
| 2507AshExt6a | AshExt | July | Coleoptera | Scarabaeidae | Dialytes |
| 2507AshExt6a | AshExt | July | Coleoptera | Scarabaeidae | Melanocanthon |
| 2507AshExt6a | AshExt | July | Coleoptera | Scirtidae | Contacyphon |
| 2507AshExt6a | AshExt | July | Diptera | Agromyzidae | Calycomyza |
| 2507AshExt6a | AshExt | July | Diptera | Agromyzidae | Cerodontha |
| 2507AshExt6a | AshExt | July | Diptera | Agromyzidae | Japanagromyza |
| 2507AshExt6a | AshExt | July | Diptera | Agromyzidae | Liriomyza |
| 2507AshExt6a | AshExt | July | Diptera | Agromyzidae | Nemorimyza |
| 2507AshExt6a | AshExt | July | Diptera | Agromyzidae | Phytomyza |
| 2507AshExt6a | AshExt | July | Diptera | Anisopodidae | Mycetobia |
| 2507AshExt6a | AshExt | July | Diptera | Calliphoridae | Lucilia |
| 2507AshExt6a | AshExt | July | Diptera | Cecidomyiidae | Other |
| 2507AshExt6a | AshExt | July | Diptera | Chaoboridae | Chaoborus |
| 2507AshExt6a | AshExt | July | Diptera | Chaoboridae | Other |
| 2507AshExt6a | AshExt | July | Diptera | Chironomidae | Ablabesmyia |
| 2507AshExt6a | AshExt | July | Diptera | Chironomidae | Chironomus |
| 2507AshExt6a | AshExt | July | Diptera | Chironomidae | Dicrotendipes |
| 2507AshExt6a | AshExt | July | Diptera | Chironomidae | Endochironomus |
| 2507AshExt6a | AshExt | July | Diptera | Chironomidae | Glyptotendipes |
| 2507AshExt6a | AshExt | July | Diptera | Chironomidae | Microtendipes |

| | | | | | |
|---|---|---|---|---|---|
| 2507AshExt6a | AshExt | July | Diptera | Chironomidae | Orthocladius |
| 2507AshExt6a | AshExt | July | Diptera | Chironomidae | Other |
| 2507AshExt6a | AshExt | July | Diptera | Chironomidae | Other |
| 2507AshExt6a | AshExt | July | Diptera | Chironomidae | Other |
| 2507AshExt6a | AshExt | July | Diptera | Chironomidae | Other |
| 2507AshExt6a | AshExt | July | Diptera | Chironomidae | Parachironomus |
| 2507AshExt6a | AshExt | July | Diptera | Chironomidae | Polypedilum |
| 2507AshExt6a | AshExt | July | Diptera | Chironomidae | Pseudochironomus |
| 2507AshExt6a | AshExt | July | Diptera | Chironomidae | Tanytarsus |
| 2507AshExt6a | AshExt | July | Diptera | Chironomidae | Xenochironomus |
| 2507AshExt6a | AshExt | July | Diptera | Chloropidae | Apotropina |
| 2507AshExt6a | AshExt | July | Diptera | Chloropidae | Elachiptera |
| 2507AshExt6a | AshExt | July | Diptera | Chloropidae | Liohippelates |
| 2507AshExt6a | AshExt | July | Diptera | Chloropidae | Oscinella |
| 2507AshExt6a | AshExt | July | Diptera | Culicidae | Coquillettidia |
| 2507AshExt6a | AshExt | July | Diptera | Culicidae | Culex |
| 2507AshExt6a | AshExt | July | Diptera | Culicidae | Uranotaenia |
| 2507AshExt6a | AshExt | July | Diptera | Drosophilidae | Drosophila |
| 2507AshExt6a | AshExt | July | Diptera | Empididae | Other |
| 2507AshExt6a | AshExt | July | Diptera | Hybotidae | Platypalpus |
| 2507AshExt6a | AshExt | July | Diptera | Hybotidae | Syndyas |
| 2507AshExt6a | AshExt | July | Diptera | Keroplatidae | Other |
| 2507AshExt6a | AshExt | July | Diptera | Limoniidae | Dicranomyia |
| 2507AshExt6a | AshExt | July | Diptera | Limoniidae | Dicranophragma |
| 2507AshExt6a | AshExt | July | Diptera | Limoniidae | Epiphragma |
| 2507AshExt6a | AshExt | July | Diptera | Limoniidae | Erioptera |
| 2507AshExt6a | AshExt | July | Diptera | Limoniidae | Metalimnobia |
| 2507AshExt6a | AshExt | July | Diptera | Limoniidae | Ormosia |
| 2507AshExt6a | AshExt | July | Diptera | Limoniidae | Other |
| 2507AshExt6a | AshExt | July | Diptera | Limoniidae | Other |
| 2507AshExt6a | AshExt | July | Diptera | Limoniidae | Pseudolimnophila |

| | | | | | |
|---|---|---|---|---|---|
| 2507AshExt6a | AshExt | July | Diptera | Muscidae | Mydaea |
| 2507AshExt6a | AshExt | July | Diptera | Mycetophilidae | Mycomya |
| 2507AshExt6a | AshExt | July | Diptera | Mycetophilidae | Neoempheria |
| 2507AshExt6a | AshExt | July | Diptera | Opomyzidae | Geomyza |
| 2507AshExt6a | AshExt | July | Diptera | Pallopteridae | Toxonevra |
| 2507AshExt6a | AshExt | July | Diptera | Phoridae | Diplonevra |
| 2507AshExt6a | AshExt | July | Diptera | Polleniidae | Pollenia |
| 2507AshExt6a | AshExt | July | Diptera | Psychodidae | Clogmia |
| 2507AshExt6a | AshExt | July | Diptera | Psychodidae | Psychoda |
| 2507AshExt6a | AshExt | July | Diptera | Sarcophagidae | Ravinia |
| 2507AshExt6a | AshExt | July | Diptera | Sciaridae | Claustropyga |
| 2507AshExt6a | AshExt | July | Diptera | Sciaridae | Corynoptera |
| 2507AshExt6a | AshExt | July | Diptera | Sciaridae | Epidapus |
| 2507AshExt6a | AshExt | July | Diptera | Sciaridae | Other |
| 2507AshExt6a | AshExt | July | Diptera | Sphaeroceridae | Leptocera |
| 2507AshExt6a | AshExt | July | Diptera | Syrphidae | Chalcosyrphus |
| 2507AshExt6a | AshExt | July | Diptera | Syrphidae | Palpada |
| 2507AshExt6a | AshExt | July | Diptera | Tabanidae | Hybomitra |
| 2507AshExt6a | AshExt | July | Diptera | Tabanidae | Other |
| 2507AshExt6a | AshExt | July | Diptera | Tachinidae | Cryptomeigenia |
| 2507AshExt6a | AshExt | July | Diptera | Tachinidae | Cyzenis |
| 2507AshExt6a | AshExt | July | Diptera | Tachinidae | Zaira |
| 2507AshExt6a | AshExt | July | Diptera | Tipulidae | Dolichopeza |
| 2507AshExt6a | AshExt | July | Diptera | Tipulidae | Tipula |
| 2507AshExt6a | AshExt | July | Ephemeroptera | Caenidae | Caenis |
| 2507AshExt6a | AshExt | July | Ephemeroptera | Ephemeridae | Hexagenia |
| 2507AshExt6a | AshExt | July | Ephemeroptera | Heptageniidae | Epeorus |
| 2507AshExt6a | AshExt | July | Hemiptera | Clastopteridae | Clastoptera |
| 2507AshExt6a | AshExt | July | Hemiptera | Lygaeidae | Kleidocerys |
| 2507AshExt6a | AshExt | July | Hymenoptera | Apidae | Triepeolus |
| 2507AshExt6a | AshExt | July | Hymenoptera | Braconidae | Bracon |

| | | | | | |
|---|---|---|---|---|---|
| 2507AshExt6a | AshExt | July | Hymenoptera | Braconidae | Chorebus |
| 2507AshExt6a | AshExt | July | Hymenoptera | Ichneumonidae | Megastylus |
| 2507AshExt6a | AshExt | July | Hymenoptera | Ichneumonidae | Polyblastus |
| 2507AshExt6a | AshExt | July | Hymenoptera | Other | Other |
| 2507AshExt6a | AshExt | July | Hymenoptera | Tenthredinidae | Dolerus |
| 2507AshExt6a | AshExt | July | Hymenoptera | Tenthredinidae | Nematus |
| 2507AshExt6a | AshExt | July | Hymenoptera | Trigonalidae | Orthogonalys |
| 2507AshExt6a | AshExt | July | Lepidoptera | Crambidae | Blepharomastix |
| 2507AshExt6a | AshExt | July | Lepidoptera | Crambidae | Parapoynx |
| 2507AshExt6a | AshExt | July | Lepidoptera | Elachistidae | Elachista |
| 2507AshExt6a | AshExt | July | Lepidoptera | Erebidae | Panopoda |
| 2507AshExt6a | AshExt | July | Lepidoptera | Gelechiidae | Chionodes |
| 2507AshExt6a | AshExt | July | Lepidoptera | Noctuidae | Condica |
| 2507AshExt6a | AshExt | July | Lepidoptera | Noctuidae | Leuconycta |
| 2507AshExt6a | AshExt | July | Lepidoptera | Noctuidae | Pyreferra |
| 2507AshExt6a | AshExt | July | Lepidoptera | Notodontidae | Heterocampa |
| 2507AshExt6a | AshExt | July | Lepidoptera | Notodontidae | Hyparpax |
| 2507AshExt6a | AshExt | July | Lepidoptera | Nymphalidae | Oeneis |
| 2507AshExt6a | AshExt | July | Lepidoptera | Pyralidae | Atheloca |
| 2507AshExt6a | AshExt | July | Lepidoptera | Pyralidae | Elasmopalpus |
| 2507AshExt6a | AshExt | July | Lepidoptera | Tischeriidae | Coptotriche |
| 2507AshExt6a | AshExt | July | Lepidoptera | Tortricidae | Acleris |
| 2507AshExt6a | AshExt | July | Lepidoptera | Tortricidae | Argyrotaenia |
| 2507AshExt6a | AshExt | July | Lepidoptera | Tortricidae | Rhyacionia |
| 2507AshExt6a | AshExt | July | Megaloptera | Corydalidae | Corydalus |
| 2507AshExt6a | AshExt | July | Neuroptera | Hemerobiidae | Hemerobius |
| 2507AshExt6a | AshExt | July | Odonata | Libellulidae | Brachymesia |
| 2507AshExt6a | AshExt | July | Orthoptera | Tettigoniidae | Conocephalus |
| 2507AshExt6a | AshExt | July | Other | Other | Other |
| 2507AshExt6a | AshExt | July | Other | Other | Other |
| 2507AshExt6a | AshExt | July | Plecoptera | Chloroperlidae | Sweltsa |

| | | | | | |
|---|---|---|---|---|---|
| 2507AshExt6a | AshExt | July | Psocodea | Peripsocidae | Peripsocus |
| 2507AshExt6a | AshExt | July | Sarcoptiformes | Oripodidae | Other |
| 2507AshExt6a | AshExt | July | Trichoptera | Dipseudopsidae | Phylocentropus |
| 2507AshExt6a | AshExt | July | Trichoptera | Hydroptilidae | Orthotrichia |
| 2507AshExt6a | AshExt | July | Trichoptera | Leptoceridae | Oecetis |
| 2507AshExt6a | AshExt | July | Trombidiformes | Arrenuridae | Arrenurus |
| 2507AshExt6a | AshExt | July | Trombidiformes | Aturidae | Aturus |
| 2507AshExt6a | AshExt | July | Trombidiformes | Axonopsidae | Brachypoda |
| 2507AshExt6a | AshExt | July | Trombidiformes | Eupodidae | Other |
| 2507AshExt6a | AshExt | July | Trombidiformes | Feltriidae | Feltria |
| 2507AshExt6a | AshExt | July | Trombidiformes | Hydryphantidae | Hydryphantes |
| 2507AshExt6a | AshExt | July | Trombidiformes | Hydryphantidae | Parathyas |
| 2507AshExt6a | AshExt | July | Trombidiformes | Lebertiidae | Lebertia |
| 2507AshExt6a | AshExt | July | Trombidiformes | Mideopsidae | Mideopsis |
| 2507AshExt6a | AshExt | July | Trombidiformes | Pionidae | Piona |
| 2507AshExt6a | AshExt | July | Trombidiformes | Pionidae | Tiphys |
| 2507AshExt6a | AshExt | July | Trombidiformes | Sperchontidae | Sperchon |
| 2507AshExt6a | AshExt | July | Trombidiformes | Tarsonemidae | Phytonemus |
| 2507AshExt6a | AshExt | July | Trombidiformes | Torrenticolidae | Testudacarus |
| 2507AshExt6a | AshExt | July | Trombidiformes | Torrenticolidae | Torrenticola |
| 2507AshExt6a | AshExt | July | Trombidiformes | Unionicolidae | Koenikea |
| 2507AshExt6c | AshExt | July | Araneae | Agelenidae | Coras |
| 2507AshExt6c | AshExt | July | Araneae | Amaurobiidae | Callobius |
| 2507AshExt6c | AshExt | July | Araneae | Anyphaenidae | Anyphaena |
| 2507AshExt6c | AshExt | July | Araneae | Cicurinidae | Cicurina |
| 2507AshExt6c | AshExt | July | Araneae | Dictynidae | Dictyna |
| 2507AshExt6c | AshExt | July | Araneae | Gnaphosidae | Gnaphosa |
| 2507AshExt6c | AshExt | July | Araneae | Linyphiidae | Grammonota |
| 2507AshExt6c | AshExt | July | Araneae | Lycosidae | Rabidosa |
| 2507AshExt6c | AshExt | July | Araneae | Lycosidae | Schizocosa |
| 2507AshExt6c | AshExt | July | Araneae | Phrurolithidae | Scotinella |

| | | | | | |
|---|---|---|---|---|---|
| 2507AshExt6c | AshExt | July | Araneae | Salticidae | Maevia |
| 2507AshExt6c | AshExt | July | Araneae | Salticidae | Sibianor |
| 2507AshExt6c | AshExt | July | Araneae | Theridiidae | Canalidion |
| 2507AshExt6c | AshExt | July | Araneae | Theridiidae | Rugathodes |
| 2507AshExt6c | AshExt | July | Araneae | Theridiidae | Spintharus |
| 2507AshExt6c | AshExt | July | Araneae | Thomisidae | Mecaphesa |
| 2507AshExt6c | AshExt | July | Blattodea | Ectobiidae | Parcoblatta |
| 2507AshExt6c | AshExt | July | Coleoptera | Buprestidae | Pachyschelus |
| 2507AshExt6c | AshExt | July | Coleoptera | Carabidae | Apenes |
| 2507AshExt6c | AshExt | July | Coleoptera | Carabidae | Harpalus |
| 2507AshExt6c | AshExt | July | Coleoptera | Carabidae | Other |
| 2507AshExt6c | AshExt | July | Coleoptera | Carabidae | Polyderis |
| 2507AshExt6c | AshExt | July | Coleoptera | Curculionidae | Conotrachelus |
| 2507AshExt6c | AshExt | July | Coleoptera | Curculionidae | Cyrtepistomus |
| 2507AshExt6c | AshExt | July | Coleoptera | Curculionidae | Listronotus |
| 2507AshExt6c | AshExt | July | Coleoptera | Dytiscidae | Acilius |
| 2507AshExt6c | AshExt | July | Coleoptera | Lampyridae | Photinus |
| 2507AshExt6c | AshExt | July | Coleoptera | Scarabaeidae | Dialytes |
| 2507AshExt6c | AshExt | July | Coleoptera | Scarabaeidae | Melanocanthon |
| 2507AshExt6c | AshExt | July | Coleoptera | Scirtidae | Contacyphon |
| 2507AshExt6c | AshExt | July | Diptera | Agromyzidae | Calycomyza |
| 2507AshExt6c | AshExt | July | Diptera | Agromyzidae | Cerodontha |
| 2507AshExt6c | AshExt | July | Diptera | Agromyzidae | Japanagromyza |
| 2507AshExt6c | AshExt | July | Diptera | Agromyzidae | Liriomyza |
| 2507AshExt6c | AshExt | July | Diptera | Agromyzidae | Nemorimyza |
| 2507AshExt6c | AshExt | July | Diptera | Agromyzidae | Phytomyza |
| 2507AshExt6c | AshExt | July | Diptera | Anisopodidae | Mycetobia |
| 2507AshExt6c | AshExt | July | Diptera | Calliphoridae | Lucilia |
| 2507AshExt6c | AshExt | July | Diptera | Cecidomyiidae | Other |
| 2507AshExt6c | AshExt | July | Diptera | Chaoboridae | Chaoborus |
| 2507AshExt6c | AshExt | July | Diptera | Chaoboridae | Other |

| | | | | | |
|---|---|---|---|---|---|
| 2507AshExt6c | AshExt | July | Diptera | Chironomidae | Ablabesmyia |
| 2507AshExt6c | AshExt | July | Diptera | Chironomidae | Chironomus |
| 2507AshExt6c | AshExt | July | Diptera | Chironomidae | Dicrotendipes |
| 2507AshExt6c | AshExt | July | Diptera | Chironomidae | Endochironomus |
| 2507AshExt6c | AshExt | July | Diptera | Chironomidae | Glyptotendipes |
| 2507AshExt6c | AshExt | July | Diptera | Chironomidae | Microtendipes |
| 2507AshExt6c | AshExt | July | Diptera | Chironomidae | Orthocladius |
| 2507AshExt6c | AshExt | July | Diptera | Chironomidae | Other |
| 2507AshExt6c | AshExt | July | Diptera | Chironomidae | Other |
| 2507AshExt6c | AshExt | July | Diptera | Chironomidae | Other |
| 2507AshExt6c | AshExt | July | Diptera | Chironomidae | Other |
| 2507AshExt6c | AshExt | July | Diptera | Chironomidae | Parachironomus |
| 2507AshExt6c | AshExt | July | Diptera | Chironomidae | Polypedilum |
| 2507AshExt6c | AshExt | July | Diptera | Chironomidae | Pseudochironomus |
| 2507AshExt6c | AshExt | July | Diptera | Chironomidae | Tanytarsus |
| 2507AshExt6c | AshExt | July | Diptera | Chironomidae | Xenochironomus |
| 2507AshExt6c | AshExt | July | Diptera | Chloropidae | Apotropina |
| 2507AshExt6c | AshExt | July | Diptera | Chloropidae | Elachiptera |
| 2507AshExt6c | AshExt | July | Diptera | Chloropidae | Liohippelates |
| 2507AshExt6c | AshExt | July | Diptera | Chloropidae | Oscinella |
| 2507AshExt6c | AshExt | July | Diptera | Culicidae | Coquillettidia |
| 2507AshExt6c | AshExt | July | Diptera | Culicidae | Culex |
| 2507AshExt6c | AshExt | July | Diptera | Culicidae | Uranotaenia |
| 2507AshExt6c | AshExt | July | Diptera | Drosophilidae | Drosophila |
| 2507AshExt6c | AshExt | July | Diptera | Empididae | Other |
| 2507AshExt6c | AshExt | July | Diptera | Hybotidae | Platypalpus |
| 2507AshExt6c | AshExt | July | Diptera | Hybotidae | Syndyas |
| 2507AshExt6c | AshExt | July | Diptera | Keroplatidae | Other |
| 2507AshExt6c | AshExt | July | Diptera | Limoniidae | Dicranomyia |
| 2507AshExt6c | AshExt | July | Diptera | Limoniidae | Dicranophragma |
| 2507AshExt6c | AshExt | July | Diptera | Limoniidae | Epiphragma |

| | | | | | |
|---|---|---|---|---|---|
| 2507AshExt6c | AshExt | July | Diptera | Limoniidae | Erioptera |
| 2507AshExt6c | AshExt | July | Diptera | Limoniidae | Metalimnobia |
| 2507AshExt6c | AshExt | July | Diptera | Limoniidae | Ormosia |
| 2507AshExt6c | AshExt | July | Diptera | Limoniidae | Other |
| 2507AshExt6c | AshExt | July | Diptera | Limoniidae | Other |
| 2507AshExt6c | AshExt | July | Diptera | Limoniidae | Pseudolimnophila |
| 2507AshExt6c | AshExt | July | Diptera | Muscidae | Mydaea |
| 2507AshExt6c | AshExt | July | Diptera | Mycetophilidae | Mycomya |
| 2507AshExt6c | AshExt | July | Diptera | Mycetophilidae | Neoempheria |
| 2507AshExt6c | AshExt | July | Diptera | Opomyzidae | Geomyza |
| 2507AshExt6c | AshExt | July | Diptera | Pallopteridae | Toxonevra |
| 2507AshExt6c | AshExt | July | Diptera | Phoridae | Diplonevra |
| 2507AshExt6c | AshExt | July | Diptera | Polleniidae | Pollenia |
| 2507AshExt6c | AshExt | July | Diptera | Psychodidae | Clogmia |
| 2507AshExt6c | AshExt | July | Diptera | Psychodidae | Psychoda |
| 2507AshExt6c | AshExt | July | Diptera | Sarcophagidae | Ravinia |
| 2507AshExt6c | AshExt | July | Diptera | Sciaridae | Claustropyga |
| 2507AshExt6c | AshExt | July | Diptera | Sciaridae | Corynoptera |
| 2507AshExt6c | AshExt | July | Diptera | Sciaridae | Epidapus |
| 2507AshExt6c | AshExt | July | Diptera | Sciaridae | Other |
| 2507AshExt6c | AshExt | July | Diptera | Sphaeroceridae | Leptocera |
| 2507AshExt6c | AshExt | July | Diptera | Syrphidae | Chalcosyrphus |
| 2507AshExt6c | AshExt | July | Diptera | Syrphidae | Palpada |
| 2507AshExt6c | AshExt | July | Diptera | Tabanidae | Hybomitra |
| 2507AshExt6c | AshExt | July | Diptera | Tabanidae | Other |
| 2507AshExt6c | AshExt | July | Diptera | Tachinidae | Cryptomeigenia |
| 2507AshExt6c | AshExt | July | Diptera | Tachinidae | Cyzenis |
| 2507AshExt6c | AshExt | July | Diptera | Tachinidae | Zaira |
| 2507AshExt6c | AshExt | July | Diptera | Tipulidae | Dolichopeza |
| 2507AshExt6c | AshExt | July | Diptera | Tipulidae | Tipula |
| 2507AshExt6c | AshExt | July | Ephemeroptera | Caenidae | Caenis |

| | | | | | |
|---|---|---|---|---|---|
| 2507AshExt6c | AshExt | July | Ephemeroptera | Ephemeridae | Hexagenia |
| 2507AshExt6c | AshExt | July | Ephemeroptera | Heptageniidae | Epeorus |
| 2507AshExt6c | AshExt | July | Hemiptera | Clastopteridae | Clastoptera |
| 2507AshExt6c | AshExt | July | Hemiptera | Lygaeidae | Kleidocerys |
| 2507AshExt6c | AshExt | July | Hymenoptera | Apidae | Triepeolus |
| 2507AshExt6c | AshExt | July | Hymenoptera | Braconidae | Bracon |
| 2507AshExt6c | AshExt | July | Hymenoptera | Braconidae | Chorebus |
| 2507AshExt6c | AshExt | July | Hymenoptera | Ichneumonidae | Megastylus |
| 2507AshExt6c | AshExt | July | Hymenoptera | Ichneumonidae | Polyblastus |
| 2507AshExt6c | AshExt | July | Hymenoptera | Other | Other |
| 2507AshExt6c | AshExt | July | Hymenoptera | Tenthredinidae | Dolerus |
| 2507AshExt6c | AshExt | July | Hymenoptera | Tenthredinidae | Nematus |
| 2507AshExt6c | AshExt | July | Hymenoptera | Trigonalidae | Orthogonalys |
| 2507AshExt6c | AshExt | July | Lepidoptera | Crambidae | Blepharomastix |
| 2507AshExt6c | AshExt | July | Lepidoptera | Crambidae | Parapoynx |
| 2507AshExt6c | AshExt | July | Lepidoptera | Elachistidae | Elachista |
| 2507AshExt6c | AshExt | July | Lepidoptera | Erebidae | Panopoda |
| 2507AshExt6c | AshExt | July | Lepidoptera | Gelechiidae | Chionodes |
| 2507AshExt6c | AshExt | July | Lepidoptera | Noctuidae | Condica |
| 2507AshExt6c | AshExt | July | Lepidoptera | Noctuidae | Leuconycta |
| 2507AshExt6c | AshExt | July | Lepidoptera | Noctuidae | Pyreferra |
| 2507AshExt6c | AshExt | July | Lepidoptera | Notodontidae | Heterocampa |
| 2507AshExt6c | AshExt | July | Lepidoptera | Notodontidae | Hyparpax |
| 2507AshExt6c | AshExt | July | Lepidoptera | Nymphalidae | Oeneis |
| 2507AshExt6c | AshExt | July | Lepidoptera | Pyralidae | Atheloca |
| 2507AshExt6c | AshExt | July | Lepidoptera | Pyralidae | Elasmopalpus |
| 2507AshExt6c | AshExt | July | Lepidoptera | Tischeriidae | Coptotriche |
| 2507AshExt6c | AshExt | July | Lepidoptera | Tortricidae | Acleris |
| 2507AshExt6c | AshExt | July | Lepidoptera | Tortricidae | Argyrotaenia |
| 2507AshExt6c | AshExt | July | Lepidoptera | Tortricidae | Rhyacionia |
| 2507AshExt6c | AshExt | July | Megaloptera | Corydalidae | Corydalus |

| | | | | | |
|---|---|---|---|---|---|
| 2507AshExt6c | AshExt | July | Neuroptera | Hemerobiidae | Hemerobius |
| 2507AshExt6c | AshExt | July | Odonata | Libellulidae | Brachymesia |
| 2507AshExt6c | AshExt | July | Orthoptera | Tettigoniidae | Conocephalus |
| 2507AshExt6c | AshExt | July | Other | Other | Other |
| 2507AshExt6c | AshExt | July | Other | Other | Other |
| 2507AshExt6c | AshExt | July | Plecoptera | Chloroperlidae | Sweltsa |
| 2507AshExt6c | AshExt | July | Psocodea | Peripsocidae | Peripsocus |
| 2507AshExt6c | AshExt | July | Sarcoptiformes | Oripodidae | Other |
| 2507AshExt6c | AshExt | July | Trichoptera | Dipseudopsidae | Phylocentropus |
| 2507AshExt6c | AshExt | July | Trichoptera | Hydroptilidae | Orthotrichia |
| 2507AshExt6c | AshExt | July | Trichoptera | Leptoceridae | Oecetis |
| 2507AshExt6c | AshExt | July | Trombidiformes | Arrenuridae | Arrenurus |
| 2507AshExt6c | AshExt | July | Trombidiformes | Aturidae | Aturus |
| 2507AshExt6c | AshExt | July | Trombidiformes | Axonopsidae | Brachypoda |
| 2507AshExt6c | AshExt | July | Trombidiformes | Eupodidae | Other |
| 2507AshExt6c | AshExt | July | Trombidiformes | Feltriidae | Feltria |
| 2507AshExt6c | AshExt | July | Trombidiformes | Hydryphantidae | Hydryphantes |
| 2507AshExt6c | AshExt | July | Trombidiformes | Hydryphantidae | Parathyas |
| 2507AshExt6c | AshExt | July | Trombidiformes | Lebertiidae | Lebertia |
| 2507AshExt6c | AshExt | July | Trombidiformes | Mideopsidae | Mideopsis |
| 2507AshExt6c | AshExt | July | Trombidiformes | Pionidae | Piona |
| 2507AshExt6c | AshExt | July | Trombidiformes | Pionidae | Tiphys |
| 2507AshExt6c | AshExt | July | Trombidiformes | Sperchontidae | Sperchon |
| 2507AshExt6c | AshExt | July | Trombidiformes | Tarsonemidae | Phytonemus |
| 2507AshExt6c | AshExt | July | Trombidiformes | Torrenticolidae | Testudacarus |
| 2507AshExt6c | AshExt | July | Trombidiformes | Torrenticolidae | Torrenticola |
| 2507AshExt6c | AshExt | July | Trombidiformes | Unionicolidae | Koenikea |
| 2508AshExt6a | AshExt | August | Araneae | Agelenidae | Coras |
| 2508AshExt6a | AshExt | August | Araneae | Amaurobiidae | Callobius |
| 2508AshExt6a | AshExt | August | Araneae | Anyphaenidae | Anyphaena |
| 2508AshExt6a | AshExt | August | Araneae | Cicurinidae | Cicurina |

| | | | | | |
|---|---|---|---|---|---|
| 2508AshExt6a | AshExt | August | Araneae | Dictynidae | Dictyna |
| 2508AshExt6a | AshExt | August | Araneae | Gnaphosidae | Gnaphosa |
| 2508AshExt6a | AshExt | August | Araneae | Linyphiidae | Grammonota |
| 2508AshExt6a | AshExt | August | Araneae | Lycosidae | Rabidosa |
| 2508AshExt6a | AshExt | August | Araneae | Lycosidae | Schizocosa |
| 2508AshExt6a | AshExt | August | Araneae | Phrurolithidae | Scotinella |
| 2508AshExt6a | AshExt | August | Araneae | Salticidae | Maevia |
| 2508AshExt6a | AshExt | August | Araneae | Salticidae | Sibianor |
| 2508AshExt6a | AshExt | August | Araneae | Theridiidae | Canalidion |
| 2508AshExt6a | AshExt | August | Araneae | Theridiidae | Rugathodes |
| 2508AshExt6a | AshExt | August | Araneae | Theridiidae | Spintharus |
| 2508AshExt6a | AshExt | August | Araneae | Thomisidae | Mecaphesa |
| 2508AshExt6a | AshExt | August | Blattodea | Ectobiidae | Parcoblatta |
| 2508AshExt6a | AshExt | August | Coleoptera | Buprestidae | Pachyschelus |
| 2508AshExt6a | AshExt | August | Coleoptera | Carabidae | Apenes |
| 2508AshExt6a | AshExt | August | Coleoptera | Carabidae | Harpalus |
| 2508AshExt6a | AshExt | August | Coleoptera | Carabidae | Other |
| 2508AshExt6a | AshExt | August | Coleoptera | Carabidae | Polyderis |
| 2508AshExt6a | AshExt | August | Coleoptera | Curculionidae | Conotrachelus |
| 2508AshExt6a | AshExt | August | Coleoptera | Curculionidae | Cyrtepistomus |
| 2508AshExt6a | AshExt | August | Coleoptera | Curculionidae | Listronotus |
| 2508AshExt6a | AshExt | August | Coleoptera | Dytiscidae | Acilius |
| 2508AshExt6a | AshExt | August | Coleoptera | Lampyridae | Photinus |
| 2508AshExt6a | AshExt | August | Coleoptera | Scarabaeidae | Dialytes |
| 2508AshExt6a | AshExt | August | Coleoptera | Scarabaeidae | Melanocanthon |
| 2508AshExt6a | AshExt | August | Coleoptera | Scirtidae | Contacyphon |
| 2508AshExt6a | AshExt | August | Diptera | Agromyzidae | Calycomyza |
| 2508AshExt6a | AshExt | August | Diptera | Agromyzidae | Cerodontha |
| 2508AshExt6a | AshExt | August | Diptera | Agromyzidae | Japanagromyza |
| 2508AshExt6a | AshExt | August | Diptera | Agromyzidae | Liriomyza |
| 2508AshExt6a | AshExt | August | Diptera | Agromyzidae | Nemorimyza |

| | | | | | |
|---|---|---|---|---|---|
| 2508AshExt6a | AshExt | August | Diptera | Agromyzidae | Phytomyza |
| 2508AshExt6a | AshExt | August | Diptera | Anisopodidae | Mycetobia |
| 2508AshExt6a | AshExt | August | Diptera | Calliphoridae | Lucilia |
| 2508AshExt6a | AshExt | August | Diptera | Cecidomyiidae | Other |
| 2508AshExt6a | AshExt | August | Diptera | Chaoboridae | Chaoborus |
| 2508AshExt6a | AshExt | August | Diptera | Chaoboridae | Other |
| 2508AshExt6a | AshExt | August | Diptera | Chironomidae | Ablabesmyia |
| 2508AshExt6a | AshExt | August | Diptera | Chironomidae | Chironomus |
| 2508AshExt6a | AshExt | August | Diptera | Chironomidae | Dicrotendipes |
| 2508AshExt6a | AshExt | August | Diptera | Chironomidae | Endochironomus |
| 2508AshExt6a | AshExt | August | Diptera | Chironomidae | Glyptotendipes |
| 2508AshExt6a | AshExt | August | Diptera | Chironomidae | Microtendipes |
| 2508AshExt6a | AshExt | August | Diptera | Chironomidae | Orthocladius |
| 2508AshExt6a | AshExt | August | Diptera | Chironomidae | Other |
| 2508AshExt6a | AshExt | August | Diptera | Chironomidae | Other |
| 2508AshExt6a | AshExt | August | Diptera | Chironomidae | Other |
| 2508AshExt6a | AshExt | August | Diptera | Chironomidae | Other |
| 2508AshExt6a | AshExt | August | Diptera | Chironomidae | Parachironomus |
| 2508AshExt6a | AshExt | August | Diptera | Chironomidae | Polypedilum |
| 2508AshExt6a | AshExt | August | Diptera | Chironomidae | Pseudochironomus |
| 2508AshExt6a | AshExt | August | Diptera | Chironomidae | Tanytarsus |
| 2508AshExt6a | AshExt | August | Diptera | Chironomidae | Xenochironomus |
| 2508AshExt6a | AshExt | August | Diptera | Chloropidae | Apotropina |
| 2508AshExt6a | AshExt | August | Diptera | Chloropidae | Elachiptera |
| 2508AshExt6a | AshExt | August | Diptera | Chloropidae | Liohippelates |
| 2508AshExt6a | AshExt | August | Diptera | Chloropidae | Oscinella |
| 2508AshExt6a | AshExt | August | Diptera | Culicidae | Coquillettidia |
| 2508AshExt6a | AshExt | August | Diptera | Culicidae | Culex |
| 2508AshExt6a | AshExt | August | Diptera | Culicidae | Uranotaenia |
| 2508AshExt6a | AshExt | August | Diptera | Drosophilidae | Drosophila |
| 2508AshExt6a | AshExt | August | Diptera | Empididae | Other |

| | | | | | |
|---|---|---|---|---|---|
| 2508AshExt6a | AshExt | August | Diptera | Hybotidae | Platypalpus |
| 2508AshExt6a | AshExt | August | Diptera | Hybotidae | Syndyas |
| 2508AshExt6a | AshExt | August | Diptera | Keroplatidae | Other |
| 2508AshExt6a | AshExt | August | Diptera | Limoniidae | Dicranomyia |
| 2508AshExt6a | AshExt | August | Diptera | Limoniidae | Dicranophragma |
| 2508AshExt6a | AshExt | August | Diptera | Limoniidae | Epiphragma |
| 2508AshExt6a | AshExt | August | Diptera | Limoniidae | Erioptera |
| 2508AshExt6a | AshExt | August | Diptera | Limoniidae | Metalimnobia |
| 2508AshExt6a | AshExt | August | Diptera | Limoniidae | Ormosia |
| 2508AshExt6a | AshExt | August | Diptera | Limoniidae | Other |
| 2508AshExt6a | AshExt | August | Diptera | Limoniidae | Other |
| 2508AshExt6a | AshExt | August | Diptera | Limoniidae | Pseudolimnophila |
| 2508AshExt6a | AshExt | August | Diptera | Muscidae | Mydaea |
| 2508AshExt6a | AshExt | August | Diptera | Mycetophilidae | Mycomya |
| 2508AshExt6a | AshExt | August | Diptera | Mycetophilidae | Neoempheria |
| 2508AshExt6a | AshExt | August | Diptera | Opomyzidae | Geomyza |
| 2508AshExt6a | AshExt | August | Diptera | Pallopteridae | Toxonevra |
| 2508AshExt6a | AshExt | August | Diptera | Phoridae | Diplonevra |
| 2508AshExt6a | AshExt | August | Diptera | Polleniidae | Pollenia |
| 2508AshExt6a | AshExt | August | Diptera | Psychodidae | Clogmia |
| 2508AshExt6a | AshExt | August | Diptera | Psychodidae | Psychoda |
| 2508AshExt6a | AshExt | August | Diptera | Sarcophagidae | Ravinia |
| 2508AshExt6a | AshExt | August | Diptera | Sciaridae | Claustropyga |
| 2508AshExt6a | AshExt | August | Diptera | Sciaridae | Corynoptera |
| 2508AshExt6a | AshExt | August | Diptera | Sciaridae | Epidapus |
| 2508AshExt6a | AshExt | August | Diptera | Sciaridae | Other |
| 2508AshExt6a | AshExt | August | Diptera | Sphaeroceridae | Leptocera |
| 2508AshExt6a | AshExt | August | Diptera | Syrphidae | Chalcosyrphus |
| 2508AshExt6a | AshExt | August | Diptera | Syrphidae | Palpada |
| 2508AshExt6a | AshExt | August | Diptera | Tabanidae | Hybomitra |
| 2508AshExt6a | AshExt | August | Diptera | Tabanidae | Other |

| | | | | | |
|---|---|---|---|---|---|
| 2508AshExt6a | AshExt | August | Diptera | Tachinidae | Cryptomeigenia |
| 2508AshExt6a | AshExt | August | Diptera | Tachinidae | Cyzenis |
| 2508AshExt6a | AshExt | August | Diptera | Tachinidae | Zaira |
| 2508AshExt6a | AshExt | August | Diptera | Tipulidae | Dolichopeza |
| 2508AshExt6a | AshExt | August | Diptera | Tipulidae | Tipula |
| 2508AshExt6a | AshExt | August | Ephemeroptera | Caenidae | Caenis |
| 2508AshExt6a | AshExt | August | Ephemeroptera | Ephemeridae | Hexagenia |
| 2508AshExt6a | AshExt | August | Ephemeroptera | Heptageniidae | Epeorus |
| 2508AshExt6a | AshExt | August | Hemiptera | Clastopteridae | Clastoptera |
| 2508AshExt6a | AshExt | August | Hemiptera | Lygaeidae | Kleidocerys |
| 2508AshExt6a | AshExt | August | Hymenoptera | Apidae | Triepeolus |
| 2508AshExt6a | AshExt | August | Hymenoptera | Braconidae | Bracon |
| 2508AshExt6a | AshExt | August | Hymenoptera | Braconidae | Chorebus |
| 2508AshExt6a | AshExt | August | Hymenoptera | Ichneumonidae | Megastylus |
| 2508AshExt6a | AshExt | August | Hymenoptera | Ichneumonidae | Polyblastus |
| 2508AshExt6a | AshExt | August | Hymenoptera | Other | Other |
| 2508AshExt6a | AshExt | August | Hymenoptera | Tenthredinidae | Dolerus |
| 2508AshExt6a | AshExt | August | Hymenoptera | Tenthredinidae | Nematus |
| 2508AshExt6a | AshExt | August | Hymenoptera | Trigonalidae | Orthogonalys |
| 2508AshExt6a | AshExt | August | Lepidoptera | Crambidae | Blepharomastix |
| 2508AshExt6a | AshExt | August | Lepidoptera | Crambidae | Parapoynx |
| 2508AshExt6a | AshExt | August | Lepidoptera | Elachistidae | Elachista |
| 2508AshExt6a | AshExt | August | Lepidoptera | Erebidae | Panopoda |
| 2508AshExt6a | AshExt | August | Lepidoptera | Gelechiidae | Chionodes |
| 2508AshExt6a | AshExt | August | Lepidoptera | Noctuidae | Condica |
| 2508AshExt6a | AshExt | August | Lepidoptera | Noctuidae | Leuconycta |
| 2508AshExt6a | AshExt | August | Lepidoptera | Noctuidae | Pyreferra |
| 2508AshExt6a | AshExt | August | Lepidoptera | Notodontidae | Heterocampa |
| 2508AshExt6a | AshExt | August | Lepidoptera | Notodontidae | Hyparpax |
| 2508AshExt6a | AshExt | August | Lepidoptera | Nymphalidae | Oeneis |
| 2508AshExt6a | AshExt | August | Lepidoptera | Pyralidae | Atheloca |

| | | | | | |
|---|---|---|---|---|---|
| 2508AshExt6a | AshExt | August | Lepidoptera | Pyralidae | Elasmopalpus |
| 2508AshExt6a | AshExt | August | Lepidoptera | Tischeriidae | Coptotriche |
| 2508AshExt6a | AshExt | August | Lepidoptera | Tortricidae | Acleris |
| 2508AshExt6a | AshExt | August | Lepidoptera | Tortricidae | Argyrotaenia |
| 2508AshExt6a | AshExt | August | Lepidoptera | Tortricidae | Rhyacionia |
| 2508AshExt6a | AshExt | August | Megaloptera | Corydalidae | Corydalus |
| 2508AshExt6a | AshExt | August | Neuroptera | Hemerobiidae | Hemerobius |
| 2508AshExt6a | AshExt | August | Odonata | Libellulidae | Brachymesia |
| 2508AshExt6a | AshExt | August | Orthoptera | Tettigoniidae | Conocephalus |
| 2508AshExt6a | AshExt | August | Other | Other | Other |
| 2508AshExt6a | AshExt | August | Other | Other | Other |
| 2508AshExt6a | AshExt | August | Plecoptera | Chloroperlidae | Sweltsa |
| 2508AshExt6a | AshExt | August | Psocodea | Peripsocidae | Peripsocus |
| 2508AshExt6a | AshExt | August | Sarcoptiformes | Oripodidae | Other |
| 2508AshExt6a | AshExt | August | Trichoptera | Dipseudopsidae | Phylocentropus |
| 2508AshExt6a | AshExt | August | Trichoptera | Hydroptilidae | Orthotrichia |
| 2508AshExt6a | AshExt | August | Trichoptera | Leptoceridae | Oecetis |
| 2508AshExt6a | AshExt | August | Trombidiformes | Arrenuridae | Arrenurus |
| 2508AshExt6a | AshExt | August | Trombidiformes | Aturidae | Aturus |
| 2508AshExt6a | AshExt | August | Trombidiformes | Axonopsidae | Brachypoda |
| 2508AshExt6a | AshExt | August | Trombidiformes | Eupodidae | Other |
| 2508AshExt6a | AshExt | August | Trombidiformes | Feltriidae | Feltria |
| 2508AshExt6a | AshExt | August | Trombidiformes | Hydryphantidae | Hydryphantes |
| 2508AshExt6a | AshExt | August | Trombidiformes | Hydryphantidae | Parathyas |
| 2508AshExt6a | AshExt | August | Trombidiformes | Lebertiidae | Lebertia |
| 2508AshExt6a | AshExt | August | Trombidiformes | Mideopsidae | Mideopsis |
| 2508AshExt6a | AshExt | August | Trombidiformes | Pionidae | Piona |
| 2508AshExt6a | AshExt | August | Trombidiformes | Pionidae | Tiphys |
| 2508AshExt6a | AshExt | August | Trombidiformes | Sperchontidae | Sperchon |
| 2508AshExt6a | AshExt | August | Trombidiformes | Tarsonemidae | Phytonemus |
| 2508AshExt6a | AshExt | August | Trombidiformes | Torrenticolidae | Testudacarus |

| | | | | | |
|---|---|---|---|---|---|
| 2508AshExt6a | AshExt | August | Trombidiformes | Torrenticolidae | Torrenticola |
| 2508AshExt6a | AshExt | August | Trombidiformes | Unionicolidae | Koenikea |
| 2508AshExt6b | AshExt | August | Araneae | Agelenidae | Coras |
| 2508AshExt6b | AshExt | August | Araneae | Amaurobiidae | Callobius |
| 2508AshExt6b | AshExt | August | Araneae | Anyphaenidae | Anyphaena |
| 2508AshExt6b | AshExt | August | Araneae | Cicurinidae | Cicurina |
| 2508AshExt6b | AshExt | August | Araneae | Dictynidae | Dictyna |
| 2508AshExt6b | AshExt | August | Araneae | Gnaphosidae | Gnaphosa |
| 2508AshExt6b | AshExt | August | Araneae | Linyphiidae | Grammonota |
| 2508AshExt6b | AshExt | August | Araneae | Lycosidae | Rabidosa |
| 2508AshExt6b | AshExt | August | Araneae | Lycosidae | Schizocosa |
| 2508AshExt6b | AshExt | August | Araneae | Phrurolithidae | Scotinella |
| 2508AshExt6b | AshExt | August | Araneae | Salticidae | Maevia |
| 2508AshExt6b | AshExt | August | Araneae | Salticidae | Sibianor |
| 2508AshExt6b | AshExt | August | Araneae | Theridiidae | Canalidion |
| 2508AshExt6b | AshExt | August | Araneae | Theridiidae | Rugathodes |
| 2508AshExt6b | AshExt | August | Araneae | Theridiidae | Spintharus |
| 2508AshExt6b | AshExt | August | Araneae | Thomisidae | Mecaphesa |
| 2508AshExt6b | AshExt | August | Blattodea | Ectobiidae | Parcoblatta |
| 2508AshExt6b | AshExt | August | Coleoptera | Buprestidae | Pachyschelus |
| 2508AshExt6b | AshExt | August | Coleoptera | Carabidae | Apenes |
| 2508AshExt6b | AshExt | August | Coleoptera | Carabidae | Harpalus |
| 2508AshExt6b | AshExt | August | Coleoptera | Carabidae | Other |
| 2508AshExt6b | AshExt | August | Coleoptera | Carabidae | Polyderis |
| 2508AshExt6b | AshExt | August | Coleoptera | Curculionidae | Conotrachelus |
| 2508AshExt6b | AshExt | August | Coleoptera | Curculionidae | Cyrtepistomus |
| 2508AshExt6b | AshExt | August | Coleoptera | Curculionidae | Listronotus |
| 2508AshExt6b | AshExt | August | Coleoptera | Dytiscidae | Acilius |
| 2508AshExt6b | AshExt | August | Coleoptera | Lampyridae | Photinus |
| 2508AshExt6b | AshExt | August | Coleoptera | Scarabaeidae | Dialytes |
| 2508AshExt6b | AshExt | August | Coleoptera | Scarabaeidae | Melanocanthon |

| | | | | | |
|---|---|---|---|---|---|
| 2508AshExt6b | AshExt | August | Coleoptera | Scirtidae | Contacyphon |
| 2508AshExt6b | AshExt | August | Diptera | Agromyzidae | Calycomyza |
| 2508AshExt6b | AshExt | August | Diptera | Agromyzidae | Cerodontha |
| 2508AshExt6b | AshExt | August | Diptera | Agromyzidae | Japanagromyza |
| 2508AshExt6b | AshExt | August | Diptera | Agromyzidae | Liriomyza |
| 2508AshExt6b | AshExt | August | Diptera | Agromyzidae | Nemorimyza |
| 2508AshExt6b | AshExt | August | Diptera | Agromyzidae | Phytomyza |
| 2508AshExt6b | AshExt | August | Diptera | Anisopodidae | Mycetobia |
| 2508AshExt6b | AshExt | August | Diptera | Calliphoridae | Lucilia |
| 2508AshExt6b | AshExt | August | Diptera | Cecidomyiidae | Other |
| 2508AshExt6b | AshExt | August | Diptera | Chaoboridae | Chaoborus |
| 2508AshExt6b | AshExt | August | Diptera | Chaoboridae | Other |
| 2508AshExt6b | AshExt | August | Diptera | Chironomidae | Ablabesmyia |
| 2508AshExt6b | AshExt | August | Diptera | Chironomidae | Chironomus |
| 2508AshExt6b | AshExt | August | Diptera | Chironomidae | Dicrotendipes |
| 2508AshExt6b | AshExt | August | Diptera | Chironomidae | Endochironomus |
| 2508AshExt6b | AshExt | August | Diptera | Chironomidae | Glyptotendipes |
| 2508AshExt6b | AshExt | August | Diptera | Chironomidae | Microtendipes |
| 2508AshExt6b | AshExt | August | Diptera | Chironomidae | Orthocladius |
| 2508AshExt6b | AshExt | August | Diptera | Chironomidae | Other |
| 2508AshExt6b | AshExt | August | Diptera | Chironomidae | Other |
| 2508AshExt6b | AshExt | August | Diptera | Chironomidae | Other |
| 2508AshExt6b | AshExt | August | Diptera | Chironomidae | Other |
| 2508AshExt6b | AshExt | August | Diptera | Chironomidae | Parachironomus |
| 2508AshExt6b | AshExt | August | Diptera | Chironomidae | Polypedilum |
| 2508AshExt6b | AshExt | August | Diptera | Chironomidae | Pseudochironomus |
| 2508AshExt6b | AshExt | August | Diptera | Chironomidae | Tanytarsus |
| 2508AshExt6b | AshExt | August | Diptera | Chironomidae | Xenochironomus |
| 2508AshExt6b | AshExt | August | Diptera | Chloropidae | Apotropina |
| 2508AshExt6b | AshExt | August | Diptera | Chloropidae | Elachiptera |
| 2508AshExt6b | AshExt | August | Diptera | Chloropidae | Liohippelates |

| | | | | | |
|---|---|---|---|---|---|
| 2508AshExt6b | AshExt | August | Diptera | Chloropidae | Oscinella |
| 2508AshExt6b | AshExt | August | Diptera | Culicidae | Coquillettidia |
| 2508AshExt6b | AshExt | August | Diptera | Culicidae | Culex |
| 2508AshExt6b | AshExt | August | Diptera | Culicidae | Uranotaenia |
| 2508AshExt6b | AshExt | August | Diptera | Drosophilidae | Drosophila |
| 2508AshExt6b | AshExt | August | Diptera | Empididae | Other |
| 2508AshExt6b | AshExt | August | Diptera | Hybotidae | Platypalpus |
| 2508AshExt6b | AshExt | August | Diptera | Hybotidae | Syndyas |
| 2508AshExt6b | AshExt | August | Diptera | Keroplatidae | Other |
| 2508AshExt6b | AshExt | August | Diptera | Limoniidae | Dicranomyia |
| 2508AshExt6b | AshExt | August | Diptera | Limoniidae | Dicranophragma |
| 2508AshExt6b | AshExt | August | Diptera | Limoniidae | Epiphragma |
| 2508AshExt6b | AshExt | August | Diptera | Limoniidae | Erioptera |
| 2508AshExt6b | AshExt | August | Diptera | Limoniidae | Metalimnobia |
| 2508AshExt6b | AshExt | August | Diptera | Limoniidae | Ormosia |
| 2508AshExt6b | AshExt | August | Diptera | Limoniidae | Other |
| 2508AshExt6b | AshExt | August | Diptera | Limoniidae | Other |
| 2508AshExt6b | AshExt | August | Diptera | Limoniidae | Pseudolimnophila |
| 2508AshExt6b | AshExt | August | Diptera | Muscidae | Mydaea |
| 2508AshExt6b | AshExt | August | Diptera | Mycetophilidae | Mycomya |
| 2508AshExt6b | AshExt | August | Diptera | Mycetophilidae | Neoempheria |
| 2508AshExt6b | AshExt | August | Diptera | Opomyzidae | Geomyza |
| 2508AshExt6b | AshExt | August | Diptera | Pallopteridae | Toxonevra |
| 2508AshExt6b | AshExt | August | Diptera | Phoridae | Diplonevra |
| 2508AshExt6b | AshExt | August | Diptera | Polleniidae | Pollenia |
| 2508AshExt6b | AshExt | August | Diptera | Psychodidae | Clogmia |
| 2508AshExt6b | AshExt | August | Diptera | Psychodidae | Psychoda |
| 2508AshExt6b | AshExt | August | Diptera | Sarcophagidae | Ravinia |
| 2508AshExt6b | AshExt | August | Diptera | Sciaridae | Claustropyga |
| 2508AshExt6b | AshExt | August | Diptera | Sciaridae | Corynoptera |
| 2508AshExt6b | AshExt | August | Diptera | Sciaridae | Epidapus |

| | | | | | |
|---|---|---|---|---|---|
| 2508AshExt6b | AshExt | August | Diptera | Sciaridae | Other |
| 2508AshExt6b | AshExt | August | Diptera | Sphaeroceridae | Leptocera |
| 2508AshExt6b | AshExt | August | Diptera | Syrphidae | Chalcosyrphus |
| 2508AshExt6b | AshExt | August | Diptera | Syrphidae | Palpada |
| 2508AshExt6b | AshExt | August | Diptera | Tabanidae | Hybomitra |
| 2508AshExt6b | AshExt | August | Diptera | Tabanidae | Other |
| 2508AshExt6b | AshExt | August | Diptera | Tachinidae | Cryptomeigenia |
| 2508AshExt6b | AshExt | August | Diptera | Tachinidae | Cyzenis |
| 2508AshExt6b | AshExt | August | Diptera | Tachinidae | Zaira |
| 2508AshExt6b | AshExt | August | Diptera | Tipulidae | Dolichopeza |
| 2508AshExt6b | AshExt | August | Diptera | Tipulidae | Tipula |
| 2508AshExt6b | AshExt | August | Ephemeroptera | Caenidae | Caenis |
| 2508AshExt6b | AshExt | August | Ephemeroptera | Ephemeridae | Hexagenia |
| 2508AshExt6b | AshExt | August | Ephemeroptera | Heptageniidae | Epeorus |
| 2508AshExt6b | AshExt | August | Hemiptera | Clastopteridae | Clastoptera |
| 2508AshExt6b | AshExt | August | Hemiptera | Lygaeidae | Kleidocerys |
| 2508AshExt6b | AshExt | August | Hymenoptera | Apidae | Triepeolus |
| 2508AshExt6b | AshExt | August | Hymenoptera | Braconidae | Bracon |
| 2508AshExt6b | AshExt | August | Hymenoptera | Braconidae | Chorebus |
| 2508AshExt6b | AshExt | August | Hymenoptera | Ichneumonidae | Megastylus |
| 2508AshExt6b | AshExt | August | Hymenoptera | Ichneumonidae | Polyblastus |
| 2508AshExt6b | AshExt | August | Hymenoptera | Other | Other |
| 2508AshExt6b | AshExt | August | Hymenoptera | Tenthredinidae | Dolerus |
| 2508AshExt6b | AshExt | August | Hymenoptera | Tenthredinidae | Nematus |
| 2508AshExt6b | AshExt | August | Hymenoptera | Trigonalidae | Orthogonalys |
| 2508AshExt6b | AshExt | August | Lepidoptera | Crambidae | Blepharomastix |
| 2508AshExt6b | AshExt | August | Lepidoptera | Crambidae | Parapoynx |
| 2508AshExt6b | AshExt | August | Lepidoptera | Elachistidae | Elachista |
| 2508AshExt6b | AshExt | August | Lepidoptera | Erebidae | Panopoda |
| 2508AshExt6b | AshExt | August | Lepidoptera | Gelechiidae | Chionodes |
| 2508AshExt6b | AshExt | August | Lepidoptera | Noctuidae | Condica |

| | | | | | |
|---|---|---|---|---|---|
| 2508AshExt6b | AshExt | August | Lepidoptera | Noctuidae | Leuconycta |
| 2508AshExt6b | AshExt | August | Lepidoptera | Noctuidae | Pyreferra |
| 2508AshExt6b | AshExt | August | Lepidoptera | Notodontidae | Heterocampa |
| 2508AshExt6b | AshExt | August | Lepidoptera | Notodontidae | Hyparpax |
| 2508AshExt6b | AshExt | August | Lepidoptera | Nymphalidae | Oeneis |
| 2508AshExt6b | AshExt | August | Lepidoptera | Pyralidae | Atheloca |
| 2508AshExt6b | AshExt | August | Lepidoptera | Pyralidae | Elasmopalpus |
| 2508AshExt6b | AshExt | August | Lepidoptera | Tischeriidae | Coptotriche |
| 2508AshExt6b | AshExt | August | Lepidoptera | Tortricidae | Acleris |
| 2508AshExt6b | AshExt | August | Lepidoptera | Tortricidae | Argyrotaenia |
| 2508AshExt6b | AshExt | August | Lepidoptera | Tortricidae | Rhyacionia |
| 2508AshExt6b | AshExt | August | Megaloptera | Corydalidae | Corydalus |
| 2508AshExt6b | AshExt | August | Neuroptera | Hemerobiidae | Hemerobius |
| 2508AshExt6b | AshExt | August | Odonata | Libellulidae | Brachymesia |
| 2508AshExt6b | AshExt | August | Orthoptera | Tettigoniidae | Conocephalus |
| 2508AshExt6b | AshExt | August | Other | Other | Other |
| 2508AshExt6b | AshExt | August | Other | Other | Other |
| 2508AshExt6b | AshExt | August | Plecoptera | Chloroperlidae | Sweltsa |
| 2508AshExt6b | AshExt | August | Psocodea | Peripsocidae | Peripsocus |
| 2508AshExt6b | AshExt | August | Sarcoptiformes | Oripodidae | Other |
| 2508AshExt6b | AshExt | August | Trichoptera | Dipseudopsidae | Phylocentropus |
| 2508AshExt6b | AshExt | August | Trichoptera | Hydroptilidae | Orthotrichia |
| 2508AshExt6b | AshExt | August | Trichoptera | Leptoceridae | Oecetis |
| 2508AshExt6b | AshExt | August | Trombidiformes | Arrenuridae | Arrenurus |
| 2508AshExt6b | AshExt | August | Trombidiformes | Aturidae | Aturus |
| 2508AshExt6b | AshExt | August | Trombidiformes | Axonopsidae | Brachypoda |
| 2508AshExt6b | AshExt | August | Trombidiformes | Eupodidae | Other |
| 2508AshExt6b | AshExt | August | Trombidiformes | Feltriidae | Feltria |
| 2508AshExt6b | AshExt | August | Trombidiformes | Hydryphantidae | Hydryphantes |
| 2508AshExt6b | AshExt | August | Trombidiformes | Hydryphantidae | Parathyas |
| 2508AshExt6b | AshExt | August | Trombidiformes | Lebertiidae | Lebertia |

| | | | | | |
|---|---|---|---|---|---|
| 2508AshExt6b | AshExt | August | Trombidiformes | Mideopsidae | Mideopsis |
| 2508AshExt6b | AshExt | August | Trombidiformes | Pionidae | Piona |
| 2508AshExt6b | AshExt | August | Trombidiformes | Pionidae | Tiphys |
| 2508AshExt6b | AshExt | August | Trombidiformes | Sperchontidae | Sperchon |
| 2508AshExt6b | AshExt | August | Trombidiformes | Tarsonemidae | Phytonemus |
| 2508AshExt6b | AshExt | August | Trombidiformes | Torrenticolidae | Testudacarus |
| 2508AshExt6b | AshExt | August | Trombidiformes | Torrenticolidae | Torrenticola |
| 2508AshExt6b | AshExt | August | Trombidiformes | Unionicolidae | Koenikea |
| 2508AshExt6c | AshExt | August | Araneae | Agelenidae | Coras |
| 2508AshExt6c | AshExt | August | Araneae | Amaurobiidae | Callobius |
| 2508AshExt6c | AshExt | August | Araneae | Anyphaenidae | Anyphaena |
| 2508AshExt6c | AshExt | August | Araneae | Cicurinidae | Cicurina |
| 2508AshExt6c | AshExt | August | Araneae | Dictynidae | Dictyna |
| 2508AshExt6c | AshExt | August | Araneae | Gnaphosidae | Gnaphosa |
| 2508AshExt6c | AshExt | August | Araneae | Linyphiidae | Grammonota |
| 2508AshExt6c | AshExt | August | Araneae | Lycosidae | Rabidosa |
| 2508AshExt6c | AshExt | August | Araneae | Lycosidae | Schizocosa |
| 2508AshExt6c | AshExt | August | Araneae | Phrurolithidae | Scotinella |
| 2508AshExt6c | AshExt | August | Araneae | Salticidae | Maevia |
| 2508AshExt6c | AshExt | August | Araneae | Salticidae | Sibianor |
| 2508AshExt6c | AshExt | August | Araneae | Theridiidae | Canalidion |
| 2508AshExt6c | AshExt | August | Araneae | Theridiidae | Rugathodes |
| 2508AshExt6c | AshExt | August | Araneae | Theridiidae | Spintharus |
| 2508AshExt6c | AshExt | August | Araneae | Thomisidae | Mecaphesa |
| 2508AshExt6c | AshExt | August | Blattodea | Ectobiidae | Parcoblatta |
| 2508AshExt6c | AshExt | August | Coleoptera | Buprestidae | Pachyschelus |
| 2508AshExt6c | AshExt | August | Coleoptera | Carabidae | Apenes |
| 2508AshExt6c | AshExt | August | Coleoptera | Carabidae | Harpalus |
| 2508AshExt6c | AshExt | August | Coleoptera | Carabidae | Other |
| 2508AshExt6c | AshExt | August | Coleoptera | Carabidae | Polyderis |
| 2508AshExt6c | AshExt | August | Coleoptera | Curculionidae | Conotrachelus |

| | | | | | |
|---|---|---|---|---|---|
| 2508AshExt6c | AshExt | August | Coleoptera | Curculionidae | Cyrtepistomus |
| 2508AshExt6c | AshExt | August | Coleoptera | Curculionidae | Listronotus |
| 2508AshExt6c | AshExt | August | Coleoptera | Dytiscidae | Acilius |
| 2508AshExt6c | AshExt | August | Coleoptera | Lampyridae | Photinus |
| 2508AshExt6c | AshExt | August | Coleoptera | Scarabaeidae | Dialytes |
| 2508AshExt6c | AshExt | August | Coleoptera | Scarabaeidae | Melanocanthon |
| 2508AshExt6c | AshExt | August | Coleoptera | Scirtidae | Contacyphon |
| 2508AshExt6c | AshExt | August | Diptera | Agromyzidae | Calycomyza |
| 2508AshExt6c | AshExt | August | Diptera | Agromyzidae | Cerodontha |
| 2508AshExt6c | AshExt | August | Diptera | Agromyzidae | Japanagromyza |
| 2508AshExt6c | AshExt | August | Diptera | Agromyzidae | Liriomyza |
| 2508AshExt6c | AshExt | August | Diptera | Agromyzidae | Nemorimyza |
| 2508AshExt6c | AshExt | August | Diptera | Agromyzidae | Phytomyza |
| 2508AshExt6c | AshExt | August | Diptera | Anisopodidae | Mycetobia |
| 2508AshExt6c | AshExt | August | Diptera | Calliphoridae | Lucilia |
| 2508AshExt6c | AshExt | August | Diptera | Cecidomyiidae | Other |
| 2508AshExt6c | AshExt | August | Diptera | Chaoboridae | Chaoborus |
| 2508AshExt6c | AshExt | August | Diptera | Chaoboridae | Other |
| 2508AshExt6c | AshExt | August | Diptera | Chironomidae | Ablabesmyia |
| 2508AshExt6c | AshExt | August | Diptera | Chironomidae | Chironomus |
| 2508AshExt6c | AshExt | August | Diptera | Chironomidae | Dicrotendipes |
| 2508AshExt6c | AshExt | August | Diptera | Chironomidae | Endochironomus |
| 2508AshExt6c | AshExt | August | Diptera | Chironomidae | Glyptotendipes |
| 2508AshExt6c | AshExt | August | Diptera | Chironomidae | Microtendipes |
| 2508AshExt6c | AshExt | August | Diptera | Chironomidae | Orthocladius |
| 2508AshExt6c | AshExt | August | Diptera | Chironomidae | Other |
| 2508AshExt6c | AshExt | August | Diptera | Chironomidae | Other |
| 2508AshExt6c | AshExt | August | Diptera | Chironomidae | Other |
| 2508AshExt6c | AshExt | August | Diptera | Chironomidae | Other |
| 2508AshExt6c | AshExt | August | Diptera | Chironomidae | Parachironomus |
| 2508AshExt6c | AshExt | August | Diptera | Chironomidae | Polypedilum |

| | | | | | |
|---|---|---|---|---|---|
| 2508AshExt6c | AshExt | August | Diptera | Chironomidae | Pseudochironomus |
| 2508AshExt6c | AshExt | August | Diptera | Chironomidae | Tanytarsus |
| 2508AshExt6c | AshExt | August | Diptera | Chironomidae | Xenochironomus |
| 2508AshExt6c | AshExt | August | Diptera | Chloropidae | Apotropina |
| 2508AshExt6c | AshExt | August | Diptera | Chloropidae | Elachiptera |
| 2508AshExt6c | AshExt | August | Diptera | Chloropidae | Liohippelates |
| 2508AshExt6c | AshExt | August | Diptera | Chloropidae | Oscinella |
| 2508AshExt6c | AshExt | August | Diptera | Culicidae | Coquillettidia |
| 2508AshExt6c | AshExt | August | Diptera | Culicidae | Culex |
| 2508AshExt6c | AshExt | August | Diptera | Culicidae | Uranotaenia |
| 2508AshExt6c | AshExt | August | Diptera | Drosophilidae | Drosophila |
| 2508AshExt6c | AshExt | August | Diptera | Empididae | Other |
| 2508AshExt6c | AshExt | August | Diptera | Hybotidae | Platypalpus |
| 2508AshExt6c | AshExt | August | Diptera | Hybotidae | Syndyas |
| 2508AshExt6c | AshExt | August | Diptera | Keroplatidae | Other |
| 2508AshExt6c | AshExt | August | Diptera | Limoniidae | Dicranomyia |
| 2508AshExt6c | AshExt | August | Diptera | Limoniidae | Dicranophragma |
| 2508AshExt6c | AshExt | August | Diptera | Limoniidae | Epiphragma |
| 2508AshExt6c | AshExt | August | Diptera | Limoniidae | Erioptera |
| 2508AshExt6c | AshExt | August | Diptera | Limoniidae | Metalimnobia |
| 2508AshExt6c | AshExt | August | Diptera | Limoniidae | Ormosia |
| 2508AshExt6c | AshExt | August | Diptera | Limoniidae | Other |
| 2508AshExt6c | AshExt | August | Diptera | Limoniidae | Other |
| 2508AshExt6c | AshExt | August | Diptera | Limoniidae | Pseudolimnophila |
| 2508AshExt6c | AshExt | August | Diptera | Muscidae | Mydaea |
| 2508AshExt6c | AshExt | August | Diptera | Mycetophilidae | Mycomya |
| 2508AshExt6c | AshExt | August | Diptera | Mycetophilidae | Neoempheria |
| 2508AshExt6c | AshExt | August | Diptera | Opomyzidae | Geomyza |
| 2508AshExt6c | AshExt | August | Diptera | Pallopteridae | Toxonevra |
| 2508AshExt6c | AshExt | August | Diptera | Phoridae | Diplonevra |
| 2508AshExt6c | AshExt | August | Diptera | Polleniidae | Pollenia |

| | | | | | |
|---|---|---|---|---|---|
| 2508AshExt6c | AshExt | August | Diptera | Psychodidae | Clogmia |
| 2508AshExt6c | AshExt | August | Diptera | Psychodidae | Psychoda |
| 2508AshExt6c | AshExt | August | Diptera | Sarcophagidae | Ravinia |
| 2508AshExt6c | AshExt | August | Diptera | Sciaridae | Claustropyga |
| 2508AshExt6c | AshExt | August | Diptera | Sciaridae | Corynoptera |
| 2508AshExt6c | AshExt | August | Diptera | Sciaridae | Epidapus |
| 2508AshExt6c | AshExt | August | Diptera | Sciaridae | Other |
| 2508AshExt6c | AshExt | August | Diptera | Sphaeroceridae | Leptocera |
| 2508AshExt6c | AshExt | August | Diptera | Syrphidae | Chalcosyrphus |
| 2508AshExt6c | AshExt | August | Diptera | Syrphidae | Palpada |
| 2508AshExt6c | AshExt | August | Diptera | Tabanidae | Hybomitra |
| 2508AshExt6c | AshExt | August | Diptera | Tabanidae | Other |
| 2508AshExt6c | AshExt | August | Diptera | Tachinidae | Cryptomeigenia |
| 2508AshExt6c | AshExt | August | Diptera | Tachinidae | Cyzenis |
| 2508AshExt6c | AshExt | August | Diptera | Tachinidae | Zaira |
| 2508AshExt6c | AshExt | August | Diptera | Tipulidae | Dolichopeza |
| 2508AshExt6c | AshExt | August | Diptera | Tipulidae | Tipula |
| 2508AshExt6c | AshExt | August | Ephemeroptera | Caenidae | Caenis |
| 2508AshExt6c | AshExt | August | Ephemeroptera | Ephemeridae | Hexagenia |
| 2508AshExt6c | AshExt | August | Ephemeroptera | Heptageniidae | Epeorus |
| 2508AshExt6c | AshExt | August | Hemiptera | Clastopteridae | Clastoptera |
| 2508AshExt6c | AshExt | August | Hemiptera | Lygaeidae | Kleidocerys |
| 2508AshExt6c | AshExt | August | Hymenoptera | Apidae | Triepeolus |
| 2508AshExt6c | AshExt | August | Hymenoptera | Braconidae | Bracon |
| 2508AshExt6c | AshExt | August | Hymenoptera | Braconidae | Chorebus |
| 2508AshExt6c | AshExt | August | Hymenoptera | Ichneumonidae | Megastylus |
| 2508AshExt6c | AshExt | August | Hymenoptera | Ichneumonidae | Polyblastus |
| 2508AshExt6c | AshExt | August | Hymenoptera | Other | Other |
| 2508AshExt6c | AshExt | August | Hymenoptera | Tenthredinidae | Dolerus |
| 2508AshExt6c | AshExt | August | Hymenoptera | Tenthredinidae | Nematus |
| 2508AshExt6c | AshExt | August | Hymenoptera | Trigonalidae | Orthogonalys |

| | | | | | |
|---|---|---|---|---|---|
| 2508AshExt6c | AshExt | August | Lepidoptera | Crambidae | Blepharomastix |
| 2508AshExt6c | AshExt | August | Lepidoptera | Crambidae | Parapoynx |
| 2508AshExt6c | AshExt | August | Lepidoptera | Elachistidae | Elachista |
| 2508AshExt6c | AshExt | August | Lepidoptera | Erebidae | Panopoda |
| 2508AshExt6c | AshExt | August | Lepidoptera | Gelechiidae | Chionodes |
| 2508AshExt6c | AshExt | August | Lepidoptera | Noctuidae | Condica |
| 2508AshExt6c | AshExt | August | Lepidoptera | Noctuidae | Leuconycta |
| 2508AshExt6c | AshExt | August | Lepidoptera | Noctuidae | Pyreferra |
| 2508AshExt6c | AshExt | August | Lepidoptera | Notodontidae | Heterocampa |
| 2508AshExt6c | AshExt | August | Lepidoptera | Notodontidae | Hyparpax |
| 2508AshExt6c | AshExt | August | Lepidoptera | Nymphalidae | Oeneis |
| 2508AshExt6c | AshExt | August | Lepidoptera | Pyralidae | Atheloca |
| 2508AshExt6c | AshExt | August | Lepidoptera | Pyralidae | Elasmopalpus |
| 2508AshExt6c | AshExt | August | Lepidoptera | Tischeriidae | Coptotriche |
| 2508AshExt6c | AshExt | August | Lepidoptera | Tortricidae | Acleris |
| 2508AshExt6c | AshExt | August | Lepidoptera | Tortricidae | Argyrotaenia |
| 2508AshExt6c | AshExt | August | Lepidoptera | Tortricidae | Rhyacionia |
| 2508AshExt6c | AshExt | August | Megaloptera | Corydalidae | Corydalus |
| 2508AshExt6c | AshExt | August | Neuroptera | Hemerobiidae | Hemerobius |
| 2508AshExt6c | AshExt | August | Odonata | Libellulidae | Brachymesia |
| 2508AshExt6c | AshExt | August | Orthoptera | Tettigoniidae | Conocephalus |
| 2508AshExt6c | AshExt | August | Other | Other | Other |
| 2508AshExt6c | AshExt | August | Other | Other | Other |
| 2508AshExt6c | AshExt | August | Plecoptera | Chloroperlidae | Sweltsa |
| 2508AshExt6c | AshExt | August | Psocodea | Peripsocidae | Peripsocus |
| 2508AshExt6c | AshExt | August | Sarcoptiformes | Oripodidae | Other |
| 2508AshExt6c | AshExt | August | Trichoptera | Dipseudopsidae | Phylocentropus |
| 2508AshExt6c | AshExt | August | Trichoptera | Hydroptilidae | Orthotrichia |
| 2508AshExt6c | AshExt | August | Trichoptera | Leptoceridae | Oecetis |
| 2508AshExt6c | AshExt | August | Trombidiformes | Arrenuridae | Arrenurus |
| 2508AshExt6c | AshExt | August | Trombidiformes | Aturidae | Aturus |

| | | | | | |
|---|---|---|---|---|---|
| 2508AshExt6c | AshExt | August | Trombidiformes | Axonopsidae | Brachypoda |
| 2508AshExt6c | AshExt | August | Trombidiformes | Eupodidae | Other |
| 2508AshExt6c | AshExt | August | Trombidiformes | Feltriidae | Feltria |
| 2508AshExt6c | AshExt | August | Trombidiformes | Hydryphantidae | Hydryphantes |
| 2508AshExt6c | AshExt | August | Trombidiformes | Hydryphantidae | Parathyas |
| 2508AshExt6c | AshExt | August | Trombidiformes | Lebertiidae | Lebertia |
| 2508AshExt6c | AshExt | August | Trombidiformes | Mideopsidae | Mideopsis |
| 2508AshExt6c | AshExt | August | Trombidiformes | Pionidae | Piona |
| 2508AshExt6c | AshExt | August | Trombidiformes | Pionidae | Tiphys |
| 2508AshExt6c | AshExt | August | Trombidiformes | Sperchontidae | Sperchon |
| 2508AshExt6c | AshExt | August | Trombidiformes | Tarsonemidae | Phytonemus |
| 2508AshExt6c | AshExt | August | Trombidiformes | Torrenticolidae | Testudacarus |
| 2508AshExt6c | AshExt | August | Trombidiformes | Torrenticolidae | Torrenticola |
| 2508AshExt6c | AshExt | August | Trombidiformes | Unionicolidae | Koenikea |
| 2505Wmcc6a | WMCC | May | Araneae | Agelenidae | Coras |
| 2505Wmcc6a | WMCC | May | Araneae | Amaurobiidae | Callobius |
| 2505Wmcc6a | WMCC | May | Araneae | Anyphaenidae | Anyphaena |
| 2505Wmcc6a | WMCC | May | Araneae | Cicurinidae | Cicurina |
| 2505Wmcc6a | WMCC | May | Araneae | Dictynidae | Dictyna |
| 2505Wmcc6a | WMCC | May | Araneae | Gnaphosidae | Gnaphosa |
| 2505Wmcc6a | WMCC | May | Araneae | Linyphiidae | Grammonota |
| 2505Wmcc6a | WMCC | May | Araneae | Lycosidae | Rabidosa |
| 2505Wmcc6a | WMCC | May | Araneae | Lycosidae | Schizocosa |
| 2505Wmcc6a | WMCC | May | Araneae | Phrurolithidae | Scotinella |
| 2505Wmcc6a | WMCC | May | Araneae | Salticidae | Maevia |
| 2505Wmcc6a | WMCC | May | Araneae | Salticidae | Sibianor |
| 2505Wmcc6a | WMCC | May | Araneae | Theridiidae | Canalidion |
| 2505Wmcc6a | WMCC | May | Araneae | Theridiidae | Rugathodes |
| 2505Wmcc6a | WMCC | May | Araneae | Theridiidae | Spintharus |
| 2505Wmcc6a | WMCC | May | Araneae | Thomisidae | Mecaphesa |
| 2505Wmcc6a | WMCC | May | Blattodea | Ectobiidae | Parcoblatta |

| | | | | | |
|---|---|---|---|---|---|
| 2505Wmcc6a | WMCC | May | Coleoptera | Buprestidae | Pachyschelus |
| 2505Wmcc6a | WMCC | May | Coleoptera | Carabidae | Apenes |
| 2505Wmcc6a | WMCC | May | Coleoptera | Carabidae | Harpalus |
| 2505Wmcc6a | WMCC | May | Coleoptera | Carabidae | Other |
| 2505Wmcc6a | WMCC | May | Coleoptera | Carabidae | Polyderis |
| 2505Wmcc6a | WMCC | May | Coleoptera | Curculionidae | Conotrachelus |
| 2505Wmcc6a | WMCC | May | Coleoptera | Curculionidae | Cyrtepistomus |
| 2505Wmcc6a | WMCC | May | Coleoptera | Curculionidae | Listronotus |
| 2505Wmcc6a | WMCC | May | Coleoptera | Dytiscidae | Acilius |
| 2505Wmcc6a | WMCC | May | Coleoptera | Lampyridae | Photinus |
| 2505Wmcc6a | WMCC | May | Coleoptera | Scarabaeidae | Dialytes |
| 2505Wmcc6a | WMCC | May | Coleoptera | Scarabaeidae | Melanocanthon |
| 2505Wmcc6a | WMCC | May | Coleoptera | Scirtidae | Contacyphon |
| 2505Wmcc6a | WMCC | May | Diptera | Agromyzidae | Calycomyza |
| 2505Wmcc6a | WMCC | May | Diptera | Agromyzidae | Cerodontha |
| 2505Wmcc6a | WMCC | May | Diptera | Agromyzidae | Japanagromyza |
| 2505Wmcc6a | WMCC | May | Diptera | Agromyzidae | Liriomyza |
| 2505Wmcc6a | WMCC | May | Diptera | Agromyzidae | Nemorimyza |
| 2505Wmcc6a | WMCC | May | Diptera | Agromyzidae | Phytomyza |
| 2505Wmcc6a | WMCC | May | Diptera | Anisopodidae | Mycetobia |
| 2505Wmcc6a | WMCC | May | Diptera | Calliphoridae | Lucilia |
| 2505Wmcc6a | WMCC | May | Diptera | Cecidomyiidae | Other |
| 2505Wmcc6a | WMCC | May | Diptera | Chaoboridae | Chaoborus |
| 2505Wmcc6a | WMCC | May | Diptera | Chaoboridae | Other |
| 2505Wmcc6a | WMCC | May | Diptera | Chironomidae | Ablabesmyia |
| 2505Wmcc6a | WMCC | May | Diptera | Chironomidae | Chironomus |
| 2505Wmcc6a | WMCC | May | Diptera | Chironomidae | Dicrotendipes |
| 2505Wmcc6a | WMCC | May | Diptera | Chironomidae | Endochironomus |
| 2505Wmcc6a | WMCC | May | Diptera | Chironomidae | Glyptotendipes |
| 2505Wmcc6a | WMCC | May | Diptera | Chironomidae | Microtendipes |
| 2505Wmcc6a | WMCC | May | Diptera | Chironomidae | Orthocladius |

| | | | | | |
|---|---|---|---|---|---|
| 2505Wmcc6a | WMCC | May | Diptera | Chironomidae | Other |
| 2505Wmcc6a | WMCC | May | Diptera | Chironomidae | Other |
| 2505Wmcc6a | WMCC | May | Diptera | Chironomidae | Other |
| 2505Wmcc6a | WMCC | May | Diptera | Chironomidae | Other |
| 2505Wmcc6a | WMCC | May | Diptera | Chironomidae | Parachironomus |
| 2505Wmcc6a | WMCC | May | Diptera | Chironomidae | Polypedilum |
| 2505Wmcc6a | WMCC | May | Diptera | Chironomidae | Pseudochironomus |
| 2505Wmcc6a | WMCC | May | Diptera | Chironomidae | Tanytarsus |
| 2505Wmcc6a | WMCC | May | Diptera | Chironomidae | Xenochironomus |
| 2505Wmcc6a | WMCC | May | Diptera | Chloropidae | Apotropina |
| 2505Wmcc6a | WMCC | May | Diptera | Chloropidae | Elachiptera |
| 2505Wmcc6a | WMCC | May | Diptera | Chloropidae | Liohippelates |
| 2505Wmcc6a | WMCC | May | Diptera | Chloropidae | Oscinella |
| 2505Wmcc6a | WMCC | May | Diptera | Culicidae | Coquillettidia |
| 2505Wmcc6a | WMCC | May | Diptera | Culicidae | Culex |
| 2505Wmcc6a | WMCC | May | Diptera | Culicidae | Uranotaenia |
| 2505Wmcc6a | WMCC | May | Diptera | Drosophilidae | Drosophila |
| 2505Wmcc6a | WMCC | May | Diptera | Empididae | Other |
| 2505Wmcc6a | WMCC | May | Diptera | Hybotidae | Platypalpus |
| 2505Wmcc6a | WMCC | May | Diptera | Hybotidae | Syndyas |
| 2505Wmcc6a | WMCC | May | Diptera | Keroplatidae | Other |
| 2505Wmcc6a | WMCC | May | Diptera | Limoniidae | Dicranomyia |
| 2505Wmcc6a | WMCC | May | Diptera | Limoniidae | Dicranophragma |
| 2505Wmcc6a | WMCC | May | Diptera | Limoniidae | Epiphragma |
| 2505Wmcc6a | WMCC | May | Diptera | Limoniidae | Erioptera |
| 2505Wmcc6a | WMCC | May | Diptera | Limoniidae | Metalimnobia |
| 2505Wmcc6a | WMCC | May | Diptera | Limoniidae | Ormosia |
| 2505Wmcc6a | WMCC | May | Diptera | Limoniidae | Other |
| 2505Wmcc6a | WMCC | May | Diptera | Limoniidae | Other |
| 2505Wmcc6a | WMCC | May | Diptera | Limoniidae | Pseudolimnophila |
| 2505Wmcc6a | WMCC | May | Diptera | Muscidae | Mydaea |

| | | | | | |
|---|---|---|---|---|---|
| 2505Wmcc6a | WMCC | May | Diptera | Mycetophilidae | Mycomya |
| 2505Wmcc6a | WMCC | May | Diptera | Mycetophilidae | Neoempheria |
| 2505Wmcc6a | WMCC | May | Diptera | Opomyzidae | Geomyza |
| 2505Wmcc6a | WMCC | May | Diptera | Pallopteridae | Toxonevra |
| 2505Wmcc6a | WMCC | May | Diptera | Phoridae | Diplonevra |
| 2505Wmcc6a | WMCC | May | Diptera | Polleniidae | Pollenia |
| 2505Wmcc6a | WMCC | May | Diptera | Psychodidae | Clogmia |
| 2505Wmcc6a | WMCC | May | Diptera | Psychodidae | Psychoda |
| 2505Wmcc6a | WMCC | May | Diptera | Sarcophagidae | Ravinia |
| 2505Wmcc6a | WMCC | May | Diptera | Sciaridae | Claustropyga |
| 2505Wmcc6a | WMCC | May | Diptera | Sciaridae | Corynoptera |
| 2505Wmcc6a | WMCC | May | Diptera | Sciaridae | Epidapus |
| 2505Wmcc6a | WMCC | May | Diptera | Sciaridae | Other |
| 2505Wmcc6a | WMCC | May | Diptera | Sphaeroceridae | Leptocera |
| 2505Wmcc6a | WMCC | May | Diptera | Syrphidae | Chalcosyrphus |
| 2505Wmcc6a | WMCC | May | Diptera | Syrphidae | Palpada |
| 2505Wmcc6a | WMCC | May | Diptera | Tabanidae | Hybomitra |
| 2505Wmcc6a | WMCC | May | Diptera | Tabanidae | Other |
| 2505Wmcc6a | WMCC | May | Diptera | Tachinidae | Cryptomeigenia |
| 2505Wmcc6a | WMCC | May | Diptera | Tachinidae | Cyzenis |
| 2505Wmcc6a | WMCC | May | Diptera | Tachinidae | Zaira |
| 2505Wmcc6a | WMCC | May | Diptera | Tipulidae | Dolichopeza |
| 2505Wmcc6a | WMCC | May | Diptera | Tipulidae | Tipula |
| 2505Wmcc6a | WMCC | May | Ephemeroptera | Caenidae | Caenis |
| 2505Wmcc6a | WMCC | May | Ephemeroptera | Ephemeridae | Hexagenia |
| 2505Wmcc6a | WMCC | May | Ephemeroptera | Heptageniidae | Epeorus |
| 2505Wmcc6a | WMCC | May | Hemiptera | Clastopteridae | Clastoptera |
| 2505Wmcc6a | WMCC | May | Hemiptera | Lygaeidae | Kleidocerys |
| 2505Wmcc6a | WMCC | May | Hymenoptera | Apidae | Triepeolus |
| 2505Wmcc6a | WMCC | May | Hymenoptera | Braconidae | Bracon |
| 2505Wmcc6a | WMCC | May | Hymenoptera | Braconidae | Chorebus |

| | | | | | |
|---|---|---|---|---|---|
| 2505Wmcc6a | WMCC | May | Hymenoptera | Ichneumonidae | Megastylus |
| 2505Wmcc6a | WMCC | May | Hymenoptera | Ichneumonidae | Polyblastus |
| 2505Wmcc6a | WMCC | May | Hymenoptera | Other | Other |
| 2505Wmcc6a | WMCC | May | Hymenoptera | Tenthredinidae | Dolerus |
| 2505Wmcc6a | WMCC | May | Hymenoptera | Tenthredinidae | Nematus |
| 2505Wmcc6a | WMCC | May | Hymenoptera | Trigonalidae | Orthogonalys |
| 2505Wmcc6a | WMCC | May | Lepidoptera | Crambidae | Blepharomastix |
| 2505Wmcc6a | WMCC | May | Lepidoptera | Crambidae | Parapoynx |
| 2505Wmcc6a | WMCC | May | Lepidoptera | Elachistidae | Elachista |
| 2505Wmcc6a | WMCC | May | Lepidoptera | Erebidae | Panopoda |
| 2505Wmcc6a | WMCC | May | Lepidoptera | Gelechiidae | Chionodes |
| 2505Wmcc6a | WMCC | May | Lepidoptera | Noctuidae | Condica |
| 2505Wmcc6a | WMCC | May | Lepidoptera | Noctuidae | Leuconycta |
| 2505Wmcc6a | WMCC | May | Lepidoptera | Noctuidae | Pyreferra |
| 2505Wmcc6a | WMCC | May | Lepidoptera | Notodontidae | Heterocampa |
| 2505Wmcc6a | WMCC | May | Lepidoptera | Notodontidae | Hyparpax |
| 2505Wmcc6a | WMCC | May | Lepidoptera | Nymphalidae | Oeneis |
| 2505Wmcc6a | WMCC | May | Lepidoptera | Pyralidae | Atheloca |
| 2505Wmcc6a | WMCC | May | Lepidoptera | Pyralidae | Elasmopalpus |
| 2505Wmcc6a | WMCC | May | Lepidoptera | Tischeriidae | Coptotriche |
| 2505Wmcc6a | WMCC | May | Lepidoptera | Tortricidae | Acleris |
| 2505Wmcc6a | WMCC | May | Lepidoptera | Tortricidae | Argyrotaenia |
| 2505Wmcc6a | WMCC | May | Lepidoptera | Tortricidae | Rhyacionia |
| 2505Wmcc6a | WMCC | May | Megaloptera | Corydalidae | Corydalus |
| 2505Wmcc6a | WMCC | May | Neuroptera | Hemerobiidae | Hemerobius |
| 2505Wmcc6a | WMCC | May | Odonata | Libellulidae | Brachymesia |
| 2505Wmcc6a | WMCC | May | Orthoptera | Tettigoniidae | Conocephalus |
| 2505Wmcc6a | WMCC | May | Other | Other | Other |
| 2505Wmcc6a | WMCC | May | Other | Other | Other |
| 2505Wmcc6a | WMCC | May | Plecoptera | Chloroperlidae | Sweltsa |
| 2505Wmcc6a | WMCC | May | Psocodea | Peripsocidae | Peripsocus |

| | | | | | |
|---|---|---|---|---|---|
| 2505Wmcc6a | WMCC | May | Sarcoptiformes | Oripodidae | Other |
| 2505Wmcc6a | WMCC | May | Trichoptera | Dipseudopsidae | Phylocentropus |
| 2505Wmcc6a | WMCC | May | Trichoptera | Hydroptilidae | Orthotrichia |
| 2505Wmcc6a | WMCC | May | Trichoptera | Leptoceridae | Oecetis |
| 2505Wmcc6a | WMCC | May | Trombidiformes | Arrenuridae | Arrenurus |
| 2505Wmcc6a | WMCC | May | Trombidiformes | Aturidae | Aturus |
| 2505Wmcc6a | WMCC | May | Trombidiformes | Axonopsidae | Brachypoda |
| 2505Wmcc6a | WMCC | May | Trombidiformes | Eupodidae | Other |
| 2505Wmcc6a | WMCC | May | Trombidiformes | Feltriidae | Feltria |
| 2505Wmcc6a | WMCC | May | Trombidiformes | Hydryphantidae | Hydryphantes |
| 2505Wmcc6a | WMCC | May | Trombidiformes | Hydryphantidae | Parathyas |
| 2505Wmcc6a | WMCC | May | Trombidiformes | Lebertiidae | Lebertia |
| 2505Wmcc6a | WMCC | May | Trombidiformes | Mideopsidae | Mideopsis |
| 2505Wmcc6a | WMCC | May | Trombidiformes | Pionidae | Piona |
| 2505Wmcc6a | WMCC | May | Trombidiformes | Pionidae | Tiphys |
| 2505Wmcc6a | WMCC | May | Trombidiformes | Sperchontidae | Sperchon |
| 2505Wmcc6a | WMCC | May | Trombidiformes | Tarsonemidae | Phytonemus |
| 2505Wmcc6a | WMCC | May | Trombidiformes | Torrenticolidae | Testudacarus |
| 2505Wmcc6a | WMCC | May | Trombidiformes | Torrenticolidae | Torrenticola |
| 2505Wmcc6a | WMCC | May | Trombidiformes | Unionicolidae | Koenikea |
| 2505Wmcc6c | WMCC | May | Araneae | Agelenidae | Coras |
| 2505Wmcc6c | WMCC | May | Araneae | Amaurobiidae | Callobius |
| 2505Wmcc6c | WMCC | May | Araneae | Anyphaenidae | Anyphaena |
| 2505Wmcc6c | WMCC | May | Araneae | Cicurinidae | Cicurina |
| 2505Wmcc6c | WMCC | May | Araneae | Dictynidae | Dictyna |
| 2505Wmcc6c | WMCC | May | Araneae | Gnaphosidae | Gnaphosa |
| 2505Wmcc6c | WMCC | May | Araneae | Linyphiidae | Grammonota |
| 2505Wmcc6c | WMCC | May | Araneae | Lycosidae | Rabidosa |
| 2505Wmcc6c | WMCC | May | Araneae | Lycosidae | Schizocosa |
| 2505Wmcc6c | WMCC | May | Araneae | Phrurolithidae | Scotinella |
| 2505Wmcc6c | WMCC | May | Araneae | Salticidae | Maevia |

| | | | | | |
|---|---|---|---|---|---|
| 2505Wmcc6c | WMCC | May | Araneae | Salticidae | Sibianor |
| 2505Wmcc6c | WMCC | May | Araneae | Theridiidae | Canalidion |
| 2505Wmcc6c | WMCC | May | Araneae | Theridiidae | Rugathodes |
| 2505Wmcc6c | WMCC | May | Araneae | Theridiidae | Spintharus |
| 2505Wmcc6c | WMCC | May | Araneae | Thomisidae | Mecaphesa |
| 2505Wmcc6c | WMCC | May | Blattodea | Ectobiidae | Parcoblatta |
| 2505Wmcc6c | WMCC | May | Coleoptera | Buprestidae | Pachyschelus |
| 2505Wmcc6c | WMCC | May | Coleoptera | Carabidae | Apenes |
| 2505Wmcc6c | WMCC | May | Coleoptera | Carabidae | Harpalus |
| 2505Wmcc6c | WMCC | May | Coleoptera | Carabidae | Other |
| 2505Wmcc6c | WMCC | May | Coleoptera | Carabidae | Polyderis |
| 2505Wmcc6c | WMCC | May | Coleoptera | Curculionidae | Conotrachelus |
| 2505Wmcc6c | WMCC | May | Coleoptera | Curculionidae | Cyrtepistomus |
| 2505Wmcc6c | WMCC | May | Coleoptera | Curculionidae | Listronotus |
| 2505Wmcc6c | WMCC | May | Coleoptera | Dytiscidae | Acilius |
| 2505Wmcc6c | WMCC | May | Coleoptera | Lampyridae | Photinus |
| 2505Wmcc6c | WMCC | May | Coleoptera | Scarabaeidae | Dialytes |
| 2505Wmcc6c | WMCC | May | Coleoptera | Scarabaeidae | Melanocanthon |
| 2505Wmcc6c | WMCC | May | Coleoptera | Scirtidae | Contacyphon |
| 2505Wmcc6c | WMCC | May | Diptera | Agromyzidae | Calycomyza |
| 2505Wmcc6c | WMCC | May | Diptera | Agromyzidae | Cerodontha |
| 2505Wmcc6c | WMCC | May | Diptera | Agromyzidae | Japanagromyza |
| 2505Wmcc6c | WMCC | May | Diptera | Agromyzidae | Liriomyza |
| 2505Wmcc6c | WMCC | May | Diptera | Agromyzidae | Nemorimyza |
| 2505Wmcc6c | WMCC | May | Diptera | Agromyzidae | Phytomyza |
| 2505Wmcc6c | WMCC | May | Diptera | Anisopodidae | Mycetobia |
| 2505Wmcc6c | WMCC | May | Diptera | Calliphoridae | Lucilia |
| 2505Wmcc6c | WMCC | May | Diptera | Cecidomyiidae | Other |
| 2505Wmcc6c | WMCC | May | Diptera | Chaoboridae | Chaoborus |
| 2505Wmcc6c | WMCC | May | Diptera | Chaoboridae | Other |
| 2505Wmcc6c | WMCC | May | Diptera | Chironomidae | Ablabesmyia |

| | | | | | |
|---|---|---|---|---|---|
| 2505Wmcc6c | WMCC | May | Diptera | Chironomidae | Chironomus |
| 2505Wmcc6c | WMCC | May | Diptera | Chironomidae | Dicrotendipes |
| 2505Wmcc6c | WMCC | May | Diptera | Chironomidae | Endochironomus |
| 2505Wmcc6c | WMCC | May | Diptera | Chironomidae | Glyptotendipes |
| 2505Wmcc6c | WMCC | May | Diptera | Chironomidae | Microtendipes |
| 2505Wmcc6c | WMCC | May | Diptera | Chironomidae | Orthocladius |
| 2505Wmcc6c | WMCC | May | Diptera | Chironomidae | Other |
| 2505Wmcc6c | WMCC | May | Diptera | Chironomidae | Other |
| 2505Wmcc6c | WMCC | May | Diptera | Chironomidae | Other |
| 2505Wmcc6c | WMCC | May | Diptera | Chironomidae | Other |
| 2505Wmcc6c | WMCC | May | Diptera | Chironomidae | Parachironomus |
| 2505Wmcc6c | WMCC | May | Diptera | Chironomidae | Polypedilum |
| 2505Wmcc6c | WMCC | May | Diptera | Chironomidae | Pseudochironomus |
| 2505Wmcc6c | WMCC | May | Diptera | Chironomidae | Tanytarsus |
| 2505Wmcc6c | WMCC | May | Diptera | Chironomidae | Xenochironomus |
| 2505Wmcc6c | WMCC | May | Diptera | Chloropidae | Apotropina |
| 2505Wmcc6c | WMCC | May | Diptera | Chloropidae | Elachiptera |
| 2505Wmcc6c | WMCC | May | Diptera | Chloropidae | Liohippelates |
| 2505Wmcc6c | WMCC | May | Diptera | Chloropidae | Oscinella |
| 2505Wmcc6c | WMCC | May | Diptera | Culicidae | Coquillettidia |
| 2505Wmcc6c | WMCC | May | Diptera | Culicidae | Culex |
| 2505Wmcc6c | WMCC | May | Diptera | Culicidae | Uranotaenia |
| 2505Wmcc6c | WMCC | May | Diptera | Drosophilidae | Drosophila |
| 2505Wmcc6c | WMCC | May | Diptera | Empididae | Other |
| 2505Wmcc6c | WMCC | May | Diptera | Hybotidae | Platypalpus |
| 2505Wmcc6c | WMCC | May | Diptera | Hybotidae | Syndyas |
| 2505Wmcc6c | WMCC | May | Diptera | Keroplatidae | Other |
| 2505Wmcc6c | WMCC | May | Diptera | Limoniidae | Dicranomyia |
| 2505Wmcc6c | WMCC | May | Diptera | Limoniidae | Dicranophragma |
| 2505Wmcc6c | WMCC | May | Diptera | Limoniidae | Epiphragma |
| 2505Wmcc6c | WMCC | May | Diptera | Limoniidae | Erioptera |

| | | | | | |
|---|---|---|---|---|---|
| 2505Wmcc6c | WMCC | May | Diptera | Limoniidae | Metalimnobia |
| 2505Wmcc6c | WMCC | May | Diptera | Limoniidae | Ormosia |
| 2505Wmcc6c | WMCC | May | Diptera | Limoniidae | Other |
| 2505Wmcc6c | WMCC | May | Diptera | Limoniidae | Other |
| 2505Wmcc6c | WMCC | May | Diptera | Limoniidae | Pseudolimnophila |
| 2505Wmcc6c | WMCC | May | Diptera | Muscidae | Mydaea |
| 2505Wmcc6c | WMCC | May | Diptera | Mycetophilidae | Mycomya |
| 2505Wmcc6c | WMCC | May | Diptera | Mycetophilidae | Neoempheria |
| 2505Wmcc6c | WMCC | May | Diptera | Opomyzidae | Geomyza |
| 2505Wmcc6c | WMCC | May | Diptera | Pallopteridae | Toxonevra |
| 2505Wmcc6c | WMCC | May | Diptera | Phoridae | Diplonevra |
| 2505Wmcc6c | WMCC | May | Diptera | Polleniidae | Pollenia |
| 2505Wmcc6c | WMCC | May | Diptera | Psychodidae | Clogmia |
| 2505Wmcc6c | WMCC | May | Diptera | Psychodidae | Psychoda |
| 2505Wmcc6c | WMCC | May | Diptera | Sarcophagidae | Ravinia |
| 2505Wmcc6c | WMCC | May | Diptera | Sciaridae | Claustropyga |
| 2505Wmcc6c | WMCC | May | Diptera | Sciaridae | Corynoptera |
| 2505Wmcc6c | WMCC | May | Diptera | Sciaridae | Epidapus |
| 2505Wmcc6c | WMCC | May | Diptera | Sciaridae | Other |
| 2505Wmcc6c | WMCC | May | Diptera | Sphaeroceridae | Leptocera |
| 2505Wmcc6c | WMCC | May | Diptera | Syrphidae | Chalcosyrphus |
| 2505Wmcc6c | WMCC | May | Diptera | Syrphidae | Palpada |
| 2505Wmcc6c | WMCC | May | Diptera | Tabanidae | Hybomitra |
| 2505Wmcc6c | WMCC | May | Diptera | Tabanidae | Other |
| 2505Wmcc6c | WMCC | May | Diptera | Tachinidae | Cryptomeigenia |
| 2505Wmcc6c | WMCC | May | Diptera | Tachinidae | Cyzenis |
| 2505Wmcc6c | WMCC | May | Diptera | Tachinidae | Zaira |
| 2505Wmcc6c | WMCC | May | Diptera | Tipulidae | Dolichopeza |
| 2505Wmcc6c | WMCC | May | Diptera | Tipulidae | Tipula |
| 2505Wmcc6c | WMCC | May | Ephemeroptera | Caenidae | Caenis |
| 2505Wmcc6c | WMCC | May | Ephemeroptera | Ephemeridae | Hexagenia |

| | | | | | |
|---|---|---|---|---|---|
| 2505Wmcc6c | WMCC | May | Ephemeroptera | Heptageniidae | Epeorus |
| 2505Wmcc6c | WMCC | May | Hemiptera | Clastopteridae | Clastoptera |
| 2505Wmcc6c | WMCC | May | Hemiptera | Lygaeidae | Kleidocerys |
| 2505Wmcc6c | WMCC | May | Hymenoptera | Apidae | Triepeolus |
| 2505Wmcc6c | WMCC | May | Hymenoptera | Braconidae | Bracon |
| 2505Wmcc6c | WMCC | May | Hymenoptera | Braconidae | Chorebus |
| 2505Wmcc6c | WMCC | May | Hymenoptera | Ichneumonidae | Megastylus |
| 2505Wmcc6c | WMCC | May | Hymenoptera | Ichneumonidae | Polyblastus |
| 2505Wmcc6c | WMCC | May | Hymenoptera | Other | Other |
| 2505Wmcc6c | WMCC | May | Hymenoptera | Tenthredinidae | Dolerus |
| 2505Wmcc6c | WMCC | May | Hymenoptera | Tenthredinidae | Nematus |
| 2505Wmcc6c | WMCC | May | Hymenoptera | Trigonalidae | Orthogonalys |
| 2505Wmcc6c | WMCC | May | Lepidoptera | Crambidae | Blepharomastix |
| 2505Wmcc6c | WMCC | May | Lepidoptera | Crambidae | Parapoynx |
| 2505Wmcc6c | WMCC | May | Lepidoptera | Elachistidae | Elachista |
| 2505Wmcc6c | WMCC | May | Lepidoptera | Erebidae | Panopoda |
| 2505Wmcc6c | WMCC | May | Lepidoptera | Gelechiidae | Chionodes |
| 2505Wmcc6c | WMCC | May | Lepidoptera | Noctuidae | Condica |
| 2505Wmcc6c | WMCC | May | Lepidoptera | Noctuidae | Leuconycta |
| 2505Wmcc6c | WMCC | May | Lepidoptera | Noctuidae | Pyreferra |
| 2505Wmcc6c | WMCC | May | Lepidoptera | Notodontidae | Heterocampa |
| 2505Wmcc6c | WMCC | May | Lepidoptera | Notodontidae | Hyparpax |
| 2505Wmcc6c | WMCC | May | Lepidoptera | Nymphalidae | Oeneis |
| 2505Wmcc6c | WMCC | May | Lepidoptera | Pyralidae | Atheloca |
| 2505Wmcc6c | WMCC | May | Lepidoptera | Pyralidae | Elasmopalpus |
| 2505Wmcc6c | WMCC | May | Lepidoptera | Tischeriidae | Coptotriche |
| 2505Wmcc6c | WMCC | May | Lepidoptera | Tortricidae | Acleris |
| 2505Wmcc6c | WMCC | May | Lepidoptera | Tortricidae | Argyrotaenia |
| 2505Wmcc6c | WMCC | May | Lepidoptera | Tortricidae | Rhyacionia |
| 2505Wmcc6c | WMCC | May | Megaloptera | Corydalidae | Corydalus |
| 2505Wmcc6c | WMCC | May | Neuroptera | Hemerobiidae | Hemerobius |

| | | | | | |
|---|---|---|---|---|---|
| 2505Wmcc6c | WMCC | May | Odonata | Libellulidae | Brachymesia |
| 2505Wmcc6c | WMCC | May | Orthoptera | Tettigoniidae | Conocephalus |
| 2505Wmcc6c | WMCC | May | Other | Other | Other |
| 2505Wmcc6c | WMCC | May | Other | Other | Other |
| 2505Wmcc6c | WMCC | May | Plecoptera | Chloroperlidae | Sweltsa |
| 2505Wmcc6c | WMCC | May | Psocodea | Peripsocidae | Peripsocus |
| 2505Wmcc6c | WMCC | May | Sarcoptiformes | Oripodidae | Other |
| 2505Wmcc6c | WMCC | May | Trichoptera | Dipseudopsidae | Phylocentropus |
| 2505Wmcc6c | WMCC | May | Trichoptera | Hydroptilidae | Orthotrichia |
| 2505Wmcc6c | WMCC | May | Trichoptera | Leptoceridae | Oecetis |
| 2505Wmcc6c | WMCC | May | Trombidiformes | Arrenuridae | Arrenurus |
| 2505Wmcc6c | WMCC | May | Trombidiformes | Aturidae | Aturus |
| 2505Wmcc6c | WMCC | May | Trombidiformes | Axonopsidae | Brachypoda |
| 2505Wmcc6c | WMCC | May | Trombidiformes | Eupodidae | Other |
| 2505Wmcc6c | WMCC | May | Trombidiformes | Feltriidae | Feltria |
| 2505Wmcc6c | WMCC | May | Trombidiformes | Hydryphantidae | Hydryphantes |
| 2505Wmcc6c | WMCC | May | Trombidiformes | Hydryphantidae | Parathyas |
| 2505Wmcc6c | WMCC | May | Trombidiformes | Lebertiidae | Lebertia |
| 2505Wmcc6c | WMCC | May | Trombidiformes | Mideopsidae | Mideopsis |
| 2505Wmcc6c | WMCC | May | Trombidiformes | Pionidae | Piona |
| 2505Wmcc6c | WMCC | May | Trombidiformes | Pionidae | Tiphys |
| 2505Wmcc6c | WMCC | May | Trombidiformes | Sperchontidae | Sperchon |
| 2505Wmcc6c | WMCC | May | Trombidiformes | Tarsonemidae | Phytonemus |
| 2505Wmcc6c | WMCC | May | Trombidiformes | Torrenticolidae | Testudacarus |
| 2505Wmcc6c | WMCC | May | Trombidiformes | Torrenticolidae | Torrenticola |
| 2505Wmcc6c | WMCC | May | Trombidiformes | Unionicolidae | Koenikea |
| 2506Wmcc6b | WMCC | June | Araneae | Agelenidae | Coras |
| 2506Wmcc6b | WMCC | June | Araneae | Amaurobiidae | Callobius |
| 2506Wmcc6b | WMCC | June | Araneae | Anyphaenidae | Anyphaena |
| 2506Wmcc6b | WMCC | June | Araneae | Cicurinidae | Cicurina |
| 2506Wmcc6b | WMCC | June | Araneae | Dictynidae | Dictyna |

| | | | | | |
|---|---|---|---|---|---|
| 2506Wmcc6b | WMCC | June | Araneae | Gnaphosidae | Gnaphosa |
| 2506Wmcc6b | WMCC | June | Araneae | Linyphiidae | Grammonota |
| 2506Wmcc6b | WMCC | June | Araneae | Lycosidae | Rabidosa |
| 2506Wmcc6b | WMCC | June | Araneae | Lycosidae | Schizocosa |
| 2506Wmcc6b | WMCC | June | Araneae | Phrurolithidae | Scotinella |
| 2506Wmcc6b | WMCC | June | Araneae | Salticidae | Maevia |
| 2506Wmcc6b | WMCC | June | Araneae | Salticidae | Sibianor |
| 2506Wmcc6b | WMCC | June | Araneae | Theridiidae | Canalidion |
| 2506Wmcc6b | WMCC | June | Araneae | Theridiidae | Rugathodes |
| 2506Wmcc6b | WMCC | June | Araneae | Theridiidae | Spintharus |
| 2506Wmcc6b | WMCC | June | Araneae | Thomisidae | Mecaphesa |
| 2506Wmcc6b | WMCC | June | Blattodea | Ectobiidae | Parcoblatta |
| 2506Wmcc6b | WMCC | June | Coleoptera | Buprestidae | Pachyschelus |
| 2506Wmcc6b | WMCC | June | Coleoptera | Carabidae | Apenes |
| 2506Wmcc6b | WMCC | June | Coleoptera | Carabidae | Harpalus |
| 2506Wmcc6b | WMCC | June | Coleoptera | Carabidae | Other |
| 2506Wmcc6b | WMCC | June | Coleoptera | Carabidae | Polyderis |
| 2506Wmcc6b | WMCC | June | Coleoptera | Curculionidae | Conotrachelus |
| 2506Wmcc6b | WMCC | June | Coleoptera | Curculionidae | Cyrtepistomus |
| 2506Wmcc6b | WMCC | June | Coleoptera | Curculionidae | Listronotus |
| 2506Wmcc6b | WMCC | June | Coleoptera | Dytiscidae | Acilius |
| 2506Wmcc6b | WMCC | June | Coleoptera | Lampyridae | Photinus |
| 2506Wmcc6b | WMCC | June | Coleoptera | Scarabaeidae | Dialytes |
| 2506Wmcc6b | WMCC | June | Coleoptera | Scarabaeidae | Melanocanthon |
| 2506Wmcc6b | WMCC | June | Coleoptera | Scirtidae | Contacyphon |
| 2506Wmcc6b | WMCC | June | Diptera | Agromyzidae | Calycomyza |
| 2506Wmcc6b | WMCC | June | Diptera | Agromyzidae | Cerodontha |
| 2506Wmcc6b | WMCC | June | Diptera | Agromyzidae | Japanagromyza |
| 2506Wmcc6b | WMCC | June | Diptera | Agromyzidae | Liriomyza |
| 2506Wmcc6b | WMCC | June | Diptera | Agromyzidae | Nemorimyza |
| 2506Wmcc6b | WMCC | June | Diptera | Agromyzidae | Phytomyza |

| | | | | | |
|---|---|---|---|---|---|
| 2506Wmcc6b | WMCC | June | Diptera | Anisopodidae | Mycetobia |
| 2506Wmcc6b | WMCC | June | Diptera | Calliphoridae | Lucilia |
| 2506Wmcc6b | WMCC | June | Diptera | Cecidomyiidae | Other |
| 2506Wmcc6b | WMCC | June | Diptera | Chaoboridae | Chaoborus |
| 2506Wmcc6b | WMCC | June | Diptera | Chaoboridae | Other |
| 2506Wmcc6b | WMCC | June | Diptera | Chironomidae | Ablabesmyia |
| 2506Wmcc6b | WMCC | June | Diptera | Chironomidae | Chironomus |
| 2506Wmcc6b | WMCC | June | Diptera | Chironomidae | Dicrotendipes |
| 2506Wmcc6b | WMCC | June | Diptera | Chironomidae | Endochironomus |
| 2506Wmcc6b | WMCC | June | Diptera | Chironomidae | Glyptotendipes |
| 2506Wmcc6b | WMCC | June | Diptera | Chironomidae | Microtendipes |
| 2506Wmcc6b | WMCC | June | Diptera | Chironomidae | Orthocladius |
| 2506Wmcc6b | WMCC | June | Diptera | Chironomidae | Other |
| 2506Wmcc6b | WMCC | June | Diptera | Chironomidae | Other |
| 2506Wmcc6b | WMCC | June | Diptera | Chironomidae | Other |
| 2506Wmcc6b | WMCC | June | Diptera | Chironomidae | Other |
| 2506Wmcc6b | WMCC | June | Diptera | Chironomidae | Parachironomus |
| 2506Wmcc6b | WMCC | June | Diptera | Chironomidae | Polypedilum |
| 2506Wmcc6b | WMCC | June | Diptera | Chironomidae | Pseudochironomus |
| 2506Wmcc6b | WMCC | June | Diptera | Chironomidae | Tanytarsus |
| 2506Wmcc6b | WMCC | June | Diptera | Chironomidae | Xenochironomus |
| 2506Wmcc6b | WMCC | June | Diptera | Chloropidae | Apotropina |
| 2506Wmcc6b | WMCC | June | Diptera | Chloropidae | Elachiptera |
| 2506Wmcc6b | WMCC | June | Diptera | Chloropidae | Liohippelates |
| 2506Wmcc6b | WMCC | June | Diptera | Chloropidae | Oscinella |
| 2506Wmcc6b | WMCC | June | Diptera | Culicidae | Coquillettidia |
| 2506Wmcc6b | WMCC | June | Diptera | Culicidae | Culex |
| 2506Wmcc6b | WMCC | June | Diptera | Culicidae | Uranotaenia |
| 2506Wmcc6b | WMCC | June | Diptera | Drosophilidae | Drosophila |
| 2506Wmcc6b | WMCC | June | Diptera | Empididae | Other |
| 2506Wmcc6b | WMCC | June | Diptera | Hybotidae | Platypalpus |

| | | | | | |
|---|---|---|---|---|---|
| 2506Wmcc6b | WMCC | June | Diptera | Hybotidae | Syndyas |
| 2506Wmcc6b | WMCC | June | Diptera | Keroplatidae | Other |
| 2506Wmcc6b | WMCC | June | Diptera | Limoniidae | Dicranomyia |
| 2506Wmcc6b | WMCC | June | Diptera | Limoniidae | Dicranophragma |
| 2506Wmcc6b | WMCC | June | Diptera | Limoniidae | Epiphragma |
| 2506Wmcc6b | WMCC | June | Diptera | Limoniidae | Erioptera |
| 2506Wmcc6b | WMCC | June | Diptera | Limoniidae | Metalimnobia |
| 2506Wmcc6b | WMCC | June | Diptera | Limoniidae | Ormosia |
| 2506Wmcc6b | WMCC | June | Diptera | Limoniidae | Other |
| 2506Wmcc6b | WMCC | June | Diptera | Limoniidae | Other |
| 2506Wmcc6b | WMCC | June | Diptera | Limoniidae | Pseudolimnophila |
| 2506Wmcc6b | WMCC | June | Diptera | Muscidae | Mydaea |
| 2506Wmcc6b | WMCC | June | Diptera | Mycetophilidae | Mycomya |
| 2506Wmcc6b | WMCC | June | Diptera | Mycetophilidae | Neoempheria |
| 2506Wmcc6b | WMCC | June | Diptera | Opomyzidae | Geomyza |
| 2506Wmcc6b | WMCC | June | Diptera | Pallopteridae | Toxonevra |
| 2506Wmcc6b | WMCC | June | Diptera | Phoridae | Diplonevra |
| 2506Wmcc6b | WMCC | June | Diptera | Polleniidae | Pollenia |
| 2506Wmcc6b | WMCC | June | Diptera | Psychodidae | Clogmia |
| 2506Wmcc6b | WMCC | June | Diptera | Psychodidae | Psychoda |
| 2506Wmcc6b | WMCC | June | Diptera | Sarcophagidae | Ravinia |
| 2506Wmcc6b | WMCC | June | Diptera | Sciaridae | Claustropyga |
| 2506Wmcc6b | WMCC | June | Diptera | Sciaridae | Corynoptera |
| 2506Wmcc6b | WMCC | June | Diptera | Sciaridae | Epidapus |
| 2506Wmcc6b | WMCC | June | Diptera | Sciaridae | Other |
| 2506Wmcc6b | WMCC | June | Diptera | Sphaeroceridae | Leptocera |
| 2506Wmcc6b | WMCC | June | Diptera | Syrphidae | Chalcosyrphus |
| 2506Wmcc6b | WMCC | June | Diptera | Syrphidae | Palpada |
| 2506Wmcc6b | WMCC | June | Diptera | Tabanidae | Hybomitra |
| 2506Wmcc6b | WMCC | June | Diptera | Tabanidae | Other |
| 2506Wmcc6b | WMCC | June | Diptera | Tachinidae | Cryptomeigenia |

| | | | | | |
|---|---|---|---|---|---|
| 2506Wmcc6b | WMCC | June | Diptera | Tachinidae | Cyzenis |
| 2506Wmcc6b | WMCC | June | Diptera | Tachinidae | Zaira |
| 2506Wmcc6b | WMCC | June | Diptera | Tipulidae | Dolichopeza |
| 2506Wmcc6b | WMCC | June | Diptera | Tipulidae | Tipula |
| 2506Wmcc6b | WMCC | June | Ephemeroptera | Caenidae | Caenis |
| 2506Wmcc6b | WMCC | June | Ephemeroptera | Ephemeridae | Hexagenia |
| 2506Wmcc6b | WMCC | June | Ephemeroptera | Heptageniidae | Epeorus |
| 2506Wmcc6b | WMCC | June | Hemiptera | Clastopteridae | Clastoptera |
| 2506Wmcc6b | WMCC | June | Hemiptera | Lygaeidae | Kleidocerys |
| 2506Wmcc6b | WMCC | June | Hymenoptera | Apidae | Triepeolus |
| 2506Wmcc6b | WMCC | June | Hymenoptera | Braconidae | Bracon |
| 2506Wmcc6b | WMCC | June | Hymenoptera | Braconidae | Chorebus |
| 2506Wmcc6b | WMCC | June | Hymenoptera | Ichneumonidae | Megastylus |
| 2506Wmcc6b | WMCC | June | Hymenoptera | Ichneumonidae | Polyblastus |
| 2506Wmcc6b | WMCC | June | Hymenoptera | Other | Other |
| 2506Wmcc6b | WMCC | June | Hymenoptera | Tenthredinidae | Dolerus |
| 2506Wmcc6b | WMCC | June | Hymenoptera | Tenthredinidae | Nematus |
| 2506Wmcc6b | WMCC | June | Hymenoptera | Trigonalidae | Orthogonalys |
| 2506Wmcc6b | WMCC | June | Lepidoptera | Crambidae | Blepharomastix |
| 2506Wmcc6b | WMCC | June | Lepidoptera | Crambidae | Parapoynx |
| 2506Wmcc6b | WMCC | June | Lepidoptera | Elachistidae | Elachista |
| 2506Wmcc6b | WMCC | June | Lepidoptera | Erebidae | Panopoda |
| 2506Wmcc6b | WMCC | June | Lepidoptera | Gelechiidae | Chionodes |
| 2506Wmcc6b | WMCC | June | Lepidoptera | Noctuidae | Condica |
| 2506Wmcc6b | WMCC | June | Lepidoptera | Noctuidae | Leuconycta |
| 2506Wmcc6b | WMCC | June | Lepidoptera | Noctuidae | Pyreferra |
| 2506Wmcc6b | WMCC | June | Lepidoptera | Notodontidae | Heterocampa |
| 2506Wmcc6b | WMCC | June | Lepidoptera | Notodontidae | Hyparpax |
| 2506Wmcc6b | WMCC | June | Lepidoptera | Nymphalidae | Oeneis |
| 2506Wmcc6b | WMCC | June | Lepidoptera | Pyralidae | Atheloca |
| 2506Wmcc6b | WMCC | June | Lepidoptera | Pyralidae | Elasmopalpus |

| | | | | | |
|---|---|---|---|---|---|
| 2506Wmcc6b | WMCC | June | Lepidoptera | Tischeriidae | Coptotriche |
| 2506Wmcc6b | WMCC | June | Lepidoptera | Tortricidae | Acleris |
| 2506Wmcc6b | WMCC | June | Lepidoptera | Tortricidae | Argyrotaenia |
| 2506Wmcc6b | WMCC | June | Lepidoptera | Tortricidae | Rhyacionia |
| 2506Wmcc6b | WMCC | June | Megaloptera | Corydalidae | Corydalus |
| 2506Wmcc6b | WMCC | June | Neuroptera | Hemerobiidae | Hemerobius |
| 2506Wmcc6b | WMCC | June | Odonata | Libellulidae | Brachymesia |
| 2506Wmcc6b | WMCC | June | Orthoptera | Tettigoniidae | Conocephalus |
| 2506Wmcc6b | WMCC | June | Other | Other | Other |
| 2506Wmcc6b | WMCC | June | Other | Other | Other |
| 2506Wmcc6b | WMCC | June | Plecoptera | Chloroperlidae | Sweltsa |
| 2506Wmcc6b | WMCC | June | Psocodea | Peripsocidae | Peripsocus |
| 2506Wmcc6b | WMCC | June | Sarcoptiformes | Oripodidae | Other |
| 2506Wmcc6b | WMCC | June | Trichoptera | Dipseudopsidae | Phylocentropus |
| 2506Wmcc6b | WMCC | June | Trichoptera | Hydroptilidae | Orthotrichia |
| 2506Wmcc6b | WMCC | June | Trichoptera | Leptoceridae | Oecetis |
| 2506Wmcc6b | WMCC | June | Trombidiformes | Arrenuridae | Arrenurus |
| 2506Wmcc6b | WMCC | June | Trombidiformes | Aturidae | Aturus |
| 2506Wmcc6b | WMCC | June | Trombidiformes | Axonopsidae | Brachypoda |
| 2506Wmcc6b | WMCC | June | Trombidiformes | Eupodidae | Other |
| 2506Wmcc6b | WMCC | June | Trombidiformes | Feltriidae | Feltria |
| 2506Wmcc6b | WMCC | June | Trombidiformes | Hydryphantidae | Hydryphantes |
| 2506Wmcc6b | WMCC | June | Trombidiformes | Hydryphantidae | Parathyas |
| 2506Wmcc6b | WMCC | June | Trombidiformes | Lebertiidae | Lebertia |
| 2506Wmcc6b | WMCC | June | Trombidiformes | Mideopsidae | Mideopsis |
| 2506Wmcc6b | WMCC | June | Trombidiformes | Pionidae | Piona |
| 2506Wmcc6b | WMCC | June | Trombidiformes | Pionidae | Tiphys |
| 2506Wmcc6b | WMCC | June | Trombidiformes | Sperchontidae | Sperchon |
| 2506Wmcc6b | WMCC | June | Trombidiformes | Tarsonemidae | Phytonemus |
| 2506Wmcc6b | WMCC | June | Trombidiformes | Torrenticolidae | Testudacarus |
| 2506Wmcc6b | WMCC | June | Trombidiformes | Torrenticolidae | Torrenticola |

| | | | | | |
|---|---|---|---|---|---|
| 2506Wmcc6b | WMCC | June | Trombidiformes | Unionicolidae | Koenikea |
| 2506Wmcc6c | WMCC | June | Araneae | Agelenidae | Coras |
| 2506Wmcc6c | WMCC | June | Araneae | Amaurobiidae | Callobius |
| 2506Wmcc6c | WMCC | June | Araneae | Anyphaenidae | Anyphaena |
| 2506Wmcc6c | WMCC | June | Araneae | Cicurinidae | Cicurina |
| 2506Wmcc6c | WMCC | June | Araneae | Dictynidae | Dictyna |
| 2506Wmcc6c | WMCC | June | Araneae | Gnaphosidae | Gnaphosa |
| 2506Wmcc6c | WMCC | June | Araneae | Linyphiidae | Grammonota |
| 2506Wmcc6c | WMCC | June | Araneae | Lycosidae | Rabidosa |
| 2506Wmcc6c | WMCC | June | Araneae | Lycosidae | Schizocosa |
| 2506Wmcc6c | WMCC | June | Araneae | Phrurolithidae | Scotinella |
| 2506Wmcc6c | WMCC | June | Araneae | Salticidae | Maevia |
| 2506Wmcc6c | WMCC | June | Araneae | Salticidae | Sibianor |
| 2506Wmcc6c | WMCC | June | Araneae | Theridiidae | Canalidion |
| 2506Wmcc6c | WMCC | June | Araneae | Theridiidae | Rugathodes |
| 2506Wmcc6c | WMCC | June | Araneae | Theridiidae | Spintharus |
| 2506Wmcc6c | WMCC | June | Araneae | Thomisidae | Mecaphesa |
| 2506Wmcc6c | WMCC | June | Blattodea | Ectobiidae | Parcoblatta |
| 2506Wmcc6c | WMCC | June | Coleoptera | Buprestidae | Pachyschelus |
| 2506Wmcc6c | WMCC | June | Coleoptera | Carabidae | Apenes |
| 2506Wmcc6c | WMCC | June | Coleoptera | Carabidae | Harpalus |
| 2506Wmcc6c | WMCC | June | Coleoptera | Carabidae | Other |
| 2506Wmcc6c | WMCC | June | Coleoptera | Carabidae | Polyderis |
| 2506Wmcc6c | WMCC | June | Coleoptera | Curculionidae | Conotrachelus |
| 2506Wmcc6c | WMCC | June | Coleoptera | Curculionidae | Cyrtepistomus |
| 2506Wmcc6c | WMCC | June | Coleoptera | Curculionidae | Listronotus |
| 2506Wmcc6c | WMCC | June | Coleoptera | Dytiscidae | Acilius |
| 2506Wmcc6c | WMCC | June | Coleoptera | Lampyridae | Photinus |
| 2506Wmcc6c | WMCC | June | Coleoptera | Scarabaeidae | Dialytes |
| 2506Wmcc6c | WMCC | June | Coleoptera | Scarabaeidae | Melanocanthon |
| 2506Wmcc6c | WMCC | June | Coleoptera | Scirtidae | Contacyphon |

| | | | | | |
|---|---|---|---|---|---|
| 2506Wmcc6c | WMCC | June | Diptera | Agromyzidae | Calycomyza |
| 2506Wmcc6c | WMCC | June | Diptera | Agromyzidae | Cerodontha |
| 2506Wmcc6c | WMCC | June | Diptera | Agromyzidae | Japanagromyza |
| 2506Wmcc6c | WMCC | June | Diptera | Agromyzidae | Liriomyza |
| 2506Wmcc6c | WMCC | June | Diptera | Agromyzidae | Nemorimyza |
| 2506Wmcc6c | WMCC | June | Diptera | Agromyzidae | Phytomyza |
| 2506Wmcc6c | WMCC | June | Diptera | Anisopodidae | Mycetobia |
| 2506Wmcc6c | WMCC | June | Diptera | Calliphoridae | Lucilia |
| 2506Wmcc6c | WMCC | June | Diptera | Cecidomyiidae | Other |
| 2506Wmcc6c | WMCC | June | Diptera | Chaoboridae | Chaoborus |
| 2506Wmcc6c | WMCC | June | Diptera | Chaoboridae | Other |
| 2506Wmcc6c | WMCC | June | Diptera | Chironomidae | Ablabesmyia |
| 2506Wmcc6c | WMCC | June | Diptera | Chironomidae | Chironomus |
| 2506Wmcc6c | WMCC | June | Diptera | Chironomidae | Dicrotendipes |
| 2506Wmcc6c | WMCC | June | Diptera | Chironomidae | Endochironomus |
| 2506Wmcc6c | WMCC | June | Diptera | Chironomidae | Glyptotendipes |
| 2506Wmcc6c | WMCC | June | Diptera | Chironomidae | Microtendipes |
| 2506Wmcc6c | WMCC | June | Diptera | Chironomidae | Orthocladius |
| 2506Wmcc6c | WMCC | June | Diptera | Chironomidae | Other |
| 2506Wmcc6c | WMCC | June | Diptera | Chironomidae | Other |
| 2506Wmcc6c | WMCC | June | Diptera | Chironomidae | Other |
| 2506Wmcc6c | WMCC | June | Diptera | Chironomidae | Other |
| 2506Wmcc6c | WMCC | June | Diptera | Chironomidae | Parachironomus |
| 2506Wmcc6c | WMCC | June | Diptera | Chironomidae | Polypedilum |
| 2506Wmcc6c | WMCC | June | Diptera | Chironomidae | Pseudochironomus |
| 2506Wmcc6c | WMCC | June | Diptera | Chironomidae | Tanytarsus |
| 2506Wmcc6c | WMCC | June | Diptera | Chironomidae | Xenochironomus |
| 2506Wmcc6c | WMCC | June | Diptera | Chloropidae | Apotropina |
| 2506Wmcc6c | WMCC | June | Diptera | Chloropidae | Elachiptera |
| 2506Wmcc6c | WMCC | June | Diptera | Chloropidae | Liohippelates |
| 2506Wmcc6c | WMCC | June | Diptera | Chloropidae | Oscinella |

| | | | | | |
|---|---|---|---|---|---|
| 2506Wmcc6c | WMCC | June | Diptera | Culicidae | Coquillettidia |
| 2506Wmcc6c | WMCC | June | Diptera | Culicidae | Culex |
| 2506Wmcc6c | WMCC | June | Diptera | Culicidae | Uranotaenia |
| 2506Wmcc6c | WMCC | June | Diptera | Drosophilidae | Drosophila |
| 2506Wmcc6c | WMCC | June | Diptera | Empididae | Other |
| 2506Wmcc6c | WMCC | June | Diptera | Hybotidae | Platypalpus |
| 2506Wmcc6c | WMCC | June | Diptera | Hybotidae | Syndyas |
| 2506Wmcc6c | WMCC | June | Diptera | Keroplatidae | Other |
| 2506Wmcc6c | WMCC | June | Diptera | Limoniidae | Dicranomyia |
| 2506Wmcc6c | WMCC | June | Diptera | Limoniidae | Dicranophragma |
| 2506Wmcc6c | WMCC | June | Diptera | Limoniidae | Epiphragma |
| 2506Wmcc6c | WMCC | June | Diptera | Limoniidae | Erioptera |
| 2506Wmcc6c | WMCC | June | Diptera | Limoniidae | Metalimnobia |
| 2506Wmcc6c | WMCC | June | Diptera | Limoniidae | Ormosia |
| 2506Wmcc6c | WMCC | June | Diptera | Limoniidae | Other |
| 2506Wmcc6c | WMCC | June | Diptera | Limoniidae | Other |
| 2506Wmcc6c | WMCC | June | Diptera | Limoniidae | Pseudolimnophila |
| 2506Wmcc6c | WMCC | June | Diptera | Muscidae | Mydaea |
| 2506Wmcc6c | WMCC | June | Diptera | Mycetophilidae | Mycomya |
| 2506Wmcc6c | WMCC | June | Diptera | Mycetophilidae | Neoempheria |
| 2506Wmcc6c | WMCC | June | Diptera | Opomyzidae | Geomyza |
| 2506Wmcc6c | WMCC | June | Diptera | Pallopteridae | Toxonevra |
| 2506Wmcc6c | WMCC | June | Diptera | Phoridae | Diplonevra |
| 2506Wmcc6c | WMCC | June | Diptera | Polleniidae | Pollenia |
| 2506Wmcc6c | WMCC | June | Diptera | Psychodidae | Clogmia |
| 2506Wmcc6c | WMCC | June | Diptera | Psychodidae | Psychoda |
| 2506Wmcc6c | WMCC | June | Diptera | Sarcophagidae | Ravinia |
| 2506Wmcc6c | WMCC | June | Diptera | Sciaridae | Claustropyga |
| 2506Wmcc6c | WMCC | June | Diptera | Sciaridae | Corynoptera |
| 2506Wmcc6c | WMCC | June | Diptera | Sciaridae | Epidapus |
| 2506Wmcc6c | WMCC | June | Diptera | Sciaridae | Other |

| | | | | | |
|---|---|---|---|---|---|
| 2506Wmcc6c | WMCC | June | Diptera | Sphaeroceridae | Leptocera |
| 2506Wmcc6c | WMCC | June | Diptera | Syrphidae | Chalcosyrphus |
| 2506Wmcc6c | WMCC | June | Diptera | Syrphidae | Palpada |
| 2506Wmcc6c | WMCC | June | Diptera | Tabanidae | Hybomitra |
| 2506Wmcc6c | WMCC | June | Diptera | Tabanidae | Other |
| 2506Wmcc6c | WMCC | June | Diptera | Tachinidae | Cryptomeigenia |
| 2506Wmcc6c | WMCC | June | Diptera | Tachinidae | Cyzenis |
| 2506Wmcc6c | WMCC | June | Diptera | Tachinidae | Zaira |
| 2506Wmcc6c | WMCC | June | Diptera | Tipulidae | Dolichopeza |
| 2506Wmcc6c | WMCC | June | Diptera | Tipulidae | Tipula |
| 2506Wmcc6c | WMCC | June | Ephemeroptera | Caenidae | Caenis |
| 2506Wmcc6c | WMCC | June | Ephemeroptera | Ephemeridae | Hexagenia |
| 2506Wmcc6c | WMCC | June | Ephemeroptera | Heptageniidae | Epeorus |
| 2506Wmcc6c | WMCC | June | Hemiptera | Clastopteridae | Clastoptera |
| 2506Wmcc6c | WMCC | June | Hemiptera | Lygaeidae | Kleidocerys |
| 2506Wmcc6c | WMCC | June | Hymenoptera | Apidae | Triepeolus |
| 2506Wmcc6c | WMCC | June | Hymenoptera | Braconidae | Bracon |
| 2506Wmcc6c | WMCC | June | Hymenoptera | Braconidae | Chorebus |
| 2506Wmcc6c | WMCC | June | Hymenoptera | Ichneumonidae | Megastylus |
| 2506Wmcc6c | WMCC | June | Hymenoptera | Ichneumonidae | Polyblastus |
| 2506Wmcc6c | WMCC | June | Hymenoptera | Other | Other |
| 2506Wmcc6c | WMCC | June | Hymenoptera | Tenthredinidae | Dolerus |
| 2506Wmcc6c | WMCC | June | Hymenoptera | Tenthredinidae | Nematus |
| 2506Wmcc6c | WMCC | June | Hymenoptera | Trigonalidae | Orthogonalys |
| 2506Wmcc6c | WMCC | June | Lepidoptera | Crambidae | Blepharomastix |
| 2506Wmcc6c | WMCC | June | Lepidoptera | Crambidae | Parapoynx |
| 2506Wmcc6c | WMCC | June | Lepidoptera | Elachistidae | Elachista |
| 2506Wmcc6c | WMCC | June | Lepidoptera | Erebidae | Panopoda |
| 2506Wmcc6c | WMCC | June | Lepidoptera | Gelechiidae | Chionodes |
| 2506Wmcc6c | WMCC | June | Lepidoptera | Noctuidae | Condica |
| 2506Wmcc6c | WMCC | June | Lepidoptera | Noctuidae | Leuconycta |

| | | | | | |
|---|---|---|---|---|---|
| 2506Wmcc6c | WMCC | June | Lepidoptera | Noctuidae | Pyreferra |
| 2506Wmcc6c | WMCC | June | Lepidoptera | Notodontidae | Heterocampa |
| 2506Wmcc6c | WMCC | June | Lepidoptera | Notodontidae | Hyparpax |
| 2506Wmcc6c | WMCC | June | Lepidoptera | Nymphalidae | Oeneis |
| 2506Wmcc6c | WMCC | June | Lepidoptera | Pyralidae | Atheloca |
| 2506Wmcc6c | WMCC | June | Lepidoptera | Pyralidae | Elasmopalpus |
| 2506Wmcc6c | WMCC | June | Lepidoptera | Tischeriidae | Coptotriche |
| 2506Wmcc6c | WMCC | June | Lepidoptera | Tortricidae | Acleris |
| 2506Wmcc6c | WMCC | June | Lepidoptera | Tortricidae | Argyrotaenia |
| 2506Wmcc6c | WMCC | June | Lepidoptera | Tortricidae | Rhyacionia |
| 2506Wmcc6c | WMCC | June | Megaloptera | Corydalidae | Corydalus |
| 2506Wmcc6c | WMCC | June | Neuroptera | Hemerobiidae | Hemerobius |
| 2506Wmcc6c | WMCC | June | Odonata | Libellulidae | Brachymesia |
| 2506Wmcc6c | WMCC | June | Orthoptera | Tettigoniidae | Conocephalus |
| 2506Wmcc6c | WMCC | June | Other | Other | Other |
| 2506Wmcc6c | WMCC | June | Other | Other | Other |
| 2506Wmcc6c | WMCC | June | Plecoptera | Chloroperlidae | Sweltsa |
| 2506Wmcc6c | WMCC | June | Psocodea | Peripsocidae | Peripsocus |
| 2506Wmcc6c | WMCC | June | Sarcoptiformes | Oripodidae | Other |
| 2506Wmcc6c | WMCC | June | Trichoptera | Dipseudopsidae | Phylocentropus |
| 2506Wmcc6c | WMCC | June | Trichoptera | Hydroptilidae | Orthotrichia |
| 2506Wmcc6c | WMCC | June | Trichoptera | Leptoceridae | Oecetis |
| 2506Wmcc6c | WMCC | June | Trombidiformes | Arrenuridae | Arrenurus |
| 2506Wmcc6c | WMCC | June | Trombidiformes | Aturidae | Aturus |
| 2506Wmcc6c | WMCC | June | Trombidiformes | Axonopsidae | Brachypoda |
| 2506Wmcc6c | WMCC | June | Trombidiformes | Eupodidae | Other |
| 2506Wmcc6c | WMCC | June | Trombidiformes | Feltriidae | Feltria |
| 2506Wmcc6c | WMCC | June | Trombidiformes | Hydryphantidae | Hydryphantes |
| 2506Wmcc6c | WMCC | June | Trombidiformes | Hydryphantidae | Parathyas |
| 2506Wmcc6c | WMCC | June | Trombidiformes | Lebertiidae | Lebertia |
| 2506Wmcc6c | WMCC | June | Trombidiformes | Mideopsidae | Mideopsis |

| | | | | | |
|---|---|---|---|---|---|
| 2506Wmcc6c | WMCC | June | Trombidiformes | Pionidae | Piona |
| 2506Wmcc6c | WMCC | June | Trombidiformes | Pionidae | Tiphys |
| 2506Wmcc6c | WMCC | June | Trombidiformes | Sperchontidae | Sperchon |
| 2506Wmcc6c | WMCC | June | Trombidiformes | Tarsonemidae | Phytonemus |
| 2506Wmcc6c | WMCC | June | Trombidiformes | Torrenticolidae | Testudacarus |
| 2506Wmcc6c | WMCC | June | Trombidiformes | Torrenticolidae | Torrenticola |
| 2506Wmcc6c | WMCC | June | Trombidiformes | Unionicolidae | Koenikea |
| 2507Wmcc6b | WMCC | July | Araneae | Agelenidae | Coras |
| 2507Wmcc6b | WMCC | July | Araneae | Amaurobiidae | Callobius |
| 2507Wmcc6b | WMCC | July | Araneae | Anyphaenidae | Anyphaena |
| 2507Wmcc6b | WMCC | July | Araneae | Cicurinidae | Cicurina |
| 2507Wmcc6b | WMCC | July | Araneae | Dictynidae | Dictyna |
| 2507Wmcc6b | WMCC | July | Araneae | Gnaphosidae | Gnaphosa |
| 2507Wmcc6b | WMCC | July | Araneae | Linyphiidae | Grammonota |
| 2507Wmcc6b | WMCC | July | Araneae | Lycosidae | Rabidosa |
| 2507Wmcc6b | WMCC | July | Araneae | Lycosidae | Schizocosa |
| 2507Wmcc6b | WMCC | July | Araneae | Phrurolithidae | Scotinella |
| 2507Wmcc6b | WMCC | July | Araneae | Salticidae | Maevia |
| 2507Wmcc6b | WMCC | July | Araneae | Salticidae | Sibianor |
| 2507Wmcc6b | WMCC | July | Araneae | Theridiidae | Canalidion |
| 2507Wmcc6b | WMCC | July | Araneae | Theridiidae | Rugathodes |
| 2507Wmcc6b | WMCC | July | Araneae | Theridiidae | Spintharus |
| 2507Wmcc6b | WMCC | July | Araneae | Thomisidae | Mecaphesa |
| 2507Wmcc6b | WMCC | July | Blattodea | Ectobiidae | Parcoblatta |
| 2507Wmcc6b | WMCC | July | Coleoptera | Buprestidae | Pachyschelus |
| 2507Wmcc6b | WMCC | July | Coleoptera | Carabidae | Apenes |
| 2507Wmcc6b | WMCC | July | Coleoptera | Carabidae | Harpalus |
| 2507Wmcc6b | WMCC | July | Coleoptera | Carabidae | Other |
| 2507Wmcc6b | WMCC | July | Coleoptera | Carabidae | Polyderis |
| 2507Wmcc6b | WMCC | July | Coleoptera | Curculionidae | Conotrachelus |
| 2507Wmcc6b | WMCC | July | Coleoptera | Curculionidae | Cyrtepistomus |

| | | | | | |
|---|---|---|---|---|---|
| 2507Wmcc6b | WMCC | July | Coleoptera | Curculionidae | Listronotus |
| 2507Wmcc6b | WMCC | July | Coleoptera | Dytiscidae | Acilius |
| 2507Wmcc6b | WMCC | July | Coleoptera | Lampyridae | Photinus |
| 2507Wmcc6b | WMCC | July | Coleoptera | Scarabaeidae | Dialytes |
| 2507Wmcc6b | WMCC | July | Coleoptera | Scarabaeidae | Melanocanthon |
| 2507Wmcc6b | WMCC | July | Coleoptera | Scirtidae | Contacyphon |
| 2507Wmcc6b | WMCC | July | Diptera | Agromyzidae | Calycomyza |
| 2507Wmcc6b | WMCC | July | Diptera | Agromyzidae | Cerodontha |
| 2507Wmcc6b | WMCC | July | Diptera | Agromyzidae | Japanagromyza |
| 2507Wmcc6b | WMCC | July | Diptera | Agromyzidae | Liriomyza |
| 2507Wmcc6b | WMCC | July | Diptera | Agromyzidae | Nemorimyza |
| 2507Wmcc6b | WMCC | July | Diptera | Agromyzidae | Phytomyza |
| 2507Wmcc6b | WMCC | July | Diptera | Anisopodidae | Mycetobia |
| 2507Wmcc6b | WMCC | July | Diptera | Calliphoridae | Lucilia |
| 2507Wmcc6b | WMCC | July | Diptera | Cecidomyiidae | Other |
| 2507Wmcc6b | WMCC | July | Diptera | Chaoboridae | Chaoborus |
| 2507Wmcc6b | WMCC | July | Diptera | Chaoboridae | Other |
| 2507Wmcc6b | WMCC | July | Diptera | Chironomidae | Ablabesmyia |
| 2507Wmcc6b | WMCC | July | Diptera | Chironomidae | Chironomus |
| 2507Wmcc6b | WMCC | July | Diptera | Chironomidae | Dicrotendipes |
| 2507Wmcc6b | WMCC | July | Diptera | Chironomidae | Endochironomus |
| 2507Wmcc6b | WMCC | July | Diptera | Chironomidae | Glyptotendipes |
| 2507Wmcc6b | WMCC | July | Diptera | Chironomidae | Microtendipes |
| 2507Wmcc6b | WMCC | July | Diptera | Chironomidae | Orthocladius |
| 2507Wmcc6b | WMCC | July | Diptera | Chironomidae | Other |
| 2507Wmcc6b | WMCC | July | Diptera | Chironomidae | Other |
| 2507Wmcc6b | WMCC | July | Diptera | Chironomidae | Other |
| 2507Wmcc6b | WMCC | July | Diptera | Chironomidae | Other |
| 2507Wmcc6b | WMCC | July | Diptera | Chironomidae | Parachironomus |
| 2507Wmcc6b | WMCC | July | Diptera | Chironomidae | Polypedilum |
| 2507Wmcc6b | WMCC | July | Diptera | Chironomidae | Pseudochironomus |

| | | | | | |
|---|---|---|---|---|---|
| 2507Wmcc6b | WMCC | July | Diptera | Chironomidae | Tanytarsus |
| 2507Wmcc6b | WMCC | July | Diptera | Chironomidae | Xenochironomus |
| 2507Wmcc6b | WMCC | July | Diptera | Chloropidae | Apotropina |
| 2507Wmcc6b | WMCC | July | Diptera | Chloropidae | Elachiptera |
| 2507Wmcc6b | WMCC | July | Diptera | Chloropidae | Liohippelates |
| 2507Wmcc6b | WMCC | July | Diptera | Chloropidae | Oscinella |
| 2507Wmcc6b | WMCC | July | Diptera | Culicidae | Coquillettidia |
| 2507Wmcc6b | WMCC | July | Diptera | Culicidae | Culex |
| 2507Wmcc6b | WMCC | July | Diptera | Culicidae | Uranotaenia |
| 2507Wmcc6b | WMCC | July | Diptera | Drosophilidae | Drosophila |
| 2507Wmcc6b | WMCC | July | Diptera | Empididae | Other |
| 2507Wmcc6b | WMCC | July | Diptera | Hybotidae | Platypalpus |
| 2507Wmcc6b | WMCC | July | Diptera | Hybotidae | Syndyas |
| 2507Wmcc6b | WMCC | July | Diptera | Keroplatidae | Other |
| 2507Wmcc6b | WMCC | July | Diptera | Limoniidae | Dicranomyia |
| 2507Wmcc6b | WMCC | July | Diptera | Limoniidae | Dicranophragma |
| 2507Wmcc6b | WMCC | July | Diptera | Limoniidae | Epiphragma |
| 2507Wmcc6b | WMCC | July | Diptera | Limoniidae | Erioptera |
| 2507Wmcc6b | WMCC | July | Diptera | Limoniidae | Metalimnobia |
| 2507Wmcc6b | WMCC | July | Diptera | Limoniidae | Ormosia |
| 2507Wmcc6b | WMCC | July | Diptera | Limoniidae | Other |
| 2507Wmcc6b | WMCC | July | Diptera | Limoniidae | Other |
| 2507Wmcc6b | WMCC | July | Diptera | Limoniidae | Pseudolimnophila |
| 2507Wmcc6b | WMCC | July | Diptera | Muscidae | Mydaea |
| 2507Wmcc6b | WMCC | July | Diptera | Mycetophilidae | Mycomya |
| 2507Wmcc6b | WMCC | July | Diptera | Mycetophilidae | Neoempheria |
| 2507Wmcc6b | WMCC | July | Diptera | Opomyzidae | Geomyza |
| 2507Wmcc6b | WMCC | July | Diptera | Pallopteridae | Toxonevra |
| 2507Wmcc6b | WMCC | July | Diptera | Phoridae | Diplonevra |
| 2507Wmcc6b | WMCC | July | Diptera | Polleniidae | Pollenia |
| 2507Wmcc6b | WMCC | July | Diptera | Psychodidae | Clogmia |

| | | | | | |
|---|---|---|---|---|---|
| 2507Wmcc6b | WMCC | July | Diptera | Psychodidae | Psychoda |
| 2507Wmcc6b | WMCC | July | Diptera | Sarcophagidae | Ravinia |
| 2507Wmcc6b | WMCC | July | Diptera | Sciaridae | Claustropyga |
| 2507Wmcc6b | WMCC | July | Diptera | Sciaridae | Corynoptera |
| 2507Wmcc6b | WMCC | July | Diptera | Sciaridae | Epidapus |
| 2507Wmcc6b | WMCC | July | Diptera | Sciaridae | Other |
| 2507Wmcc6b | WMCC | July | Diptera | Sphaeroceridae | Leptocera |
| 2507Wmcc6b | WMCC | July | Diptera | Syrphidae | Chalcosyrphus |
| 2507Wmcc6b | WMCC | July | Diptera | Syrphidae | Palpada |
| 2507Wmcc6b | WMCC | July | Diptera | Tabanidae | Hybomitra |
| 2507Wmcc6b | WMCC | July | Diptera | Tabanidae | Other |
| 2507Wmcc6b | WMCC | July | Diptera | Tachinidae | Cryptomeigenia |
| 2507Wmcc6b | WMCC | July | Diptera | Tachinidae | Cyzenis |
| 2507Wmcc6b | WMCC | July | Diptera | Tachinidae | Zaira |
| 2507Wmcc6b | WMCC | July | Diptera | Tipulidae | Dolichopeza |
| 2507Wmcc6b | WMCC | July | Diptera | Tipulidae | Tipula |
| 2507Wmcc6b | WMCC | July | Ephemeroptera | Caenidae | Caenis |
| 2507Wmcc6b | WMCC | July | Ephemeroptera | Ephemeridae | Hexagenia |
| 2507Wmcc6b | WMCC | July | Ephemeroptera | Heptageniidae | Epeorus |
| 2507Wmcc6b | WMCC | July | Hemiptera | Clastopteridae | Clastoptera |
| 2507Wmcc6b | WMCC | July | Hemiptera | Lygaeidae | Kleidocerys |
| 2507Wmcc6b | WMCC | July | Hymenoptera | Apidae | Triepeolus |
| 2507Wmcc6b | WMCC | July | Hymenoptera | Braconidae | Bracon |
| 2507Wmcc6b | WMCC | July | Hymenoptera | Braconidae | Chorebus |
| 2507Wmcc6b | WMCC | July | Hymenoptera | Ichneumonidae | Megastylus |
| 2507Wmcc6b | WMCC | July | Hymenoptera | Ichneumonidae | Polyblastus |
| 2507Wmcc6b | WMCC | July | Hymenoptera | Other | Other |
| 2507Wmcc6b | WMCC | July | Hymenoptera | Tenthredinidae | Dolerus |
| 2507Wmcc6b | WMCC | July | Hymenoptera | Tenthredinidae | Nematus |
| 2507Wmcc6b | WMCC | July | Hymenoptera | Trigonalidae | Orthogonalys |
| 2507Wmcc6b | WMCC | July | Lepidoptera | Crambidae | Blepharomastix |

| | | | | | |
|---|---|---|---|---|---|
| 2507Wmcc6b | WMCC | July | Lepidoptera | Crambidae | Parapoynx |
| 2507Wmcc6b | WMCC | July | Lepidoptera | Elachistidae | Elachista |
| 2507Wmcc6b | WMCC | July | Lepidoptera | Erebidae | Panopoda |
| 2507Wmcc6b | WMCC | July | Lepidoptera | Gelechiidae | Chionodes |
| 2507Wmcc6b | WMCC | July | Lepidoptera | Noctuidae | Condica |
| 2507Wmcc6b | WMCC | July | Lepidoptera | Noctuidae | Leuconycta |
| 2507Wmcc6b | WMCC | July | Lepidoptera | Noctuidae | Pyreferra |
| 2507Wmcc6b | WMCC | July | Lepidoptera | Notodontidae | Heterocampa |
| 2507Wmcc6b | WMCC | July | Lepidoptera | Notodontidae | Hyparpax |
| 2507Wmcc6b | WMCC | July | Lepidoptera | Nymphalidae | Oeneis |
| 2507Wmcc6b | WMCC | July | Lepidoptera | Pyralidae | Atheloca |
| 2507Wmcc6b | WMCC | July | Lepidoptera | Pyralidae | Elasmopalpus |
| 2507Wmcc6b | WMCC | July | Lepidoptera | Tischeriidae | Coptotriche |
| 2507Wmcc6b | WMCC | July | Lepidoptera | Tortricidae | Acleris |
| 2507Wmcc6b | WMCC | July | Lepidoptera | Tortricidae | Argyrotaenia |
| 2507Wmcc6b | WMCC | July | Lepidoptera | Tortricidae | Rhyacionia |
| 2507Wmcc6b | WMCC | July | Megaloptera | Corydalidae | Corydalus |
| 2507Wmcc6b | WMCC | July | Neuroptera | Hemerobiidae | Hemerobius |
| 2507Wmcc6b | WMCC | July | Odonata | Libellulidae | Brachymesia |
| 2507Wmcc6b | WMCC | July | Orthoptera | Tettigoniidae | Conocephalus |
| 2507Wmcc6b | WMCC | July | Other | Other | Other |
| 2507Wmcc6b | WMCC | July | Other | Other | Other |
| 2507Wmcc6b | WMCC | July | Plecoptera | Chloroperlidae | Sweltsa |
| 2507Wmcc6b | WMCC | July | Psocodea | Peripsocidae | Peripsocus |
| 2507Wmcc6b | WMCC | July | Sarcoptiformes | Oripodidae | Other |
| 2507Wmcc6b | WMCC | July | Trichoptera | Dipseudopsidae | Phylocentropus |
| 2507Wmcc6b | WMCC | July | Trichoptera | Hydroptilidae | Orthotrichia |
| 2507Wmcc6b | WMCC | July | Trichoptera | Leptoceridae | Oecetis |
| 2507Wmcc6b | WMCC | July | Trombidiformes | Arrenuridae | Arrenurus |
| 2507Wmcc6b | WMCC | July | Trombidiformes | Aturidae | Aturus |
| 2507Wmcc6b | WMCC | July | Trombidiformes | Axonopsidae | Brachypoda |

| | | | | | |
|---|---|---|---|---|---|
| 2507Wmcc6b | WMCC | July | Trombidiformes | Eupodidae | Other |
| 2507Wmcc6b | WMCC | July | Trombidiformes | Feltriidae | Feltria |
| 2507Wmcc6b | WMCC | July | Trombidiformes | Hydryphantidae | Hydryphantes |
| 2507Wmcc6b | WMCC | July | Trombidiformes | Hydryphantidae | Parathyas |
| 2507Wmcc6b | WMCC | July | Trombidiformes | Lebertiidae | Lebertia |
| 2507Wmcc6b | WMCC | July | Trombidiformes | Mideopsidae | Mideopsis |
| 2507Wmcc6b | WMCC | July | Trombidiformes | Pionidae | Piona |
| 2507Wmcc6b | WMCC | July | Trombidiformes | Pionidae | Tiphys |
| 2507Wmcc6b | WMCC | July | Trombidiformes | Sperchontidae | Sperchon |
| 2507Wmcc6b | WMCC | July | Trombidiformes | Tarsonemidae | Phytonemus |
| 2507Wmcc6b | WMCC | July | Trombidiformes | Torrenticolidae | Testudacarus |
| 2507Wmcc6b | WMCC | July | Trombidiformes | Torrenticolidae | Torrenticola |
| 2507Wmcc6b | WMCC | July | Trombidiformes | Unionicolidae | Koenikea |
| 2507Wmcc6c | WMCC | July | Araneae | Agelenidae | Coras |
| 2507Wmcc6c | WMCC | July | Araneae | Amaurobiidae | Callobius |
| 2507Wmcc6c | WMCC | July | Araneae | Anyphaenidae | Anyphaena |
| 2507Wmcc6c | WMCC | July | Araneae | Cicurinidae | Cicurina |
| 2507Wmcc6c | WMCC | July | Araneae | Dictynidae | Dictyna |
| 2507Wmcc6c | WMCC | July | Araneae | Gnaphosidae | Gnaphosa |
| 2507Wmcc6c | WMCC | July | Araneae | Linyphiidae | Grammonota |
| 2507Wmcc6c | WMCC | July | Araneae | Lycosidae | Rabidosa |
| 2507Wmcc6c | WMCC | July | Araneae | Lycosidae | Schizocosa |
| 2507Wmcc6c | WMCC | July | Araneae | Phrurolithidae | Scotinella |
| 2507Wmcc6c | WMCC | July | Araneae | Salticidae | Maevia |
| 2507Wmcc6c | WMCC | July | Araneae | Salticidae | Sibianor |
| 2507Wmcc6c | WMCC | July | Araneae | Theridiidae | Canalidion |
| 2507Wmcc6c | WMCC | July | Araneae | Theridiidae | Rugathodes |
| 2507Wmcc6c | WMCC | July | Araneae | Theridiidae | Spintharus |
| 2507Wmcc6c | WMCC | July | Araneae | Thomisidae | Mecaphesa |
| 2507Wmcc6c | WMCC | July | Blattodea | Ectobiidae | Parcoblatta |
| 2507Wmcc6c | WMCC | July | Coleoptera | Buprestidae | Pachyschelus |

| | | | | | |
|---|---|---|---|---|---|
| 2507Wmcc6c | WMCC | July | Coleoptera | Carabidae | Apenes |
| 2507Wmcc6c | WMCC | July | Coleoptera | Carabidae | Harpalus |
| 2507Wmcc6c | WMCC | July | Coleoptera | Carabidae | Other |
| 2507Wmcc6c | WMCC | July | Coleoptera | Carabidae | Polyderis |
| 2507Wmcc6c | WMCC | July | Coleoptera | Curculionidae | Conotrachelus |
| 2507Wmcc6c | WMCC | July | Coleoptera | Curculionidae | Cyrtepistomus |
| 2507Wmcc6c | WMCC | July | Coleoptera | Curculionidae | Listronotus |
| 2507Wmcc6c | WMCC | July | Coleoptera | Dytiscidae | Acilius |
| 2507Wmcc6c | WMCC | July | Coleoptera | Lampyridae | Photinus |
| 2507Wmcc6c | WMCC | July | Coleoptera | Scarabaeidae | Dialytes |
| 2507Wmcc6c | WMCC | July | Coleoptera | Scarabaeidae | Melanocanthon |
| 2507Wmcc6c | WMCC | July | Coleoptera | Scirtidae | Contacyphon |
| 2507Wmcc6c | WMCC | July | Diptera | Agromyzidae | Calycomyza |
| 2507Wmcc6c | WMCC | July | Diptera | Agromyzidae | Cerodontha |
| 2507Wmcc6c | WMCC | July | Diptera | Agromyzidae | Japanagromyza |
| 2507Wmcc6c | WMCC | July | Diptera | Agromyzidae | Liriomyza |
| 2507Wmcc6c | WMCC | July | Diptera | Agromyzidae | Nemorimyza |
| 2507Wmcc6c | WMCC | July | Diptera | Agromyzidae | Phytomyza |
| 2507Wmcc6c | WMCC | July | Diptera | Anisopodidae | Mycetobia |
| 2507Wmcc6c | WMCC | July | Diptera | Calliphoridae | Lucilia |
| 2507Wmcc6c | WMCC | July | Diptera | Cecidomyiidae | Other |
| 2507Wmcc6c | WMCC | July | Diptera | Chaoboridae | Chaoborus |
| 2507Wmcc6c | WMCC | July | Diptera | Chaoboridae | Other |
| 2507Wmcc6c | WMCC | July | Diptera | Chironomidae | Ablabesmyia |
| 2507Wmcc6c | WMCC | July | Diptera | Chironomidae | Chironomus |
| 2507Wmcc6c | WMCC | July | Diptera | Chironomidae | Dicrotendipes |
| 2507Wmcc6c | WMCC | July | Diptera | Chironomidae | Endochironomus |
| 2507Wmcc6c | WMCC | July | Diptera | Chironomidae | Glyptotendipes |
| 2507Wmcc6c | WMCC | July | Diptera | Chironomidae | Microtendipes |
| 2507Wmcc6c | WMCC | July | Diptera | Chironomidae | Orthocladius |
| 2507Wmcc6c | WMCC | July | Diptera | Chironomidae | Other |

| | | | | | |
|---|---|---|---|---|---|
| 2507Wmcc6c | WMCC | July | Diptera | Chironomidae | Other |
| 2507Wmcc6c | WMCC | July | Diptera | Chironomidae | Other |
| 2507Wmcc6c | WMCC | July | Diptera | Chironomidae | Other |
| 2507Wmcc6c | WMCC | July | Diptera | Chironomidae | Parachironomus |
| 2507Wmcc6c | WMCC | July | Diptera | Chironomidae | Polypedilum |
| 2507Wmcc6c | WMCC | July | Diptera | Chironomidae | Pseudochironomus |
| 2507Wmcc6c | WMCC | July | Diptera | Chironomidae | Tanytarsus |
| 2507Wmcc6c | WMCC | July | Diptera | Chironomidae | Xenochironomus |
| 2507Wmcc6c | WMCC | July | Diptera | Chloropidae | Apotropina |
| 2507Wmcc6c | WMCC | July | Diptera | Chloropidae | Elachiptera |
| 2507Wmcc6c | WMCC | July | Diptera | Chloropidae | Liohippelates |
| 2507Wmcc6c | WMCC | July | Diptera | Chloropidae | Oscinella |
| 2507Wmcc6c | WMCC | July | Diptera | Culicidae | Coquillettidia |
| 2507Wmcc6c | WMCC | July | Diptera | Culicidae | Culex |
| 2507Wmcc6c | WMCC | July | Diptera | Culicidae | Uranotaenia |
| 2507Wmcc6c | WMCC | July | Diptera | Drosophilidae | Drosophila |
| 2507Wmcc6c | WMCC | July | Diptera | Empididae | Other |
| 2507Wmcc6c | WMCC | July | Diptera | Hybotidae | Platypalpus |
| 2507Wmcc6c | WMCC | July | Diptera | Hybotidae | Syndyas |
| 2507Wmcc6c | WMCC | July | Diptera | Keroplatidae | Other |
| 2507Wmcc6c | WMCC | July | Diptera | Limoniidae | Dicranomyia |
| 2507Wmcc6c | WMCC | July | Diptera | Limoniidae | Dicranophragma |
| 2507Wmcc6c | WMCC | July | Diptera | Limoniidae | Epiphragma |
| 2507Wmcc6c | WMCC | July | Diptera | Limoniidae | Erioptera |
| 2507Wmcc6c | WMCC | July | Diptera | Limoniidae | Metalimnobia |
| 2507Wmcc6c | WMCC | July | Diptera | Limoniidae | Ormosia |
| 2507Wmcc6c | WMCC | July | Diptera | Limoniidae | Other |
| 2507Wmcc6c | WMCC | July | Diptera | Limoniidae | Other |
| 2507Wmcc6c | WMCC | July | Diptera | Limoniidae | Pseudolimnophila |
| 2507Wmcc6c | WMCC | July | Diptera | Muscidae | Mydaea |
| 2507Wmcc6c | WMCC | July | Diptera | Mycetophilidae | Mycomya |

| | | | | | |
|---|---|---|---|---|---|
| 2507Wmcc6c | WMCC | July | Diptera | Mycetophilidae | Neoempheria |
| 2507Wmcc6c | WMCC | July | Diptera | Opomyzidae | Geomyza |
| 2507Wmcc6c | WMCC | July | Diptera | Pallopteridae | Toxonevra |
| 2507Wmcc6c | WMCC | July | Diptera | Phoridae | Diplonevra |
| 2507Wmcc6c | WMCC | July | Diptera | Polleniidae | Pollenia |
| 2507Wmcc6c | WMCC | July | Diptera | Psychodidae | Clogmia |
| 2507Wmcc6c | WMCC | July | Diptera | Psychodidae | Psychoda |
| 2507Wmcc6c | WMCC | July | Diptera | Sarcophagidae | Ravinia |
| 2507Wmcc6c | WMCC | July | Diptera | Sciaridae | Claustropyga |
| 2507Wmcc6c | WMCC | July | Diptera | Sciaridae | Corynoptera |
| 2507Wmcc6c | WMCC | July | Diptera | Sciaridae | Epidapus |
| 2507Wmcc6c | WMCC | July | Diptera | Sciaridae | Other |
| 2507Wmcc6c | WMCC | July | Diptera | Sphaeroceridae | Leptocera |
| 2507Wmcc6c | WMCC | July | Diptera | Syrphidae | Chalcosyrphus |
| 2507Wmcc6c | WMCC | July | Diptera | Syrphidae | Palpada |
| 2507Wmcc6c | WMCC | July | Diptera | Tabanidae | Hybomitra |
| 2507Wmcc6c | WMCC | July | Diptera | Tabanidae | Other |
| 2507Wmcc6c | WMCC | July | Diptera | Tachinidae | Cryptomeigenia |
| 2507Wmcc6c | WMCC | July | Diptera | Tachinidae | Cyzenis |
| 2507Wmcc6c | WMCC | July | Diptera | Tachinidae | Zaira |
| 2507Wmcc6c | WMCC | July | Diptera | Tipulidae | Dolichopeza |
| 2507Wmcc6c | WMCC | July | Diptera | Tipulidae | Tipula |
| 2507Wmcc6c | WMCC | July | Ephemeroptera | Caenidae | Caenis |
| 2507Wmcc6c | WMCC | July | Ephemeroptera | Ephemeridae | Hexagenia |
| 2507Wmcc6c | WMCC | July | Ephemeroptera | Heptageniidae | Epeorus |
| 2507Wmcc6c | WMCC | July | Hemiptera | Clastopteridae | Clastoptera |
| 2507Wmcc6c | WMCC | July | Hemiptera | Lygaeidae | Kleidocerys |
| 2507Wmcc6c | WMCC | July | Hymenoptera | Apidae | Triepeolus |
| 2507Wmcc6c | WMCC | July | Hymenoptera | Braconidae | Bracon |
| 2507Wmcc6c | WMCC | July | Hymenoptera | Braconidae | Chorebus |
| 2507Wmcc6c | WMCC | July | Hymenoptera | Ichneumonidae | Megastylus |

| | | | | | |
|---|---|---|---|---|---|
| 2507Wmcc6c | WMCC | July | Hymenoptera | Ichneumonidae | Polyblastus |
| 2507Wmcc6c | WMCC | July | Hymenoptera | Other | Other |
| 2507Wmcc6c | WMCC | July | Hymenoptera | Tenthredinidae | Dolerus |
| 2507Wmcc6c | WMCC | July | Hymenoptera | Tenthredinidae | Nematus |
| 2507Wmcc6c | WMCC | July | Hymenoptera | Trigonalidae | Orthogonalys |
| 2507Wmcc6c | WMCC | July | Lepidoptera | Crambidae | Blepharomastix |
| 2507Wmcc6c | WMCC | July | Lepidoptera | Crambidae | Parapoynx |
| 2507Wmcc6c | WMCC | July | Lepidoptera | Elachistidae | Elachista |
| 2507Wmcc6c | WMCC | July | Lepidoptera | Erebidae | Panopoda |
| 2507Wmcc6c | WMCC | July | Lepidoptera | Gelechiidae | Chionodes |
| 2507Wmcc6c | WMCC | July | Lepidoptera | Noctuidae | Condica |
| 2507Wmcc6c | WMCC | July | Lepidoptera | Noctuidae | Leuconycta |
| 2507Wmcc6c | WMCC | July | Lepidoptera | Noctuidae | Pyreferra |
| 2507Wmcc6c | WMCC | July | Lepidoptera | Notodontidae | Heterocampa |
| 2507Wmcc6c | WMCC | July | Lepidoptera | Notodontidae | Hyparpax |
| 2507Wmcc6c | WMCC | July | Lepidoptera | Nymphalidae | Oeneis |
| 2507Wmcc6c | WMCC | July | Lepidoptera | Pyralidae | Atheloca |
| 2507Wmcc6c | WMCC | July | Lepidoptera | Pyralidae | Elasmopalpus |
| 2507Wmcc6c | WMCC | July | Lepidoptera | Tischeriidae | Coptotriche |
| 2507Wmcc6c | WMCC | July | Lepidoptera | Tortricidae | Acleris |
| 2507Wmcc6c | WMCC | July | Lepidoptera | Tortricidae | Argyrotaenia |
| 2507Wmcc6c | WMCC | July | Lepidoptera | Tortricidae | Rhyacionia |
| 2507Wmcc6c | WMCC | July | Megaloptera | Corydalidae | Corydalus |
| 2507Wmcc6c | WMCC | July | Neuroptera | Hemerobiidae | Hemerobius |
| 2507Wmcc6c | WMCC | July | Odonata | Libellulidae | Brachymesia |
| 2507Wmcc6c | WMCC | July | Orthoptera | Tettigoniidae | Conocephalus |
| 2507Wmcc6c | WMCC | July | Other | Other | Other |
| 2507Wmcc6c | WMCC | July | Other | Other | Other |
| 2507Wmcc6c | WMCC | July | Plecoptera | Chloroperlidae | Sweltsa |
| 2507Wmcc6c | WMCC | July | Psocodea | Peripsocidae | Peripsocus |
| 2507Wmcc6c | WMCC | July | Sarcoptiformes | Oripodidae | Other |

| | | | | | |
|---|---|---|---|---|---|
| 2507Wmcc6c | WMCC | July | Trichoptera | Dipseudopsidae | Phylocentropus |
| 2507Wmcc6c | WMCC | July | Trichoptera | Hydroptilidae | Orthotrichia |
| 2507Wmcc6c | WMCC | July | Trichoptera | Leptoceridae | Oecetis |
| 2507Wmcc6c | WMCC | July | Trombidiformes | Arrenuridae | Arrenurus |
| 2507Wmcc6c | WMCC | July | Trombidiformes | Aturidae | Aturus |
| 2507Wmcc6c | WMCC | July | Trombidiformes | Axonopsidae | Brachypoda |
| 2507Wmcc6c | WMCC | July | Trombidiformes | Eupodidae | Other |
| 2507Wmcc6c | WMCC | July | Trombidiformes | Feltriidae | Feltria |
| 2507Wmcc6c | WMCC | July | Trombidiformes | Hydryphantidae | Hydryphantes |
| 2507Wmcc6c | WMCC | July | Trombidiformes | Hydryphantidae | Parathyas |
| 2507Wmcc6c | WMCC | July | Trombidiformes | Lebertiidae | Lebertia |
| 2507Wmcc6c | WMCC | July | Trombidiformes | Mideopsidae | Mideopsis |
| 2507Wmcc6c | WMCC | July | Trombidiformes | Pionidae | Piona |
| 2507Wmcc6c | WMCC | July | Trombidiformes | Pionidae | Tiphys |
| 2507Wmcc6c | WMCC | July | Trombidiformes | Sperchontidae | Sperchon |
| 2507Wmcc6c | WMCC | July | Trombidiformes | Tarsonemidae | Phytonemus |
| 2507Wmcc6c | WMCC | July | Trombidiformes | Torrenticolidae | Testudacarus |
| 2507Wmcc6c | WMCC | July | Trombidiformes | Torrenticolidae | Torrenticola |
| 2507Wmcc6c | WMCC | July | Trombidiformes | Unionicolidae | Koenikea |
| 2508Wmcc6b | WMCC | August | Araneae | Agelenidae | Coras |
| 2508Wmcc6b | WMCC | August | Araneae | Amaurobiidae | Callobius |
| 2508Wmcc6b | WMCC | August | Araneae | Anyphaenidae | Anyphaena |
| 2508Wmcc6b | WMCC | August | Araneae | Cicurinidae | Cicurina |
| 2508Wmcc6b | WMCC | August | Araneae | Dictynidae | Dictyna |
| 2508Wmcc6b | WMCC | August | Araneae | Gnaphosidae | Gnaphosa |
| 2508Wmcc6b | WMCC | August | Araneae | Linyphiidae | Grammonota |
| 2508Wmcc6b | WMCC | August | Araneae | Lycosidae | Rabidosa |
| 2508Wmcc6b | WMCC | August | Araneae | Lycosidae | Schizocosa |
| 2508Wmcc6b | WMCC | August | Araneae | Phrurolithidae | Scotinella |
| 2508Wmcc6b | WMCC | August | Araneae | Salticidae | Maevia |
| 2508Wmcc6b | WMCC | August | Araneae | Salticidae | Sibianor |

| | | | | | |
|---|---|---|---|---|---|
| 2508Wmcc6b | WMCC | August | Araneae | Theridiidae | Canalidion |
| 2508Wmcc6b | WMCC | August | Araneae | Theridiidae | Rugathodes |
| 2508Wmcc6b | WMCC | August | Araneae | Theridiidae | Spintharus |
| 2508Wmcc6b | WMCC | August | Araneae | Thomisidae | Mecaphesa |
| 2508Wmcc6b | WMCC | August | Blattodea | Ectobiidae | Parcoblatta |
| 2508Wmcc6b | WMCC | August | Coleoptera | Buprestidae | Pachyschelus |
| 2508Wmcc6b | WMCC | August | Coleoptera | Carabidae | Apenes |
| 2508Wmcc6b | WMCC | August | Coleoptera | Carabidae | Harpalus |
| 2508Wmcc6b | WMCC | August | Coleoptera | Carabidae | Other |
| 2508Wmcc6b | WMCC | August | Coleoptera | Carabidae | Polyderis |
| 2508Wmcc6b | WMCC | August | Coleoptera | Curculionidae | Conotrachelus |
| 2508Wmcc6b | WMCC | August | Coleoptera | Curculionidae | Cyrtepistomus |
| 2508Wmcc6b | WMCC | August | Coleoptera | Curculionidae | Listronotus |
| 2508Wmcc6b | WMCC | August | Coleoptera | Dytiscidae | Acilius |
| 2508Wmcc6b | WMCC | August | Coleoptera | Lampyridae | Photinus |
| 2508Wmcc6b | WMCC | August | Coleoptera | Scarabaeidae | Dialytes |
| 2508Wmcc6b | WMCC | August | Coleoptera | Scarabaeidae | Melanocanthon |
| 2508Wmcc6b | WMCC | August | Coleoptera | Scirtidae | Contacyphon |
| 2508Wmcc6b | WMCC | August | Diptera | Agromyzidae | Calycomyza |
| 2508Wmcc6b | WMCC | August | Diptera | Agromyzidae | Cerodontha |
| 2508Wmcc6b | WMCC | August | Diptera | Agromyzidae | Japanagromyza |
| 2508Wmcc6b | WMCC | August | Diptera | Agromyzidae | Liriomyza |
| 2508Wmcc6b | WMCC | August | Diptera | Agromyzidae | Nemorimyza |
| 2508Wmcc6b | WMCC | August | Diptera | Agromyzidae | Phytomyza |
| 2508Wmcc6b | WMCC | August | Diptera | Anisopodidae | Mycetobia |
| 2508Wmcc6b | WMCC | August | Diptera | Calliphoridae | Lucilia |
| 2508Wmcc6b | WMCC | August | Diptera | Cecidomyiidae | Other |
| 2508Wmcc6b | WMCC | August | Diptera | Chaoboridae | Chaoborus |
| 2508Wmcc6b | WMCC | August | Diptera | Chaoboridae | Other |
| 2508Wmcc6b | WMCC | August | Diptera | Chironomidae | Ablabesmyia |
| 2508Wmcc6b | WMCC | August | Diptera | Chironomidae | Chironomus |

| | | | | | |
|---|---|---|---|---|---|
| 2508Wmcc6b | WMCC | August | Diptera | Chironomidae | Dicrotendipes |
| 2508Wmcc6b | WMCC | August | Diptera | Chironomidae | Endochironomus |
| 2508Wmcc6b | WMCC | August | Diptera | Chironomidae | Glyptotendipes |
| 2508Wmcc6b | WMCC | August | Diptera | Chironomidae | Microtendipes |
| 2508Wmcc6b | WMCC | August | Diptera | Chironomidae | Orthocladius |
| 2508Wmcc6b | WMCC | August | Diptera | Chironomidae | Other |
| 2508Wmcc6b | WMCC | August | Diptera | Chironomidae | Other |
| 2508Wmcc6b | WMCC | August | Diptera | Chironomidae | Other |
| 2508Wmcc6b | WMCC | August | Diptera | Chironomidae | Other |
| 2508Wmcc6b | WMCC | August | Diptera | Chironomidae | Parachironomus |
| 2508Wmcc6b | WMCC | August | Diptera | Chironomidae | Polypedilum |
| 2508Wmcc6b | WMCC | August | Diptera | Chironomidae | Pseudochironomus |
| 2508Wmcc6b | WMCC | August | Diptera | Chironomidae | Tanytarsus |
| 2508Wmcc6b | WMCC | August | Diptera | Chironomidae | Xenochironomus |
| 2508Wmcc6b | WMCC | August | Diptera | Chloropidae | Apotropina |
| 2508Wmcc6b | WMCC | August | Diptera | Chloropidae | Elachiptera |
| 2508Wmcc6b | WMCC | August | Diptera | Chloropidae | Liohippelates |
| 2508Wmcc6b | WMCC | August | Diptera | Chloropidae | Oscinella |
| 2508Wmcc6b | WMCC | August | Diptera | Culicidae | Coquillettidia |
| 2508Wmcc6b | WMCC | August | Diptera | Culicidae | Culex |
| 2508Wmcc6b | WMCC | August | Diptera | Culicidae | Uranotaenia |
| 2508Wmcc6b | WMCC | August | Diptera | Drosophilidae | Drosophila |
| 2508Wmcc6b | WMCC | August | Diptera | Empididae | Other |
| 2508Wmcc6b | WMCC | August | Diptera | Hybotidae | Platypalpus |
| 2508Wmcc6b | WMCC | August | Diptera | Hybotidae | Syndyas |
| 2508Wmcc6b | WMCC | August | Diptera | Keroplatidae | Other |
| 2508Wmcc6b | WMCC | August | Diptera | Limoniidae | Dicranomyia |
| 2508Wmcc6b | WMCC | August | Diptera | Limoniidae | Dicranophragma |
| 2508Wmcc6b | WMCC | August | Diptera | Limoniidae | Epiphragma |
| 2508Wmcc6b | WMCC | August | Diptera | Limoniidae | Erioptera |
| 2508Wmcc6b | WMCC | August | Diptera | Limoniidae | Metalimnobia |

| | | | | | |
|---|---|---|---|---|---|
| 2508Wmcc6b | WMCC | August | Diptera | Limoniidae | Ormosia |
| 2508Wmcc6b | WMCC | August | Diptera | Limoniidae | Other |
| 2508Wmcc6b | WMCC | August | Diptera | Limoniidae | Other |
| 2508Wmcc6b | WMCC | August | Diptera | Limoniidae | Pseudolimnophila |
| 2508Wmcc6b | WMCC | August | Diptera | Muscidae | Mydaea |
| 2508Wmcc6b | WMCC | August | Diptera | Mycetophilidae | Mycomya |
| 2508Wmcc6b | WMCC | August | Diptera | Mycetophilidae | Neoempheria |
| 2508Wmcc6b | WMCC | August | Diptera | Opomyzidae | Geomyza |
| 2508Wmcc6b | WMCC | August | Diptera | Pallopteridae | Toxonevra |
| 2508Wmcc6b | WMCC | August | Diptera | Phoridae | Diplonevra |
| 2508Wmcc6b | WMCC | August | Diptera | Polleniidae | Pollenia |
| 2508Wmcc6b | WMCC | August | Diptera | Psychodidae | Clogmia |
| 2508Wmcc6b | WMCC | August | Diptera | Psychodidae | Psychoda |
| 2508Wmcc6b | WMCC | August | Diptera | Sarcophagidae | Ravinia |
| 2508Wmcc6b | WMCC | August | Diptera | Sciaridae | Claustropyga |
| 2508Wmcc6b | WMCC | August | Diptera | Sciaridae | Corynoptera |
| 2508Wmcc6b | WMCC | August | Diptera | Sciaridae | Epidapus |
| 2508Wmcc6b | WMCC | August | Diptera | Sciaridae | Other |
| 2508Wmcc6b | WMCC | August | Diptera | Sphaeroceridae | Leptocera |
| 2508Wmcc6b | WMCC | August | Diptera | Syrphidae | Chalcosyrphus |
| 2508Wmcc6b | WMCC | August | Diptera | Syrphidae | Palpada |
| 2508Wmcc6b | WMCC | August | Diptera | Tabanidae | Hybomitra |
| 2508Wmcc6b | WMCC | August | Diptera | Tabanidae | Other |
| 2508Wmcc6b | WMCC | August | Diptera | Tachinidae | Cryptomeigenia |
| 2508Wmcc6b | WMCC | August | Diptera | Tachinidae | Cyzenis |
| 2508Wmcc6b | WMCC | August | Diptera | Tachinidae | Zaira |
| 2508Wmcc6b | WMCC | August | Diptera | Tipulidae | Dolichopeza |
| 2508Wmcc6b | WMCC | August | Diptera | Tipulidae | Tipula |
| 2508Wmcc6b | WMCC | August | Ephemeroptera | Caenidae | Caenis |
| 2508Wmcc6b | WMCC | August | Ephemeroptera | Ephemeridae | Hexagenia |
| 2508Wmcc6b | WMCC | August | Ephemeroptera | Heptageniidae | Epeorus |

| | | | | | |
|---|---|---|---|---|---|
| 2508Wmcc6b | WMCC | August | Hemiptera | Clastopteridae | Clastoptera |
| 2508Wmcc6b | WMCC | August | Hemiptera | Lygaeidae | Kleidocerys |
| 2508Wmcc6b | WMCC | August | Hymenoptera | Apidae | Triepeolus |
| 2508Wmcc6b | WMCC | August | Hymenoptera | Braconidae | Bracon |
| 2508Wmcc6b | WMCC | August | Hymenoptera | Braconidae | Chorebus |
| 2508Wmcc6b | WMCC | August | Hymenoptera | Ichneumonidae | Megastylus |
| 2508Wmcc6b | WMCC | August | Hymenoptera | Ichneumonidae | Polyblastus |
| 2508Wmcc6b | WMCC | August | Hymenoptera | Other | Other |
| 2508Wmcc6b | WMCC | August | Hymenoptera | Tenthredinidae | Dolerus |
| 2508Wmcc6b | WMCC | August | Hymenoptera | Tenthredinidae | Nematus |
| 2508Wmcc6b | WMCC | August | Hymenoptera | Trigonalidae | Orthogonalys |
| 2508Wmcc6b | WMCC | August | Lepidoptera | Crambidae | Blepharomastix |
| 2508Wmcc6b | WMCC | August | Lepidoptera | Crambidae | Parapoynx |
| 2508Wmcc6b | WMCC | August | Lepidoptera | Elachistidae | Elachista |
| 2508Wmcc6b | WMCC | August | Lepidoptera | Erebidae | Panopoda |
| 2508Wmcc6b | WMCC | August | Lepidoptera | Gelechiidae | Chionodes |
| 2508Wmcc6b | WMCC | August | Lepidoptera | Noctuidae | Condica |
| 2508Wmcc6b | WMCC | August | Lepidoptera | Noctuidae | Leuconycta |
| 2508Wmcc6b | WMCC | August | Lepidoptera | Noctuidae | Pyreferra |
| 2508Wmcc6b | WMCC | August | Lepidoptera | Notodontidae | Heterocampa |
| 2508Wmcc6b | WMCC | August | Lepidoptera | Notodontidae | Hyparpax |
| 2508Wmcc6b | WMCC | August | Lepidoptera | Nymphalidae | Oeneis |
| 2508Wmcc6b | WMCC | August | Lepidoptera | Pyralidae | Atheloca |
| 2508Wmcc6b | WMCC | August | Lepidoptera | Pyralidae | Elasmopalpus |
| 2508Wmcc6b | WMCC | August | Lepidoptera | Tischeriidae | Coptotriche |
| 2508Wmcc6b | WMCC | August | Lepidoptera | Tortricidae | Acleris |
| 2508Wmcc6b | WMCC | August | Lepidoptera | Tortricidae | Argyrotaenia |
| 2508Wmcc6b | WMCC | August | Lepidoptera | Tortricidae | Rhyacionia |
| 2508Wmcc6b | WMCC | August | Megaloptera | Corydalidae | Corydalus |
| 2508Wmcc6b | WMCC | August | Neuroptera | Hemerobiidae | Hemerobius |
| 2508Wmcc6b | WMCC | August | Odonata | Libellulidae | Brachymesia |

| | | | | | |
|---|---|---|---|---|---|
| 2508Wmcc6b | WMCC | August | Orthoptera | Tettigoniidae | Conocephalus |
| 2508Wmcc6b | WMCC | August | Other | Other | Other |
| 2508Wmcc6b | WMCC | August | Other | Other | Other |
| 2508Wmcc6b | WMCC | August | Plecoptera | Chloroperlidae | Sweltsa |
| 2508Wmcc6b | WMCC | August | Psocodea | Peripsocidae | Peripsocus |
| 2508Wmcc6b | WMCC | August | Sarcoptiformes | Oripodidae | Other |
| 2508Wmcc6b | WMCC | August | Trichoptera | Dipseudopsidae | Phylocentropus |
| 2508Wmcc6b | WMCC | August | Trichoptera | Hydroptilidae | Orthotrichia |
| 2508Wmcc6b | WMCC | August | Trichoptera | Leptoceridae | Oecetis |
| 2508Wmcc6b | WMCC | August | Trombidiformes | Arrenuridae | Arrenurus |
| 2508Wmcc6b | WMCC | August | Trombidiformes | Aturidae | Aturus |
| 2508Wmcc6b | WMCC | August | Trombidiformes | Axonopsidae | Brachypoda |
| 2508Wmcc6b | WMCC | August | Trombidiformes | Eupodidae | Other |
| 2508Wmcc6b | WMCC | August | Trombidiformes | Feltriidae | Feltria |
| 2508Wmcc6b | WMCC | August | Trombidiformes | Hydryphantidae | Hydryphantes |
| 2508Wmcc6b | WMCC | August | Trombidiformes | Hydryphantidae | Parathyas |
| 2508Wmcc6b | WMCC | August | Trombidiformes | Lebertiidae | Lebertia |
| 2508Wmcc6b | WMCC | August | Trombidiformes | Mideopsidae | Mideopsis |
| 2508Wmcc6b | WMCC | August | Trombidiformes | Pionidae | Piona |
| 2508Wmcc6b | WMCC | August | Trombidiformes | Pionidae | Tiphys |
| 2508Wmcc6b | WMCC | August | Trombidiformes | Sperchontidae | Sperchon |
| 2508Wmcc6b | WMCC | August | Trombidiformes | Tarsonemidae | Phytonemus |
| 2508Wmcc6b | WMCC | August | Trombidiformes | Torrenticolidae | Testudacarus |
| 2508Wmcc6b | WMCC | August | Trombidiformes | Torrenticolidae | Torrenticola |
| 2508Wmcc6b | WMCC | August | Trombidiformes | Unionicolidae | Koenikea |
| 2508Wmcc6c | WMCC | August | Araneae | Agelenidae | Coras |
| 2508Wmcc6c | WMCC | August | Araneae | Amaurobiidae | Callobius |
| 2508Wmcc6c | WMCC | August | Araneae | Anyphaenidae | Anyphaena |
| 2508Wmcc6c | WMCC | August | Araneae | Cicurinidae | Cicurina |
| 2508Wmcc6c | WMCC | August | Araneae | Dictynidae | Dictyna |
| 2508Wmcc6c | WMCC | August | Araneae | Gnaphosidae | Gnaphosa |

| | | | | | |
|---|---|---|---|---|---|
| 2508Wmcc6c | WMCC | August | Araneae | Linyphiidae | Grammonota |
| 2508Wmcc6c | WMCC | August | Araneae | Lycosidae | Rabidosa |
| 2508Wmcc6c | WMCC | August | Araneae | Lycosidae | Schizocosa |
| 2508Wmcc6c | WMCC | August | Araneae | Phrurolithidae | Scotinella |
| 2508Wmcc6c | WMCC | August | Araneae | Salticidae | Maevia |
| 2508Wmcc6c | WMCC | August | Araneae | Salticidae | Sibianor |
| 2508Wmcc6c | WMCC | August | Araneae | Theridiidae | Canalidion |
| 2508Wmcc6c | WMCC | August | Araneae | Theridiidae | Rugathodes |
| 2508Wmcc6c | WMCC | August | Araneae | Theridiidae | Spintharus |
| 2508Wmcc6c | WMCC | August | Araneae | Thomisidae | Mecaphesa |
| 2508Wmcc6c | WMCC | August | Blattodea | Ectobiidae | Parcoblatta |
| 2508Wmcc6c | WMCC | August | Coleoptera | Buprestidae | Pachyschelus |
| 2508Wmcc6c | WMCC | August | Coleoptera | Carabidae | Apenes |
| 2508Wmcc6c | WMCC | August | Coleoptera | Carabidae | Harpalus |
| 2508Wmcc6c | WMCC | August | Coleoptera | Carabidae | Other |
| 2508Wmcc6c | WMCC | August | Coleoptera | Carabidae | Polyderis |
| 2508Wmcc6c | WMCC | August | Coleoptera | Curculionidae | Conotrachelus |
| 2508Wmcc6c | WMCC | August | Coleoptera | Curculionidae | Cyrtepistomus |
| 2508Wmcc6c | WMCC | August | Coleoptera | Curculionidae | Listronotus |
| 2508Wmcc6c | WMCC | August | Coleoptera | Dytiscidae | Acilius |
| 2508Wmcc6c | WMCC | August | Coleoptera | Lampyridae | Photinus |
| 2508Wmcc6c | WMCC | August | Coleoptera | Scarabaeidae | Dialytes |
| 2508Wmcc6c | WMCC | August | Coleoptera | Scarabaeidae | Melanocanthon |
| 2508Wmcc6c | WMCC | August | Coleoptera | Scirtidae | Contacyphon |
| 2508Wmcc6c | WMCC | August | Diptera | Agromyzidae | Calycomyza |
| 2508Wmcc6c | WMCC | August | Diptera | Agromyzidae | Cerodontha |
| 2508Wmcc6c | WMCC | August | Diptera | Agromyzidae | Japanagromyza |
| 2508Wmcc6c | WMCC | August | Diptera | Agromyzidae | Liriomyza |
| 2508Wmcc6c | WMCC | August | Diptera | Agromyzidae | Nemorimyza |
| 2508Wmcc6c | WMCC | August | Diptera | Agromyzidae | Phytomyza |
| 2508Wmcc6c | WMCC | August | Diptera | Anisopodidae | Mycetobia |

| | | | | | |
|---|---|---|---|---|---|
| 2508Wmcc6c | WMCC | August | Diptera | Calliphoridae | Lucilia |
| 2508Wmcc6c | WMCC | August | Diptera | Cecidomyiidae | Other |
| 2508Wmcc6c | WMCC | August | Diptera | Chaoboridae | Chaoborus |
| 2508Wmcc6c | WMCC | August | Diptera | Chaoboridae | Other |
| 2508Wmcc6c | WMCC | August | Diptera | Chironomidae | Ablabesmyia |
| 2508Wmcc6c | WMCC | August | Diptera | Chironomidae | Chironomus |
| 2508Wmcc6c | WMCC | August | Diptera | Chironomidae | Dicrotendipes |
| 2508Wmcc6c | WMCC | August | Diptera | Chironomidae | Endochironomus |
| 2508Wmcc6c | WMCC | August | Diptera | Chironomidae | Glyptotendipes |
| 2508Wmcc6c | WMCC | August | Diptera | Chironomidae | Microtendipes |
| 2508Wmcc6c | WMCC | August | Diptera | Chironomidae | Orthocladius |
| 2508Wmcc6c | WMCC | August | Diptera | Chironomidae | Other |
| 2508Wmcc6c | WMCC | August | Diptera | Chironomidae | Other |
| 2508Wmcc6c | WMCC | August | Diptera | Chironomidae | Other |
| 2508Wmcc6c | WMCC | August | Diptera | Chironomidae | Other |
| 2508Wmcc6c | WMCC | August | Diptera | Chironomidae | Parachironomus |
| 2508Wmcc6c | WMCC | August | Diptera | Chironomidae | Polypedilum |
| 2508Wmcc6c | WMCC | August | Diptera | Chironomidae | Pseudochironomus |
| 2508Wmcc6c | WMCC | August | Diptera | Chironomidae | Tanytarsus |
| 2508Wmcc6c | WMCC | August | Diptera | Chironomidae | Xenochironomus |
| 2508Wmcc6c | WMCC | August | Diptera | Chloropidae | Apotropina |
| 2508Wmcc6c | WMCC | August | Diptera | Chloropidae | Elachiptera |
| 2508Wmcc6c | WMCC | August | Diptera | Chloropidae | Liohippelates |
| 2508Wmcc6c | WMCC | August | Diptera | Chloropidae | Oscinella |
| 2508Wmcc6c | WMCC | August | Diptera | Culicidae | Coquillettidia |
| 2508Wmcc6c | WMCC | August | Diptera | Culicidae | Culex |
| 2508Wmcc6c | WMCC | August | Diptera | Culicidae | Uranotaenia |
| 2508Wmcc6c | WMCC | August | Diptera | Drosophilidae | Drosophila |
| 2508Wmcc6c | WMCC | August | Diptera | Empididae | Other |
| 2508Wmcc6c | WMCC | August | Diptera | Hybotidae | Platypalpus |
| 2508Wmcc6c | WMCC | August | Diptera | Hybotidae | Syndyas |

| | | | | | |
|---|---|---|---|---|---|
| 2508Wmcc6c | WMCC | August | Diptera | Keroplatidae | Other |
| 2508Wmcc6c | WMCC | August | Diptera | Limoniidae | Dicranomyia |
| 2508Wmcc6c | WMCC | August | Diptera | Limoniidae | Dicranophragma |
| 2508Wmcc6c | WMCC | August | Diptera | Limoniidae | Epiphragma |
| 2508Wmcc6c | WMCC | August | Diptera | Limoniidae | Erioptera |
| 2508Wmcc6c | WMCC | August | Diptera | Limoniidae | Metalimnobia |
| 2508Wmcc6c | WMCC | August | Diptera | Limoniidae | Ormosia |
| 2508Wmcc6c | WMCC | August | Diptera | Limoniidae | Other |
| 2508Wmcc6c | WMCC | August | Diptera | Limoniidae | Other |
| 2508Wmcc6c | WMCC | August | Diptera | Limoniidae | Pseudolimnophila |
| 2508Wmcc6c | WMCC | August | Diptera | Muscidae | Mydaea |
| 2508Wmcc6c | WMCC | August | Diptera | Mycetophilidae | Mycomya |
| 2508Wmcc6c | WMCC | August | Diptera | Mycetophilidae | Neoempheria |
| 2508Wmcc6c | WMCC | August | Diptera | Opomyzidae | Geomyza |
| 2508Wmcc6c | WMCC | August | Diptera | Pallopteridae | Toxonevra |
| 2508Wmcc6c | WMCC | August | Diptera | Phoridae | Diplonevra |
| 2508Wmcc6c | WMCC | August | Diptera | Polleniidae | Pollenia |
| 2508Wmcc6c | WMCC | August | Diptera | Psychodidae | Clogmia |
| 2508Wmcc6c | WMCC | August | Diptera | Psychodidae | Psychoda |
| 2508Wmcc6c | WMCC | August | Diptera | Sarcophagidae | Ravinia |
| 2508Wmcc6c | WMCC | August | Diptera | Sciaridae | Claustropyga |
| 2508Wmcc6c | WMCC | August | Diptera | Sciaridae | Corynoptera |
| 2508Wmcc6c | WMCC | August | Diptera | Sciaridae | Epidapus |
| 2508Wmcc6c | WMCC | August | Diptera | Sciaridae | Other |
| 2508Wmcc6c | WMCC | August | Diptera | Sphaeroceridae | Leptocera |
| 2508Wmcc6c | WMCC | August | Diptera | Syrphidae | Chalcosyrphus |
| 2508Wmcc6c | WMCC | August | Diptera | Syrphidae | Palpada |
| 2508Wmcc6c | WMCC | August | Diptera | Tabanidae | Hybomitra |
| 2508Wmcc6c | WMCC | August | Diptera | Tabanidae | Other |
| 2508Wmcc6c | WMCC | August | Diptera | Tachinidae | Cryptomeigenia |
| 2508Wmcc6c | WMCC | August | Diptera | Tachinidae | Cyzenis |

| | | | | | |
|---|---|---|---|---|---|
| 2508Wmcc6c | WMCC | August | Diptera | Tachinidae | Zaira |
| 2508Wmcc6c | WMCC | August | Diptera | Tipulidae | Dolichopeza |
| 2508Wmcc6c | WMCC | August | Diptera | Tipulidae | Tipula |
| 2508Wmcc6c | WMCC | August | Ephemeroptera | Caenidae | Caenis |
| 2508Wmcc6c | WMCC | August | Ephemeroptera | Ephemeridae | Hexagenia |
| 2508Wmcc6c | WMCC | August | Ephemeroptera | Heptageniidae | Epeorus |
| 2508Wmcc6c | WMCC | August | Hemiptera | Clastopteridae | Clastoptera |
| 2508Wmcc6c | WMCC | August | Hemiptera | Lygaeidae | Kleidocerys |
| 2508Wmcc6c | WMCC | August | Hymenoptera | Apidae | Triepeolus |
| 2508Wmcc6c | WMCC | August | Hymenoptera | Braconidae | Bracon |
| 2508Wmcc6c | WMCC | August | Hymenoptera | Braconidae | Chorebus |
| 2508Wmcc6c | WMCC | August | Hymenoptera | Ichneumonidae | Megastylus |
| 2508Wmcc6c | WMCC | August | Hymenoptera | Ichneumonidae | Polyblastus |
| 2508Wmcc6c | WMCC | August | Hymenoptera | Other | Other |
| 2508Wmcc6c | WMCC | August | Hymenoptera | Tenthredinidae | Dolerus |
| 2508Wmcc6c | WMCC | August | Hymenoptera | Tenthredinidae | Nematus |
| 2508Wmcc6c | WMCC | August | Hymenoptera | Trigonalidae | Orthogonalys |
| 2508Wmcc6c | WMCC | August | Lepidoptera | Crambidae | Blepharomastix |
| 2508Wmcc6c | WMCC | August | Lepidoptera | Crambidae | Parapoynx |
| 2508Wmcc6c | WMCC | August | Lepidoptera | Elachistidae | Elachista |
| 2508Wmcc6c | WMCC | August | Lepidoptera | Erebidae | Panopoda |
| 2508Wmcc6c | WMCC | August | Lepidoptera | Gelechiidae | Chionodes |
| 2508Wmcc6c | WMCC | August | Lepidoptera | Noctuidae | Condica |
| 2508Wmcc6c | WMCC | August | Lepidoptera | Noctuidae | Leuconycta |
| 2508Wmcc6c | WMCC | August | Lepidoptera | Noctuidae | Pyreferra |
| 2508Wmcc6c | WMCC | August | Lepidoptera | Notodontidae | Heterocampa |
| 2508Wmcc6c | WMCC | August | Lepidoptera | Notodontidae | Hyparpax |
| 2508Wmcc6c | WMCC | August | Lepidoptera | Nymphalidae | Oeneis |
| 2508Wmcc6c | WMCC | August | Lepidoptera | Pyralidae | Atheloca |
| 2508Wmcc6c | WMCC | August | Lepidoptera | Pyralidae | Elasmopalpus |
| 2508Wmcc6c | WMCC | August | Lepidoptera | Tischeriidae | Coptotriche |

| | | | | | |
|---|---|---|---|---|---|
| 2508Wmcc6c | WMCC | August | Lepidoptera | Tortricidae | Acleris |
| 2508Wmcc6c | WMCC | August | Lepidoptera | Tortricidae | Argyrotaenia |
| 2508Wmcc6c | WMCC | August | Lepidoptera | Tortricidae | Rhyacionia |
| 2508Wmcc6c | WMCC | August | Megaloptera | Corydalidae | Corydalus |
| 2508Wmcc6c | WMCC | August | Neuroptera | Hemerobiidae | Hemerobius |
| 2508Wmcc6c | WMCC | August | Odonata | Libellulidae | Brachymesia |
| 2508Wmcc6c | WMCC | August | Orthoptera | Tettigoniidae | Conocephalus |
| 2508Wmcc6c | WMCC | August | Other | Other | Other |
| 2508Wmcc6c | WMCC | August | Other | Other | Other |
| 2508Wmcc6c | WMCC | August | Plecoptera | Chloroperlidae | Sweltsa |
| 2508Wmcc6c | WMCC | August | Psocodea | Peripsocidae | Peripsocus |
| 2508Wmcc6c | WMCC | August | Sarcoptiformes | Oripodidae | Other |
| 2508Wmcc6c | WMCC | August | Trichoptera | Dipseudopsidae | Phylocentropus |
| 2508Wmcc6c | WMCC | August | Trichoptera | Hydroptilidae | Orthotrichia |
| 2508Wmcc6c | WMCC | August | Trichoptera | Leptoceridae | Oecetis |
| 2508Wmcc6c | WMCC | August | Trombidiformes | Arrenuridae | Arrenurus |
| 2508Wmcc6c | WMCC | August | Trombidiformes | Aturidae | Aturus |
| 2508Wmcc6c | WMCC | August | Trombidiformes | Axonopsidae | Brachypoda |
| 2508Wmcc6c | WMCC | August | Trombidiformes | Eupodidae | Other |
| 2508Wmcc6c | WMCC | August | Trombidiformes | Feltriidae | Feltria |
| 2508Wmcc6c | WMCC | August | Trombidiformes | Hydryphantidae | Hydryphantes |
| 2508Wmcc6c | WMCC | August | Trombidiformes | Hydryphantidae | Parathyas |
| 2508Wmcc6c | WMCC | August | Trombidiformes | Lebertiidae | Lebertia |
| 2508Wmcc6c | WMCC | August | Trombidiformes | Mideopsidae | Mideopsis |
| 2508Wmcc6c | WMCC | August | Trombidiformes | Pionidae | Piona |
| 2508Wmcc6c | WMCC | August | Trombidiformes | Pionidae | Tiphys |
| 2508Wmcc6c | WMCC | August | Trombidiformes | Sperchontidae | Sperchon |
| 2508Wmcc6c | WMCC | August | Trombidiformes | Tarsonemidae | Phytonemus |
| 2508Wmcc6c | WMCC | August | Trombidiformes | Torrenticolidae | Testudacarus |
| 2508Wmcc6c | WMCC | August | Trombidiformes | Torrenticolidae | Torrenticola |
| 2508Wmcc6c | WMCC | August | Trombidiformes | Unionicolidae | Koenikea |

| Relative Abundance |
|---:|
| 0.0001 |
| 0.0000 |
| 0.0017 |
| 0.0004 |
| 0.0000 |
| 0.0000 |
| 0.0000 |
| 0.0000 |
| 0.0006 |
| 0.0001 |
| 0.0002 |
| 0.0000 |
| 0.0000 |
| 0.0000 |
| 0.0014 |
| 0.0002 |
| 0.0000 |
| 0.0000 |
| 0.0002 |
| 0.0004 |
| 0.0000 |
| 0.0024 |
| 0.0000 |
| 0.0014 |
| 0.0000 |
| 0.0000 |
| 0.0001 |
| 0.0002 |
| 0.0000 |
| 0.0000 |
| 0.0001 |

| |
|---|
| 0.0000 |
| 0.0000 |
| 0.0000 |
| 0.0006 |
| 0.0011 |
| 0.1083 |
| 0.0016 |
| 0.0071 |
| 0.0095 |
| 0.0000 |
| 0.0012 |
| 0.0397 |
| 0.0006 |
| 0.0008 |
| 0.0000 |
| 0.0000 |
| 0.0000 |
| 0.0026 |
| 0.0010 |
| 0.0000 |
| 0.0212 |
| 0.0032 |
| 0.0196 |
| 0.0000 |
| 0.0020 |
| 0.0062 |
| 0.0010 |
| 0.0000 |
| 0.0004 |
| 0.0002 |
| 0.0000 |

| |
|---|
| 0.0000 |
| 0.0000 |
| 0.0002 |
| 0.0001 |
| 0.0000 |
| 0.0000 |
| 0.0000 |
| 0.0000 |
| 0.0000 |
| 0.0000 |
| 0.0000 |
| 0.0002 |
| 0.0000 |
| 0.0000 |
| 0.0000 |
| 0.0000 |
| 0.0001 |
| 0.0000 |
| 0.0000 |
| 0.0000 |
| 0.0000 |
| 0.0005 |
| 0.0008 |
| 0.0000 |
| 0.0000 |
| 0.0000 |
| 0.0000 |
| 0.0000 |
| 0.0000 |
| 0.0019 |
| 0.0002 |

| |
|---|
| 0.0000 |
| 0.0000 |
| 0.0000 |
| 0.0000 |
| 0.0449 |
| 0.0000 |
| 0.0000 |
| 0.0000 |
| 0.0000 |
| 0.0000 |
| 0.0000 |
| 0.0000 |
| 0.0000 |
| 0.0000 |
| 0.0001 |
| 0.0002 |
| 0.0004 |
| 0.0000 |
| 0.0000 |
| 0.0027 |
| 0.0009 |
| 0.0000 |
| 0.0002 |
| 0.0000 |
| 0.0000 |
| 0.0000 |
| 0.0000 |
| 0.0000 |
| 0.0000 |
| 0.0008 |
| 0.0001 |

| |
|---|
| 0.0000 |
| 0.0000 |
| 0.0000 |
| 0.0010 |
| 0.0026 |
| 0.0000 |
| 0.0006 |
| 0.0000 |
| 0.0003 |
| 0.0000 |
| 0.0001 |
| 0.0000 |
| 0.0027 |
| 0.0014 |
| 0.0014 |
| 0.0000 |
| 0.0000 |
| 0.0009 |
| 0.0000 |
| 0.0000 |
| 0.0000 |
| 0.0000 |
| 0.0000 |
| 0.0000 |
| 0.0002 |
| 0.0004 |
| 0.0009 |
| 0.0000 |
| 0.0014 |
| 0.0000 |
| 0.6938 |

| |
|---|
| 0.0000 |
| 0.0004 |
| 0.0000 |
| 0.0002 |
| 0.0000 |
| 0.0000 |
| 0.0000 |
| 0.0000 |
| 0.0000 |
| 0.0000 |
| 0.0000 |
| 0.0000 |
| 0.0000 |
| 0.0000 |
| 0.0000 |
| 0.0000 |
| 0.0000 |
| 0.0000 |
| 0.0000 |
| 0.0000 |
| 0.0032 |
| 0.0076 |
| 0.0000 |
| 0.0000 |
| 0.0000 |
| 0.0155 |
| 0.0000 |
| 0.0063 |
| 0.0000 |
| 0.0000 |
| 0.0000 |

| |
|---|
| 0.0000 |
| 0.0000 |
| 0.0000 |
| 0.0000 |
| 0.0000 |
| 0.0000 |
| 0.0000 |
| 0.0000 |
| 0.0000 |
| 0.0000 |
| 0.0066 |
| 0.1012 |
| 0.0000 |
| 0.0228 |
| 0.3794 |
| 0.0000 |
| 0.0000 |
| 0.0379 |
| 0.0000 |
| 0.0000 |
| 0.0000 |
| 0.1012 |
| 0.0000 |
| 0.1170 |
| 0.0727 |
| 0.0006 |
| 0.0000 |
| 0.0000 |
| 0.0190 |
| 0.0000 |
| 0.0000 |

| |
|---|
| 0.0006 |
| 0.0000 |
| 0.0000 |
| 0.0032 |
| 0.0032 |
| 0.0000 |
| 0.0000 |
| 0.0000 |
| 0.0000 |
| 0.0057 |
| 0.0000 |
| 0.0000 |
| 0.0000 |
| 0.0000 |
| 0.0000 |
| 0.0190 |
| 0.0000 |
| 0.0000 |
| 0.0000 |
| 0.0000 |
| 0.0000 |
| 0.0000 |
| 0.0000 |
| 0.0000 |
| 0.0000 |
| 0.0000 |
| 0.0000 |
| 0.0028 |
| 0.0000 |
| 0.0000 |
| 0.0000 |

| |
|---|
| 0.0000 |
| 0.0000 |
| 0.0000 |
| 0.0000 |
| 0.0085 |
| 0.0000 |
| 0.0000 |
| 0.0000 |
| 0.0000 |
| 0.0000 |
| 0.0000 |
| 0.0000 |
| 0.0000 |
| 0.0000 |
| 0.0000 |
| 0.0000 |
| 0.0000 |
| 0.0000 |
| 0.0000 |
| 0.0000 |
| 0.0000 |
| 0.0000 |
| 0.0000 |
| 0.0000 |
| 0.0000 |
| 0.0016 |
| 0.0000 |
| 0.0000 |
| 0.0000 |
| 0.0000 |
| 0.0000 |

| |
|---|
| 0.0000 |
| 0.0000 |
| 0.0000 |
| 0.0000 |
| 0.0063 |
| 0.0000 |
| 0.0000 |
| 0.0000 |
| 0.0000 |
| 0.0000 |
| 0.0000 |
| 0.0000 |
| 0.0000 |
| 0.0000 |
| 0.0016 |
| 0.0000 |
| 0.0000 |
| 0.0000 |
| 0.0000 |
| 0.0028 |
| 0.0028 |
| 0.0000 |
| 0.0000 |
| 0.0000 |
| 0.0000 |
| 0.0000 |
| 0.0000 |
| 0.0000 |
| 0.0000 |
| 0.0000 |
| 0.0032 |

| |
|---:|
| 0.0000 |
| 0.0000 |
| 0.0000 |
| 0.0035 |
| 0.0000 |
| 0.0032 |
| 0.0411 |
| 0.0000 |
| 0.0000 |
| 0.0000 |
| 0.0000 |
| 0.0000 |
| 0.0000 |
| 0.0000 |
| 0.0128 |
| 0.0000 |
| 0.0000 |
| 0.0000 |
| 0.0000 |
| 0.0000 |
| 0.0000 |
| 0.0000 |
| 0.0000 |
| 0.0000 |
| 0.0000 |
| 0.0000 |
| 0.0000 |
| 0.0034 |
| 0.0000 |
| 0.0000 |
| 0.0000 |

| |
|---|
| 0.0000 |
| 0.0000 |
| 0.0043 |
| 0.0000 |
| 0.0000 |
| 0.0000 |
| 0.0000 |
| 0.0000 |
| 0.0000 |
| 0.0000 |
| 0.0000 |
| 0.0000 |
| 0.0000 |
| 0.0000 |
| 0.0000 |
| 0.0000 |
| 0.0004 |
| 0.4062 |
| 0.0000 |
| 0.0077 |
| 0.0214 |
| 0.0000 |
| 0.0000 |
| 0.1009 |
| 0.0043 |
| 0.0214 |
| 0.0000 |
| 0.0043 |
| 0.0086 |
| 0.0043 |
| 0.0599 |

| |
|---|
| 0.0000 |
| 0.0385 |
| 0.0000 |
| 0.0043 |
| 0.0000 |
| 0.0000 |
| 0.0000 |
| 0.0000 |
| 0.0000 |
| 0.0000 |
| 0.0000 |
| 0.0000 |
| 0.0000 |
| 0.0000 |
| 0.0000 |
| 0.0000 |
| 0.0171 |
| 0.0000 |
| 0.0000 |
| 0.0000 |
| 0.0000 |
| 0.0000 |
| 0.0000 |
| 0.0000 |
| 0.0043 |
| 0.0000 |
| 0.0000 |
| 0.0000 |
| 0.0000 |
| 0.0000 |
| 0.0000 |

| |
|---|
| 0.0000 |
| 0.0000 |
| 0.0000 |
| 0.0000 |
| 0.0000 |
| 0.0000 |
| 0.0000 |
| 0.0000 |
| 0.0000 |
| 0.0000 |
| 0.0038 |
| 0.0000 |
| 0.0000 |
| 0.0000 |
| 0.0000 |
| 0.0000 |
| 0.0000 |
| 0.0299 |
| 0.0000 |
| 0.0000 |
| 0.0128 |
| 0.0000 |
| 0.0000 |
| 0.0513 |
| 0.0000 |
| 0.0000 |
| 0.0000 |
| 0.0000 |
| 0.0064 |
| 0.0000 |
| 0.0000 |

| |
|---:|
| 0.0000 |
| 0.0000 |
| 0.0000 |
| 0.0000 |
| 0.0000 |
| 0.0000 |
| 0.0000 |
| 0.0000 |
| 0.0000 |
| 0.0000 |
| 0.0000 |
| 0.0000 |
| 0.0000 |
| 0.0000 |
| 0.0000 |
| 0.0000 |
| 0.0000 |
| 0.0000 |
| 0.0000 |
| 0.0000 |
| 0.0000 |
| 0.0000 |
| 0.0000 |
| 0.0000 |
| 0.0000 |
| 0.0000 |
| 0.0000 |
| 0.0000 |
| 0.0000 |
| 0.0000 |
| 0.0000 |

| |
|---|
| 0.0000 |
| 0.0000 |
| 0.0000 |
| 0.0000 |
| 0.0000 |
| 0.0000 |
| 0.0000 |
| 0.0051 |
| 0.0000 |
| 0.1368 |
| 0.0000 |
| 0.0000 |
| 0.0000 |
| 0.0000 |
| 0.0000 |
| 0.0000 |
| 0.0299 |
| 0.0000 |
| 0.0000 |
| 0.0000 |
| 0.1571 |
| 0.0000 |
| 0.0000 |
| 0.0000 |
| 0.0000 |
| 0.0000 |
| 0.0000 |
| 0.0000 |
| 0.0000 |
| 0.0000 |
| 0.0000 |

| |
|---|
| 0.0000 |
| 0.0000 |
| 0.0419 |
| 0.0000 |
| 0.0000 |
| 0.0524 |
| 0.0000 |
| 0.0000 |
| 0.0000 |
| 0.0000 |
| 0.0000 |
| 0.0000 |
| 0.0000 |
| 0.0000 |
| 0.0000 |
| 0.0000 |
| 0.0000 |
| 0.0000 |
| 0.0000 |
| 0.0000 |
| 0.0000 |
| 0.0000 |
| 0.0000 |
| 0.0000 |
| 0.0000 |
| 0.3141 |
| 0.0000 |
| 0.0000 |
| 0.0000 |
| 0.0000 |
| 0.0000 |

| | |
|---|---|
| | 0.0000 |
| | 0.0000 |
| | 0.0000 |
| | 0.0000 |
| | 0.0000 |
| | 0.0000 |
| | 0.0000 |
| | 0.0000 |
| | 0.0000 |
| | 0.0000 |
| | 0.0000 |
| | 0.0000 |
| | 0.0000 |
| | 0.0000 |
| | 0.0000 |
| | 0.0000 |
| | 0.0000 |
| | 0.0000 |
| | 0.0000 |
| | 0.0000 |
| | 0.0000 |
| | 0.0000 |
| | 0.0000 |
| | 0.0000 |
| | 0.0000 |
| | 0.0000 |
| | 0.0000 |
| | 0.0000 |
| | 0.0000 |
| | 0.0000 |
| | 0.0000 |

| |
|---|
| 0.0000 |
| 0.0000 |
| 0.0000 |
| 0.0000 |
| 0.0000 |
| 0.0000 |
| 0.0000 |
| 0.0000 |
| 0.0000 |
| 0.0000 |
| 0.0000 |
| 0.0000 |
| 0.0000 |
| 0.0000 |
| 0.0000 |
| 0.0000 |
| 0.0471 |
| 0.0000 |
| 0.0000 |
| 0.0000 |
| 0.0000 |
| 0.0000 |
| 0.0000 |
| 0.0000 |
| 0.0000 |
| 0.0000 |
| 0.0000 |
| 0.0000 |
| 0.0000 |
| 0.0000 |
| 0.0000 |

| |
|---|
| 0.0000 |
| 0.0314 |
| 0.0000 |
| 0.0000 |
| 0.0000 |
| 0.0052 |
| 0.0000 |
| 0.0000 |
| 0.0000 |
| 0.0000 |
| 0.0000 |
| 0.0000 |
| 0.0000 |
| 0.0000 |
| 0.0000 |
| 0.0000 |
| 0.0000 |
| 0.0000 |
| 0.0000 |
| 0.0000 |
| 0.0000 |
| 0.0000 |
| 0.0000 |
| 0.0000 |
| 0.0000 |
| 0.0000 |
| 0.0000 |
| 0.0000 |
| 0.0000 |
| 0.0000 |
| 0.0000 |

| |
|---:|
| 0.1414 |
| 0.0000 |
| 0.0000 |
| 0.0000 |
| 0.0000 |
| 0.0000 |
| 0.0000 |
| 0.0000 |
| 0.0000 |
| 0.0000 |
| 0.0524 |
| 0.0000 |
| 0.0000 |
| 0.0000 |
| 0.0000 |
| 0.1571 |
| 0.0000 |
| 0.0000 |
| 0.0000 |
| 0.0000 |
| 0.0000 |
| 0.0000 |
| 0.0000 |
| 0.0000 |
| 0.0000 |
| 0.0000 |
| 0.0038 |
| 0.0000 |
| 0.0000 |
| 0.0000 |
| 0.0000 |

| |
|---|
| 0.0000 |
| 0.0000 |
| 0.0000 |
| 0.0000 |
| 0.0000 |
| 0.0000 |
| 0.0000 |
| 0.0000 |
| 0.0065 |
| 0.0553 |
| 0.0000 |
| 0.0038 |
| 0.0000 |
| 0.0000 |
| 0.0115 |
| 0.0000 |
| 0.0000 |
| 0.0000 |
| 0.0000 |
| 0.0000 |
| 0.0000 |
| 0.0000 |
| 0.0004 |
| 0.0000 |
| 0.0000 |
| 0.0000 |
| 0.0000 |
| 0.0000 |
| 0.0000 |
| 0.0461 |
| 0.0000 |

| |
|---|
| 0.0104 |
| 0.0730 |
| 0.0000 |
| 0.0000 |
| 0.2756 |
| 0.0138 |
| 0.0000 |
| 0.0000 |
| 0.0000 |
| 0.0000 |
| 0.0000 |
| 0.0000 |
| 0.0027 |
| 0.0000 |
| 0.0000 |
| 0.0000 |
| 0.0000 |
| 0.0000 |
| 0.0023 |
| 0.0000 |
| 0.0000 |
| 0.0000 |
| 0.0000 |
| 0.0000 |
| 0.0000 |
| 0.0000 |
| 0.0000 |
| 0.0000 |
| 0.0000 |
| 0.0000 |
| 0.0077 |

| |
|---|
| 0.0000 |
| 0.0000 |
| 0.0000 |
| 0.0000 |
| 0.0000 |
| 0.0000 |
| 0.0038 |
| 0.0000 |
| 0.0000 |
| 0.0000 |
| 0.0000 |
| 0.0000 |
| 0.0000 |
| 0.0000 |
| 0.0000 |
| 0.0000 |
| 0.0000 |
| 0.0000 |
| 0.0000 |
| 0.0000 |
| 0.0000 |
| 0.0000 |
| 0.0035 |
| 0.0000 |
| 0.0038 |
| 0.0000 |
| 0.0000 |
| 0.0000 |
| 0.0038 |
| 0.0000 |
| 0.0000 |

| |
|---|
| 0.0000 |
| 0.0000 |
| 0.0000 |
| 0.0000 |
| 0.0000 |
| 0.0000 |
| 0.0000 |
| 0.0000 |
| 0.0000 |
| 0.0000 |
| 0.0000 |
| 0.0000 |
| 0.0000 |
| 0.0000 |
| 0.0000 |
| 0.0000 |
| 0.0000 |
| 0.0000 |
| 0.0000 |
| 0.0000 |
| 0.0000 |
| 0.0000 |
| 0.0038 |
| 0.0000 |
| 0.0000 |
| 0.0807 |
| 0.0000 |
| 0.0000 |
| 0.0000 |
| 0.0000 |
| 0.0038 |

| |
|---|
| 0.0000 |
| 0.0000 |
| 0.0000 |
| 0.0000 |
| 0.0000 |
| 0.0000 |
| 0.0035 |
| 0.0035 |
| 0.0000 |
| 0.0000 |
| 0.0000 |
| 0.0184 |
| 0.0000 |
| 0.0000 |
| 0.0000 |
| 0.0000 |
| 0.0038 |
| 0.0000 |
| 0.0000 |
| 0.0000 |
| 0.0000 |
| 0.0738 |
| 0.0000 |
| 0.0000 |
| 0.2806 |
| 0.0000 |
| 0.0000 |
| 0.0000 |
| 0.0000 |
| 0.0000 |
| 0.0000 |

| |
|---|
| 0.0000 |
| 0.0085 |
| 0.0000 |
| 0.0000 |
| 0.0000 |
| 0.0000 |
| 0.0000 |
| 0.0000 |
| 0.0000 |
| 0.0000 |
| 0.0000 |
| 0.0000 |
| 0.0000 |
| 0.0000 |
| 0.0000 |
| 0.0537 |
| 0.0000 |
| 0.0000 |
| 0.0000 |
| 0.0000 |
| 0.0000 |
| 0.0000 |
| 0.0000 |
| 0.0000 |
| 0.0000 |
| 0.0000 |
| 0.0000 |
| 0.0000 |
| 0.0000 |
| 0.0000 |
| 0.0000 |

| |
|---|
| 0.0000 |
| 0.0000 |
| 0.0000 |
| 0.0000 |
| 0.3922 |
| 0.0000 |
| 0.0162 |
| 0.1023 |
| 0.0000 |
| 0.0000 |
| 0.0000 |
| 0.0000 |
| 0.0000 |
| 0.0000 |
| 0.0000 |
| 0.0000 |
| 0.0000 |
| 0.0000 |
| 0.0000 |
| 0.0000 |
| 0.0000 |
| 0.0000 |
| 0.0000 |
| 0.0000 |
| 0.0000 |
| 0.0000 |
| 0.0000 |
| 0.0085 |
| 0.0000 |
| 0.0000 |
| 0.0000 |

| |
|---|
| 0.0000 |
| 0.0000 |
| 0.0000 |
| 0.0000 |
| 0.0000 |
| 0.0000 |
| 0.0000 |
| 0.0000 |
| 0.0000 |
| 0.0000 |
| 0.0000 |
| 0.0000 |
| 0.0000 |
| 0.0000 |
| 0.0000 |
| 0.0000 |
| 0.0000 |
| 0.0000 |
| 0.0000 |
| 0.0000 |
| 0.0000 |
| 0.0000 |
| 0.0000 |
| 0.0000 |
| 0.0000 |
| 0.0000 |
| 0.0000 |
| 0.0000 |
| 0.0000 |
| 0.0000 |
| 0.0000 |

| |
|---|
| 0.0000 |
| 0.0000 |
| 0.0000 |
| 0.0000 |
| 0.0000 |
| 0.0000 |
| 0.0000 |
| 0.0000 |
| 0.0000 |
| 0.0000 |
| 0.0000 |
| 0.0000 |
| 0.0000 |
| 0.0000 |
| 0.0000 |
| 0.0000 |
| 0.0000 |
| 0.0009 |
| 0.0000 |
| 0.0000 |
| 0.0000 |
| 0.0000 |
| 0.0000 |
| 0.0000 |
| 0.0000 |
| 0.0000 |
| 0.0000 |
| 0.0000 |
| 0.0000 |
| 0.0000 |
| 0.0000 |

| |
|---|
| 0.0000 |
| 0.0000 |
| 0.0000 |
| 0.0000 |
| 0.0000 |
| 0.0000 |
| 0.0085 |
| 0.0000 |
| 0.0000 |
| 0.0000 |
| 0.0000 |
| 0.0000 |
| 0.0000 |
| 0.0000 |
| 0.0000 |
| 0.0000 |
| 0.0000 |
| 0.0000 |
| 0.0000 |
| 0.0000 |
| 0.4092 |
| 0.0000 |
| 0.0000 |
| 0.0000 |
| 0.0000 |
| 0.0000 |
| 0.0000 |
| 0.0000 |
| 0.0000 |
| 0.0000 |
| 0.0000 |

| |
|---:|
| 0.0000 |
| 0.0000 |
| 0.0000 |
| 0.0000 |
| 0.0000 |
| 0.0000 |
| 0.0000 |
| 0.0000 |
| 0.0000 |
| 0.0000 |
| 0.0000 |
| 0.0000 |
| 0.0000 |
| 0.0000 |
| 0.0000 |
| 0.0000 |
| 0.0000 |
| 0.0000 |
| 0.0000 |
| 0.0000 |
| 0.0000 |
| 0.0000 |
| 0.0000 |
| 0.0000 |
| 0.0000 |
| 0.0000 |
| 0.0000 |
| 0.0000 |
| 0.0000 |
| 0.0000 |
| 0.0000 |

| |
|---|
| 0.0000 |
| 0.0000 |
| 0.0000 |
| 0.0000 |
| 0.0000 |
| 0.0000 |
| 0.0000 |
| 0.0000 |
| 0.0000 |
| 0.0000 |
| 0.0709 |
| 0.0000 |
| 0.1277 |
| 0.0000 |
| 0.0000 |
| 0.0000 |
| 0.0000 |
| 0.0000 |
| 0.0000 |
| 0.0000 |
| 0.0000 |
| 0.0000 |
| 0.0000 |
| 0.0000 |
| 0.0000 |
| 0.0000 |
| 0.0000 |
| 0.0000 |
| 0.0000 |
| 0.0000 |
| 0.0000 |

| |
|---|
| 0.0000 |
| 0.0000 |
| 0.0000 |
| 0.0000 |
| 0.0000 |
| 0.0000 |
| 0.0000 |
| 0.0000 |
| 0.0000 |
| 0.0000 |
| 0.0000 |
| 0.0000 |
| 0.0000 |
| 0.0000 |
| 0.0000 |
| 0.0000 |
| 0.0000 |
| 0.0000 |
| 0.0000 |
| 0.0000 |
| 0.0000 |
| 0.0000 |
| 0.0000 |
| 0.0000 |
| 0.0000 |
| 0.0000 |
| 0.0000 |
| 0.0000 |
| 0.0000 |
| 0.0000 |
| 0.0000 |

| |
|---|
| 0.0000 |
| 0.0000 |
| 0.0000 |
| 0.0000 |
| 0.0000 |
| 0.0000 |
| 0.0000 |
| 0.0709 |
| 0.0000 |
| 0.0000 |
| 0.0000 |
| 0.0000 |
| 0.0000 |
| 0.0000 |
| 0.0000 |
| 0.0000 |
| 0.0000 |
| 0.0000 |
| 0.0000 |
| 0.0000 |
| 0.0000 |
| 0.0000 |
| 0.0000 |
| 0.0000 |
| 0.0000 |
| 0.0000 |
| 0.0000 |
| 0.0000 |
| 0.0000 |
| 0.0000 |
| 0.0000 |

| |
|---|
| 0.0284 |
| 0.0000 |
| 0.0000 |
| 0.0000 |
| 0.0000 |
| 0.0000 |
| 0.0000 |
| 0.0000 |
| 0.0000 |
| 0.0000 |
| 0.0000 |
| 0.0000 |
| 0.0000 |
| 0.0000 |
| 0.0000 |
| 0.0000 |
| 0.0000 |
| 0.0000 |
| 0.0638 |
| 0.0000 |
| 0.0000 |
| 0.0000 |
| 0.0000 |
| 0.0000 |
| 0.0000 |
| 0.0000 |
| 0.3546 |
| 0.0000 |
| 0.0000 |
| 0.0000 |
| 0.0000 |

| |
|---|
| 0.0000 |
| 0.0000 |
| 0.0000 |
| 0.0000 |
| 0.0000 |
| 0.0000 |
| 0.0000 |
| 0.0000 |
| 0.0000 |
| 0.2837 |
| 0.0000 |
| 0.0000 |
| 0.0000 |
| 0.0149 |
| 0.0000 |
| 0.0000 |
| 0.0000 |
| 0.0000 |
| 0.0000 |
| 0.0000 |
| 0.0016 |
| 0.0000 |
| 0.0000 |
| 0.0000 |
| 0.0000 |
| 0.0000 |
| 0.0063 |
| 0.0000 |
| 0.0000 |
| 0.0000 |
| 0.0000 |

| |
|---|
| 0.0000 |
| 0.0079 |
| 0.0000 |
| 0.0393 |
| 0.0000 |
| 0.0000 |
| 0.0000 |
| 0.0063 |
| 0.0000 |
| 0.0000 |
| 0.0000 |
| 0.0000 |
| 0.0000 |
| 0.0000 |
| 0.0000 |
| 0.0149 |
| 0.0306 |
| 0.0000 |
| 0.0848 |
| 0.0157 |
| 0.0000 |
| 0.0000 |
| 0.0000 |
| 0.0000 |
| 0.0000 |
| 0.0000 |
| 0.0000 |
| 0.0000 |
| 0.0000 |
| 0.0079 |
| 0.0047 |

| |
|---|
| 0.0000 |
| 0.0000 |
| 0.0000 |
| 0.0000 |
| 0.0000 |
| 0.0000 |
| 0.0079 |
| 0.0000 |
| 0.0000 |
| 0.0079 |
| 0.0000 |
| 0.0000 |
| 0.0000 |
| 0.0000 |
| 0.0000 |
| 0.0000 |
| 0.0000 |
| 0.0000 |
| 0.0000 |
| 0.0000 |
| 0.0000 |
| 0.0000 |
| 0.6991 |
| 0.0000 |
| 0.0000 |
| 0.0000 |
| 0.0000 |
| 0.0000 |
| 0.0000 |
| 0.0000 |
| 0.0000 |

| |
|---|
| 0.0000 |
| 0.0000 |
| 0.0000 |
| 0.0000 |
| 0.0000 |
| 0.0000 |
| 0.0000 |
| 0.0000 |
| 0.0000 |
| 0.0079 |
| 0.0000 |
| 0.0000 |
| 0.0000 |
| 0.0000 |
| 0.0000 |
| 0.0000 |
| 0.0000 |
| 0.0000 |
| 0.0000 |
| 0.0000 |
| 0.0000 |
| 0.0000 |
| 0.0000 |
| 0.0000 |
| 0.0000 |
| 0.0000 |
| 0.0000 |
| 0.0039 |
| 0.0000 |
| 0.0008 |
| 0.0000 |

| |
|---|
| 0.0000 |
| 0.0000 |
| 0.0000 |
| 0.0000 |
| 0.0000 |
| 0.0000 |
| 0.0000 |
| 0.0000 |
| 0.0000 |
| 0.0000 |
| 0.0000 |
| 0.0000 |
| 0.0000 |
| 0.0000 |
| 0.0000 |
| 0.0000 |
| 0.0000 |
| 0.0000 |
| 0.0000 |
| 0.0000 |
| 0.0000 |
| 0.0000 |
| 0.0000 |
| 0.0000 |
| 0.0071 |
| 0.0071 |
| 0.0000 |
| 0.0000 |
| 0.0000 |
| 0.0000 |
| 0.0000 |

| |
|---|
| 0.0000 |
| 0.0000 |
| 0.0000 |
| 0.0000 |
| 0.0000 |
| 0.0000 |
| 0.0000 |
| 0.0000 |
| 0.0157 |
| 0.0079 |
| 0.0000 |
| 0.0000 |
| 0.0000 |
| 0.0000 |
| 0.0000 |
| 0.0000 |
| 0.0000 |
| 0.0000 |
| 0.0000 |
| 0.0338 |
| 0.0000 |
| 0.0000 |
| 0.0000 |
| 0.0000 |
| 0.0000 |
| 0.0000 |
| 0.0169 |
| 0.0000 |
| 0.0000 |
| 0.0000 |
| 0.0000 |

| |
|---|
| 0.0000 |
| 0.0450 |
| 0.0000 |
| 0.0000 |
| 0.0000 |
| 0.0000 |
| 0.0000 |
| 0.0188 |
| 0.0000 |
| 0.0000 |
| 0.0000 |
| 0.0000 |
| 0.0113 |
| 0.0000 |
| 0.0938 |
| 0.0000 |
| 0.0000 |
| 0.0000 |
| 0.0000 |
| 0.0000 |
| 0.0000 |
| 0.0000 |
| 0.0188 |
| 0.0000 |
| 0.1013 |
| 0.2064 |
| 0.0000 |
| 0.0000 |
| 0.0375 |
| 0.0000 |
| 0.0188 |

| |
|---|
| 0.0000 |
| 0.0000 |
| 0.0000 |
| 0.0000 |
| 0.0188 |
| 0.0000 |
| 0.0000 |
| 0.0000 |
| 0.0938 |
| 0.0000 |
| 0.0000 |
| 0.0000 |
| 0.0000 |
| 0.0000 |
| 0.0000 |
| 0.0000 |
| 0.0375 |
| 0.0000 |
| 0.0000 |
| 0.0000 |
| 0.0000 |
| 0.0000 |
| 0.0000 |
| 0.0000 |
| 0.0000 |
| 0.0000 |
| 0.0000 |
| 0.0000 |
| 0.0000 |
| 0.0000 |
| 0.0000 |

| |
|---|
| 0.0000 |
| 0.0000 |
| 0.0000 |
| 0.0000 |
| 0.0000 |
| 0.0000 |
| 0.0000 |
| 0.0169 |
| 0.0000 |
| 0.0000 |
| 0.0000 |
| 0.0000 |
| 0.0000 |
| 0.0000 |
| 0.0000 |
| 0.0000 |
| 0.0000 |
| 0.0000 |
| 0.0000 |
| 0.0000 |
| 0.0000 |
| 0.0000 |
| 0.0000 |
| 0.0000 |
| 0.0000 |
| 0.0000 |
| 0.0000 |
| 0.0000 |
| 0.0000 |
| 0.0000 |
| 0.0000 |

| |
|---|
| 0.0000 |
| 0.0000 |
| 0.0000 |
| 0.0000 |
| 0.0000 |
| 0.0000 |
| 0.0150 |
| 0.0000 |
| 0.0000 |
| 0.0000 |
| 0.0000 |
| 0.0000 |
| 0.0000 |
| 0.0000 |
| 0.0000 |
| 0.0000 |
| 0.0000 |
| 0.0000 |
| 0.0000 |
| 0.0000 |
| 0.0000 |
| 0.0000 |
| 0.0188 |
| 0.0000 |
| 0.0000 |
| 0.0094 |
| 0.0000 |
| 0.0000 |
| 0.0000 |
| 0.0000 |
| 0.0000 |

| |
| ---: |
| 0.0000 |
| 0.0000 |
| 0.0000 |
| 0.0000 |
| 0.0000 |
| 0.0000 |
| 0.0000 |
| 0.0000 |
| 0.0000 |
| 0.0000 |
| 0.0000 |
| 0.0000 |
| 0.0000 |
| 0.0000 |
| 0.0188 |
| 0.0000 |
| 0.0375 |
| 0.0000 |
| 0.0000 |
| 0.0000 |
| 0.0000 |
| 0.0000 |
| 0.1313 |
| 0.0000 |
| 0.0000 |
| 0.0036 |
| 0.0000 |
| 0.0000 |
| 0.0013 |
| 0.0000 |
| 0.0000 |

| |
|---|
| 0.0000 |
| 0.0049 |
| 0.0000 |
| 0.0000 |
| 0.0025 |
| 0.0000 |
| 0.0013 |
| 0.0000 |
| 0.0000 |
| 0.0000 |
| 0.0000 |
| 0.0000 |
| 0.0503 |
| 0.0013 |
| 0.0050 |
| 0.8737 |
| 0.0006 |
| 0.0000 |
| 0.0000 |
| 0.0000 |
| 0.0000 |
| 0.0000 |
| 0.0000 |
| 0.0000 |
| 0.0000 |
| 0.0000 |
| 0.0000 |
| 0.0000 |
| 0.0063 |
| 0.0000 |
| 0.0103 |

| |
|---|
| 0.0025 |
| 0.0000 |
| 0.0000 |
| 0.0013 |
| 0.0000 |
| 0.0000 |
| 0.0000 |
| 0.0000 |
| 0.0000 |
| 0.0013 |
| 0.0013 |
| 0.0000 |
| 0.0000 |
| 0.0000 |
| 0.0000 |
| 0.0000 |
| 0.0000 |
| 0.0000 |
| 0.0000 |
| 0.0000 |
| 0.0000 |
| 0.0000 |
| 0.0025 |
| 0.0000 |
| 0.0000 |
| 0.0010 |
| 0.0000 |
| 0.0000 |
| 0.0000 |
| 0.0000 |
| 0.0000 |

| |
|---|
| 0.0038 |
| 0.0000 |
| 0.0000 |
| 0.0000 |
| 0.0000 |
| 0.0000 |
| 0.0000 |
| 0.0000 |
| 0.0000 |
| 0.0000 |
| 0.0000 |
| 0.0000 |
| 0.0000 |
| 0.0000 |
| 0.0000 |
| 0.0000 |
| 0.0000 |
| 0.0000 |
| 0.0000 |
| 0.0000 |
| 0.0000 |
| 0.0011 |
| 0.0000 |
| 0.0000 |
| 0.0000 |
| 0.0000 |
| 0.0000 |
| 0.0000 |
| 0.0000 |
| 0.0000 |
| 0.0000 |

| |
|---|
| 0.0000 |
| 0.0000 |
| 0.0000 |
| 0.0000 |
| 0.0000 |
| 0.0000 |
| 0.0000 |
| 0.0000 |
| 0.0000 |
| 0.0050 |
| 0.0000 |
| 0.0000 |
| 0.0010 |
| 0.0000 |
| 0.0000 |
| 0.0000 |
| 0.0000 |
| 0.0000 |
| 0.0000 |
| 0.0000 |
| 0.0000 |
| 0.0000 |
| 0.0000 |
| 0.0000 |
| 0.0000 |
| 0.0000 |
| 0.0000 |
| 0.0000 |
| 0.0000 |
| 0.0000 |
| 0.0000 |

| |
|---|
| 0.0006 |
| 0.0045 |
| 0.0000 |
| 0.0000 |
| 0.0000 |
| 0.0068 |
| 0.0011 |
| 0.0000 |
| 0.0000 |
| 0.0000 |
| 0.0000 |
| 0.0000 |
| 0.0000 |
| 0.0000 |
| 0.0000 |
| 0.0000 |
| 0.0000 |
| 0.0000 |
| 0.0000 |
| 0.0000 |
| 0.0013 |
| 0.0000 |
| 0.0013 |
| 0.0000 |
| 0.0000 |
| 0.0025 |
| 0.0000 |
| 0.0000 |
| 0.0000 |
| 0.0000 |
| 0.0002 |

| |
|---|
| 0.0015 |
| 0.0000 |
| 0.0000 |
| 0.0001 |
| 0.0002 |
| 0.0004 |
| 0.0000 |
| 0.0002 |
| 0.0004 |
| 0.0000 |
| 0.0000 |
| 0.0010 |
| 0.0008 |
| 0.0005 |
| 0.0000 |
| 0.0000 |
| 0.0012 |
| 0.0000 |
| 0.0000 |
| 0.0025 |
| 0.0000 |
| 0.8277 |
| 0.0004 |
| 0.0006 |
| 0.0001 |
| 0.0954 |
| 0.0007 |
| 0.0000 |
| 0.0000 |
| 0.0001 |
| 0.0000 |

| |
| ---: |
| 0.0000 |
| 0.0000 |
| 0.0000 |
| 0.0150 |
| 0.0004 |
| 0.0056 |
| 0.0024 |
| 0.0000 |
| 0.0000 |
| 0.0006 |
| 0.0001 |
| 0.0000 |
| 0.0000 |
| 0.0006 |
| 0.0000 |
| 0.0001 |
| 0.0018 |
| 0.0000 |
| 0.0006 |
| 0.0000 |
| 0.0004 |
| 0.0000 |
| 0.0000 |
| 0.0000 |
| 0.0008 |
| 0.0000 |
| 0.0002 |
| 0.0001 |
| 0.0008 |
| 0.0000 |
| 0.0004 |

| |
|---|
| 0.0013 |
| 0.0000 |
| 0.0000 |
| 0.0000 |
| 0.0000 |
| 0.0002 |
| 0.0014 |
| 0.0000 |
| 0.0006 |
| 0.0024 |
| 0.0008 |
| 0.0002 |
| 0.0000 |
| 0.0000 |
| 0.0000 |
| 0.0004 |
| 0.0001 |
| 0.0004 |
| 0.0000 |
| 0.0000 |
| 0.0000 |
| 0.0018 |
| 0.0000 |
| 0.0001 |
| 0.0001 |
| 0.0000 |
| 0.0000 |
| 0.0017 |
| 0.0000 |
| 0.0001 |
| 0.0002 |

| |
|---|
| 0.0000 |
| 0.0000 |
| 0.0001 |
| 0.0000 |
| 0.0001 |
| 0.0000 |
| 0.0005 |
| 0.0000 |
| 0.0065 |
| 0.0000 |
| 0.0000 |
| 0.0000 |
| 0.0000 |
| 0.0000 |
| 0.0001 |
| 0.0000 |
| 0.0000 |
| 0.0000 |
| 0.0004 |
| 0.0006 |
| 0.0001 |
| 0.0000 |
| 0.0000 |
| 0.0001 |
| 0.0000 |
| 0.0000 |
| 0.0000 |
| 0.0000 |
| 0.0001 |
| 0.0000 |
| 0.0000 |

| |
|---|
| 0.0000 |
| 0.0000 |
| 0.0002 |
| 0.0000 |
| 0.0000 |
| 0.0000 |
| 0.0002 |
| 0.0012 |
| 0.0000 |
| 0.0000 |
| 0.0000 |
| 0.0011 |
| 0.0060 |
| 0.0000 |
| 0.0000 |
| 0.0000 |
| 0.0000 |
| 0.0000 |
| 0.0000 |
| 0.0000 |
| 0.0000 |
| 0.0004 |
| 0.0001 |
| 0.0000 |
| 0.0000 |
| 0.0000 |
| 0.0011 |
| 0.0002 |
| 0.0005 |
| 0.0000 |
| 0.0000 |

| |
|---|
| 0.0000 |
| 0.0000 |
| 0.0002 |
| 0.0038 |
| 0.0000 |
| 0.0000 |
| 0.0962 |
| 0.0000 |
| 0.0000 |
| 0.0000 |
| 0.0000 |
| 0.0000 |
| 0.0000 |
| 0.0000 |
| 0.0000 |
| 0.0000 |
| 0.0000 |
| 0.0000 |
| 0.0000 |
| 0.0000 |
| 0.0000 |
| 0.0000 |
| 0.0000 |
| 0.0000 |
| 0.0000 |
| 0.0000 |
| 0.0000 |
| 0.0000 |
| 0.0000 |
| 0.0000 |
| 0.0000 |

| |
|---|
| 0.0000 |
| 0.0000 |
| 0.0000 |
| 0.0000 |
| 0.0000 |
| 0.0000 |
| 0.0000 |
| 0.0000 |
| 0.0000 |
| 0.0000 |
| 0.0000 |
| 0.4327 |
| 0.1923 |
| 0.0000 |
| 0.0000 |
| 0.0000 |
| 0.0000 |
| 0.0000 |
| 0.0000 |
| 0.0000 |
| 0.0000 |
| 0.0000 |
| 0.0000 |
| 0.0000 |
| 0.0000 |
| 0.0000 |
| 0.0000 |
| 0.0000 |
| 0.0000 |
| 0.0000 |
| 0.0000 |

| |
|---|
| 0.0000 |
| 0.0000 |
| 0.0000 |
| 0.0000 |
| 0.0000 |
| 0.0000 |
| 0.0000 |
| 0.0000 |
| 0.0000 |
| 0.0000 |
| 0.0000 |
| 0.0000 |
| 0.0000 |
| 0.0000 |
| 0.0000 |
| 0.0000 |
| 0.0000 |
| 0.0000 |
| 0.0000 |
| 0.0000 |
| 0.0000 |
| 0.0000 |
| 0.0000 |
| 0.0000 |
| 0.0000 |
| 0.0000 |
| 0.0000 |
| 0.0000 |
| 0.0000 |
| 0.0000 |
| 0.0000 |

| |
|---|
| 0.0000 |
| 0.0000 |
| 0.0000 |
| 0.0000 |
| 0.0000 |
| 0.0000 |
| 0.0000 |
| 0.0000 |
| 0.0000 |
| 0.0000 |
| 0.0000 |
| 0.0000 |
| 0.0000 |
| 0.0000 |
| 0.0000 |
| 0.0000 |
| 0.0000 |
| 0.0000 |
| 0.0000 |
| 0.0000 |
| 0.0000 |
| 0.0000 |
| 0.0000 |
| 0.0000 |
| 0.0000 |
| 0.0000 |
| 0.0000 |
| 0.0000 |
| 0.0000 |
| 0.0000 |
| 0.0000 |

| |
|---|
| 0.0000 |
| 0.0000 |
| 0.0000 |
| 0.0000 |
| 0.0000 |
| 0.0000 |
| 0.0000 |
| 0.0000 |
| 0.0000 |
| 0.0000 |
| 0.0000 |
| 0.0000 |
| 0.0000 |
| 0.0000 |
| 0.0000 |
| 0.0000 |
| 0.0000 |
| 0.0865 |
| 0.0000 |
| 0.0000 |
| 0.0000 |
| 0.0000 |
| 0.0000 |
| 0.0000 |
| 0.0000 |
| 0.0000 |
| 0.0000 |
| 0.0000 |
| 0.0000 |
| 0.0000 |
| 0.0000 |

| |
|---|
| 0.0000 |
| 0.1923 |
| 0.0000 |
| 0.0000 |
| 0.0000 |
| 0.0000 |
| 0.0000 |
| 0.0000 |
| 0.0000 |
| 0.0000 |
| 0.0000 |
| 0.0000 |
| 0.0000 |
| 0.0000 |
| 0.0000 |
| 0.0000 |
| 0.0000 |
| 0.0000 |
| 0.0000 |
| 0.0000 |
| 0.0000 |
| 0.0000 |
| 0.0000 |
| 0.0000 |
| 0.0000 |
| 0.0000 |
| 0.0000 |
| 0.0000 |
| 0.0000 |
| 0.0000 |
| 0.0000 |

| |
|---:|
| 0.1149 |
| 0.0000 |
| 0.0000 |
| 0.0000 |
| 0.0000 |
| 0.0000 |
| 0.0000 |
| 0.0000 |
| 0.0000 |
| 0.0000 |
| 0.0000 |
| 0.0000 |
| 0.0000 |
| 0.0000 |
| 0.0000 |
| 0.1149 |
| 0.0000 |
| 0.2069 |
| 0.0000 |
| 0.0000 |
| 0.0000 |
| 0.1034 |
| 0.0000 |
| 0.0000 |
| 0.0000 |
| 0.0000 |
| 0.3448 |
| 0.0000 |
| 0.1149 |
| 0.0000 |
| 0.0000 |

| |
|---:|
| 0.0000 |
| 0.0000 |
| 0.0000 |
| 0.0000 |
| 0.0000 |
| 0.0000 |
| 0.0000 |
| 0.0000 |
| 0.0000 |
| 0.0000 |
| 0.0000 |
| 0.0000 |
| 0.0000 |
| 0.0000 |
| 0.0000 |
| 0.0000 |
| 0.0000 |
| 0.0000 |
| 0.0000 |
| 0.0000 |
| 0.0000 |
| 0.0000 |
| 0.0000 |
| 0.0000 |
| 0.0000 |
| 0.0000 |
| 0.0000 |
| 0.0000 |
| 0.0000 |
| 0.0000 |
| 0.0000 |

| | |
|---|---|
| | 0.0000 |
| | 0.0000 |
| | 0.0000 |
| | 0.0000 |
| | 0.0000 |
| | 0.0000 |
| | 0.0000 |
| | 0.0000 |
| | 0.0000 |
| | 0.0000 |
| | 0.0000 |
| | 0.0000 |
| | 0.0000 |
| | 0.0000 |
| | 0.0000 |
| | 0.0000 |
| | 0.0000 |
| | 0.0000 |
| | 0.0000 |
| | 0.0000 |
| | 0.0000 |
| | 0.0000 |
| | 0.0000 |
| | 0.0000 |
| | 0.0000 |
| | 0.0000 |
| | 0.0000 |
| | 0.0000 |
| | 0.0000 |
| | 0.0000 |
| | 0.0000 |

| |
|---|
| 0.0000 |
| 0.0000 |
| 0.0000 |
| 0.0000 |
| 0.0000 |
| 0.0000 |
| 0.0000 |
| 0.0000 |
| 0.0000 |
| 0.0000 |
| 0.0000 |
| 0.0000 |
| 0.0000 |
| 0.0000 |
| 0.0000 |
| 0.0000 |
| 0.0000 |
| 0.0000 |
| 0.0000 |
| 0.0000 |
| 0.0000 |
| 0.0000 |
| 0.0000 |
| 0.0000 |
| 0.0000 |
| 0.0000 |
| 0.0000 |
| 0.0000 |
| 0.0000 |
| 0.0000 |
| 0.0000 |

| |
|---|
| 0.0000 |
| 0.0000 |
| 0.0000 |
| 0.0000 |
| 0.0000 |
| 0.0000 |
| 0.0000 |
| 0.0000 |
| 0.0000 |
| 0.0000 |
| 0.0000 |
| 0.0000 |
| 0.0000 |
| 0.0000 |
| 0.0000 |
| 0.0000 |
| 0.0000 |
| 0.0000 |
| 0.0000 |
| 0.0000 |
| 0.0000 |
| 0.0000 |
| 0.0000 |
| 0.0000 |
| 0.0000 |
| 0.0000 |
| 0.0000 |
| 0.0000 |
| 0.0000 |
| 0.0000 |
| 0.0980 |

| |
|---|
| 0.0000 |
| 0.0000 |
| 0.0000 |
| 0.0000 |
| 0.0000 |
| 0.0000 |
| 0.0000 |
| 0.0000 |
| 0.1961 |
| 0.0000 |
| 0.0000 |
| 0.0000 |
| 0.0000 |
| 0.0000 |
| 0.0000 |
| 0.0000 |
| 0.0000 |
| 0.0000 |
| 0.0000 |
| 0.0000 |
| 0.0000 |
| 0.2941 |
| 0.0000 |
| 0.0882 |
| 0.0000 |
| 0.0000 |
| 0.0000 |
| 0.0000 |
| 0.0000 |
| 0.0000 |
| 0.0000 |

| |
|---:|
| 0.0000 |
| 0.0000 |
| 0.0000 |
| 0.0000 |
| 0.0000 |
| 0.0000 |
| 0.0000 |
| 0.0000 |
| 0.0000 |
| 0.0000 |
| 0.0000 |
| 0.0000 |
| 0.0000 |
| 0.0000 |
| 0.0000 |
| 0.0000 |
| 0.0000 |
| 0.0000 |
| 0.0000 |
| 0.0000 |
| 0.0000 |
| 0.0000 |
| 0.0000 |
| 0.0000 |
| 0.0000 |
| 0.0000 |
| 0.0000 |
| 0.0000 |
| 0.0000 |
| 0.0000 |
| 0.0000 |

| |
|---|
| 0.0000 |
| 0.0000 |
| 0.0000 |
| 0.0000 |
| 0.0000 |
| 0.0000 |
| 0.0000 |
| 0.0000 |
| 0.0000 |
| 0.0000 |
| 0.0000 |
| 0.0000 |
| 0.0000 |
| 0.0000 |
| 0.0882 |
| 0.0000 |
| 0.0000 |
| 0.0000 |
| 0.0000 |
| 0.0000 |
| 0.0000 |
| 0.0000 |
| 0.0000 |
| 0.0000 |
| 0.0000 |
| 0.0000 |
| 0.0000 |
| 0.0000 |
| 0.0000 |
| 0.0000 |
| 0.0000 |

| |
|---:|
| 0.0000 |
| 0.0000 |
| 0.0000 |
| 0.0000 |
| 0.0000 |
| 0.0000 |
| 0.0000 |
| 0.0000 |
| 0.0000 |
| 0.0000 |
| 0.0000 |
| 0.0000 |
| 0.0000 |
| 0.0000 |
| 0.0000 |
| 0.0000 |
| 0.0000 |
| 0.0000 |
| 0.0000 |
| 0.0000 |
| 0.0000 |
| 0.0000 |
| 0.0000 |
| 0.0000 |
| 0.0490 |
| 0.0000 |
| 0.0000 |
| 0.0000 |
| 0.0000 |
| 0.0882 |
| 0.0000 |

| |
|---|
| 0.0000 |
| 0.0000 |
| 0.0000 |
| 0.0000 |
| 0.0000 |
| 0.0000 |
| 0.0000 |
| 0.0000 |
| 0.0000 |
| 0.0000 |
| 0.0000 |
| 0.0000 |
| 0.0000 |
| 0.0000 |
| 0.0980 |
| 0.0000 |
| 0.0000 |
| 0.0000 |
| 0.0000 |
| 0.0000 |
| 0.0000 |
| 0.0000 |
| 0.0000 |
| 0.0000 |
| 0.2308 |
| 0.0000 |
| 0.0000 |
| 0.0000 |
| 0.0000 |
| 0.0000 |
| 0.0000 |

| | |
|---|---|
| | 0.0000 |
| | 0.0000 |
| | 0.0000 |
| | 0.0000 |
| | 0.0000 |
| | 0.0000 |
| | 0.0000 |
| | 0.0000 |
| | 0.0000 |
| | 0.0000 |
| | 0.0000 |
| | 0.0000 |
| | 0.0000 |
| | 0.0000 |
| | 0.0000 |
| | 0.0000 |
| | 0.0000 |
| | 0.0000 |
| | 0.0000 |
| | 0.0000 |
| | 0.0000 |
| | 0.0000 |
| | 0.0000 |
| | 0.0000 |
| | 0.0000 |
| | 0.0000 |
| | 0.0000 |
| | 0.0000 |
| | 0.0000 |
| | 0.0000 |
| | 0.0000 |

| |
|---|
| 0.0000 |
| 0.0000 |
| 0.0000 |
| 0.0000 |
| 0.0000 |
| 0.0000 |
| 0.0000 |
| 0.0000 |
| 0.0000 |
| 0.0000 |
| 0.0000 |
| 0.0000 |
| 0.0000 |
| 0.0000 |
| 0.0000 |
| 0.0000 |
| 0.0000 |
| 0.0000 |
| 0.0000 |
| 0.0000 |
| 0.0000 |
| 0.0000 |
| 0.0000 |
| 0.0000 |
| 0.0000 |
| 0.0000 |
| 0.0000 |
| 0.0000 |
| 0.0000 |
| 0.0000 |
| 0.0000 |

| |
|---|
| 0.0000 |
| 0.0000 |
| 0.0000 |
| 0.0000 |
| 0.0000 |
| 0.0000 |
| 0.0000 |
| 0.0000 |
| 0.0000 |
| 0.0000 |
| 0.0000 |
| 0.0000 |
| 0.0000 |
| 0.0000 |
| 0.0000 |
| 0.0000 |
| 0.0000 |
| 0.0000 |
| 0.0000 |
| 0.0000 |
| 0.0000 |
| 0.0000 |
| 0.0000 |
| 0.0000 |
| 0.0000 |
| 0.0000 |
| 0.0000 |
| 0.0000 |
| 0.0000 |
| 0.0000 |
| 0.0000 |

| |
|---|
| 0.0000 |
| 0.0000 |
| 0.0000 |
| 0.0000 |
| 0.0000 |
| 0.0000 |
| 0.0000 |
| 0.0000 |
| 0.0000 |
| 0.0000 |
| 0.0000 |
| 0.0000 |
| 0.0000 |
| 0.0000 |
| 0.0000 |
| 0.0000 |
| 0.0000 |
| 0.0000 |
| 0.0000 |
| 0.0000 |
| 0.0000 |
| 0.0000 |
| 0.0000 |
| 0.0000 |
| 0.0000 |
| 0.0000 |
| 0.0000 |
| 0.0000 |
| 0.0000 |
| 0.2564 |
| 0.0000 |

| |
|---|
| 0.0000 |
| 0.0000 |
| 0.0000 |
| 0.0000 |
| 0.0000 |
| 0.0000 |
| 0.0000 |
| 0.0000 |
| 0.0000 |
| 0.0000 |
| 0.0000 |
| 0.0000 |
| 0.0000 |
| 0.0000 |
| 0.0000 |
| 0.0000 |
| 0.0000 |
| 0.0000 |
| 0.0000 |
| 0.0000 |
| 0.0000 |
| 0.0000 |
| 0.5128 |
| 0.0000 |
| 0.0000 |
| 0.0000 |
| 0.0000 |
| 0.0000 |
| 0.0000 |
| 0.0000 |
| 0.0112 |

| |
|---|
| 0.0000 |
| 0.0000 |
| 0.0000 |
| 0.0000 |
| 0.0000 |
| 0.0000 |
| 0.0000 |
| 0.0011 |
| 0.0000 |
| 0.0000 |
| 0.0000 |
| 0.0000 |
| 0.0019 |
| 0.0020 |
| 0.0000 |
| 0.0000 |
| 0.0000 |
| 0.0000 |
| 0.0000 |
| 0.0000 |
| 0.0011 |
| 0.0000 |
| 0.0000 |
| 0.0000 |
| 0.0000 |
| 0.0000 |
| 0.0000 |
| 0.0000 |
| 0.0000 |
| 0.0000 |
| 0.0001 |

| |
|---|
| 0.0000 |
| 0.0000 |
| 0.0011 |
| 0.0000 |
| 0.0080 |
| 0.0046 |
| 0.0000 |
| 0.0000 |
| 0.0011 |
| 0.0000 |
| 0.0000 |
| 0.0000 |
| 0.0000 |
| 0.0000 |
| 0.0000 |
| 0.0000 |
| 0.0000 |
| 0.0000 |
| 0.0000 |
| 0.0000 |
| 0.0000 |
| 0.0000 |
| 0.0000 |
| 0.0000 |
| 0.0000 |
| 0.0091 |
| 0.0000 |
| 0.0000 |
| 0.0011 |
| 0.0000 |
| 0.0000 |

| |
|---:|
| 0.0000 |
| 0.0000 |
| 0.0000 |
| 0.0000 |
| 0.0000 |
| 0.0000 |
| 0.0000 |
| 0.0000 |
| 0.0034 |
| 0.0000 |
| 0.0000 |
| 0.0000 |
| 0.0000 |
| 0.0000 |
| 0.0000 |
| 0.0000 |
| 0.0000 |
| 0.0000 |
| 0.0000 |
| 0.0000 |
| 0.0000 |
| 0.0000 |
| 0.0000 |
| 0.0000 |
| 0.0000 |
| 0.0000 |
| 0.0020 |
| 0.0000 |
| 0.0000 |
| 0.0000 |
| 0.0000 |

| |
|---|
| 0.0000 |
| 0.0000 |
| 0.0000 |
| 0.0000 |
| 0.0000 |
| 0.0000 |
| 0.0000 |
| 0.0000 |
| 0.0000 |
| 0.0000 |
| 0.0000 |
| 0.0000 |
| 0.0000 |
| 0.0000 |
| 0.0000 |
| 0.0000 |
| 0.0000 |
| 0.0000 |
| 0.0000 |
| 0.0000 |
| 0.0000 |
| 0.0000 |
| 0.0000 |
| 0.0000 |
| 0.0000 |
| 0.0000 |
| 0.0000 |
| 0.0000 |
| 0.0001 |
| 0.0000 |
| 0.0005 |

| |
|---|
| 0.0000 |
| 0.0000 |
| 0.0000 |
| 0.4157 |
| 0.0023 |
| 0.0000 |
| 0.0000 |
| 0.0000 |
| 0.0000 |
| 0.0000 |
| 0.0051 |
| 0.0022 |
| 0.0000 |
| 0.0000 |
| 0.0000 |
| 0.0091 |
| 0.0000 |
| 0.0011 |
| 0.0002 |
| 0.0000 |
| 0.0000 |
| 0.0000 |
| 0.0000 |
| 0.0000 |
| 0.0146 |
| 0.0011 |
| 0.4988 |
| 0.0011 |
| 0.0000 |
| 0.0000 |
| 0.0000 |

| |
|---|
| 0.0000 |
| 0.0000 |
| 0.0000 |
| 0.0000 |
| 0.0000 |
| 0.0000 |
| 0.0000 |
| 0.0000 |
| 0.0000 |
| 0.0000 |
| 0.0000 |
| 0.0000 |
| 0.0000 |
| 0.0000 |
| 0.0000 |
| 0.0000 |
| 0.0000 |
| 0.0000 |
| 0.0000 |
| 0.0359 |
| 0.0000 |
| 0.0200 |
| 0.0000 |
| 0.0000 |
| 0.0000 |
| 0.0000 |
| 0.0000 |
| 0.0000 |
| 0.0000 |
| 0.0000 |
| 0.0000 |

| |
|---|
| 0.0000 |
| 0.0000 |
| 0.0000 |
| 0.0000 |
| 0.0000 |
| 0.0000 |
| 0.0000 |
| 0.0000 |
| 0.1597 |
| 0.0000 |
| 0.0000 |
| 0.2794 |
| 0.0000 |
| 0.0000 |
| 0.0978 |
| 0.0000 |
| 0.0000 |
| 0.0000 |
| 0.0000 |
| 0.0000 |
| 0.0000 |
| 0.0000 |
| 0.0639 |
| 0.0599 |
| 0.0000 |
| 0.0000 |
| 0.0000 |
| 0.0000 |
| 0.0000 |
| 0.0000 |
| 0.0000 |

| |
|---|
| 0.0000 |
| 0.0000 |
| 0.0000 |
| 0.0000 |
| 0.0000 |
| 0.0000 |
| 0.0000 |
| 0.0000 |
| 0.0000 |
| 0.0000 |
| 0.0000 |
| 0.0000 |
| 0.0000 |
| 0.0000 |
| 0.0000 |
| 0.0000 |
| 0.0000 |
| 0.0000 |
| 0.0000 |
| 0.0000 |
| 0.0000 |
| 0.0000 |
| 0.0000 |
| 0.0000 |
| 0.0000 |
| 0.0000 |
| 0.0000 |
| 0.0000 |
| 0.0000 |
| 0.0000 |
| 0.0000 |

| |
|---|
| 0.0000 |
| 0.0180 |
| 0.0000 |
| 0.0000 |
| 0.0000 |
| 0.0000 |
| 0.0000 |
| 0.0000 |
| 0.0000 |
| 0.0000 |
| 0.0000 |
| 0.0000 |
| 0.0399 |
| 0.0000 |
| 0.0000 |
| 0.0000 |
| 0.0000 |
| 0.0000 |
| 0.0000 |
| 0.0000 |
| 0.0000 |
| 0.0000 |
| 0.0100 |
| 0.0000 |
| 0.0000 |
| 0.0000 |
| 0.0000 |
| 0.0000 |
| 0.0000 |
| 0.0000 |
| 0.0000 |

| |
|---|
| 0.0000 |
| 0.0000 |
| 0.0000 |
| 0.0000 |
| 0.0000 |
| 0.0000 |
| 0.0000 |
| 0.0160 |
| 0.0200 |
| 0.0000 |
| 0.0000 |
| 0.0000 |
| 0.0000 |
| 0.0000 |
| 0.0000 |
| 0.0000 |
| 0.0000 |
| 0.0000 |
| 0.0000 |
| 0.0000 |
| 0.0000 |
| 0.0000 |
| 0.0000 |
| 0.0000 |
| 0.1796 |
| 0.0000 |
| 0.0000 |
| 0.0000 |
| 0.0000 |
| 0.0000 |
| 0.0000 |

| |
|---|
| 0.0000 |
| 0.0000 |
| 0.0000 |
| 0.0000 |
| 0.0000 |
| 0.0000 |
| 0.0000 |
| 0.0000 |
| 0.0000 |
| 0.0000 |
| 0.0000 |
| 0.0000 |
| 0.0000 |
| 0.0000 |
| 0.0000 |
| 0.0000 |
| 0.0000 |
| 0.0000 |
| 0.0000 |
| 0.0000 |
| 0.0429 |
| 0.0000 |
| 0.0000 |
| 0.0000 |
| 0.1963 |
| 0.0000 |
| 0.0000 |
| 0.0000 |
| 0.0000 |
| 0.0000 |
| 0.0000 |

| |
|---|
| 0.0000 |
| 0.0613 |
| 0.0000 |
| 0.0000 |
| 0.0000 |
| 0.0000 |
| 0.0000 |
| 0.0061 |
| 0.0000 |
| 0.0000 |
| 0.0000 |
| 0.0000 |
| 0.0000 |
| 0.0000 |
| 0.0000 |
| 0.0000 |
| 0.4847 |
| 0.0000 |
| 0.0000 |
| 0.0000 |
| 0.0613 |
| 0.0000 |
| 0.0000 |
| 0.0000 |
| 0.0000 |
| 0.0000 |
| 0.0000 |
| 0.0000 |
| 0.0000 |
| 0.0000 |
| 0.0000 |

| |
|---|
| 0.0000 |
| 0.0000 |
| 0.0000 |
| 0.0000 |
| 0.0000 |
| 0.0000 |
| 0.0000 |
| 0.0000 |
| 0.0000 |
| 0.0000 |
| 0.0000 |
| 0.0000 |
| 0.0000 |
| 0.0000 |
| 0.0000 |
| 0.0000 |
| 0.0000 |
| 0.0000 |
| 0.0000 |
| 0.0000 |
| 0.0000 |
| 0.0000 |
| 0.0000 |
| 0.0000 |
| 0.0000 |
| 0.0000 |
| 0.0000 |
| 0.0000 |
| 0.0000 |
| 0.0000 |
| 0.0000 |

| |
|---|
| 0.0000 |
| 0.0000 |
| 0.0000 |
| 0.0000 |
| 0.0000 |
| 0.0000 |
| 0.0000 |
| 0.0000 |
| 0.0000 |
| 0.0000 |
| 0.0000 |
| 0.0000 |
| 0.0000 |
| 0.0000 |
| 0.0000 |
| 0.0000 |
| 0.0000 |
| 0.0000 |
| 0.0000 |
| 0.0000 |
| 0.0000 |
| 0.0000 |
| 0.0000 |
| 0.0000 |
| 0.0000 |
| 0.0000 |
| 0.0000 |
| 0.0000 |
| 0.0000 |
| 0.0491 |
| 0.0000 |

| |
|---|
| 0.0368 |
| 0.0000 |
| 0.0000 |
| 0.0000 |
| 0.0000 |
| 0.0000 |
| 0.0000 |
| 0.0000 |
| 0.0000 |
| 0.0000 |
| 0.0000 |
| 0.0000 |
| 0.0000 |
| 0.0000 |
| 0.0000 |
| 0.0000 |
| 0.0000 |
| 0.0000 |
| 0.0000 |
| 0.0000 |
| 0.0000 |
| 0.0000 |
| 0.0000 |
| 0.0000 |
| 0.0000 |
| 0.0000 |
| 0.0000 |
| 0.0000 |
| 0.0000 |
| 0.0000 |
| 0.0000 |

| |
|---|
| 0.0000 |
| 0.0613 |
| 0.0000 |
| 0.0000 |
| 0.0000 |
| 0.0000 |
| 0.0000 |
| 0.0000 |
| 0.0000 |
| 0.0000 |
| 0.0000 |
| 0.0000 |
| 0.0000 |
| 0.0000 |
| 0.0000 |
| 0.0000 |
| 0.0000 |
| 0.0000 |
| 0.0000 |
| 0.0000 |
| 0.0000 |
| 0.0000 |
| 0.0000 |
| 0.0000 |
| 0.0000 |
| 0.0000 |
| 0.0000 |
| 0.0000 |
| 0.0000 |
| 0.0000 |
| 0.0000 |

| |
|---|
| 0.0000 |
| 0.0000 |
| 0.0000 |
| 0.0000 |
| 0.0000 |
| 0.2222 |
| 0.0000 |
| 0.2222 |
| 0.0000 |
| 0.0000 |
| 0.0000 |
| 0.0000 |
| 0.0000 |
| 0.0000 |
| 0.0000 |
| 0.0000 |
| 0.0000 |
| 0.0000 |
| 0.0000 |
| 0.0000 |
| 0.0000 |
| 0.0000 |
| 0.3333 |
| 0.0000 |
| 0.0000 |
| 0.0000 |
| 0.0000 |
| 0.0000 |
| 0.0000 |
| 0.0000 |
| 0.0000 |

| |
|---|
| 0.0000 |
| 0.0000 |
| 0.0000 |
| 0.0000 |
| 0.0000 |
| 0.0000 |
| 0.0000 |
| 0.0000 |
| 0.0000 |
| 0.0000 |
| 0.0000 |
| 0.0000 |
| 0.0000 |
| 0.0000 |
| 0.0000 |
| 0.0000 |
| 0.0000 |
| 0.0000 |
| 0.0000 |
| 0.0000 |
| 0.0000 |
| 0.0000 |
| 0.0000 |
| 0.0000 |
| 0.0000 |
| 0.0000 |
| 0.0000 |
| 0.0000 |
| 0.0000 |
| 0.0000 |
| 0.0000 |

| | |
|---|---|
| | 0.0000 |
| | 0.0000 |
| | 0.0000 |
| | 0.0000 |
| | 0.0000 |
| | 0.0000 |
| | 0.0000 |
| | 0.0000 |
| | 0.0000 |
| | 0.0000 |
| | 0.0000 |
| | 0.0000 |
| | 0.0000 |
| | 0.0000 |
| | 0.0000 |
| | 0.0000 |
| | 0.0000 |
| | 0.0000 |
| | 0.0000 |
| | 0.0000 |
| | 0.0000 |
| | 0.0000 |
| | 0.0000 |
| | 0.0000 |
| | 0.0000 |
| | 0.0000 |
| | 0.0000 |
| | 0.0000 |
| | 0.0000 |
| | 0.0000 |
| | 0.0000 |

| |
|---|
| 0.0000 |
| 0.0000 |
| 0.0000 |
| 0.0000 |
| 0.0000 |
| 0.0000 |
| 0.0000 |
| 0.0000 |
| 0.0000 |
| 0.0000 |
| 0.0000 |
| 0.0000 |
| 0.0000 |
| 0.0000 |
| 0.0000 |
| 0.0000 |
| 0.0000 |
| 0.0000 |
| 0.0000 |
| 0.0000 |
| 0.0000 |
| 0.0000 |
| 0.0000 |
| 0.0000 |
| 0.0000 |
| 0.0000 |
| 0.0000 |
| 0.0000 |
| 0.0000 |
| 0.0000 |
| 0.0000 |

| |
|---|
| 0.0000 |
| 0.0000 |
| 0.0000 |
| 0.0000 |
| 0.0000 |
| 0.0000 |
| 0.0000 |
| 0.0000 |
| 0.2222 |
| 0.0000 |
| 0.0000 |
| 0.0000 |
| 0.0000 |
| 0.0000 |
| 0.0000 |
| 0.0000 |
| 0.0000 |
| 0.0000 |
| 0.0000 |
| 0.0000 |
| 0.0000 |
| 0.0000 |
| 0.0000 |
| 0.1957 |
| 0.0000 |
| 0.0000 |
| 0.0000 |
| 0.0000 |
| 0.0000 |
| 0.0000 |
| 0.0000 |

| |
|---|
| 0.0000 |
| 0.0000 |
| 0.0000 |
| 0.0000 |
| 0.2174 |
| 0.0000 |
| 0.0000 |
| 0.0000 |
| 0.0000 |
| 0.0000 |
| 0.0000 |
| 0.0000 |
| 0.0000 |
| 0.0000 |
| 0.0000 |
| 0.0000 |
| 0.0000 |
| 0.0000 |
| 0.0000 |
| 0.0000 |
| 0.0000 |
| 0.0000 |
| 0.0000 |
| 0.0000 |
| 0.0000 |
| 0.0000 |
| 0.0000 |
| 0.0000 |
| 0.3913 |
| 0.0000 |
| 0.0000 |

| |
|---|
| 0.0000 |
| 0.0000 |
| 0.0000 |
| 0.0000 |
| 0.0000 |
| 0.0000 |
| 0.0000 |
| 0.0000 |
| 0.0000 |
| 0.0000 |
| 0.0000 |
| 0.0000 |
| 0.0000 |
| 0.0000 |
| 0.0000 |
| 0.0000 |
| 0.0000 |
| 0.0000 |
| 0.0000 |
| 0.0000 |
| 0.0000 |
| 0.0000 |
| 0.0000 |
| 0.0000 |
| 0.0000 |
| 0.0000 |
| 0.0000 |
| 0.0000 |
| 0.0000 |
| 0.0000 |
| 0.0000 |

| |
|---|
| 0.0000 |
| 0.0000 |
| 0.0000 |
| 0.0000 |
| 0.0000 |
| 0.0000 |
| 0.0000 |
| 0.0000 |
| 0.0000 |
| 0.0000 |
| 0.0000 |
| 0.0000 |
| 0.0000 |
| 0.0000 |
| 0.0000 |
| 0.0000 |
| 0.0000 |
| 0.0000 |
| 0.0000 |
| 0.1957 |
| 0.0000 |
| 0.0000 |
| 0.0000 |
| 0.0000 |
| 0.0000 |
| 0.0000 |
| 0.0000 |
| 0.0000 |
| 0.0000 |
| 0.0000 |
| 0.0000 |

| |
|---|
| 0.0000 |
| 0.0000 |
| 0.0000 |
| 0.0000 |
| 0.0000 |
| 0.0000 |
| 0.0000 |
| 0.0000 |
| 0.0000 |
| 0.0000 |
| 0.0000 |
| 0.0000 |
| 0.0000 |
| 0.0000 |
| 0.0000 |
| 0.0000 |
| 0.0000 |
| 0.0000 |
| 0.0000 |
| 0.0000 |
| 0.0000 |
| 0.0000 |
| 0.0000 |
| 0.0000 |
| 0.0000 |
| 0.0000 |
| 0.0000 |
| 0.0000 |
| 0.0000 |
| 0.0000 |
| 0.0000 |

| |
|---|
| 0.0000 |
| 0.0000 |
| 0.0000 |
| 0.0000 |
| 0.0000 |
| 0.0000 |
| 0.0000 |
| 0.0000 |
| 0.0000 |
| 0.0000 |
| 0.0000 |
| 0.0000 |
| 0.0000 |
| 0.0000 |
| 0.0000 |
| 0.0000 |
| 0.0000 |
| 0.0000 |
| 0.0000 |
| 0.0000 |
| 0.0000 |
| 0.0000 |
| 0.0000 |
| 0.0000 |
| 0.0000 |
| 0.0000 |
| 0.0000 |
| 0.0000 |
| 0.0000 |
| 0.0000 |
| 0.0000 |

| |
| ---: |
| 0.0260 |
| 0.0000 |
| 0.0000 |
| 0.0000 |
| 0.0000 |
| 0.0000 |
| 0.0000 |
| 0.0000 |
| 0.0000 |
| 0.0000 |
| 0.0000 |
| 0.3506 |
| 0.0000 |
| 0.0000 |
| 0.0000 |
| 0.0000 |
| 0.0000 |
| 0.1299 |
| 0.0000 |
| 0.0000 |
| 0.0000 |
| 0.0000 |
| 0.0000 |
| 0.0000 |
| 0.0000 |
| 0.0130 |
| 0.0000 |
| 0.0000 |
| 0.0000 |
| 0.0000 |
| 0.0000 |

| |
|---|
| 0.0000 |
| 0.0000 |
| 0.0000 |
| 0.2338 |
| 0.0000 |
| 0.0000 |
| 0.0000 |
| 0.0000 |
| 0.0000 |
| 0.0000 |
| 0.0000 |
| 0.0000 |
| 0.0000 |
| 0.0000 |
| 0.0000 |
| 0.0000 |
| 0.0000 |
| 0.0000 |
| 0.0000 |
| 0.0000 |
| 0.0000 |
| 0.0000 |
| 0.0000 |
| 0.0000 |
| 0.0000 |
| 0.0000 |
| 0.0000 |
| 0.0000 |
| 0.0000 |
| 0.0000 |
| 0.0000 |

| |
|---|
| 0.0000 |
| 0.0000 |
| 0.0000 |
| 0.0000 |
| 0.0000 |
| 0.0000 |
| 0.0000 |
| 0.0000 |
| 0.0000 |
| 0.0000 |
| 0.0000 |
| 0.0000 |
| 0.0000 |
| 0.0000 |
| 0.0000 |
| 0.0000 |
| 0.0000 |
| 0.0000 |
| 0.0000 |
| 0.0000 |
| 0.0000 |
| 0.0000 |
| 0.0000 |
| 0.0000 |
| 0.0000 |
| 0.1169 |
| 0.0000 |
| 0.0000 |
| 0.0000 |
| 0.0000 |
| 0.0000 |

| |
|---|
| 0.0000 |
| 0.0000 |
| 0.0000 |
| 0.0000 |
| 0.0000 |
| 0.0000 |
| 0.0000 |
| 0.0000 |
| 0.0000 |
| 0.0000 |
| 0.0000 |
| 0.0000 |
| 0.0000 |
| 0.0000 |
| 0.0000 |
| 0.0000 |
| 0.0000 |
| 0.0000 |
| 0.0000 |
| 0.0000 |
| 0.0000 |
| 0.0000 |
| 0.0000 |
| 0.0000 |
| 0.0000 |
| 0.0000 |
| 0.0000 |
| 0.0000 |
| 0.0000 |
| 0.0000 |
| 0.0000 |

| |
|---:|
| 0.0000 |
| 0.0000 |
| 0.0000 |
| 0.0000 |
| 0.0000 |
| 0.0000 |
| 0.0000 |
| 0.0000 |
| 0.0000 |
| 0.0000 |
| 0.0000 |
| 0.0000 |
| 0.0000 |
| 0.0000 |
| 0.0000 |
| 0.0000 |
| 0.0000 |
| 0.0000 |
| 0.0000 |
| 0.0000 |
| 0.0000 |
| 0.0000 |
| 0.0000 |
| 0.0000 |
| 0.1299 |
| 0.0000 |
| 0.0000 |
| 0.0000 |
| 0.0000 |
| 0.0000 |
| 0.0000 |

| |
|---|
| 0.0000 |
| 0.0000 |
| 0.0000 |
| 0.0000 |
| 0.2843 |
| 0.0000 |
| 0.0000 |
| 0.0000 |
| 0.0000 |
| 0.0000 |
| 0.0000 |
| 0.0000 |
| 0.0000 |
| 0.0000 |
| 0.0000 |
| 0.0000 |
| 0.0000 |
| 0.0784 |
| 0.0000 |
| 0.0000 |
| 0.0000 |
| 0.0000 |
| 0.0000 |
| 0.0000 |
| 0.0000 |
| 0.0000 |
| 0.0000 |
| 0.0000 |
| 0.0000 |
| 0.0000 |
| 0.0000 |

| |
|---|
| 0.0000 |
| 0.0000 |
| 0.0000 |
| 0.0000 |
| 0.0000 |
| 0.0000 |
| 0.0000 |
| 0.0000 |
| 0.0000 |
| 0.3529 |
| 0.0000 |
| 0.0000 |
| 0.0000 |
| 0.0000 |
| 0.0000 |
| 0.0000 |
| 0.0000 |
| 0.0000 |
| 0.0000 |
| 0.0000 |
| 0.0000 |
| 0.0000 |
| 0.0000 |
| 0.0000 |
| 0.0000 |
| 0.0000 |
| 0.0000 |
| 0.0000 |
| 0.0000 |
| 0.0000 |
| 0.0000 |

| |
|---|
| 0.0000 |
| 0.0000 |
| 0.0000 |
| 0.0000 |
| 0.0000 |
| 0.0000 |
| 0.0000 |
| 0.0000 |
| 0.0000 |
| 0.0000 |
| 0.0000 |
| 0.0000 |
| 0.0000 |
| 0.0000 |
| 0.0000 |
| 0.0000 |
| 0.0000 |
| 0.0000 |
| 0.0000 |
| 0.0000 |
| 0.0000 |
| 0.0000 |
| 0.0000 |
| 0.0000 |
| 0.0000 |
| 0.0000 |
| 0.0000 |
| 0.0000 |
| 0.0000 |
| 0.0000 |
| 0.0000 |

| |
|---|
| 0.0000 |
| 0.0000 |
| 0.0000 |
| 0.0000 |
| 0.0000 |
| 0.0000 |
| 0.0000 |
| 0.0000 |
| 0.0000 |
| 0.0000 |
| 0.0000 |
| 0.0000 |
| 0.0000 |
| 0.0000 |
| 0.0000 |
| 0.0000 |
| 0.0000 |
| 0.0000 |
| 0.0000 |
| 0.0000 |
| 0.0000 |
| 0.0000 |
| 0.0000 |
| 0.0000 |
| 0.0000 |
| 0.0000 |
| 0.0000 |
| 0.0000 |
| 0.0000 |
| 0.0000 |
| 0.0000 |

| |
|---:|
| 0.0000 |
| 0.0000 |
| 0.0000 |
| 0.0000 |
| 0.0000 |
| 0.0000 |
| 0.0000 |
| 0.0000 |
| 0.0000 |
| 0.0000 |
| 0.0000 |
| 0.0000 |
| 0.0000 |
| 0.0000 |
| 0.0000 |
| 0.0882 |
| 0.0000 |
| 0.0000 |
| 0.0000 |
| 0.0000 |
| 0.0000 |
| 0.0000 |
| 0.0000 |
| 0.0000 |
| 0.0000 |
| 0.0000 |
| 0.0000 |
| 0.0000 |
| 0.0000 |
| 0.0000 |
| 0.1961 |

| |
|---:|
| 0.0000 |
| 0.0000 |
| 0.0000 |
| 0.0000 |
| 0.0000 |
| 0.0000 |
| 0.0000 |
| 0.0000 |
| 0.0000 |
| 0.0000 |
| 0.0074 |
| 0.0000 |
| 0.0004 |
| 0.0000 |
| 0.0000 |
| 0.0000 |
| 0.0000 |
| 0.4371 |
| 0.0039 |
| 0.0000 |
| 0.0000 |
| 0.0000 |
| 0.0059 |
| 0.0031 |
| 0.0000 |
| 0.0000 |
| 0.0000 |
| 0.0000 |
| 0.0000 |
| 0.0020 |
| 0.0000 |

| |
|---|
| 0.0998 |
| 0.0000 |
| 0.0000 |
| 0.0012 |
| 0.0000 |
| 0.0020 |
| 0.0000 |
| 0.0000 |
| 0.0000 |
| 0.0018 |
| 0.0002 |
| 0.0000 |
| 0.0000 |
| 0.0352 |
| 0.0000 |
| 0.0206 |
| 0.0176 |
| 0.0000 |
| 0.0000 |
| 0.0039 |
| 0.0000 |
| 0.0000 |
| 0.0000 |
| 0.0000 |
| 0.0000 |
| 0.0000 |
| 0.0059 |
| 0.0000 |
| 0.0039 |
| 0.0000 |
| 0.0098 |

| |
|---|
| 0.0000 |
| 0.0000 |
| 0.0000 |
| 0.0039 |
| 0.0000 |
| 0.0000 |
| 0.0000 |
| 0.0020 |
| 0.0000 |
| 0.0000 |
| 0.0002 |
| 0.0000 |
| 0.0000 |
| 0.0000 |
| 0.0000 |
| 0.0000 |
| 0.0098 |
| 0.0000 |
| 0.0000 |
| 0.0254 |
| 0.0000 |
| 0.1879 |
| 0.0039 |
| 0.0039 |
| 0.0000 |
| 0.0000 |
| 0.0000 |
| 0.0000 |
| 0.0000 |
| 0.0000 |
| 0.0000 |

| |
|---|
| 0.0000 |
| 0.0000 |
| 0.0000 |
| 0.0000 |
| 0.0023 |
| 0.0006 |
| 0.0053 |
| 0.0000 |
| 0.0020 |
| 0.0000 |
| 0.0196 |
| 0.0059 |
| 0.0000 |
| 0.0000 |
| 0.0000 |
| 0.0215 |
| 0.0002 |
| 0.0000 |
| 0.0000 |
| 0.0000 |
| 0.0078 |
| 0.0014 |
| 0.0000 |
| 0.0000 |
| 0.0000 |
| 0.0000 |
| 0.0000 |
| 0.0020 |
| 0.0000 |
| 0.0000 |
| 0.0000 |

| |
|---|
| 0.0000 |
| 0.0000 |
| 0.0000 |
| 0.0000 |
| 0.0059 |
| 0.0000 |
| 0.0000 |
| 0.0014 |
| 0.0000 |
| 0.0000 |
| 0.0000 |
| 0.0000 |
| 0.0000 |
| 0.0000 |
| 0.0000 |
| 0.0000 |
| 0.0010 |
| 0.0000 |
| 0.0000 |
| 0.0000 |
| 0.0000 |
| 0.0035 |
| 0.0055 |
| 0.0000 |
| 0.0000 |
| 0.0000 |
| 0.0000 |
| 0.0000 |
| 0.0000 |
| 0.0000 |
| 0.0000 |

| |
|---|
| 0.0039 |
| 0.0000 |
| 0.0000 |
| 0.0000 |
| 0.0000 |
| 0.0078 |
| 0.0000 |
| 0.0000 |
| 0.0000 |
| 0.0039 |
| 0.0000 |
| 0.0000 |
| 0.0000 |
| 0.0000 |
| 0.0000 |
| 0.0000 |
| 0.0000 |
| 0.0000 |
| 0.0000 |
| 0.0000 |
| 0.0000 |
| 0.0000 |
| 0.0000 |
| 0.0004 |
| 0.0085 |
| 0.0000 |
| 0.0000 |
| 0.0000 |
| 0.0000 |
| 0.0000 |
| 0.0000 |

| |
|---|
| 0.0038 |
| 0.0000 |
| 0.0000 |
| 0.0000 |
| 0.0085 |
| 0.0000 |
| 0.0021 |
| 0.0000 |
| 0.0000 |
| 0.0000 |
| 0.0000 |
| 0.0000 |
| 0.0000 |
| 0.0000 |
| 0.0000 |
| 0.0000 |
| 0.0000 |
| 0.0000 |
| 0.0129 |
| 0.7038 |
| 0.0000 |
| 0.0038 |
| 0.0063 |
| 0.0000 |
| 0.0000 |
| 0.0233 |
| 0.0021 |
| 0.0042 |
| 0.0762 |
| 0.0000 |
| 0.0000 |

| |
|---|
| 0.0000 |
| 0.0042 |
| 0.0000 |
| 0.0000 |
| 0.0000 |
| 0.0000 |
| 0.0106 |
| 0.0000 |
| 0.0000 |
| 0.0000 |
| 0.0004 |
| 0.0021 |
| 0.0021 |
| 0.0000 |
| 0.0000 |
| 0.0000 |
| 0.0004 |
| 0.0000 |
| 0.0000 |
| 0.0042 |
| 0.0000 |
| 0.0000 |
| 0.0000 |
| 0.0021 |
| 0.0000 |
| 0.0000 |
| 0.0000 |
| 0.0000 |
| 0.0000 |
| 0.0000 |
| 0.0000 |

| |
|---|
| 0.0000 |
| 0.0000 |
| 0.0000 |
| 0.0000 |
| 0.0076 |
| 0.0000 |
| 0.0000 |
| 0.0000 |
| 0.0000 |
| 0.0000 |
| 0.0000 |
| 0.0000 |
| 0.0074 |
| 0.0000 |
| 0.0000 |
| 0.0000 |
| 0.0000 |
| 0.0000 |
| 0.0000 |
| 0.0000 |
| 0.0000 |
| 0.0000 |
| 0.0021 |
| 0.0000 |
| 0.0000 |
| 0.0000 |
| 0.0000 |
| 0.0000 |
| 0.0000 |
| 0.0000 |
| 0.0000 |

| |
|---|
| 0.0000 |
| 0.0002 |
| 0.0074 |
| 0.0000 |
| 0.0000 |
| 0.0000 |
| 0.0000 |
| 0.0000 |
| 0.0000 |
| 0.0000 |
| 0.0000 |
| 0.0000 |
| 0.0000 |
| 0.0000 |
| 0.0000 |
| 0.0000 |
| 0.0000 |
| 0.0000 |
| 0.0237 |
| 0.0000 |
| 0.0000 |
| 0.0000 |
| 0.0000 |
| 0.0000 |
| 0.0000 |
| 0.0000 |
| 0.0000 |
| 0.0019 |
| 0.0021 |
| 0.0042 |
| 0.0000 |

| |
|---|
| 0.0000 |
| 0.0000 |
| 0.0000 |
| 0.0000 |
| 0.0000 |
| 0.0000 |
| 0.0000 |
| 0.0548 |
| 0.0000 |
| 0.0000 |
| 0.0000 |
| 0.0042 |
| 0.0000 |
| 0.0000 |
| 0.0000 |
| 0.0000 |
| 0.0000 |
| 0.0000 |
| 0.0021 |
| 0.0000 |
| 0.0044 |
| 0.0000 |
| 0.0211 |
| 0.0000 |
| 0.0022 |
| 0.0000 |
| 0.0000 |
| 0.0000 |
| 0.0000 |
| 0.0000 |
| 0.0111 |

| |
|---|
| 0.0000 |
| 0.0000 |
| 0.0000 |
| 0.0111 |
| 0.0356 |
| 0.0000 |
| 0.0000 |
| 0.0000 |
| 0.0000 |
| 0.0000 |
| 0.0333 |
| 0.0000 |
| 0.0222 |
| 0.0000 |
| 0.0000 |
| 0.0000 |
| 0.0000 |
| 0.0000 |
| 0.0000 |
| 0.0000 |
| 0.0000 |
| 0.0000 |
| 0.0000 |
| 0.0000 |
| 0.0000 |
| 0.0889 |
| 0.0000 |
| 0.3789 |
| 0.0333 |
| 0.0000 |
| 0.0000 |

| |
|---|
| 0.1667 |
| 0.0000 |
| 0.0000 |
| 0.0000 |
| 0.0000 |
| 0.0000 |
| 0.0000 |
| 0.0000 |
| 0.0000 |
| 0.0000 |
| 0.0000 |
| 0.0000 |
| 0.0000 |
| 0.0000 |
| 0.0000 |
| 0.0111 |
| 0.0000 |
| 0.0000 |
| 0.0000 |
| 0.0000 |
| 0.0000 |
| 0.0000 |
| 0.0000 |
| 0.0000 |
| 0.0000 |
| 0.0000 |
| 0.0000 |
| 0.0000 |
| 0.0000 |
| 0.0000 |
| 0.0000 |

| |
|---|
| 0.0000 |
| 0.0000 |
| 0.0000 |
| 0.0000 |
| 0.0000 |
| 0.0000 |
| 0.0000 |
| 0.0000 |
| 0.0000 |
| 0.0000 |
| 0.0000 |
| 0.0000 |
| 0.0000 |
| 0.0000 |
| 0.0000 |
| 0.0111 |
| 0.0000 |
| 0.0000 |
| 0.0178 |
| 0.0000 |
| 0.0000 |
| 0.0000 |
| 0.0000 |
| 0.0000 |
| 0.0000 |
| 0.0000 |
| 0.0000 |
| 0.0000 |
| 0.0011 |
| 0.0000 |
| 0.0000 |

| |
| ---: |
| 0.0000 |
| 0.0000 |
| 0.0000 |
| 0.0000 |
| 0.0100 |
| 0.0000 |
| 0.0000 |
| 0.0011 |
| 0.0000 |
| 0.0000 |
| 0.0000 |
| 0.0000 |
| 0.0000 |
| 0.0000 |
| 0.0000 |
| 0.0000 |
| 0.0000 |
| 0.0000 |
| 0.0000 |
| 0.0000 |
| 0.0000 |
| 0.0000 |
| 0.0000 |
| 0.0000 |
| 0.0000 |
| 0.0000 |
| 0.0000 |
| 0.0000 |
| 0.0000 |
| 0.0000 |
| 0.0000 |

| |
|---|
| 0.0000 |
| 0.0000 |
| 0.0400 |
| 0.0211 |
| 0.0000 |
| 0.0000 |
| 0.0000 |
| 0.0000 |
| 0.0000 |
| 0.0000 |
| 0.0000 |
| 0.0000 |
| 0.0000 |
| 0.0000 |
| 0.0000 |
| 0.0000 |
| 0.0000 |
| 0.0778 |
| 0.0000 |
| 0.0000 |
| 0.0000 |
| 0.0000 |
| 0.0000 |
| 0.0000 |
| 0.0000 |
| 0.0000 |
| 0.0000 |
| 0.0000 |
| 0.0213 |
| 0.0000 |
| 0.0000 |

| |
|---|
| 0.0000 |
| 0.0000 |
| 0.0000 |
| 0.0000 |
| 0.0000 |
| 0.0000 |
| 0.0000 |
| 0.0000 |
| 0.0000 |
| 0.0000 |
| 0.0180 |
| 0.0000 |
| 0.0000 |
| 0.0000 |
| 0.0000 |
| 0.0000 |
| 0.0337 |
| 0.0000 |
| 0.0112 |
| 0.0000 |
| 0.0000 |
| 0.0000 |
| 0.0000 |
| 0.0000 |
| 0.0000 |
| 0.0000 |
| 0.0000 |
| 0.0000 |
| 0.0000 |
| 0.0000 |
| 0.0000 |

| |
|---|
| 0.0000 |
| 0.0000 |
| 0.0202 |
| 0.0449 |
| 0.0809 |
| 0.0000 |
| 0.0000 |
| 0.0000 |
| 0.0000 |
| 0.0000 |
| 0.0000 |
| 0.0000 |
| 0.0000 |
| 0.0000 |
| 0.0000 |
| 0.0000 |
| 0.0000 |
| 0.0000 |
| 0.0000 |
| 0.0000 |
| 0.0000 |
| 0.0000 |
| 0.0000 |
| 0.0000 |
| 0.0000 |
| 0.0000 |
| 0.0000 |
| 0.0000 |
| 0.0000 |
| 0.0000 |
| 0.0000 |

| |
|---|
| 0.0000 |
| 0.0000 |
| 0.0000 |
| 0.0000 |
| 0.0000 |
| 0.0000 |
| 0.0000 |
| 0.0000 |
| 0.0000 |
| 0.0000 |
| 0.0000 |
| 0.0000 |
| 0.0000 |
| 0.0112 |
| 0.0000 |
| 0.0000 |
| 0.0000 |
| 0.0000 |
| 0.0000 |
| 0.3933 |
| 0.0000 |
| 0.0000 |
| 0.0000 |
| 0.0000 |
| 0.0101 |
| 0.0000 |
| 0.0112 |
| 0.0000 |
| 0.0000 |
| 0.0000 |
| 0.0000 |

| |
|---|
| 0.0000 |
| 0.0000 |
| 0.0000 |
| 0.0000 |
| 0.0000 |
| 0.0225 |
| 0.0000 |
| 0.0000 |
| 0.0000 |
| 0.0000 |
| 0.0101 |
| 0.0000 |
| 0.0000 |
| 0.0000 |
| 0.0000 |
| 0.0000 |
| 0.0000 |
| 0.0000 |
| 0.0000 |
| 0.0449 |
| 0.0000 |
| 0.0000 |
| 0.0000 |
| 0.0000 |
| 0.0000 |
| 0.0000 |
| 0.0000 |
| 0.0000 |
| 0.0000 |
| 0.0000 |
| 0.0000 |

| |
|---|
| 0.0112 |
| 0.0112 |
| 0.0000 |
| 0.0112 |
| 0.0000 |
| 0.0000 |
| 0.0000 |
| 0.0000 |
| 0.0303 |
| 0.0000 |
| 0.0000 |
| 0.0000 |
| 0.0000 |
| 0.0000 |
| 0.1910 |
| 0.0000 |
| 0.0000 |
| 0.0000 |
| 0.0000 |
| 0.0000 |
| 0.0000 |
| 0.0000 |
| 0.0000 |
| 0.0112 |
| 0.0000 |
| 0.0000 |
| 0.0000 |
| 0.0000 |
| 0.0000 |
| 0.0000 |
| 0.0000 |

| |
|---|
| 0.0000 |
| 0.0000 |
| 0.0000 |
| 0.1091 |
| 0.0000 |
| 0.0000 |
| 0.0000 |
| 0.0000 |
| 0.0000 |
| 0.0000 |
| 0.0121 |
| 0.0000 |
| 0.0000 |
| 0.0000 |
| 0.0000 |
| 0.0000 |
| 0.0485 |
| 0.0000 |
| 0.0000 |
| 0.1212 |
| 0.0000 |
| 0.0000 |
| 0.0000 |
| 0.0000 |
| 0.0000 |
| 0.0000 |
| 0.0000 |
| 0.0000 |
| 0.0000 |
| 0.0000 |
| 0.0000 |

| |
|---|
| 0.0000 |
| 0.0000 |
| 0.0000 |
| 0.0000 |
| 0.0000 |
| 0.0000 |
| 0.0606 |
| 0.0000 |
| 0.2727 |
| 0.1818 |
| 0.0000 |
| 0.0000 |
| 0.0000 |
| 0.0000 |
| 0.0000 |
| 0.0000 |
| 0.0000 |
| 0.0000 |
| 0.0000 |
| 0.0000 |
| 0.0000 |
| 0.0000 |
| 0.0000 |
| 0.0000 |
| 0.0000 |
| 0.0000 |
| 0.0000 |
| 0.0000 |
| 0.0000 |
| 0.0000 |
| 0.0000 |

| |
|---|
| 0.0000 |
| 0.0000 |
| 0.0000 |
| 0.0000 |
| 0.0000 |
| 0.0000 |
| 0.0000 |
| 0.0000 |
| 0.0000 |
| 0.0000 |
| 0.0000 |
| 0.0000 |
| 0.0000 |
| 0.0000 |
| 0.0000 |
| 0.0000 |
| 0.0000 |
| 0.0000 |
| 0.0000 |
| 0.0000 |
| 0.0000 |
| 0.0000 |
| 0.0000 |
| 0.0000 |
| 0.0000 |
| 0.0000 |
| 0.0000 |
| 0.0000 |
| 0.0000 |
| 0.0000 |
| 0.0000 |

| |
|---|
| 0.0000 |
| 0.0000 |
| 0.0000 |
| 0.0000 |
| 0.0000 |
| 0.0000 |
| 0.0000 |
| 0.0000 |
| 0.0000 |
| 0.0000 |
| 0.0000 |
| 0.0000 |
| 0.0000 |
| 0.0000 |
| 0.0000 |
| 0.0000 |
| 0.0000 |
| 0.0000 |
| 0.0000 |
| 0.0061 |
| 0.0000 |
| 0.0485 |
| 0.0000 |
| 0.0000 |
| 0.0000 |
| 0.0000 |
| 0.0000 |
| 0.0000 |
| 0.0000 |
| 0.0000 |
| 0.0000 |

| |
|---|
| 0.0000 |
| 0.0000 |
| 0.0000 |
| 0.0000 |
| 0.0000 |
| 0.0242 |
| 0.0000 |
| 0.0000 |
| 0.0000 |
| 0.0000 |
| 0.0000 |
| 0.0000 |
| 0.0000 |
| 0.0000 |
| 0.0545 |
| 0.0000 |
| 0.0000 |
| 0.0000 |
| 0.0000 |
| 0.0000 |
| 0.0000 |
| 0.0000 |
| 0.0000 |
| 0.0000 |
| 0.0000 |
| 0.0000 |
| 0.0000 |
| 0.0000 |
| 0.0000 |
| 0.0606 |
| 0.0000 |

| |
|---|
| 0.0000 |
| 0.0000 |
| 0.0000 |
| 0.0000 |
| 0.0000 |
| 0.0000 |
| 0.0000 |
| 0.0000 |
| 0.0000 |
| 0.0031 |
| 0.0000 |
| 0.0000 |
| 0.0000 |
| 0.0000 |
| 0.0000 |
| 0.0000 |
| 0.0000 |
| 0.0000 |
| 0.0000 |
| 0.0000 |
| 0.0000 |
| 0.0031 |
| 0.0000 |
| 0.0000 |
| 0.0000 |
| 0.0000 |
| 0.0000 |
| 0.0000 |
| 0.0000 |
| 0.0000 |
| 0.0031 |

| |
|---|
| 0.0000 |
| 0.0000 |
| 0.0000 |
| 0.0000 |
| 0.0000 |
| 0.0000 |
| 0.0000 |
| 0.0000 |
| 0.0000 |
| 0.0009 |
| 0.0000 |
| 0.0000 |
| 0.0000 |
| 0.0000 |
| 0.0225 |
| 0.5116 |
| 0.0000 |
| 0.0000 |
| 0.0059 |
| 0.0000 |
| 0.0000 |
| 0.0000 |
| 0.0000 |
| 0.0000 |
| 0.0000 |
| 0.0000 |
| 0.0000 |
| 0.0000 |
| 0.0000 |
| 0.0000 |
| 0.0000 |

| |
|---|
| 0.0000 |
| 0.0000 |
| 0.0031 |
| 0.0000 |
| 0.0000 |
| 0.0000 |
| 0.0000 |
| 0.0000 |
| 0.0000 |
| 0.0000 |
| 0.0000 |
| 0.0000 |
| 0.0000 |
| 0.0000 |
| 0.0000 |
| 0.0000 |
| 0.0000 |
| 0.0000 |
| 0.0031 |
| 0.0000 |
| 0.0000 |
| 0.0000 |
| 0.0000 |
| 0.0000 |
| 0.0000 |
| 0.0000 |
| 0.0000 |
| 0.0000 |
| 0.0000 |
| 0.0000 |
| 0.0000 |

| |
|---|
| 0.0000 |
| 0.0000 |
| 0.0000 |
| 0.0000 |
| 0.0000 |
| 0.0028 |
| 0.0000 |
| 0.0000 |
| 0.0000 |
| 0.0000 |
| 0.0000 |
| 0.0000 |
| 0.0000 |
| 0.0000 |
| 0.0000 |
| 0.0000 |
| 0.0000 |
| 0.0031 |
| 0.0000 |
| 0.0000 |
| 0.0000 |
| 0.0000 |
| 0.0000 |
| 0.0031 |
| 0.0000 |
| 0.0000 |
| 0.0000 |
| 0.0000 |
| 0.0000 |
| 0.0000 |
| 0.0000 |

| |
|---|
| 0.0000 |
| 0.0000 |
| 0.0000 |
| 0.0000 |
| 0.0000 |
| 0.0000 |
| 0.0000 |
| 0.0000 |
| 0.0000 |
| 0.0000 |
| 0.0000 |
| 0.0000 |
| 0.0000 |
| 0.0000 |
| 0.0000 |
| 0.0000 |
| 0.0000 |
| 0.0000 |
| 0.0000 |
| 0.0000 |
| 0.0028 |
| 0.0000 |
| 0.0000 |
| 0.0000 |
| 0.0000 |
| 0.0000 |
| 0.0000 |
| 0.4253 |
| 0.0000 |
| 0.0000 |
| 0.0000 |

| |
|---|
| 0.0000 |
| 0.0000 |
| 0.0000 |
| 0.0000 |
| 0.0031 |
| 0.0000 |
| 0.0031 |
| 0.0000 |
| 0.0000 |
| 0.0000 |
| 0.0000 |
| 0.0000 |
| 0.0000 |
| 0.0000 |
| 0.0006 |
| 0.0017 |
| 0.0000 |
| 0.0000 |
| 0.0006 |
| 0.0000 |
| 0.0000 |
| 0.0000 |
| 0.0000 |
| 0.0000 |
| 0.0000 |
| 0.0000 |
| 0.0000 |
| 0.0013 |
| 0.0010 |
| 0.0000 |
| 0.0000 |

| |
|---|
| 0.0000 |
| 0.0000 |
| 0.0000 |
| 0.0006 |
| 0.0000 |
| 0.0013 |
| 0.0000 |
| 0.0000 |
| 0.0000 |
| 0.0000 |
| 0.0000 |
| 0.0000 |
| 0.0000 |
| 0.0000 |
| 0.0000 |
| 0.0019 |
| 0.0000 |
| 0.0160 |
| 0.0025 |
| 0.0000 |
| 0.0097 |
| 0.9149 |
| 0.0000 |
| 0.0000 |
| 0.0000 |
| 0.0000 |
| 0.0000 |
| 0.0000 |
| 0.0000 |
| 0.0000 |
| 0.0000 |

| |
|---|
| 0.0000 |
| 0.0000 |
| 0.0013 |
| 0.0000 |
| 0.0000 |
| 0.0000 |
| 0.0000 |
| 0.0000 |
| 0.0006 |
| 0.0000 |
| 0.0000 |
| 0.0000 |
| 0.0000 |
| 0.0000 |
| 0.0000 |
| 0.0000 |
| 0.0000 |
| 0.0000 |
| 0.0000 |
| 0.0000 |
| 0.0000 |
| 0.0000 |
| 0.0000 |
| 0.0000 |
| 0.0000 |
| 0.0000 |
| 0.0000 |
| 0.0000 |
| 0.0000 |
| 0.0000 |
| 0.0000 |

| |
|---:|
| 0.0006 |
| 0.0000 |
| 0.0019 |
| 0.0000 |
| 0.0000 |
| 0.0000 |
| 0.0000 |
| 0.0000 |
| 0.0000 |
| 0.0000 |
| 0.0000 |
| 0.0015 |
| 0.0000 |
| 0.0000 |
| 0.0000 |
| 0.0000 |
| 0.0000 |
| 0.0000 |
| 0.0000 |
| 0.0000 |
| 0.0000 |
| 0.0000 |
| 0.0000 |
| 0.0000 |
| 0.0000 |
| 0.0000 |
| 0.0004 |
| 0.0000 |
| 0.0000 |
| 0.0000 |
| 0.0000 |

| |
|---|
| 0.0000 |
| 0.0000 |
| 0.0005 |
| 0.0000 |
| 0.0008 |
| 0.0029 |
| 0.0000 |
| 0.0000 |
| 0.0000 |
| 0.0000 |
| 0.0000 |
| 0.0000 |
| 0.0000 |
| 0.0000 |
| 0.0000 |
| 0.0000 |
| 0.0000 |
| 0.0000 |
| 0.0000 |
| 0.0006 |
| 0.0000 |
| 0.0003 |
| 0.0000 |
| 0.0000 |
| 0.0000 |
| 0.0000 |
| 0.0028 |
| 0.0006 |
| 0.0000 |
| 0.0287 |
| 0.0000 |

| |
|---|
| 0.0006 |
| 0.0000 |
| 0.0000 |
| 0.0000 |
| 0.0000 |
| 0.0000 |
| 0.0000 |
| 0.0000 |
| 0.0000 |
| 0.0000 |
| 0.0031 |
| 0.0000 |
| 0.0006 |
| 0.0000 |
| 0.0000 |
| 0.0000 |
| 0.0000 |
| 0.0000 |
| 0.0000 |
| 0.0000 |
| 0.0000 |
| 0.0000 |
| 0.0000 |
| 0.0000 |
| 0.0000 |
| 0.0000 |
| 0.0000 |
| 0.0000 |
| 0.0000 |
| 0.0000 |
| 0.0000 |

| |
|---|
| 0.0000 |
| 0.0000 |
| 0.0000 |
| 0.0000 |
| 0.0000 |
| 0.0000 |
| 0.0000 |
| 0.0000 |
| 0.0000 |
| 0.0000 |
| 0.0000 |
| 0.0000 |
| 0.0000 |
| 0.0000 |
| 0.0000 |
| 0.0000 |
| 0.0000 |
| 0.0000 |
| 0.0000 |
| 0.0000 |
| 0.0000 |
| 0.0000 |
| 0.0000 |
| 0.0000 |
| 0.0000 |
| 0.0000 |
| 0.2195 |
| 0.2927 |
| 0.0000 |
| 0.0000 |
| 0.0000 |

| |
|---|
| 0.0000 |
| 0.0000 |
| 0.0000 |
| 0.0000 |
| 0.0000 |
| 0.0000 |
| 0.0000 |
| 0.0000 |
| 0.0000 |
| 0.0000 |
| 0.0000 |
| 0.0000 |
| 0.0000 |
| 0.0000 |
| 0.2439 |
| 0.0000 |
| 0.0000 |
| 0.0000 |
| 0.0000 |
| 0.0000 |
| 0.0000 |
| 0.0000 |
| 0.0000 |
| 0.0000 |
| 0.0000 |
| 0.0000 |
| 0.0000 |
| 0.0000 |
| 0.0000 |
| 0.0000 |
| 0.0000 |

| |
|---|
| 0.0000 |
| 0.0000 |
| 0.0000 |
| 0.0000 |
| 0.0000 |
| 0.0000 |
| 0.0000 |
| 0.0000 |
| 0.0000 |
| 0.0000 |
| 0.0000 |
| 0.0000 |
| 0.0000 |
| 0.0000 |
| 0.0000 |
| 0.0000 |
| 0.0000 |
| 0.0000 |
| 0.0000 |
| 0.0000 |
| 0.0000 |
| 0.0000 |
| 0.0000 |
| 0.0000 |
| 0.0000 |
| 0.0000 |
| 0.0000 |
| 0.0000 |
| 0.0000 |
| 0.0000 |
| 0.0000 |

| |
|---|
| 0.0000 |
| 0.0000 |
| 0.0000 |
| 0.0000 |
| 0.0000 |
| 0.0000 |
| 0.0000 |
| 0.0000 |
| 0.0000 |
| 0.0000 |
| 0.0000 |
| 0.0000 |
| 0.0000 |
| 0.0000 |
| 0.0000 |
| 0.0000 |
| 0.0000 |
| 0.0000 |
| 0.0000 |
| 0.0000 |
| 0.0000 |
| 0.0000 |
| 0.0000 |
| 0.0000 |
| 0.0000 |
| 0.0000 |
| 0.0000 |
| 0.0000 |
| 0.0000 |
| 0.0000 |
| 0.0000 |

| |
|---|
| 0.0000 |
| 0.0000 |
| 0.0000 |
| 0.0000 |
| 0.0000 |
| 0.0000 |
| 0.0000 |
| 0.0000 |
| 0.0000 |
| 0.0000 |
| 0.0000 |
| 0.0000 |
| 0.0000 |
| 0.0000 |
| 0.0000 |
| 0.0000 |
| 0.0000 |
| 0.0000 |
| 0.0000 |
| 0.0000 |
| 0.2439 |
| 0.0000 |
| 0.0000 |
| 0.0000 |
| 0.0000 |
| 0.0000 |
| 0.0000 |
| 0.0000 |
| 0.0000 |
| 0.0000 |
| 0.0000 |

| |
|---|
| 0.0000 |
| 0.0000 |
| 0.0000 |
| 0.0000 |
| 0.0000 |
| 0.0000 |
| 0.0000 |
| 0.0000 |
| 0.0000 |
| 0.0000 |
| 0.0000 |
| 0.0000 |
| 0.0000 |
| 0.0000 |
| 0.0000 |
| 0.0000 |
| 0.0000 |
| 0.0000 |
| 0.0000 |
| 0.0000 |
| 0.0000 |
| 0.0000 |
| 0.0000 |
| 0.0000 |
| 0.0000 |
| 0.0000 |
| 0.0000 |
| 0.0000 |
| 0.0000 |
| 0.0000 |
| 0.0000 |

| |
|---|
| 0.0000 |
| 0.0056 |
| 0.1444 |
| 0.0000 |
| 0.0000 |
| 0.0024 |
| 0.0000 |
| 0.0000 |
| 0.0000 |
| 0.0000 |
| 0.0000 |
| 0.0000 |
| 0.0000 |
| 0.0000 |
| 0.0000 |
| 0.0000 |
| 0.0000 |
| 0.0000 |
| 0.0000 |
| 0.0000 |
| 0.0000 |
| 0.0000 |
| 0.0000 |
| 0.0000 |
| 0.0000 |
| 0.0000 |
| 0.0000 |
| 0.0000 |
| 0.0000 |
| 0.0000 |
| 0.0000 |

| |
|---:|
| 0.0000 |
| 0.0000 |
| 0.0000 |
| 0.0000 |
| 0.0000 |
| 0.0000 |
| 0.0000 |
| 0.0000 |
| 0.0000 |
| 0.0000 |
| 0.0000 |
| 0.0000 |
| 0.0000 |
| 0.0000 |
| 0.0000 |
| 0.0000 |
| 0.0000 |
| 0.0000 |
| 0.0000 |
| 0.0000 |
| 0.0000 |
| 0.0000 |
| 0.0000 |
| 0.0000 |
| 0.0000 |
| 0.0000 |
| 0.0000 |
| 0.0000 |
| 0.0000 |
| 0.0000 |
| 0.0000 |

| |
|---|
| 0.0000 |
| 0.0000 |
| 0.0000 |
| 0.0000 |
| 0.0000 |
| 0.0000 |
| 0.0000 |
| 0.0000 |
| 0.0000 |
| 0.0000 |
| 0.0000 |
| 0.0000 |
| 0.0000 |
| 0.0005 |
| 0.0000 |
| 0.0000 |
| 0.0000 |
| 0.0000 |
| 0.0000 |
| 0.0000 |
| 0.0000 |
| 0.0000 |
| 0.0000 |
| 0.0000 |
| 0.0000 |
| 0.0000 |
| 0.0000 |
| 0.0000 |
| 0.0000 |
| 0.0000 |
| 0.0000 |

| |
|---|
| 0.0000 |
| 0.0000 |
| 0.0012 |
| 0.0000 |
| 0.0000 |
| 0.0000 |
| 0.0000 |
| 0.0022 |
| 0.0022 |
| 0.0000 |
| 0.0000 |
| 0.0000 |
| 0.0000 |
| 0.0000 |
| 0.0000 |
| 0.0012 |
| 0.0122 |
| 0.0024 |
| 0.0017 |
| 0.0000 |
| 0.0000 |
| 0.0000 |
| 0.7830 |
| 0.0000 |
| 0.0000 |
| 0.0000 |
| 0.0000 |
| 0.0000 |
| 0.0244 |
| 0.0166 |
| 0.0000 |
