## Supplementary material for "Microbial, dietary insect, and pathogen communities in fresh and decomposing guano of anthropic little brown bat (Myotis lucifugus) maternity colonies": Table S18

| Site | Month | Order | Mean Relative Abundance | Site / Month Total |
|---|---|---|---|---|
| AshInt | May | Araneae | 0.0106 | 1.0000 |
| AshInt | May | Blattodea | 0.0000 | 1.0000 |
| AshInt | May | Coleoptera | 0.0103 | 1.0000 |
| AshInt | May | Diptera | 0.6426 | 1.0000 |
| AshInt | May | Ephemeroptera | 0.0171 | 1.0000 |
| AshInt | May | Hemiptera | 0.0000 | 1.0000 |
| AshInt | May | Hymenoptera | 0.0042 | 1.0000 |
| AshInt | May | Lepidoptera | 0.0044 | 1.0000 |
| AshInt | May | Megaloptera | 0.0000 | 1.0000 |
| AshInt | May | Neuroptera | 0.0000 | 1.0000 |
| AshInt | May | Odonata | 0.0000 | 1.0000 |
| AshInt | May | Orthoptera | 0.0009 | 1.0000 |
| AshInt | May | Other | 0.0028 | 1.0000 |
| AshInt | May | Plecoptera | 0.0000 | 1.0000 |
| AshInt | May | Psocodea | 0.0000 | 1.0000 |
| AshInt | May | Sarcoptiformes | 0.0003 | 1.0000 |
| AshInt | May | Trichoptera | 0.0000 | 1.0000 |
| AshInt | May | Trombidiformes | 0.3067 | 1.0000 |
| AshInt | June | Araneae | 0.0726 | 1.0000 |
| AshInt | June | Blattodea | 0.0364 | 1.0000 |
| AshInt | June | Coleoptera | 0.0227 | 1.0000 |
| AshInt | June | Diptera | 0.4423 | 1.0000 |
| AshInt | June | Ephemeroptera | 0.0000 | 1.0000 |
| AshInt | June | Hemiptera | 0.0000 | 1.0000 |
| AshInt | June | Hymenoptera | 0.0125 | 1.0000 |
| AshInt | June | Lepidoptera | 0.0323 | 1.0000 |
| AshInt | June | Megaloptera | 0.0000 | 1.0000 |
| AshInt | June | Neuroptera | 0.0000 | 1.0000 |
| AshInt | June | Odonata | 0.0000 | 1.0000 |
| AshInt | June | Orthoptera | 0.0000 | 1.0000 |
| AshInt | June | Other | 0.0494 | 1.0000 |

| | | | | |
|---|---|---|---|---|
| AshInt | June | Plecoptera | 0.0000 | 1.0000 |
| AshInt | June | Psocodea | 0.0000 | 1.0000 |
| AshInt | June | Sarcoptiformes | 0.0000 | 1.0000 |
| AshInt | June | Trichoptera | 0.0061 | 1.0000 |
| AshInt | June | Trombidiformes | 0.3256 | 1.0000 |
| AshInt | July | Araneae | 0.0395 | 1.0000 |
| AshInt | July | Blattodea | 0.0000 | 1.0000 |
| AshInt | July | Coleoptera | 0.0591 | 1.0000 |
| AshInt | July | Diptera | 0.5669 | 1.0000 |
| AshInt | July | Ephemeroptera | 0.0000 | 1.0000 |
| AshInt | July | Hemiptera | 0.0000 | 1.0000 |
| AshInt | July | Hymenoptera | 0.0066 | 1.0000 |
| AshInt | July | Lepidoptera | 0.0188 | 1.0000 |
| AshInt | July | Megaloptera | 0.0000 | 1.0000 |
| AshInt | July | Neuroptera | 0.0000 | 1.0000 |
| AshInt | July | Odonata | 0.0000 | 1.0000 |
| AshInt | July | Orthoptera | 0.0000 | 1.0000 |
| AshInt | July | Other | 0.0260 | 1.0000 |
| AshInt | July | Plecoptera | 0.0000 | 1.0000 |
| AshInt | July | Psocodea | 0.0000 | 1.0000 |
| AshInt | July | Sarcoptiformes | 0.0000 | 1.0000 |
| AshInt | July | Trichoptera | 0.0000 | 1.0000 |
| AshInt | July | Trombidiformes | 0.2832 | 1.0000 |
| AshInt | August | Araneae | 0.0384 | 1.0000 |
| AshInt | August | Blattodea | 0.0000 | 1.0000 |
| AshInt | August | Coleoptera | 0.6199 | 1.0000 |
| AshInt | August | Diptera | 0.2330 | 1.0000 |
| AshInt | August | Ephemeroptera | 0.0022 | 1.0000 |
| AshInt | August | Hemiptera | 0.0000 | 1.0000 |
| AshInt | August | Hymenoptera | 0.0024 | 1.0000 |
| AshInt | August | Lepidoptera | 0.0005 | 1.0000 |

| | | | | |
|---|---|---|---|---|
| AshInt | August | Megaloptera | 0.0019 | 1.0000 |
| AshInt | August | Neuroptera | 0.0000 | 1.0000 |
| AshInt | August | Odonata | 0.0000 | 1.0000 |
| AshInt | August | Orthoptera | 0.0000 | 1.0000 |
| AshInt | August | Other | 0.0339 | 1.0000 |
| AshInt | August | Plecoptera | 0.0000 | 1.0000 |
| AshInt | August | Psocodea | 0.0000 | 1.0000 |
| AshInt | August | Sarcoptiformes | 0.0000 | 1.0000 |
| AshInt | August | Trichoptera | 0.0000 | 1.0000 |
| AshInt | August | Trombidiformes | 0.0679 | 1.0000 |
| AshExt | May | Araneae | 0.1096 | 1.0000 |
| AshExt | May | Blattodea | 0.0000 | 1.0000 |
| AshExt | May | Coleoptera | 0.1037 | 1.0000 |
| AshExt | May | Diptera | 0.4519 | 1.0000 |
| AshExt | May | Ephemeroptera | 0.0000 | 1.0000 |
| AshExt | May | Hemiptera | 0.0000 | 1.0000 |
| AshExt | May | Hymenoptera | 0.0000 | 1.0000 |
| AshExt | May | Lepidoptera | 0.1018 | 1.0000 |
| AshExt | May | Megaloptera | 0.0000 | 1.0000 |
| AshExt | May | Neuroptera | 0.0000 | 1.0000 |
| AshExt | May | Odonata | 0.0000 | 1.0000 |
| AshExt | May | Orthoptera | 0.0000 | 1.0000 |
| AshExt | May | Other | 0.0294 | 1.0000 |
| AshExt | May | Plecoptera | 0.0000 | 1.0000 |
| AshExt | May | Psocodea | 0.0000 | 1.0000 |
| AshExt | May | Sarcoptiformes | 0.0000 | 1.0000 |
| AshExt | May | Trichoptera | 0.0000 | 1.0000 |
| AshExt | May | Trombidiformes | 0.2036 | 1.0000 |
| AshExt | June | Araneae | 0.0845 | 1.0000 |
| AshExt | June | Blattodea | 0.0127 | 1.0000 |
| AshExt | June | Coleoptera | 0.0295 | 1.0000 |

| | | | | |
|---|---|---|---|---|
| AshExt | June | Diptera | 0.4184 | 1.0000 |
| AshExt | June | Ephemeroptera | 0.0133 | 1.0000 |
| AshExt | June | Hemiptera | 0.0000 | 1.0000 |
| AshExt | June | Hymenoptera | 0.0320 | 1.0000 |
| AshExt | June | Lepidoptera | 0.1515 | 1.0000 |
| AshExt | June | Megaloptera | 0.0000 | 1.0000 |
| AshExt | June | Neuroptera | 0.0000 | 1.0000 |
| AshExt | June | Odonata | 0.0000 | 1.0000 |
| AshExt | June | Orthoptera | 0.0000 | 1.0000 |
| AshExt | June | Other | 0.0024 | 1.0000 |
| AshExt | June | Plecoptera | 0.0000 | 1.0000 |
| AshExt | June | Psocodea | 0.0000 | 1.0000 |
| AshExt | June | Sarcoptiformes | 0.0000 | 1.0000 |
| AshExt | June | Trichoptera | 0.0034 | 1.0000 |
| AshExt | June | Trombidiformes | 0.2523 | 1.0000 |
| AshExt | July | Araneae | 0.2065 | 1.0000 |
| AshExt | July | Blattodea | 0.0000 | 1.0000 |
| AshExt | July | Coleoptera | 0.2222 | 1.0000 |
| AshExt | July | Diptera | 0.4601 | 1.0000 |
| AshExt | July | Ephemeroptera | 0.0000 | 1.0000 |
| AshExt | July | Hemiptera | 0.0000 | 1.0000 |
| AshExt | July | Hymenoptera | 0.0000 | 1.0000 |
| AshExt | July | Lepidoptera | 0.0000 | 1.0000 |
| AshExt | July | Megaloptera | 0.0000 | 1.0000 |
| AshExt | July | Neuroptera | 0.0000 | 1.0000 |
| AshExt | July | Odonata | 0.0000 | 1.0000 |
| AshExt | July | Orthoptera | 0.0000 | 1.0000 |
| AshExt | July | Other | 0.0000 | 1.0000 |
| AshExt | July | Plecoptera | 0.0000 | 1.0000 |
| AshExt | July | Psocodea | 0.0000 | 1.0000 |
| AshExt | July | Sarcoptiformes | 0.0000 | 1.0000 |

| | | | | |
|---|---|---|---|---|
| AshExt | July | Trichoptera | 0.0000 | 1.0000 |
| AshExt | July | Trombidiformes | 0.1111 | 1.0000 |
| AshExt | August | Araneae | 0.3991 | 1.0000 |
| AshExt | August | Blattodea | 0.0000 | 1.0000 |
| AshExt | August | Coleoptera | 0.0826 | 1.0000 |
| AshExt | August | Diptera | 0.3656 | 1.0000 |
| AshExt | August | Ephemeroptera | 0.0000 | 1.0000 |
| AshExt | August | Hemiptera | 0.0031 | 1.0000 |
| AshExt | August | Hymenoptera | 0.0007 | 1.0000 |
| AshExt | August | Lepidoptera | 0.0027 | 1.0000 |
| AshExt | August | Megaloptera | 0.0000 | 1.0000 |
| AshExt | August | Neuroptera | 0.0000 | 1.0000 |
| AshExt | August | Odonata | 0.0000 | 1.0000 |
| AshExt | August | Orthoptera | 0.0000 | 1.0000 |
| AshExt | August | Other | 0.0324 | 1.0000 |
| AshExt | August | Plecoptera | 0.0000 | 1.0000 |
| AshExt | August | Psocodea | 0.0000 | 1.0000 |
| AshExt | August | Sarcoptiformes | 0.0000 | 1.0000 |
| AshExt | August | Trichoptera | 0.0000 | 1.0000 |
| AshExt | August | Trombidiformes | 0.1139 | 1.0000 |
| WMCC | May | Araneae | 0.0472 | 1.0000 |
| WMCC | May | Blattodea | 0.0000 | 1.0000 |
| WMCC | May | Coleoptera | 0.0350 | 1.0000 |
| WMCC | May | Diptera | 0.7924 | 1.0000 |
| WMCC | May | Ephemeroptera | 0.0000 | 1.0000 |
| WMCC | May | Hemiptera | 0.0000 | 1.0000 |
| WMCC | May | Hymenoptera | 0.0094 | 1.0000 |
| WMCC | May | Lepidoptera | 0.0118 | 1.0000 |
| WMCC | May | Megaloptera | 0.0000 | 1.0000 |
| WMCC | May | Neuroptera | 0.0000 | 1.0000 |
| WMCC | May | Odonata | 0.0000 | 1.0000 |

| | | | | |
|---|---|---|---|---|
| WMCC | May | Orthoptera | 0.0000 | 1.0000 |
| WMCC | May | Other | 0.0326 | 1.0000 |
| WMCC | May | Plecoptera | 0.0021 | 1.0000 |
| WMCC | May | Psocodea | 0.0000 | 1.0000 |
| WMCC | May | Sarcoptiformes | 0.0000 | 1.0000 |
| WMCC | May | Trichoptera | 0.0000 | 1.0000 |
| WMCC | May | Trombidiformes | 0.0695 | 1.0000 |
| WMCC | June | Araneae | 0.1045 | 1.0000 |
| WMCC | June | Blattodea | 0.0000 | 1.0000 |
| WMCC | June | Coleoptera | 0.0831 | 1.0000 |
| WMCC | June | Diptera | 0.5435 | 1.0000 |
| WMCC | June | Ephemeroptera | 0.0112 | 1.0000 |
| WMCC | June | Hemiptera | 0.0000 | 1.0000 |
| WMCC | June | Hymenoptera | 0.0323 | 1.0000 |
| WMCC | June | Lepidoptera | 0.0514 | 1.0000 |
| WMCC | June | Megaloptera | 0.0000 | 1.0000 |
| WMCC | June | Neuroptera | 0.0000 | 1.0000 |
| WMCC | June | Odonata | 0.0000 | 1.0000 |
| WMCC | June | Orthoptera | 0.0000 | 1.0000 |
| WMCC | June | Other | 0.0424 | 1.0000 |
| WMCC | June | Plecoptera | 0.0000 | 1.0000 |
| WMCC | June | Psocodea | 0.0000 | 1.0000 |
| WMCC | June | Sarcoptiformes | 0.0000 | 1.0000 |
| WMCC | June | Trichoptera | 0.0955 | 1.0000 |
| WMCC | June | Trombidiformes | 0.0359 | 1.0000 |
| WMCC | July | Araneae | 0.0057 | 1.0000 |
| WMCC | July | Blattodea | 0.0000 | 1.0000 |
| WMCC | July | Coleoptera | 0.0025 | 1.0000 |
| WMCC | July | Diptera | 0.7505 | 1.0000 |
| WMCC | July | Ephemeroptera | 0.0016 | 1.0000 |
| WMCC | July | Hemiptera | 0.0002 | 1.0000 |

| | | | | |
|---|---|---|---|---|
| WMCC | July | Hymenoptera | 0.0022 | 1.0000 |
| WMCC | July | Lepidoptera | 0.0019 | 1.0000 |
| WMCC | July | Megaloptera | 0.0000 | 1.0000 |
| WMCC | July | Neuroptera | 0.0000 | 1.0000 |
| WMCC | July | Odonata | 0.0000 | 1.0000 |
| WMCC | July | Orthoptera | 0.0000 | 1.0000 |
| WMCC | July | Other | 0.0031 | 1.0000 |
| WMCC | July | Plecoptera | 0.0000 | 1.0000 |
| WMCC | July | Psocodea | 0.0144 | 1.0000 |
| WMCC | July | Sarcoptiformes | 0.0000 | 1.0000 |
| WMCC | July | Trichoptera | 0.2129 | 1.0000 |
| WMCC | July | Trombidiformes | 0.0050 | 1.0000 |
| WMCC | August | Araneae | 0.0000 | 1.0000 |
| WMCC | August | Blattodea | 0.0000 | 1.0000 |
| WMCC | August | Coleoptera | 0.0000 | 1.0000 |
| WMCC | August | Diptera | 0.4542 | 1.0000 |
| WMCC | August | Ephemeroptera | 0.0000 | 1.0000 |
| WMCC | August | Hemiptera | 0.0000 | 1.0000 |
| WMCC | August | Hymenoptera | 0.0002 | 1.0000 |
| WMCC | August | Lepidoptera | 0.0006 | 1.0000 |
| WMCC | August | Megaloptera | 0.0000 | 1.0000 |
| WMCC | August | Neuroptera | 0.0000 | 1.0000 |
| WMCC | August | Odonata | 0.0000 | 1.0000 |
| WMCC | August | Orthoptera | 0.0000 | 1.0000 |
| WMCC | August | Other | 0.0022 | 1.0000 |
| WMCC | August | Plecoptera | 0.0000 | 1.0000 |
| WMCC | August | Psocodea | 0.0000 | 1.0000 |
| WMCC | August | Sarcoptiformes | 0.0000 | 1.0000 |
| WMCC | August | Trichoptera | 0.0000 | 1.0000 |
| WMCC | August | Trombidiformes | 0.5427 | 1.0000 |
