## Supplementary material for "Microbial, dietary insect, and pathogen communities in fresh and decomposing guano of anthropic little brown bat (Myotis lucifugus) maternity colonies": Table S19

| Site | Month | Order | Family | Mean Relative Abundance | Site / Month Total |
|---|---|---|---|---|---|
| AshInt | May | Araneae | Agelenidae | 0.0000 | 1.0000 |
| AshInt | May | Araneae | Amaurobiidae | 0.0000 | 1.0000 |
| AshInt | May | Araneae | Anyphaenidae | 0.0049 | 1.0000 |
| AshInt | May | Araneae | Cicurinidae | 0.0001 | 1.0000 |
| AshInt | May | Araneae | Dictynidae | 0.0000 | 1.0000 |
| AshInt | May | Araneae | Gnaphosidae | 0.0000 | 1.0000 |
| AshInt | May | Araneae | Linyphiidae | 0.0000 | 1.0000 |
| AshInt | May | Araneae | Lycosidae | 0.0002 | 1.0000 |
| AshInt | May | Araneae | Phrurolithidae | 0.0000 | 1.0000 |
| AshInt | May | Araneae | Salticidae | 0.0001 | 1.0000 |
| AshInt | May | Araneae | Theridiidae | 0.0015 | 1.0000 |
| AshInt | May | Araneae | Thomisidae | 0.0037 | 1.0000 |
| AshInt | May | Blattodea | Ectobiidae | 0.0000 | 1.0000 |
| AshInt | May | Coleoptera | Buprestidae | 0.0000 | 1.0000 |
| AshInt | May | Coleoptera | Carabidae | 0.0097 | 1.0000 |
| AshInt | May | Coleoptera | Curculionidae | 0.0005 | 1.0000 |
| AshInt | May | Coleoptera | Dytiscidae | 0.0000 | 1.0000 |
| AshInt | May | Coleoptera | Lampyridae | 0.0000 | 1.0000 |
| AshInt | May | Coleoptera | Scarabaeidae | 0.0001 | 1.0000 |
| AshInt | May | Coleoptera | Scirtidae | 0.0000 | 1.0000 |
| AshInt | May | Diptera | Agromyzidae | 0.0029 | 1.0000 |
| AshInt | May | Diptera | Anisopodidae | 0.2052 | 1.0000 |
| AshInt | May | Diptera | Calliphoridae | 0.0005 | 1.0000 |
| AshInt | May | Diptera | Cecidomyiidae | 0.0125 | 1.0000 |
| AshInt | May | Diptera | Chaoboridae | 0.1368 | 1.0000 |
| AshInt | May | Diptera | Chironomidae | 0.2311 | 1.0000 |
| AshInt | May | Diptera | Chloropidae | 0.0026 | 1.0000 |
| AshInt | May | Diptera | Culicidae | 0.0000 | 1.0000 |
| AshInt | May | Diptera | Drosophilidae | 0.0001 | 1.0000 |
| AshInt | May | Diptera | Empididae | 0.0019 | 1.0000 |
| AshInt | May | Diptera | Hybotidae | 0.0057 | 1.0000 |

| | | | | | |
|---|---|---|---|---|---|
| AshInt | May | Diptera | Keroplatidae | 0.0000 | 1.0000 |
| AshInt | May | Diptera | Limoniidae | 0.0078 | 1.0000 |
| AshInt | May | Diptera | Muscidae | 0.0000 | 1.0000 |
| AshInt | May | Diptera | Mycetophilidae | 0.0000 | 1.0000 |
| AshInt | May | Diptera | Opomyzidae | 0.0000 | 1.0000 |
| AshInt | May | Diptera | Pallopteridae | 0.0000 | 1.0000 |
| AshInt | May | Diptera | Phoridae | 0.0011 | 1.0000 |
| AshInt | May | Diptera | Polleniidae | 0.0003 | 1.0000 |
| AshInt | May | Diptera | Psychodidae | 0.0000 | 1.0000 |
| AshInt | May | Diptera | Sarcophagidae | 0.0000 | 1.0000 |
| AshInt | May | Diptera | Sciaridae | 0.0048 | 1.0000 |
| AshInt | May | Diptera | Sphaeroceridae | 0.0001 | 1.0000 |
| AshInt | May | Diptera | Syrphidae | 0.0000 | 1.0000 |
| AshInt | May | Diptera | Tabanidae | 0.0000 | 1.0000 |
| AshInt | May | Diptera | Tachinidae | 0.0249 | 1.0000 |
| AshInt | May | Diptera | Tipulidae | 0.0043 | 1.0000 |
| AshInt | May | Ephemeroptera | Caenidae | 0.0000 | 1.0000 |
| AshInt | May | Ephemeroptera | Ephemeridae | 0.0000 | 1.0000 |
| AshInt | May | Ephemeroptera | Heptageniidae | 0.0171 | 1.0000 |
| AshInt | May | Hemiptera | Clastopteridae | 0.0000 | 1.0000 |
| AshInt | May | Hemiptera | Lygaeidae | 0.0000 | 1.0000 |
| AshInt | May | Hymenoptera | Apidae | 0.0000 | 1.0000 |
| AshInt | May | Hymenoptera | Braconidae | 0.0023 | 1.0000 |
| AshInt | May | Hymenoptera | Ichneumonidae | 0.0000 | 1.0000 |
| AshInt | May | Hymenoptera | Other | 0.0014 | 1.0000 |
| AshInt | May | Hymenoptera | Tenthredinidae | 0.0003 | 1.0000 |
| AshInt | May | Hymenoptera | Trigonalidae | 0.0001 | 1.0000 |
| AshInt | May | Lepidoptera | Crambidae | 0.0000 | 1.0000 |
| AshInt | May | Lepidoptera | Elachistidae | 0.0000 | 1.0000 |
| AshInt | May | Lepidoptera | Erebidae | 0.0000 | 1.0000 |
| AshInt | May | Lepidoptera | Gelechiidae | 0.0000 | 1.0000 |

| | | | | | |
|---|---|---|---|---|---|
| AshInt | May | Lepidoptera | Noctuidae | 0.0024 | 1.0000 |
| AshInt | May | Lepidoptera | Notodontidae | 0.0000 | 1.0000 |
| AshInt | May | Lepidoptera | Nymphalidae | 0.0000 | 1.0000 |
| AshInt | May | Lepidoptera | Pyralidae | 0.0012 | 1.0000 |
| AshInt | May | Lepidoptera | Tischeriidae | 0.0000 | 1.0000 |
| AshInt | May | Lepidoptera | Tortricidae | 0.0008 | 1.0000 |
| AshInt | May | Megaloptera | Corydalidae | 0.0000 | 1.0000 |
| AshInt | May | Neuroptera | Hemerobiidae | 0.0000 | 1.0000 |
| AshInt | May | Odonata | Libellulidae | 0.0000 | 1.0000 |
| AshInt | May | Orthoptera | Tettigoniidae | 0.0009 | 1.0000 |
| AshInt | May | Other | Other | 0.0028 | 1.0000 |
| AshInt | May | Plecoptera | Chloroperlidae | 0.0000 | 1.0000 |
| AshInt | May | Psocodea | Peripsocidae | 0.0000 | 1.0000 |
| AshInt | May | Sarcoptiformes | Oripodidae | 0.0003 | 1.0000 |
| AshInt | May | Trichoptera | Dipseudopsidae | 0.0000 | 1.0000 |
| AshInt | May | Trichoptera | Hydroptilidae | 0.0000 | 1.0000 |
| AshInt | May | Trichoptera | Leptoceridae | 0.0000 | 1.0000 |
| AshInt | May | Trombidiformes | Arrenuridae | 0.0000 | 1.0000 |
| AshInt | May | Trombidiformes | Aturidae | 0.0000 | 1.0000 |
| AshInt | May | Trombidiformes | Axonopsidae | 0.0000 | 1.0000 |
| AshInt | May | Trombidiformes | Eupodidae | 0.0011 | 1.0000 |
| AshInt | May | Trombidiformes | Feltriidae | 0.0001 | 1.0000 |
| AshInt | May | Trombidiformes | Hydryphantidae | 0.0020 | 1.0000 |
| AshInt | May | Trombidiformes | Lebertiidae | 0.0472 | 1.0000 |
| AshInt | May | Trombidiformes | Mideopsidae | 0.0000 | 1.0000 |
| AshInt | May | Trombidiformes | Pionidae | 0.2460 | 1.0000 |
| AshInt | May | Trombidiformes | Sperchontidae | 0.0001 | 1.0000 |
| AshInt | May | Trombidiformes | Tarsonemidae | 0.0000 | 1.0000 |
| AshInt | May | Trombidiformes | Torrenticolidae | 0.0100 | 1.0000 |
| AshInt | May | Trombidiformes | Unionicolidae | 0.0000 | 1.0000 |
| AshInt | June | Araneae | Agelenidae | 0.0000 | 1.0000 |

| | | | | | |
|---|---|---|---|---|---|
| AshInt | June | Araneae | Amaurobiidae | 0.0000 | 1.0000 |
| AshInt | June | Araneae | Anyphaenidae | 0.0565 | 1.0000 |
| AshInt | June | Araneae | Cicurinidae | 0.0000 | 1.0000 |
| AshInt | June | Araneae | Dictynidae | 0.0000 | 1.0000 |
| AshInt | June | Araneae | Gnaphosidae | 0.0000 | 1.0000 |
| AshInt | June | Araneae | Linyphiidae | 0.0000 | 1.0000 |
| AshInt | June | Araneae | Lycosidae | 0.0000 | 1.0000 |
| AshInt | June | Araneae | Phrurolithidae | 0.0000 | 1.0000 |
| AshInt | June | Araneae | Salticidae | 0.0000 | 1.0000 |
| AshInt | June | Araneae | Theridiidae | 0.0000 | 1.0000 |
| AshInt | June | Araneae | Thomisidae | 0.0161 | 1.0000 |
| AshInt | June | Blattodea | Ectobiidae | 0.0364 | 1.0000 |
| AshInt | June | Coleoptera | Buprestidae | 0.0000 | 1.0000 |
| AshInt | June | Coleoptera | Carabidae | 0.0226 | 1.0000 |
| AshInt | June | Coleoptera | Curculionidae | 0.0000 | 1.0000 |
| AshInt | June | Coleoptera | Dytiscidae | 0.0000 | 1.0000 |
| AshInt | June | Coleoptera | Lampyridae | 0.0000 | 1.0000 |
| AshInt | June | Coleoptera | Scarabaeidae | 0.0000 | 1.0000 |
| AshInt | June | Coleoptera | Scirtidae | 0.0001 | 1.0000 |
| AshInt | June | Diptera | Agromyzidae | 0.0000 | 1.0000 |
| AshInt | June | Diptera | Anisopodidae | 0.1461 | 1.0000 |
| AshInt | June | Diptera | Calliphoridae | 0.0000 | 1.0000 |
| AshInt | June | Diptera | Cecidomyiidae | 0.1136 | 1.0000 |
| AshInt | June | Diptera | Chaoboridae | 0.0584 | 1.0000 |
| AshInt | June | Diptera | Chironomidae | 0.0981 | 1.0000 |
| AshInt | June | Diptera | Chloropidae | 0.0028 | 1.0000 |
| AshInt | June | Diptera | Culicidae | 0.0000 | 1.0000 |
| AshInt | June | Diptera | Drosophilidae | 0.0000 | 1.0000 |
| AshInt | June | Diptera | Empididae | 0.0000 | 1.0000 |
| AshInt | June | Diptera | Hybotidae | 0.0000 | 1.0000 |
| AshInt | June | Diptera | Keroplatidae | 0.0026 | 1.0000 |

| | | | | | |
|---|---|---|---|---|---|
| AshInt | June | Diptera | Limoniidae | 0.0013 | 1.0000 |
| AshInt | June | Diptera | Muscidae | 0.0000 | 1.0000 |
| AshInt | June | Diptera | Mycetophilidae | 0.0000 | 1.0000 |
| AshInt | June | Diptera | Opomyzidae | 0.0000 | 1.0000 |
| AshInt | June | Diptera | Pallopteridae | 0.0000 | 1.0000 |
| AshInt | June | Diptera | Phoridae | 0.0000 | 1.0000 |
| AshInt | June | Diptera | Polleniidae | 0.0000 | 1.0000 |
| AshInt | June | Diptera | Psychodidae | 0.0000 | 1.0000 |
| AshInt | June | Diptera | Sarcophagidae | 0.0000 | 1.0000 |
| AshInt | June | Diptera | Sciaridae | 0.0169 | 1.0000 |
| AshInt | June | Diptera | Sphaeroceridae | 0.0000 | 1.0000 |
| AshInt | June | Diptera | Syrphidae | 0.0013 | 1.0000 |
| AshInt | June | Diptera | Tabanidae | 0.0000 | 1.0000 |
| AshInt | June | Diptera | Tachinidae | 0.0013 | 1.0000 |
| AshInt | June | Diptera | Tipulidae | 0.0000 | 1.0000 |
| AshInt | June | Ephemeroptera | Caenidae | 0.0000 | 1.0000 |
| AshInt | June | Ephemeroptera | Ephemeridae | 0.0000 | 1.0000 |
| AshInt | June | Ephemeroptera | Heptageniidae | 0.0000 | 1.0000 |
| AshInt | June | Hemiptera | Clastopteridae | 0.0000 | 1.0000 |
| AshInt | June | Hemiptera | Lygaeidae | 0.0000 | 1.0000 |
| AshInt | June | Hymenoptera | Apidae | 0.0105 | 1.0000 |
| AshInt | June | Hymenoptera | Braconidae | 0.0000 | 1.0000 |
| AshInt | June | Hymenoptera | Ichneumonidae | 0.0020 | 1.0000 |
| AshInt | June | Hymenoptera | Other | 0.0000 | 1.0000 |
| AshInt | June | Hymenoptera | Tenthredinidae | 0.0000 | 1.0000 |
| AshInt | June | Hymenoptera | Trigonalidae | 0.0000 | 1.0000 |
| AshInt | June | Lepidoptera | Crambidae | 0.0000 | 1.0000 |
| AshInt | June | Lepidoptera | Elachistidae | 0.0000 | 1.0000 |
| AshInt | June | Lepidoptera | Erebidae | 0.0000 | 1.0000 |
| AshInt | June | Lepidoptera | Gelechiidae | 0.0000 | 1.0000 |
| AshInt | June | Lepidoptera | Noctuidae | 0.0013 | 1.0000 |

| | | | | | |
|---|---|---|---|---|---|
| AshInt | June | Lepidoptera | Notodontidae | 0.0269 | 1.0000 |
| AshInt | June | Lepidoptera | Nymphalidae | 0.0000 | 1.0000 |
| AshInt | June | Lepidoptera | Pyralidae | 0.0000 | 1.0000 |
| AshInt | June | Lepidoptera | Tischeriidae | 0.0000 | 1.0000 |
| AshInt | June | Lepidoptera | Tortricidae | 0.0041 | 1.0000 |
| AshInt | June | Megaloptera | Corydalidae | 0.0000 | 1.0000 |
| AshInt | June | Neuroptera | Hemerobiidae | 0.0000 | 1.0000 |
| AshInt | June | Odonata | Libellulidae | 0.0000 | 1.0000 |
| AshInt | June | Orthoptera | Tettigoniidae | 0.0000 | 1.0000 |
| AshInt | June | Other | Other | 0.0494 | 1.0000 |
| AshInt | June | Plecoptera | Chloroperlidae | 0.0000 | 1.0000 |
| AshInt | June | Psocodea | Peripsocidae | 0.0000 | 1.0000 |
| AshInt | June | Sarcoptiformes | Oripodidae | 0.0000 | 1.0000 |
| AshInt | June | Trichoptera | Dipseudopsidae | 0.0061 | 1.0000 |
| AshInt | June | Trichoptera | Hydroptilidae | 0.0000 | 1.0000 |
| AshInt | June | Trichoptera | Leptoceridae | 0.0000 | 1.0000 |
| AshInt | June | Trombidiformes | Arrenuridae | 0.1364 | 1.0000 |
| AshInt | June | Trombidiformes | Aturidae | 0.0000 | 1.0000 |
| AshInt | June | Trombidiformes | Axonopsidae | 0.0187 | 1.0000 |
| AshInt | June | Trombidiformes | Eupodidae | 0.0000 | 1.0000 |
| AshInt | June | Trombidiformes | Feltriidae | 0.0000 | 1.0000 |
| AshInt | June | Trombidiformes | Hydryphantidae | 0.0000 | 1.0000 |
| AshInt | June | Trombidiformes | Lebertiidae | 0.0770 | 1.0000 |
| AshInt | June | Trombidiformes | Mideopsidae | 0.0000 | 1.0000 |
| AshInt | June | Trombidiformes | Pionidae | 0.0935 | 1.0000 |
| AshInt | June | Trombidiformes | Sperchontidae | 0.0000 | 1.0000 |
| AshInt | June | Trombidiformes | Tarsonemidae | 0.0000 | 1.0000 |
| AshInt | June | Trombidiformes | Torrenticolidae | 0.0000 | 1.0000 |
| AshInt | June | Trombidiformes | Unionicolidae | 0.0000 | 1.0000 |
| AshInt | July | Araneae | Agelenidae | 0.0000 | 1.0000 |
| AshInt | July | Araneae | Amaurobiidae | 0.0000 | 1.0000 |

| | | | | | |
|---|---|---|---|---|---|
| AshInt | July | Araneae | Anyphaenidae | 0.0162 | 1.0000 |
| AshInt | July | Araneae | Cicurinidae | 0.0000 | 1.0000 |
| AshInt | July | Araneae | Dictynidae | 0.0000 | 1.0000 |
| AshInt | July | Araneae | Gnaphosidae | 0.0000 | 1.0000 |
| AshInt | July | Araneae | Linyphiidae | 0.0000 | 1.0000 |
| AshInt | July | Araneae | Lycosidae | 0.0000 | 1.0000 |
| AshInt | July | Araneae | Phrurolithidae | 0.0062 | 1.0000 |
| AshInt | July | Araneae | Salticidae | 0.0000 | 1.0000 |
| AshInt | July | Araneae | Theridiidae | 0.0000 | 1.0000 |
| AshInt | July | Araneae | Thomisidae | 0.0171 | 1.0000 |
| AshInt | July | Blattodea | Ectobiidae | 0.0000 | 1.0000 |
| AshInt | July | Coleoptera | Buprestidae | 0.0000 | 1.0000 |
| AshInt | July | Coleoptera | Carabidae | 0.0089 | 1.0000 |
| AshInt | July | Coleoptera | Curculionidae | 0.0131 | 1.0000 |
| AshInt | July | Coleoptera | Dytiscidae | 0.0000 | 1.0000 |
| AshInt | July | Coleoptera | Lampyridae | 0.0038 | 1.0000 |
| AshInt | July | Coleoptera | Scarabaeidae | 0.0334 | 1.0000 |
| AshInt | July | Coleoptera | Scirtidae | 0.0000 | 1.0000 |
| AshInt | July | Diptera | Agromyzidae | 0.0050 | 1.0000 |
| AshInt | July | Diptera | Anisopodidae | 0.0401 | 1.0000 |
| AshInt | July | Diptera | Calliphoridae | 0.0000 | 1.0000 |
| AshInt | July | Diptera | Cecidomyiidae | 0.1046 | 1.0000 |
| AshInt | July | Diptera | Chaoboridae | 0.0740 | 1.0000 |
| AshInt | July | Diptera | Chironomidae | 0.0605 | 1.0000 |
| AshInt | July | Diptera | Chloropidae | 0.0052 | 1.0000 |
| AshInt | July | Diptera | Culicidae | 0.0125 | 1.0000 |
| AshInt | July | Diptera | Drosophilidae | 0.0000 | 1.0000 |
| AshInt | July | Diptera | Empididae | 0.0000 | 1.0000 |
| AshInt | July | Diptera | Hybotidae | 0.0000 | 1.0000 |
| AshInt | July | Diptera | Keroplatidae | 0.0000 | 1.0000 |
| AshInt | July | Diptera | Limoniidae | 0.2330 | 1.0000 |

| | | | | | |
|---|---|---|---|---|---|
| AshInt | July | Diptera | Muscidae | 0.0000 | 1.0000 |
| AshInt | July | Diptera | Mycetophilidae | 0.0000 | 1.0000 |
| AshInt | July | Diptera | Opomyzidae | 0.0000 | 1.0000 |
| AshInt | July | Diptera | Pallopteridae | 0.0000 | 1.0000 |
| AshInt | July | Diptera | Phoridae | 0.0056 | 1.0000 |
| AshInt | July | Diptera | Polleniidae | 0.0000 | 1.0000 |
| AshInt | July | Diptera | Psychodidae | 0.0000 | 1.0000 |
| AshInt | July | Diptera | Sarcophagidae | 0.0000 | 1.0000 |
| AshInt | July | Diptera | Sciaridae | 0.0026 | 1.0000 |
| AshInt | July | Diptera | Sphaeroceridae | 0.0000 | 1.0000 |
| AshInt | July | Diptera | Syrphidae | 0.0000 | 1.0000 |
| AshInt | July | Diptera | Tabanidae | 0.0236 | 1.0000 |
| AshInt | July | Diptera | Tachinidae | 0.0000 | 1.0000 |
| AshInt | July | Diptera | Tipulidae | 0.0000 | 1.0000 |
| AshInt | July | Ephemeroptera | Caenidae | 0.0000 | 1.0000 |
| AshInt | July | Ephemeroptera | Ephemeridae | 0.0000 | 1.0000 |
| AshInt | July | Ephemeroptera | Heptageniidae | 0.0000 | 1.0000 |
| AshInt | July | Hemiptera | Clastopteridae | 0.0000 | 1.0000 |
| AshInt | July | Hemiptera | Lygaeidae | 0.0000 | 1.0000 |
| AshInt | July | Hymenoptera | Apidae | 0.0000 | 1.0000 |
| AshInt | July | Hymenoptera | Braconidae | 0.0013 | 1.0000 |
| AshInt | July | Hymenoptera | Ichneumonidae | 0.0003 | 1.0000 |
| AshInt | July | Hymenoptera | Other | 0.0000 | 1.0000 |
| AshInt | July | Hymenoptera | Tenthredinidae | 0.0050 | 1.0000 |
| AshInt | July | Hymenoptera | Trigonalidae | 0.0000 | 1.0000 |
| AshInt | July | Lepidoptera | Crambidae | 0.0000 | 1.0000 |
| AshInt | July | Lepidoptera | Elachistidae | 0.0000 | 1.0000 |
| AshInt | July | Lepidoptera | Erebidae | 0.0095 | 1.0000 |
| AshInt | July | Lepidoptera | Gelechiidae | 0.0000 | 1.0000 |
| AshInt | July | Lepidoptera | Noctuidae | 0.0000 | 1.0000 |
| AshInt | July | Lepidoptera | Notodontidae | 0.0000 | 1.0000 |

| | | | | | |
|---|---|---|---|---|---|
| AshInt | July | Lepidoptera | Nymphalidae | 0.0000 | 1.0000 |
| AshInt | July | Lepidoptera | Pyralidae | 0.0000 | 1.0000 |
| AshInt | July | Lepidoptera | Tischeriidae | 0.0063 | 1.0000 |
| AshInt | July | Lepidoptera | Tortricidae | 0.0031 | 1.0000 |
| AshInt | July | Megaloptera | Corydalidae | 0.0000 | 1.0000 |
| AshInt | July | Neuroptera | Hemerobiidae | 0.0000 | 1.0000 |
| AshInt | July | Odonata | Libellulidae | 0.0000 | 1.0000 |
| AshInt | July | Orthoptera | Tettigoniidae | 0.0000 | 1.0000 |
| AshInt | July | Other | Other | 0.0260 | 1.0000 |
| AshInt | July | Plecoptera | Chloroperlidae | 0.0000 | 1.0000 |
| AshInt | July | Psocodea | Peripsocidae | 0.0000 | 1.0000 |
| AshInt | July | Sarcoptiformes | Oripodidae | 0.0000 | 1.0000 |
| AshInt | July | Trichoptera | Dipseudopsidae | 0.0000 | 1.0000 |
| AshInt | July | Trichoptera | Hydroptilidae | 0.0000 | 1.0000 |
| AshInt | July | Trichoptera | Leptoceridae | 0.0000 | 1.0000 |
| AshInt | July | Trombidiformes | Arrenuridae | 0.1182 | 1.0000 |
| AshInt | July | Trombidiformes | Aturidae | 0.0000 | 1.0000 |
| AshInt | July | Trombidiformes | Axonopsidae | 0.0000 | 1.0000 |
| AshInt | July | Trombidiformes | Eupodidae | 0.0000 | 1.0000 |
| AshInt | July | Trombidiformes | Feltriidae | 0.0000 | 1.0000 |
| AshInt | July | Trombidiformes | Hydryphantidae | 0.0000 | 1.0000 |
| AshInt | July | Trombidiformes | Lebertiidae | 0.0115 | 1.0000 |
| AshInt | July | Trombidiformes | Mideopsidae | 0.0026 | 1.0000 |
| AshInt | July | Trombidiformes | Pionidae | 0.0125 | 1.0000 |
| AshInt | July | Trombidiformes | Sperchontidae | 0.0000 | 1.0000 |
| AshInt | July | Trombidiformes | Tarsonemidae | 0.0000 | 1.0000 |
| AshInt | July | Trombidiformes | Torrenticolidae | 0.0946 | 1.0000 |
| AshInt | July | Trombidiformes | Unionicolidae | 0.0438 | 1.0000 |
| AshInt | August | Araneae | Agelenidae | 0.0000 | 1.0000 |
| AshInt | August | Araneae | Amaurobiidae | 0.0001 | 1.0000 |
| AshInt | August | Araneae | Anyphaenidae | 0.0338 | 1.0000 |

| | | | | | |
|---|---|---|---|---|---|
| AshInt | August | Araneae | Cicurinidae | 0.0000 | 1.0000 |
| AshInt | August | Araneae | Dictynidae | 0.0000 | 1.0000 |
| AshInt | August | Araneae | Gnaphosidae | 0.0005 | 1.0000 |
| AshInt | August | Araneae | Linyphiidae | 0.0001 | 1.0000 |
| AshInt | August | Araneae | Lycosidae | 0.0001 | 1.0000 |
| AshInt | August | Araneae | Phrurolithidae | 0.0017 | 1.0000 |
| AshInt | August | Araneae | Salticidae | 0.0001 | 1.0000 |
| AshInt | August | Araneae | Theridiidae | 0.0019 | 1.0000 |
| AshInt | August | Araneae | Thomisidae | 0.0002 | 1.0000 |
| AshInt | August | Blattodea | Ectobiidae | 0.0000 | 1.0000 |
| AshInt | August | Coleoptera | Buprestidae | 0.0000 | 1.0000 |
| AshInt | August | Coleoptera | Carabidae | 0.0184 | 1.0000 |
| AshInt | August | Coleoptera | Curculionidae | 0.5692 | 1.0000 |
| AshInt | August | Coleoptera | Dytiscidae | 0.0002 | 1.0000 |
| AshInt | August | Coleoptera | Lampyridae | 0.0000 | 1.0000 |
| AshInt | August | Coleoptera | Scarabaeidae | 0.0320 | 1.0000 |
| AshInt | August | Coleoptera | Scirtidae | 0.0000 | 1.0000 |
| AshInt | August | Diptera | Agromyzidae | 0.0000 | 1.0000 |
| AshInt | August | Diptera | Anisopodidae | 0.0071 | 1.0000 |
| AshInt | August | Diptera | Calliphoridae | 0.0001 | 1.0000 |
| AshInt | August | Diptera | Cecidomyiidae | 0.1495 | 1.0000 |
| AshInt | August | Diptera | Chaoboridae | 0.0657 | 1.0000 |
| AshInt | August | Diptera | Chironomidae | 0.0027 | 1.0000 |
| AshInt | August | Diptera | Chloropidae | 0.0004 | 1.0000 |
| AshInt | August | Diptera | Culicidae | 0.0012 | 1.0000 |
| AshInt | August | Diptera | Drosophilidae | 0.0008 | 1.0000 |
| AshInt | August | Diptera | Empididae | 0.0000 | 1.0000 |
| AshInt | August | Diptera | Hybotidae | 0.0000 | 1.0000 |
| AshInt | August | Diptera | Keroplatidae | 0.0000 | 1.0000 |
| AshInt | August | Diptera | Limoniidae | 0.0032 | 1.0000 |
| AshInt | August | Diptera | Muscidae | 0.0000 | 1.0000 |

| | | | | | |
|---|---|---|---|---|---|
| AshInt | August | Diptera | Mycetophilidae | 0.0002 | 1.0000 |
| AshInt | August | Diptera | Opomyzidae | 0.0001 | 1.0000 |
| AshInt | August | Diptera | Pallopteridae | 0.0000 | 1.0000 |
| AshInt | August | Diptera | Phoridae | 0.0000 | 1.0000 |
| AshInt | August | Diptera | Polleniidae | 0.0000 | 1.0000 |
| AshInt | August | Diptera | Psychodidae | 0.0006 | 1.0000 |
| AshInt | August | Diptera | Sarcophagidae | 0.0000 | 1.0000 |
| AshInt | August | Diptera | Sciaridae | 0.0010 | 1.0000 |
| AshInt | August | Diptera | Sphaeroceridae | 0.0000 | 1.0000 |
| AshInt | August | Diptera | Syrphidae | 0.0001 | 1.0000 |
| AshInt | August | Diptera | Tabanidae | 0.0000 | 1.0000 |
| AshInt | August | Diptera | Tachinidae | 0.0001 | 1.0000 |
| AshInt | August | Diptera | Tipulidae | 0.0002 | 1.0000 |
| AshInt | August | Ephemeroptera | Caenidae | 0.0000 | 1.0000 |
| AshInt | August | Ephemeroptera | Ephemeridae | 0.0022 | 1.0000 |
| AshInt | August | Ephemeroptera | Heptageniidae | 0.0000 | 1.0000 |
| AshInt | August | Hemiptera | Clastopteridae | 0.0000 | 1.0000 |
| AshInt | August | Hemiptera | Lygaeidae | 0.0000 | 1.0000 |
| AshInt | August | Hymenoptera | Apidae | 0.0000 | 1.0000 |
| AshInt | August | Hymenoptera | Braconidae | 0.0000 | 1.0000 |
| AshInt | August | Hymenoptera | Ichneumonidae | 0.0017 | 1.0000 |
| AshInt | August | Hymenoptera | Other | 0.0000 | 1.0000 |
| AshInt | August | Hymenoptera | Tenthredinidae | 0.0007 | 1.0000 |
| AshInt | August | Hymenoptera | Trigonalidae | 0.0000 | 1.0000 |
| AshInt | August | Lepidoptera | Crambidae | 0.0000 | 1.0000 |
| AshInt | August | Lepidoptera | Elachistidae | 0.0000 | 1.0000 |
| AshInt | August | Lepidoptera | Erebidae | 0.0000 | 1.0000 |
| AshInt | August | Lepidoptera | Gelechiidae | 0.0000 | 1.0000 |
| AshInt | August | Lepidoptera | Noctuidae | 0.0000 | 1.0000 |
| AshInt | August | Lepidoptera | Notodontidae | 0.0000 | 1.0000 |
| AshInt | August | Lepidoptera | Nymphalidae | 0.0000 | 1.0000 |

| | | | | | |
|---|---|---|---|---|---|
| AshInt | August | Lepidoptera | Pyralidae | 0.0001 | 1.0000 |
| AshInt | August | Lepidoptera | Tischeriidae | 0.0000 | 1.0000 |
| AshInt | August | Lepidoptera | Tortricidae | 0.0003 | 1.0000 |
| AshInt | August | Megaloptera | Corydalidae | 0.0019 | 1.0000 |
| AshInt | August | Neuroptera | Hemerobiidae | 0.0000 | 1.0000 |
| AshInt | August | Odonata | Libellulidae | 0.0000 | 1.0000 |
| AshInt | August | Orthoptera | Tettigoniidae | 0.0000 | 1.0000 |
| AshInt | August | Other | Other | 0.0339 | 1.0000 |
| AshInt | August | Plecoptera | Chloroperlidae | 0.0000 | 1.0000 |
| AshInt | August | Psocodea | Peripsocidae | 0.0000 | 1.0000 |
| AshInt | August | Sarcoptiformes | Oripodidae | 0.0000 | 1.0000 |
| AshInt | August | Trichoptera | Dipseudopsidae | 0.0000 | 1.0000 |
| AshInt | August | Trichoptera | Hydroptilidae | 0.0000 | 1.0000 |
| AshInt | August | Trichoptera | Leptoceridae | 0.0000 | 1.0000 |
| AshInt | August | Trombidiformes | Arrenuridae | 0.0000 | 1.0000 |
| AshInt | August | Trombidiformes | Aturidae | 0.0000 | 1.0000 |
| AshInt | August | Trombidiformes | Axonopsidae | 0.0001 | 1.0000 |
| AshInt | August | Trombidiformes | Eupodidae | 0.0000 | 1.0000 |
| AshInt | August | Trombidiformes | Feltriidae | 0.0000 | 1.0000 |
| AshInt | August | Trombidiformes | Hydryphantidae | 0.0000 | 1.0000 |
| AshInt | August | Trombidiformes | Lebertiidae | 0.0649 | 1.0000 |
| AshInt | August | Trombidiformes | Mideopsidae | 0.0001 | 1.0000 |
| AshInt | August | Trombidiformes | Pionidae | 0.0006 | 1.0000 |
| AshInt | August | Trombidiformes | Sperchontidae | 0.0000 | 1.0000 |
| AshInt | August | Trombidiformes | Tarsonemidae | 0.0008 | 1.0000 |
| AshInt | August | Trombidiformes | Torrenticolidae | 0.0001 | 1.0000 |
| AshInt | August | Trombidiformes | Unionicolidae | 0.0013 | 1.0000 |
| AshExt | May | Araneae | Agelenidae | 0.0000 | 1.0000 |
| AshExt | May | Araneae | Amaurobiidae | 0.0000 | 1.0000 |
| AshExt | May | Araneae | Anyphaenidae | 0.0769 | 1.0000 |
| AshExt | May | Araneae | Cicurinidae | 0.0000 | 1.0000 |

| | | | | | |
|---|---|---|---|---|---|
| AshExt | May | Araneae | Dictynidae | 0.0000 | 1.0000 |
| AshExt | May | Araneae | Gnaphosidae | 0.0000 | 1.0000 |
| AshExt | May | Araneae | Linyphiidae | 0.0000 | 1.0000 |
| AshExt | May | Araneae | Lycosidae | 0.0000 | 1.0000 |
| AshExt | May | Araneae | Phrurolithidae | 0.0000 | 1.0000 |
| AshExt | May | Araneae | Salticidae | 0.0000 | 1.0000 |
| AshExt | May | Araneae | Theridiidae | 0.0327 | 1.0000 |
| AshExt | May | Araneae | Thomisidae | 0.0000 | 1.0000 |
| AshExt | May | Blattodea | Ectobiidae | 0.0000 | 1.0000 |
| AshExt | May | Coleoptera | Buprestidae | 0.0000 | 1.0000 |
| AshExt | May | Coleoptera | Carabidae | 0.0383 | 1.0000 |
| AshExt | May | Coleoptera | Curculionidae | 0.0654 | 1.0000 |
| AshExt | May | Coleoptera | Dytiscidae | 0.0000 | 1.0000 |
| AshExt | May | Coleoptera | Lampyridae | 0.0000 | 1.0000 |
| AshExt | May | Coleoptera | Scarabaeidae | 0.0000 | 1.0000 |
| AshExt | May | Coleoptera | Scirtidae | 0.0000 | 1.0000 |
| AshExt | May | Diptera | Agromyzidae | 0.0000 | 1.0000 |
| AshExt | May | Diptera | Anisopodidae | 0.1364 | 1.0000 |
| AshExt | May | Diptera | Calliphoridae | 0.0000 | 1.0000 |
| AshExt | May | Diptera | Cecidomyiidae | 0.0984 | 1.0000 |
| AshExt | May | Diptera | Chaoboridae | 0.0000 | 1.0000 |
| AshExt | May | Diptera | Chironomidae | 0.1877 | 1.0000 |
| AshExt | May | Diptera | Chloropidae | 0.0000 | 1.0000 |
| AshExt | May | Diptera | Culicidae | 0.0000 | 1.0000 |
| AshExt | May | Diptera | Drosophilidae | 0.0000 | 1.0000 |
| AshExt | May | Diptera | Empididae | 0.0000 | 1.0000 |
| AshExt | May | Diptera | Hybotidae | 0.0000 | 1.0000 |
| AshExt | May | Diptera | Keroplatidae | 0.0000 | 1.0000 |
| AshExt | May | Diptera | Limoniidae | 0.0000 | 1.0000 |
| AshExt | May | Diptera | Muscidae | 0.0000 | 1.0000 |
| AshExt | May | Diptera | Mycetophilidae | 0.0000 | 1.0000 |

| | | | | | |
|---|---|---|---|---|---|
| AshExt | May | Diptera | Opomyzidae | 0.0000 | 1.0000 |
| AshExt | May | Diptera | Pallopteridae | 0.0000 | 1.0000 |
| AshExt | May | Diptera | Phoridae | 0.0000 | 1.0000 |
| AshExt | May | Diptera | Polleniidae | 0.0000 | 1.0000 |
| AshExt | May | Diptera | Psychodidae | 0.0000 | 1.0000 |
| AshExt | May | Diptera | Sarcophagidae | 0.0000 | 1.0000 |
| AshExt | May | Diptera | Sciaridae | 0.0294 | 1.0000 |
| AshExt | May | Diptera | Sphaeroceridae | 0.0000 | 1.0000 |
| AshExt | May | Diptera | Syrphidae | 0.0000 | 1.0000 |
| AshExt | May | Diptera | Tabanidae | 0.0000 | 1.0000 |
| AshExt | May | Diptera | Tachinidae | 0.0000 | 1.0000 |
| AshExt | May | Diptera | Tipulidae | 0.0000 | 1.0000 |
| AshExt | May | Ephemeroptera | Caenidae | 0.0000 | 1.0000 |
| AshExt | May | Ephemeroptera | Ephemeridae | 0.0000 | 1.0000 |
| AshExt | May | Ephemeroptera | Heptageniidae | 0.0000 | 1.0000 |
| AshExt | May | Hemiptera | Clastopteridae | 0.0000 | 1.0000 |
| AshExt | May | Hemiptera | Lygaeidae | 0.0000 | 1.0000 |
| AshExt | May | Hymenoptera | Apidae | 0.0000 | 1.0000 |
| AshExt | May | Hymenoptera | Braconidae | 0.0000 | 1.0000 |
| AshExt | May | Hymenoptera | Ichneumonidae | 0.0000 | 1.0000 |
| AshExt | May | Hymenoptera | Other | 0.0000 | 1.0000 |
| AshExt | May | Hymenoptera | Tenthredinidae | 0.0000 | 1.0000 |
| AshExt | May | Hymenoptera | Trigonalidae | 0.0000 | 1.0000 |
| AshExt | May | Lepidoptera | Crambidae | 0.0000 | 1.0000 |
| AshExt | May | Lepidoptera | Elachistidae | 0.0000 | 1.0000 |
| AshExt | May | Lepidoptera | Erebidae | 0.0000 | 1.0000 |
| AshExt | May | Lepidoptera | Gelechiidae | 0.0000 | 1.0000 |
| AshExt | May | Lepidoptera | Noctuidae | 0.0000 | 1.0000 |
| AshExt | May | Lepidoptera | Notodontidae | 0.0000 | 1.0000 |
| AshExt | May | Lepidoptera | Nymphalidae | 0.0000 | 1.0000 |
| AshExt | May | Lepidoptera | Pyralidae | 0.0000 | 1.0000 |

| | | | | | |
|---|---|---|---|---|---|
| AshExt | May | Lepidoptera | Tischeriidae | 0.0000 | 1.0000 |
| AshExt | May | Lepidoptera | Tortricidae | 0.1018 | 1.0000 |
| AshExt | May | Megaloptera | Corydalidae | 0.0000 | 1.0000 |
| AshExt | May | Neuroptera | Hemerobiidae | 0.0000 | 1.0000 |
| AshExt | May | Odonata | Libellulidae | 0.0000 | 1.0000 |
| AshExt | May | Orthoptera | Tettigoniidae | 0.0000 | 1.0000 |
| AshExt | May | Other | Other | 0.0294 | 1.0000 |
| AshExt | May | Plecoptera | Chloroperlidae | 0.0000 | 1.0000 |
| AshExt | May | Psocodea | Peripsocidae | 0.0000 | 1.0000 |
| AshExt | May | Sarcoptiformes | Oripodidae | 0.0000 | 1.0000 |
| AshExt | May | Trichoptera | Dipseudopsidae | 0.0000 | 1.0000 |
| AshExt | May | Trichoptera | Hydroptilidae | 0.0000 | 1.0000 |
| AshExt | May | Trichoptera | Leptoceridae | 0.0000 | 1.0000 |
| AshExt | May | Trombidiformes | Arrenuridae | 0.0000 | 1.0000 |
| AshExt | May | Trombidiformes | Aturidae | 0.0000 | 1.0000 |
| AshExt | May | Trombidiformes | Axonopsidae | 0.0000 | 1.0000 |
| AshExt | May | Trombidiformes | Eupodidae | 0.0000 | 1.0000 |
| AshExt | May | Trombidiformes | Feltriidae | 0.0000 | 1.0000 |
| AshExt | May | Trombidiformes | Hydryphantidae | 0.0000 | 1.0000 |
| AshExt | May | Trombidiformes | Lebertiidae | 0.0000 | 1.0000 |
| AshExt | May | Trombidiformes | Mideopsidae | 0.0327 | 1.0000 |
| AshExt | May | Trombidiformes | Pionidae | 0.1709 | 1.0000 |
| AshExt | May | Trombidiformes | Sperchontidae | 0.0000 | 1.0000 |
| AshExt | May | Trombidiformes | Tarsonemidae | 0.0000 | 1.0000 |
| AshExt | May | Trombidiformes | Torrenticolidae | 0.0000 | 1.0000 |
| AshExt | May | Trombidiformes | Unionicolidae | 0.0000 | 1.0000 |
| AshExt | June | Araneae | Agelenidae | 0.0000 | 1.0000 |
| AshExt | June | Araneae | Amaurobiidae | 0.0000 | 1.0000 |
| AshExt | June | Araneae | Anyphaenidae | 0.0037 | 1.0000 |
| AshExt | June | Araneae | Cicurinidae | 0.0000 | 1.0000 |
| AshExt | June | Araneae | Dictynidae | 0.0000 | 1.0000 |

| | | | | | |
|---|---|---|---|---|---|
| AshExt | June | Araneae | Gnaphosidae | 0.0000 | 1.0000 |
| AshExt | June | Araneae | Linyphiidae | 0.0000 | 1.0000 |
| AshExt | June | Araneae | Lycosidae | 0.0000 | 1.0000 |
| AshExt | June | Araneae | Phrurolithidae | 0.0000 | 1.0000 |
| AshExt | June | Araneae | Salticidae | 0.0147 | 1.0000 |
| AshExt | June | Araneae | Theridiidae | 0.0000 | 1.0000 |
| AshExt | June | Araneae | Thomisidae | 0.0661 | 1.0000 |
| AshExt | June | Blattodea | Ectobiidae | 0.0127 | 1.0000 |
| AshExt | June | Coleoptera | Buprestidae | 0.0000 | 1.0000 |
| AshExt | June | Coleoptera | Carabidae | 0.0067 | 1.0000 |
| AshExt | June | Coleoptera | Curculionidae | 0.0208 | 1.0000 |
| AshExt | June | Coleoptera | Dytiscidae | 0.0000 | 1.0000 |
| AshExt | June | Coleoptera | Lampyridae | 0.0000 | 1.0000 |
| AshExt | June | Coleoptera | Scarabaeidae | 0.0000 | 1.0000 |
| AshExt | June | Coleoptera | Scirtidae | 0.0020 | 1.0000 |
| AshExt | June | Diptera | Agromyzidae | 0.0000 | 1.0000 |
| AshExt | June | Diptera | Anisopodidae | 0.0536 | 1.0000 |
| AshExt | June | Diptera | Calliphoridae | 0.0000 | 1.0000 |
| AshExt | June | Diptera | Cecidomyiidae | 0.1642 | 1.0000 |
| AshExt | June | Diptera | Chaoboridae | 0.0947 | 1.0000 |
| AshExt | June | Diptera | Chironomidae | 0.0947 | 1.0000 |
| AshExt | June | Diptera | Chloropidae | 0.0030 | 1.0000 |
| AshExt | June | Diptera | Culicidae | 0.0004 | 1.0000 |
| AshExt | June | Diptera | Drosophilidae | 0.0000 | 1.0000 |
| AshExt | June | Diptera | Empididae | 0.0000 | 1.0000 |
| AshExt | June | Diptera | Hybotidae | 0.0000 | 1.0000 |
| AshExt | June | Diptera | Keroplatidae | 0.0000 | 1.0000 |
| AshExt | June | Diptera | Limoniidae | 0.0011 | 1.0000 |
| AshExt | June | Diptera | Muscidae | 0.0000 | 1.0000 |
| AshExt | June | Diptera | Mycetophilidae | 0.0000 | 1.0000 |
| AshExt | June | Diptera | Opomyzidae | 0.0000 | 1.0000 |

| | | | | | |
|---|---|---|---|---|---|
| AshExt | June | Diptera | Pallopteridae | 0.0000 | 1.0000 |
| AshExt | June | Diptera | Phoridae | 0.0000 | 1.0000 |
| AshExt | June | Diptera | Polleniidae | 0.0000 | 1.0000 |
| AshExt | June | Diptera | Psychodidae | 0.0000 | 1.0000 |
| AshExt | June | Diptera | Sarcophagidae | 0.0000 | 1.0000 |
| AshExt | June | Diptera | Sciaridae | 0.0067 | 1.0000 |
| AshExt | June | Diptera | Sphaeroceridae | 0.0000 | 1.0000 |
| AshExt | June | Diptera | Syrphidae | 0.0000 | 1.0000 |
| AshExt | June | Diptera | Tabanidae | 0.0000 | 1.0000 |
| AshExt | June | Diptera | Tachinidae | 0.0000 | 1.0000 |
| AshExt | June | Diptera | Tipulidae | 0.0000 | 1.0000 |
| AshExt | June | Ephemeroptera | Caenidae | 0.0133 | 1.0000 |
| AshExt | June | Ephemeroptera | Ephemeridae | 0.0000 | 1.0000 |
| AshExt | June | Ephemeroptera | Heptageniidae | 0.0000 | 1.0000 |
| AshExt | June | Hemiptera | Clastopteridae | 0.0000 | 1.0000 |
| AshExt | June | Hemiptera | Lygaeidae | 0.0000 | 1.0000 |
| AshExt | June | Hymenoptera | Apidae | 0.0000 | 1.0000 |
| AshExt | June | Hymenoptera | Braconidae | 0.0000 | 1.0000 |
| AshExt | June | Hymenoptera | Ichneumonidae | 0.0000 | 1.0000 |
| AshExt | June | Hymenoptera | Other | 0.0033 | 1.0000 |
| AshExt | June | Hymenoptera | Tenthredinidae | 0.0164 | 1.0000 |
| AshExt | June | Hymenoptera | Trigonalidae | 0.0123 | 1.0000 |
| AshExt | June | Lepidoptera | Crambidae | 0.0000 | 1.0000 |
| AshExt | June | Lepidoptera | Elachistidae | 0.0000 | 1.0000 |
| AshExt | June | Lepidoptera | Erebidae | 0.0000 | 1.0000 |
| AshExt | June | Lepidoptera | Gelechiidae | 0.0000 | 1.0000 |
| AshExt | June | Lepidoptera | Noctuidae | 0.0000 | 1.0000 |
| AshExt | June | Lepidoptera | Notodontidae | 0.0000 | 1.0000 |
| AshExt | June | Lepidoptera | Nymphalidae | 0.0002 | 1.0000 |
| AshExt | June | Lepidoptera | Pyralidae | 0.0053 | 1.0000 |
| AshExt | June | Lepidoptera | Tischeriidae | 0.0067 | 1.0000 |

| | | | | | |
|---|---|---|---|---|---|
| AshExt | June | Lepidoptera | Tortricidae | 0.1393 | 1.0000 |
| AshExt | June | Megaloptera | Corydalidae | 0.0000 | 1.0000 |
| AshExt | June | Neuroptera | Hemerobiidae | 0.0000 | 1.0000 |
| AshExt | June | Odonata | Libellulidae | 0.0000 | 1.0000 |
| AshExt | June | Orthoptera | Tettigoniidae | 0.0000 | 1.0000 |
| AshExt | June | Other | Other | 0.0024 | 1.0000 |
| AshExt | June | Plecoptera | Chloroperlidae | 0.0000 | 1.0000 |
| AshExt | June | Psocodea | Peripsocidae | 0.0000 | 1.0000 |
| AshExt | June | Sarcoptiformes | Oripodidae | 0.0000 | 1.0000 |
| AshExt | June | Trichoptera | Dipseudopsidae | 0.0030 | 1.0000 |
| AshExt | June | Trichoptera | Hydroptilidae | 0.0000 | 1.0000 |
| AshExt | June | Trichoptera | Leptoceridae | 0.0004 | 1.0000 |
| AshExt | June | Trombidiformes | Arrenuridae | 0.0600 | 1.0000 |
| AshExt | June | Trombidiformes | Aturidae | 0.0000 | 1.0000 |
| AshExt | June | Trombidiformes | Axonopsidae | 0.0204 | 1.0000 |
| AshExt | June | Trombidiformes | Eupodidae | 0.0000 | 1.0000 |
| AshExt | June | Trombidiformes | Feltriidae | 0.0000 | 1.0000 |
| AshExt | June | Trombidiformes | Hydryphantidae | 0.0049 | 1.0000 |
| AshExt | June | Trombidiformes | Lebertiidae | 0.0004 | 1.0000 |
| AshExt | June | Trombidiformes | Mideopsidae | 0.1663 | 1.0000 |
| AshExt | June | Trombidiformes | Pionidae | 0.0004 | 1.0000 |
| AshExt | June | Trombidiformes | Sperchontidae | 0.0000 | 1.0000 |
| AshExt | June | Trombidiformes | Tarsonemidae | 0.0000 | 1.0000 |
| AshExt | June | Trombidiformes | Torrenticolidae | 0.0000 | 1.0000 |
| AshExt | June | Trombidiformes | Unionicolidae | 0.0000 | 1.0000 |
| AshExt | July | Araneae | Agelenidae | 0.0000 | 1.0000 |
| AshExt | July | Araneae | Amaurobiidae | 0.0000 | 1.0000 |
| AshExt | July | Araneae | Anyphaenidae | 0.0978 | 1.0000 |
| AshExt | July | Araneae | Cicurinidae | 0.0000 | 1.0000 |
| AshExt | July | Araneae | Dictynidae | 0.0000 | 1.0000 |
| AshExt | July | Araneae | Gnaphosidae | 0.0000 | 1.0000 |

| | | | | | |
|---|---|---|---|---|---|
| AshExt | July | Araneae | Linyphiidae | 0.0000 | 1.0000 |
| AshExt | July | Araneae | Lycosidae | 0.0000 | 1.0000 |
| AshExt | July | Araneae | Phrurolithidae | 0.0000 | 1.0000 |
| AshExt | July | Araneae | Salticidae | 0.0000 | 1.0000 |
| AshExt | July | Araneae | Theridiidae | 0.1087 | 1.0000 |
| AshExt | July | Araneae | Thomisidae | 0.0000 | 1.0000 |
| AshExt | July | Blattodea | Ectobiidae | 0.0000 | 1.0000 |
| AshExt | July | Coleoptera | Buprestidae | 0.0000 | 1.0000 |
| AshExt | July | Coleoptera | Carabidae | 0.1111 | 1.0000 |
| AshExt | July | Coleoptera | Curculionidae | 0.1111 | 1.0000 |
| AshExt | July | Coleoptera | Dytiscidae | 0.0000 | 1.0000 |
| AshExt | July | Coleoptera | Lampyridae | 0.0000 | 1.0000 |
| AshExt | July | Coleoptera | Scarabaeidae | 0.0000 | 1.0000 |
| AshExt | July | Coleoptera | Scirtidae | 0.0000 | 1.0000 |
| AshExt | July | Diptera | Agromyzidae | 0.0000 | 1.0000 |
| AshExt | July | Diptera | Anisopodidae | 0.0000 | 1.0000 |
| AshExt | July | Diptera | Calliphoridae | 0.0000 | 1.0000 |
| AshExt | July | Diptera | Cecidomyiidae | 0.3623 | 1.0000 |
| AshExt | July | Diptera | Chaoboridae | 0.0000 | 1.0000 |
| AshExt | July | Diptera | Chironomidae | 0.0000 | 1.0000 |
| AshExt | July | Diptera | Chloropidae | 0.0000 | 1.0000 |
| AshExt | July | Diptera | Culicidae | 0.0000 | 1.0000 |
| AshExt | July | Diptera | Drosophilidae | 0.0000 | 1.0000 |
| AshExt | July | Diptera | Empididae | 0.0000 | 1.0000 |
| AshExt | July | Diptera | Hybotidae | 0.0000 | 1.0000 |
| AshExt | July | Diptera | Keroplatidae | 0.0000 | 1.0000 |
| AshExt | July | Diptera | Limoniidae | 0.0000 | 1.0000 |
| AshExt | July | Diptera | Muscidae | 0.0000 | 1.0000 |
| AshExt | July | Diptera | Mycetophilidae | 0.0000 | 1.0000 |
| AshExt | July | Diptera | Opomyzidae | 0.0000 | 1.0000 |
| AshExt | July | Diptera | Pallopteridae | 0.0000 | 1.0000 |

| | | | | | |
|---|---|---|---|---|---|
| AshExt | July | Diptera | Phoridae | 0.0000 | 1.0000 |
| AshExt | July | Diptera | Polleniidae | 0.0000 | 1.0000 |
| AshExt | July | Diptera | Psychodidae | 0.0000 | 1.0000 |
| AshExt | July | Diptera | Sarcophagidae | 0.0000 | 1.0000 |
| AshExt | July | Diptera | Sciaridae | 0.0978 | 1.0000 |
| AshExt | July | Diptera | Sphaeroceridae | 0.0000 | 1.0000 |
| AshExt | July | Diptera | Syrphidae | 0.0000 | 1.0000 |
| AshExt | July | Diptera | Tabanidae | 0.0000 | 1.0000 |
| AshExt | July | Diptera | Tachinidae | 0.0000 | 1.0000 |
| AshExt | July | Diptera | Tipulidae | 0.0000 | 1.0000 |
| AshExt | July | Ephemeroptera | Caenidae | 0.0000 | 1.0000 |
| AshExt | July | Ephemeroptera | Ephemeridae | 0.0000 | 1.0000 |
| AshExt | July | Ephemeroptera | Heptageniidae | 0.0000 | 1.0000 |
| AshExt | July | Hemiptera | Clastopteridae | 0.0000 | 1.0000 |
| AshExt | July | Hemiptera | Lygaeidae | 0.0000 | 1.0000 |
| AshExt | July | Hymenoptera | Apidae | 0.0000 | 1.0000 |
| AshExt | July | Hymenoptera | Braconidae | 0.0000 | 1.0000 |
| AshExt | July | Hymenoptera | Ichneumonidae | 0.0000 | 1.0000 |
| AshExt | July | Hymenoptera | Other | 0.0000 | 1.0000 |
| AshExt | July | Hymenoptera | Tenthredinidae | 0.0000 | 1.0000 |
| AshExt | July | Hymenoptera | Trigonalidae | 0.0000 | 1.0000 |
| AshExt | July | Lepidoptera | Crambidae | 0.0000 | 1.0000 |
| AshExt | July | Lepidoptera | Elachistidae | 0.0000 | 1.0000 |
| AshExt | July | Lepidoptera | Erebidae | 0.0000 | 1.0000 |
| AshExt | July | Lepidoptera | Gelechiidae | 0.0000 | 1.0000 |
| AshExt | July | Lepidoptera | Noctuidae | 0.0000 | 1.0000 |
| AshExt | July | Lepidoptera | Notodontidae | 0.0000 | 1.0000 |
| AshExt | July | Lepidoptera | Nymphalidae | 0.0000 | 1.0000 |
| AshExt | July | Lepidoptera | Pyralidae | 0.0000 | 1.0000 |
| AshExt | July | Lepidoptera | Tischeriidae | 0.0000 | 1.0000 |
| AshExt | July | Lepidoptera | Tortricidae | 0.0000 | 1.0000 |

| | | | | | |
|---|---|---|---|---|---|
| AshExt | July | Megaloptera | Corydalidae | 0.0000 | 1.0000 |
| AshExt | July | Neuroptera | Hemerobiidae | 0.0000 | 1.0000 |
| AshExt | July | Odonata | Libellulidae | 0.0000 | 1.0000 |
| AshExt | July | Orthoptera | Tettigoniidae | 0.0000 | 1.0000 |
| AshExt | July | Other | Other | 0.0000 | 1.0000 |
| AshExt | July | Plecoptera | Chloroperlidae | 0.0000 | 1.0000 |
| AshExt | July | Psocodea | Peripsocidae | 0.0000 | 1.0000 |
| AshExt | July | Sarcoptiformes | Oripodidae | 0.0000 | 1.0000 |
| AshExt | July | Trichoptera | Dipseudopsidae | 0.0000 | 1.0000 |
| AshExt | July | Trichoptera | Hydroptilidae | 0.0000 | 1.0000 |
| AshExt | July | Trichoptera | Leptoceridae | 0.0000 | 1.0000 |
| AshExt | July | Trombidiformes | Arrenuridae | 0.0000 | 1.0000 |
| AshExt | July | Trombidiformes | Aturidae | 0.0000 | 1.0000 |
| AshExt | July | Trombidiformes | Axonopsidae | 0.0000 | 1.0000 |
| AshExt | July | Trombidiformes | Eupodidae | 0.1111 | 1.0000 |
| AshExt | July | Trombidiformes | Feltriidae | 0.0000 | 1.0000 |
| AshExt | July | Trombidiformes | Hydryphantidae | 0.0000 | 1.0000 |
| AshExt | July | Trombidiformes | Lebertiidae | 0.0000 | 1.0000 |
| AshExt | July | Trombidiformes | Mideopsidae | 0.0000 | 1.0000 |
| AshExt | July | Trombidiformes | Pionidae | 0.0000 | 1.0000 |
| AshExt | July | Trombidiformes | Sperchontidae | 0.0000 | 1.0000 |
| AshExt | July | Trombidiformes | Tarsonemidae | 0.0000 | 1.0000 |
| AshExt | July | Trombidiformes | Torrenticolidae | 0.0000 | 1.0000 |
| AshExt | July | Trombidiformes | Unionicolidae | 0.0000 | 1.0000 |
| AshExt | August | Araneae | Agelenidae | 0.0000 | 1.0000 |
| AshExt | August | Araneae | Amaurobiidae | 0.0000 | 1.0000 |
| AshExt | August | Araneae | Anyphaenidae | 0.0973 | 1.0000 |
| AshExt | August | Araneae | Cicurinidae | 0.0000 | 1.0000 |
| AshExt | August | Araneae | Dictynidae | 0.0088 | 1.0000 |
| AshExt | August | Araneae | Gnaphosidae | 0.0000 | 1.0000 |
| AshExt | August | Araneae | Linyphiidae | 0.0000 | 1.0000 |

| | | | | | |
|---|---|---|---|---|---|
| AshExt | August | Araneae | Lycosidae | 0.0000 | 1.0000 |
| AshExt | August | Araneae | Phrurolithidae | 0.1457 | 1.0000 |
| AshExt | August | Araneae | Salticidae | 0.0013 | 1.0000 |
| AshExt | August | Araneae | Theridiidae | 0.0020 | 1.0000 |
| AshExt | August | Araneae | Thomisidae | 0.1441 | 1.0000 |
| AshExt | August | Blattodea | Ectobiidae | 0.0000 | 1.0000 |
| AshExt | August | Coleoptera | Buprestidae | 0.0000 | 1.0000 |
| AshExt | August | Coleoptera | Carabidae | 0.0439 | 1.0000 |
| AshExt | August | Coleoptera | Curculionidae | 0.0333 | 1.0000 |
| AshExt | August | Coleoptera | Dytiscidae | 0.0000 | 1.0000 |
| AshExt | August | Coleoptera | Lampyridae | 0.0004 | 1.0000 |
| AshExt | August | Coleoptera | Scarabaeidae | 0.0007 | 1.0000 |
| AshExt | August | Coleoptera | Scirtidae | 0.0043 | 1.0000 |
| AshExt | August | Diptera | Agromyzidae | 0.0007 | 1.0000 |
| AshExt | August | Diptera | Anisopodidae | 0.0117 | 1.0000 |
| AshExt | August | Diptera | Calliphoridae | 0.0000 | 1.0000 |
| AshExt | August | Diptera | Cecidomyiidae | 0.2024 | 1.0000 |
| AshExt | August | Diptera | Chaoboridae | 0.0059 | 1.0000 |
| AshExt | August | Diptera | Chironomidae | 0.0078 | 1.0000 |
| AshExt | August | Diptera | Chloropidae | 0.0013 | 1.0000 |
| AshExt | August | Diptera | Culicidae | 0.0007 | 1.0000 |
| AshExt | August | Diptera | Drosophilidae | 0.0001 | 1.0000 |
| AshExt | August | Diptera | Empididae | 0.0000 | 1.0000 |
| AshExt | August | Diptera | Hybotidae | 0.0000 | 1.0000 |
| AshExt | August | Diptera | Keroplatidae | 0.0000 | 1.0000 |
| AshExt | August | Diptera | Limoniidae | 0.0770 | 1.0000 |
| AshExt | August | Diptera | Muscidae | 0.0000 | 1.0000 |
| AshExt | August | Diptera | Mycetophilidae | 0.0000 | 1.0000 |
| AshExt | August | Diptera | Opomyzidae | 0.0000 | 1.0000 |
| AshExt | August | Diptera | Pallopteridae | 0.0000 | 1.0000 |
| AshExt | August | Diptera | Phoridae | 0.0000 | 1.0000 |

| | | | | | |
|---|---|---|---|---|---|
| AshExt | August | Diptera | Polleniidae | 0.0000 | 1.0000 |
| AshExt | August | Diptera | Psychodidae | 0.0000 | 1.0000 |
| AshExt | August | Diptera | Sarcophagidae | 0.0000 | 1.0000 |
| AshExt | August | Diptera | Sciaridae | 0.0417 | 1.0000 |
| AshExt | August | Diptera | Sphaeroceridae | 0.0000 | 1.0000 |
| AshExt | August | Diptera | Syrphidae | 0.0007 | 1.0000 |
| AshExt | August | Diptera | Tabanidae | 0.0085 | 1.0000 |
| AshExt | August | Diptera | Tachinidae | 0.0000 | 1.0000 |
| AshExt | August | Diptera | Tipulidae | 0.0072 | 1.0000 |
| AshExt | August | Ephemeroptera | Caenidae | 0.0000 | 1.0000 |
| AshExt | August | Ephemeroptera | Ephemeridae | 0.0000 | 1.0000 |
| AshExt | August | Ephemeroptera | Heptageniidae | 0.0000 | 1.0000 |
| AshExt | August | Hemiptera | Clastopteridae | 0.0026 | 1.0000 |
| AshExt | August | Hemiptera | Lygaeidae | 0.0005 | 1.0000 |
| AshExt | August | Hymenoptera | Apidae | 0.0000 | 1.0000 |
| AshExt | August | Hymenoptera | Braconidae | 0.0000 | 1.0000 |
| AshExt | August | Hymenoptera | Ichneumonidae | 0.0000 | 1.0000 |
| AshExt | August | Hymenoptera | Other | 0.0007 | 1.0000 |
| AshExt | August | Hymenoptera | Tenthredinidae | 0.0000 | 1.0000 |
| AshExt | August | Hymenoptera | Trigonalidae | 0.0000 | 1.0000 |
| AshExt | August | Lepidoptera | Crambidae | 0.0000 | 1.0000 |
| AshExt | August | Lepidoptera | Elachistidae | 0.0000 | 1.0000 |
| AshExt | August | Lepidoptera | Erebidae | 0.0000 | 1.0000 |
| AshExt | August | Lepidoptera | Gelechiidae | 0.0020 | 1.0000 |
| AshExt | August | Lepidoptera | Noctuidae | 0.0005 | 1.0000 |
| AshExt | August | Lepidoptera | Notodontidae | 0.0000 | 1.0000 |
| AshExt | August | Lepidoptera | Nymphalidae | 0.0000 | 1.0000 |
| AshExt | August | Lepidoptera | Pyralidae | 0.0000 | 1.0000 |
| AshExt | August | Lepidoptera | Tischeriidae | 0.0000 | 1.0000 |
| AshExt | August | Lepidoptera | Tortricidae | 0.0003 | 1.0000 |
| AshExt | August | Megaloptera | Corydalidae | 0.0000 | 1.0000 |

| | | | | | |
|---|---|---|---|---|---|
| AshExt | August | Neuroptera | Hemerobiidae | 0.0000 | 1.0000 |
| AshExt | August | Odonata | Libellulidae | 0.0000 | 1.0000 |
| AshExt | August | Orthoptera | Tettigoniidae | 0.0000 | 1.0000 |
| AshExt | August | Other | Other | 0.0324 | 1.0000 |
| AshExt | August | Plecoptera | Chloroperlidae | 0.0000 | 1.0000 |
| AshExt | August | Psocodea | Peripsocidae | 0.0000 | 1.0000 |
| AshExt | August | Sarcoptiformes | Oripodidae | 0.0000 | 1.0000 |
| AshExt | August | Trichoptera | Dipseudopsidae | 0.0000 | 1.0000 |
| AshExt | August | Trichoptera | Hydroptilidae | 0.0000 | 1.0000 |
| AshExt | August | Trichoptera | Leptoceridae | 0.0000 | 1.0000 |
| AshExt | August | Trombidiformes | Arrenuridae | 0.0000 | 1.0000 |
| AshExt | August | Trombidiformes | Aturidae | 0.0000 | 1.0000 |
| AshExt | August | Trombidiformes | Axonopsidae | 0.0013 | 1.0000 |
| AshExt | August | Trombidiformes | Eupodidae | 0.0000 | 1.0000 |
| AshExt | August | Trombidiformes | Feltriidae | 0.0000 | 1.0000 |
| AshExt | August | Trombidiformes | Hydryphantidae | 0.0000 | 1.0000 |
| AshExt | August | Trombidiformes | Lebertiidae | 0.1113 | 1.0000 |
| AshExt | August | Trombidiformes | Mideopsidae | 0.0000 | 1.0000 |
| AshExt | August | Trombidiformes | Pionidae | 0.0000 | 1.0000 |
| AshExt | August | Trombidiformes | Sperchontidae | 0.0013 | 1.0000 |
| AshExt | August | Trombidiformes | Tarsonemidae | 0.0000 | 1.0000 |
| AshExt | August | Trombidiformes | Torrenticolidae | 0.0000 | 1.0000 |
| AshExt | August | Trombidiformes | Unionicolidae | 0.0000 | 1.0000 |
| WMCC | May | Araneae | Agelenidae | 0.0022 | 1.0000 |
| WMCC | May | Araneae | Amaurobiidae | 0.0000 | 1.0000 |
| WMCC | May | Araneae | Anyphaenidae | 0.0106 | 1.0000 |
| WMCC | May | Araneae | Cicurinidae | 0.0000 | 1.0000 |
| WMCC | May | Araneae | Dictynidae | 0.0011 | 1.0000 |
| WMCC | May | Araneae | Gnaphosidae | 0.0000 | 1.0000 |
| WMCC | May | Araneae | Linyphiidae | 0.0000 | 1.0000 |
| WMCC | May | Araneae | Lycosidae | 0.0000 | 1.0000 |

| | | | | | |
|---|---|---|---|---|---|
| WMCC | May | Araneae | Phrurolithidae | 0.0002 | 1.0000 |
| WMCC | May | Araneae | Salticidae | 0.0098 | 1.0000 |
| WMCC | May | Araneae | Theridiidae | 0.0056 | 1.0000 |
| WMCC | May | Araneae | Thomisidae | 0.0178 | 1.0000 |
| WMCC | May | Blattodea | Ectobiidae | 0.0000 | 1.0000 |
| WMCC | May | Coleoptera | Buprestidae | 0.0019 | 1.0000 |
| WMCC | May | Coleoptera | Carabidae | 0.0209 | 1.0000 |
| WMCC | May | Coleoptera | Curculionidae | 0.0122 | 1.0000 |
| WMCC | May | Coleoptera | Dytiscidae | 0.0000 | 1.0000 |
| WMCC | May | Coleoptera | Lampyridae | 0.0000 | 1.0000 |
| WMCC | May | Coleoptera | Scarabaeidae | 0.0000 | 1.0000 |
| WMCC | May | Coleoptera | Scirtidae | 0.0000 | 1.0000 |
| WMCC | May | Diptera | Agromyzidae | 0.0065 | 1.0000 |
| WMCC | May | Diptera | Anisopodidae | 0.3963 | 1.0000 |
| WMCC | May | Diptera | Calliphoridae | 0.0000 | 1.0000 |
| WMCC | May | Diptera | Cecidomyiidae | 0.1913 | 1.0000 |
| WMCC | May | Diptera | Chaoboridae | 0.0198 | 1.0000 |
| WMCC | May | Diptera | Chironomidae | 0.1436 | 1.0000 |
| WMCC | May | Diptera | Chloropidae | 0.0079 | 1.0000 |
| WMCC | May | Diptera | Culicidae | 0.0000 | 1.0000 |
| WMCC | May | Diptera | Drosophilidae | 0.0002 | 1.0000 |
| WMCC | May | Diptera | Empididae | 0.0000 | 1.0000 |
| WMCC | May | Diptera | Hybotidae | 0.0021 | 1.0000 |
| WMCC | May | Diptera | Keroplatidae | 0.0000 | 1.0000 |
| WMCC | May | Diptera | Limoniidae | 0.0011 | 1.0000 |
| WMCC | May | Diptera | Muscidae | 0.0000 | 1.0000 |
| WMCC | May | Diptera | Mycetophilidae | 0.0000 | 1.0000 |
| WMCC | May | Diptera | Opomyzidae | 0.0000 | 1.0000 |
| WMCC | May | Diptera | Pallopteridae | 0.0000 | 1.0000 |
| WMCC | May | Diptera | Phoridae | 0.0038 | 1.0000 |
| WMCC | May | Diptera | Polleniidae | 0.0000 | 1.0000 |

| | | | | | |
|---|---|---|---|---|---|
| WMCC | May | Diptera | Psychodidae | 0.0000 | 1.0000 |
| WMCC | May | Diptera | Sarcophagidae | 0.0000 | 1.0000 |
| WMCC | May | Diptera | Sciaridae | 0.0181 | 1.0000 |
| WMCC | May | Diptera | Sphaeroceridae | 0.0000 | 1.0000 |
| WMCC | May | Diptera | Syrphidae | 0.0000 | 1.0000 |
| WMCC | May | Diptera | Tabanidae | 0.0000 | 1.0000 |
| WMCC | May | Diptera | Tachinidae | 0.0000 | 1.0000 |
| WMCC | May | Diptera | Tipulidae | 0.0016 | 1.0000 |
| WMCC | May | Ephemeroptera | Caenidae | 0.0000 | 1.0000 |
| WMCC | May | Ephemeroptera | Ephemeridae | 0.0000 | 1.0000 |
| WMCC | May | Ephemeroptera | Heptageniidae | 0.0000 | 1.0000 |
| WMCC | May | Hemiptera | Clastopteridae | 0.0000 | 1.0000 |
| WMCC | May | Hemiptera | Lygaeidae | 0.0000 | 1.0000 |
| WMCC | May | Hymenoptera | Apidae | 0.0000 | 1.0000 |
| WMCC | May | Hymenoptera | Braconidae | 0.0050 | 1.0000 |
| WMCC | May | Hymenoptera | Ichneumonidae | 0.0007 | 1.0000 |
| WMCC | May | Hymenoptera | Other | 0.0037 | 1.0000 |
| WMCC | May | Hymenoptera | Tenthredinidae | 0.0000 | 1.0000 |
| WMCC | May | Hymenoptera | Trigonalidae | 0.0000 | 1.0000 |
| WMCC | May | Lepidoptera | Crambidae | 0.0000 | 1.0000 |
| WMCC | May | Lepidoptera | Elachistidae | 0.0000 | 1.0000 |
| WMCC | May | Lepidoptera | Erebidae | 0.0000 | 1.0000 |
| WMCC | May | Lepidoptera | Gelechiidae | 0.0000 | 1.0000 |
| WMCC | May | Lepidoptera | Noctuidae | 0.0000 | 1.0000 |
| WMCC | May | Lepidoptera | Notodontidae | 0.0000 | 1.0000 |
| WMCC | May | Lepidoptera | Nymphalidae | 0.0000 | 1.0000 |
| WMCC | May | Lepidoptera | Pyralidae | 0.0118 | 1.0000 |
| WMCC | May | Lepidoptera | Tischeriidae | 0.0000 | 1.0000 |
| WMCC | May | Lepidoptera | Tortricidae | 0.0000 | 1.0000 |
| WMCC | May | Megaloptera | Corydalidae | 0.0000 | 1.0000 |
| WMCC | May | Neuroptera | Hemerobiidae | 0.0000 | 1.0000 |

| | | | | | |
|---|---|---|---|---|---|
| WMCC | May | Odonata | Libellulidae | 0.0000 | 1.0000 |
| WMCC | May | Orthoptera | Tettigoniidae | 0.0000 | 1.0000 |
| WMCC | May | Other | Other | 0.0326 | 1.0000 |
| WMCC | May | Plecoptera | Chloroperlidae | 0.0021 | 1.0000 |
| WMCC | May | Psocodea | Peripsocidae | 0.0000 | 1.0000 |
| WMCC | May | Sarcoptiformes | Oripodidae | 0.0000 | 1.0000 |
| WMCC | May | Trichoptera | Dipseudopsidae | 0.0000 | 1.0000 |
| WMCC | May | Trichoptera | Hydroptilidae | 0.0000 | 1.0000 |
| WMCC | May | Trichoptera | Leptoceridae | 0.0000 | 1.0000 |
| WMCC | May | Trombidiformes | Arrenuridae | 0.0000 | 1.0000 |
| WMCC | May | Trombidiformes | Aturidae | 0.0000 | 1.0000 |
| WMCC | May | Trombidiformes | Axonopsidae | 0.0000 | 1.0000 |
| WMCC | May | Trombidiformes | Eupodidae | 0.0274 | 1.0000 |
| WMCC | May | Trombidiformes | Feltriidae | 0.0000 | 1.0000 |
| WMCC | May | Trombidiformes | Hydryphantidae | 0.0000 | 1.0000 |
| WMCC | May | Trombidiformes | Lebertiidae | 0.0410 | 1.0000 |
| WMCC | May | Trombidiformes | Mideopsidae | 0.0000 | 1.0000 |
| WMCC | May | Trombidiformes | Pionidae | 0.0000 | 1.0000 |
| WMCC | May | Trombidiformes | Sperchontidae | 0.0000 | 1.0000 |
| WMCC | May | Trombidiformes | Tarsonemidae | 0.0000 | 1.0000 |
| WMCC | May | Trombidiformes | Torrenticolidae | 0.0011 | 1.0000 |
| WMCC | May | Trombidiformes | Unionicolidae | 0.0000 | 1.0000 |
| WMCC | June | Araneae | Agelenidae | 0.0000 | 1.0000 |
| WMCC | June | Araneae | Amaurobiidae | 0.0000 | 1.0000 |
| WMCC | June | Araneae | Anyphaenidae | 0.0652 | 1.0000 |
| WMCC | June | Araneae | Cicurinidae | 0.0000 | 1.0000 |
| WMCC | June | Araneae | Dictynidae | 0.0000 | 1.0000 |
| WMCC | June | Araneae | Gnaphosidae | 0.0000 | 1.0000 |
| WMCC | June | Araneae | Linyphiidae | 0.0000 | 1.0000 |
| WMCC | June | Araneae | Lycosidae | 0.0000 | 1.0000 |
| WMCC | June | Araneae | Phrurolithidae | 0.0061 | 1.0000 |

| | | | | | |
|---|---|---|---|---|---|
| WMCC | June | Araneae | Salticidae | 0.0000 | 1.0000 |
| WMCC | June | Araneae | Theridiidae | 0.0000 | 1.0000 |
| WMCC | June | Araneae | Thomisidae | 0.0332 | 1.0000 |
| WMCC | June | Blattodea | Ectobiidae | 0.0000 | 1.0000 |
| WMCC | June | Coleoptera | Buprestidae | 0.0000 | 1.0000 |
| WMCC | June | Coleoptera | Carabidae | 0.0775 | 1.0000 |
| WMCC | June | Coleoptera | Curculionidae | 0.0056 | 1.0000 |
| WMCC | June | Coleoptera | Dytiscidae | 0.0000 | 1.0000 |
| WMCC | June | Coleoptera | Lampyridae | 0.0000 | 1.0000 |
| WMCC | June | Coleoptera | Scarabaeidae | 0.0000 | 1.0000 |
| WMCC | June | Coleoptera | Scirtidae | 0.0000 | 1.0000 |
| WMCC | June | Diptera | Agromyzidae | 0.0000 | 1.0000 |
| WMCC | June | Diptera | Anisopodidae | 0.0303 | 1.0000 |
| WMCC | June | Diptera | Calliphoridae | 0.0000 | 1.0000 |
| WMCC | June | Diptera | Cecidomyiidae | 0.1465 | 1.0000 |
| WMCC | June | Diptera | Chaoboridae | 0.1538 | 1.0000 |
| WMCC | June | Diptera | Chironomidae | 0.0000 | 1.0000 |
| WMCC | June | Diptera | Chloropidae | 0.0000 | 1.0000 |
| WMCC | June | Diptera | Culicidae | 0.0000 | 1.0000 |
| WMCC | June | Diptera | Drosophilidae | 0.0000 | 1.0000 |
| WMCC | June | Diptera | Empididae | 0.0000 | 1.0000 |
| WMCC | June | Diptera | Hybotidae | 0.0000 | 1.0000 |
| WMCC | June | Diptera | Keroplatidae | 0.0000 | 1.0000 |
| WMCC | June | Diptera | Limoniidae | 0.0000 | 1.0000 |
| WMCC | June | Diptera | Muscidae | 0.0000 | 1.0000 |
| WMCC | June | Diptera | Mycetophilidae | 0.0056 | 1.0000 |
| WMCC | June | Diptera | Opomyzidae | 0.0000 | 1.0000 |
| WMCC | June | Diptera | Pallopteridae | 0.0000 | 1.0000 |
| WMCC | June | Diptera | Phoridae | 0.0000 | 1.0000 |
| WMCC | June | Diptera | Polleniidae | 0.0000 | 1.0000 |
| WMCC | June | Diptera | Psychodidae | 0.1966 | 1.0000 |

| | | | | | |
|---|---|---|---|---|---|
| WMCC | June | Diptera | Sarcophagidae | 0.0000 | 1.0000 |
| WMCC | June | Diptera | Sciaridae | 0.0051 | 1.0000 |
| WMCC | June | Diptera | Sphaeroceridae | 0.0000 | 1.0000 |
| WMCC | June | Diptera | Syrphidae | 0.0056 | 1.0000 |
| WMCC | June | Diptera | Tabanidae | 0.0000 | 1.0000 |
| WMCC | June | Diptera | Tachinidae | 0.0000 | 1.0000 |
| WMCC | June | Diptera | Tipulidae | 0.0000 | 1.0000 |
| WMCC | June | Ephemeroptera | Caenidae | 0.0000 | 1.0000 |
| WMCC | June | Ephemeroptera | Ephemeridae | 0.0112 | 1.0000 |
| WMCC | June | Ephemeroptera | Heptageniidae | 0.0000 | 1.0000 |
| WMCC | June | Hemiptera | Clastopteridae | 0.0000 | 1.0000 |
| WMCC | June | Hemiptera | Lygaeidae | 0.0000 | 1.0000 |
| WMCC | June | Hymenoptera | Apidae | 0.0000 | 1.0000 |
| WMCC | June | Hymenoptera | Braconidae | 0.0051 | 1.0000 |
| WMCC | June | Hymenoptera | Ichneumonidae | 0.0030 | 1.0000 |
| WMCC | June | Hymenoptera | Other | 0.0000 | 1.0000 |
| WMCC | June | Hymenoptera | Tenthredinidae | 0.0242 | 1.0000 |
| WMCC | June | Hymenoptera | Trigonalidae | 0.0000 | 1.0000 |
| WMCC | June | Lepidoptera | Crambidae | 0.0225 | 1.0000 |
| WMCC | June | Lepidoptera | Elachistidae | 0.0000 | 1.0000 |
| WMCC | June | Lepidoptera | Erebidae | 0.0000 | 1.0000 |
| WMCC | June | Lepidoptera | Gelechiidae | 0.0000 | 1.0000 |
| WMCC | June | Lepidoptera | Noctuidae | 0.0000 | 1.0000 |
| WMCC | June | Lepidoptera | Notodontidae | 0.0000 | 1.0000 |
| WMCC | June | Lepidoptera | Nymphalidae | 0.0000 | 1.0000 |
| WMCC | June | Lepidoptera | Pyralidae | 0.0121 | 1.0000 |
| WMCC | June | Lepidoptera | Tischeriidae | 0.0056 | 1.0000 |
| WMCC | June | Lepidoptera | Tortricidae | 0.0112 | 1.0000 |
| WMCC | June | Megaloptera | Corydalidae | 0.0000 | 1.0000 |
| WMCC | June | Neuroptera | Hemerobiidae | 0.0000 | 1.0000 |
| WMCC | June | Odonata | Libellulidae | 0.0000 | 1.0000 |

| | | | | | |
|---|---|---|---|---|---|
| WMCC | June | Orthoptera | Tettigoniidae | 0.0000 | 1.0000 |
| WMCC | June | Other | Other | 0.0424 | 1.0000 |
| WMCC | June | Plecoptera | Chloroperlidae | 0.0000 | 1.0000 |
| WMCC | June | Psocodea | Peripsocidae | 0.0000 | 1.0000 |
| WMCC | June | Sarcoptiformes | Oripodidae | 0.0000 | 1.0000 |
| WMCC | June | Trichoptera | Dipseudopsidae | 0.0000 | 1.0000 |
| WMCC | June | Trichoptera | Hydroptilidae | 0.0955 | 1.0000 |
| WMCC | June | Trichoptera | Leptoceridae | 0.0000 | 1.0000 |
| WMCC | June | Trombidiformes | Arrenuridae | 0.0000 | 1.0000 |
| WMCC | June | Trombidiformes | Aturidae | 0.0000 | 1.0000 |
| WMCC | June | Trombidiformes | Axonopsidae | 0.0000 | 1.0000 |
| WMCC | June | Trombidiformes | Eupodidae | 0.0000 | 1.0000 |
| WMCC | June | Trombidiformes | Feltriidae | 0.0000 | 1.0000 |
| WMCC | June | Trombidiformes | Hydryphantidae | 0.0000 | 1.0000 |
| WMCC | June | Trombidiformes | Lebertiidae | 0.0359 | 1.0000 |
| WMCC | June | Trombidiformes | Mideopsidae | 0.0000 | 1.0000 |
| WMCC | June | Trombidiformes | Pionidae | 0.0000 | 1.0000 |
| WMCC | June | Trombidiformes | Sperchontidae | 0.0000 | 1.0000 |
| WMCC | June | Trombidiformes | Tarsonemidae | 0.0000 | 1.0000 |
| WMCC | June | Trombidiformes | Torrenticolidae | 0.0000 | 1.0000 |
| WMCC | June | Trombidiformes | Unionicolidae | 0.0000 | 1.0000 |
| WMCC | July | Araneae | Agelenidae | 0.0000 | 1.0000 |
| WMCC | July | Araneae | Amaurobiidae | 0.0003 | 1.0000 |
| WMCC | July | Araneae | Anyphaenidae | 0.0024 | 1.0000 |
| WMCC | July | Araneae | Cicurinidae | 0.0000 | 1.0000 |
| WMCC | July | Araneae | Dictynidae | 0.0000 | 1.0000 |
| WMCC | July | Araneae | Gnaphosidae | 0.0003 | 1.0000 |
| WMCC | July | Araneae | Linyphiidae | 0.0000 | 1.0000 |
| WMCC | July | Araneae | Lycosidae | 0.0000 | 1.0000 |
| WMCC | July | Araneae | Phrurolithidae | 0.0000 | 1.0000 |
| WMCC | July | Araneae | Salticidae | 0.0000 | 1.0000 |

| | | | | | |
|---|---|---|---|---|---|
| WMCC | July | Araneae | Theridiidae | 0.0022 | 1.0000 |
| WMCC | July | Araneae | Thomisidae | 0.0005 | 1.0000 |
| WMCC | July | Blattodea | Ectobiidae | 0.0000 | 1.0000 |
| WMCC | July | Coleoptera | Buprestidae | 0.0000 | 1.0000 |
| WMCC | July | Coleoptera | Carabidae | 0.0003 | 1.0000 |
| WMCC | July | Coleoptera | Curculionidae | 0.0022 | 1.0000 |
| WMCC | July | Coleoptera | Dytiscidae | 0.0000 | 1.0000 |
| WMCC | July | Coleoptera | Lampyridae | 0.0000 | 1.0000 |
| WMCC | July | Coleoptera | Scarabaeidae | 0.0000 | 1.0000 |
| WMCC | July | Coleoptera | Scirtidae | 0.0000 | 1.0000 |
| WMCC | July | Diptera | Agromyzidae | 0.0094 | 1.0000 |
| WMCC | July | Diptera | Anisopodidae | 0.0013 | 1.0000 |
| WMCC | July | Diptera | Calliphoridae | 0.0000 | 1.0000 |
| WMCC | July | Diptera | Cecidomyiidae | 0.0161 | 1.0000 |
| WMCC | July | Diptera | Chaoboridae | 0.7132 | 1.0000 |
| WMCC | July | Diptera | Chironomidae | 0.0036 | 1.0000 |
| WMCC | July | Diptera | Chloropidae | 0.0019 | 1.0000 |
| WMCC | July | Diptera | Culicidae | 0.0000 | 1.0000 |
| WMCC | July | Diptera | Drosophilidae | 0.0000 | 1.0000 |
| WMCC | July | Diptera | Empididae | 0.0000 | 1.0000 |
| WMCC | July | Diptera | Hybotidae | 0.0000 | 1.0000 |
| WMCC | July | Diptera | Keroplatidae | 0.0000 | 1.0000 |
| WMCC | July | Diptera | Limoniidae | 0.0016 | 1.0000 |
| WMCC | July | Diptera | Muscidae | 0.0000 | 1.0000 |
| WMCC | July | Diptera | Mycetophilidae | 0.0003 | 1.0000 |
| WMCC | July | Diptera | Opomyzidae | 0.0000 | 1.0000 |
| WMCC | July | Diptera | Pallopteridae | 0.0009 | 1.0000 |
| WMCC | July | Diptera | Phoridae | 0.0000 | 1.0000 |
| WMCC | July | Diptera | Polleniidae | 0.0000 | 1.0000 |
| WMCC | July | Diptera | Psychodidae | 0.0000 | 1.0000 |
| WMCC | July | Diptera | Sarcophagidae | 0.0000 | 1.0000 |

| | | | | | |
|---|---|---|---|---|---|
| WMCC | July | Diptera | Sciaridae | 0.0022 | 1.0000 |
| WMCC | July | Diptera | Sphaeroceridae | 0.0000 | 1.0000 |
| WMCC | July | Diptera | Syrphidae | 0.0000 | 1.0000 |
| WMCC | July | Diptera | Tabanidae | 0.0000 | 1.0000 |
| WMCC | July | Diptera | Tachinidae | 0.0000 | 1.0000 |
| WMCC | July | Diptera | Tipulidae | 0.0000 | 1.0000 |
| WMCC | July | Ephemeroptera | Caenidae | 0.0000 | 1.0000 |
| WMCC | July | Ephemeroptera | Ephemeridae | 0.0016 | 1.0000 |
| WMCC | July | Ephemeroptera | Heptageniidae | 0.0000 | 1.0000 |
| WMCC | July | Hemiptera | Clastopteridae | 0.0000 | 1.0000 |
| WMCC | July | Hemiptera | Lygaeidae | 0.0002 | 1.0000 |
| WMCC | July | Hymenoptera | Apidae | 0.0000 | 1.0000 |
| WMCC | July | Hymenoptera | Braconidae | 0.0016 | 1.0000 |
| WMCC | July | Hymenoptera | Ichneumonidae | 0.0000 | 1.0000 |
| WMCC | July | Hymenoptera | Other | 0.0000 | 1.0000 |
| WMCC | July | Hymenoptera | Tenthredinidae | 0.0003 | 1.0000 |
| WMCC | July | Hymenoptera | Trigonalidae | 0.0004 | 1.0000 |
| WMCC | July | Lepidoptera | Crambidae | 0.0014 | 1.0000 |
| WMCC | July | Lepidoptera | Elachistidae | 0.0000 | 1.0000 |
| WMCC | July | Lepidoptera | Erebidae | 0.0000 | 1.0000 |
| WMCC | July | Lepidoptera | Gelechiidae | 0.0000 | 1.0000 |
| WMCC | July | Lepidoptera | Noctuidae | 0.0000 | 1.0000 |
| WMCC | July | Lepidoptera | Notodontidae | 0.0000 | 1.0000 |
| WMCC | July | Lepidoptera | Nymphalidae | 0.0000 | 1.0000 |
| WMCC | July | Lepidoptera | Pyralidae | 0.0000 | 1.0000 |
| WMCC | July | Lepidoptera | Tischeriidae | 0.0000 | 1.0000 |
| WMCC | July | Lepidoptera | Tortricidae | 0.0005 | 1.0000 |
| WMCC | July | Megaloptera | Corydalidae | 0.0000 | 1.0000 |
| WMCC | July | Neuroptera | Hemerobiidae | 0.0000 | 1.0000 |
| WMCC | July | Odonata | Libellulidae | 0.0000 | 1.0000 |
| WMCC | July | Orthoptera | Tettigoniidae | 0.0000 | 1.0000 |

| | | | | | |
|---|---|---|---|---|---|
| WMCC | July | Other | Other | 0.0031 | 1.0000 |
| WMCC | July | Plecoptera | Chloroperlidae | 0.0000 | 1.0000 |
| WMCC | July | Psocodea | Peripsocidae | 0.0144 | 1.0000 |
| WMCC | July | Sarcoptiformes | Oripodidae | 0.0000 | 1.0000 |
| WMCC | July | Trichoptera | Dipseudopsidae | 0.0003 | 1.0000 |
| WMCC | July | Trichoptera | Hydroptilidae | 0.0000 | 1.0000 |
| WMCC | July | Trichoptera | Leptoceridae | 0.2126 | 1.0000 |
| WMCC | July | Trombidiformes | Arrenuridae | 0.0000 | 1.0000 |
| WMCC | July | Trombidiformes | Aturidae | 0.0000 | 1.0000 |
| WMCC | July | Trombidiformes | Axonopsidae | 0.0000 | 1.0000 |
| WMCC | July | Trombidiformes | Eupodidae | 0.0000 | 1.0000 |
| WMCC | July | Trombidiformes | Feltriidae | 0.0000 | 1.0000 |
| WMCC | July | Trombidiformes | Hydryphantidae | 0.0000 | 1.0000 |
| WMCC | July | Trombidiformes | Lebertiidae | 0.0031 | 1.0000 |
| WMCC | July | Trombidiformes | Mideopsidae | 0.0000 | 1.0000 |
| WMCC | July | Trombidiformes | Pionidae | 0.0019 | 1.0000 |
| WMCC | July | Trombidiformes | Sperchontidae | 0.0000 | 1.0000 |
| WMCC | July | Trombidiformes | Tarsonemidae | 0.0000 | 1.0000 |
| WMCC | July | Trombidiformes | Torrenticolidae | 0.0000 | 1.0000 |
| WMCC | July | Trombidiformes | Unionicolidae | 0.0000 | 1.0000 |
| WMCC | August | Araneae | Agelenidae | 0.0000 | 1.0000 |
| WMCC | August | Araneae | Amaurobiidae | 0.0000 | 1.0000 |
| WMCC | August | Araneae | Anyphaenidae | 0.0000 | 1.0000 |
| WMCC | August | Araneae | Cicurinidae | 0.0000 | 1.0000 |
| WMCC | August | Araneae | Dictynidae | 0.0000 | 1.0000 |
| WMCC | August | Araneae | Gnaphosidae | 0.0000 | 1.0000 |
| WMCC | August | Araneae | Linyphiidae | 0.0000 | 1.0000 |
| WMCC | August | Araneae | Lycosidae | 0.0000 | 1.0000 |
| WMCC | August | Araneae | Phrurolithidae | 0.0000 | 1.0000 |
| WMCC | August | Araneae | Salticidae | 0.0000 | 1.0000 |
| WMCC | August | Araneae | Theridiidae | 0.0000 | 1.0000 |

| | | | | | |
|---|---|---|---|---|---|
| WMCC | August | Araneae | Thomisidae | 0.0000 | 1.0000 |
| WMCC | August | Blattodea | Ectobiidae | 0.0000 | 1.0000 |
| WMCC | August | Coleoptera | Buprestidae | 0.0000 | 1.0000 |
| WMCC | August | Coleoptera | Carabidae | 0.0000 | 1.0000 |
| WMCC | August | Coleoptera | Curculionidae | 0.0000 | 1.0000 |
| WMCC | August | Coleoptera | Dytiscidae | 0.0000 | 1.0000 |
| WMCC | August | Coleoptera | Lampyridae | 0.0000 | 1.0000 |
| WMCC | August | Coleoptera | Scarabaeidae | 0.0000 | 1.0000 |
| WMCC | August | Coleoptera | Scirtidae | 0.0000 | 1.0000 |
| WMCC | August | Diptera | Agromyzidae | 0.0000 | 1.0000 |
| WMCC | August | Diptera | Anisopodidae | 0.0000 | 1.0000 |
| WMCC | August | Diptera | Calliphoridae | 0.0000 | 1.0000 |
| WMCC | August | Diptera | Cecidomyiidae | 0.1126 | 1.0000 |
| WMCC | August | Diptera | Chaoboridae | 0.2185 | 1.0000 |
| WMCC | August | Diptera | Chironomidae | 0.0012 | 1.0000 |
| WMCC | August | Diptera | Chloropidae | 0.1220 | 1.0000 |
| WMCC | August | Diptera | Culicidae | 0.0000 | 1.0000 |
| WMCC | August | Diptera | Drosophilidae | 0.0000 | 1.0000 |
| WMCC | August | Diptera | Empididae | 0.0000 | 1.0000 |
| WMCC | August | Diptera | Hybotidae | 0.0000 | 1.0000 |
| WMCC | August | Diptera | Keroplatidae | 0.0000 | 1.0000 |
| WMCC | August | Diptera | Limoniidae | 0.0000 | 1.0000 |
| WMCC | August | Diptera | Muscidae | 0.0000 | 1.0000 |
| WMCC | August | Diptera | Mycetophilidae | 0.0000 | 1.0000 |
| WMCC | August | Diptera | Opomyzidae | 0.0000 | 1.0000 |
| WMCC | August | Diptera | Pallopteridae | 0.0000 | 1.0000 |
| WMCC | August | Diptera | Phoridae | 0.0000 | 1.0000 |
| WMCC | August | Diptera | Polleniidae | 0.0000 | 1.0000 |
| WMCC | August | Diptera | Psychodidae | 0.0000 | 1.0000 |
| WMCC | August | Diptera | Sarcophagidae | 0.0000 | 1.0000 |
| WMCC | August | Diptera | Sciaridae | 0.0000 | 1.0000 |

| | | | | | |
|---|---|---|---|---|---|
| WMCC | August | Diptera | Sphaeroceridae | 0.0000 | 1.0000 |
| WMCC | August | Diptera | Syrphidae | 0.0000 | 1.0000 |
| WMCC | August | Diptera | Tabanidae | 0.0000 | 1.0000 |
| WMCC | August | Diptera | Tachinidae | 0.0000 | 1.0000 |
| WMCC | August | Diptera | Tipulidae | 0.0000 | 1.0000 |
| WMCC | August | Ephemeroptera | Caenidae | 0.0000 | 1.0000 |
| WMCC | August | Ephemeroptera | Ephemeridae | 0.0000 | 1.0000 |
| WMCC | August | Ephemeroptera | Heptageniidae | 0.0000 | 1.0000 |
| WMCC | August | Hemiptera | Clastopteridae | 0.0000 | 1.0000 |
| WMCC | August | Hemiptera | Lygaeidae | 0.0000 | 1.0000 |
| WMCC | August | Hymenoptera | Apidae | 0.0000 | 1.0000 |
| WMCC | August | Hymenoptera | Braconidae | 0.0000 | 1.0000 |
| WMCC | August | Hymenoptera | Ichneumonidae | 0.0000 | 1.0000 |
| WMCC | August | Hymenoptera | Other | 0.0002 | 1.0000 |
| WMCC | August | Hymenoptera | Tenthredinidae | 0.0000 | 1.0000 |
| WMCC | August | Hymenoptera | Trigonalidae | 0.0000 | 1.0000 |
| WMCC | August | Lepidoptera | Crambidae | 0.0000 | 1.0000 |
| WMCC | August | Lepidoptera | Elachistidae | 0.0000 | 1.0000 |
| WMCC | August | Lepidoptera | Erebidae | 0.0000 | 1.0000 |
| WMCC | August | Lepidoptera | Gelechiidae | 0.0000 | 1.0000 |
| WMCC | August | Lepidoptera | Noctuidae | 0.0000 | 1.0000 |
| WMCC | August | Lepidoptera | Notodontidae | 0.0000 | 1.0000 |
| WMCC | August | Lepidoptera | Nymphalidae | 0.0000 | 1.0000 |
| WMCC | August | Lepidoptera | Pyralidae | 0.0000 | 1.0000 |
| WMCC | August | Lepidoptera | Tischeriidae | 0.0000 | 1.0000 |
| WMCC | August | Lepidoptera | Tortricidae | 0.0006 | 1.0000 |
| WMCC | August | Megaloptera | Corydalidae | 0.0000 | 1.0000 |
| WMCC | August | Neuroptera | Hemerobiidae | 0.0000 | 1.0000 |
| WMCC | August | Odonata | Libellulidae | 0.0000 | 1.0000 |
| WMCC | August | Orthoptera | Tettigoniidae | 0.0000 | 1.0000 |
| WMCC | August | Other | Other | 0.0022 | 1.0000 |

| | | | | | |
|---|---|---|---|---|---|
| WMCC | August | Plecoptera | Chloroperlidae | 0.0000 | 1.0000 |
| WMCC | August | Psocodea | Peripsocidae | 0.0000 | 1.0000 |
| WMCC | August | Sarcoptiformes | Oripodidae | 0.0000 | 1.0000 |
| WMCC | August | Trichoptera | Dipseudopsidae | 0.0000 | 1.0000 |
| WMCC | August | Trichoptera | Hydroptilidae | 0.0000 | 1.0000 |
| WMCC | August | Trichoptera | Leptoceridae | 0.0000 | 1.0000 |
| WMCC | August | Trombidiformes | Arrenuridae | 0.0006 | 1.0000 |
| WMCC | August | Trombidiformes | Aturidae | 0.0061 | 1.0000 |
| WMCC | August | Trombidiformes | Axonopsidae | 0.0012 | 1.0000 |
| WMCC | August | Trombidiformes | Eupodidae | 0.0009 | 1.0000 |
| WMCC | August | Trombidiformes | Feltriidae | 0.0000 | 1.0000 |
| WMCC | August | Trombidiformes | Hydryphantidae | 0.0000 | 1.0000 |
| WMCC | August | Trombidiformes | Lebertiidae | 0.3915 | 1.0000 |
| WMCC | August | Trombidiformes | Mideopsidae | 0.0000 | 1.0000 |
| WMCC | August | Trombidiformes | Pionidae | 0.0000 | 1.0000 |
| WMCC | August | Trombidiformes | Sperchontidae | 0.1220 | 1.0000 |
| WMCC | August | Trombidiformes | Tarsonemidae | 0.0000 | 1.0000 |
| WMCC | August | Trombidiformes | Torrenticolidae | 0.0205 | 1.0000 |
| WMCC | August | Trombidiformes | Unionicolidae | 0.0000 | 1.0000 |
