## Supplementary material for "Microbial, dietary insect, and pathogen communities in fresh and decomposing guano of anthropic little brown bat (Myotis lucifugus) maternity colonies": Table S20

| barcode | well | sample ID | Combined Sample | Total reads | Total reads per combined sample | N50 (Raw) | Bat Alignment Rate (%) | Bat aln rate combined | Total Reads (after bat removal) | # Reads bat (removed) |
|---|---|---|---|---|---|---|---|---|---|---|
| barcode12 | D02 | 2507AshExt6W4aMetagenome | | 2,343,591.0 | | 2837.0 | 0.79 | | 2,325,042.00 | 18,549.0 |
| barcode14 | F02 | 2507AshExt6W4bMetagenome | | 4,182,608.0 | | 2921.0 | 0.96 | | 4,142,313.00 | 40,295.0 |
| barcode22 | F03 | 2507AshExt6W4cMetagenome | 2507AshExt6W4 | 2,738,664.0 | 9264863.0 | 2569.0 | 0.44 | 0.73 | 2,726,545.00 | 12,119.0 |
| barcode09 | A05 | 2507AshInt6W4aMetagenome | | 4,098,350.0 | | 621.0 | 50.39 | | 2,033,176.00 | 2,065,174.0 |
| barcode04 | D01 | 2507AshInt6W4cMetagenome | 2507AshInt6 | 1,456,039.0 | 5554389.0 | 591.0 | 72.08 | 61.24 | 406,545.00 | 1,049,494.0 |
| barcode07 | G04 | 2507Wmcc6W4aMetagenome | | 5,828,586.0 | | 1736.0 | 0.61 | | 5,792,908.00 | 35,678.0 |
| barcode02 | B01 | 2507Wmcc6W4bMetagenome | | 3,128,482.0 | | 2513.0 | 0.37 | | 3,116,772.00 | 11,710.0 |
| barcode06 | F04 | 2507Wmcc6W4cMetagenome | 2507Wmcc6W4 | 5,973,254.0 | 14930322.0 | 2352.0 | 0.62 | 0.53 | 5,936,325.00 | 36,929.0 |
| barcode01 | A04 | 2508AshExt6W8aMetagenome | | 9,101,291.0 | | 2190.0 | 0.53 | | 9,052,788.00 | 48,503.0 |
| barcode23 | G03 | 2508AshExt6W8bMetagenome | | 6,968,626.0 | | 1872.0 | 5.61 | | 6,577,343.00 | 391,283.0 |
| barcode11 | C02 | 2508AshExt6W8cMetagenome | 2508AshExt6W8 | 4,129,821.0 | 20199738.0 | 1493.0 | 0.38 | 2.17 | 4,114,109.00 | 15,712.0 |
| barcode03 | C01 | 2508AshInt6W8aMetagenome | | 791,670.0 | | 571.0 | 66.05 | | 268,734.00 | 522,936.0 |
| barcode07 | G01 | 2508AshInt6W8aMetagenome | | 733,896.0 | | 578.0 | 65.42 | | 253,768.00 | 480,128.0 |
| barcode20 | D06 | 2508AshInt6W8bMetagenome | | 2,637,041.0 | | 490.0 | 46.84 | | 1,401,827.00 | 1,235,214.0 |
| barcode22 | F06 | 2508AshInt6W8cMetagenome | 2508AshInt6W8 | 2,022,612.0 | 6185219.0 | 633.0 | 59.99 | 59.58 | 809,311.00 | 1,213,301.0 |
| barcode10 | B02 | 2508Wmcc6W8aMetagenome | | 3,058,193.0 | | 2992.0 | 0.41 | | 3,045,739.00 | 12,454.0 |
| barcode13 | E02 | 2508Wmcc6W8bMetagenome | | 2,535,088.0 | | 2865.0 | 0.50 | | 2,522,457.00 | 12,631.0 |
| barcode09 | A02 | 2508Wmcc6W8cMetagenome | 2508Wmcc6W8 | 4,607,855.0 | 10201136.0 | 3432.0 | 0.46 | 0.46 | 4,586,695.00 | 21,160.0 |
| barcode15 | G02 | 2509AshExt6W12aMetagenome | | 2,177,821.0 | | 2925.0 | 0.50 | | 2,166,922.00 | 10,899.0 |
| barcode24 | H03 | 2509AshExt6W12bMetagenome | | 2,822,762.0 | | 2480.0 | 0.43 | | 2,810,494.00 | 12,268.0 |
| barcode19 | C03 | 2509AshExt6W12cMetagenome | 2509AshExt6W12 | 2,006,489.0 | 7007072.0 | 3249.0 | 0.57 | 0.50 | 1,995,147.00 | 11,342.0 |
| barcode14 | F05 | 2509AshExt6W4aMetagenome | | 4,270,311.0 | | 672.0 | 2.15 | | 4,178,705.00 | 91,606.0 |
| barcode24 | H06 | 2509AshExt6W4bMetagenome | | 956,748.0 | | 1519.0 | 2.09 | | 936,767.00 | 19,981.0 |
| barcode10 | B05 | 2509AshExt6W4cMetagenome | 2509AshExt6W4 | 6,046,150.0 | 11273209.0 | 525.0 | 3.68 | 2.64 | 5,823,658.00 | 222,492.0 |
| barcode13 | E05 | 2509AshInt6W12aMetagenome | | 4,480,045.0 | | 434.0 | 28.63 | | 3,197,282.00 | 1,282,763.0 |
| barcode17 | A06 | 2509AshInt6W12bMetagenome | | 3,945,024.0 | | 451.0 | 61.56 | | 1,516,397.00 | 2,428,627.0 |
| barcode18 | B06 | 2509AshInt6W12cMetagenome | 2509AshInt6W12 | 3,958,255.0 | 12383324.0 | 434.0 | 65.51 | 51.90 | 1,365,270.00 | 2,592,985.0 |
| barcode16 | H05 | 2509AshInt6W4aMetagenome | | 2,945,006.0 | | 610.0 | 93.27 | | 198,342.00 | 2,746,664.0 |
| barcode11 | C05 | 2509AshInt6W4bMetagenome | | 4,443,792.0 | | 621.0 | 91.83 | | 363,140.00 | 4,080,652.0 |
| barcode20 | D03 | 2509Wmcc6W12aMetagenome | 2509AshInt6W4 | 3,057,227.0 | 10446025.0 | 2980.0 | 0.47 | 61.86 | 3,042,878.00 | 14,349.0 |
| barcode04 | D04 | 2509Wmcc6W12bMetagenome | | 6,941,849.0 | | 2214.0 | 0.62 | | 6,898,794.00 | 43,055.0 |
| barcode17 | A03 | 2509Wmcc6W12cMetagenome | | 3,100,267.0 | | 2934.0 | 0.48 | | 3,085,455.00 | 14,812.0 |
| barcode08 | H04 | 2509Wmcc6W4bMetagenome | 2509Wmcc6W12 | 4,910,593.0 | 14952709.0 | 2613.0 | 0.70 | 0.60 | 4,876,094.00 | 34,499.0 |
