## Supplementary material for "Microbial, dietary insect, and pathogen communities in fresh and decomposing guano of anthropic little brown bat (Myotis lucifugus) maternity colonies": Table S21

| Sample | Rename | N50 | Largest Contig (bp) | # Primary Contigs | # alt/dup contigs | # possibly circular contigs >= 1M | # contigs >= 1M | # contigs >= 100k | Total bases within assembly |
|---|---|---|---|---|---|---|---|---|---|
| 2507AshExt6W4 | 2506AshExt6W4 | 30763 | 4687867 | 37670 | 798 and 52 | 4 | 54 | 482 | 594242971 |
| 2507AshInt6W4 | 2506AshInt6W4 | 6254 | 244415 | 4256 | 45 and 3 | 0 | 0 | 4 | 22241673 |
| 2507Wmcc6W4 | 2506Wmcc6W4 | 37875 | 4904260 | 43532 | 2039 and 23 | 3 | 51 | 624 | 650300496 |
| 2508AshExt6W8 | 2506AshExt6W8 | 21298 | 4801812 | 36885 | 415 and 32 | 9 | 36 | 337 | 462734828 |
| 2508AshInt6W8 | 2506AshInt6W8 | 6345 | 114874 | 3275 | 4 and 16 | 0 | 0 | 1 | 16656912 |
| 2508Wmcc6W8 | 2506Wmcc6W8 | 27337 | 4807570 | 68668 | 881 and 36 | 4 | 75 | 932 | 1033709455 |
| 2509AshExt6W12 | 2506AshExt6W12 | 23776 | 8684598 | 36930 | 341 and 21 | 8 | 37 | 417 | 575011260 |
| 2509AshExt6W4 | 2508AshExt6W4 | 7280 | 456962 | 21805 | 161 and 5 | 0 | 0 | 41 | 128300257 |
| 2509AshInt6W12 | 2506AshInt6W12 | 3474 | 53401 | 2148 | 28 and 3 | 0 | 0 | 0 | 6893672 |
| 2509AshInt6W4 | 2508AshInt6W4 | 4592 | 17666 | 755 | 2 and 0 | 0 | 0 | 0 | 3206144 |
| 2509Wmcc6W12 | 2506Wmcc6W12 | 18814 | 4610326 | 75724 | 1645 and 31 | 9 | 51 | 721 | 983218969 |
| 2509Wmcc6W4b | 2508Wmcc6W4b | 26192 | 4696756 | 27028 | 306 and 4 | 4 | 29 | 284 | 443087801 |
