## Supplementary material for "Microbial, dietary insect, and pathogen communities in fresh and decomposing guano of anthropic little brown bat (Myotis lucifugus) maternity colonies": Table S22

| Name | Max Hit | 2507AshEt6W4 | 2507AshInt6W4 | 2507Wmcc6W4 | 2508AshEt6W8 | 2508AshInt6W8 | 2508Wmcc6W8 | 2509AshEt6W12 | 2509AshEt6W4 | 2509AshInt6W1 | 2509AshInt6W4 | 2509Wmcc6W12 | 2509Wmcc6W4 | 2510AshEtW8 | 2510WmccW8 | Lineage |
|---|---|---|---|---|---|---|---|---|---|---|---|---|---|---|---|---|
| Acadianvirus reprobate | 3 | 2 | | 2 | | | 2 | 3 | 2 | | | 2 | | 2 | 3 | Viruses>Duplodnaviria>Heunggongvirae>Uroviricota>Caudoviricetes>Bclasvirinae>Acadianvirus |
| Aguilavirus mEp213 | 2 | | | | 1 | 1 | | | | 1 | | | | | | Viruses>Duplodnaviria>Heunggongvirae>Uroviricota>Caudoviricetes>Aguilavirus |
| Alegriavirus av2B8 | 2 | | 1 | 1 | | 2 | | | | 1 | | 1 | | | | Viruses>Duplodnaviria>Heunggongvirae>Uroviricota>Caudoviricetes>Alegriavirus |
| Alphabaculovirus alterchrincludentis | 3 | | | 2 | | 1 | 1 | 1 | 1 | 1 | 1 | 3 | | 1 | 2 | Viruses>Naldaviricetes>Lefavirales>Baculoviridae>Alphabaculovirus |
| Alphabaculovirus cycundantis | 2 | 1 | | 1 | | 1 | 1 | 1 | 1 | | | 1 | | 1 | 2 | Viruses>Naldaviricetes>Lefavirales>Baculoviridae>Alphabaculovirus |
| Alphabaculovirus helarmigerae | 3 | 2 | 2 | 2 | | 2 | | 2 | | 3 | | 2 | | | 2 | Viruses>Naldaviricetes>Lefavirales>Baculoviridae>Alphabaculovirus |
| Alphabaculovirus oxochraceae | 2 | | 1 | 1 | 1 | 2 | 1 | 1 | | 1 | | | | | | Viruses>Naldaviricetes>Lefavirales>Baculoviridae>Alphabaculovirus |
| Amginevirus amohnition | 2 | | | 1 | | | | 1 | | | | 1 | | 1 | 2 | Viruses>Duplodnaviria>Heunggongvirae>Uroviricota>Caudoviricetes>Weiservirinae>Amginevirus |
| Appavirus appa | 2 | 1 | 1 | 1 | 1 | 1 | 2 | 1 | 1 | 2 | | 1 | 1 | 1 | 1 | Viruses>Duplodnaviria>Heunggongvirae>Uroviricota>Caudoviricetes>Appavirus |
| Ascovirus TnAV2a | 3 | | | | | 1 | | 2 | | 2 | | 3 | | 3 | 1 | Viruses>Varidnaviria>Bamfordvirae>Nucleocytoviricota>Megaviricetes>Pimascovirales>Ascoviridae>Ascovirus |
| Attisvirus attis | 3 | 2 | | 2 | 1 | 1 | | 3 | 2 | | | 3 | | 2 | | Viruses>Duplodnaviria>Heunggongvirae>Uroviricota>Caudoviricetes>Attisvirus |
| Attisvirus lamberg | 2 | | | | 1 | | 2 | 1 | | | | 1 | | 1 | | Viruses>Duplodnaviria>Heunggongvirae>Uroviricota>Caudoviricetes>Attisvirus |
| Bat mastadenovirus C | 3 | | 1 | 2 | | | | 1 | | 1 | | | | 3 | | Viruses>Varidnaviria>Bamfordvirae>Preplasmiviricota>Tectiliviricetes>Rowavirales>Adenoviridae>Mastadenovirus |
| Betabaculovirus cypomonellae | 2 | 1 | | 1 | 1 | 1 | 1 | 1 | 1 | 1 | 1 | 1 | | 1 | 2 | Viruses>Naldaviricetes>Lefavirales>Baculoviridae>Betabaculovirus |
| Betanudivirus hezeae | 3 | 2 | | 2 | 1 | | 2 | 3 | | 2 | | 1 | | 3 | 3 | Viruses>Naldaviricetes>Lefavirales>Nudiviridae>Betanudivirus |
| Biavirus raunefjordenense | 2 | 1 | 1 | 1 | 1 | 2 | 1 | 2 | 1 | 1 | 1 | 1 | 1 | 1 | 1 | Viruses>Varidnaviria>Bamfordvirae>Nucleocytoviricota>Megaviricetes>Imitervirales>Schizomimiviridae>Biavirus |
| Bievrevirus bv4A7 | 2 | | | | | 2 | | | | 1 | | | | | | Viruses>Duplodnaviria>Heunggongvirae>Uroviricota>Caudoviricetes>Bievrevirus |
| Bixzunavirus Bxz1 | 4 | | 1 | | 1 | 1 | 1 | 4 | 1 | | | 4 | | | 2 | Viruses>Duplodnaviria>Heunggongvirae>Uroviricota>Caudoviricetes>Ceeclamvirinae>Bixzunavirus |
| Canine mastadenovirus A | 3 | | 1 | | | 3 | | | 1 | 3 | | | | | | Viruses>Varidnaviria>Bamfordvirae>Preplasmiviricota>Tectiliviricetes>Rowavirales>Adenoviridae>Mastadenovirus |
| Corndogvirus firecracker | 3 | | | 3 | | | 2 | 1 | | 2 | | 2 | | 2 | | Viruses>Duplodnaviria>Heunggongvirae>Uroviricota>Caudoviricetes>Corndogvirus |
| Cotia virus | 2 | | 1 | 1 | 1 | 2 | 1 | | 1 | 1 | | 1 | | 1 | | Viruses>Varidnaviria>Bamfordvirae>Nucleocytoviricota>Pokkesviricetes>Chitovirales>Poxviridae>Chordopoxvirinae>Oryzopoxvirus |
| Cotonvirus japonicum | 2 | 1 | 1 | 1 | 1 | 1 | 1 | 1 | 1 | 1 | 1 | 1 | 1 | 2 | 1 | Viruses>Varidnaviria>Bamfordvirae>Nucleocytoviricota>Megaviricetes>Imitervirales>Mimiviridae>Megamimivirinae>Cotonvirus |
| Demosthenesvirus demosthenes | 2 | 1 | | 2 | | 1 | 1 | | | 1 | | 1 | 1 | | 1 | Viruses>Duplodnaviria>Heunggongvirae>Uroviricota>Caudoviricetes>Demosthenesvirus |
| Demosthenesvirus katyusha | 3 | 3 | | 3 | 2 | 2 | 2 | 2 | 1 | 2 | | 2 | 2 | 2 | 2 | Viruses>Duplodnaviria>Heunggongvirae>Uroviricota>Caudoviricetes>Demosthenesvirus |
| Derbicusvirus derbicus | 3 | 3 | 3 | 2 | 2 | | 2 | 2 | 2 | 2 | | 2 | | 2 | 2 | Viruses>Duplodnaviria>Heunggongvirae>Uroviricota>Caudoviricetes>Derbicusvirus |
| Donellivirus gee | 2 | 1 | 1 | 1 | 1 | 1 | | 2 | 1 | 1 | | 1 | | 1 | 2 | Viruses>Duplodnaviria>Heunggongvirae>Uroviricota>Caudoviricetes>Donellivirus |
| Fipvunavirus Fpv4 | 3 | | | 2 | | 2 | | 2 | 2 | 3 | | 2 | | | | Viruses>Duplodnaviria>Heunggongvirae>Uroviricota>Caudoviricetes>Fipvunavirus |
| Fletchervirus NCTC12673 | 3 | 2 | 3 | 2 | 2 | 2 | 2 | 3 | 2 | 3 | | 2 | 3 | | 3 | Viruses>Duplodnaviria>Heunggongvirae>Uroviricota>Caudoviricetes>Eucampyvirinae>Fletchervirus |
| Fromanvirus packman | 3 | 3 | | 3 | 3 | | 2 | 2 | | | | 2 | | 2 | 2 | Viruses>Duplodnaviria>Heunggongvirae>Uroviricota>Caudoviricetes>Fromanvirus |
| Gemsvirus gv5004652 | 2 | | 2 | | | 2 | | | | 1 | | | | | | Viruses>Duplodnaviria>Heunggongvirae>Uroviricota>Caudoviricetes>Peduoviridae>Gemsvirus |
| Glaedevirus gv2H10 | 2 | | 1 | | | 2 | | | | 2 | | | | | | Viruses>Duplodnaviria>Heunggongvirae>Uroviricota>Caudoviricetes>Glaedevirus |
| Gordtnkvirus gordtnk2 | 3 | | | 1 | | | | 3 | | | | 3 | | | 2 | Viruses>Duplodnaviria>Heunggongvirae>Uroviricota>Caudoviricetes>Gordtnkvirus |
| Human mastadenovirus B | 3 | | 2 | | | | 1 | | | | | | | | 3 | Viruses>Varidnaviria>Bamfordvirae>Preplasmiviricota>Tectiliviricetes>Rowavirales>Adenoviridae>Mastadenovirus |
| Human mastadenovirus C | 4 | 1 | | 4 | | | 4 | | 4 | 3 | | | | 4 | | Viruses>Varidnaviria>Bamfordvirae>Preplasmiviricota>Tectiliviricetes>Rowavirales>Adenoviridae>Mastadenovirus |
| Human mastadenovirus E | 3 | 2 | 3 | | 1 | | | 3 | 1 | 1 | | 1 | 1 | | 3 | Viruses>Varidnaviria>Bamfordvirae>Preplasmiviricota>Tectiliviricetes>Rowavirales>Adenoviridae>Mastadenovirus |
| Jacevirus jace | 2 | 1 | | 2 | 2 | | 1 | 1 | | | 1 | 1 | 1 | 1 | 1 | Viruses>Duplodnaviria>Heunggongvirae>Uroviricota>Caudoviricetes>Jacevirus |
| Japarudivirus SBRV1 | 2 | 1 | 1 | | | 1 | | 1 | 1 | 1 | | | | | | Viruses>Adnaviria>Zilligvirae>Taleaviricota>Tokiviricetes>Ligamenvirales>Rudiviridae>Japarudivirus |
| Jouyvirus ev207 | 2 | 1 | | | | 2 | | | | 1 | | | | | | Viruses>Duplodnaviria>Heunggongvirae>Uroviricota>Caudoviricetes>Jouyvirus |
| Jouyvirus jv1H12 | 2 | | 2 | | | 2 | | | | 2 | | | | | | Viruses>Duplodnaviria>Heunggongvirae>Uroviricota>Caudoviricetes>Jouyvirus |
| Kayvirus G1 | 4 | 3 | | 2 | | 2 | | 1 | 3 | 3 | 4 | | | 4 | 3 | Viruses>Duplodnaviria>Heunggongvirae>Uroviricota>Caudoviricetes>Herelleviridae>Twortvirinae>Kayvirus |
| Kochikohdavirus EF24C | 3 | 2 | 2 | | | 2 | | 3 | | 2 | | 2 | | | 2 | Viruses>Duplodnaviria>Heunggongvirae>Uroviricota>Caudoviricetes>Herelleviridae>Brockvirinae>Kochikohdavirus |
| Latrobevirus FNU1 | 2 | | 2 | | 1 | 1 | | 1 | 1 | 2 | | 1 | | 1 | | Viruses>Duplodnaviria>Heunggongvirae>Uroviricota>Caudoviricetes>Latrobevirus |
| Lessievirus bcep22 | 2 | 1 | | 2 | 1 | | 2 | 1 | 1 | | 2 | 1 | 2 | 2 | 2 | Viruses>Duplodnaviria>Heunggongvirae>Uroviricota>Caudoviricetes>Lessievirus |
| Lessievirus bcepil02 | 2 | | | 1 | | 1 | 1 | 1 | 1 | 1 | 2 | 1 | 2 | 2 | 2 | Viruses>Duplodnaviria>Heunggongvirae>Uroviricota>Caudoviricetes>Lessievirus |
| Lessievirus DC1 | 2 | 1 | | 1 | | | 1 | 1 | 1 | | 2 | 1 | 2 | 1 | 2 | Viruses>Duplodnaviria>Heunggongvirae>Uroviricota>Caudoviricetes>Lessievirus |
| Mardivirus columbidalpha1 | 3 | 2 | 2 | 2 | 2 | 2 | 2 | 2 | 2 | 3 | 2 | 2 | 2 | 2 | 2 | Viruses>Duplodnaviria>Heunggongvirae>Peploviricota>Herviviricetes>Herpesvirales>Orthoherpesviridae>Alphaherpesvirinae>Mardivirus |
| Marienburgvirus BP4795 | 2 | | 1 | | | 2 | | | | 1 | | | | | | Viruses>Duplodnaviria>Heunggongvirae>Uroviricota>Caudoviricetes>Marienburgvirus |
| Marienburgvirus JLK2012 | 2 | | 1 | | | 2 | | | | 1 | | | | | | Viruses>Duplodnaviria>Heunggongvirae>Uroviricota>Caudoviricetes>Marienburgvirus |
| Menderavirus IMESM1 | 3 | 3 | 2 | 1 | 2 | 1 | 1 | 2 | 2 | 1 | | | | 3 | | Viruses>Duplodnaviria>Heunggongvirae>Uroviricota>Caudoviricetes>Menderavirus |
| Microwolfvirus JHC117 | 3 | 3 | | 2 | | | 3 | 1 | | | | | | 2 | | Viruses>Duplodnaviria>Heunggongvirae>Uroviricota>Caudoviricetes>Microwolfvirus |
| Montyvirus monty | 3 | | | 2 | | | 2 | 1 | | | | | 3 | | 3 | Viruses>Duplodnaviria>Heunggongvirae>Uroviricota>Caudoviricetes>Montyvirus |
| Mooglevirus mordin | 3 | 1 | | 1 | | | 1 | | 3 | | | | | | | Viruses>Duplodnaviria>Heunggongvirae>Uroviricota>Caudoviricetes>Ounavirinae>Mooglevirus |

| | | | | | | | | | | | | | | | | |
|---|---|---|---|---|---|---|---|---|---|---|---|---|---|---|---|---|
| Mosigvirus 0157tp3 | 3 | 1 | | 1 | | | 1 | | | | | 1 | | | 3 | Viruses>Duplodnaviria>Heunggongvirae>Uroviricota>Caudoviricetes>Straboviridae>Tevenvirinae>Mosigvirus |
| Moumouvirus australiense | 2 | 1 | 1 | 1 | 1 | 2 | 1 | 1 | 1 | 1 | 1 | 1 | | 1 | 1 | Viruses>Varidnaviria>Bamfordvirae>Nucleocytoviricota>Megaviricetes>Imitervirales>Mimiviridae>Megamimivirinae>Moumouvirus |
| Muminvirus snusmum | 2 | | 1 | | | 1 | | 1 | | 2 | | | | 1 | | Viruses>Duplodnaviria>Heunggongvirae>Uroviricota>Caudoviricetes>Muminvirus |
| Muscavirus musdomesticae | 2 | | | 1 | | 1 | 1 | 2 | | 1 | | 1 | | | 1 | Viruses>Naldaviricetes>Lefavirales>Hytrosaviridae>Muscavirus |
| Nankokuvirus PAKP3 | 3 | 1 | 1 | | | 3 | | 2 | 1 | | | 3 | | | | Viruses>Duplodnaviria>Heunggongvirae>Uroviricota>Caudoviricetes>Nankokuvirus |
| Nesevirus ev243 | 2 | | 1 | | | 2 | | | | 1 | | | | | | Viruses>Duplodnaviria>Heunggongvirae>Uroviricota>Caudoviricetes>Nesevirus |
| Nesevirus nv2G7b | 2 | | 1 | | | 2 | | | | | | | | | | Viruses>Duplodnaviria>Heunggongvirae>Uroviricota>Caudoviricetes>Nesevirus |
| Oslovirus PA2 | 3 | | 3 | | | 1 | | | | 3 | | | | | | Viruses>Duplodnaviria>Heunggongvirae>Uroviricota>Caudoviricetes>Sepvirinae>Oslovirus |
| Otagovirus Psa374 | 8 | 2 | | | | 2 | 1 | 5 | 2 | | | 3 | 8 | 2 | 7 | Viruses>Duplodnaviria>Heunggongvirae>Uroviricota>Caudoviricetes>Otagovirus |
| Pacinivirus VCO139 | 3 | | | 3 | 2 | | | 1 | | | | 2 | | | 2 | Viruses>Duplodnaviria>Heunggongvirae>Uroviricota>Caudoviricetes>Schitoviridae>Pacinivirus |
| Pecentumvirus LMSP25 | 3 | 2 | | 2 | | 3 | | 1 | | 2 | | 2 | | 2 | | Viruses>Duplodnaviria>Heunggongvirae>Uroviricota>Caudoviricetes>Herelleviridae>Jasinkavirinae>Pecentumvirus |
| Peduovirus magyaro | 2 | | 2 | | 1 | 2 | | | | 1 | | | | | | Viruses>Duplodnaviria>Heunggongvirae>Uroviricota>Caudoviricetes>Peduoviridae>Peduovirus |
| Peduovirus P2 | 2 | | 2 | | | 2 | 1 | | | 2 | | | | 2 | | Viruses>Duplodnaviria>Heunggongvirae>Uroviricota>Caudoviricetes>Peduoviridae>Peduovirus |
| Peduovirus P37 | 2 | | 2 | 1 | | 2 | | | | 2 | | | | | | Viruses>Duplodnaviria>Heunggongvirae>Uroviricota>Caudoviricetes>Peduoviridae>Peduovirus |
| Peduovirus pv12474III | 2 | | 2 | | | 2 | | | | 1 | | | | | | Viruses>Duplodnaviria>Heunggongvirae>Uroviricota>Caudoviricetes>Peduoviridae>Peduovirus |
| Peduovirus R18C | 2 | | 1 | | 1 | 2 | | | 1 | 1 | | | | | | Viruses>Duplodnaviria>Heunggongvirae>Uroviricota>Caudoviricetes>Peduoviridae>Peduovirus |
| Peduovirus STYP1 | 2 | | 1 | | | 2 | | | | 1 | | | | | | Viruses>Duplodnaviria>Heunggongvirae>Uroviricota>Caudoviricetes>Peduoviridae>Peduovirus |
| Peduovirus YPM22 | 2 | | 2 | | | 1 | | | | 1 | | | | | | Viruses>Duplodnaviria>Heunggongvirae>Uroviricota>Caudoviricetes>Peduoviridae>Peduovirus |
| Peduovirus YPM50 | 2 | | 2 | | | 2 | | | | 1 | | | | | | Viruses>Duplodnaviria>Heunggongvirae>Uroviricota>Caudoviricetes>Peduoviridae>Peduovirus |
| Pegunavirus suffolk | 6 | 1 | | 6 | 1 | | 4 | | 1 | | | | | | | Viruses>Duplodnaviria>Heunggongvirae>Uroviricota>Caudoviricetes>Bclasvirinae>Pegunavirus |
| Plotvirus plot | 3 | 1 | | | | | 2 | 2 | | | | 3 | | | | Viruses>Duplodnaviria>Heunggongvirae>Uroviricota>Caudoviricetes>Dclasvirinae>Plotvirus |
| Pseudotevenvirus miller | 3 | 3 | | 2 | 3 | 2 | | 2 | 3 | 2 | | 2 | | 2 | | Viruses>Duplodnaviria>Heunggongvirae>Uroviricota>Caudoviricetes>Straboviridae>Pseudotevenvirus |
| Pseudotevenvirus RB16 | 2 | 2 | | 1 | 2 | | 1 | | 2 | 1 | | 1 | | | 1 | Viruses>Duplodnaviria>Heunggongvirae>Uroviricota>Caudoviricetes>Straboviridae>Pseudotevenvirus |
| Pseudotevenvirus RB43 | 2 | 2 | | 1 | 2 | | | 1 | 2 | | | 1 | | | | Viruses>Duplodnaviria>Heunggongvirae>Uroviricota>Caudoviricetes>Straboviridae>Pseudotevenvirus |
| Pulverervirus PFR1 | 3 | 2 | | 3 | 1 | | 2 | 2 | | | | 2 | | 3 | 1 | Viruses>Duplodnaviria>Heunggongvirae>Uroviricota>Caudoviricetes>Pulverervirus |
| Punavirus P1 | 3 | | 3 | | | 3 | 1 | | 1 | 3 | | | | | 1 | Viruses>Duplodnaviria>Heunggongvirae>Uroviricota>Caudoviricetes>Punavirus |
| Punavirus RCS47 | 2 | | 2 | | | 2 | 1 | | | 2 | | 1 | | 1 | 1 | Viruses>Duplodnaviria>Heunggongvirae>Uroviricota>Caudoviricetes>Punavirus |
| Radostvirus ev099 | 2 | | 1 | | | 2 | | 1 | 1 | | | | | | | Viruses>Duplodnaviria>Heunggongvirae>Uroviricota>Caudoviricetes>Radostvirus |
| Rosebushvirus godines | 3 | 2 | | 1 | | | 1 | 2 | 2 | 2 | | 3 | | 3 | 2 | Viruses>Duplodnaviria>Heunggongvirae>Uroviricota>Caudoviricetes>Bclasvirinae>Rosebushvirus |
| Roskildevirus cronus | 2 | 2 | | 1 | | 1 | | 1 | 1 | | | | | | 1 | Viruses>Duplodnaviria>Heunggongvirae>Uroviricota>Caudoviricetes>Straboviridae>Tevenvirinae>Roskildevirus |
| Samwavirus samW | 3 | | | 2 | 1 | | | 3 | | | | 3 | 1 | | 3 | Viruses>Duplodnaviria>Heunggongvirae>Uroviricota>Caudoviricetes>Samwavirus |
| Saphexavirus SAP6 | 2 | | 2 | | | 2 | | 1 | | 1 | 1 | | | | | Viruses>Duplodnaviria>Heunggongvirae>Uroviricota>Caudoviricetes>Saphexavirus |
| Simian mastadenovirus B | 3 | | 1 | 3 | | | 3 | 1 | | 1 | | 1 | | | | Viruses>Varidnaviria>Bamfordvirae>Preplasmiviricota>Tectiliviricetes>Rowavirales>Adenoviridae>Mastadenovirus |
| Soupsvirus soups | 3 | 3 | | 3 | 3 | | 1 | 3 | | 2 | | 2 | | 3 | 1 | Viruses>Duplodnaviria>Heunggongvirae>Uroviricota>Caudoviricetes>Soupsvirus |
| Soupsvirus strosahl | 3 | 3 | | 3 | 3 | | 3 | 3 | | | | 2 | | 1 | 3 | Viruses>Duplodnaviria>Heunggongvirae>Uroviricota>Caudoviricetes>Soupsvirus |
| Takahashivirus PBS1 | 2 | 2 | 2 | 2 | | 2 | 2 | 1 | | 2 | 2 | | 1 | 2 | 2 | Viruses>Duplodnaviria>Heunggongvirae>Uroviricota>Caudoviricetes>Takahashivirus |
| Tequatrovirus slur02 | 2 | | | | | | | | | | | | | | | Viruses>Duplodnaviria>Heunggongvirae>Uroviricota>Caudoviricetes>Straboviridae>Tevenvirinae>Tequatrovirus |
| Timquatrovirus zoeJ | 2 | 1 | | 1 | 1 | | 1 | 1 | 1 | | | 2 | | | 1 | Viruses>Duplodnaviria>Heunggongvirae>Uroviricota>Caudoviricetes>Weiservirinae>Timquatrovirus |
| Tinduovirus TIN2 | 2 | 1 | | 1 | 1 | 1 | | 1 | | 1 | | | 2 | 2 | 1 | Viruses>Duplodnaviria>Heunggongvirae>Uroviricota>Caudoviricetes>Tinduovirus |
| Tinduovirus TIN3 | 3 | 1 | | 3 | 1 | | 3 | | | | | 3 | 3 | 3 | 3 | Viruses>Duplodnaviria>Heunggongvirae>Uroviricota>Caudoviricetes>Tinduovirus |
| Wellingtonvirus wellington | 3 | 2 | | 2 | 1 | | 2 | 1 | 3 | | | 3 | | 2 | | Viruses>Duplodnaviria>Heunggongvirae>Uroviricota>Caudoviricetes>Wellingtonvirus |
| Winklervirus xtwenty | 2 | | | 2 | 1 | 1 | 1 | | | | 1 | 1 | | 1 | | Viruses>Duplodnaviria>Heunggongvirae>Uroviricota>Caudoviricetes>Straboviridae>Tevenvirinae>Winklervirus |
| Wroclawvirus PA5oct | 2 | 2 | 1 | 1 | 1 | 1 | | 1 | 1 | 1 | | | 1 | 1 | 1 | Viruses>Duplodnaviria>Heunggongvirae>Uroviricota>Caudoviricetes>Arenbergviridae>Wroclawvirus |
| Xuanwuvirus P88 | 2 | | | | | 2 | | | | 1 | | | | | | Viruses>Duplodnaviria>Heunggongvirae>Uroviricota>Caudoviricetes>Peduoviridae>Xuanwuvirus |
| Xuanwuvirus xv503458 | 2 | | 1 | | | 2 | | | | 1 | | | | | | Viruses>Duplodnaviria>Heunggongvirae>Uroviricota>Caudoviricetes>Peduoviridae>Xuanwuvirus |
| Xuanwuvirus xv520873 | 2 | | | | | 2 | | | | 2 | | | | | 1 | Viruses>Duplodnaviria>Heunggongvirae>Uroviricota>Caudoviricetes>Peduoviridae>Xuanwuvirus |
| Yihwangvirus BCPST | 2 | | 1 | | | 2 | 1 | | | 1 | | | | | | Viruses>Duplodnaviria>Heunggongvirae>Uroviricota>Caudoviricetes>Sejongvirinae>Yihwangvirus |
| Yonseivirus N137 | 4 | 3 | | 3 | 4 | | | 3 | 3 | | | | | 3 | | Viruses>Duplodnaviria>Heunggongvirae>Uroviricota>Caudoviricetes>Casjensviridae>Yonseivirus |
