## Supplementary material for "Microbial, dietary insect, and pathogen communities in fresh and decomposing guano of anthropic little brown bat (Myotis lucifugus) maternity colonies": Table S23

| Sample | n | Order | Relative_abundance |
|---|---|---|---|
| 2506AshExt6W4 | 6 | Imitervirales | 0.285714 |
| 2506AshExt6W4 | 1 | Magrovirales | 0.047619 |
| 2506AshExt6W4 | 2 | Ortervirales | 0.095238 |
| 2506AshExt6W4 | 2 | Piccovirales | 0.095238 |
| 2506AshExt6W4 | 1 | Pimascovirales | 0.047619 |
| 2506AshExt6W4 | 1 | Priklausovirales | 0.047619 |
| 2506AshExt6W4 | 8 | Tubulavirales | 0.380952 |
| 2506AshInt6W4 | 2 | Piccovirales | 0.666667 |
| 2506AshInt6W4 | 1 | Rowavirales | 0.333333 |
| 2506Wmcc6W4 | 1 | Chitovirales | 0.076923 |
| 2506Wmcc6W4 | 1 | Crassvirales | 0.076923 |
| 2506Wmcc6W4 | 1 | Imitervirales | 0.076923 |
| 2506Wmcc6W4 | 1 | Lautamovirales | 0.076923 |
| 2506Wmcc6W4 | 2 | Ortervirales | 0.153846 |
| 2506Wmcc6W4 | 6 | Tubulavirales | 0.461538 |
| 2506Wmcc6W4 | 1 | Vinavirales | 0.076923 |
| 2506AshExt6W8 | 3 | Algavirales | 0.25 |
| 2506AshExt6W8 | 1 | Chitovirales | 0.083333 |
| 2506AshExt6W8 | 1 | Halopanivirales | 0.083333 |
| 2506AshExt6W8 | 1 | Imitervirales | 0.083333 |
| 2506AshExt6W8 | 1 | Petitvirales | 0.083333 |
| 2506AshExt6W8 | 5 | Tubulavirales | 0.416667 |
| 2506AshInt6W8 | 1 | Algavirales | 1 |
| 2506Wmcc6W8 | 1 | Algavirales | 0.066667 |
| 2506Wmcc6W8 | 1 | Chitovirales | 0.066667 |
| 2506Wmcc6W8 | 2 | Crassvirales | 0.133333 |
| 2506Wmcc6W8 | 1 | Imitervirales | 0.066667 |
| 2506Wmcc6W8 | 2 | Ortervirales | 0.133333 |
| 2506Wmcc6W8 | 1 | Pimascovirales | 0.066667 |
| 2506Wmcc6W8 | 4 | Tubulavirales | 0.266667 |
| 2506Wmcc6W8 | 3 | Vinavirales | 0.2 |
| 2506AshExt6W12 | 3 | Algavirales | 0.142857 |
| 2506AshExt6W12 | 1 | Chitovirales | 0.047619 |
| 2506AshExt6W12 | 1 | Crassvirales | 0.047619 |
| 2506AshExt6W12 | 1 | Halopanivirales | 0.047619 |
| 2506AshExt6W12 | 2 | Hareavirales | 0.095238 |
| 2506AshExt6W12 | 1 | Kalamavirales | 0.047619 |
| 2506AshExt6W12 | 1 | Orthopolintovira | 0.047619 |
| 2506AshExt6W12 | 1 | Petitvirales | 0.047619 |
| 2506AshExt6W12 | 10 | Tubulavirales | 0.47619 |
| 2508AshExt6W4 | 1 | Halopanivirales | 0.166667 |
| 2508AshExt6W4 | 1 | Kalamavirales | 0.166667 |
| 2508AshExt6W4 | 1 | Priklausovirales | 0.166667 |
| 2508AshExt6W4 | 3 | Tubulavirales | 0.5 |
| 2508AshInt6W4 | 1 | Tubulavirales | 1 |
| 2506Wmcc6W12 | 1 | Chitovirales | 0.066667 |
| 2506Wmcc6W12 | 2 | Crassvirales | 0.133333 |
| 2506Wmcc6W12 | 4 | Imitervirales | 0.266667 |
| 2506Wmcc6W12 | 3 | Ortervirales | 0.2 |
| 2506Wmcc6W12 | 4 | Tubulavirales | 0.266667 |
| 2506Wmcc6W12 | 1 | Vinavirales | 0.066667 |
| 2508Wmcc6W4b | 3 | Algavirales | 0.375 |
| 2508Wmcc6W4b | 1 | Crassvirales | 0.125 |
| 2508Wmcc6W4b | 2 | Imitervirales | 0.25 |
| 2508Wmcc6W4b | 1 | Kalamavirales | 0.125 |
| 2508Wmcc6W4b | 1 | Tubulavirales | 0.125 |
