## Supplementary material for "Microbial, dietary insect, and pathogen communities in fresh and decomposing guano of anthropic little brown bat (Myotis lucifugus) maternity colonies": Table S24

| Site | Week Status | Genus | Mean Relative Abundance |
|---|---|---|---|
| ASHFExt | W4 | Acinetobacter | 39.5816667 |
| ASHFExt | Fresh | Clostridiaceae_unclassified | 34.6016667 |
| ASHFExt | W8 | Acinetobacter | 21.0533333 |
| ASHFExt | Fresh | Candidatus_Arthromitus | 20.3916667 |
| ASHFExt | W12 | Massilia | 16.17 |
| ASHFExt | W12 | Acinetobacter | 15.45 |
| ASHFExt | Fresh | Plesiomonas | 12.6708333 |
| ASHFExt | W8 | Massilia | 9.165 |
| ASHFExt | W4 | Massilia | 8.41666667 |
| ASHFExt | W12 | Nocardiaceae_unclassified | 8.32 |
| ASHFExt | W12 | Sphingobacterium | 7.34 |
| ASHFExt | W12 | Carnobacterium | 7.3 |
| ASHFExt | W4 | Pseudomonas | 7.13166667 |
| ASHFExt | Fresh | Enterobacteriaceae_unclassified | 7.12416667 |
| ASHFExt | W12 | Chryseobacterium | 6.80666667 |
| ASHFExt | W8 | Nocardiaceae_unclassified | 6.07666667 |
| ASHFExt | W4 | Enterobacteriaceae_unclassified | 5.74333333 |
| ASHFExt | W4 | Chryseobacterium | 5.33666667 |
| ASHFExt | W8 | Chryseobacterium | 5.03166667 |
| ASHFExt | W8 | Comamonadaceae_unclassified | 4.495 |
| ASHFExt | W12 | Micrococcaceae_unclassified | 4.40333333 |
| ASHFExt | W8 | Sphingobacterium | 4.35 |
| ASHFExt | W4 | Sphingobacterium | 4.10666667 |
| ASHFExt | Fresh | Bacillaceae_unclassified | 3.95 |
| ASHFExt | Fresh | Aeromonas | 3.72416667 |
| ASHFExt | W4 | Alkanindiges | 3.32666667 |
| ASHFExt | W8 | Enterobacteriaceae_unclassified | 3.29833333 |
| ASHFExt | Fresh | Enterococcus | 3.275 |

| | | | |
|---|---|---|---|
| ASHFExt | W8 | Carnobacterium | 2.81 |
| ASHFExt | W8 | Alkanindiges | 2.79 |
| ASHFExt | W12 | Planococcaceae_unclassified | 2.77 |
| ASHFExt | W12 | Enterobacteriaceae_unclassified | 2.75 |
| ASHFExt | W8 | Micrococcaceae_unclassified | 2.625 |
| ASHFExt | W8 | Pseudomonas | 2.60833333 |
| ASHFExt | W8 | Dietzia | 2.58 |
| ASHFExt | W4 | Planococcaceae_unclassified | 2.42 |
| ASHFExt | W12 | Alkanindiges | 2.26666667 |
| ASHFExt | W4 | Nocardiaceae_unclassified | 2.245 |
| ASHFExt | W4 | Enterobacterales_unclassified | 2.14333333 |
| ASHFExt | W12 | Dietzia | 2.14 |
| ASHFExt | Fresh | Hafnia-Obesumbacterium | 2.1375 |
| ASHFExt | W4 | Carnobacterium | 1.97166667 |
| ASHFExt | W8 | Kocuria | 1.95166667 |
| ASHFExt | W8 | Planococcaceae_unclassified | 1.77833333 |
| ASHFExt | W8 | Pedobacter | 1.72666667 |
| ASHFExt | W8 | Enterobacterales_unclassified | 1.65833333 |
| ASHFExt | W12 | Paenarthrobacter | 1.65666667 |
| ASHFExt | W12 | Pseudomonas | 1.56 |
| ASHFExt | W12 | Comamonadaceae_unclassified | 1.49666667 |
| ASHFExt | W4 | Micrococcaceae_unclassified | 1.445 |
| ASHFExt | W4 | Lactococcus | 1.405 |
| ASHFExt | Fresh | Nosocomiicoccus | 1.35666667 |
| ASHFExt | W12 | Kocuria | 1.27666667 |
| ASHFExt | W8 | Deinococcus | 1.26833333 |
| ASHFExt | Fresh | Lactococcus | 1.2625 |
| ASHFExt | W4 | Comamonadaceae_unclassified | 1.22666667 |
| ASHFExt | W8 | Micrococcales_unclassified | 1.21666667 |

| | | | |
|---|---|---|---|
| ASHFExt | W12 | Pedobacter | 1.19333333 |
| ASHFExt | Fresh | Peptostreptococcaceae_unclassified | 1.085 |
| ASHFExt | W12 | Corallococcus | 1.04666667 |
| ASHFExt | W12 | Enterobacterales_unclassified | 0.96666667 |
| ASHFExt | W8 | Oxalobacteraceae_unclassified | 0.94666667 |
| ASHFExt | Fresh | Ureaplasma | 0.92333333 |
| ASHFExt | W12 | Psychrobacter | 0.91666667 |
| ASHFExt | W12 | Leucobacter | 0.87666667 |
| ASHFExt | W12 | Lactococcus | 0.83 |
| ASHFExt | W4 | Plesiomonas | 0.82166667 |
| ASHFExt | W8 | Edaphobaculum | 0.79833333 |
| ASHFExt | W4 | Enterococcus | 0.79833333 |
| ASHFExt | W8 | Cystobacter | 0.79 |
| ASHFExt | W8 | Leucobacter | 0.76333333 |
| ASHFExt | W4 | Psychrobacter | 0.75666667 |
| ASHFExt | W8 | Rhizobiaceae_unclassified | 0.755 |
| ASHFExt | W12 | Enterococcus | 0.74333333 |
| ASHFExt | W8 | Carnobacteriaceae_unclassified | 0.74166667 |
| ASHFExt | W4 | Peptostreptococcaceae_unclassified | 0.73166667 |
| ASHFExt | W8 | Psychrobacter | 0.72166667 |
| ASHFExt | W8 | Corallococcus | 0.715 |
| ASHFExt | W4 | Dietzia | 0.69166667 |
| ASHFExt | W8 | Paenibacillus | 0.68 |
| ASHFExt | Fresh | Enterobacterales_unclassified | 0.67166667 |
| ASHFExt | Fresh | Tyzzerella | 0.67083333 |
| ASHFExt | W12 | Plesiomonas | 0.67 |
| ASHFExt | W8 | Paenarthrobacter | 0.66666667 |
| ASHFExt | W4 | Oxalobacteraceae_unclassified | 0.64166667 |
| ASHFExt | W12 | Flavobacterium | 0.64 |

| | | | |
|---|---|---|---|
| ASHFExt | W8 | uncultured | 0.63333333 |
| ASHFExt | Fresh | Corynebacterium | 0.63083333 |
| ASHFExt | W4 | Carnobacteriaceae_unclassified | 0.62333333 |
| ASHFExt | W4 | Leucobacter | 0.605 |
| ASHFExt | Fresh | Diplorickettsiaceae_unclassified | 0.59416667 |
| ASHFExt | W8 | Stenotrophomonas | 0.58666667 |
| ASHFExt | W8 | Flavobacterium | 0.57166667 |
| ASHFExt | W8 | Paracoccus | 0.56833333 |
| ASHFExt | W4 | Kocuria | 0.52833333 |
| ASHFExt | W12 | Micrococcales_unclassified | 0.48666667 |
| ASHFExt | W8 | Chitinophagaceae_unclassified | 0.48666667 |
| ASHFExt | W12 | Rhodococcus | 0.46666667 |
| ASHFExt | W8 | Adhaeribacter | 0.43166667 |
| ASHFExt | W4 | Pedobacter | 0.42833333 |
| ASHFExt | W8 | Rhodococcus | 0.42666667 |
| ASHFExt | W8 | Dyadobacter | 0.425 |
| ASHFExt | W8 | Rhodobacteraceae_unclassified | 0.42333333 |
| ASHFExt | W8 | Devosiaceae_unclassified | 0.4 |
| ASHFExt | W8 | Roseomonas | 0.38833333 |
| ASHFExt | W4 | Hafnia-Obesumbacterium | 0.38666667 |
| ASHFExt | W12 | Deinococcus | 0.37333333 |
| ASHFExt | W12 | Curtobacterium | 0.36666667 |
| ASHFExt | W8 | Enterococcus | 0.35833333 |
| ASHFExt | Fresh | Rickettsiella | 0.35333333 |
| ASHFExt | Fresh | Bacteria_unclassified | 0.34833333 |
| ASHFExt | W12 | Sphingomonas | 0.34333333 |
| ASHFExt | W12 | Stenotrophomonas | 0.33333333 |
| ASHFExt | W4 | Rhodococcus | 0.32166667 |
| ASHFExt | W8 | Abditibacterium | 0.31666667 |

| | | | |
|---|---|---|---|
| ASHFExt | W8 | Curtobacterium | 0.31666667 |
| ASHFExt | W12 | Oxalobacteraceae_unclassified | 0.31666667 |
| ASHFExt | W8 | Caulobacteraceae_unclassified | 0.315 |
| ASHFExt | W8 | Sphingomonas | 0.31166667 |
| ASHFExt | W8 | Devosia | 0.29666667 |
| ASHFExt | W4 | Exiguobacterium | 0.29 |
| ASHFExt | W4 | Candidatus_Arthromitus | 0.28833333 |
| ASHFExt | W4 | Paenarthrobacter | 0.28 |
| ASHFExt | W8 | Alcaligenaceae_unclassified | 0.27833333 |
| ASHFExt | W4 | Rummeliibacillus | 0.27833333 |
| ASHFExt | W12 | Dyadobacter | 0.27666667 |
| ASHFExt | W8 | Chthoniobacter | 0.26833333 |
| ASHFExt | W12 | Rhizobiaceae_unclassified | 0.26666667 |
| ASHFExt | W8 | Fluviicola | 0.265 |
| ASHFExt | Fresh | Corynebacteriales_unclassified | 0.26333333 |
| ASHFExt | Fresh | Cetobacterium | 0.26333333 |
| ASHFExt | W12 | Roseomonas | 0.26 |
| ASHFExt | Fresh | Synergistaceae_unclassified | 0.25916667 |
| ASHFExt | W8 | Hymenobacter | 0.255 |
| ASHFExt | W8 | Arcticibacter | 0.25166667 |
| ASHFExt | W4 | Paracoccus | 0.25166667 |
| ASHFExt | W8 | Brachybacterium | 0.25 |
| ASHFExt | W4 | Deinococcus | 0.25 |
| ASHFExt | W8 | JG30-KF-CM45_ge | 0.245 |
| ASHFExt | W12 | Methylobacterium-Methylorubrum | 0.24333333 |
| ASHFExt | W8 | Aminobacter | 0.24 |
| ASHFExt | Fresh | Desulfovibrio | 0.23 |
| ASHFExt | W8 | Sphingobacteriaceae_unclassified | 0.23 |
| ASHFExt | W8 | Blastococcus | 0.22666667 |

| | | | |
|---|---|---|---|
| ASHFExt | W12 | Exiguobacterium | 0.22666667 |
| ASHFExt | Fresh | Fusobacterium | 0.22333333 |
| ASHFExt | W12 | Aminobacter | 0.22333333 |
| ASHFExt | W12 | Edaphobaculum | 0.22333333 |
| ASHFExt | W4 | Stenotrophomonas | 0.22333333 |
| ASHFExt | Fresh | Clostridium_sensu_stricto_1 | 0.21833333 |
| ASHFExt | W12 | Brachybacterium | 0.21666667 |
| ASHFExt | W8 | Microvirga | 0.21 |
| ASHFExt | W12 | Kineococcus | 0.21 |
| ASHFExt | W12 | Hymenobacter | 0.20666667 |
| ASHFExt | Fresh | Rickettsia | 0.205 |
| ASHFExt | W12 | Novosphingobium | 0.20333333 |
| ASHFExt | W12 | Paenibacillus | 0.20333333 |
| ASHFExt | W4 | Paenibacillus | 0.20166667 |
| ASHFExt | W8 | Kineococcus | 0.19333333 |
| ASHFExt | Fresh | Salinisphaera | 0.19 |
| ASHFExt | Fresh | Atopostipes | 0.18916667 |
| ASHFExt | W8 | Rummeliibacillus | 0.18666667 |
| ASHFExt | W4 | Moraxellaceae_unclassified | 0.18333333 |
| ASHFExt | Fresh | Mycoplasma | 0.1825 |
| ASHFExt | W4 | Hymenobacter | 0.18166667 |
| ASHFExt | Fresh | Alcaligenaceae_ge | 0.18 |
| ASHFExt | W8 | Exiguobacterium | 0.175 |
| ASHFExt | W8 | Nocardioides | 0.17333333 |
| ASHFExt | W12 | Paracoccus | 0.17333333 |
| ASHFExt | W12 | Bacillales_unclassified | 0.17 |
| ASHFExt | W4 | Curtobacterium | 0.16833333 |
| ASHFExt | W8 | Leifsonia | 0.16333333 |
| ASHFExt | W8 | Aeromicrobium | 0.16 |

| | | | |
|---|---|---|---|
| ASHFExt | Fresh | Bacteroidia_unclassified | 0.15916667 |
| ASHFExt | W8 | Methylobacterium-Methylorubrum | 0.15833333 |
| ASHFExt | W8 | Sericytochromatia_ge | 0.15666667 |
| ASHFExt | W12 | Arcticibacter | 0.15 |
| ASHFExt | W4 | Edaphobaculum | 0.15 |
| ASHFExt | W12 | Leifsonia | 0.14333333 |
| ASHFExt | W12 | Microvirga | 0.14333333 |
| ASHFExt | W4 | Dyadobacter | 0.14 |
| ASHFExt | W8 | Pontibacter | 0.14 |
| ASHFExt | W4 | uncultured | 0.13833333 |
| ASHFExt | Fresh | Paraclostridium | 0.13833333 |
| ASHFExt | W12 | Blastococcus | 0.13666667 |
| ASHFExt | W12 | Moraxellaceae_unclassified | 0.13666667 |
| ASHFExt | W12 | Sphingomonadaceae_unclassified | 0.13666667 |
| ASHFExt | W4 | Adhaeribacter | 0.135 |
| ASHFExt | W8 | Sphingomonadaceae_unclassified | 0.12833333 |
| ASHFExt | W12 | Actinobacteria_unclassified | 0.12666667 |
| ASHFExt | W12 | Brevundimonas | 0.12666667 |
| ASHFExt | W4 | Roseomonas | 0.125 |
| ASHFExt | W12 | Actinoplanes | 0.12333333 |
| ASHFExt | W8 | Actinoplanes | 0.12166667 |
| ASHFExt | W4 | Rhodobacteraceae_unclassified | 0.12166667 |
| ASHFExt | W8 | Bacteriovoracaceae_unclassified | 0.11833333 |
| ASHFExt | W12 | Devosia | 0.11666667 |
| ASHFExt | W4 | Rhizobiaceae_unclassified | 0.11666667 |
| ASHFExt | W8 | Brevundimonas | 0.11166667 |
| ASHFExt | W8 | Lysinibacillus | 0.11 |
| ASHFExt | W8 | Moraxellaceae_unclassified | 0.10833333 |
| ASHFExt | Fresh | Micrococcaceae_unclassified | 0.10833333 |

| | | | |
|---|---|---|---|
| ASHFExt | W8 | Actinobacteria_unclassified | 0.10666667 |
| ASHFExt | W8 | Novosphingobium | 0.10666667 |
| ASHFExt | W4 | Sphingomonas | 0.105 |
| ASHFExt | W4 | Gammaproteobacteria_unclassified | 0.10333333 |
| ASHFExt | W12 | Alcaligenaceae_unclassified | 0.10333333 |
| ASHFExt | W8 | Bacillales_unclassified | 0.10166667 |
| ASHFExt | W8 | Shinella | 0.09833333 |
| ASHFExt | W12 | Devosiaceae_unclassified | 0.09666667 |
| ASHFExt | W12 | JG30-KF-CM45_ge | 0.09666667 |
| ASHFExt | W4 | Alcaligenaceae_unclassified | 0.095 |
| ASHFExt | W4 | Flavobacterium | 0.095 |
| ASHFExt | W8 | Candidatus_Arthromitus | 0.09333333 |
| ASHFExt | W8 | Mycobacterium | 0.09333333 |
| ASHFExt | Fresh | Lacticaseibacillus | 0.09083333 |
| ASHFExt | W12 | Bacilli_unclassified | 0.09 |
| ASHFExt | W12 | Mucilaginibacter | 0.09 |
| ASHFExt | W12 | Rhodobacteraceae_unclassified | 0.09 |
| ASHFExt | W12 | Sphingobacteriaceae_unclassified | 0.09 |
| ASHFExt | W12 | Adhaeribacter | 0.09 |
| ASHFExt | W12 | uncultured | 0.09 |
| ASHFExt | W8 | Bdellovibrio | 0.08833333 |
| ASHFExt | W8 | Peredibacter | 0.085 |
| ASHFExt | W4 | Micrococcales_unclassified | 0.085 |
| ASHFExt | W12 | Aeromicrobium | 0.08333333 |
| ASHFExt | W12 | Micromonosporaceae_unclassified | 0.08 |
| ASHFExt | W8 | Aureimonas | 0.08 |
| ASHFExt | W4 | Staphylococcus | 0.07833333 |
| ASHFExt | W8 | Bosea | 0.07666667 |
| ASHFExt | W12 | Nocardioides | 0.07666667 |

| | | | |
|---|---|---|---|
| ASHFExt | W8 | Bacteriovorax | 0.075 |
| ASHFExt | W8 | Fimbriimonadaceae_ge | 0.075 |
| ASHFExt | W12 | Abditibacterium | 0.07333333 |
| ASHFExt | W12 | Carnobacteriaceae_unclassified | 0.07333333 |
| ASHFExt | W12 | Solibacillus | 0.07333333 |
| ASHFExt | W8 | Gammaproteobacteria_unclassified | 0.07166667 |
| ASHFExt | W12 | Chitinophagaceae_unclassified | 0.07 |
| ASHFExt | W4 | Orbus | 0.06833333 |
| ASHFExt | W8 | Marmoricola | 0.06666667 |
| ASHFExt | W8 | Nocardiaceae_ge | 0.06666667 |
| ASHFExt | W12 | Corynebacterium | 0.06666667 |
| ASHFExt | W12 | Allorhizobium-Neorhizobium-Pararhizobium- | 0.06333333 |
| ASHFExt | W4 | Bacillales_unclassified | 0.06333333 |
| ASHFExt | W8 | Gemmatimonadaceae_unclassified | 0.06333333 |
| ASHFExt | W4 | Sericytochromatia_ge | 0.06166667 |
| ASHFExt | W4 | Abditibacterium | 0.06166667 |
| ASHFExt | W4 | Clostridium_sensu_stricto_1 | 0.06166667 |
| ASHFExt | W4 | Leifsonia | 0.06166667 |
| ASHFExt | W4 | Methylobacterium-Methylorubrum | 0.06166667 |
| ASHFExt | W4 | Bacilli_unclassified | 0.06 |
| ASHFExt | W4 | Saccharibacillus | 0.05833333 |
| ASHFExt | W8 | Azospirillum | 0.05666667 |
| ASHFExt | W12 | Burkholderiales_unclassified | 0.05666667 |
| ASHFExt | W12 | Microbacteriaceae_unclassified | 0.05666667 |
| ASHFExt | W12 | Nocardiaceae_ge | 0.05666667 |
| ASHFExt | W4 | Brevundimonas | 0.055 |
| ASHFExt | Fresh | Helicobacter | 0.05416667 |
| ASHFExt | W12 | Aureimonas | 0.05333333 |
| ASHFExt | W4 | Devosia | 0.05333333 |

| | | | |
|---|---|---|---|
| ASHFExt | W12 | Gammaproteobacteria_unclassified | 0.05333333 |
| ASHFExt | Fresh | Leuconostoc | 0.05333333 |
| ASHFExt | W8 | Microbacteriaceae_unclassified | 0.05333333 |
| ASHFExt | W8 | Segetibacter | 0.05333333 |
| ASHFExt | W4 | Devosiaceae_unclassified | 0.05166667 |
| ASHFExt | W8 | Larkinella | 0.05166667 |
| ASHFExt | W8 | Bacteria_unclassified | 0.05 |
| ASHFExt | W12 | Bacillaceae_unclassified | 0.05 |
| ASHFExt | Fresh | Bartonella | 0.05 |
| ASHFExt | W12 | Rummeliibacillus | 0.05 |
| ASHFExt | W8 | Staphylococcus | 0.05 |
| ASHFExt | W4 | Lysinibacillus | 0.04833333 |
| ASHFExt | W8 | Iamia | 0.04833333 |
| ASHFExt | Fresh | uncultured_ge | 0.0475 |
| ASHFExt | Fresh | Alkanindiges | 0.04666667 |
| ASHFExt | W8 | Bacilli_unclassified | 0.04666667 |
| ASHFExt | W4 | Microvirga | 0.04666667 |
| ASHFExt | W8 | WD2101_soil_group_ge | 0.04666667 |
| ASHFExt | W8 | Georgenia | 0.045 |
| ASHFExt | W4 | Lactobacillales_unclassified | 0.045 |
| ASHFExt | W4 | Pontibacter | 0.045 |
| ASHFExt | W8 | Brevibacterium | 0.04333333 |
| ASHFExt | W12 | Cellvibrio | 0.04333333 |
| ASHFExt | W12 | Modestobacter | 0.04333333 |
| ASHFExt | W4 | Nocardiaceae_ge | 0.04166667 |
| ASHFExt | W4 | Lacticaseibacillus | 0.04166667 |
| ASHFExt | W8 | Thermomonas | 0.04166667 |
| ASHFExt | Fresh | Flavobacteriales_unclassified | 0.04083333 |
| ASHFExt | W8 | Phaselicystis | 0.04 |

| | | | |
|---|---|---|---|
| ASHFExt | W12 | Pontibacter | 0.04 |
| ASHFExt | W4 | Pseudomonadaceae_unclassified | 0.04 |
| ASHFExt | W12 | Sphingobium | 0.04 |
| ASHFExt | W12 | Cystobacter | 0.04 |
| ASHFExt | Fresh | Commensalibacter | 0.03916667 |
| ASHFExt | W4 | Aminobacter | 0.03833333 |
| ASHFExt | W4 | Kineococcus | 0.03833333 |
| ASHFExt | Fresh | Orbus | 0.03833333 |
| ASHFExt | W4 | Novosphingobium | 0.03833333 |
| ASHFExt | W4 | Sphingobacteriaceae_unclassified | 0.03666667 |
| ASHFExt | W8 | Gemmata | 0.03666667 |
| ASHFExt | W8 | Lysobacter | 0.03666667 |
| ASHFExt | W8 | Micromonosporaceae_unclassified | 0.03666667 |
| ASHFExt | W12 | Shinella | 0.03666667 |
| ASHFExt | Fresh | Staphylococcus | 0.03583333 |
| ASHFExt | Fresh | Aquabacterium | 0.035 |
| ASHFExt | W8 | Corynebacteriales_unclassified | 0.035 |
| ASHFExt | W8 | Gordonia | 0.035 |
| ASHFExt | W8 | Solirubrobacter | 0.035 |
| ASHFExt | W8 | Taibaiella | 0.035 |
| ASHFExt | Fresh | Bogoriella | 0.03333333 |
| ASHFExt | W12 | Ochrobactrum | 0.03333333 |
| ASHFExt | W4 | Solibacillus | 0.03333333 |
| ASHFExt | W4 | Brachybacterium | 0.03333333 |
| ASHFExt | W8 | Cellvibrio | 0.03333333 |
| ASHFExt | W12 | Chthoniobacter | 0.03333333 |
| ASHFExt | W12 | Marmoricola | 0.03333333 |
| ASHFExt | Fresh | Pectobacteriaceae_unclassified | 0.03333333 |
| ASHFExt | W8 | Burkholderiales_unclassified | 0.03166667 |

| | | | |
|---|---|---|---|
| ASHFExt | W8 | Mucilaginibacter | 0.03166667 |
| ASHFExt | W8 | Cohnella | 0.03166667 |
| ASHFExt | W8 | Rubrivirga | 0.03166667 |
| ASHFExt | W8 | Allorhizobium-Neorhizobium-Pararhizobium- | 0.03 |
| ASHFExt | W4 | Bacillaceae_unclassified | 0.03 |
| ASHFExt | W8 | Clostridiaceae_unclassified | 0.03 |
| ASHFExt | W12 | Intrasporangiaceae_unclassified | 0.03 |
| ASHFExt | W12 | Lactobacillales_unclassified | 0.03 |
| ASHFExt | W4 | Nocardioides | 0.03 |
| ASHFExt | W8 | Persicitalea | 0.03 |
| ASHFExt | W8 | Saccharibacillus | 0.03 |
| ASHFExt | Fresh | Firmicutes_unclassified | 0.02916667 |
| ASHFExt | W4 | Actinobacteria_unclassified | 0.02833333 |
| ASHFExt | W8 | Alphaproteobacteria_unclassified | 0.02833333 |
| ASHFExt | W8 | Belnapia | 0.02833333 |
| ASHFExt | W4 | Blastococcus | 0.02833333 |
| ASHFExt | W8 | Caulobacter | 0.02833333 |
| ASHFExt | W4 | Cystobacter | 0.02833333 |
| ASHFExt | W8 | Psychroglaciecola | 0.02833333 |
| ASHFExt | Fresh | Lachnospiraceae_unclassified | 0.02833333 |
| ASHFExt | W8 | Sphingobium | 0.02833333 |
| ASHFExt | W12 | Alcaligenaceae_ge | 0.02666667 |
| ASHFExt | W4 | Bacteria_unclassified | 0.02666667 |
| ASHFExt | W8 | Bacteroidia_unclassified | 0.02666667 |
| ASHFExt | W12 | Caulobacteraceae_unclassified | 0.02666667 |
| ASHFExt | Fresh | Micrococcales_unclassified | 0.02666667 |
| ASHFExt | Fresh | Paeniclostridium | 0.02583333 |
| ASHFExt | Fresh | Tomitella | 0.02583333 |
| ASHFExt | W8 | Lactobacillales_unclassified | 0.025 |

| | | | |
|---|---|---|---|
| ASHFExt | W8 | Bacillaceae_unclassified | 0.02333333 |
| ASHFExt | W12 | Belnapia | 0.02333333 |
| ASHFExt | W12 | Burkholderia-Caballeronia-Paraburkholderia | 0.02333333 |
| ASHFExt | Fresh | Lactobacillales_unclassified | 0.02333333 |
| ASHFExt | W12 | Myxococcus | 0.02333333 |
| ASHFExt | W8 | Oligoflexus | 0.02333333 |
| ASHFExt | W12 | Saccharibacillus | 0.02333333 |
| ASHFExt | Fresh | Sebaldella | 0.02333333 |
| ASHFExt | W4 | Sphingomonadaceae_unclassified | 0.02333333 |
| ASHFExt | W8 | Tepidisphaerales_unclassified | 0.02333333 |
| ASHFExt | Fresh | Asaia | 0.0225 |
| ASHFExt | W4 | Microbacteriaceae_unclassified | 0.02166667 |
| ASHFExt | W8 | Kaistia | 0.02166667 |
| ASHFExt | Fresh | Ruminococcaceae_unclassified | 0.02083333 |
| ASHFExt | Fresh | Erysipelatoclostridium | 0.02083333 |
| ASHFExt | W4 | Burkholderiales_unclassified | 0.02 |
| ASHFExt | W4 | Clostridiaceae_unclassified | 0.02 |
| ASHFExt | W8 | Flavitalea | 0.02 |
| ASHFExt | W8 | Rhodopseudomonas | 0.02 |
| ASHFExt | W12 | Skermanella | 0.02 |
| ASHFExt | W8 | Sumerlaea | 0.02 |
| ASHFExt | W8 | uncultured_ge | 0.02 |
| ASHFExt | W12 | Psychroglaciecola | 0.02 |
| ASHFExt | W8 | AKIW781_ge | 0.01833333 |
| ASHFExt | Fresh | Methylobacterium-Methylorubrum | 0.01833333 |
| ASHFExt | W4 | Persicitalea | 0.01833333 |
| ASHFExt | Fresh | 1174-901-12 | 0.01833333 |
| ASHFExt | W8 | Absconditabacteriales_(SR1)_ge | 0.01833333 |
| ASHFExt | W4 | Arcticibacter | 0.01833333 |

| | | | |
|---|---|---|---|
| ASHFExt | W8 | Fontibacillus | 0.01833333 |
| ASHFExt | W12 | Bacteria_unclassified | 0.01666667 |
| ASHFExt | W12 | Corynebacteriales_unclassified | 0.01666667 |
| ASHFExt | W4 | Georgenia | 0.01666667 |
| ASHFExt | W12 | Lysinibacillus | 0.01666667 |
| ASHFExt | W12 | Segetibacter | 0.01666667 |
| ASHFExt | W8 | Silvanigrellaceae_unclassified | 0.01666667 |
| ASHFExt | W12 | Thermomonas | 0.01666667 |
| ASHFExt | W12 | Yersiniaceae_unclassified | 0.01666667 |
| ASHFExt | W12 | Alphaproteobacteria_unclassified | 0.01666667 |
| ASHFExt | W12 | Azospirillum | 0.01666667 |
| ASHFExt | W12 | Bosea | 0.01666667 |
| ASHFExt | Fresh | Carnobacterium | 0.01666667 |
| ASHFExt | W12 | Cohnella | 0.01666667 |
| ASHFExt | W8 | Cryptosporangium | 0.01666667 |
| ASHFExt | W12 | Flavisolibacter | 0.01666667 |
| ASHFExt | W12 | Georgenia | 0.01666667 |
| ASHFExt | W12 | Lachnospiraceae_unclassified | 0.01666667 |
| ASHFExt | W4 | Phaselicystis | 0.01666667 |
| ASHFExt | W8 | Streptomyces | 0.01666667 |
| ASHFExt | Fresh | uncultured | 0.01666667 |
| ASHFExt | Fresh | Bacilli_unclassified | 0.01583333 |
| ASHFExt | Fresh | Vagococcus | 0.01583333 |
| ASHFExt | W8 | Camelimonas | 0.015 |
| ASHFExt | W4 | Chthoniobacter | 0.015 |
| ASHFExt | W4 | Corallococcus | 0.015 |
| ASHFExt | W8 | Isosphaeraceae_unclassified | 0.015 |
| ASHFExt | W8 | Jatrophihabitans | 0.015 |
| ASHFExt | Fresh | Microbacteriaceae_unclassified | 0.015 |

| | | | |
|---|---|---|---|
| ASHFExt | Fresh | Mycoplasmataceae_unclassified | 0.015 |
| ASHFExt | W8 | Nocardia | 0.015 |
| ASHFExt | W8 | Patulibacter | 0.015 |
| ASHFExt | W8 | Spirosoma | 0.015 |
| ASHFExt | W4 | Duganella | 0.015 |
| ASHFExt | W8 | Geminicoccus | 0.015 |
| ASHFExt | Fresh | Mycobacterium | 0.01416667 |
| ASHFExt | Fresh | Providencia | 0.01416667 |
| ASHFExt | Fresh | Flavobacteriaceae_unclassified | 0.01416667 |
| ASHFExt | W12 | 67-14_ge | 0.01333333 |
| ASHFExt | W4 | Actinoplanes | 0.01333333 |
| ASHFExt | W4 | Aeromicrobium | 0.01333333 |
| ASHFExt | W8 | Antricoccus | 0.01333333 |
| ASHFExt | W8 | Aquabacterium | 0.01333333 |
| ASHFExt | W12 | Beijerinckiaceae_unclassified | 0.01333333 |
| ASHFExt | W12 | Conexibacter | 0.01333333 |
| ASHFExt | W8 | Intrasporangiaceae_unclassified | 0.01333333 |
| ASHFExt | W8 | Lactococcus | 0.01333333 |
| ASHFExt | W12 | Larkinella | 0.01333333 |
| ASHFExt | W12 | Lysobacter | 0.01333333 |
| ASHFExt | W8 | Nakamurella | 0.01333333 |
| ASHFExt | W8 | Phenylobacterium | 0.01333333 |
| ASHFExt | W8 | Polyangiaceae_unclassified | 0.01333333 |
| ASHFExt | W8 | Pseudomonadaceae_unclassified | 0.01333333 |
| ASHFExt | W12 | Rhizobiales_unclassified | 0.01333333 |
| ASHFExt | W12 | Sericytochromatia_ge | 0.01333333 |
| ASHFExt | W4 | Shinella | 0.01333333 |
| ASHFExt | W4 | Sphingobium | 0.01333333 |
| ASHFExt | W12 | Staphylococcus | 0.01333333 |

| | | | |
|---|---|---|---|
| ASHFExt | W8 | Tundrisphaera | 0.01333333 |
| ASHFExt | W12 | Williamsia | 0.01333333 |
| ASHFExt | W12 | Cryptosporangium | 0.01333333 |
| ASHFExt | W12 | Mycobacterium | 0.01333333 |
| ASHFExt | Fresh | Rhodococcus | 0.01333333 |
| ASHFExt | Fresh | Morganella | 0.0125 |
| ASHFExt | Fresh | Paenalcaligenes | 0.0125 |
| ASHFExt | W4 | Allorhizobium-Neorhizobium-Pararhizobium- | 0.01166667 |
| ASHFExt | Fresh | Beijerinckiaceae_unclassified | 0.01166667 |
| ASHFExt | W4 | Chitinophagaceae_unclassified | 0.01166667 |
| ASHFExt | W8 | Geodermatophilaceae_unclassified | 0.01166667 |
| ASHFExt | W8 | Ochrobactrum | 0.01166667 |
| ASHFExt | Fresh | Rhodobacteraceae_unclassified | 0.01166667 |
| ASHFExt | W8 | Skermanella | 0.01166667 |
| ASHFExt | W4 | Alcaligenaceae_ge | 0.01166667 |
| ASHFExt | W8 | Alsobacter | 0.01166667 |
| ASHFExt | Fresh | Bacillales_unclassified | 0.01083333 |
| ASHFExt | Fresh | Sporosarcina | 0.01083333 |
| ASHFExt | W12 | Geodermatophilaceae_unclassified | 0.01 |
| ASHFExt | W12 | Solirubrobacter | 0.01 |
| ASHFExt | W8 | 0319-6G20_ge | 0.01 |
| ASHFExt | W8 | Acetobacteraceae_unclassified | 0.01 |
| ASHFExt | W12 | Actinomycetospora | 0.01 |
| ASHFExt | W8 | Alcaligenaceae_ge | 0.01 |
| ASHFExt | W12 | Alsobacter | 0.01 |
| ASHFExt | W4 | Bdellovibrio | 0.01 |
| ASHFExt | W8 | Beijerinckiaceae_unclassified | 0.01 |
| ASHFExt | W4 | Bosea | 0.01 |
| ASHFExt | W8 | Clostridium_sensu_stricto_1 | 0.01 |

| | | | |
|---|---|---|---|
| ASHFExt | W4 | Corynebacteriales_unclassified | 0.01 |
| ASHFExt | W12 | Geminicoccus | 0.01 |
| ASHFExt | W4 | JG30-KF-CM45_ge | 0.01 |
| ASHFExt | W8 | Lachnospiraceae_unclassified | 0.01 |
| ASHFExt | W4 | Larkinella | 0.01 |
| ASHFExt | W8 | Morganella | 0.01 |
| ASHFExt | W12 | Nakamurella | 0.01 |
| ASHFExt | W4 | Rickettsiaceae_unclassified | 0.01 |
| ASHFExt | W8 | Saccharimonadales_unclassified | 0.01 |
| ASHFExt | W4 | Segetibacter | 0.01 |
| ASHFExt | W8 | Solibacillus | 0.01 |
| ASHFExt | W12 | Jeotgalicoccus | 0.01 |
| ASHFExt | W12 | Paeniglutamicibacter | 0.01 |
| ASHFExt | Fresh | Propionibacteriales_unclassified | 0.01 |
| ASHFExt | W12 | Spirosoma | 0.01 |
| ASHFExt | W12 | Staphylococcaceae_unclassified | 0.01 |
| ASHFExt | Fresh | JG30-KF-CM45_ge | 0.00916667 |
| ASHFExt | W8 | 67-14_ge | 0.00833333 |
| ASHFExt | W8 | Blastocatella | 0.00833333 |
| ASHFExt | W8 | Burkholderia-Caballeronia-Paraburkholderia | 0.00833333 |
| ASHFExt | Fresh | Curtobacterium | 0.00833333 |
| ASHFExt | W8 | Ellin6055 | 0.00833333 |
| ASHFExt | W4 | Fimbriimonadaceae_ge | 0.00833333 |
| ASHFExt | W8 | Flavisolibacter | 0.00833333 |
| ASHFExt | Fresh | Iamia | 0.00833333 |
| ASHFExt | Fresh | Leucobacter | 0.00833333 |
| ASHFExt | W4 | Mycobacterium | 0.00833333 |
| ASHFExt | W8 | Rickettsiaceae_unclassified | 0.00833333 |
| ASHFExt | W8 | Staphylococcaceae_unclassified | 0.00833333 |

| | | | |
|---|---|---|---|
| ASHFExt | W8 | Tepidisphaera | 0.00833333 |
| ASHFExt | W8 | Verrucomicrobium | 0.00833333 |
| ASHFExt | W8 | Xanthobacteraceae_unclassified | 0.00833333 |
| ASHFExt | W8 | mle1-27_ge | 0.00833333 |
| ASHFExt | W4 | uncultured_ge | 0.00833333 |
| ASHFExt | W4 | Bacteriovoracaceae_unclassified | 0.00833333 |
| ASHFExt | W8 | Duganella | 0.00833333 |
| ASHFExt | W8 | Williamsia | 0.00833333 |
| ASHFExt | Fresh | Pragia | 0.0075 |
| ASHFExt | Fresh | Brevinema | 0.0075 |
| ASHFExt | W12 | AKIW781_ge | 0.00666667 |
| ASHFExt | Fresh | Actinomycetospora | 0.00666667 |
| ASHFExt | W12 | Altererythrobacter | 0.00666667 |
| ASHFExt | W4 | Antricoccus | 0.00666667 |
| ASHFExt | W12 | Arthrobacter | 0.00666667 |
| ASHFExt | W8 | Atopostipes | 0.00666667 |
| ASHFExt | W4 | Aureimonas | 0.00666667 |
| ASHFExt | W4 | Azospirillum | 0.00666667 |
| ASHFExt | W12 | Blastocatella | 0.00666667 |
| ASHFExt | W12 | Bradyrhizobium | 0.00666667 |
| ASHFExt | W12 | Brevibacterium | 0.00666667 |
| ASHFExt | W4 | Cohnella | 0.00666667 |
| ASHFExt | W4 | Corynebacterium | 0.00666667 |
| ASHFExt | W12 | Fimbriimonadaceae_unclassified | 0.00666667 |
| ASHFExt | W8 | Firmicutes_unclassified | 0.00666667 |
| ASHFExt | W12 | Fontibacillus | 0.00666667 |
| ASHFExt | W12 | Frankiales_unclassified | 0.00666667 |
| ASHFExt | W12 | Gemmatimonadaceae_unclassified | 0.00666667 |
| ASHFExt | W12 | Gemmatimonas | 0.00666667 |

| | | | |
|---|---|---|---|
| ASHFExt | W12 | Gemmatirosa | 0.00666667 |
| ASHFExt | W12 | Jatrophihabitans | 0.00666667 |
| ASHFExt | W8 | LD29 | 0.00666667 |
| ASHFExt | Fresh | Lactobacillaceae_unclassified | 0.00666667 |
| ASHFExt | W12 | Lautropia | 0.00666667 |
| ASHFExt | W12 | Luteolibacter | 0.00666667 |
| ASHFExt | W8 | Luteolibacter | 0.00666667 |
| ASHFExt | W4 | Lysobacter | 0.00666667 |
| ASHFExt | W8 | Paenibacillaceae_unclassified | 0.00666667 |
| ASHFExt | W8 | Paeniclostridium | 0.00666667 |
| ASHFExt | W12 | Pajaroellobacter | 0.00666667 |
| ASHFExt | W8 | Parcubacteria_unclassified | 0.00666667 |
| ASHFExt | W12 | Propionibacteriales_unclassified | 0.00666667 |
| ASHFExt | W12 | Pseudomonadaceae_unclassified | 0.00666667 |
| ASHFExt | W12 | Rickettsiaceae_unclassified | 0.00666667 |
| ASHFExt | W12 | Saccharimonadales_unclassified | 0.00666667 |
| ASHFExt | W8 | Solirubrobacteraceae_unclassified | 0.00666667 |
| ASHFExt | W12 | Sphingoaurantiacus | 0.00666667 |
| ASHFExt | W4 | Staphylococcaceae_unclassified | 0.00666667 |
| ASHFExt | W8 | Steroidobacter | 0.00666667 |
| ASHFExt | W4 | Sumerlaea | 0.00666667 |
| ASHFExt | W12 | Taibaiella | 0.00666667 |
| ASHFExt | W4 | Tepidisphaerales_unclassified | 0.00666667 |
| ASHFExt | W12 | WD2101_soil_group_ge | 0.00666667 |
| ASHFExt | Fresh | Dysgonomonas | 0.00666667 |
| ASHFExt | Fresh | Truepera | 0.00666667 |
| ASHFExt | Fresh | Aeromicrobium | 0.00583333 |
| ASHFExt | Fresh | Brevibacterium | 0.00583333 |
| ASHFExt | Fresh | Proteus | 0.00583333 |

| | | | |
|---|---|---|---|
| ASHFExt | W4 | 0319-6G20_ge | 0.005 |
| ASHFExt | W8 | A0839_ge | 0.005 |
| ASHFExt | W8 | Actinomycetospora | 0.005 |
| ASHFExt | W8 | Agromyces | 0.005 |
| ASHFExt | W8 | Anaeromyxobacter | 0.005 |
| ASHFExt | W8 | Bacteroides | 0.005 |
| ASHFExt | W4 | Beijerinckiaceae_unclassified | 0.005 |
| ASHFExt | W4 | Belnapia | 0.005 |
| ASHFExt | W4 | Caulobacteraceae_unclassified | 0.005 |
| ASHFExt | W4 | Conexibacter | 0.005 |
| ASHFExt | W8 | D05-2_ge | 0.005 |
| ASHFExt | W8 | Frankiales_unclassified | 0.005 |
| ASHFExt | W4 | Geminicoccus | 0.005 |
| ASHFExt | W4 | Gemmatimonadaceae_unclassified | 0.005 |
| ASHFExt | W4 | Intrasporangiaceae_unclassified | 0.005 |
| ASHFExt | W8 | Isosphaera | 0.005 |
| ASHFExt | W4 | Kineosporiaceae_unclassified | 0.005 |
| ASHFExt | W8 | Microbacterium | 0.005 |
| ASHFExt | W4 | Mucilaginibacter | 0.005 |
| ASHFExt | W8 | Myxococcaceae_unclassified | 0.005 |
| ASHFExt | W8 | Parapusillimonas | 0.005 |
| ASHFExt | W8 | Peptostreptococcaceae_unclassified | 0.005 |
| ASHFExt | W8 | Plesiomonas | 0.005 |
| ASHFExt | W4 | Psychroglaciecola | 0.005 |
| ASHFExt | W8 | Pusillimonas | 0.005 |
| ASHFExt | W8 | Rhizobiales_unclassified | 0.005 |
| ASHFExt | W8 | Rhodospirillales_unclassified | 0.005 |
| ASHFExt | W8 | SM2D12_ge | 0.005 |
| ASHFExt | W4 | Saccharimonadales_unclassified | 0.005 |

| | | | |
|---|---|---|---|
| ASHFExt | Fresh | Silvanigrella | 0.005 |
| ASHFExt | W8 | Sporosarcina | 0.005 |
| ASHFExt | W4 | Streptococcaceae_unclassified | 0.005 |
| ASHFExt | W8 | Weeksellaceae_unclassified | 0.005 |
| ASHFExt | Fresh | Pseudomonadales_unclassified | 0.005 |
| ASHFExt | Fresh | Raoultibacter | 0.005 |
| ASHFExt | Fresh | Streptococcaceae_unclassified | 0.005 |
| ASHFExt | Fresh | Clostridia_unclassified | 0.00416667 |
| ASHFExt | Fresh | Pusillimonas | 0.00416667 |
| ASHFExt | Fresh | Actinobacteria_unclassified | 0.00416667 |
| ASHFExt | Fresh | Alcaligenaceae_unclassified | 0.00416667 |
| ASHFExt | Fresh | Membranicola | 0.00416667 |
| ASHFExt | Fresh | Rickettsiaceae_unclassified | 0.00416667 |
| ASHFExt | Fresh | Liquorilactobacillus | 0.00416667 |
| ASHFExt | Fresh | Paracoccus | 0.00416667 |
| ASHFExt | W12 | 0319-6G20_ge | 0.00333333 |
| ASHFExt | W8 | A4b_ge | 0.00333333 |
| ASHFExt | W12 | Absconditabacteriales_(SR1)_ge | 0.00333333 |
| ASHFExt | W12 | Acidiphilium | 0.00333333 |
| ASHFExt | W8 | Acidiphilium | 0.00333333 |
| ASHFExt | W12 | Actinobacteriota_unclassified | 0.00333333 |
| ASHFExt | W12 | Actinocatenispora | 0.00333333 |
| ASHFExt | W12 | Actinocorallia | 0.00333333 |
| ASHFExt | W12 | Agathobacter | 0.00333333 |
| ASHFExt | W4 | Alphaproteobacteria_unclassified | 0.00333333 |
| ASHFExt | W4 | Alsobacter | 0.00333333 |
| ASHFExt | W4 | Altererythrobacter | 0.00333333 |
| ASHFExt | W8 | Altererythrobacter | 0.00333333 |
| ASHFExt | W12 | Anaerostipes | 0.00333333 |

| | | | |
|---|---|---|---|
| ASHFExt | W12 | Antricoccus | 0.00333333 |
| ASHFExt | W12 | Aquabacterium | 0.00333333 |
| ASHFExt | W12 | Armatimonadales_unclassified | 0.00333333 |
| ASHFExt | W4 | Arthrobacter | 0.00333333 |
| ASHFExt | W4 | Asaia | 0.00333333 |
| ASHFExt | W4 | Aurantisolimonas | 0.00333333 |
| ASHFExt | W4 | Brevibacterium | 0.00333333 |
| ASHFExt | W12 | Camelimonas | 0.00333333 |
| ASHFExt | W12 | Candidatus_Arthromitus | 0.00333333 |
| ASHFExt | W12 | Caulobacter | 0.00333333 |
| ASHFExt | W4 | Caulobacter | 0.00333333 |
| ASHFExt | W12 | Cetobacterium | 0.00333333 |
| ASHFExt | W12 | Chitiniphilus | 0.00333333 |
| ASHFExt | W4 | Chlamydiales_unclassified | 0.00333333 |
| ASHFExt | W8 | Chlamydiales_unclassified | 0.00333333 |
| ASHFExt | W8 | Chloroflexales_unclassified | 0.00333333 |
| ASHFExt | W12 | Clostridiaceae_unclassified | 0.00333333 |
| ASHFExt | W12 | Clostridium_sensu_stricto_1 | 0.00333333 |
| ASHFExt | Fresh | Comamonadaceae_unclassified | 0.00333333 |
| ASHFExt | W8 | Corynebacterium | 0.00333333 |
| ASHFExt | Fresh | Dietzia | 0.00333333 |
| ASHFExt | W12 | Duganella | 0.00333333 |
| ASHFExt | W8 | Entomoplasma | 0.00333333 |
| ASHFExt | W12 | Erysipelatoclostridium | 0.00333333 |
| ASHFExt | W12 | Ferruginibacter | 0.00333333 |
| ASHFExt | W8 | Ferruginibacter | 0.00333333 |
| ASHFExt | W12 | Fimbriimonadaceae_ge | 0.00333333 |
| ASHFExt | W12 | Flavitalea | 0.00333333 |
| ASHFExt | W12 | Flexivirga | 0.00333333 |

| | | | |
|---|---|---|---|
| ASHFExt | W8 | Flexivirga | 0.00333333 |
| ASHFExt | W4 | Frankiales_unclassified | 0.00333333 |
| ASHFExt | W12 | Gemmata | 0.00333333 |
| ASHFExt | W8 | Gemmataceae_unclassified | 0.00333333 |
| ASHFExt | W12 | Glutamicibacter | 0.00333333 |
| ASHFExt | W12 | Gordonia | 0.00333333 |
| ASHFExt | W4 | Gordonia | 0.00333333 |
| ASHFExt | W12 | Herbaspirillum | 0.00333333 |
| ASHFExt | W12 | Iamia | 0.00333333 |
| ASHFExt | W12 | Isosphaeraceae_unclassified | 0.00333333 |
| ASHFExt | W12 | JCM_18997 | 0.00333333 |
| ASHFExt | W4 | Janthinobacterium | 0.00333333 |
| ASHFExt | W4 | Jatrophihabitans | 0.00333333 |
| ASHFExt | W8 | Jeotgalicoccus | 0.00333333 |
| ASHFExt | W4 | Kineosporia | 0.00333333 |
| ASHFExt | W8 | Klenkia | 0.00333333 |
| ASHFExt | W12 | Kribbella | 0.00333333 |
| ASHFExt | W12 | Lachnospiraceae_NK4A136_group | 0.00333333 |
| ASHFExt | W4 | Leuconostoc | 0.00333333 |
| ASHFExt | W8 | Longimicrobiaceae_ge | 0.00333333 |
| ASHFExt | W12 | Luteibacter | 0.00333333 |
| ASHFExt | W12 | Luteimonas | 0.00333333 |
| ASHFExt | W12 | Microbacterium | 0.00333333 |
| ASHFExt | W4 | Microbacterium | 0.00333333 |
| ASHFExt | W12 | Microgenomatia_ge | 0.00333333 |
| ASHFExt | W8 | Microgenomatia_ge | 0.00333333 |
| ASHFExt | W4 | Micromonosporaceae_unclassified | 0.00333333 |
| ASHFExt | W12 | Microtrichales_unclassified | 0.00333333 |
| ASHFExt | W8 | Modestobacter | 0.00333333 |

| | | | |
|---|---|---|---|
| ASHFExt | W12 | Morganellaceae_unclassified | 0.00333333 |
| ASHFExt | W8 | Mycoplasmataceae_unclassified | 0.00333333 |
| ASHFExt | W12 | Myxococcaceae_unclassified | 0.00333333 |
| ASHFExt | Fresh | Nocardioides | 0.00333333 |
| ASHFExt | W12 | Nubsella | 0.00333333 |
| ASHFExt | W4 | Ochrobactrum | 0.00333333 |
| ASHFExt | W12 | Oryzihumus | 0.00333333 |
| ASHFExt | W8 | Oryzihumus | 0.00333333 |
| ASHFExt | W12 | Oscillospiraceae_unclassified | 0.00333333 |
| ASHFExt | W12 | Paenibacillaceae_unclassified | 0.00333333 |
| ASHFExt | W4 | Paenibacillaceae_unclassified | 0.00333333 |
| ASHFExt | W12 | Patulibacter | 0.00333333 |
| ASHFExt | W4 | Patulibacter | 0.00333333 |
| ASHFExt | W12 | Peredibacter | 0.00333333 |
| ASHFExt | W12 | Persicitalea | 0.00333333 |
| ASHFExt | W12 | Phenylobacterium | 0.00333333 |
| ASHFExt | W8 | Pirellula | 0.00333333 |
| ASHFExt | W12 | Promicromonosporaceae_unclassified | 0.00333333 |
| ASHFExt | W12 | Pseudonocardiaceae_unclassified | 0.00333333 |
| ASHFExt | W8 | Rhodothermaceae_unclassified | 0.00333333 |
| ASHFExt | W12 | Rickettsiella | 0.00333333 |
| ASHFExt | W12 | Saccharimonadaceae_unclassified | 0.00333333 |
| ASHFExt | W4 | Salinisphaera | 0.00333333 |
| ASHFExt | W12 | Sanguibacter-Flavimobilis | 0.00333333 |
| ASHFExt | W12 | Sediminibacterium | 0.00333333 |
| ASHFExt | W4 | Skermanella | 0.00333333 |
| ASHFExt | W8 | Sphingoaurantiacus | 0.00333333 |
| ASHFExt | Fresh | Staphylococcaceae_unclassified | 0.00333333 |
| ASHFExt | W12 | Steroidobacter | 0.00333333 |

| | | | |
|---|---|---|---|
| ASHFExt | W12 | Streptococcaceae_unclassified | 0.00333333 |
| ASHFExt | W12 | Streptomyces | 0.00333333 |
| ASHFExt | W4 | Streptomyces | 0.00333333 |
| ASHFExt | W12 | Tepidisphaerales_unclassified | 0.00333333 |
| ASHFExt | W8 | Thermoleophilia_unclassified | 0.00333333 |
| ASHFExt | W8 | Truepera | 0.00333333 |
| ASHFExt | W12 | Tundrisphaera | 0.00333333 |
| ASHFExt | W4 | Weeksellaceae_unclassified | 0.00333333 |
| ASHFExt | W12 | Xanthobacteraceae_unclassified | 0.00333333 |
| ASHFExt | Fresh | 67-14_ge | 0.00333333 |
| ASHFExt | Fresh | Alphaproteobacteria_unclassified | 0.00333333 |
| ASHFExt | Fresh | Breznakia | 0.00333333 |
| ASHFExt | Fresh | Chitinibacter | 0.00333333 |
| ASHFExt | Fresh | Eubacterium | 0.00333333 |
| ASHFExt | Fresh | WPS-2_ge | 0.00333333 |
| ASHFExt | Fresh | Yersiniaceae_unclassified | 0.00333333 |
| ASHFExt | Fresh | Aequorivita | 0.0025 |
| ASHFExt | Fresh | Bacteroides | 0.0025 |
| ASHFExt | Fresh | Candidatus_Soleaferrea | 0.0025 |
| ASHFExt | Fresh | Cyanobacteriia_unclassified | 0.0025 |
| ASHFExt | Fresh | Oscillospirales_unclassified | 0.0025 |
| ASHFExt | Fresh | Planococcaceae_unclassified | 0.0025 |
| ASHFExt | Fresh | Acetobacteraceae_unclassified | 0.0025 |
| ASHFExt | Fresh | Chlamydia | 0.0025 |
| ASHFExt | Fresh | Leptotrichia | 0.0025 |
| ASHFExt | Fresh | Streptomyces | 0.0025 |
| ASHFExt | Fresh | Weissella | 0.0025 |
| ASHFExt | Fresh | Xanthomonadales_unclassified | 0.0025 |
| ASHFExt | W4 | Absconditabacteriales_(SR1)_ge | 0.00166667 |

| | | | |
|---|---|---|---|
| ASHFExt | W4 | Acetobacteraceae_unclassified | 0.00166667 |
| ASHFExt | W8 | Acidimicrobiia_unclassified | 0.00166667 |
| ASHFExt | W4 | Acidobacteriales_unclassified | 0.00166667 |
| ASHFExt | W8 | Aliterella | 0.00166667 |
| ASHFExt | W8 | Angustibacter | 0.00166667 |
| ASHFExt | W4 | Aquabacterium | 0.00166667 |
| ASHFExt | W8 | Aquamicrobium | 0.00166667 |
| ASHFExt | W8 | Armatimonadales_ge | 0.00166667 |
| ASHFExt | W8 | Azospirillaceae_unclassified | 0.00166667 |
| ASHFExt | W8 | Bogoriella | 0.00166667 |
| ASHFExt | Fresh | Bradyrhizobium | 0.00166667 |
| ASHFExt | W8 | Bradyrhizobium | 0.00166667 |
| ASHFExt | Fresh | Burkholderia-Caballeronia-Paraburkholderia | 0.00166667 |
| ASHFExt | W4 | Burkholderia-Caballeronia-Paraburkholderia | 0.00166667 |
| ASHFExt | W8 | Caenimonas | 0.00166667 |
| ASHFExt | W8 | Candidatus_Megaira | 0.00166667 |
| ASHFExt | W8 | Candidatus_Soleaferrea | 0.00166667 |
| ASHFExt | W8 | Cellulomonadaceae_unclassified | 0.00166667 |
| ASHFExt | W8 | Cellulomonas | 0.00166667 |
| ASHFExt | W4 | Commensalibacter | 0.00166667 |
| ASHFExt | W8 | Conexibacter | 0.00166667 |
| ASHFExt | W8 | Constrictibacter | 0.00166667 |
| ASHFExt | W8 | Cyanobacteriia_unclassified | 0.00166667 |
| ASHFExt | W4 | D05-2_ge | 0.00166667 |
| ASHFExt | W8 | Dermabacteraceae_unclassified | 0.00166667 |
| ASHFExt | Fresh | Dickeya | 0.00166667 |
| ASHFExt | W4 | Diplorickettsiaceae_unclassified | 0.00166667 |
| ASHFExt | W8 | Diplorickettsiaceae_unclassified | 0.00166667 |
| ASHFExt | W8 | Erysipelatoclostridium | 0.00166667 |

| | | | |
|---|---|---|---|
| ASHFExt | W4 | Ferruginibacter | 0.00166667 |
| ASHFExt | W8 | Fimbriimonadaceae_unclassified | 0.00166667 |
| ASHFExt | W8 | Flavobacteriaceae_unclassified | 0.00166667 |
| ASHFExt | W4 | Flexivirga | 0.00166667 |
| ASHFExt | W8 | Geodermatophilus | 0.00166667 |
| ASHFExt | W4 | Herpetosiphon | 0.00166667 |
| ASHFExt | W8 | Herpetosiphon | 0.00166667 |
| ASHFExt | W8 | Hirschia | 0.00166667 |
| ASHFExt | W4 | Iamia | 0.00166667 |
| ASHFExt | W4 | Jeotgalicoccus | 0.00166667 |
| ASHFExt | W4 | Kaistia | 0.00166667 |
| ASHFExt | W8 | Kineosporia | 0.00166667 |
| ASHFExt | W8 | LWQ8_ge | 0.00166667 |
| ASHFExt | W8 | Labrys | 0.00166667 |
| ASHFExt | W8 | Limnobacter | 0.00166667 |
| ASHFExt | W8 | Luteimonas | 0.00166667 |
| ASHFExt | W8 | Mariniradius | 0.00166667 |
| ASHFExt | W4 | Marmoricola | 0.00166667 |
| ASHFExt | W4 | Modestobacter | 0.00166667 |
| ASHFExt | W8 | Mycobacteriaceae_unclassified | 0.00166667 |
| ASHFExt | W8 | Myxococcus | 0.00166667 |
| ASHFExt | W4 | Nakamurella | 0.00166667 |
| ASHFExt | W8 | Nitrososphaeraceae_unclassified | 0.00166667 |
| ASHFExt | W8 | Nubsella | 0.00166667 |
| ASHFExt | W4 | Oligoflexus | 0.00166667 |
| ASHFExt | W4 | Oryzihumus | 0.00166667 |
| ASHFExt | W4 | Paeniglutamicibacter | 0.00166667 |
| ASHFExt | W8 | Paludisphaera | 0.00166667 |
| ASHFExt | W4 | Parafilimonas | 0.00166667 |

| | | | |
|---|---|---|---|
| ASHFExt | W8 | Parviterribacter | 0.00166667 |
| ASHFExt | W4 | Pedococcus-Phycicoccus | 0.00166667 |
| ASHFExt | W8 | Peptostreptococcales-Tissierellales_unclassifie | 0.00166667 |
| ASHFExt | W4 | Phenylobacterium | 0.00166667 |
| ASHFExt | W8 | Planococcus | 0.00166667 |
| ASHFExt | W4 | Promicromonosporaceae_unclassified | 0.00166667 |
| ASHFExt | W8 | Propionibacteriales_unclassified | 0.00166667 |
| ASHFExt | W4 | Proteobacteria_unclassified | 0.00166667 |
| ASHFExt | W8 | Pseudactinotalea | 0.00166667 |
| ASHFExt | W4 | Pseudoduganella | 0.00166667 |
| ASHFExt | W4 | Pseudomonadales_unclassified | 0.00166667 |
| ASHFExt | W8 | Pseudomonadales_unclassified | 0.00166667 |
| ASHFExt | Fresh | Pseudomonas | 0.00166667 |
| ASHFExt | W8 | Pseudonocardia | 0.00166667 |
| ASHFExt | W4 | Quadrisphaera | 0.00166667 |
| ASHFExt | W4 | Rhodanobacter | 0.00166667 |
| ASHFExt | W8 | Rhodanobacter | 0.00166667 |
| ASHFExt | W4 | Rhodopseudomonas | 0.00166667 |
| ASHFExt | W4 | Rhodothermaceae_ge | 0.00166667 |
| ASHFExt | W8 | Rickettsiales_unclassified | 0.00166667 |
| ASHFExt | W4 | Rickettsiella | 0.00166667 |
| ASHFExt | W8 | Roseiflexaceae_unclassified | 0.00166667 |
| ASHFExt | W8 | Saccharimonadaceae_unclassified | 0.00166667 |
| ASHFExt | W4 | Solirubrobacter | 0.00166667 |
| ASHFExt | W4 | Spirosoma | 0.00166667 |
| ASHFExt | W8 | Sporacetigenium | 0.00166667 |
| ASHFExt | W8 | Steroidobacteraceae_unclassified | 0.00166667 |
| ASHFExt | W4 | Streptococcus | 0.00166667 |
| ASHFExt | W4 | Streptomycetaceae_unclassified | 0.00166667 |

| | | | |
|---|---|---|---|
| ASHFExt | W4 | Tepidisphaeraceae_ge | 0.00166667 |
| ASHFExt | Fresh | Thermoleophilia_unclassified | 0.00166667 |
| ASHFExt | W4 | Truepera | 0.00166667 |
| ASHFExt | W8 | Tyzzerella | 0.00166667 |
| ASHFExt | W8 | Vampirovibrionales_unclassified | 0.00166667 |
| ASHFExt | W4 | Wolbachia | 0.00166667 |
| ASHFExt | W4 | mle1-27_ge | 0.00166667 |
| ASHFExt | Fresh | Anaerovoracaceae_unclassified | 0.00166667 |
| ASHFExt | Fresh | Blattabacterium | 0.00166667 |
| ASHFExt | Fresh | Brachybacterium | 0.00166667 |
| ASHFExt | Fresh | Desulfovibrionaceae_unclassified | 0.00166667 |
| ASHFExt | Fresh | Eggerthellaceae_unclassified | 0.00166667 |
| ASHFExt | Fresh | Frankiales_unclassified | 0.00166667 |
| ASHFExt | Fresh | Gordonia | 0.00166667 |
| ASHFExt | Fresh | Incertae_Sedis | 0.00166667 |
| ASHFExt | Fresh | Intrasporangiaceae_unclassified | 0.00166667 |
| ASHFExt | Fresh | Lactiplantibacillus | 0.00166667 |
| ASHFExt | Fresh | Limibaculum | 0.00166667 |
| ASHFExt | Fresh | Neorickettsia | 0.00166667 |
| ASHFExt | Fresh | Nocardia | 0.00166667 |
| ASHFExt | Fresh | Nocardiaceae_unclassified | 0.00166667 |
| ASHFExt | Fresh | Nocardiopsis | 0.00166667 |
| ASHFExt | Fresh | Oxalobacter | 0.00166667 |
| ASHFExt | Fresh | Patulibacter | 0.00166667 |
| ASHFExt | Fresh | RF39_ge | 0.00166667 |
| ASHFExt | Fresh | Rhizobiaceae_unclassified | 0.00166667 |
| ASHFExt | Fresh | Salicola | 0.00166667 |
| ASHFExt | Fresh | Shimwellia | 0.00166667 |
| ASHFExt | Fresh | Spiroplasma | 0.00166667 |

| | | | |
|---|---|---|---|
| ASHFExt | Fresh | Tissierella | 0.00166667 |
| ASHFExt | Fresh | Tundrisphaera | 0.00166667 |
| ASHFExt | Fresh | Alsobacter | 0.00083333 |
| ASHFExt | Fresh | Anaerofustis | 0.00083333 |
| ASHFExt | Fresh | Bacteroidales_unclassified | 0.00083333 |
| ASHFExt | Fresh | CL500-29_marine_group | 0.00083333 |
| ASHFExt | Fresh | Campylobacterales_unclassified | 0.00083333 |
| ASHFExt | Fresh | Candidatus_Omnitrophus | 0.00083333 |
| ASHFExt | Fresh | Candidatus_Rhabdochlamydia | 0.00083333 |
| ASHFExt | Fresh | Candidatus_Tammella | 0.00083333 |
| ASHFExt | Fresh | Carnobacteriaceae_unclassified | 0.00083333 |
| ASHFExt | Fresh | Chitinophagaceae_unclassified | 0.00083333 |
| ASHFExt | Fresh | Chloroflexi_unclassified | 0.00083333 |
| ASHFExt | Fresh | Christensenellaceae_R-7_group | 0.00083333 |
| ASHFExt | Fresh | Chryseobacterium | 0.00083333 |
| ASHFExt | Fresh | Crenobacter | 0.00083333 |
| ASHFExt | Fresh | Desulfobacterales_unclassified | 0.00083333 |
| ASHFExt | Fresh | Desulfobacterota_unclassified | 0.00083333 |
| ASHFExt | Fresh | Epulopiscium | 0.00083333 |
| ASHFExt | Fresh | Erysipelotrichaceae_unclassified | 0.00083333 |
| ASHFExt | Fresh | Flavobacterium | 0.00083333 |
| ASHFExt | Fresh | Gammaproteobacteria_unclassified | 0.00083333 |
| ASHFExt | Fresh | Haloparvum | 0.00083333 |
| ASHFExt | Fresh | Hyphomicrobiaceae_unclassified | 0.00083333 |
| ASHFExt | Fresh | Isosphaeraceae_unclassified | 0.00083333 |
| ASHFExt | Fresh | Jeotgalicoccus | 0.00083333 |
| ASHFExt | Fresh | Leifsonia | 0.00083333 |
| ASHFExt | Fresh | Luteolibacter | 0.00083333 |
| ASHFExt | Fresh | Marinomonas | 0.00083333 |

| | | | |
|---|---|---|---|
| ASHFExt | Fresh | Marmoricola | 0.00083333 |
| ASHFExt | Fresh | Massilia | 0.00083333 |
| ASHFExt | Fresh | Microtrichales_unclassified | 0.00083333 |
| ASHFExt | Fresh | Myxococcaceae_unclassified | 0.00083333 |
| ASHFExt | Fresh | Oscillospiraceae_unclassified | 0.00083333 |
| ASHFExt | Fresh | Oxalobacteraceae_unclassified | 0.00083333 |
| ASHFExt | Fresh | Paludicola | 0.00083333 |
| ASHFExt | Fresh | Parapusillimonas | 0.00083333 |
| ASHFExt | Fresh | Pasteurellaceae_unclassified | 0.00083333 |
| ASHFExt | Fresh | Pedobacter | 0.00083333 |
| ASHFExt | Fresh | Pseudogracilibacillus | 0.00083333 |
| ASHFExt | Fresh | Psychroglaciecola | 0.00083333 |
| ASHFExt | Fresh | Rhodoplanes | 0.00083333 |
| ASHFExt | Fresh | Rhodospirillales_unclassified | 0.00083333 |
| ASHFExt | Fresh | Rs-K70_termite_group_ge | 0.00083333 |
| ASHFExt | Fresh | RsaHf231_ge | 0.00083333 |
| ASHFExt | Fresh | SJA-15_ge | 0.00083333 |
| ASHFExt | Fresh | Solirubrobacteraceae_unclassified | 0.00083333 |
| ASHFExt | Fresh | Solirubrobacterales_unclassified | 0.00083333 |
| ASHFExt | Fresh | Streptomycetaceae_unclassified | 0.00083333 |
| ASHFExt | Fresh | TM7a | 0.00083333 |
| ASHFExt | Fresh | Taibaiella | 0.00083333 |
| ASHFExt | Fresh | Telmatospirillum | 0.00083333 |
| ASHFExt | Fresh | Wolbachia | 0.00083333 |
| ASHFExt | Fresh | endosymbionts | 0.00083333 |
| ASHFExt | Fresh | vadinHA49_ge | 0.00083333 |
| ASHFInt | W8 | Clostridiaceae_unclassified | 56.5966667 |
| ASHFInt | W12 | Clostridiaceae_unclassified | 50.5266667 |
| ASHFInt | W4 | Clostridiaceae_unclassified | 36.4416667 |

| | | | |
|---|---|---|---|
| ASHFInt | Fresh | Clostridiaceae_unclassified | 32.4575 |
| ASHFInt | W8 | Plesiomonas | 19.6066667 |
| ASHFInt | W4 | Plesiomonas | 16.8 |
| ASHFInt | Fresh | Candidatus_Arthromitus | 14.3041667 |
| ASHFInt | Fresh | Plesiomonas | 13.4933333 |
| ASHFInt | W4 | Candidatus_Arthromitus | 13.1016667 |
| ASHFInt | W12 | Plesiomonas | 11.8866667 |
| ASHFInt | W12 | Salinisphaera | 7.74333333 |
| ASHFInt | Fresh | Rickettsiella | 7.1175 |
| ASHFInt | W4 | Hafnia-Obesumbacterium | 6.1 |
| ASHFInt | W12 | Aeromonas | 5.07666667 |
| ASHFInt | W12 | Candidatus_Arthromitus | 4.43 |
| ASHFInt | W8 | Enterococcus | 4.32 |
| ASHFInt | W12 | Enterococcus | 4.28666667 |
| ASHFInt | W8 | Enterobacteriaceae_unclassified | 4.27333333 |
| ASHFInt | Fresh | Enterobacteriaceae_unclassified | 4.2725 |
| ASHFInt | Fresh | Enterococcus | 4.23083333 |
| ASHFInt | W4 | Enterobacteriaceae_unclassified | 3.96333333 |
| ASHFInt | W4 | Ureaplasma | 3.94333333 |
| ASHFInt | Fresh | Bacillaceae_unclassified | 3.395 |
| ASHFInt | Fresh | Cetobacterium | 3.24 |
| ASHFInt | W8 | Candidatus_Arthromitus | 3.15 |
| ASHFInt | W4 | Enterococcus | 3.02333333 |
| ASHFInt | W4 | Clostridium_sensu_stricto_1 | 3.01 |
| ASHFInt | Fresh | Lactococcus | 2.83083333 |
| ASHFInt | Fresh | Candidatus_Rhabdochlamydia | 2.55666667 |
| ASHFInt | Fresh | Hafnia-Obesumbacterium | 2.535 |
| ASHFInt | W8 | Paraclostridium | 2.41333333 |
| ASHFInt | W4 | Peptostreptococcaceae_unclassified | 2.22833333 |

| | | | |
|---|---|---|---|
| ASHFInt | W4 | Morganella | 2.08666667 |
| ASHFInt | W12 | Bacillaceae_unclassified | 1.9 |
| ASHFInt | W8 | Clostridium_sensu_stricto_1 | 1.89 |
| ASHFInt | W12 | Clostridium_sensu_stricto_1 | 1.61 |
| ASHFInt | W4 | Paeniclostridium | 1.48333333 |
| ASHFInt | W12 | Hafnia-Obesumbacterium | 1.43 |
| ASHFInt | Fresh | Clostridium_sensu_stricto_1 | 1.26166667 |
| ASHFInt | W8 | Mycoplasmataceae_unclassified | 1.11 |
| ASHFInt | W4 | Bacillaceae_unclassified | 1.105 |
| ASHFInt | W4 | Salinisphaera | 1.095 |
| ASHFInt | W12 | Staphylococcus | 1.06 |
| ASHFInt | W4 | Lactococcus | 0.96 |
| ASHFInt | Fresh | Nosocomiicoccus | 0.94333333 |
| ASHFInt | W8 | Bacillaceae_unclassified | 0.93666667 |
| ASHFInt | W8 | Cetobacterium | 0.92666667 |
| ASHFInt | W8 | Hafnia-Obesumbacterium | 0.87 |
| ASHFInt | W12 | Morganella | 0.80666667 |
| ASHFInt | W12 | Nosocomiicoccus | 0.79 |
| ASHFInt | W12 | Enterobacteriaceae_unclassified | 0.72666667 |
| ASHFInt | W4 | Paraclostridium | 0.685 |
| ASHFInt | Fresh | Micrococcales_unclassified | 0.68333333 |
| ASHFInt | W4 | Aeromonas | 0.67666667 |
| ASHFInt | W12 | Paraclostridium | 0.65333333 |
| ASHFInt | W12 | Peptostreptococcaceae_unclassified | 0.65333333 |
| ASHFInt | W12 | Lachnospiraceae_unclassified | 0.64666667 |
| ASHFInt | W12 | Enterobacterales_unclassified | 0.63666667 |
| ASHFInt | Fresh | Mycoplasmataceae_unclassified | 0.61166667 |
| ASHFInt | Fresh | Corynebacteriales_unclassified | 0.605 |
| ASHFInt | Fresh | Salinisphaera | 0.59833333 |

| | | | |
|---|---|---|---|
| ASHFInt | W8 | Paeniclostridium | 0.51666667 |
| ASHFInt | W4 | Orbus | 0.50833333 |
| ASHFInt | W12 | Bacteroidia_unclassified | 0.48 |
| ASHFInt | W8 | Peptostreptococcaceae_unclassified | 0.47333333 |
| ASHFInt | Fresh | Enterobacterales_unclassified | 0.44833333 |
| ASHFInt | W4 | Nosocomiicoccus | 0.44 |
| ASHFInt | W4 | Cetobacterium | 0.40666667 |
| ASHFInt | W8 | Nosocomiicoccus | 0.38666667 |
| ASHFInt | W12 | Mycoplasmataceae_unclassified | 0.38333333 |
| ASHFInt | Fresh | Helicobacter | 0.37583333 |
| ASHFInt | Fresh | Bacteroidia_unclassified | 0.36166667 |
| ASHFInt | W8 | Salinisphaera | 0.35666667 |
| ASHFInt | W8 | Ureaplasma | 0.35 |
| ASHFInt | Fresh | Rickettsia | 0.34166667 |
| ASHFInt | W12 | Synergistaceae_unclassified | 0.32666667 |
| ASHFInt | W12 | Ureaplasma | 0.32333333 |
| ASHFInt | W12 | Firmicutes_unclassified | 0.31333333 |
| ASHFInt | W12 | Corynebacteriales_unclassified | 0.31 |
| ASHFInt | W4 | Enterobacterales_unclassified | 0.29833333 |
| ASHFInt | Fresh | Atopostipes | 0.27416667 |
| ASHFInt | Fresh | Aeromonas | 0.245 |
| ASHFInt | Fresh | Alcaligenaceae_ge | 0.22333333 |
| ASHFInt | W8 | Corynebacteriales_unclassified | 0.22 |
| ASHFInt | Fresh | Diplorickettsiaceae_unclassified | 0.215 |
| ASHFInt | Fresh | Bartonella | 0.2 |
| ASHFInt | Fresh | Ruminococcaceae_unclassified | 0.19833333 |
| ASHFInt | W12 | Sebaldella | 0.19666667 |
| ASHFInt | W8 | Ruminococcaceae_unclassified | 0.19333333 |
| ASHFInt | W4 | Corynebacteriales_unclassified | 0.19166667 |

| | | | |
|---|---|---|---|
| ASHFInt | Fresh | Corynebacterium | 0.19 |
| ASHFInt | W12 | Paeniclostridium | 0.18666667 |
| ASHFInt | W12 | Cetobacterium | 0.16666667 |
| ASHFInt | W12 | Fusobacterium | 0.16666667 |
| ASHFInt | W12 | Atopostipes | 0.15666667 |
| ASHFInt | W4 | Mycoplasma | 0.14666667 |
| ASHFInt | W12 | Alcaligenaceae_ge | 0.13333333 |
| ASHFInt | W12 | Flavobacteriaceae_unclassified | 0.13 |
| ASHFInt | Fresh | Paraclostridium | 0.1275 |
| ASHFInt | W8 | Lachnospiraceae_unclassified | 0.12333333 |
| ASHFInt | W8 | Bacteroidia_unclassified | 0.11666667 |
| ASHFInt | Fresh | Carnobacterium | 0.11416667 |
| ASHFInt | W12 | Mycoplasma | 0.10666667 |
| ASHFInt | Fresh | Ureaplasma | 0.10333333 |
| ASHFInt | W4 | Lacticaseibacillus | 0.1 |
| ASHFInt | W4 | Alcaligenaceae_ge | 0.09666667 |
| ASHFInt | W8 | Synergistaceae_unclassified | 0.09666667 |
| ASHFInt | W8 | Enterobacterales_unclassified | 0.09 |
| ASHFInt | Fresh | Micrococcaceae_unclassified | 0.0875 |
| ASHFInt | W12 | Bogoriella | 0.08666667 |
| ASHFInt | W4 | Tyzzerella | 0.085 |
| ASHFInt | W4 | Romboutsia | 0.08333333 |
| ASHFInt | Fresh | Peptostreptococcaceae_unclassified | 0.08 |
| ASHFInt | W4 | Atopostipes | 0.07833333 |
| ASHFInt | W12 | Flavobacteriales_unclassified | 0.07666667 |
| ASHFInt | W12 | Bacilli_unclassified | 0.07333333 |
| ASHFInt | W8 | Fusobacterium | 0.07333333 |
| ASHFInt | W12 | Ruminococcaceae_unclassified | 0.07333333 |
| ASHFInt | W8 | Arsenophonus | 0.07 |

| | | | |
|---|---|---|---|
| ASHFInt | W8 | Atopostipes | 0.06666667 |
| ASHFInt | W12 | Lactococcus | 0.06333333 |
| ASHFInt | W4 | Epulopiscium | 0.06166667 |
| ASHFInt | W12 | Alkanindiges | 0.06 |
| ASHFInt | W4 | Terrisporobacter | 0.06 |
| ASHFInt | W12 | Carnobacterium | 0.06 |
| ASHFInt | Fresh | Commensalibacter | 0.05916667 |
| ASHFInt | W12 | Candidatus_Rhabdochlamydia | 0.05666667 |
| ASHFInt | W12 | Truepera | 0.05666667 |
| ASHFInt | W12 | Lactobacillales_unclassified | 0.05333333 |
| ASHFInt | W4 | Mycobacterium | 0.05333333 |
| ASHFInt | Fresh | Aquabacterium | 0.0525 |
| ASHFInt | W12 | Bacteria_unclassified | 0.05 |
| ASHFInt | Fresh | Bogoriella | 0.05 |
| ASHFInt | W8 | Lactobacillales_unclassified | 0.05 |
| ASHFInt | W12 | Micrococcaceae_unclassified | 0.05 |
| ASHFInt | W8 | Morganella | 0.05 |
| ASHFInt | W12 | Pragia | 0.05 |
| ASHFInt | W4 | Ruminococcaceae_unclassified | 0.05 |
| ASHFInt | Fresh | Morganella | 0.04583333 |
| ASHFInt | Fresh | Microbacteriaceae_unclassified | 0.045 |
| ASHFInt | W4 | Micrococcaceae_unclassified | 0.04333333 |
| ASHFInt | Fresh | Bacteria_unclassified | 0.04166667 |
| ASHFInt | W4 | Tomitella | 0.04166667 |
| ASHFInt | W4 | Bartonella | 0.04 |
| ASHFInt | W8 | Lactococcus | 0.04 |
| ASHFInt | W8 | Staphylococcus | 0.04 |
| ASHFInt | W12 | uncultured | 0.04 |
| ASHFInt | Fresh | Leuconostoc | 0.04 |

| | | | |
|---|---|---|---|
| ASHFInt | Fresh | Lachnospiraceae_unclassified | 0.03916667 |
| ASHFInt | Fresh | uncultured | 0.03916667 |
| ASHFInt | W4 | Staphylococcus | 0.03833333 |
| ASHFInt | Fresh | Lactobacillales_unclassified | 0.0375 |
| ASHFInt | W12 | Micrococcales_unclassified | 0.03666667 |
| ASHFInt | W12 | Desulfovibrio | 0.03666667 |
| ASHFInt | W12 | Raoultibacter | 0.03666667 |
| ASHFInt | W8 | Sebaldella | 0.03666667 |
| ASHFInt | Fresh | Mycobacterium | 0.035 |
| ASHFInt | Fresh | Nocardioides | 0.03333333 |
| ASHFInt | Fresh | Lactobacillaceae_unclassified | 0.03083333 |
| ASHFInt | Fresh | Tyzzerella | 0.03083333 |
| ASHFInt | W12 | Alcaligenaceae_unclassified | 0.03 |
| ASHFInt | W12 | Intestinimonas | 0.03 |
| ASHFInt | W8 | Raoultibacter | 0.03 |
| ASHFInt | Fresh | Tomitella | 0.03 |
| ASHFInt | W12 | Christensenellaceae_R-7_group | 0.03 |
| ASHFInt | W12 | Oscillospirales_unclassified | 0.03 |
| ASHFInt | W12 | Pusillimonas | 0.03 |
| ASHFInt | Fresh | Flavobacteriales_unclassified | 0.02916667 |
| ASHFInt | Fresh | Firmicutes_unclassified | 0.0275 |
| ASHFInt | W8 | Alcaligenaceae_ge | 0.02666667 |
| ASHFInt | W4 | Corynebacterium | 0.02666667 |
| ASHFInt | W8 | Corynebacterium | 0.02666667 |
| ASHFInt | W12 | Erysipelatoclostridiaceae_unclassified | 0.02666667 |
| ASHFInt | W8 | uncultured | 0.02666667 |
| ASHFInt | W12 | Membranicola | 0.02666667 |
| ASHFInt | W8 | Bogoriella | 0.02333333 |
| ASHFInt | W4 | Firmicutes_unclassified | 0.02333333 |

| | | | |
|---|---|---|---|
| ASHFInt | W8 | Firmicutes_unclassified | 0.02333333 |
| ASHFInt | W12 | Oscillospiraceae_unclassified | 0.02333333 |
| ASHFInt | W8 | Oscillospirales_unclassified | 0.02333333 |
| ASHFInt | W12 | Rhodobacteraceae_unclassified | 0.02333333 |
| ASHFInt | W12 | Rs-K70_termite_group_ge | 0.02333333 |
| ASHFInt | W12 | Tomitella | 0.02333333 |
| ASHFInt | W12 | Microbacteriaceae_unclassified | 0.02333333 |
| ASHFInt | W12 | Rhizobiaceae_unclassified | 0.02333333 |
| ASHFInt | Fresh | Bacilli_unclassified | 0.0225 |
| ASHFInt | Fresh | Paeniclostridium | 0.0225 |
| ASHFInt | Fresh | RsaHf231_ge | 0.02166667 |
| ASHFInt | Fresh | JG30-KF-CM45_ge | 0.02083333 |
| ASHFInt | W4 | Bacilli_unclassified | 0.02 |
| ASHFInt | W8 | Clostridia_unclassified | 0.02 |
| ASHFInt | W12 | Iamia | 0.02 |
| ASHFInt | W8 | Iamia | 0.02 |
| ASHFInt | W8 | Micrococcaceae_unclassified | 0.02 |
| ASHFInt | W8 | Oscillospiraceae_unclassified | 0.02 |
| ASHFInt | Fresh | Truepera | 0.01916667 |
| ASHFInt | Fresh | Alkanindiges | 0.01833333 |
| ASHFInt | W4 | Bacteroidia_unclassified | 0.01833333 |
| ASHFInt | Fresh | Curtobacterium | 0.01833333 |
| ASHFInt | W12 | Candidatus_Soleaferrea | 0.01666667 |
| ASHFInt | W12 | JG30-KF-CM45_ge | 0.01666667 |
| ASHFInt | W12 | Orbus | 0.01666667 |
| ASHFInt | Fresh | Bacillales_unclassified | 0.01666667 |
| ASHFInt | W8 | JG30-KF-CM45_ge | 0.01666667 |
| ASHFInt | W4 | Lachnospiraceae_unclassified | 0.01666667 |
| ASHFInt | W12 | Sporosarcina | 0.01666667 |

| | | | |
|---|---|---|---|
| ASHFInt | Fresh | Pectobacteriaceae_unclassified | 0.01666667 |
| ASHFInt | Fresh | Flavobacteriaceae_unclassified | 0.01583333 |
| ASHFInt | W4 | Carnobacterium | 0.015 |
| ASHFInt | W4 | Lactobacillales_unclassified | 0.015 |
| ASHFInt | Fresh | uncultured_ge | 0.015 |
| ASHFInt | W8 | Alkanindiges | 0.01333333 |
| ASHFInt | W8 | Anaerovoracaceae_unclassified | 0.01333333 |
| ASHFInt | W12 | Aquabacterium | 0.01333333 |
| ASHFInt | W4 | Aquabacterium | 0.01333333 |
| ASHFInt | W8 | Bacteria_unclassified | 0.01333333 |
| ASHFInt | W12 | Dysgonomonas | 0.01333333 |
| ASHFInt | W12 | Helicobacter | 0.01333333 |
| ASHFInt | W8 | Microbacteriaceae_unclassified | 0.01333333 |
| ASHFInt | W4 | Pasteurellaceae_unclassified | 0.01333333 |
| ASHFInt | W12 | Propionibacteriales_unclassified | 0.01333333 |
| ASHFInt | W4 | Providencia | 0.01333333 |
| ASHFInt | W12 | Pseudomonas | 0.01333333 |
| ASHFInt | W8 | Sarcina | 0.01333333 |
| ASHFInt | W4 | Sporosarcina | 0.01333333 |
| ASHFInt | W12 | uncultured_ge | 0.01333333 |
| ASHFInt | Fresh | Alcaligenaceae_unclassified | 0.0125 |
| ASHFInt | Fresh | Iamia | 0.0125 |
| ASHFInt | Fresh | Streptomyces | 0.0125 |
| ASHFInt | W4 | Aeromicrobium | 0.01166667 |
| ASHFInt | W4 | Bogoriella | 0.01166667 |
| ASHFInt | W4 | uncultured_ge | 0.01166667 |
| ASHFInt | Fresh | Oscillospirales_unclassified | 0.01083333 |
| ASHFInt | W8 | Bacilli_unclassified | 0.01 |
| ASHFInt | W12 | Staphylococcaceae_unclassified | 0.01 |

| | | | |
|---|---|---|---|
| ASHFInt | W12 | Acidaminococcaceae_unclassified | 0.01 |
| ASHFInt | W12 | Anaerovoracaceae_unclassified | 0.01 |
| ASHFInt | W12 | Bartonella | 0.01 |
| ASHFInt | W12 | Brevinema | 0.01 |
| ASHFInt | W8 | Christensenellaceae_R-7_group | 0.01 |
| ASHFInt | W8 | Incertae_Sedis | 0.01 |
| ASHFInt | W12 | Leucobacter | 0.01 |
| ASHFInt | Fresh | Mycoplasma | 0.01 |
| ASHFInt | W4 | Rhizobiaceae_unclassified | 0.01 |
| ASHFInt | W4 | Sebaldella | 0.01 |
| ASHFInt | Fresh | Breznakia | 0.01 |
| ASHFInt | W8 | Candidatus_Soleaferrea | 0.01 |
| ASHFInt | W8 | Flavobacteriales_unclassified | 0.01 |
| ASHFInt | W12 | Pseudomonadales_unclassified | 0.01 |
| ASHFInt | Fresh | Raoultibacter | 0.01 |
| ASHFInt | W12 | Rickettsiaceae_ge | 0.01 |
| ASHFInt | W8 | Sphingobacteriaceae_unclassified | 0.01 |
| ASHFInt | W12 | Taibaiella | 0.01 |
| ASHFInt | W12 | Thiopseudomonas | 0.01 |
| ASHFInt | W12 | Xanthomonadales_unclassified | 0.01 |
| ASHFInt | Fresh | Aeromicrobium | 0.00916667 |
| ASHFInt | Fresh | Leucobacter | 0.00916667 |
| ASHFInt | Fresh | Weissella | 0.00916667 |
| ASHFInt | W4 | Candidatus_Rhabdochlamydia | 0.00833333 |
| ASHFInt | W4 | Dysgonomonas | 0.00833333 |
| ASHFInt | W4 | Microbacteriaceae_unclassified | 0.00833333 |
| ASHFInt | W4 | Micrococcales_unclassified | 0.00833333 |
| ASHFInt | Fresh | Rhizobiaceae_unclassified | 0.00833333 |
| ASHFInt | Fresh | Oscillospiraceae_unclassified | 0.00833333 |

| | | | |
|---|---|---|---|
| ASHFInt | W4 | Oscillospiraceae_unclassified | 0.00833333 |
| ASHFInt | Fresh | Comamonadaceae_unclassified | 0.0075 |
| ASHFInt | Fresh | Camelimonas | 0.0075 |
| ASHFInt | Fresh | Clostridia_unclassified | 0.0075 |
| ASHFInt | Fresh | Wolbachia | 0.0075 |
| ASHFInt | W12 | Aeromicrobium | 0.00666667 |
| ASHFInt | W8 | Alcaligenaceae_unclassified | 0.00666667 |
| ASHFInt | Fresh | Asaia | 0.00666667 |
| ASHFInt | W8 | Bacillales_unclassified | 0.00666667 |
| ASHFInt | W12 | Bacteroides | 0.00666667 |
| ASHFInt | W8 | Bartonella | 0.00666667 |
| ASHFInt | W12 | Clostridia_unclassified | 0.00666667 |
| ASHFInt | W4 | Clostridiales_unclassified | 0.00666667 |
| ASHFInt | W12 | Corynebacterium | 0.00666667 |
| ASHFInt | W12 | Cryomorpha | 0.00666667 |
| ASHFInt | W8 | Dietzia | 0.00666667 |
| ASHFInt | W4 | Gammaproteobacteria_unclassified | 0.00666667 |
| ASHFInt | W8 | Mariniradius | 0.00666667 |
| ASHFInt | W12 | Mycobacterium | 0.00666667 |
| ASHFInt | W4 | Mycoplasmataceae_unclassified | 0.00666667 |
| ASHFInt | W8 | Pragia | 0.00666667 |
| ASHFInt | Fresh | Propionibacteriales_unclassified | 0.00666667 |
| ASHFInt | W4 | Propionibacteriales_unclassified | 0.00666667 |
| ASHFInt | W8 | Pusillimonas | 0.00666667 |
| ASHFInt | W8 | Rhizobiaceae_unclassified | 0.00666667 |
| ASHFInt | W12 | Rhodanobacteraceae_unclassified | 0.00666667 |
| ASHFInt | W12 | Rickettsiella | 0.00666667 |
| ASHFInt | W12 | Sphingobacteriaceae_unclassified | 0.00666667 |
| ASHFInt | W8 | Taibaiella | 0.00666667 |

| | | | |
|---|---|---|---|
| ASHFInt | W4 | Turicibacter | 0.00666667 |
| ASHFInt | Fresh | Gammaproteobacteria_unclassified | 0.00666667 |
| ASHFInt | Fresh | Orbus | 0.00666667 |
| ASHFInt | Fresh | Pusillimonas | 0.00666667 |
| ASHFInt | Fresh | Rhodobacteraceae_unclassified | 0.00666667 |
| ASHFInt | Fresh | Fusobacterium | 0.00583333 |
| ASHFInt | Fresh | Paracoccus | 0.00583333 |
| ASHFInt | W4 | Aequorivita | 0.005 |
| ASHFInt | W4 | Alkanindiges | 0.005 |
| ASHFInt | W4 | Peptostreptococcales-Tissierellales_unclassifie | 0.005 |
| ASHFInt | W4 | Planococcaceae_unclassified | 0.005 |
| ASHFInt | W4 | Pseudomonas | 0.005 |
| ASHFInt | Fresh | Rickettsiaceae_unclassified | 0.005 |
| ASHFInt | W4 | Truepera | 0.005 |
| ASHFInt | W4 | uncultured | 0.005 |
| ASHFInt | Fresh | Pedobacter | 0.005 |
| ASHFInt | Fresh | Staphylococcus | 0.005 |
| ASHFInt | Fresh | endosymbionts | 0.005 |
| ASHFInt | Fresh | Providencia | 0.00416667 |
| ASHFInt | Fresh | Membranicola | 0.00416667 |
| ASHFInt | Fresh | Neorickettsia | 0.00416667 |
| ASHFInt | Fresh | Nocardiaceae_unclassified | 0.00416667 |
| ASHFInt | Fresh | Rhodococcus | 0.00416667 |
| ASHFInt | Fresh | Robinsoniella | 0.00416667 |
| ASHFInt | Fresh | Streptomycetaceae_unclassified | 0.00416667 |
| ASHFInt | Fresh | Turicibacter | 0.00416667 |
| ASHFInt | W12 | 1174-901-12 | 0.00333333 |
| ASHFInt | W8 | Acetobacter | 0.00333333 |
| ASHFInt | W4 | Actinobacteria_unclassified | 0.00333333 |

| | | | |
|---|---|---|---|
| ASHFInt | W4 | Actinomycetospora | 0.00333333 |
| ASHFInt | W12 | Aequorivita | 0.00333333 |
| ASHFInt | W8 | Aeromicrobium | 0.00333333 |
| ASHFInt | W12 | Asaia | 0.00333333 |
| ASHFInt | W12 | Bacillales_unclassified | 0.00333333 |
| ASHFInt | W4 | Bacillales_unclassified | 0.00333333 |
| ASHFInt | W4 | Bacteria_unclassified | 0.00333333 |
| ASHFInt | W4 | Beijerinckiaceae_unclassified | 0.00333333 |
| ASHFInt | W8 | Brachybacterium | 0.00333333 |
| ASHFInt | W12 | Bradymonadaceae_ge | 0.00333333 |
| ASHFInt | W12 | Bradymonadaceae_unclassified | 0.00333333 |
| ASHFInt | W12 | Brevibacterium | 0.00333333 |
| ASHFInt | W4 | Brevinema | 0.00333333 |
| ASHFInt | W8 | Brevinema | 0.00333333 |
| ASHFInt | W12 | Breznakia | 0.00333333 |
| ASHFInt | W8 | Breznakia | 0.00333333 |
| ASHFInt | W12 | Brumimicrobium | 0.00333333 |
| ASHFInt | W4 | Budvicia | 0.00333333 |
| ASHFInt | W8 | Burkholderiales_unclassified | 0.00333333 |
| ASHFInt | W12 | Candidatus_Fritschea | 0.00333333 |
| ASHFInt | W8 | Candidatus_Fritschea | 0.00333333 |
| ASHFInt | W12 | Caulobacteraceae_unclassified | 0.00333333 |
| ASHFInt | W4 | Cellulosilyticum | 0.00333333 |
| ASHFInt | Fresh | Christensenellaceae_R-7_group | 0.00333333 |
| ASHFInt | W4 | Chryseobacterium | 0.00333333 |
| ASHFInt | W12 | Coprococcus | 0.00333333 |
| ASHFInt | W8 | Curtobacterium | 0.00333333 |
| ASHFInt | W12 | Desulfobacterales_unclassified | 0.00333333 |
| ASHFInt | W8 | Desulfovibrio | 0.00333333 |

| | | | |
|---|---|---|---|
| ASHFInt | W12 | Desulfovibrionaceae_unclassified | 0.00333333 |
| ASHFInt | W12 | Devosiaceae_unclassified | 0.00333333 |
| ASHFInt | W8 | Erysipelatoclostridiaceae_unclassified | 0.00333333 |
| ASHFInt | Fresh | Escherichia-Shigella | 0.00333333 |
| ASHFInt | W12 | Eubacterium | 0.00333333 |
| ASHFInt | W12 | Family_XI_unclassified | 0.00333333 |
| ASHFInt | W4 | Family_XI_unclassified | 0.00333333 |
| ASHFInt | W4 | Flavobacteriaceae_unclassified | 0.00333333 |
| ASHFInt | W8 | Flavobacteriaceae_unclassified | 0.00333333 |
| ASHFInt | W12 | Fusicatenibacter | 0.00333333 |
| ASHFInt | W4 | Helicobacter | 0.00333333 |
| ASHFInt | W12 | Incertae_Sedis | 0.00333333 |
| ASHFInt | W8 | Isosphaera | 0.00333333 |
| ASHFInt | W4 | JG30-KF-CM45_ge | 0.00333333 |
| ASHFInt | W8 | Lachnospiraceae_UCG-010 | 0.00333333 |
| ASHFInt | W12 | Lactobacillaceae_unclassified | 0.00333333 |
| ASHFInt | W8 | Leuconostoc | 0.00333333 |
| ASHFInt | W4 | Marmoricola | 0.00333333 |
| ASHFInt | W8 | Massilia | 0.00333333 |
| ASHFInt | W8 | Mycobacterium | 0.00333333 |
| ASHFInt | W12 | Nocardiaceae_unclassified | 0.00333333 |
| ASHFInt | W4 | Nocardioides | 0.00333333 |
| ASHFInt | Fresh | Nocardiopsis | 0.00333333 |
| ASHFInt | W8 | Oscillibacter | 0.00333333 |
| ASHFInt | W8 | Paenalcaligenes | 0.00333333 |
| ASHFInt | W12 | Paenibacillaceae_unclassified | 0.00333333 |
| ASHFInt | W12 | Pajaroellobacter | 0.00333333 |
| ASHFInt | W8 | Patulibacter | 0.00333333 |
| ASHFInt | W8 | Pedobacter | 0.00333333 |

| | | | |
|---|---|---|---|
| ASHFInt | W12 | Peptostreptococcales-Tissierellales_unclassifie | 0.00333333 |
| ASHFInt | W4 | Peredibacter | 0.00333333 |
| ASHFInt | W12 | Planococcaceae_unclassified | 0.00333333 |
| ASHFInt | W8 | Planococcaceae_unclassified | 0.00333333 |
| ASHFInt | W4 | Pseudactinotalea | 0.00333333 |
| ASHFInt | W8 | Psychrobacter | 0.00333333 |
| ASHFInt | W12 | RF39_ge | 0.00333333 |
| ASHFInt | W8 | RF39_ge | 0.00333333 |
| ASHFInt | W8 | Rhodanobacteraceae_unclassified | 0.00333333 |
| ASHFInt | W4 | Rhodococcus | 0.00333333 |
| ASHFInt | W12 | Rickettsia | 0.00333333 |
| ASHFInt | W8 | Rs-K70_termite_group_ge | 0.00333333 |
| ASHFInt | W12 | Sanguibacter-Flavimobilis | 0.00333333 |
| ASHFInt | W8 | Sporosarcina | 0.00333333 |
| ASHFInt | W12 | Streptococcus | 0.00333333 |
| ASHFInt | W12 | Streptomyces | 0.00333333 |
| ASHFInt | W4 | Streptomyces | 0.00333333 |
| ASHFInt | W12 | Terrisporobacter | 0.00333333 |
| ASHFInt | W8 | Tomitella | 0.00333333 |
| ASHFInt | W12 | Tuzzerella | 0.00333333 |
| ASHFInt | W12 | Tyzzerella | 0.00333333 |
| ASHFInt | W12 | WPS-2_ge | 0.00333333 |
| ASHFInt | W8 | WPS-2_ge | 0.00333333 |
| ASHFInt | W12 | vadinHA49_ge | 0.00333333 |
| ASHFInt | Fresh | Gordonia | 0.00333333 |
| ASHFInt | Fresh | Pasteurellaceae_unclassified | 0.00333333 |
| ASHFInt | Fresh | Phyllobacterium | 0.00333333 |
| ASHFInt | Fresh | Vagococcus | 0.00333333 |
| ASHFInt | Fresh | 67-14_ge | 0.0025 |

| | | | |
|---|---|---|---|
| ASHFInt | Fresh | Actinobacteria_unclassified | 0.0025 |
| ASHFInt | Fresh | Anaerovoracaceae_unclassified | 0.0025 |
| ASHFInt | Fresh | Brevibacterium | 0.0025 |
| ASHFInt | Fresh | Candidatus_Soleaferrea | 0.0025 |
| ASHFInt | Fresh | Georgenia | 0.0025 |
| ASHFInt | Fresh | Pseudogracilibacillus | 0.0025 |
| ASHFInt | Fresh | RF39_ge | 0.0025 |
| ASHFInt | Fresh | Acetobacteraceae_unclassified | 0.0025 |
| ASHFInt | Fresh | Actinobacteriota_unclassified | 0.0025 |
| ASHFInt | Fresh | Apibacter | 0.0025 |
| ASHFInt | Fresh | Gluconobacter | 0.0025 |
| ASHFInt | Fresh | Helicobacteraceae_unclassified | 0.0025 |
| ASHFInt | Fresh | Incertae_Sedis | 0.0025 |
| ASHFInt | Fresh | Lacticaseibacillus | 0.0025 |
| ASHFInt | Fresh | Nocardioidaceae_unclassified | 0.0025 |
| ASHFInt | Fresh | Sporosarcina | 0.0025 |
| ASHFInt | Fresh | Streptococcus | 0.0025 |
| ASHFInt | Fresh | Dickeya | 0.0025 |
| ASHFInt | W4 | 67-14_ge | 0.00166667 |
| ASHFInt | W4 | Actinocatenispora | 0.00166667 |
| ASHFInt | W4 | Actinospica | 0.00166667 |
| ASHFInt | W4 | Asaia | 0.00166667 |
| ASHFInt | Fresh | Beijerinckiaceae_unclassified | 0.00166667 |
| ASHFInt | Fresh | Brevinema | 0.00166667 |
| ASHFInt | W4 | Burkholderia-Caballeronia-Paraburkholderia | 0.00166667 |
| ASHFInt | W4 | Catenulispora | 0.00166667 |
| ASHFInt | W4 | Cohnella | 0.00166667 |
| ASHFInt | W4 | Comamonadaceae_unclassified | 0.00166667 |
| ASHFInt | W4 | Crossiella | 0.00166667 |

| | | | |
|---|---|---|---|
| ASHFInt | W4 | Curtobacterium | 0.00166667 |
| ASHFInt | W4 | Flavobacteriales_unclassified | 0.00166667 |
| ASHFInt | W4 | Fusobacterium | 0.00166667 |
| ASHFInt | W4 | Gaiella | 0.00166667 |
| ASHFInt | W4 | Iamia | 0.00166667 |
| ASHFInt | W4 | Leifsonia | 0.00166667 |
| ASHFInt | W4 | Leucobacter | 0.00166667 |
| ASHFInt | W4 | Limibaculum | 0.00166667 |
| ASHFInt | W4 | Lysinibacillus | 0.00166667 |
| ASHFInt | W4 | Massilia | 0.00166667 |
| ASHFInt | W4 | Membranicola | 0.00166667 |
| ASHFInt | Fresh | Methylobacterium-Methylorubrum | 0.00166667 |
| ASHFInt | Fresh | Nakamurella | 0.00166667 |
| ASHFInt | W4 | Nitrososphaeraceae_ge | 0.00166667 |
| ASHFInt | W4 | Nitrososphaeraceae_unclassified | 0.00166667 |
| ASHFInt | W4 | Paenibacillus | 0.00166667 |
| ASHFInt | W4 | Pectobacteriaceae_unclassified | 0.00166667 |
| ASHFInt | W4 | Pedobacter | 0.00166667 |
| ASHFInt | W4 | Planomonospora | 0.00166667 |
| ASHFInt | Fresh | Pseudomonas | 0.00166667 |
| ASHFInt | W4 | Pusillimonas | 0.00166667 |
| ASHFInt | W4 | Rs-K70_termite_group_ge | 0.00166667 |
| ASHFInt | W4 | Sarcina | 0.00166667 |
| ASHFInt | W4 | Solitalea | 0.00166667 |
| ASHFInt | W4 | Stenotrophomonas | 0.00166667 |
| ASHFInt | W4 | Streptomycetaceae_unclassified | 0.00166667 |
| ASHFInt | W4 | Thermoleophilia_unclassified | 0.00166667 |
| ASHFInt | W4 | Xanthomonadaceae_unclassified | 0.00166667 |
| ASHFInt | Fresh | 1174-901-12 | 0.00166667 |

| | | | |
|---|---|---|---|
| ASHFInt | Fresh | Acidiphilium | 0.00166667 |
| ASHFInt | Fresh | Aequorivita | 0.00166667 |
| ASHFInt | Fresh | Allorhizobium-Neorhizobium-Pararhizobium- | 0.00166667 |
| ASHFInt | Fresh | Brachybacterium | 0.00166667 |
| ASHFInt | Fresh | Catenulispora | 0.00166667 |
| ASHFInt | Fresh | Chitinophagaceae_unclassified | 0.00166667 |
| ASHFInt | Fresh | Dietzia | 0.00166667 |
| ASHFInt | Fresh | Dyadobacter | 0.00166667 |
| ASHFInt | Fresh | Exiguobacterium | 0.00166667 |
| ASHFInt | Fresh | Labilithrix | 0.00166667 |
| ASHFInt | Fresh | Limibaculum | 0.00166667 |
| ASHFInt | Fresh | Paludicola | 0.00166667 |
| ASHFInt | Fresh | Patulibacter | 0.00166667 |
| ASHFInt | Fresh | Phormidium_IAM_M-71 | 0.00166667 |
| ASHFInt | Fresh | Proteus | 0.00166667 |
| ASHFInt | Fresh | Romboutsia | 0.00166667 |
| ASHFInt | Fresh | Rummeliibacillus | 0.00166667 |
| ASHFInt | Fresh | Stenotrophomonas | 0.00166667 |
| ASHFInt | Fresh | Synergistaceae_unclassified | 0.00166667 |
| ASHFInt | Fresh | Terrisporobacter | 0.00166667 |
| ASHFInt | Fresh | Verticiella | 0.00166667 |
| ASHFInt | Fresh | 966-1 | 0.00083333 |
| ASHFInt | Fresh | Bradymonadaceae_ge | 0.00083333 |
| ASHFInt | Fresh | Brucella | 0.00083333 |
| ASHFInt | Fresh | Budvicia | 0.00083333 |
| ASHFInt | Fresh | Burkholderiales_unclassified | 0.00083333 |
| ASHFInt | Fresh | Cyanobium_PCC-6307 | 0.00083333 |
| ASHFInt | Fresh | Desulfovibrio | 0.00083333 |
| ASHFInt | Fresh | Desulfuromonadia_unclassified | 0.00083333 |

| | | | |
|---|---|---|---|
| ASHFInt | Fresh | Edwardsiella | 0.00083333 |
| ASHFInt | Fresh | Erysipelatoclostridiaceae_unclassified | 0.00083333 |
| ASHFInt | Fresh | Erysipelatoclostridium | 0.00083333 |
| ASHFInt | Fresh | Family_XI_unclassified | 0.00083333 |
| ASHFInt | Fresh | Gemmobacter | 0.00083333 |
| ASHFInt | Fresh | Gottschalkia | 0.00083333 |
| ASHFInt | Fresh | Hyphomicrobiaceae_unclassified | 0.00083333 |
| ASHFInt | Fresh | IMCC26207 | 0.00083333 |
| ASHFInt | Fresh | Intrasporangiaceae_unclassified | 0.00083333 |
| ASHFInt | Fresh | Isosphaeraceae_unclassified | 0.00083333 |
| ASHFInt | Fresh | Legionella | 0.00083333 |
| ASHFInt | Fresh | Microbacterium | 0.00083333 |
| ASHFInt | Fresh | Microtrichales_unclassified | 0.00083333 |
| ASHFInt | Fresh | Natronomonas | 0.00083333 |
| ASHFInt | Fresh | Novosphingobium | 0.00083333 |
| ASHFInt | Fresh | Peptostreptococcales-Tissierellales_unclassifie | 0.00083333 |
| ASHFInt | Fresh | Pragia | 0.00083333 |
| ASHFInt | Fresh | Pseudomonadales_unclassified | 0.00083333 |
| ASHFInt | Fresh | Pseudonocardiaceae_unclassified | 0.00083333 |
| ASHFInt | Fresh | Smaragdicoccus | 0.00083333 |
| ASHFInt | Fresh | TM7a | 0.00083333 |
| ASHFInt | Fresh | Taibaiella | 0.00083333 |
| ASHFInt | Fresh | Thermoleophilia_unclassified | 0.00083333 |
| ASHFInt | Fresh | Tissierella | 0.00083333 |
| ASHFInt | Fresh | Tsukamurella | 0.00083333 |
| ASHFInt | Fresh | Williamsia | 0.00083333 |
| ASHFInt | Fresh | Xanthomonas | 0.00083333 |
| ASHFInt | Fresh | Yersiniaceae_unclassified | 0.00083333 |
| WMCC | Fresh | Clostridiaceae_unclassified | 31.6576923 |

| | | | |
|---|---|---|---|
| WMCC | W4 | Massilia | 25.2283333 |
| WMCC | W12 | Nocardiaceae_unclassified | 24.0166667 |
| WMCC | Fresh | Enterococcus | 15.1038462 |
| WMCC | W8 | Massilia | 11.9157143 |
| WMCC | W4 | Pseudomonas | 11.5566667 |
| WMCC | W8 | Rhodococcus | 11.2485714 |
| WMCC | W12 | Massilia | 11.11 |
| WMCC | Fresh | Candidatus_Arthromitus | 9.18 |
| WMCC | Fresh | Lactococcus | 8.73692308 |
| WMCC | W8 | Nocardiaceae_unclassified | 8.24714286 |
| WMCC | W4 | Chryseobacterium | 7.63833333 |
| WMCC | W12 | Rhodococcus | 7.35666667 |
| WMCC | W4 | Nocardiaceae_unclassified | 7.13 |
| WMCC | W4 | Rhodococcus | 6.99833333 |
| WMCC | W8 | Pseudomonas | 6.78 |
| WMCC | Fresh | Aeromonas | 6.18923077 |
| WMCC | Fresh | Plesiomonas | 5.76615385 |
| WMCC | W4 | Sphingobacterium | 5.70333333 |
| WMCC | W12 | Pedobacter | 5.11666667 |
| WMCC | W8 | Pedobacter | 5.01142857 |
| WMCC | W12 | Paenarthrobacter | 4.93666667 |
| WMCC | W8 | Chryseobacterium | 4.87857143 |
| WMCC | W4 | Enterobacteriaceae_unclassified | 4.68333333 |
| WMCC | W8 | Sphingobacterium | 4.44 |
| WMCC | Fresh | Candidatus_Rhabdochlamydia | 4.38461538 |
| WMCC | W4 | Stenotrophomonas | 4.29666667 |
| WMCC | W8 | Stenotrophomonas | 3.95285714 |
| WMCC | Fresh | Clostridium_sensu_stricto_1 | 3.90538462 |
| WMCC | W12 | Enterobacteriaceae_unclassified | 3.65 |

| | | | |
|---|---|---|---|
| WMCC | W8 | Comamonadaceae_unclassified | 3.19 |
| WMCC | W8 | Rubritaleaceae_unclassified | 2.84571429 |
| WMCC | W4 | Alkanindiges | 2.83 |
| WMCC | W8 | Enterobacteriaceae_unclassified | 2.78 |
| WMCC | W12 | Burkholderia-Caballeronia-Paraburkholderia | 2.69333333 |
| WMCC | W4 | Enterobacterales_unclassified | 2.61833333 |
| WMCC | W12 | Sphingobacterium | 2.52 |
| WMCC | W4 | Paenibacillus | 2.49833333 |
| WMCC | W12 | Allorhizobium-Neorhizobium-Pararhizobium- | 2.46 |
| WMCC | W4 | Pedobacter | 2.34 |
| WMCC | W12 | Curtobacterium | 2.16666667 |
| WMCC | W8 | Allorhizobium-Neorhizobium-Pararhizobium- | 1.91285714 |
| WMCC | Fresh | Mycoplasma | 1.90923077 |
| WMCC | W8 | Paenibacillus | 1.89285714 |
| WMCC | W8 | Rhizobiaceae_unclassified | 1.84 |
| WMCC | W4 | Duganella | 1.83833333 |
| WMCC | W12 | Methylobacterium-Methylorubrum | 1.8 |
| WMCC | W12 | Pseudomonas | 1.77333333 |
| WMCC | Fresh | Desulfovibrio | 1.77230769 |
| WMCC | W12 | Rhizobiaceae_unclassified | 1.76666667 |
| WMCC | W4 | Curtobacterium | 1.72333333 |
| WMCC | Fresh | Hafnia-Obesumbacterium | 1.64615385 |
| WMCC | W4 | Comamonadaceae_unclassified | 1.64333333 |
| WMCC | W8 | Curtobacterium | 1.59857143 |
| WMCC | W12 | Acinetobacter | 1.54333333 |
| WMCC | W12 | Chryseobacterium | 1.54333333 |
| WMCC | W4 | Acinetobacter | 1.525 |
| WMCC | W12 | Leifsonia | 1.51666667 |
| WMCC | W8 | Patulibacter | 1.50142857 |

| | | | |
|---|---|---|---|
| WMCC | W8 | Paenarthrobacter | 1.47857143 |
| WMCC | W8 | Acinetobacter | 1.43 |
| WMCC | Fresh | Enterobacteriaceae_unclassified | 1.37615385 |
| WMCC | W12 | Mucilaginibacter | 1.31 |
| WMCC | W12 | Luteibacter | 1.30666667 |
| WMCC | W8 | Sphingomonas | 1.25142857 |
| WMCC | W8 | Streptomyces | 1.18142857 |
| WMCC | W8 | Alkanindiges | 1.16714286 |
| WMCC | W12 | Sphingomonas | 1.12 |
| WMCC | W12 | Stenotrophomonas | 1.08666667 |
| WMCC | W12 | Paenibacillus | 1.08 |
| WMCC | W12 | Patulibacter | 1.04666667 |
| WMCC | W12 | Enterobacterales_unclassified | 1.01666667 |
| WMCC | Fresh | Providencia | 0.93846154 |
| WMCC | Fresh | Peptostreptococcaceae_unclassified | 0.92384615 |
| WMCC | W8 | Chitinophaga | 0.90571429 |
| WMCC | W12 | Chitinibacteraceae_unclassified | 0.86666667 |
| WMCC | W8 | Micrococcaceae_unclassified | 0.86571429 |
| WMCC | W8 | Carnobacterium | 0.84428571 |
| WMCC | W12 | Carnobacterium | 0.82 |
| WMCC | W4 | Micrococcaceae_unclassified | 0.72333333 |
| WMCC | W8 | Leifsonia | 0.71571429 |
| WMCC | W4 | Streptomyces | 0.69166667 |
| WMCC | W12 | Comamonadaceae_unclassified | 0.68 |
| WMCC | W4 | Paenarthrobacter | 0.68 |
| WMCC | W12 | Abditibacterium | 0.67333333 |
| WMCC | Fresh | Rickettsiella | 0.66461538 |
| WMCC | W12 | Chitinophaga | 0.64333333 |
| WMCC | W4 | Enterococcus | 0.64333333 |

| | | | |
|---|---|---|---|
| WMCC | W8 | Methylobacterium-Methylorubrum | 0.64 |
| WMCC | W12 | Micrococcaceae_unclassified | 0.64 |
| WMCC | W4 | Rhizobiaceae_unclassified | 0.62833333 |
| WMCC | W8 | Burkholderia-Caballeronia-Paraburkholderia | 0.61428571 |
| WMCC | W12 | Micromonosporaceae_unclassified | 0.6 |
| WMCC | W8 | Flavobacterium | 0.59857143 |
| WMCC | W4 | Allorhizobium-Neorhizobium-Pararhizobium- | 0.59833333 |
| WMCC | W8 | Enterobacterales_unclassified | 0.59 |
| WMCC | W12 | Nocardioides | 0.55 |
| WMCC | W12 | Xanthobacteraceae_unclassified | 0.54666667 |
| WMCC | Fresh | Mycoplasmataceae_unclassified | 0.53384615 |
| WMCC | W8 | Duganella | 0.52857143 |
| WMCC | Fresh | Alcaligenaceae_ge | 0.50538462 |
| WMCC | W8 | Luteibacter | 0.48142857 |
| WMCC | W8 | Psychrobacter | 0.48142857 |
| WMCC | W12 | Williamsia | 0.45 |
| WMCC | W8 | Mucilaginibacter | 0.45 |
| WMCC | W8 | Aeromicrobium | 0.43142857 |
| WMCC | W8 | Dyadobacter | 0.42428571 |
| WMCC | W12 | Kocuria | 0.41666667 |
| WMCC | W12 | Mycobacterium | 0.40666667 |
| WMCC | Fresh | Bacillaceae_unclassified | 0.40615385 |
| WMCC | Fresh | Enterobacterales_unclassified | 0.40461538 |
| WMCC | W4 | Alcaligenaceae_unclassified | 0.40333333 |
| WMCC | W12 | Rhodopseudomonas | 0.39333333 |
| WMCC | W8 | Carnobacteriaceae_unclassified | 0.39 |
| WMCC | W12 | Sphingobacteriaceae_unclassified | 0.37666667 |
| WMCC | W4 | Sphingomonas | 0.36 |
| WMCC | W4 | Burkholderia-Caballeronia-Paraburkholderia | 0.345 |

| | | | |
|---|---|---|---|
| WMCC | Fresh | Nosocomiicoccus | 0.34230769 |
| WMCC | W8 | Brevundimonas | 0.34142857 |
| WMCC | Fresh | Salinisphaera | 0.33846154 |
| WMCC | W12 | Actinobacteria_unclassified | 0.32333333 |
| WMCC | W12 | Azospirillum | 0.31333333 |
| WMCC | W12 | Alcaligenaceae_unclassified | 0.31 |
| WMCC | W8 | Cohnella | 0.31 |
| WMCC | W8 | Nocardioides | 0.30571429 |
| WMCC | W12 | Oligoflexus | 0.28333333 |
| WMCC | W4 | Leifsonia | 0.28166667 |
| WMCC | W8 | Nosocomiicoccus | 0.27714286 |
| WMCC | W4 | Methylobacterium-Methylorubrum | 0.27666667 |
| WMCC | W8 | Alcaligenaceae_unclassified | 0.26714286 |
| WMCC | W12 | Caulobacter | 0.26666667 |
| WMCC | W8 | uncultured | 0.26571429 |
| WMCC | W8 | Devosia | 0.26 |
| WMCC | W4 | Micrococcales_unclassified | 0.26 |
| WMCC | W12 | Dyadobacter | 0.25666667 |
| WMCC | Fresh | Paeniclostridium | 0.25615385 |
| WMCC | W12 | Micrococcales_unclassified | 0.25 |
| WMCC | W12 | Sphingomonadaceae_unclassified | 0.25 |
| WMCC | W8 | Micrococcales_unclassified | 0.24571429 |
| WMCC | W8 | Rhodopseudomonas | 0.24428571 |
| WMCC | W8 | Micromonosporaceae_unclassified | 0.24428571 |
| WMCC | W8 | Clostridiaceae_unclassified | 0.24 |
| WMCC | W4 | Nocardioides | 0.23833333 |
| WMCC | W8 | Chitinibacteraceae_unclassified | 0.22428571 |
| WMCC | W12 | Bosea | 0.22333333 |
| WMCC | W4 | Carnobacterium | 0.21833333 |

| | | | |
|---|---|---|---|
| WMCC | Fresh | Ruminococcaceae_unclassified | 0.21307692 |
| WMCC | Fresh | Ureaplasma | 0.21076923 |
| WMCC | W4 | Brevundimonas | 0.20666667 |
| WMCC | W12 | Devosiaceae_unclassified | 0.20333333 |
| WMCC | W8 | uncultured_ge | 0.20285714 |
| WMCC | W12 | Phenylobacterium | 0.19333333 |
| WMCC | W8 | Leucobacter | 0.19285714 |
| WMCC | W4 | Luteibacter | 0.185 |
| WMCC | W4 | Mucilaginibacter | 0.18333333 |
| WMCC | W8 | Dietzia | 0.17714286 |
| WMCC | W12 | uncultured | 0.16666667 |
| WMCC | Fresh | Cetobacterium | 0.16461538 |
| WMCC | W8 | Enterococcus | 0.15857143 |
| WMCC | W8 | Actinobacteria_unclassified | 0.15571429 |
| WMCC | W8 | Sphingomonadaceae_unclassified | 0.15571429 |
| WMCC | W12 | Roseomonas | 0.15333333 |
| WMCC | W12 | Kaistia | 0.15 |
| WMCC | W4 | Oxalobacteraceae_unclassified | 0.14666667 |
| WMCC | W12 | Nocardiaceae_ge | 0.14 |
| WMCC | W12 | Spirosoma | 0.14 |
| WMCC | W8 | Oxalobacteraceae_unclassified | 0.13857143 |
| WMCC | W8 | Taibaiella | 0.13571429 |
| WMCC | W8 | Azospirillum | 0.13428571 |
| WMCC | W12 | Actinoplanes | 0.13333333 |
| WMCC | Fresh | Tyzzerella | 0.13153846 |
| WMCC | W12 | Beijerinckiaceae_unclassified | 0.13 |
| WMCC | W12 | Burkholderiales_unclassified | 0.13 |
| WMCC | W4 | Cohnella | 0.13 |
| WMCC | W12 | Phaselicystis | 0.13 |

| | | | |
|---|---|---|---|
| WMCC | W8 | Kaistia | 0.12857143 |
| WMCC | W8 | Fluviicola | 0.12857143 |
| WMCC | W8 | Bacteria_unclassified | 0.12857143 |
| WMCC | W4 | Hymenobacter | 0.12833333 |
| WMCC | W8 | Aureimonas | 0.12714286 |
| WMCC | W12 | Alkanindiges | 0.12666667 |
| WMCC | W12 | Deinococcus | 0.12666667 |
| WMCC | W8 | Hymenobacter | 0.12571429 |
| WMCC | W8 | Candidatus_Udaeobacter | 0.12428571 |
| WMCC | W12 | Taibaiella | 0.12333333 |
| WMCC | W12 | Nakamurella | 0.12333333 |
| WMCC | W8 | Roseomonas | 0.12142857 |
| WMCC | W12 | Aureimonas | 0.12 |
| WMCC | W12 | Cohnella | 0.12 |
| WMCC | W8 | Rummeliibacillus | 0.11714286 |
| WMCC | W12 | Jatrophihabitans | 0.11666667 |
| WMCC | W12 | Novosphingobium | 0.11 |
| WMCC | Fresh | Pseudomonas | 0.10846154 |
| WMCC | W4 | Patulibacter | 0.10666667 |
| WMCC | W8 | Bosea | 0.10571429 |
| WMCC | W8 | Mycobacterium | 0.10571429 |
| WMCC | Fresh | Morganella | 0.10307692 |
| WMCC | W8 | Sphingobacteriaceae_unclassified | 0.10285714 |
| WMCC | W8 | Williamsia | 0.10285714 |
| WMCC | Fresh | Lachnoclostridium | 0.09923077 |
| WMCC | W8 | Ochrobactrum | 0.09714286 |
| WMCC | W12 | Microbacteriaceae_unclassified | 0.09666667 |
| WMCC | W12 | Cellvibrio | 0.09666667 |
| WMCC | W12 | Rubritaleaceae_unclassified | 0.09666667 |

| | | | |
|---|---|---|---|
| WMCC | W8 | Luteolibacter | 0.09428571 |
| WMCC | W12 | Rummeliibacillus | 0.09333333 |
| WMCC | W12 | Arthrobacter | 0.09 |
| WMCC | W12 | Bacteria_unclassified | 0.09 |
| WMCC | W4 | Rummeliibacillus | 0.09 |
| WMCC | W4 | Exiguobacterium | 0.08833333 |
| WMCC | W8 | Novosphingobium | 0.08714286 |
| WMCC | W8 | Caulobacter | 0.08714286 |
| WMCC | W8 | Nakamurella | 0.08714286 |
| WMCC | W12 | Belnapia | 0.08666667 |
| WMCC | W4 | Chitinophaga | 0.08666667 |
| WMCC | W8 | Abditibacterium | 0.08571429 |
| WMCC | Fresh | Aquabacterium | 0.08461538 |
| WMCC | W12 | 1174-901-12 | 0.08333333 |
| WMCC | W12 | Hymenobacter | 0.08333333 |
| WMCC | W4 | Flavobacterium | 0.08333333 |
| WMCC | W8 | Bradyrhizobium | 0.08285714 |
| WMCC | W8 | Microbacteriaceae_unclassified | 0.08142857 |
| WMCC | W8 | Solirubrobacter | 0.07857143 |
| WMCC | W4 | Burkholderiales_unclassified | 0.07666667 |
| WMCC | W4 | Dyadobacter | 0.07666667 |
| WMCC | W12 | Streptomyces | 0.07666667 |
| WMCC | Fresh | Lactobacillales_unclassified | 0.07615385 |
| WMCC | Fresh | Paraclostridium | 0.07538462 |
| WMCC | W4 | Ochrobactrum | 0.075 |
| WMCC | W4 | Pseudomonadaceae_unclassified | 0.07333333 |
| WMCC | W4 | Nocardiaceae_ge | 0.07333333 |
| WMCC | W8 | Janthinobacterium | 0.07285714 |
| WMCC | Fresh | Leuconostoc | 0.07230769 |

| | | | |
|---|---|---|---|
| WMCC | W8 | Planococcaceae_unclassified | 0.07142857 |
| WMCC | W12 | Bdellovibrio | 0.07 |
| WMCC | W12 | Conexibacter | 0.07 |
| WMCC | W8 | Aminobacter | 0.06857143 |
| WMCC | W8 | Chthoniobacter | 0.06857143 |
| WMCC | W4 | Firmicutes_unclassified | 0.06833333 |
| WMCC | W8 | Marmoricola | 0.06714286 |
| WMCC | W8 | Xanthobacteraceae_unclassified | 0.06714286 |
| WMCC | Fresh | Unknown_Family_unclassified | 0.06692308 |
| WMCC | W12 | Fimbriimonadaceae_ge | 0.06666667 |
| WMCC | W4 | Staphylococcaceae_unclassified | 0.065 |
| WMCC | Fresh | Staphylococcaceae_unclassified | 0.06461538 |
| WMCC | W8 | Spirosoma | 0.06428571 |
| WMCC | W4 | Chitinibacteraceae_unclassified | 0.06333333 |
| WMCC | W4 | Xanthomonas | 0.06166667 |
| WMCC | W4 | Gammaproteobacteria_unclassified | 0.06 |
| WMCC | W12 | Rhodanobacteraceae_unclassified | 0.06 |
| WMCC | W8 | Bacteriovorax | 0.06 |
| WMCC | W4 | Microbacteriaceae_unclassified | 0.05833333 |
| WMCC | W8 | Sphingobium | 0.05714286 |
| WMCC | W8 | 67-14_ge | 0.05714286 |
| WMCC | W4 | Devosia | 0.05666667 |
| WMCC | W4 | Kocuria | 0.05666667 |
| WMCC | W12 | Leucobacter | 0.05666667 |
| WMCC | Fresh | Lachnospiraceae_unclassified | 0.05615385 |
| WMCC | W8 | Gammaproteobacteria_unclassified | 0.05571429 |
| WMCC | W4 | Clostridiaceae_unclassified | 0.055 |
| WMCC | W8 | Burkholderiales_unclassified | 0.05428571 |
| WMCC | W8 | Flavobacteriaceae_unclassified | 0.05428571 |

| | | | |
|---|---|---|---|
| WMCC | W8 | Pseudomonadaceae_unclassified | 0.05428571 |
| WMCC | W12 | Arcticibacter | 0.05333333 |
| WMCC | W12 | Chthoniobacter | 0.05333333 |
| WMCC | W12 | Solirubrobacter | 0.05333333 |
| WMCC | W8 | Bdellovibrio | 0.05285714 |
| WMCC | Fresh | Orbus | 0.05230769 |
| WMCC | Fresh | Rhodococcus | 0.05230769 |
| WMCC | W8 | Nocardiaceae_ge | 0.05142857 |
| WMCC | W8 | IMCC26256_ge | 0.05142857 |
| WMCC | W12 | Acetobacteraceae_unclassified | 0.05 |
| WMCC | W12 | Rurimicrobium | 0.05 |
| WMCC | W12 | Sphingobium | 0.05 |
| WMCC | W12 | Tepidisphaerales_unclassified | 0.05 |
| WMCC | W8 | Phenylobacterium | 0.04857143 |
| WMCC | W4 | Leucobacter | 0.04833333 |
| WMCC | W8 | Edaphobaculum | 0.04714286 |
| WMCC | W12 | Fluviicola | 0.04666667 |
| WMCC | W4 | Micromonosporaceae_unclassified | 0.04666667 |
| WMCC | W12 | Psychroglaciecola | 0.04666667 |
| WMCC | W8 | Kocuria | 0.04571429 |
| WMCC | W8 | Rhizobiales_unclassified | 0.04571429 |
| WMCC | W4 | Actinobacteria_unclassified | 0.045 |
| WMCC | W8 | Actinoplanes | 0.04428571 |
| WMCC | W8 | Aquabacterium | 0.04428571 |
| WMCC | W8 | Deinococcus | 0.04428571 |
| WMCC | W8 | Rhodanobacteraceae_unclassified | 0.04428571 |
| WMCC | W8 | WD2101_soil_group_ge | 0.04428571 |
| WMCC | Fresh | Helicobacter | 0.04384615 |
| WMCC | W12 | Corynebacteriales_unclassified | 0.04333333 |

| | | | |
|---|---|---|---|
| WMCC | W4 | Nosocomiicoccus | 0.04333333 |
| WMCC | Fresh | Pedobacter | 0.04230769 |
| WMCC | Fresh | Lyngbya_PCC-7419 | 0.04230769 |
| WMCC | W8 | Gaiella | 0.04142857 |
| WMCC | W12 | Oxalobacteraceae_unclassified | 0.04 |
| WMCC | W8 | Phaselicystis | 0.04 |
| WMCC | W12 | Rhizobiales_unclassified | 0.04 |
| WMCC | W12 | Solirubrobacteraceae_unclassified | 0.04 |
| WMCC | Fresh | Silvanigrella | 0.04 |
| WMCC | W8 | Jatrophihabitans | 0.03714286 |
| WMCC | W8 | Ferruginibacter | 0.03714286 |
| WMCC | Fresh | Epulopiscium | 0.03692308 |
| WMCC | W4 | Janthinobacterium | 0.03666667 |
| WMCC | W12 | Devosia | 0.03666667 |
| WMCC | W12 | Terriglobus | 0.03666667 |
| WMCC | Fresh | Bacillales_unclassified | 0.03615385 |
| WMCC | W8 | Devosiaceae_unclassified | 0.03571429 |
| WMCC | W8 | Candidatus_Arthromitus | 0.03428571 |
| WMCC | W8 | Staphylococcaceae_unclassified | 0.03428571 |
| WMCC | W8 | Blastococcus | 0.03428571 |
| WMCC | W8 | KD4-96_ge | 0.03428571 |
| WMCC | W12 | Enterococcus | 0.03333333 |
| WMCC | W8 | Kineococcus | 0.03285714 |
| WMCC | Fresh | uncultured | 0.03230769 |
| WMCC | W8 | Conexibacter | 0.03142857 |
| WMCC | Fresh | Firmicutes_unclassified | 0.03076923 |
| WMCC | W8 | 1174-901-12 | 0.03 |
| WMCC | W12 | Asticcacaulis | 0.03 |
| WMCC | Fresh | Curtobacterium | 0.03 |

| | | | |
|---|---|---|---|
| WMCC | Fresh | Stenotrophomonas | 0.02923077 |
| WMCC | W8 | Plesiomonas | 0.02857143 |
| WMCC | W8 | Solirubrobacteraceae_unclassified | 0.02857143 |
| WMCC | W4 | Azospirillum | 0.02833333 |
| WMCC | W4 | Williamsia | 0.02833333 |
| WMCC | Fresh | Staphylococcus | 0.02769231 |
| WMCC | W8 | Nocardia | 0.02714286 |
| WMCC | W12 | 0319-6G20_ge | 0.02666667 |
| WMCC | W4 | Candidatus_Arthromitus | 0.02666667 |
| WMCC | W12 | Caulobacteraceae_unclassified | 0.02666667 |
| WMCC | W12 | Clostridiaceae_unclassified | 0.02666667 |
| WMCC | W12 | Edaphobaculum | 0.02666667 |
| WMCC | W12 | Flavitalea | 0.02666667 |
| WMCC | W12 | TM7a | 0.02666667 |
| WMCC | W8 | 0319-6G20_ge | 0.02571429 |
| WMCC | W8 | Rhodoplanes | 0.02571429 |
| WMCC | Fresh | Comamonadaceae_unclassified | 0.02538462 |
| WMCC | W4 | Alcaligenaceae_ge | 0.025 |
| WMCC | W4 | Lachnospiraceae_unclassified | 0.025 |
| WMCC | W4 | Sphingobacteriaceae_unclassified | 0.025 |
| WMCC | Fresh | Massilia | 0.02461538 |
| WMCC | W8 | Oligoflexus | 0.02428571 |
| WMCC | W8 | Beijerinckiaceae_unclassified | 0.02428571 |
| WMCC | W8 | Belnapia | 0.02428571 |
| WMCC | W8 | Streptomycetaceae_unclassified | 0.02428571 |
| WMCC | Fresh | Bacteria_unclassified | 0.02384615 |
| WMCC | Fresh | Brevinema | 0.02384615 |
| WMCC | Fresh | Methylobacterium-Methylorubrum | 0.02384615 |
| WMCC | W12 | Candidatus_Protochlamydia | 0.02333333 |

| | | | |
|---|---|---|---|
| WMCC | W12 | Kineosporia | 0.02333333 |
| WMCC | W4 | Novosphingobium | 0.02333333 |
| WMCC | W12 | uncultured_ge | 0.02333333 |
| WMCC | W12 | Fimbriimonadaceae_unclassified | 0.02333333 |
| WMCC | Fresh | Snowella_0TU37S04 | 0.02307692 |
| WMCC | W8 | Caulobacteraceae_unclassified | 0.02285714 |
| WMCC | W8 | Frankiales_unclassified | 0.02285714 |
| WMCC | W8 | Paenibacillaceae_unclassified | 0.02285714 |
| WMCC | W8 | Gaiellales_unclassified | 0.02285714 |
| WMCC | W8 | Pseudonocardia | 0.02285714 |
| WMCC | W4 | Tyzzerella | 0.02166667 |
| WMCC | Fresh | Pasteurellaceae_unclassified | 0.02153846 |
| WMCC | Fresh | Patulibacter | 0.02153846 |
| WMCC | Fresh | Rickettsiaceae_unclassified | 0.02153846 |
| WMCC | W8 | Acetobacteraceae_unclassified | 0.02142857 |
| WMCC | W8 | Chitinophagaceae_unclassified | 0.02142857 |
| WMCC | W8 | Vicinamibacteraceae_ge | 0.02142857 |
| WMCC | Fresh | endosymbionts | 0.02076923 |
| WMCC | W4 | Aeromicrobium | 0.02 |
| WMCC | W12 | Bacteroidia_unclassified | 0.02 |
| WMCC | W12 | Lachnospiraceae_unclassified | 0.02 |
| WMCC | W12 | Marmoricola | 0.02 |
| WMCC | W4 | Moraxellaceae_unclassified | 0.02 |
| WMCC | W12 | Pseudomonadaceae_unclassified | 0.02 |
| WMCC | W12 | Psychrobacter | 0.02 |
| WMCC | W4 | Roseomonas | 0.02 |
| WMCC | W12 | Tepidisphaera | 0.02 |
| WMCC | W12 | Vampirovibrionaceae_ge | 0.02 |
| WMCC | W12 | Aeromicrobium | 0.02 |

| | | | |
|---|---|---|---|
| WMCC | W8 | Arcticibacter | 0.02 |
| WMCC | W8 | Blastocatellaceae_unclassified | 0.02 |
| WMCC | W12 | Chitinophagaceae_unclassified | 0.02 |
| WMCC | W8 | Ellin6055 | 0.02 |
| WMCC | W8 | Sericytochromatia_ge | 0.01857143 |
| WMCC | W8 | Xanthomonadaceae_unclassified | 0.01857143 |
| WMCC | W8 | Saccharimonadales_unclassified | 0.01857143 |
| WMCC | Fresh | Rhizobiaceae_unclassified | 0.01846154 |
| WMCC | W4 | Corynebacteriales_unclassified | 0.01833333 |
| WMCC | W4 | uncultured | 0.01833333 |
| WMCC | W8 | Enterorhabdus | 0.01714286 |
| WMCC | W8 | Intrasporangiaceae_unclassified | 0.01714286 |
| WMCC | W8 | Kineosporia | 0.01714286 |
| WMCC | W8 | Klenkia | 0.01714286 |
| WMCC | W8 | Microvirga | 0.01714286 |
| WMCC | W8 | Parafilimonas | 0.01714286 |
| WMCC | W8 | Psychroglaciecola | 0.01714286 |
| WMCC | W8 | Tepidisphaerales_unclassified | 0.01714286 |
| WMCC | W4 | Arcticibacter | 0.01666667 |
| WMCC | W4 | Lactococcus | 0.01666667 |
| WMCC | W4 | Staphylococcus | 0.01666667 |
| WMCC | W12 | Actinomycetospora | 0.01666667 |
| WMCC | W12 | Aminobacter | 0.01666667 |
| WMCC | W4 | Bacteria_unclassified | 0.01666667 |
| WMCC | W4 | Kineococcus | 0.01666667 |
| WMCC | Fresh | Chryseobacterium | 0.01615385 |
| WMCC | Fresh | Microcystis_PCC-7914 | 0.01615385 |
| WMCC | W8 | P3OB-42 | 0.01571429 |
| WMCC | W8 | Shinella | 0.01571429 |

| | | | |
|---|---|---|---|
| WMCC | Fresh | Corynebacterium | 0.01538462 |
| WMCC | Fresh | Gammaproteobacteria_unclassified | 0.01538462 |
| WMCC | Fresh | Streptococcaceae_unclassified | 0.01538462 |
| WMCC | Fresh | Lacticaseibacillus | 0.01538462 |
| WMCC | W4 | Mycobacterium | 0.015 |
| WMCC | W4 | Sphingomonadaceae_unclassified | 0.015 |
| WMCC | Fresh | Aureimonas | 0.01461538 |
| WMCC | Fresh | uncultured_ge | 0.01461538 |
| WMCC | Fresh | Sphingomonas | 0.01461538 |
| WMCC | W8 | Alphaproteobacteria_unclassified | 0.01428571 |
| WMCC | W8 | Burkholderiaceae_unclassified | 0.01428571 |
| WMCC | W8 | Corynebacteriales_unclassified | 0.01428571 |
| WMCC | W8 | Gordonia | 0.01428571 |
| WMCC | W8 | Pajaroellobacter | 0.01428571 |
| WMCC | W8 | Microtrichales_unclassified | 0.01428571 |
| WMCC | W8 | Nitrososphaeraceae_unclassified | 0.01428571 |
| WMCC | W8 | Peptostreptococcaceae_unclassified | 0.01428571 |
| WMCC | W8 | SC-I-84_ge | 0.01428571 |
| WMCC | W8 | TK10_ge | 0.01428571 |
| WMCC | W12 | Burkholderiaceae_unclassified | 0.01333333 |
| WMCC | W4 | Caulobacter | 0.01333333 |
| WMCC | W4 | Devosiaceae_unclassified | 0.01333333 |
| WMCC | W4 | Dietzia | 0.01333333 |
| WMCC | W12 | Gammaproteobacteria_unclassified | 0.01333333 |
| WMCC | W4 | Paenibacillaceae_unclassified | 0.01333333 |
| WMCC | W4 | Plesiomonas | 0.01333333 |
| WMCC | W4 | Psychrobacter | 0.01333333 |
| WMCC | W12 | Silvanigrella | 0.01333333 |
| WMCC | W4 | Spirosoma | 0.01333333 |

| | | | |
|---|---|---|---|
| WMCC | W12 | WD2101_soil_group_ge | 0.01333333 |
| WMCC | W12 | Klenkia | 0.01333333 |
| WMCC | W8 | Legionella | 0.01285714 |
| WMCC | W8 | Reyranella | 0.01285714 |
| WMCC | W8 | Silvanigrella | 0.01285714 |
| WMCC | W8 | Pir4_lineage | 0.01285714 |
| WMCC | Fresh | Actinomycetospora | 0.01230769 |
| WMCC | Fresh | Bacilli_unclassified | 0.01230769 |
| WMCC | Fresh | Atopostipes | 0.01230769 |
| WMCC | W4 | Aquabacterium | 0.01166667 |
| WMCC | W4 | Bacilli_unclassified | 0.01166667 |
| WMCC | W4 | Xanthobacteraceae_unclassified | 0.01166667 |
| WMCC | W4 | Actinoplanes | 0.01166667 |
| WMCC | W4 | Fluviicola | 0.01166667 |
| WMCC | Fresh | FukuN18_freshwater_group | 0.01153846 |
| WMCC | Fresh | Crossiella | 0.01153846 |
| WMCC | W8 | Gemmatimonas | 0.01142857 |
| WMCC | W8 | Kineosporiaceae_unclassified | 0.01142857 |
| WMCC | W8 | Segetibacter | 0.01142857 |
| WMCC | W8 | Lysinibacillus | 0.01142857 |
| WMCC | W12 | Chlamydiales_unclassified | 0.01 |
| WMCC | W8 | Acidibacter | 0.01 |
| WMCC | W12 | Alcaligenaceae_ge | 0.01 |
| WMCC | W8 | BIrii41_ge | 0.01 |
| WMCC | W4 | Bacillaceae_unclassified | 0.01 |
| WMCC | W4 | Bosea | 0.01 |
| WMCC | W8 | CL500-29_marine_group | 0.01 |
| WMCC | W12 | Dietzia | 0.01 |
| WMCC | W8 | Fimbriimonadaceae_ge | 0.01 |

| | | | |
|---|---|---|---|
| WMCC | W8 | Isosphaeraceae_unclassified | 0.01 |
| WMCC | W8 | JG30-KF-CM45_ge | 0.01 |
| WMCC | W12 | Kineococcus | 0.01 |
| WMCC | W12 | Micavibrio | 0.01 |
| WMCC | Fresh | Microbacteriaceae_unclassified | 0.01 |
| WMCC | W12 | Microvirga | 0.01 |
| WMCC | W4 | Oscillospiraceae_unclassified | 0.01 |
| WMCC | W12 | Pajaroellobacter | 0.01 |
| WMCC | W8 | Pirellula | 0.01 |
| WMCC | W8 | Polyangiaceae_unclassified | 0.01 |
| WMCC | W12 | Promicromonosporaceae_unclassified | 0.01 |
| WMCC | W12 | Shinella | 0.01 |
| WMCC | W12 | Tundrisphaera | 0.01 |
| WMCC | W4 | Xanthomonadaceae_unclassified | 0.01 |
| WMCC | W12 | Amantichitinum | 0.01 |
| WMCC | W12 | Corallococcus | 0.01 |
| WMCC | W8 | Firmicutes_unclassified | 0.01 |
| WMCC | W12 | Nubsella | 0.01 |
| WMCC | W12 | Ochrobactrum | 0.01 |
| WMCC | W8 | Puia | 0.01 |
| WMCC | W8 | Rurimicrobium | 0.01 |
| WMCC | W8 | Tomitella | 0.01 |
| WMCC | Fresh | Dysgonomonas | 0.01 |
| WMCC | Fresh | Streptomyces | 0.00923077 |
| WMCC | Fresh | Oscillospirales_unclassified | 0.00923077 |
| WMCC | Fresh | Synergistaceae_unclassified | 0.00923077 |
| WMCC | W8 | Bacillales_unclassified | 0.00857143 |
| WMCC | W8 | Bacilli_unclassified | 0.00857143 |
| WMCC | W8 | Fimbriimonadaceae_unclassified | 0.00857143 |

| | | | |
|---|---|---|---|
| WMCC | W8 | Flavisolibacter | 0.00857143 |
| WMCC | W8 | JGI_0001001-H03 | 0.00857143 |
| WMCC | W8 | Modestobacter | 0.00857143 |
| WMCC | W8 | Rhodobacteraceae_unclassified | 0.00857143 |
| WMCC | W8 | SM2D12_ge | 0.00857143 |
| WMCC | W8 | Thermoleophilia_unclassified | 0.00857143 |
| WMCC | W8 | Tundrisphaera | 0.00857143 |
| WMCC | W8 | Xanthomonas | 0.00857143 |
| WMCC | W8 | Cellulomonas | 0.00857143 |
| WMCC | W8 | Corallococcus | 0.00857143 |
| WMCC | W8 | MB-A2-108_ge | 0.00857143 |
| WMCC | W8 | Niabella | 0.00857143 |
| WMCC | Fresh | Micrococcaceae_unclassified | 0.00846154 |
| WMCC | Fresh | Beijerinckiaceae_unclassified | 0.00846154 |
| WMCC | Fresh | Duganella | 0.00846154 |
| WMCC | Fresh | Gluconobacter | 0.00846154 |
| WMCC | Fresh | Leifsonia | 0.00846154 |
| WMCC | Fresh | Terrisporobacter | 0.00846154 |
| WMCC | W4 | Burkholderiaceae_unclassified | 0.00833333 |
| WMCC | W4 | Saccharibacillus | 0.00833333 |
| WMCC | W4 | Aureimonas | 0.00833333 |
| WMCC | W4 | Hafnia-Obesumbacterium | 0.00833333 |
| WMCC | W4 | Rhodopseudomonas | 0.00833333 |
| WMCC | W4 | Sanguibacter-Flavimobilis | 0.00833333 |
| WMCC | W4 | Taibaiella | 0.00833333 |
| WMCC | Fresh | 1174-901-12 | 0.00769231 |
| WMCC | Fresh | Burkholderia-Caballeronia-Paraburkholderia | 0.00769231 |
| WMCC | Fresh | Psychrobacter | 0.00769231 |
| WMCC | Fresh | Alkanindiges | 0.00769231 |

| | | | |
|---|---|---|---|
| WMCC | Fresh | Micromonosporaceae_unclassified | 0.00769231 |
| WMCC | Fresh | Paenibacillus | 0.00769231 |
| WMCC | Fresh | Rhodocyclaceae_unclassified | 0.00769231 |
| WMCC | Fresh | Terrimicrobium | 0.00769231 |
| WMCC | W8 | Blastocatella | 0.00714286 |
| WMCC | W8 | Chlamydiales_unclassified | 0.00714286 |
| WMCC | W8 | Flavitalea | 0.00714286 |
| WMCC | W8 | Moraxellaceae_unclassified | 0.00714286 |
| WMCC | W8 | Paeniclostridium | 0.00714286 |
| WMCC | W8 | TM7a | 0.00714286 |
| WMCC | W8 | env.OPS_17_ge | 0.00714286 |
| WMCC | W8 | Alsobacter | 0.00714286 |
| WMCC | W8 | Bryobacter | 0.00714286 |
| WMCC | W8 | Candidatus_Koribacter | 0.00714286 |
| WMCC | W8 | Cryptosporangium | 0.00714286 |
| WMCC | W8 | Cupriavidus | 0.00714286 |
| WMCC | W8 | Exiguobacterium | 0.00714286 |
| WMCC | W8 | Gemmataceae_unclassified | 0.00714286 |
| WMCC | W8 | Hyphomicrobiaceae_unclassified | 0.00714286 |
| WMCC | W8 | Labrys | 0.00714286 |
| WMCC | W8 | Oryzihumus | 0.00714286 |
| WMCC | W8 | Pedomicrobium | 0.00714286 |
| WMCC | W8 | RB41 | 0.00714286 |
| WMCC | W8 | TRA3-20_ge | 0.00714286 |
| WMCC | Fresh | Bradyrhizobium | 0.00692308 |
| WMCC | Fresh | Candidatus_Udaeobacter | 0.00692308 |
| WMCC | Fresh | Kocuria | 0.00692308 |
| WMCC | Fresh | Neisseriaceae_unclassified | 0.00692308 |
| WMCC | Fresh | Zymobacter | 0.00692308 |

| | | | |
|---|---|---|---|
| WMCC | W4 | Abditibacterium | 0.00666667 |
| WMCC | W12 | Armatimonadales_ge | 0.00666667 |
| WMCC | W12 | Bacilli_unclassified | 0.00666667 |
| WMCC | W12 | Blastocatella | 0.00666667 |
| WMCC | W12 | Bradyrhizobium | 0.00666667 |
| WMCC | W12 | Chloroflexi_unclassified | 0.00666667 |
| WMCC | W4 | Conexibacter | 0.00666667 |
| WMCC | W12 | Cryptosporangium | 0.00666667 |
| WMCC | W12 | Duganella | 0.00666667 |
| WMCC | W12 | Firmicutes_unclassified | 0.00666667 |
| WMCC | W12 | Gordonia | 0.00666667 |
| WMCC | W4 | Kaistia | 0.00666667 |
| WMCC | W12 | Limnobacter | 0.00666667 |
| WMCC | W12 | Modestobacter | 0.00666667 |
| WMCC | W12 | Moraxellaceae_unclassified | 0.00666667 |
| WMCC | W12 | Nosocomiicoccus | 0.00666667 |
| WMCC | W12 | Oscillospiraceae_unclassified | 0.00666667 |
| WMCC | W12 | Paenibacillaceae_unclassified | 0.00666667 |
| WMCC | W4 | Paeniclostridium | 0.00666667 |
| WMCC | W4 | Rhodanobacteraceae_unclassified | 0.00666667 |
| WMCC | W12 | Rickettsiaceae_unclassified | 0.00666667 |
| WMCC | W12 | Saccharimonadales_unclassified | 0.00666667 |
| WMCC | W12 | Segetibacter | 0.00666667 |
| WMCC | Fresh | Brevundimonas | 0.00615385 |
| WMCC | Fresh | Chitinibacteraceae_unclassified | 0.00615385 |
| WMCC | Fresh | Luteibacter | 0.00615385 |
| WMCC | Fresh | Conexibacter | 0.00615385 |
| WMCC | Fresh | Cyanobium_PCC-6307 | 0.00615385 |
| WMCC | Fresh | Marmoricola | 0.00615385 |

| | | | |
|---|---|---|---|
| WMCC | Fresh | Spiroplasma | 0.00615385 |
| WMCC | W8 | Actinomycetospora | 0.00571429 |
| WMCC | W8 | Alcaligenaceae_ge | 0.00571429 |
| WMCC | W8 | Cellvibrio | 0.00571429 |
| WMCC | W8 | Clostridium_sensu_stricto_1 | 0.00571429 |
| WMCC | W8 | Eggerthellaceae_unclassified | 0.00571429 |
| WMCC | W8 | Nubsella | 0.00571429 |
| WMCC | W8 | RF39_ge | 0.00571429 |
| WMCC | W8 | Acidimicrobiia_unclassified | 0.00571429 |
| WMCC | W8 | Altererythrobacter | 0.00571429 |
| WMCC | W8 | Candidatus_Solibacter | 0.00571429 |
| WMCC | W8 | Chloroflexi_unclassified | 0.00571429 |
| WMCC | W8 | Longimicrobiaceae_ge | 0.00571429 |
| WMCC | W8 | Roseiarcus | 0.00571429 |
| WMCC | Fresh | Alphaproteobacteria_unclassified | 0.00538462 |
| WMCC | Fresh | Bartonella | 0.00538462 |
| WMCC | Fresh | IMCC26207 | 0.00538462 |
| WMCC | Fresh | Levilactobacillus | 0.00538462 |
| WMCC | Fresh | Planktothrix_NIVA-CYA_15 | 0.00538462 |
| WMCC | Fresh | Robinsoniella | 0.00538462 |
| WMCC | Fresh | Roseomonas | 0.00538462 |
| WMCC | W4 | 1174-901-12 | 0.005 |
| WMCC | W4 | Caulobacteraceae_unclassified | 0.005 |
| WMCC | W4 | Flavobacteriaceae_unclassified | 0.005 |
| WMCC | W4 | Kineosporia | 0.005 |
| WMCC | W4 | Peptostreptococcaceae_unclassified | 0.005 |
| WMCC | W4 | Planococcaceae_unclassified | 0.005 |
| WMCC | W4 | Xanthomonadales_unclassified | 0.005 |
| WMCC | Fresh | Erysipelatoclostridium | 0.00461538 |

| | | | |
|---|---|---|---|
| WMCC | Fresh | Cyanobacteriia_unclassified | 0.00461538 |
| WMCC | Fresh | Deefgea | 0.00461538 |
| WMCC | Fresh | Luteolibacter | 0.00461538 |
| WMCC | Fresh | Romboutsia | 0.00461538 |
| WMCC | Fresh | KD4-96_ge | 0.00461538 |
| WMCC | W8 | AKIW781_ge | 0.00428571 |
| WMCC | W8 | Acidothermus | 0.00428571 |
| WMCC | W8 | Aquicella | 0.00428571 |
| WMCC | W8 | Asticcacaulis | 0.00428571 |
| WMCC | W8 | Bacillaceae_unclassified | 0.00428571 |
| WMCC | W8 | Bacteroidia_unclassified | 0.00428571 |
| WMCC | W8 | Gemmata | 0.00428571 |
| WMCC | W8 | Gemmatimonadaceae_unclassified | 0.00428571 |
| WMCC | W8 | Hafnia-Obesumbacterium | 0.00428571 |
| WMCC | W8 | Ilumatobacter | 0.00428571 |
| WMCC | W8 | Ilumatobacteraceae_unclassified | 0.00428571 |
| WMCC | W8 | Nocardioidaceae_unclassified | 0.00428571 |
| WMCC | W8 | Pedosphaeraceae_ge | 0.00428571 |
| WMCC | W8 | Ruminococcus | 0.00428571 |
| WMCC | W8 | Saccharibacillus | 0.00428571 |
| WMCC | W8 | Singulisphaera | 0.00428571 |
| WMCC | W8 | Subgroup_7_ge | 0.00428571 |
| WMCC | W8 | Tyzzerella | 0.00428571 |
| WMCC | W8 | Brachybacterium | 0.00428571 |
| WMCC | W8 | Ellin517 | 0.00428571 |
| WMCC | W8 | Ellin6067 | 0.00428571 |
| WMCC | W8 | Microtrichaceae_unclassified | 0.00428571 |
| WMCC | W8 | Paracoccus | 0.00428571 |
| WMCC | W8 | Rhodomicrobium | 0.00428571 |

| | | | |
|---|---|---|---|
| WMCC | W8 | Steroidobacter | 0.00428571 |
| WMCC | Fresh | Erysipelotrichaceae_unclassified | 0.00384615 |
| WMCC | Fresh | Hungateiclostridiaceae_unclassified | 0.00384615 |
| WMCC | Fresh | Jatrophihabitans | 0.00384615 |
| WMCC | Fresh | Lysinibacillus | 0.00384615 |
| WMCC | Fresh | Nakamurella | 0.00384615 |
| WMCC | Fresh | Planococcaceae_unclassified | 0.00384615 |
| WMCC | Fresh | RsaHf231_ge | 0.00384615 |
| WMCC | Fresh | Sebaldella | 0.00384615 |
| WMCC | Fresh | Tomitella | 0.00384615 |
| WMCC | Fresh | Turicibacter | 0.00384615 |
| WMCC | Fresh | Burkholderiales_unclassified | 0.00384615 |
| WMCC | Fresh | Liquorilactobacillus | 0.00384615 |
| WMCC | Fresh | Phenylobacterium | 0.00384615 |
| WMCC | Fresh | Sphingobacterium | 0.00384615 |
| WMCC | Fresh | Vogesella | 0.00384615 |
| WMCC | W12 | Acidimicrobiia_unclassified | 0.00333333 |
| WMCC | W12 | Acidiphilium | 0.00333333 |
| WMCC | W12 | Acidothermus | 0.00333333 |
| WMCC | W12 | Alphaproteobacteria_unclassified | 0.00333333 |
| WMCC | W4 | Alphaproteobacteria_unclassified | 0.00333333 |
| WMCC | W12 | Amycolatopsis | 0.00333333 |
| WMCC | W12 | Aquabacterium | 0.00333333 |
| WMCC | W12 | Bacillaceae_unclassified | 0.00333333 |
| WMCC | W4 | Bacillales_unclassified | 0.00333333 |
| WMCC | W4 | Belnapia | 0.00333333 |
| WMCC | W12 | Bryobacter | 0.00333333 |
| WMCC | W12 | CL500-29_marine_group | 0.00333333 |
| WMCC | W12 | Candidatus_Finniella | 0.00333333 |

| | | | |
|---|---|---|---|
| WMCC | W12 | Cerasicoccus | 0.00333333 |
| WMCC | W12 | Chelatococcus | 0.00333333 |
| WMCC | W12 | Chthoniobacteraceae_unclassified | 0.00333333 |
| WMCC | W4 | Deinococcus | 0.00333333 |
| WMCC | W12 | Diplorickettsiaceae_unclassified | 0.00333333 |
| WMCC | W12 | Ellin6055 | 0.00333333 |
| WMCC | W12 | Filimonas | 0.00333333 |
| WMCC | W4 | Fimbriimonadaceae_ge | 0.00333333 |
| WMCC | W12 | Flavobacterium | 0.00333333 |
| WMCC | W12 | Frankiales_unclassified | 0.00333333 |
| WMCC | W12 | Gemmatimonadaceae_unclassified | 0.00333333 |
| WMCC | W12 | IMCC26256_ge | 0.00333333 |
| WMCC | W12 | JG30-KF-CM45_ge | 0.00333333 |
| WMCC | W4 | Lactobacillales_unclassified | 0.00333333 |
| WMCC | W12 | Luteolibacter | 0.00333333 |
| WMCC | W12 | Microbacterium | 0.00333333 |
| WMCC | W4 | Microbacterium | 0.00333333 |
| WMCC | W4 | Microvirga | 0.00333333 |
| WMCC | W12 | Parachlamydiaceae_unclassified | 0.00333333 |
| WMCC | W4 | Paraclostridium | 0.00333333 |
| WMCC | W12 | Parafilimonas | 0.00333333 |
| WMCC | W12 | Parviterribacter | 0.00333333 |
| WMCC | W12 | Plesiomonas | 0.00333333 |
| WMCC | W12 | Propionibacteriales_unclassified | 0.00333333 |
| WMCC | W4 | Proteobacteria_unclassified | 0.00333333 |
| WMCC | W12 | Pseudolabrys | 0.00333333 |
| WMCC | W12 | Raoultibacter | 0.00333333 |
| WMCC | W12 | Rhizorhapis | 0.00333333 |
| WMCC | W12 | Rhodovastum | 0.00333333 |

| | | | |
|---|---|---|---|
| WMCC | W12 | Saccharimonadaceae_unclassified | 0.00333333 |
| WMCC | W12 | Saccharimonadales_ge | 0.00333333 |
| WMCC | W12 | Schumannella | 0.00333333 |
| WMCC | W4 | Silvanigrella | 0.00333333 |
| WMCC | W12 | Sphingoaurantiacus | 0.00333333 |
| WMCC | W4 | Sphingobium | 0.00333333 |
| WMCC | W12 | Xanthomonadales_unclassified | 0.00333333 |
| WMCC | Fresh | Bacteriovorax | 0.00307692 |
| WMCC | Fresh | Chthoniobacter | 0.00307692 |
| WMCC | Fresh | Flavobacterium | 0.00307692 |
| WMCC | Fresh | Mycobacterium | 0.00307692 |
| WMCC | Fresh | Pseudonocardia | 0.00307692 |
| WMCC | Fresh | Sphingomonadaceae_unclassified | 0.00307692 |
| WMCC | Fresh | Actinoplanes | 0.00307692 |
| WMCC | Fresh | Alsobacter | 0.00307692 |
| WMCC | Fresh | Escherichia-Shigella | 0.00307692 |
| WMCC | Fresh | Gaiella | 0.00307692 |
| WMCC | Fresh | Incertae_Sedis | 0.00307692 |
| WMCC | Fresh | Methanobacterium | 0.00307692 |
| WMCC | Fresh | Paludicola | 0.00307692 |
| WMCC | Fresh | Pir4_lineage | 0.00307692 |
| WMCC | Fresh | Rhizobiales_unclassified | 0.00307692 |
| WMCC | Fresh | Solirubrobacter | 0.00307692 |
| WMCC | Fresh | Streptomycetaceae_unclassified | 0.00307692 |
| WMCC | Fresh | Fusobacterium | 0.00307692 |
| WMCC | Fresh | Sporosarcina | 0.00307692 |
| WMCC | W8 | Adlercreutzia | 0.00285714 |
| WMCC | W8 | Armatimonadales_ge | 0.00285714 |
| WMCC | W8 | Camelimonas | 0.00285714 |

| | | | |
|---|---|---|---|
| WMCC | W8 | Cyanobacteriia_unclassified | 0.00285714 |
| WMCC | W8 | DNF00809 | 0.00285714 |
| WMCC | W8 | Iamia | 0.00285714 |
| WMCC | W8 | Lacunisphaera | 0.00285714 |
| WMCC | W8 | Paeniglutamicibacter | 0.00285714 |
| WMCC | W8 | Rickettsiaceae_unclassified | 0.00285714 |
| WMCC | W8 | Xanthomonadales_unclassified | 0.00285714 |
| WMCC | W8 | 966-1 | 0.00285714 |
| WMCC | W8 | Actinobacteriota_unclassified | 0.00285714 |
| WMCC | W8 | Aurantisolimonas | 0.00285714 |
| WMCC | W8 | Chthoniobacterales_unclassified | 0.00285714 |
| WMCC | W8 | E1B-B3-114 | 0.00285714 |
| WMCC | W8 | Fimbriiglobus | 0.00285714 |
| WMCC | W8 | Fimbriimonas | 0.00285714 |
| WMCC | W8 | Geodermatophilaceae_unclassified | 0.00285714 |
| WMCC | W8 | Haliangium | 0.00285714 |
| WMCC | W8 | Hyphomicrobium | 0.00285714 |
| WMCC | W8 | LWQ8_ge | 0.00285714 |
| WMCC | W8 | Lactobacillales_unclassified | 0.00285714 |
| WMCC | W8 | Myxococcaceae_unclassified | 0.00285714 |
| WMCC | W8 | Opitutus | 0.00285714 |
| WMCC | W8 | PLTA13_ge | 0.00285714 |
| WMCC | W8 | Parcubacteria_unclassified | 0.00285714 |
| WMCC | W8 | Phyllobacterium | 0.00285714 |
| WMCC | W8 | Propionibacteriales_unclassified | 0.00285714 |
| WMCC | W8 | Pseudorhodoplanes | 0.00285714 |
| WMCC | W8 | R7C24_ge | 0.00285714 |
| WMCC | W8 | RBG-13-54-9_ge | 0.00285714 |
| WMCC | W8 | S0134_terrestrial_group_ge | 0.00285714 |

| | | | |
|---|---|---|---|
| WMCC | W8 | Saccharimonadaceae_unclassified | 0.00285714 |
| WMCC | W8 | Sanguibacter-Flavimobilis | 0.00285714 |
| WMCC | W8 | Solirubrobacterales_unclassified | 0.00285714 |
| WMCC | W8 | Sporichthya | 0.00285714 |
| WMCC | W8 | Staphylococcus | 0.00285714 |
| WMCC | W8 | Subgroup_2_ge | 0.00285714 |
| WMCC | W8 | Tepidisphaeraceae_ge | 0.00285714 |
| WMCC | W8 | Vicinamibacterales_unclassified | 0.00285714 |
| WMCC | W8 | Weeksellaceae_unclassified | 0.00285714 |
| WMCC | W8 | mle1-27_ge | 0.00285714 |
| WMCC | Fresh | Acetobacteraceae_unclassified | 0.00230769 |
| WMCC | Fresh | Actinobacteria_unclassified | 0.00230769 |
| WMCC | Fresh | Clostridia_unclassified | 0.00230769 |
| WMCC | Fresh | HOC36_ge | 0.00230769 |
| WMCC | Fresh | Mucilaginibacter | 0.00230769 |
| WMCC | Fresh | Raoultibacter | 0.00230769 |
| WMCC | Fresh | 67-14_ge | 0.00230769 |
| WMCC | Fresh | Allorhizobium-Neorhizobium-Pararhizobium- | 0.00230769 |
| WMCC | Fresh | Chitinibacter | 0.00230769 |
| WMCC | Fresh | Flavobacteriaceae_unclassified | 0.00230769 |
| WMCC | Fresh | Frankiales_unclassified | 0.00230769 |
| WMCC | Fresh | LD29 | 0.00230769 |
| WMCC | Fresh | Ochrobactrum | 0.00230769 |
| WMCC | Fresh | Rhodopseudomonas | 0.00230769 |
| WMCC | Fresh | SJA-15_ge | 0.00230769 |
| WMCC | Fresh | Williamsia | 0.00230769 |
| WMCC | Fresh | Eggerthellaceae_unclassified | 0.00230769 |
| WMCC | Fresh | Rickettsia | 0.00230769 |
| WMCC | W4 | Aeromonas | 0.00166667 |

| | | | |
|---|---|---|---|
| WMCC | W4 | Arthrobacter | 0.00166667 |
| WMCC | W4 | Bacteroidia_unclassified | 0.00166667 |
| WMCC | W4 | Bdellovibrio | 0.00166667 |
| WMCC | W4 | Beijerinckiaceae_unclassified | 0.00166667 |
| WMCC | W4 | Brachybacterium | 0.00166667 |
| WMCC | W4 | Brevibacterium | 0.00166667 |
| WMCC | W4 | Candidatus_Paracaedibacter | 0.00166667 |
| WMCC | W4 | Cerasicoccus | 0.00166667 |
| WMCC | W4 | Cryptosporangium | 0.00166667 |
| WMCC | W4 | Epulopiscium | 0.00166667 |
| WMCC | W4 | Fimbriimonadaceae_unclassified | 0.00166667 |
| WMCC | W4 | Fusicatenibacter | 0.00166667 |
| WMCC | W4 | Herbaspirillum | 0.00166667 |
| WMCC | W4 | Jatrophihabitans | 0.00166667 |
| WMCC | W4 | Klenkia | 0.00166667 |
| WMCC | W4 | Luteolibacter | 0.00166667 |
| WMCC | W4 | Marmoricola | 0.00166667 |
| WMCC | W4 | Neorhizobium | 0.00166667 |
| WMCC | W4 | Nocardia | 0.00166667 |
| WMCC | W4 | Nocardioidaceae_unclassified | 0.00166667 |
| WMCC | W4 | Pigmentiphaga | 0.00166667 |
| WMCC | W4 | Promicromonosporaceae_unclassified | 0.00166667 |
| WMCC | W4 | Raoultibacter | 0.00166667 |
| WMCC | W4 | Rickettsiales_unclassified | 0.00166667 |
| WMCC | W4 | Saccharimonadales_unclassified | 0.00166667 |
| WMCC | W4 | Shinella | 0.00166667 |
| WMCC | W4 | Solirubrobacteraceae_unclassified | 0.00166667 |
| WMCC | W4 | TM7a | 0.00166667 |
| WMCC | W4 | Tepidisphaera | 0.00166667 |

| | | | |
|---|---|---|---|
| WMCC | W4 | WD2101_soil_group_ge | 0.00166667 |
| WMCC | W4 | endosymbionts | 0.00166667 |
| WMCC | Fresh | Anaerovoracaceae_unclassified | 0.00153846 |
| WMCC | Fresh | Bacteroidia_unclassified | 0.00153846 |
| WMCC | Fresh | Bosea | 0.00153846 |
| WMCC | Fresh | Dickeya | 0.00153846 |
| WMCC | Fresh | Dyadobacter | 0.00153846 |
| WMCC | Fresh | Kineosporia | 0.00153846 |
| WMCC | Fresh | MB-A2-108_ge | 0.00153846 |
| WMCC | Fresh | Methylorosula | 0.00153846 |
| WMCC | Fresh | Nocardiaceae_unclassified | 0.00153846 |
| WMCC | Fresh | Orbaceae_unclassified | 0.00153846 |
| WMCC | Fresh | Oscillospiraceae_unclassified | 0.00153846 |
| WMCC | Fresh | Oxalobacteraceae_unclassified | 0.00153846 |
| WMCC | Fresh | PeM15_ge | 0.00153846 |
| WMCC | Fresh | Pirellula | 0.00153846 |
| WMCC | Fresh | SC-I-84_ge | 0.00153846 |
| WMCC | Fresh | Smaragdicoccus | 0.00153846 |
| WMCC | Fresh | Sphingobacteriaceae_unclassified | 0.00153846 |
| WMCC | Fresh | TK10_ge | 0.00153846 |
| WMCC | Fresh | Xanthobacteraceae_unclassified | 0.00153846 |
| WMCC | Fresh | oc32 | 0.00153846 |
| WMCC | Fresh | Bacteroidales_unclassified | 0.00153846 |
| WMCC | Fresh | Carnobacteriaceae_unclassified | 0.00153846 |
| WMCC | Fresh | Erysipelatoclostridiaceae_unclassified | 0.00153846 |
| WMCC | Fresh | IMCC26256_ge | 0.00153846 |
| WMCC | Fresh | JG30-KF-CM45_ge | 0.00153846 |
| WMCC | Fresh | Lactiplantibacillus | 0.00153846 |
| WMCC | Fresh | Lactobacillaceae_unclassified | 0.00153846 |

| | | | |
|---|---|---|---|
| WMCC | Fresh | Morganellaceae_unclassified | 0.00153846 |
| WMCC | Fresh | Nocardia | 0.00153846 |
| WMCC | Fresh | Oligoflexus | 0.00153846 |
| WMCC | Fresh | Pseudonocardiaceae_unclassified | 0.00153846 |
| WMCC | Fresh | Rhodobacteraceae_unclassified | 0.00153846 |
| WMCC | Fresh | Thermoleophilia_unclassified | 0.00153846 |
| WMCC | Fresh | Weeksellaceae_unclassified | 0.00153846 |
| WMCC | Fresh | Anaerosporobacter | 0.00153846 |
| WMCC | Fresh | Clostridium_sensu_stricto_13 | 0.00153846 |
| WMCC | Fresh | Legionella | 0.00153846 |
| WMCC | W8 | Butyricicoccus | 0.00142857 |
| WMCC | W8 | Clostridia_UCG-014_ge | 0.00142857 |
| WMCC | W8 | Clostridia_unclassified | 0.00142857 |
| WMCC | W8 | Lachnospiraceae_NK4A136_group | 0.00142857 |
| WMCC | W8 | Lachnospiraceae_unclassified | 0.00142857 |
| WMCC | W8 | Ruminococcaceae_ge | 0.00142857 |
| WMCC | W8 | A4b_ge | 0.00142857 |
| WMCC | W8 | Acidipila-Silvibacterium | 0.00142857 |
| WMCC | W8 | Actinocorallia | 0.00142857 |
| WMCC | W8 | Adhaeribacter | 0.00142857 |
| WMCC | W8 | Amnibacterium | 0.00142857 |
| WMCC | W8 | Amycolatopsis | 0.00142857 |
| WMCC | W8 | Anaerobacterium | 0.00142857 |
| WMCC | W8 | Anaeromyxobacter | 0.00142857 |
| WMCC | W8 | Aquisphaera | 0.00142857 |
| WMCC | W8 | Arenimonas | 0.00142857 |
| WMCC | W8 | Babeliaceae_ge | 0.00142857 |
| WMCC | W8 | Babeliales_unclassified | 0.00142857 |
| WMCC | W8 | Bacteroides | 0.00142857 |

| | | | |
|---|---|---|---|
| WMCC | W8 | Bauldia | 0.00142857 |
| WMCC | W8 | Brevibacterium | 0.00142857 |
| WMCC | W8 | C0119_ge | 0.00142857 |
| WMCC | W8 | CCD24_ge | 0.00142857 |
| WMCC | W8 | Caenimonas | 0.00142857 |
| WMCC | W8 | Candidatus_Protochlamydia | 0.00142857 |
| WMCC | W8 | Candidatus_Xiphinematobacter | 0.00142857 |
| WMCC | W8 | Chitiniphilus | 0.00142857 |
| WMCC | W8 | Chryseolinea | 0.00142857 |
| WMCC | W8 | Chthoniobacteraceae_unclassified | 0.00142857 |
| WMCC | W8 | Coxiella | 0.00142857 |
| WMCC | W8 | Cytophaga | 0.00142857 |
| WMCC | W8 | DS-100_ge | 0.00142857 |
| WMCC | W8 | Dactylosporangium | 0.00142857 |
| WMCC | W8 | Dokdonella | 0.00142857 |
| WMCC | W8 | Dongia | 0.00142857 |
| WMCC | W8 | Edaphobacter | 0.00142857 |
| WMCC | W8 | Flaviaesturariibacter | 0.00142857 |
| WMCC | W8 | Flaviflexus | 0.00142857 |
| WMCC | W8 | Flexivirga | 0.00142857 |
| WMCC | W8 | Granulicella | 0.00142857 |
| WMCC | W8 | Heliimonas | 0.00142857 |
| WMCC | W8 | Hirschia | 0.00142857 |
| WMCC | W8 | Hungateiclostridiaceae_unclassified | 0.00142857 |
| WMCC | W8 | IMCC26207 | 0.00142857 |
| WMCC | W8 | JCM_18997 | 0.00142857 |
| WMCC | W8 | Jeotgalicoccus | 0.00142857 |
| WMCC | W8 | Kapabacteriales_ge | 0.00142857 |
| WMCC | W8 | Lachnospiraceae_ge | 0.00142857 |

| | | | |
|---|---|---|---|
| WMCC | W8 | Lacticaseibacillus | 0.00142857 |
| WMCC | W8 | Luedemannella | 0.00142857 |
| WMCC | W8 | Lysobacter | 0.00142857 |
| WMCC | W8 | Martelella | 0.00142857 |
| WMCC | W8 | Mycobacteriaceae_unclassified | 0.00142857 |
| WMCC | W8 | Myxococcus | 0.00142857 |
| WMCC | W8 | Neochlamydia | 0.00142857 |
| WMCC | W8 | Nitrosomonadaceae_unclassified | 0.00142857 |
| WMCC | W8 | Nitrososphaeraceae_ge | 0.00142857 |
| WMCC | W8 | Nitrosospira | 0.00142857 |
| WMCC | W8 | Nitrospira | 0.00142857 |
| WMCC | W8 | Nordella | 0.00142857 |
| WMCC | W8 | OM27_clade | 0.00142857 |
| WMCC | W8 | Paenalcaligenes | 0.00142857 |
| WMCC | W8 | Parachlamydiaceae_unclassified | 0.00142857 |
| WMCC | W8 | Parviterribacter | 0.00142857 |
| WMCC | W8 | Polyangia_unclassified | 0.00142857 |
| WMCC | W8 | Polyangium | 0.00142857 |
| WMCC | W8 | Procabacter | 0.00142857 |
| WMCC | W8 | Pseudolabrys | 0.00142857 |
| WMCC | W8 | Pseudomonadales_unclassified | 0.00142857 |
| WMCC | W8 | Quadrisphaera | 0.00142857 |
| WMCC | W8 | Rhizorhapis | 0.00142857 |
| WMCC | W8 | Roseimicrobium | 0.00142857 |
| WMCC | W8 | Roseisolibacter | 0.00142857 |
| WMCC | W8 | Ruminococcaceae_unclassified | 0.00142857 |
| WMCC | W8 | SH-PL14 | 0.00142857 |
| WMCC | W8 | Shimazuella | 0.00142857 |
| WMCC | W8 | Siphonobacter | 0.00142857 |

| | | | |
|---|---|---|---|
| WMCC | W8 | Sporomusa | 0.00142857 |
| WMCC | W8 | Subgroup_17_ge | 0.00142857 |
| WMCC | W8 | Sumerlaea | 0.00142857 |
| WMCC | W8 | Tepidisphaera | 0.00142857 |
| WMCC | W8 | Thermoactinomyces | 0.00142857 |
| WMCC | W8 | Tumebacillus | 0.00142857 |
| WMCC | W8 | Vampirovibrionaceae_ge | 0.00142857 |
| WMCC | W8 | Vampirovibrionaceae_unclassified | 0.00142857 |
| WMCC | W8 | Variibacter | 0.00142857 |
| WMCC | W8 | Vermiphilaceae_ge | 0.00142857 |
| WMCC | W8 | Verrucomicrobiae_unclassified | 0.00142857 |
| WMCC | W8 | WX65_ge | 0.00142857 |
| WMCC | W8 | mle1-7 | 0.00142857 |
| WMCC | Fresh | 0319-6G20_ge | 0.00076923 |
| WMCC | Fresh | 966-1 | 0.00076923 |
| WMCC | Fresh | A4b_ge | 0.00076923 |
| WMCC | Fresh | Aeromicrobium | 0.00076923 |
| WMCC | Fresh | Aetherobacter | 0.00076923 |
| WMCC | Fresh | Agathobacter | 0.00076923 |
| WMCC | Fresh | Anaerocolumna | 0.00076923 |
| WMCC | Fresh | Anaerolineae_unclassified | 0.00076923 |
| WMCC | Fresh | Apibacter | 0.00076923 |
| WMCC | Fresh | Arthrobacter | 0.00076923 |
| WMCC | Fresh | Aurantisolimonas | 0.00076923 |
| WMCC | Fresh | BIrii41_ge | 0.00076923 |
| WMCC | Fresh | Blastococcus | 0.00076923 |
| WMCC | Fresh | Bryobacter | 0.00076923 |
| WMCC | Fresh | C0119_ge | 0.00076923 |
| WMCC | Fresh | CPR2_ge | 0.00076923 |

| | | | |
|---|---|---|---|
| WMCC | Fresh | Candidatus_Xiphinematobacter | 0.00076923 |
| WMCC | Fresh | Carnobacterium | 0.00076923 |
| WMCC | Fresh | Chitinophaga | 0.00076923 |
| WMCC | Fresh | Chitinophagaceae_unclassified | 0.00076923 |
| WMCC | Fresh | Christensenellaceae_R-7_group | 0.00076923 |
| WMCC | Fresh | Clostridium_sensu_stricto_11 | 0.00076923 |
| WMCC | Fresh | Clostridium_sensu_stricto_6 | 0.00076923 |
| WMCC | Fresh | Clostridium_sensu_stricto_8 | 0.00076923 |
| WMCC | Fresh | Cohnella | 0.00076923 |
| WMCC | Fresh | DS-100_ge | 0.00076923 |
| WMCC | Fresh | Desulfosporosinus | 0.00076923 |
| WMCC | Fresh | Devosia | 0.00076923 |
| WMCC | Fresh | Edaphobaculum | 0.00076923 |
| WMCC | Fresh | Ferruginibacter | 0.00076923 |
| WMCC | Fresh | Fimbriimonadaceae_ge | 0.00076923 |
| WMCC | Fresh | Fonticella | 0.00076923 |
| WMCC | Fresh | Gaiellales_unclassified | 0.00076923 |
| WMCC | Fresh | Gemmata | 0.00076923 |
| WMCC | Fresh | Gemmataceae_unclassified | 0.00076923 |
| WMCC | Fresh | Gemmatimonas | 0.00076923 |
| WMCC | Fresh | Geobacteraceae_unclassified | 0.00076923 |
| WMCC | Fresh | Hymenobacter | 0.00076923 |
| WMCC | Fresh | Hyphomicrobium | 0.00076923 |
| WMCC | Fresh | Iamia | 0.00076923 |
| WMCC | Fresh | Ilumatobacter | 0.00076923 |
| WMCC | Fresh | Intrasporangiaceae_unclassified | 0.00076923 |
| WMCC | Fresh | Isosphaeraceae_unclassified | 0.00076923 |
| WMCC | Fresh | Janthinobacterium | 0.00076923 |
| WMCC | Fresh | Kineococcus | 0.00076923 |

| | | | |
|---|---|---|---|
| WMCC | Fresh | Kineosporiaceae_unclassified | 0.00076923 |
| WMCC | Fresh | Ktedonobacteraceae_unclassified | 0.00076923 |
| WMCC | Fresh | Leptotrichia | 0.00076923 |
| WMCC | Fresh | Leucobacter | 0.00076923 |
| WMCC | Fresh | Methylococcaceae_unclassified | 0.00076923 |
| WMCC | Fresh | Methyloligellaceae_ge | 0.00076923 |
| WMCC | Fresh | Methylomonas | 0.00076923 |
| WMCC | Fresh | Microvirga | 0.00076923 |
| WMCC | Fresh | Natronomonas | 0.00076923 |
| WMCC | Fresh | Nitrososphaeraceae_unclassified | 0.00076923 |
| WMCC | Fresh | Nocardioides | 0.00076923 |
| WMCC | Fresh | Oenococcus | 0.00076923 |
| WMCC | Fresh | Paenarthrobacter | 0.00076923 |
| WMCC | Fresh | Pirellulaceae_unclassified | 0.00076923 |
| WMCC | Fresh | Pseudomonadaceae_unclassified | 0.00076923 |
| WMCC | Fresh | RB41 | 0.00076923 |
| WMCC | Fresh | Rhodoplanes | 0.00076923 |
| WMCC | Fresh | Rice_Cluster_II_ge | 0.00076923 |
| WMCC | Fresh | Rickettsiaceae_ge | 0.00076923 |
| WMCC | Fresh | Roseiarcus | 0.00076923 |
| WMCC | Fresh | Rubritaleaceae_unclassified | 0.00076923 |
| WMCC | Fresh | Saccharimonadales_unclassified | 0.00076923 |
| WMCC | Fresh | Solirubrobacteraceae_unclassified | 0.00076923 |
| WMCC | Fresh | Sphingobium | 0.00076923 |
| WMCC | Fresh | Sporomusaceae_unclassified | 0.00076923 |
| WMCC | Fresh | Subgroup_7_ge | 0.00076923 |
| WMCC | Fresh | Syntrophorhabdus | 0.00076923 |
| WMCC | Fresh | TRA3-20_ge | 0.00076923 |
| WMCC | Fresh | Tepidisphaerales_unclassified | 0.00076923 |

| | | | |
|---|---|---|---|
| WMCC | Fresh | Tumebacillus | 0.00076923 |
| WMCC | Fresh | Verrucomicrobiae_unclassified | 0.00076923 |
| WMCC | Fresh | Vicinamibacteraceae_ge | 0.00076923 |
| WMCC | Fresh | Zavarzinella | 0.00076923 |
| WMCC | Fresh | Acinetobacter | 0.00076923 |
| WMCC | Fresh | Allocatelliglobosispora | 0.00076923 |
| WMCC | Fresh | Amycolatopsis | 0.00076923 |
| WMCC | Fresh | Aphanizomenon_NIES81 | 0.00076923 |
| WMCC | Fresh | Arsenophonus | 0.00076923 |
| WMCC | Fresh | Bifidobacteriaceae_unclassified | 0.00076923 |
| WMCC | Fresh | Candidatus_Accumulibacter | 0.00076923 |
| WMCC | Fresh | Cellulosilyticum | 0.00076923 |
| WMCC | Fresh | Chlamydia | 0.00076923 |
| WMCC | Fresh | Clostridioides | 0.00076923 |
| WMCC | Fresh | Corynebacteriales_unclassified | 0.00076923 |
| WMCC | Fresh | Geodermatophilus | 0.00076923 |
| WMCC | Fresh | Peredibacter | 0.00076923 |
| WMCC | Fresh | Thermoactinomyces | 0.00076923 |
| WMCC | Fresh | Wolbachia | 0.00076923 |
